# BRAFV600E Expression in Mouse Neuroglial Progenitors Increase Neuronal Excitability, Cause Appearance of Balloon-like cells, Neuronal Mislocalization, and Inflammatory Immune response

**DOI:** 10.1101/544973

**Authors:** Roman U. Goz, Ari Silas, Sara Buzel, Joseph J. LoTurco

## Abstract

**BACKGROUND:** Frequent *de-novo* somatic mutations in major components (PI3KCA, AKT3, TSC1, TSC2, mTOR, BRAF) of molecular pathways crucial for cell differentiation, proliferation, growth and migration (mTOR, MAPK) has been previously implicated in malformations of cortical development (MCDs) and low-grade neuroepithelial tumors (LNETs) ^1–7^. LNETs are the most frequent tumors found in patients undergoing resective surgery for refractory epilepsy treatment. BRAFV600E is found in up to 70% of LNETs. Previous studies suggest a causal relationship between those *de-novo* somatic mutations in mTOR, MAPK pathways and seizures occurrence, even without presence of malformation or a tumor ^2, 3, 8–13^. Recently Koh and colleagues ^14^ showed that BRAFV600E mutation may cause seizures through activation of RE1-silecing transcription factor (REST). Additionally, they showed a significant downregulation of synaptic transmission and plasticity pathways and decreased expression of multiple ion channels subunits including HCN1, KCNQ3, SCN2A and SCN3B. The downregulation of those genes including GABA receptors subunits and protein expression specific to interneurons subpopulations (SST, VIP) suggests that a dysregulated inhibitory circuits are responsible for seizures in GGs. The experimental manipulation - *In-Utero* electroporation of episomal activating Cre plasmids that they used to test their hypothesis in mice however activated mutant BRAFV637 only in excitatory neurons. And the downregulated genes in mice were confirmed by qRT-PCR in the whole tissue samples. The question of how electrophysiological properties of the affected and surrounding neurons are changed were not addressed. The changes in ion conductances and neuronal circuits responsible for seizures could be only inferred from gene expression profiles. Purpose of the current work was to investigate how overactive human BRAFV600E mutated protein incorporated into the mouse genome through piggyBase transposition increase neuronal excitability in *ex-vivo* mouse cortical slices and whether it induces histopathological features and gene expression profile alteration observed in low-grade neuroepithelial tumors (LNETs).

**METHODS:** Using *In-Utero* Electroporation we have introduced human BRAFV600E protein into radial glia progenitors in mouse embryonic cortex on the background of piggyBac transposon system that allows incorporation of the DNA sequence of interest into the genome. Immunohistochemistry was used for examination of known markers in LNETs. RNA sequencing on Illumina NextSeq 500 was used to examine alterations in gene expression profiles. Whole-cell current- and voltage-clamp was used to examine changes in electrophysiological properties. Unsupervised Hierarchical Clustering Analysis was used to examine grouping of different conditions based on their gene expression profile and electrophysiological properties. Video electrocorticographic recordings were used to test whether BRAFV600E transgenic mice have spontaneous seizures.

**RESULTS:** Under GLAST driving promoter BRAFV600E induced astrogenesis, caused morphological alterations in transgenic cells akin to balloon-like cells, and delayed neuronal migration. Under NESTIN driver promoter BRAFV600E increased neurogenesis, induced balloon-like cells and caused some cells to remain close to the lateral ventricle displaying large soma size compared to neurons in the upper cortical layers. Some of the balloon-like cells were immunopositive for astroglial marker glial fibrillary acidic protein (GFAP), and for both upper and lower cortical layers markers (Cux1 and Ctip2). Gene ontology analysis for BRAFV600E gene expression profile showed that there is a tissue-wide increased inflammatory immune response, complement pathway activation, microglia recruitment and astrocytes activation, which supported increased immunoreactivity to microglial marker iba1, and to GFAP respectively. In current clamp BRAFV600E neurons have increased excitability properties including more depolarized resting membrane potential, increased input resistance, low capacitance, low rheobase, low action potential (AP) voltage threshold, and increased AP firing frequency. Additionally, BRAFV600E neurons have increased SAG and rebound excitation, indicative of increased hyperpolarization activated depolarizing conductance (I_H_), which is confirmed in voltage-clamp. The sustained potassium current sensitive to tetraethylammonium was decreased in BRAFV600E neurons.. In 4 out of 59 cells, we have also observed a post-action potential depolarizing waves, frequencies of which increased in potassium current recording when Ca^2+^ was substituted to Co^2+^ in the extracellular solution (5/24). We show that using 20 electrophysiological properties BRAFV600E neurons segregate separately from other conditions. Comparison of electrophysiological properties of those neurons with neurons bearing somatic mutations in mechanistic target of rapamycin (MTOR) pathway regulatory components, overactivation of which is been shown in malformations of cortical development (MCDs), showed that expression of PIK3CAE545K under GLAST+ promoter and TSC1 knockdown (KD) with CRISPR-Cas9 have different effects on neuronal excitability.

## 1. Introduction

Low grade neuroepithelial tumors (LNETs) are the second most frequent structural pathology in patients referred for resective surgery of intractable focal epilepsy and present in 25-80% of those cases ^14–36^. The major subtypes of LNETs - predominant in young patients ganglioglioma (GG) that represent 2-5% of pediatric brain tumors ^16, 33^, and dysembryoplastic neuroepithelial tumors (DNETs), in about 20-36% of cases are associated with focal cortical dysplasia (FCD) ^19^. FCD is considered a common cause of drug refractory epilepsy and is found in 25-46% of cases in both children and adults ^19–22, 24, 26, 27, 37–40^. In current International League Against Epilepsy FCD associated with LNETs is defined as FCD IIIb ^41^. However, in some cases other types of FCD has been found in resected cortical tissue from adjacent to LNETs areas ^42, 43^. FCD is a part of larger group of malformations of cortical development (MCDs), which also include tuberous sclerosis complex disorder (TSC), different severity megalencephalies, lisencephaly, microcephaly ^33, 44^. Inspite of abundant data on occurrence and structural pathological components with shared morphometric features in LNETs and MCDs the mechanisms that cause ictogenesis and subsequently development of epilepsy are not currently well understood. Contribution of cortical structure disruption to seizures and epilepsy may depend on specific case, the size of the affected cortical area, susceptibility of the affected cortical area to such disruptions, the population of progenitor cells involved within the boundaries of affected developmental stages ^39, 45–48^, genetic etiology, epigenetic modulation^49, 50^ and environmental effects. In some cases it may be malformation independent and originate in adjacent to malformed cortex areas ^2, 3, 8–13^. Cortical structure disruption may include dyslamination, presence of heterotopic neurons, dysmorphic cytomegalic neurons ^51, 52^, interneurons ^53^, immature misoriented small neurons ^54^, and in sever FCDs cells without clear neuronal or glial differentiation - balloon cells ^41^, giant cells in TSC ^55^, or atypical ganglion cells in LNETs ^35^. On the molecular level, mechanisms that contribute to seizures development involve genetic alterations. Genetic alterations in single components, as well as tissue-wide gene profile showed mutations in mechanistic target of rapamycin pathway (MTOR) in MCDs ^44, 56, 57^ and in mitogen activated protein kinase pathway (MAPK, also RAS-RAF-ERK) in LNETs ^1, 7, 14, 58–60^.

Recent development in DNA/RNA sequencing technologies simplified study of genetic alterations in MCDs and LNETs on a tissue-wide scale. While some studies concentrated on comparison of tens to hundreds of selected genes from microdessected heterotopic neurons, atypical ganglion cells and astrocytes from FCD and GG ^61, 62^ showing that there was a differential expression in glutamate and GABA receptors, and selected growth factors between the cells in tumor or malformation affected area and cells from control tissues, latter studies used either microarrays or RNA sequencing to interrogate global expression profiles and chromosomal reorganization in LNETs, low grade gliomas and TSC ^14, 63–71^. Selective sequencing can still be applied to discover mutations in known malformations associated genes ^4^. Those studies concentrated on discovery of additional somatic postzygotic mutations and chromosome reorganization in LNETs and MCDs, while few of them has also reported on increased inflammatory response and activation of complimentary cascade in GG ^69^ and in TSC ^63, 68^. Interestingly Stone et al. ^64^ used RNA expression profile and DNA methylation profile in LNETs (GG, DNETs, and with uncertain histologic type) to show that most of those segregate into two distinct groups, one group with astrocytic differentiation and is driven by BRAFV600E mutation and the second group had oligodendroglial differentiation and driven by FGFR1 mutation.

BRAF V600E mutation is found in up to 70% of LNETs ^1, 7, 14, 58–60^. Furthermore, recent study showed presence of this mutation in few FCD associated with LNETs cases ^72^.

In the recently developed mouse models of MCD and GG genetic alterations in MTOR and MAPK components in a small population of cortical cells was enough to disrupt cortical structure, cell morphology and cause seizures. Moreover, administration of MTOR and MAPK specific components inhibitors was enough to decrease seizures and prevent structural malformation. ^2, 3, 9, 14, 73^. However, the intrinsic electrophysiological mechanisms that may lead to seizures at the single cell level in those studies were not interrogated. This may be due to previous studies on FCD and TSC cases that showed no significant increase in intrinsic excitability of malformed components, including cytomegalic neurons, balloon cells and immature misoriented neurons ^51, 53, 54, 74–77^.

Here we hypothesized that expression of BRAFV600E mutation associated with LNETs alters gene expression in the affected cortical tissue and increase intrinsic neuronal excitability in BRAFV600E neurons, altering passive and active electrophysiological properties. To this end we used *In-utero* electroporation that allows to introduce genetic manipulation into radial glia progenitor population affecting a small percentage of cells (5-10%) ^78, 79^. This manipulation reflects the percentage of mutated alleles found in MCDs ^2, 4, 9, 73, 80, 81^ and in GG ^14^. Gene expression was examined with RNA sequencing and intrinsic neuronal properties were examined *ex-vivo* in cortical slices with whole-cell patch clamp. Gene ontology analysis of the tissue-wide expression profiles showed that there was a significant increase in immune response, as well as classic complement pathway activation in BRAFV600E cortical tissue. The decreased biological protein pathways included potassium channels. BRAFV600E expressing neurons had hyperexcitable intrinsic properties most prominent of each was increased action potential firing and low current threshold required to fire action potential (rheobase). Other electrophysiological properties that contribute to hyperexcitability of those neurons include more depolarized resting membrane potential, increased input resistance, lower capacitance, more hyperpolarized action potential voltage threshold. In current-clamp experiments significant SAG and rebound excitation in BRAFV600E neurons were observed, a phenomenon associated with hyperpolarization activated depolarizing current (IH). This was confirmed in voltage-clamp showing presence of hyperpolarization activated depolarizing currents (IH) in BRAFV600E neurons only and not in control conditions. Consistent with that SAG and rebound excitation were blocked by ZD7288. Also, in voltage-clamp experiments, we show that BRAFV600E expressing neurons had smaller sustained potassium currents sensitive to tetraethylammonium (TEA) compared to their untransfected neighbors. Finally, using unsupervised hierarchical clustering analysis on electrophysiological properties we show that most BRAFV600E neurons segregate closer together and other experimental conditions comprise the second major group. When comparing those electrophysiological properties with somatic mutations that has been found in FCD and TSC ^3, 73^ (expression of PIK3CA E545K, or CRISPR-Cas9 TSC1 KD) we show that those mutations have different effect on neuronal electrophysiology.

## 2. Materials and Methods

### 2.1 Plasmids and sgRNA sequences

pGlast-PBase and pNestin-PBase were made as previously described ^82^. “PBase was inserted downstream of the Nestin second-intron enhancer in the plasmid Nestin/hsp68-EGFP provided by Steven Goldman ^83^. This 637-bp enhancer of the second intron of rat Nestin gene (GenBank: AF004334.1) was located between bases 1162 and 1798 and is sufficient to control gene expression in the central nervous system neuroepithelial progenitor cells ^84^. For pGLAST-PBase PBase was inserted downstream of the GLAST promoter obtained from Dr. D.J. Volsky ^85^. This 1973-bp GLAST promoter was from human excitatory amino acid transporter 1 (GenBank: AF448436.1). pPBCAG-monomeric red fluorescent protein (mRFP), and pPBCAG-EGFP are constructed as previously described ^86^.” Human BRAFV600E - pBABEbleo-Flag-BRAFV600E was donated by Dr. Christopher Counter and obtained from addgene (Plasmid #53156) ^87^; and human PIK3CAE545K – pBabe-puro-HA-PIK3CAE545K was donated by Dr. Jean Zhao ^88^ and also obtained from addgene (Plasmid #12525). The BRAFV600E and PIK3CAE545K inserts were amplified with standard PCR and cloned into pPBCAG-EGFP replacing EGFP sequence using EcoRI and NotI sites. Hemagglutinin (HA), a 27 nucleotides epitope tag (5’-AGCGTAATCTGGAACATCGTATGGGTA-3’) was inserted into pPBase-BRAFV600E after BRAFV600E sequence and before NotI site. pPBase-BRAFwt was generated with quick change II XL single nucleotide site directed mutagenesis kit from Agilent according to the manufacturer protocol, to change E, a glutamic amino acid back to V - valine at position 600 restoring the mutated sequence back to its wild type. The sequence restoration to BRAFwt was confirmed with Sanger sequencing. Channelrhodopsin plasmid (pcDNA3.1hChR2-EYFP) was a gift from K Diesseroth, Stanford University, Stanford, CA, and was subcloned into the pCAG plasmid and used before ^89^. Guide RNA for TSC1 (T4 – 5’-CCATGCTGGATCCTCCACACTG-3’) and TSC2 (T7 – 5’-CCAAATCCCAGGTGTGCAGAAGG-3’) were chosen based on ^73^. These sequences were cloned into pX330 vector (Addgene, plasmid #42230)^90^ following normal cloning procedure.

### 2.2 Animals

Pregnant CD1 mice were obtained from Charles River Laboratories (Wilmington, MA, USA) and maintained at the University of Connecticut vivarium on 12 h light cycle and fed ad libitum. Animal gestational ages were determined via palpation prior to and confirmed during the surgery based on crown-ramp length ^91^. Female and male mice were used for cortical transgene delivery with *In-utero* electroporation. All procedures and experimental approaches were approved by the University of Connecticut IACUC.

### 2.3 In utero electroporation

*In-utero* electroporation was performed as previously described ^92^. Briefly, mice were anesthetized with a mixture of ketamine/xylazine (100/10 mg/kg i.p.). Metacam analgesic was administered daily at dosage of 1 mg/kg s.c. for 2 days following surgery. To visualize the plasmid during electroporation, plasmids were mixed with 2 mg/ml Fast Green (Millipore Sigma, F7252). In all conditions, pPBCAG-EGFP, pPBCAG-mRFP, pPB-BRAFV600E, pPB-BRAFwt, pPB-PIK3CAE545K, pGLAST-PBase, pNESTIN-PBase were used at the final concentration of 1.0 μg/μl. Electroporation was performed at embryonic day 14 or 15. During surgery, the uterine horns were exposed and one lateral ventricle of each embryo was pressure injected with 1-2 μl plasmid DNA. Injections were made through the uterine wall and embryonic membranes by inserting pulled glass microelectrodes (Drummond Scientific) into the lateral ventricle and injecting by pressure pulses delivered with Picospritzer II (General Valve). Electroporation was accomplished with a BTX 8300 pulse generator (BTX Harvard Apparatus) and BTX tweezertrodes. A voltage of 35-45 V was used for electroporation.

### 2.4 Image acquisition, cell counting and measurement

Images were acquired on inverted Leica TSC SP8 confocal microscope with four PMT detectors and one HyD detector equipped with 405 nm diode laser, argon (458/488/514 nm) laser, 561 nm DPSS laser and 633 nm HeNe laser. Sets of images for all the experimental and control conditions in each group (GLAST+, NESTIN+) were acquired on the same day with the same excitation power and gain settings. Some of the images were acquired with Zeiss Axiozoom.V16 with 405/488/568/647 filters and Lumencor’s SOLA SE 365 light engine with ~3.5W white light output through 3 mm dia liquid light guide (LLG) with PlanNeoFluar Z 2.3x with 0.57 n.a. lens. Axiozoom was equipped with sCMOS pco.edge 4.2 camera with CIS2020A sensor. All the images were further processed in ImageJ-Fiji package (version 1.51w, NIH, RRID: SRC_003070) ^93^. For manual cell counting and distance to pia measurement images were converted to black for EGFP and white background and pia was oriented as a horizontal plane and cells were counted with cell count plugin in Fiji by A. S., S.B. and R.G. Soma size was measured with a freehand selection tool and measure under the same brightness/contrast and color balance settings in all conditions. Balloon-like cells and aggregates were scanned and counted by S.B. and R.G. with Axiozoom Zeiss.V16. Image processing for publication was done in Fiji and Corel Draw Graphics Suite X8 (Corel, Ottawa, Canada; RRID: SCR_002865).

### 2.5 Immunohistochemistry

Animals were deeply anesthetized with isoflurane and perfused transcardially with 4% paraformaldehyde/PBS (4% PFA). Samples were post fixed overnight in 4% PFA. For immunofluorescence, brains were sectioned at 50-μ thickness on a vibratome (Leica VT 1000S). Sections were processed as free-floating and stained with rabbit polyclonal anti-GFAP (1:2000 dilution, DAKO Z0334, GenBank L19867, RRID:AB_10013482), mouse monoclonal anti-Aldehyde Dehydrogenase 1 family 1, member L1 (ALDH1L1, 1:50 dilution, NeuroMab, cat. #75-140, RRID:AB_10673448, clone N103/39, accession number P28037), goat polyclonal anti-Iba1 (1:200 dilution, Invitrogen, cat. # PIPA518039, accession number P55008), nuclear staining with Hoechst 33342, trihydrochloride, trihydrate – 10 mg/ml in water (1:3000 dilution, Molecular probes by life technologies, cat. # H3570). After blocking in PBS containing 5% of normal goat serum (Millipore Sigma, NS02L) and 0.5% Triton X-100 (Millipore Sigma, X100) for 2 h at room temperature, tissue was washed three times in PBS with 2.5% normal goat serum and 0.2% Triton X-100 (washing solution), followed by incubation with primary antibodies overnight at 4°C in the washing solution. On the following day tissue was washed again in washing solution and incubated with the appropriate secondary antibodies in washing solution (all Alexa Fluor in 1:1000, Invitrogen) for 2 h at room temperature (Alexa Fluor 568 anti-mouse IgG, Alexa Fluor 647 anti-rabbit IgG, Alexa Fluor 568 anti-goat IgG). After 2 h the tissue was washed again with washing solution once, stained with Hoechst 33342 and washed again three times. Tissue was mounted on Fisherbrand Colorfrost Plus Microscope slides (Cat #12-550-19) submerged in ProLong gold antifade (Life technologies, cat. #36930) and coverslipped with Fisherfinest premium cover glass (cat. #12-548-5P,5J,B, sizes 24×60-1, 24×40-1,22×22-1 respectively). When prolong gold antifade has cured the coverslips edges were covered with transparent nail polish.

### 2.6 RNAseq, gene ontology analysis and Gene Analytics

Mouse brains were extracted at P65 from 4 GLAST+ BRAFV600E animals, 4 GLAST+ BRAFwt animals, 4 GLAST+ control-FP animals (2 from GLAST+ BRAFV600E litter and 2 from GLAST+ BRAFwt litter) that were electroporated at E14 after deep anesthetization with isoflurane. The fluorescent EGFP area of somatosensory cortex was dissected and the white matter remains were cut out from those tissue chunks. The remaining cortical tissue chunks were further dissociated and processed with Ambien RNA extraction kit according to manufacturer protocol. The range of RNA amount for the samples was from 300-900 ng per sample measured with nanodrop-1000 spectrophotometer. Quality was assessed by RNA Integrity Numbers and values ranged from 6.6 to 8.5 for first stranded cDNA library preparation and analysis with Illumina NextSeq 500 – mid output v2 (150 cycles). Libraries were sequenced at a depth of 9.6 to 18 million reads per sample. Quality control, library preparation and sequencing were done at UCONN Center for Genome Innovation, Institute for System Genomics. For the further processing and analysis of sequencing results new tuxedo protocol was used ^94^ on UCONN High Performance Computing cluster. Fasq files containing sequencing fragments were aligned with HISAT2 (RRID:SCR_015530) ^95^ using index build for mouse genome fasta file downloaded from Ensemble data base (ftp://ftp.ensembl.org/pub/release-92/fasta/mus_musculus/dna/Mus_musculus.GRCm38.dna.primary_assembly.fa.gz), produced sequence alignment maps (sam) files were sorted with samtools (RRID:SCR_002105) outputting binary alignment maps (bam) files, that were assembled, merged with the mouse reference genome from Ensemble data base (ftp://ftp.ensembl.org/pub/release-92/gtf/mus_musculus/Mus_musculus.GRCm38.92.gtf.gz) and quantified with Stringtie ^96^. Stringtie raw counts for further differential expression analysis were extracted with Python script (http://ccb.jhu.edu/software/stringtie/dl/prepDE.py) using Python 2.7 (RRID:SCR_002918). Differential gene expression was estimated with DESeq2 1.18.1 (RRID:SCR_015687) ^97^ in R 3.4.4 (RRID:SCR_001905) ^98^ using R Studio GUI ^99^ and FDR corrected p-values (q values) of 0.05 were considered significant ^100^. The results of differentially expressed genes were further analyzed for functional enrichment with DAVID 6.8 ^101, 102^ (https://david.ncifcrf.gov/home.jsp)

Gene Analytics web-service (geneanalytics.genecards.org) ^103^ was used to analyze differentially expressed genes. Pseudorandom mouse gene list was generated in molecular biology online apps web tool (http://molbiotools.com/randomgenesetgenerator.html) using Mersenne Twister ^104^ pseudorandom number generator algorithm (from personal communication with Vladimír Čermák, the site developer).

### 2.7 Slice preparation

The P15-P70 (average P36.05, mode=36 median=35, stdev=9.86) CD1 were deeply anesthetized with isoflurane and then decapitated. Brains were rapidly removed and immersed in ice-cold oxygenated (95% oxygen and 5% carbon dioxide) dissection buffer containing (in mM/L): 83 NaCl, 2.5 KCl, 1 NaH_2_PO_4_, 26.2 NaHCO_3_, 22 dextrose, 72 sucrose 0.5 CaCl_2_, and 3.3 MgCl_2_, Coronal slices (400 μm) were cut with a vibratome (VT1200S; Leica, Nussloch, Germany), incubated in dissection buffer for 40 min at 34°C, and then stored at room temperature for the reminder of the recording day. Most of the slice recordings were performed at 34°C, besides voltage-clamp recordings of calcium and potassium currents. Slices were visualized with inversion recovery differential interference microscopy (E600FN; Nikon, Tokyo, Japan) and a CCD camera (QICAM; QImaging, Surrey, British Columbia, Canada). Individual neurons were visualized with a 40X Nikon Fluor water immersion (0.8 n.a.) objective.

### 2.8 Electrophysiology

For all experiments except potassium and calcium currents recordings, extracellular recording buffer was oxygenated (95% oxygen and 5% carbon dioxide) and contained (in mM/L): 125 NaCl, 25 NaHCO_3_, 1.25 NaH_2_PO_4_, 3 KCl, 25 dextrose, 1 MgCl_2_, and 1.3 CaCl_2_, 295-305 mOsm. For potassium currents recordings the extracellular calcium was substituted with 1.3 CoCl_2_ · 6H_2_O (Millipore Sigma, CAS number 7791-13-1). For calcium currents recordings extracellular recording buffer was oxygenated (95% oxygen and 5% carbon dioxide) and contained (in mM/L): 130 NaCl, 25 NaHCO_3_, 1.25 HaH_2_PO_4_, 3 KCL, 25 dextrose, 1 CaCl_2_, 1.3 MgCl_2_, 40 TEA, 0.001 TTX, 0.01 gabazine, 0.05 D-APV, 0.01 NBQX, 3 4AP, 0.05 ZD7288. Patch pipettes were fabricated from borosilicate glass (N51A; King Precision Glass, Claremont, California) to a resistance of 2-7 MΩ. The resultant errors were minimized with bridge balance and capacitance compensation. For current-clamp experiments, voltage-clamp recording of hyperpolarization activated currents and potassium currents pipettes were filled with an internal solution containing (in mM/L): 125 K-gluconate, 10 HEPES, 4 Mg_2_-ATP, 3 Na-GTP, 0.1 EGTA, 10 Na-phosphocreatine, 0.05% biocytin, adjusted to pH 7.3 with potassium hydroxide and to 275-285 mOsm with double-distilled water. For voltage-clamp recordings of calcium currents pipettes were filled with an internal solution containing (in mM/L): 110 CsMeSO_4_, 10 CsCl, 5 CaCl_2_, 10 EGTA, 10 HEPES, 4 Mg_2_-ATP, 0.3 Na-GTP, 10 Na-phosphocreatine, 0.05% biocytin, 25 TEA. Signals were amplified with Multiclamp 700B amplifier (Molecular Devices, Sunnyvale, CA), sampled at 20 kHz, digitized (ITC-18; HEKA instruments, Bellmore, NY) and filtered at 2 kHz with an 8-pole low-pass Bessel filter. Data were monitored, acquired, and in some cases analyzed with Axograph X software (Berkley, CA). Series resistance was monitored throughout the experiments by applying a small test voltage step and measuring the capacitive currents. Series resistance was 5 to approximately 25 MΩ, and only cells with <20% change in series resistance and holding current were included in the analysis. Reported membrane potentials and holding potentials were not corrected for liquid junction potential ~ 10 mV unless otherwise specified.

For estimation of the effect of early opening of potassium channels on AP firing frequency retigabine was used (10 μM/L; Alomone labs, Jerusalem, Israel, Cat # R-100), the 100 mM stock was prepared in DMSO and diluted into extracellular recording buffer. In specified experiments D-APV dissolved in double deionized water (ddw) to 50-100 mM stock (50 μM/L; Abcam, Cambridge, MA, cat. # ab120003, lot #GR205917) was used to block specifically NMDAR, and NBQX dissolved in DMSO to 100 mM stock (10 μM/L; Abcam, Cambridge, MA, cat #ab120045, lot #GR133243) was used to block AMPARs; SR95531 dissolved in ddw to 25 mM stock (gabazine, 10 μM/L; Abcam, Cambridge, MA, cat. #ab120042, lot #GR69200) was used to block GABA_A_R; TTX-citrate dissolved in ddw to 10 mM (1 μM/L; Abcam, Cambridge, MA, cat. #ab120055, lot #GR246757) was used to block sodium currents; 4AP dissolved in ddw to 200 mM (30 μM/L – 3 mM/L; Millipore Sigma, Burlington, MA, cat. #275875) was used to block fast activating fast inactivating potassium currents; TEA (25-40 mM/L; Abcam, Cambridge, MA, cat. #ab120275, lot #GR69136) was used to block sustained potassium currents; ZD7288 dissolved in ddw (50 μM/L; Cayman Chemicals, Ann Arbor, MI, cat. # 1522820000040, batch number 0476856-6) was used to block hyperpolarization activated depolarizing currents (I_H_).

For action potential (AP) firing frequency, input resistance measurement, 1 second current steps were applied at 10 pA increment from −40 pA to 300 pA. For TSC1/2 KD neurons those steps increased to more than 300 pA. Input resistance was measured from last 100 ms of 1 s hyperpolarizing and depolarizing subthreshold current steps and current-voltage relationship was fit with linear regression to estimate input resistance from both depolarizing and hyperpolarizing current steps. Resting membrane potential (RMP) was measured in the beginning of current-clamp protocols before application of current step pulses. SAG ratio was defined as 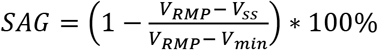 V_RMP_ – Resting Membrane Potential, V_ss_ – stable-state voltage in the last 100 ms of 1 second −40 pA pulse, V_min_ – minimal initial voltage deflection in response to 1 second −40 pA pulse. Rebound excitation was measured as an overshoot above RMP at the end of −40 pA 1 second current step, in some cells resulting in AP firing. AP voltage threshold was defined as the point at which the first derivative of voltage to time (dV/dt) crossed 50 V/s. Rheobase is the minimal current step required to elicit first AP firing. AP peak was measured from RMP.

To record potassium currents the neurons were held at −90 mV and the 500 ms voltage steps proceeded with 10 mV increments from −100 to +20 mV. For sustained potassium currents the amplitude was measured at the last 100 ms of 500 ms voltage steps. The capacitive currents were canceled with internal Multiclamp 700B compensation circuit. Cell capacitance and input resistance in those experiments was monitored and measured before compensation from +5 mV 150 ms voltage steps with Axograph X built in procedure designed to measure series resistance, membrane capacitance and membrane resistance. The measurement was done offline after offline leak current subtraction with scaled voltage steps of opposite polarity to steps that elicit potassium currents.

To record hyperpolarization activated depolarizing currents (I_H_) two protocols were used. In the main protocol, used for the analysis the neurons were held at −50 mV and 1.5 s voltage steps proceeded with 5 mV increment from −120 mV to −35 mV without capacitance and series resistance compensation. The peak measurement of I_H_ was done from the point at the beginning of the observe current to the stable state at the last 100 ms of 1.5 s voltage steps. The tail currents were measured after the voltage steps ceased. All the measurements were done offline after offline leak current subtraction with scaled voltage steps of opposite polarity to steps that elicit I_H_. The second protocol was used to increase stability of the recorded cells. In this protocol neurons were held at −70 mV and 1 s voltage steps proceeded with 5 mV increment from −100 mV to −45 mV.

To record calcium currents neurons were held at −80 mV and 200 ms voltage steps proceeded with 5 mV increment from −90 mV to 0 mV. Maximal negative deflection was used for peak current estimation at each voltage step and to construct activation curve.

Spontaneous Post-Synaptic currents (sPSCs) recording of 1-5 min was done with the chart procedure in Axograph X and was sampled at 5 kHz. The sPSCs were detected with semiautomatic sliding template method as previously described ^105^ and were visually confirmed. The parameters of the template, including amplitude, 10-90% rise time, and decay time were determined on the bases of an average of real events as well as previously reported values. The detection threshold is 2.5 times of the noise SD. The sliding template length was chosen to be 10 ms for all neurons.

### 2.9 Headmount installation and video ECoG recordings

EEG system, including headmounts, preamplifiers, amplifiers and video-ECoG recording system was purchased from pinnacle technology Inc. (Lawrence, KS). BRAFV600E and control-FP electroporated mice of at least 6 months of age were used for those experiments. Surgery was performed under general, continuous isoflurane/O_2_ anesthetic inhalation system at 1-1.5 litters/minute, with intraperitoneal injection of metacam analgesic at 5mg/kg before beginning of the procedure. The mice were stabilized in a mouse stereotaxic apparatus (Stoelting, Wood Dale, IL). The fur was shaved off from the mouse head with small trimmers and disinfected with chlorhexidine, 2% (Henry Schein Animal Health, Dublin, OH). The rostral-caudal incision in the skin was made with 25 mm cutting edge surgical scissors to allow sufficient space on the mouse skull for the headmount (about 1.5 cm). Before mounting hydrogen peroxide was applied to the surface and surgical cotton-tipped sterile q-tips were used to remove periosteum. Four pilot holes were made in the skull through the openings in the headmount with 25-gauge BD needle at the following approximate coordinates relative to bregma: −2 mm; ML: ± 1.5 mm, lambda: −2 mm. Two small pockets were made in the nuchal muscle for EMG electrodes insertion. After insertion of EMG electrodes, the headmount (#8201) with platinum and iridium leads was placed on the surface of the skull covered with cyanoacrylate glue. Four stainless steel ECoG screws, 2-0.10” in front and 2-0.12” screws in back (#8209 and #8212 respectively) were inserted through the headmount openings and manually rotated into the pilot holes. Before the screws were fully locked in place, two-part silver epoxy was applied between the screw heads and the headmount to ensure electrical conductivity. After securing the headmount with screws, dental acrylic cement was applied with a small brush dipped in acetone to the area surrounding the headmount and the base of the headmount. The dental cement cured within 2-5 minutes. One to five skin sutures were applied to close the skin incision. After the surgery mice were places in a clean cage on a warm hitting pad to recover. ECoG data acquisition started 5-7 days after the headmount surgery.

Simultaneous three mice (up to four) video ECoG recordings were performed using pinnacle #8206 data conditioning and acquisition system (DCAS) with 2 electrocorticographic (ECoG) and 1 electromyographic (EMG) channels and dome cameras with infrared light source for night time recording (#9022). The recordings continued for at least five days. Mice were housed in the circular acrylic cage on 10” × 10” base with 10” diameter/8” height. Water and food was provided ad libitum. In the beginning of the video EEG data acquisition the headmount was connected to the 3-channel mouse preamplifier (#8202-SL, the sleep configuration). The preamplifier had AGND ground connection for the animal to avoid input amplifier overcharge. This X100 preamplifier was connected to the secondary amplifier, AD/DA and filtering system – DCAS #8206 (X50.78), that was mounted on a swivel plate to allow mice to move freely. DCAS #8206 together with dome cameras was connected to the desktop PC. Sirenia acquisition software version 1.7.10 was used for simultaneous ECoG/Video recording. The ECoG data was sampled at 600 Hz and low-pass filtered with 8^th^ order progressive elliptic analogue hardware implemented filter at 25 Hz (6 dB/octave). The EMG data was sampled at 600 Hz and low-pass filtered at 100 Hz. Video recording was acquired at 20 f/s in grayscale with 60% image quality to avoid filling up hard drive capacity too fast in X1 MJPG compression at 640×480 pixels resolution. The recorded data was analyzed in Sirenia Seizure Pro software version 1.7.10.

### 2.10 Unsupervised hierarchical clustering analysis

The hierarchical clustering was done with Gene Cluster 3.0 (http://bonsai.hgc.jp/~mdehoon/software/cluster/software.htm), a freeware developed in Michael Eisen lab in Berkeley, using pairwise average linkage with Euclidean distance calculated to determine the difference between clusters of neurons by the length of the branch ^106^. We have used 20 recorded electrophysiological parameters, for which each observation was standardized by centering to the mean and dividing by standard deviation. The missing values were imputed by Non-Linear Iterative Partial Least Squares (NIPALS) algorithm in XLSTAT (https://www.xlstat.com/en/) Microsoft Excel (Microsoft, Redmond, Washington) addon ^107, 108^. The heatmap visualization for RNAseq data was done with ClustVis ^109^ (https://biit.cs.ut.ee/clustvis/). For electrophysiological data it was done in Genepattern ^110^ (https://genepattern.broadinstitute.org/, RRID:SCR_003201) with HierarchicalClusteringViewer module v.11.3.

### 2.11 Statistical analysis

All data measurements were kept in Excel (Microsoft, Redmond, Washington) and in Origin (OriginLab, Northampton, MA; RRID: SCR_002815). All the electrophysiological data was analyzed in SPSS v.24 (RRID:SCR_002865) ^111^, for large samples one-way analysis of variance (ANOVA) with Tukey posthoc correction was used, when the samples had non-homogenous variance (significant Levene test for equality of variance) Welch test with Games-Howell posthoc correction was used. For small samples from different observations independent samples two-tailed student t-test was used and depending on Levene test significance the t statistics for equal or unequal variance was reported. For measurement coming from the same neurons before and after treatment paired samples two-tailed student t-test was used. Graphical visualization of data was prepared in Origin and exported to Corel Draw Graphic Suite X8 for further processing. Arithmetical averages and SEMs were reported for all results unless otherwise specified.

## 3. Results

### 3.1 BRAFV600E alters neuronal migration and morphology

To test whether BRAFV600E is sufficient to cause developmental disruptions in cortex we introduced BRAFV600E transgenes or control-FP transgenes (mRFP) into populations of neocortical progenitors using the binary piggyBac transposon system ^82, 112–114^. We directed transgenes into either a population of GLAST+ neural progenitors that generates both pyramidal neurons and astrocytes ^82, 115^, or a NESTIN+ population that generates primarily pyramidal neurons (Figure 1A) ^116–118^. We compared the numbers, positions and morphologies of cells generated from these progenitors in three different transgene conditions (BRAFV600E, BRAFwt, and mRFP). Consistent with the upper layer laminar fates of neurons with birth dates at E14, when the transgenes were introduced, the neurons generated in all conditions expressed the upper layer pyramidal neuron marker CUX1, but not the lower layer pyramidal neuron marker CTIP2 (Figure 1C). Although positive for upper layer markers, there was a significant increase in the number of neurons in the BRAFV600E transgene conditions that were in deeper layers relative compared to control conditions (Figure 1C lower panel, Figure 2C). Interestingly, the pattern of altered neuronal positioning was different for BRAFV600E introduced into GLAST+ progenitors relative to NESTIN+ progenitors, with a subpopulation of neurons generated from NESTIN+ progenitors displaced even deeper into cortex, into the subventricular white matter. The dyslamination observed in our experiments concur the results obtained with episomal Cre expression in BRAFV637E transgenic mice ^14^. However, quantification of neuronal soma sizes indicated no significant difference in the sizes of neurons generated from GLAST+ progenitors and control conditions, whereas neurons from NESTIN+ progenitors that were also displaced into white matter had significantly larger somas with ganglion cell like morphologies (Figure 2A,B). Those results provide wider insight into the neuroglial progenitors affected population compared to Koh et al. work ^14^. Together these results indicate that BRAFV600E disrupts the normal laminar positioning attained during migration, and the different progenitor populations (NESTIN+ and GLAST+) respond differently to overactive mutant BRAF. To separate the effect of ectopic expression of human BRAF protein from the effect of BRAFV600E mutation, which has been shown in COS7 cell cultures to have 500-fold higher basal kinase activity compared to BRAFwt ^119^, we examined neurons position and morphology in cortical slices with introduced wild type human BRAF transgene. There was a smaller but significant effect compared to control-FP transgenic brains on neuronal migration. Additionally, the increase in the number of neurons in BRAFwt found in lower cortical layers was lower compared to BRAFV600E transgenic neurons (Figure 1E).

**Figure 1.**
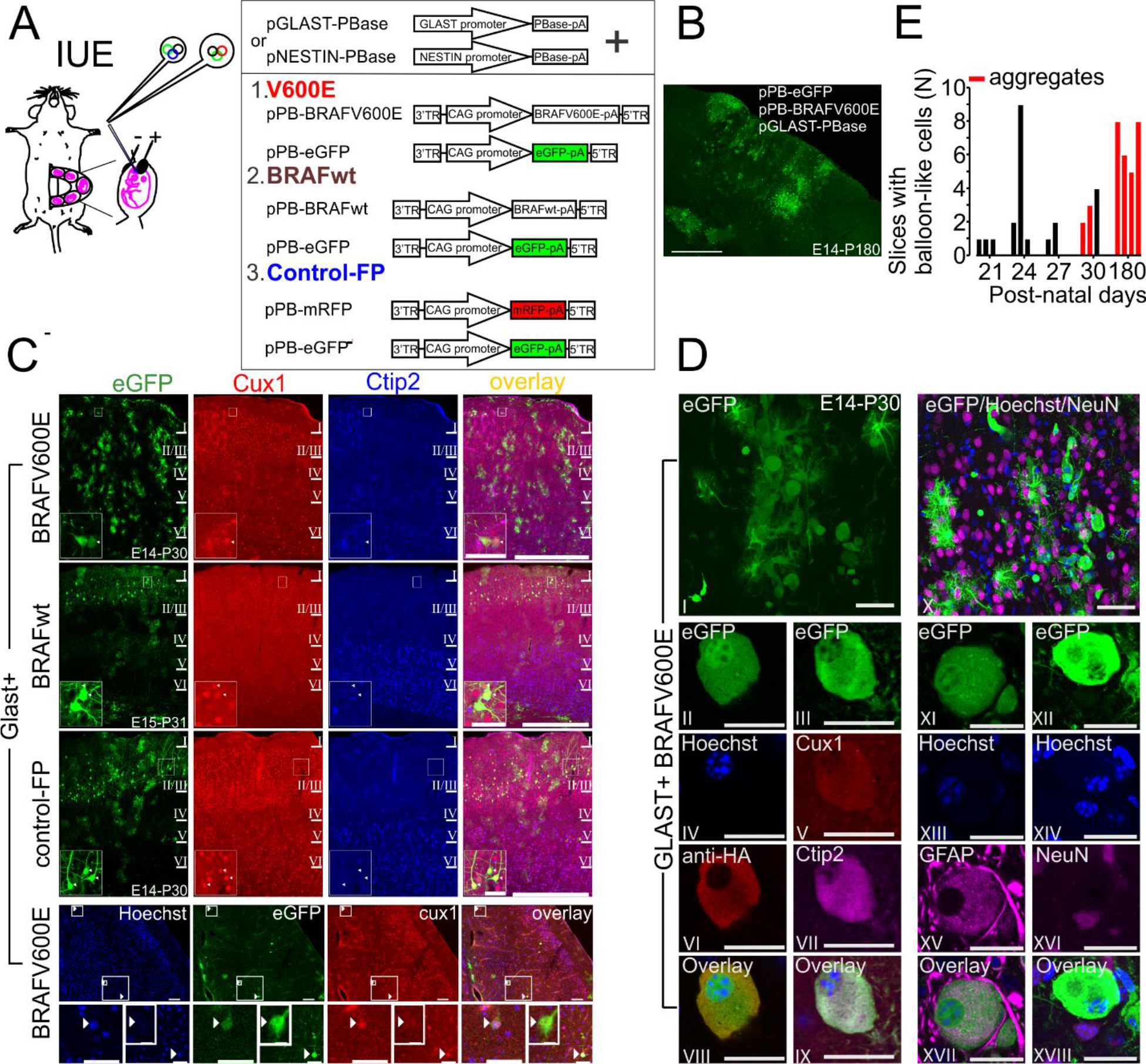
BRAFV600E expression in neuroglial progenitors at embryonic ages E14-E15 causes neuron migrational delay and cellular dysmorphogenesis. **A.** *In-Utero* Electroporation (IUE) experimental design. **B.** Representative image of GLAST+ BRAFV600E electroporated mouse brain slice at post-natal day 180 showing aggregates of balloon-like cells near the pia and aggregates of a crescent shaped cells in layers 4 and 5. **C.** Cux1, an upper cortical layer marker and Ctip2, a lower cortical layer marker shows that most of the GLAST+ BRAFV600E transfected neurons reach appropriate cortical layers relative to embryonic age at IUE. Insets show zoomed in neurons positive for Cux1, and negative for Ctip2 in upper layer 2. Lower panel shows 3 neurons located in the lower cortial layers and positive for cux1. **D.** Balloon-like cells and aggregates (panels I and II, Roman numerals – R.n.) of balloon-like cells found in Glast+ BRAFV600E transfected brains. Panels II,IV,VI,VIII (even R.n.) show anti-HA stain confirmation of pPB-BRAFV600E presence. Panels III-IX with odd R.n. show balloon-like cell stained positive for both Cux1 and Ctip2. Panels XI-XVII, odd R.n. show a balloon-like cell stained positive for Glial Fibrilary Acidic Protein (GFAP) an astrogllial markers. Panels XII-XVII, even R.n. show a ballon-like cell negative for NeuN, a neuronal marker. Panels XIX-XXI showing coronal section from Nestin+ BRAFV600E with balloon-like cells aggregates in a piriform cortex area (white arrowhead). **E.** Frequency of slices with balloon-like cells and balloon-like cells aggregates in Glast+ BRAFV600E transfected brains according to the post-natal age. Each bar represent one brain. Scale bars for B. – 500 μm; C. - 500 μm and 50 μm zoomed in images, Lower panel – 100 μm for the wide field view, 50 μm for single neurons, 10 μm for single zoomed in neuron; for D. I,X – 50 μm; II-IX, and XI-XVIII, R.n. – 20 μm; * - p<0.05, ** - p<0.01, *** - p<0.001. Error bars are ±2SEM.

**Figure 2.**
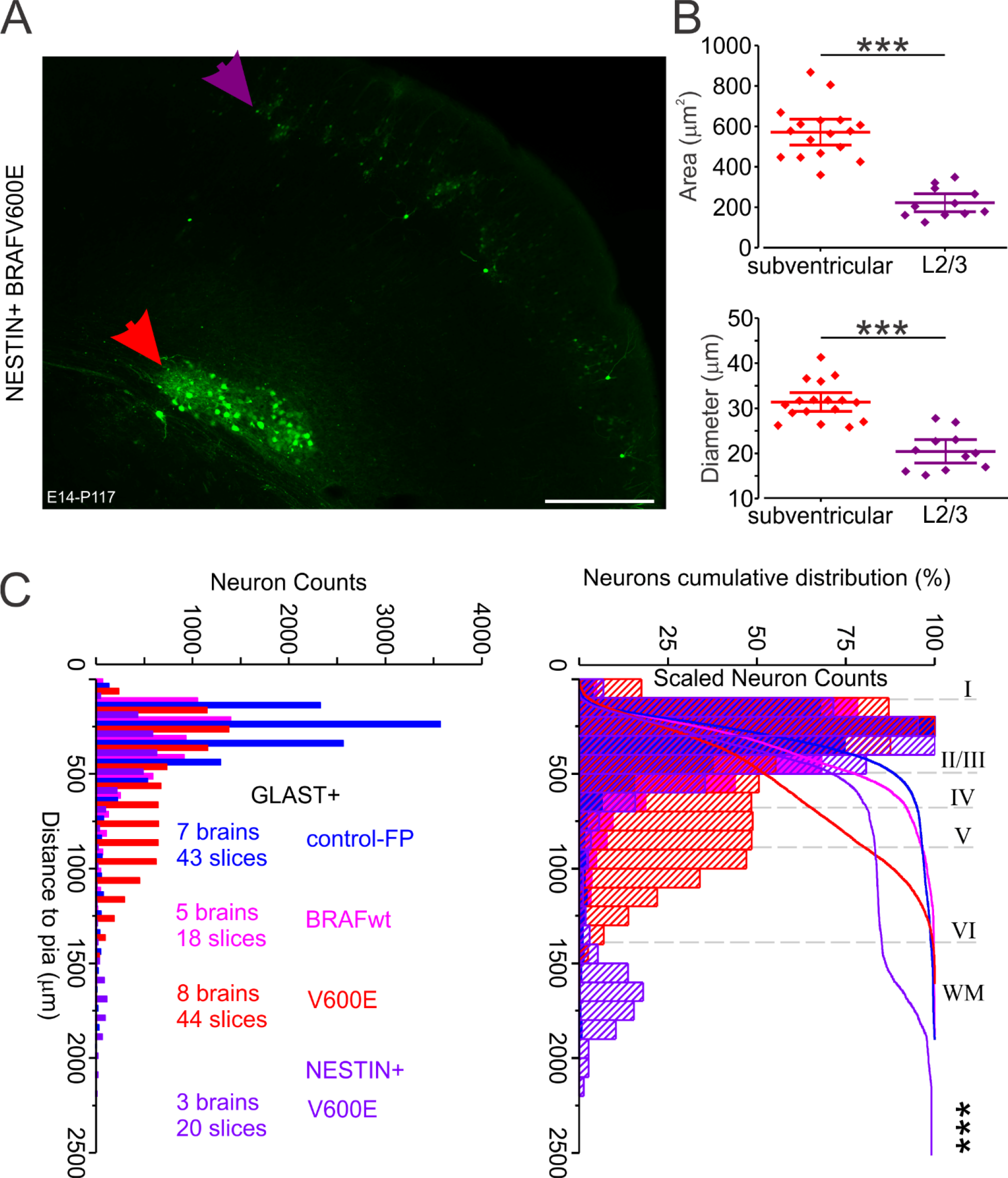
BRAFV600E expression in NESTIN+ neuroglial progenitors. **A.** Representative image of EGFP positive cells in subventricular area compared to cells in layer 2/3 of somatosensory cortex that are quantified in **B.** Area measured in NESTIN+ BRAFV600E EGFP positive cells in A in the upper panel, and diameter of those cells in the lower panel showing increased size of subventricular located cells (n=17) compared to layer 2/3 neurons (n=11, paired sample T=7.883, p<0.001 for area and T=8.224, p<0.001 for diameter). **C.** Gross neuron counts (left panel) and scaled to max neuron counts (right panel) - distance to pia measurement shows that there is a decrease in EGFP positive neuronal content in Glast+ BRAFV600E transfected mouse cortical slices and that higher number of the BRAFV600E transfected EGFP positive neurons targeted for upper cortical layers under both Glast+ and Nestin+ do not reach their deisgnated location compared to Glast+ control-FP transfected brain slices (p<0.001), and to Glast+ BRAFwt transfected brain slices (p<0.001), there was also higher number of Glast+ BRAFwt transfected neurons that did not reach designated cortical layers compared to Glast+ control-FP (p<0.001); significant difference in neuronal distance to pia was also present between Glast+ BRAFV600E and Nestin+ BRAFV600E (p=0.001) Due to non-homogenous variance - Levene test (3, 28606) = 1674.918, p<0.0001, Welch test with Games-Howell post-hoc correction were used (3, 9803.913) = 970.974, p<0.001. The results were confirmed with cumulative distribution nonparametric Mann-Whitney U test – Glast+ BRAFV600E to Glast+ control-FP, U= 31122376, p<0.001; Glast+ BRAFV600E to Glast+ BRAFwt, U= 18003030, p<0.001; Glast+ BRAFwt to Glast+ control-FP, U= 26624785, p<0.001; Nestin+ BRAFV600E to Glast+ control-FP U= 11954144.5, p<0.001; Nestin+ BRAFV600E to Glast+ BRAFwt, U= 7186563, p<0.001; Nestin+ BRAFV600E to Glast+ BRAFV600E, U= 12109290.5, p<0.001. Scale bars for A – 500 μm * - p<0.05,** - p<0.01, *** - p<0.0001. Error bars are ±2SEM.

### 3.2 BRAFV600E causes development of “Balloon-like” cells

In addition to the delayed neuronal migration observed in BRAFV600E transgenic mouse brains we tested whether there was an effect on cellular morphology. Balloon cells are a distinctive cell type characteristic of FCDIIb ^41, 120, 121^ and have also been described in FCDs in the vicinity of LNETs positive for BRAFV600E ^43, 72^. Additionally, in LNETs atypical cytomegalic ganglion cells has been shown to have like balloon cells morphology and protein expression ^61^. In cortical tubers resected from TSC patients the giant cells also show similar morphology ^122–124^. These unusual cells have distinctive morphologies and label positive for a mixture of molecular markers for neurons, glia, and neural progenitors. While not in every brain in our data set, we frequently found “balloon-like” cells in BRAFV600E transgene conditions, and these cells were positive for cux1, ctip2, GFAP, and but were negative for NeuN. There was no observed immunoreactivity to Vimentin or Nestin in balloon-like cells (data not shown). Additionally, those cells were negative for caspase-3 (data not shown), an apoptotic marker ^125^ and did not display DNA fragmentation and cellular membrane blebbing. However, we cannot exclude possibility that some of the balloon-like cells may undergo apoptosis or pyroptosis ^126^. In brains analyzed prior to P30 we found isolated balloon-like cells; however, by P30 aggregates of balloon-like cells were apparent (Figure 1B,1D). The observations indicate that BRAFV600 is sufficient to drive the development of balloon-like cells, that these cells can be generated in the lineages of either NESTIN+ or GLAST+ progenitors and appear in increasing numbers in the juvenile period, after P30. Those cells were not described in Koh et al.^14^

### 3.3 BRAFV600E causes increased astrocytogenesis and glial activation

Overactive signaling through the RAS-RAF-MEK-ERK pathway is known to increase astrocyte differentiation and proliferation ^127–130^. Consistent with this we found that BRAFV600E transgenesis in GLAST+ progenitors resulted in a significant increase in the number of astrocytes relative to neurons. In contrast, BRAFV600E transgenes introduced into NESTIN+ progenitors did not result in an appreciable number of astrocytes (Figure 3C). LNETs with BRAFV600E mutation has been shown to display high immunoreactivity to Glial Fibrillary Acidic Protein (GFAP) ^35^. Additionally, Koh et al. ^14^ showed increased immunoreactivity of glial lineage in GG patients and their mouse model. Consistent with this we found a significant increase in number of intensely positive GFAP cells in the regions of cortex containing BRAFV600E expressing neurons compared to control-FP and BRAFwt. We found that transgenesis of either NESTIN+ progenitors or GLAST+ progenitors with BRAFV600E resulted in comparable increases in the intensity of GFAP staining. Taken together this would suggest that BRAFV600E somatic mutations in proneuronal progenitors and in neurons is sufficient to result in elevated GFAP expression and potentially astrocyte activation (Figure 3A). GFAP immunoreactive cells were also positive for astrocytes marker Aldehyde Dehydrogenase 1 family member L1 (ALDH1L1) ^131^ (Figure 3B).

**Figure 3.**
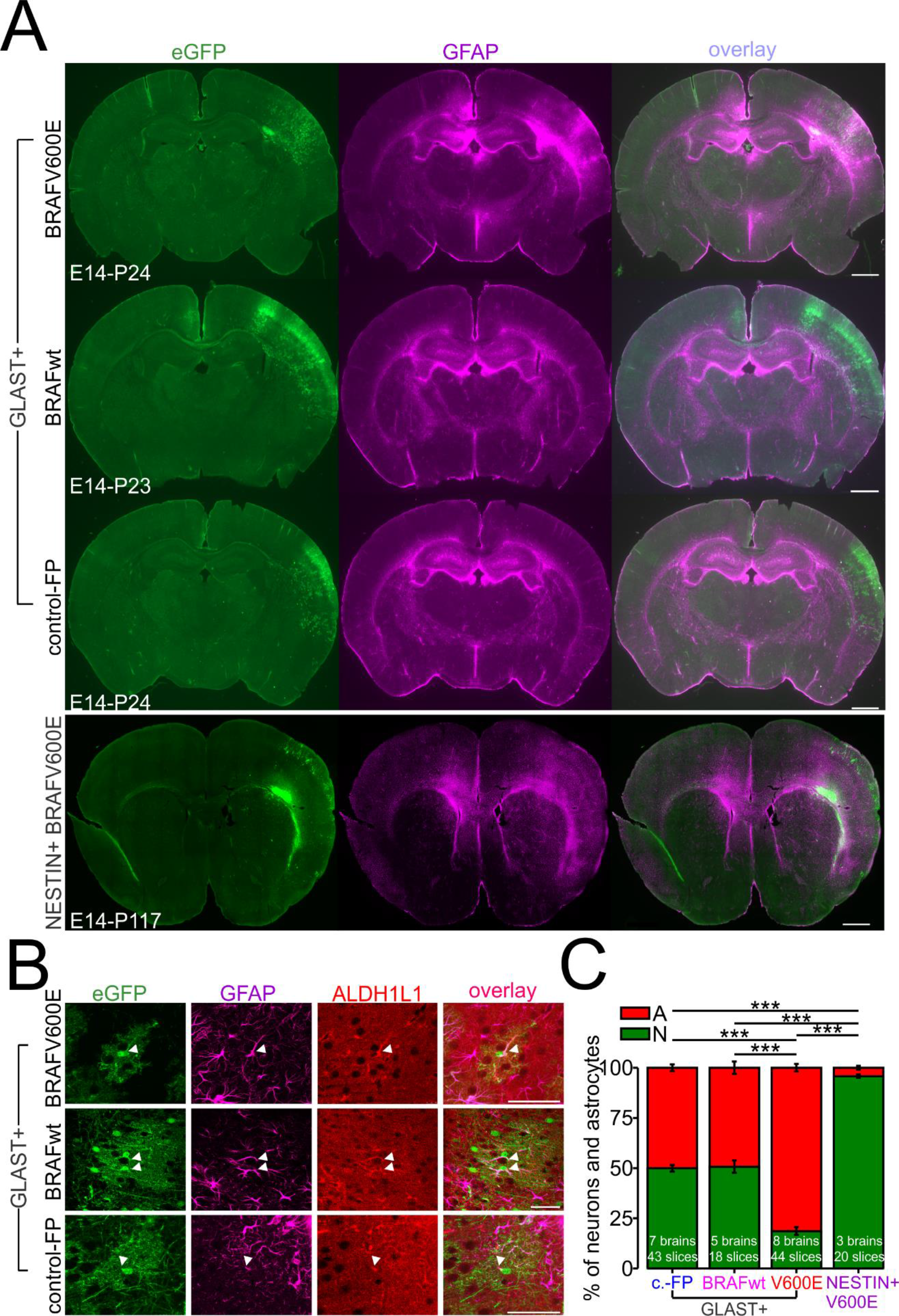
BRAFV600E transgene in GLAST+ neuroglial progenitors increases gliogenesis and induce reactive astrogliosis. **A.** Whole-slice image showing increased GFAP immunoreativity in somatosensory cortex transfected with GLAST+ BRAFV600E compared to GLAST+ control-FP and to GLAST+ BRAFwt transfected brains(upper 3 panels). Whole-slice image showing increased GFAP immunoreactivity in more frontal part of somatosensory cortex transfected with NESTIN+ BRAFV600E (lower panel). **B.** GFAP positive cells were also immunopositive to astrocytes marker ALDH1L1(white arrowheads). **C.** Neuron to astrocytes percent ratio showing increased astrogliosis (EGFP positive cells) in BRAFV600E electroporated murine cortical slices. Due to non-homogenous variance (Levene test (3,243) =4.574 and 4.575 for neurons and astrocytes respectively p=0.004), Welch test F(3, 106.49)=183.71, p<0.001 with Games-Howell posthoc correction was used for statistical comparison. Astrocytes percentage was increased in Glast+ BRAFV600E electroporated slices (n=44 slices, 8 brains) compared to Glast+ control-FP only (n=43 slices, 7 brains p<0.001); and compared to Glast+ BRAFwt electroporated slices (n=18 slices, 5 brains p<0.001). And neuronal percentage was decreased in Glast+ BRAFV600E electroporated slices compared to Glast+ control-FP (p<0.001); and compared to Glast+ BRAFwt electroporated slices (p<0.001); but not in Glast+ BRAFwt electroporated slices compared to Glast+ control-FP electroporated slices for both astrocytes and neurons percentage (p=0.996). Astrocytes percentage was decreased in Nestin+ BRAFV600E (n=20 slices, 3 brains) compared to Glast+ BRAFwt (p<0.001); to Glast+ control-FP (p<0.001); to Glast+ BRAFV600E (p<0.001). Neuronal percentage was increased in Nestin+ BRAFV600E compared to Glast+ BRAFwt (p<0.001); to Glast+ control-FP (p<0.001); and to Glast+ BRAFV600E (p<0.001). Scale bars for A(upper panel) 1 mm;(lower panel) 500 μm. B.- 50 μm; * - p<0.05,** - p<0.01, *** - p<0.0001. Error bars are ±2SEM.

To determine whether the elevated intensity of GFAP staining we observed was due to reactive gliosis and potential inflammatory responses in the regions of mutation bearing cells, we performed an RNA-seq experiment to compare the gene expression profiles of patches of cortex containing BRAFV600E, BRAFwt, or mRFP transgenes. We estimate that approximately 5-10% of cells are transfected cells bearing transgenes in a cortex ^78, 79^, and so the majority of any change in transcript is likely driven by changes in gene expression profiles in untransfected reacting cells. Using an unsupervised hierarchical clustering analysis of all genes in 12 samples, 4 in each transgene condition, we found that BRAFV600E, control-FP, and BRAFwt conditions clustered separately, except for one BRAFV600E sample which clustered with BRAFwt conditions (Figure 4A). Differential expression and gene ontology analysis indicated a significant increase in the expression of genes in the inflammatory immune response pathway (H2-Aa, CD74, H2-Ab1, CD48, CD109, Cxcl16, Ccr1), and classic complement pathway components (C3, Serpinf1, C4b, C1s1, C1ra, Serpina3i, Serpina3b, Serpina3n) (Figure 4B,C,D,E,G; Table S1). The Iba1, a microglia marker was also increased in GLAST+ BRAF V600E condition compared to control-FP, it was also increased compared to BRAFwt at p=0.011 level. A marker of microglia activation HLA-DR(CD74) was significantly increased in GLAST+ BRAFV600E compared to BRAFwt and to control-FP (Figure 4C,D). Similarly, markers of astrocyte activation, GFAP and Vimentin, were also significantly upregulated in the RNAseq profiles of the four BRAFV600E samples relative to the other transgene conditions. Overall, the pattern of gene expression changes in cortical tissue containing a subpopulation of cells expressing BRAFV600E is consistent with these cells causing a glial activation and neuroinflammatory response (Figure 4G). It is also consistent with previous studies showing increased inflammatory immune response and complement pathway activation in ganglioglioma, and, in tissue resected from epilepsy patients with tuberous sclerosis ^14, 68, 69^. Interestingly, the decreased expression of potassium channels (Figure 4G) is consistent with previous study by Aronica et al. ^69^ and Koh et al. ^14^. Additional ontology analysis with Gene Analytics web tool showed significant enrichment of genes associated with Tuberous Sclerosis (Figure 4H).

**Figure 4.**
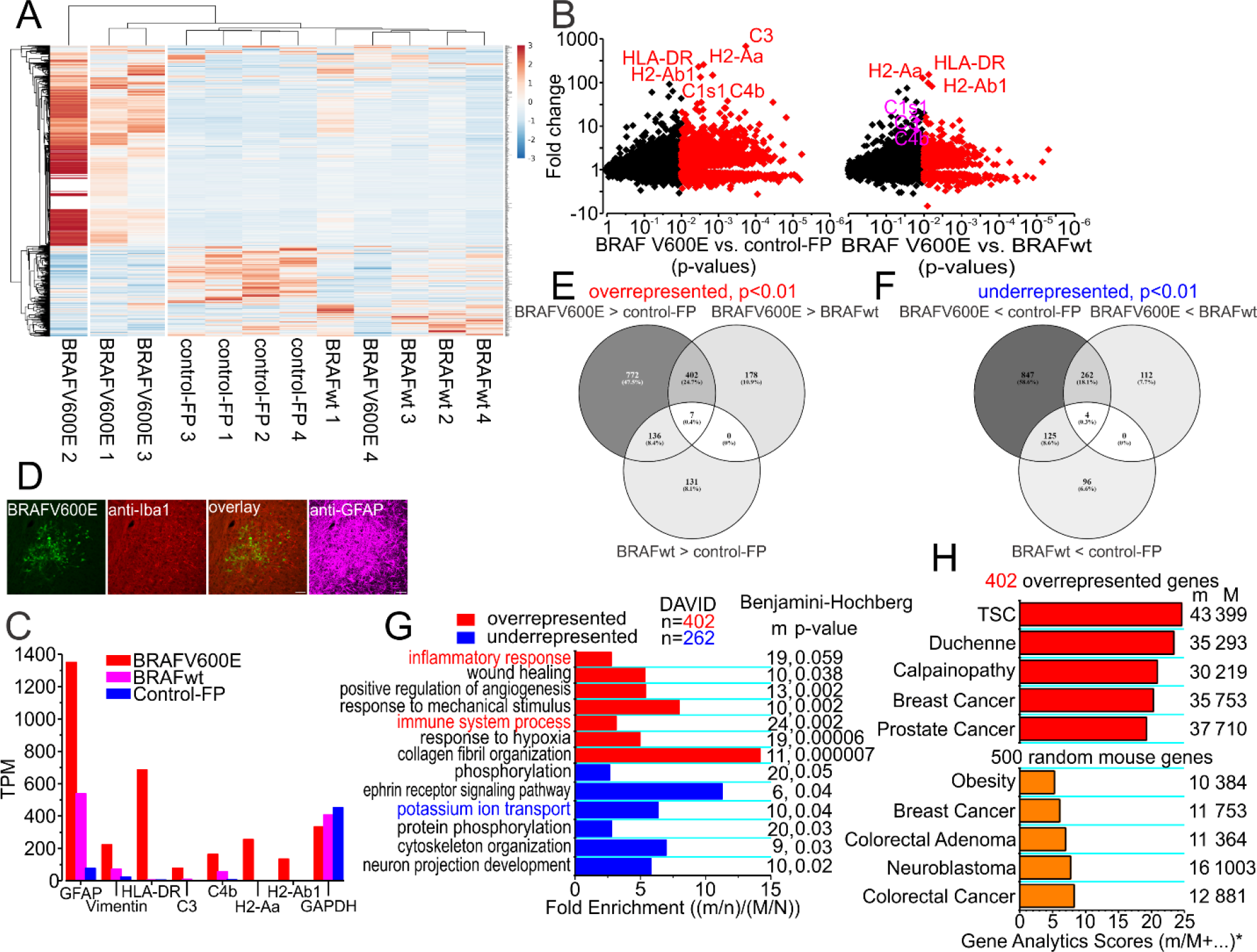
Unsupervised Hierarchical Clustering Analysis of GLAST+ BRAFV600E, GLAST+ control-FP and GLAST+ BRAFwt tissue-wide expression profile. **A.** Four clusters of three conditions with four replicates each ^100^, GLAST+ BRAFV600E, GLAST+ control-FP, GLAST+ BRAFwt, 742 genes with at least log_2_ fold change =1 were used. **B.** Scatter plot for BRAFV600E to control-FP showing fold change and p-values, data with p<0.01 is in red and was used for further functional enrichment analysis. Some of the genes with the highest fold change are shown. Scatter plot for BRAFV600E to BRAFwt showing fold change and p-values, data with p<0.01 is in red and was used for further functional enrichment analysis. Some of the genes with the highest fold change are shown. **C.** Expression vallues in transcripts per million (TPM) of GFAP and vimentin in three conditions; GFAP is increased in BRAFV600E (20.78 fold increase, p=0.000197, FDR=0.014) and BRAFwt (7.99 fold increase, p=0.0166, FDR=0.32) compared to control-FP (Table S1 and S3); BRAFV600E to BRAFwt (2.67 fold increase p=0.03066, FDR=0.185; Table S2); vimentin was also increased in BRAFV600E compared to control-FP (11.07 fold increase p=0.00008, FDR=0.014) and to BRAFwt (3.63 fold increase p=0.004, FDR=0.131); BRAFwt to control-FP (3.13 fold increase p=0.025, FDR=0.35); HLA-DR(CD74), a microglia marker was significantly increased in BRAFV600E compared to control-FP (231.76 fold change p=0.004, FDR=0.03); and in BRAFV600E compared to BRAFwt (154.39 fold change p=0.007, FDR= 0.136); C3 was significantly increased in BRAFV600E neurons compared to control-FP (681.23 fold change p= 0.0002, FDR= 0.014); BRAFV600E to BRAFwt (8.23 fold change p=0.015, FDR=0.15); BRAFwt to control-FP (84.93 fold change p=0.028, FDR=0.356); C4b was significantly increased in BRAFV600E compared to control-FP (37.75 fold change p=0.00057, FDR=0.016); H2-Aa, MHC II protein, was significantly increased in BRAFV600E compared to control-FP (258.75 fold change p=0.0025, FDR=0.027); and in BRAFV600E compared to BRAFwt (128.49 fold change p=0.011, FDR=0.14); H2-Ab1, also MHC II protein was significantly increased in BRAFV600E compared to control-FP (134.29 fold change p=0.0031, FDR=0.029); in BRAFV600E compared to BRAFwt (82.69 fold change p0.011, FDR=0.13); while GAPDH was insignificantly changed, BRAFV600E to control-FP (1.10 fold decrease p=0.025, FDR=0.175), to BRAFwt (1.03 fold decrease p=0.52, FDR=0.718); BRAFwt to control-FP (1.13 fold decrease p=0.31, FDR=0.7). **D.** Represantative microglia in BRAFV600E immunoreactive to Iba1 mixed with balloon-like cells. **E.** Venn diagram of all the upregulated genes with at least log_2_ fold change=1, and p<0.01. **F.** Venn diagram of all the downregulated genes with at least log_2_ fold change=1, and p<0.01. **G.** Fold enrichment from david functional annotation analysis of 402 overrepresented genes in BRAFV600E compared to both control-FP and BRAFwt and 262 genes underrepresented in BRAFV600E compared to both control-FP and BRAFwt. m – number of specific biological process associated genes out of 402 genes (n), M-total number of genes associated with specific biological process, N-total number of genes. **H.** Gene analytics analysis of 402 overrepresented genes in BRAFV600E compared to both control-FP and BRAFwt (upper panel). Same analysis of 500 random mouse genes. The scores are based on m/M ratio, on significantly differentially 200 upregulated or 200 downregulated genes in disease tissues from gene expression omnibus (GEO) database, or literature with at least 2-fold change and p<0.05, and genetic association to the disease based on several MalaCards data sources (ClinVar, OMIM, Orphanet, Uniprot, GeneTest), and the GeneCards-inferred relation to the disease with more frequently mentioned genes having higher scores. For each gene, the maximal score of all the above mentioned possible scores is used as the final gene score. The disease score is based on the final scores of all the matched genes. Scale bar 50 μm in D.

### 3.4 BRAFV600E increases excitability of pyramidal neurons

To test the hypothesis that neurons with BRAFV600E mutation have increased excitability we performed whole-cell patch clamp recording from pyramidal neurons in upper layers 2/3. We recorded from neurons in all three transgene conditions and in both transfected and in neighboring neurons not positive for fluorescent markers of transgenesis. In current-clamp recordings we found that BRAFV600E neurons displayed significantly higher action potential (AP) firing frequencies to 1 second depolarizing current pulses (Figure 5A upper panel, 5B, p<0.001 for 20-300 pA current steps). This significantly increased firing rate was true for neurons from both the NESTIN+ and GLAST+ progenitor populations. Neither BRAFwt nor neighboring untransfected neurons in BRAFV600E conditions showed elevated firing frequencies above fluorescent protein transfected controls (control-FP).

**Figure 5.**
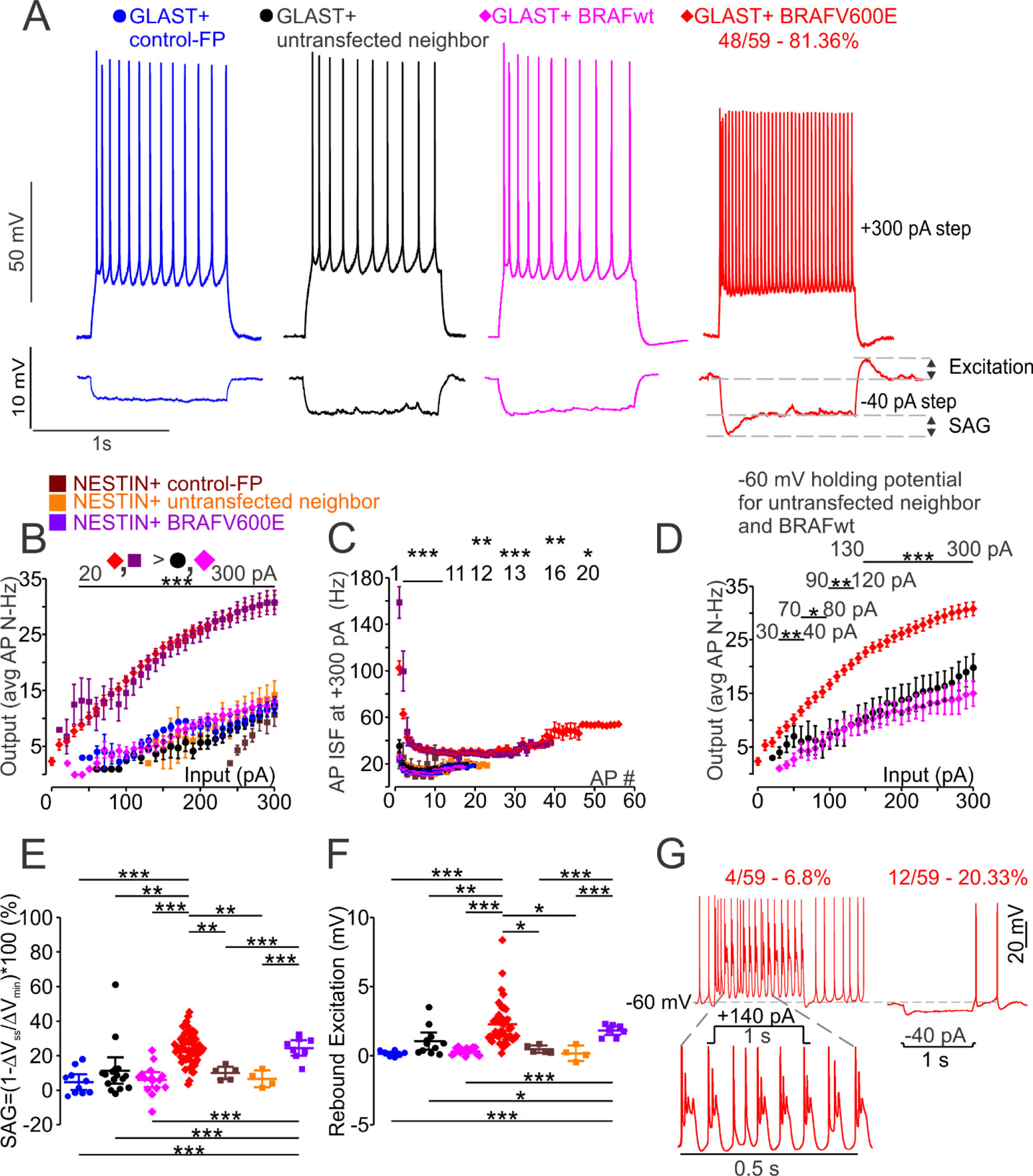
*BRAFV600E* expressing *neurons are hyperexcitable*. **A.** Represantative traces of four GLAST+ experimental conditions (upper panel). Response to −40 pA 1 sec current step showing SAG ratio and rebound excitation measurement (lower panel). **B.** Input-Output curve shows more than 2 times higher AP frequency firing in GLAST+ BRAFV600E transfected neurons (n=54 T= range of 4.37-6.73, p<0.001) and NESTIN+ BRAFV600E transfected neurons (n=8 T= range of 5.53-9.64, p<0.001) compared to all other conditions, GLAST+ untransfected neighbor (n=8), GLAST+ control-FP (n=11), GLAST+ BRAFwt (n=13), NESTIN+ control-FP (n=3, 240-300 pA), and NESTIN+ untransfected neighbor (n=5, 130-300 pA). For statistical comparison for 10 pA - 150 pA steps, due to significant difference in variances (Levene’s test (18, 149) = range of 8.816-1.68, p<0.001, p=0.013 for 140 pA and p=0.05 for 150 pA) Welch test with Games-Howell post-hoc correction was used; for 160-300 pA one-way ANOVA F(18, 150) = 13.42-16.55 with Tukey post-hoc correction was used. Only cells with 7 and more APs at 300 pA 1 sec current step are chosen for the comparison. **C.** Instantaneous frequency (ISF) of APs in the train at 300 pA 1 second depolarizing current step was significantly higher in GLAST+ BRAFV600E transfected neurons compared to all other conditions (p<0.001). AP ISF at +300 pA 1 second step due to nonhomogeneous variance for AP #1 and AP #2, Levene test (8, 132)=5.08 and 3.97 respectively, Welch test (8, 22.91 and 23.18)=27.79 and 15.82 respectively with Games-Howell posthoc correction was used, from AP #3 to AP #20 One-way ANOVA F(7-8, 82-132)=2.24-19.6, p<0.001, with Tukey posthoc correction was used; there was significant difference in ISF between GLAST+ BRAFV600E neurons (n=54) to NESTIN+ untransfected neighbor (n=5, p<0.001) and NESTIN+ control-FP (p<0.001); NESTIN+ BRAFV600E (n=8) to their untransfected neighbor (p<0.001) and NESTIN+ control-FP (p<0.001). **D.** Input-Output curve for GLAST+ BRAFV600E, GLAST+ BRAFwt held at −60 mV (n=4), and GLAST+ untransfected neighbor neurons (n=5) held at −60 mV shows still significant difference in AP firing frequency. Due to non-homogenous variance for 30 and 40 pA steps Levene (2, 53)=5.5 and 4.41, p=0.007 and p=0.017 Welch test with Games-Howell post-hoc correction was used; for steps 50-300 pA One-way ANOVA with Tukey post-hoc correction was used. For 30-40 pA steps Welch F(2,10.85) = 10.43, p=0.003 and (2, 10.22)=9.78, p=0.004 respectively. For 70-300 pA steps ANOVA F(2, 54)=3.82 – 13.84, p=0.028 and p=0.017 for 70 and 80 pA steps, p=0.007-0.001 for 90-120 pA steps; p<0.001 for 130-300 pA steps. **E.** One-way ANOVA with Tukey post-hoc correction of SAG ratio values in different conditions shows larger SAG in most of the recorded GLAST+ BRAFV600E (n=58) transfected neurons, F(3,105) = 35.98 (GLAST+ BRAFV600E to GLAST+ control-FP (n=15) – p<0.001; untransfected neighbor neurons (n=18) – p<0.01, GLAST+ BRAFwt (n=17) – p<0.001). SAG ratio was significantly larger in GLAST+ BRAFV600E (n=58) compared to NESTIN+ control-FP (n=5 T=2.94, p=0.005); to NESTIN+ untransfected neighbor (n=4 T=3.3, p=0.002); NESTIN+ BRAFV600E (n=8) to NESTIN+ control-FP (n=5 T=4.48, p<0.001); NESTIN+ BRAFV600E to their untransfected neighbor (T=4.87, p<0.001); NESTIN+ BRAFV600E to GLAST+ BRAFwt (T=5.53 p<0.001); NESTIN+BRAFV600E to GLAST+ untransfected neighbor (T=7.3 p<0.001); NESTIN+ BRAFV600E to GLAST+ control-FP (T=5.8 p<0.001). **F.** Due to non-homogenous variances (Levene’s test (3, 91) = 7.61, p<0.001) Welch test with Games-Howell post-hoc correction was used for statistical comparison of rebound excitation values and shows larger rebound excitation in GLAST+ BRAFV600E (n=42) transfected neurons compared to all non-BRAFV600E GLAST+ conditions Welch (3, 42.36) = 20.83, p<0.001; GLAST+ control-FP (n=14, p<0.001), and GLAST+ untransfected neighbor neurons (n=18, p<0.001), GLAST+ BRAFwt (n=17, p<0.001). Rebound excitation was larger in GLAST+ BRAFV600E neurons (n=42) compared to NESTIN+ untransfected neighbor (n=4 T=3.3, p=0.010); to NESTIN+ control-FP (n=5 T=2.94, p=0.013); it was increased in NESTIN+ BRAFV600E (n=8) compared to their untransfected neighbor (n=4 T=5.514, p<0.001); to NESTIN+ control-FP (n=5 T=6.035, p<0.001).BRAFV600E neurons with rebound APs were omitted from statistical comparison. **G.** Left upper panel - representative whole-cell current-clamp recording traces of GLAST+ BRAFV600E transfected neuron showing depolarization waves with smaller spikes riding on top of them following each full size AP (4 out of 58 neurons, 6.9%), this bursting was not observed in control conditions or BRAFwt condition. Left lower panel – zoomed in depolarization waves. Right panel – representative trace of rebound AP observed in 12 out of 58 (20.7%) GLAST+ BRAFV600E transfected neurons. * - p<0.05, ** - p<0.01, *** - p<0.001. Error bars are ±SEM for B, C, and D; ±2SEM for E-F.

To compare the effect of developmentally induced chronic overactivation of RAF-RAS-ERK pathway on neuronal electrophysiology to the effect of chronic overactivation of mTOR pathway we’ve performed whole-cell patch clamp experiments in cortical slices transgenic for GLAST+ PIK3CA E545K ^3^ and CRISPR-Cas9 induced mutation in TSC1 gene ^73^. Phosphatidylinositol-4,5-Bisphosphate 3-Kinase Catalytic Subunit Alpha and TSC1 are key regulatory upstream components of mTOR. Substitution of glutamic amino acid to lysine (E545K) in PIK3CA is a “hot spot” mutation and is found in FCD and cause constitutive activation. Disruption of TSC1/TSC2 complex that inhibits activation of mTOR is associated with Tuberous Sclerosis and was found in FCD too. GLAST+ progenitors were transfected with PIK3CA E545K on PiggyBac transposon background using IUE at E14-E15 in the same way as BRAFV600E, and for CRISPR-Cas9 TSC1 guide-RNA we used T4 from Lim et al. 2017. The AP firing frequency, AP ISF, rheobase, RMP and Rin was closer to control conditions in previous experiments (Figure 6A upper panel, 6B, C, D, F, G). However, AP voltage threshold was not different from GLAST+ BRAFV600E neurons. This suggest that alterations in neuronal electrophysiological properties affected differently by pathological mutations in mTOR pathway key protein components. Given those findings we decided not to pursue further inquiry into PIK3CA E545K and CRISPR-Cas9 TSC1 KD conditions.

**Figure 6.**
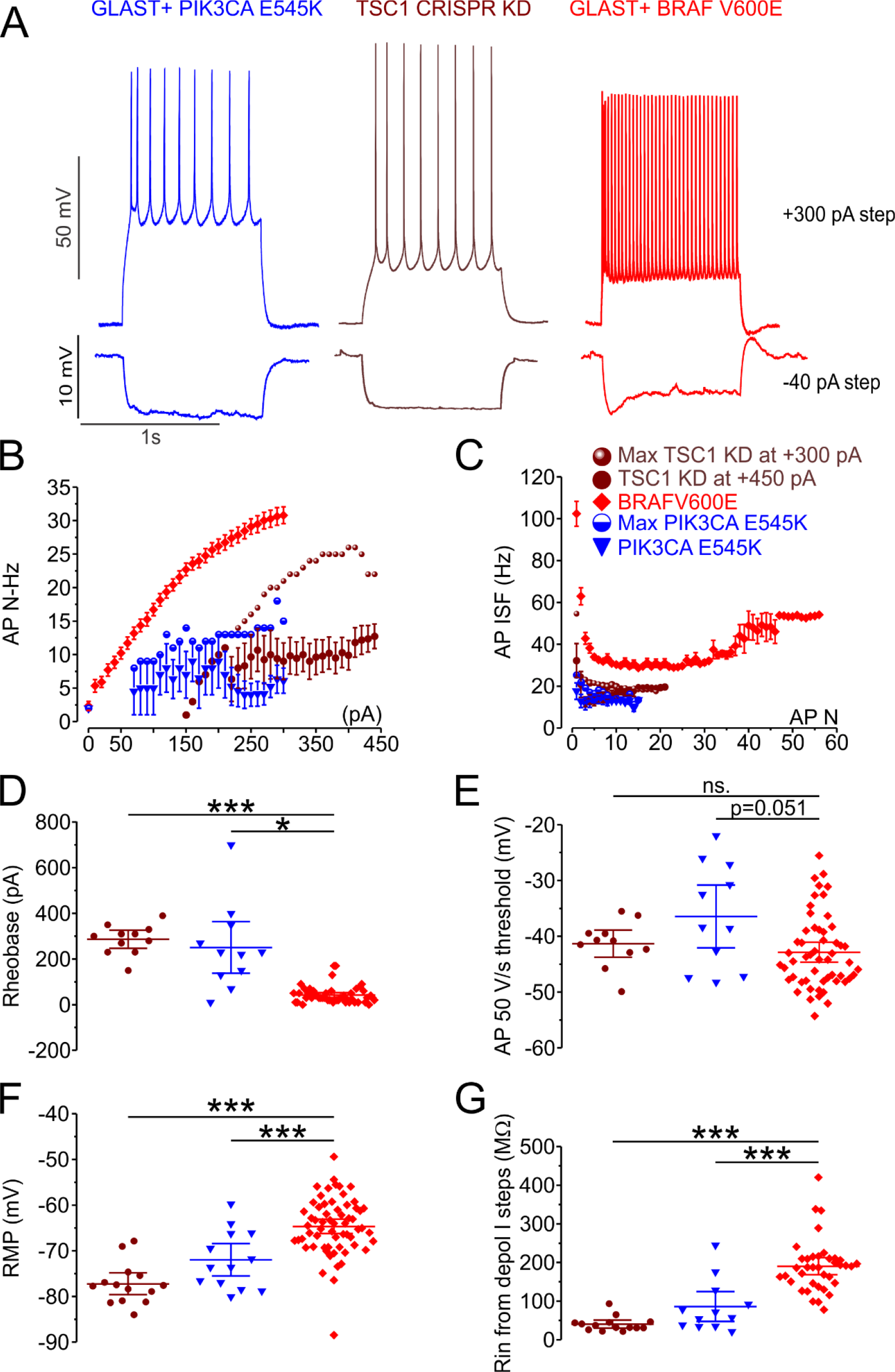
Differential effect of three experimental manipulations on whole-cell current-clamp properties. **A.** Representative traces from GLAST+ neurons expressing PIK3CA E545K (blue) and BRAFV600E mutations (red), and CRISPR knockdown of TSC1 gene showing AP firing at +300 pA (upper panel), and membrane potential response to hyperpolarizing current step of −40 pA. **B.** Average AP firing frequency in all three conditions, BRAFV600E (n=54), TSC1 KD (n=11), PIK3CA E545K (n=6). Cells with maximal values are shown for PIK3CA E545K (half-filled blue circles), and for TSC1 KD (brown spheres). **C.** AP instantaneous frequency at +300 pA current step except TSC1 KD, which is shown for +450 pA current step with maximal value cell shown for +300 pA current step (brown spheres). **D.** Rheobase for all three conditions was compared with Welch test (2, 14.38) =72.98 due to nonhomogeneous variance (Levene test (2, 68)=15.36, p<0.001), together with Games-Howell posthoc correction BRAFV600E to TSC1 KD (p<0.001), BRAFV600E to PIK3CA E545K (p=0.011, due to small sample for PIK3CA E545 student T=3.68, p=0.004 was used),. **E.** AP 50 V/s voltage threshold is not different compared with Welch (2, 19.59) =2.48, p=0.11) together with Games-Howell BRAFV600E to TSC1 KD (p=0.57), BRAFV600E to PIK3CA E545K (p=0.12); Levene test (2, 73) =4.93, p=0.01. **F.** Resting Membrane Potential (RMP recorded before application of current steps) was compared with One-way ANOVA F(2,86) =28.72, together with Tukey posthoc corrections BRAFV600E to TSC1 KD (p<0.001), and BRAFV600E to PIK3CA E545K (p<0.001). **G.** Input resistance (Rin) from depolarizing current steps (due to Ih activation in BRAFV600E) was compared with Welch (2, 25.70) =74.48, due to nonhomogeneous variance (Levene test (2, 61) =3.40, p=0.04), together with Games-Howell posthoc correction BRAFV600E to TSC1 KD (p<0.001), and BRAFV600E to PIK3CA E545K (p<0.001). * - p<0.05, ** - p<0.01, *** - p<0.001.

Instantaneous AP frequency (ISF) measured at +300 pA 1 second current step was significantly higher in BRAFV600E neurons (Figure 5C). In addition, in 4 out of 59 GLAST+ BRAFV600E neurons and in 1 out of 9 NESTIN+ BRAFV600E neurons we observed an unusual bursting pattern and post-action potential depolarization waves that were not observed in any of the non-BRAFV600E conditions (Figure 5G left panel). The number of neurons with those events was increased when recording potassium currents with Co^2+^ (1 mM) substituting Ca^2+^ (1 mM) in aCSF solution (5 out of 24). In addition to the AP firing at a significantly higher frequency BRAFV600E neurons also had lower rheobase (n=49), minimal depolarizing current step required to elicit first AP, and a lower voltage threshold to fire action potentials (Figure 7B, C, E). Passive membrane properties were also significantly different in BRAFV600E neurons. The resting membrane potential (RMP) was more depolarized in BRAFV600E neurons (n=61, −64.91 ± 0.76 mV) compared to untransfected neighbor neurons (n=23, −73.33 ± 1.55 mV, p<0.001), and input resistances measured to hyperpolarizing and depolarizing current pulses were significantly increased in BRAFV600E neurons (n=61) compared to all other non-BRAF V600E conditions (Figure 7F, G, H). The elevated resting membrane potential did not explain the increased firing rates in BRAFV600E neurons, as untransfected neighboring neurons (n=5) did not achieve AP firing frequencies similar to BRAFV600E neurons when depolarized to −60 mV, and similarly the few BRAFV600E neurons with more negative resting membrane potentials (n=14, average RMP=−70.66 ± 0.58 mV) generated high frequency trains of action potentials similar to more depolarized BRAFV600E neurons. Also, subthreshold input resistances did not correlate significantly with action potential frequencies in either BRAFV600E or control neurons. Taken together, BRAFV600E transgenes significantly alter the electrophysiological properties of pyramidal neurons making them more excitable.

**Figure 7.**
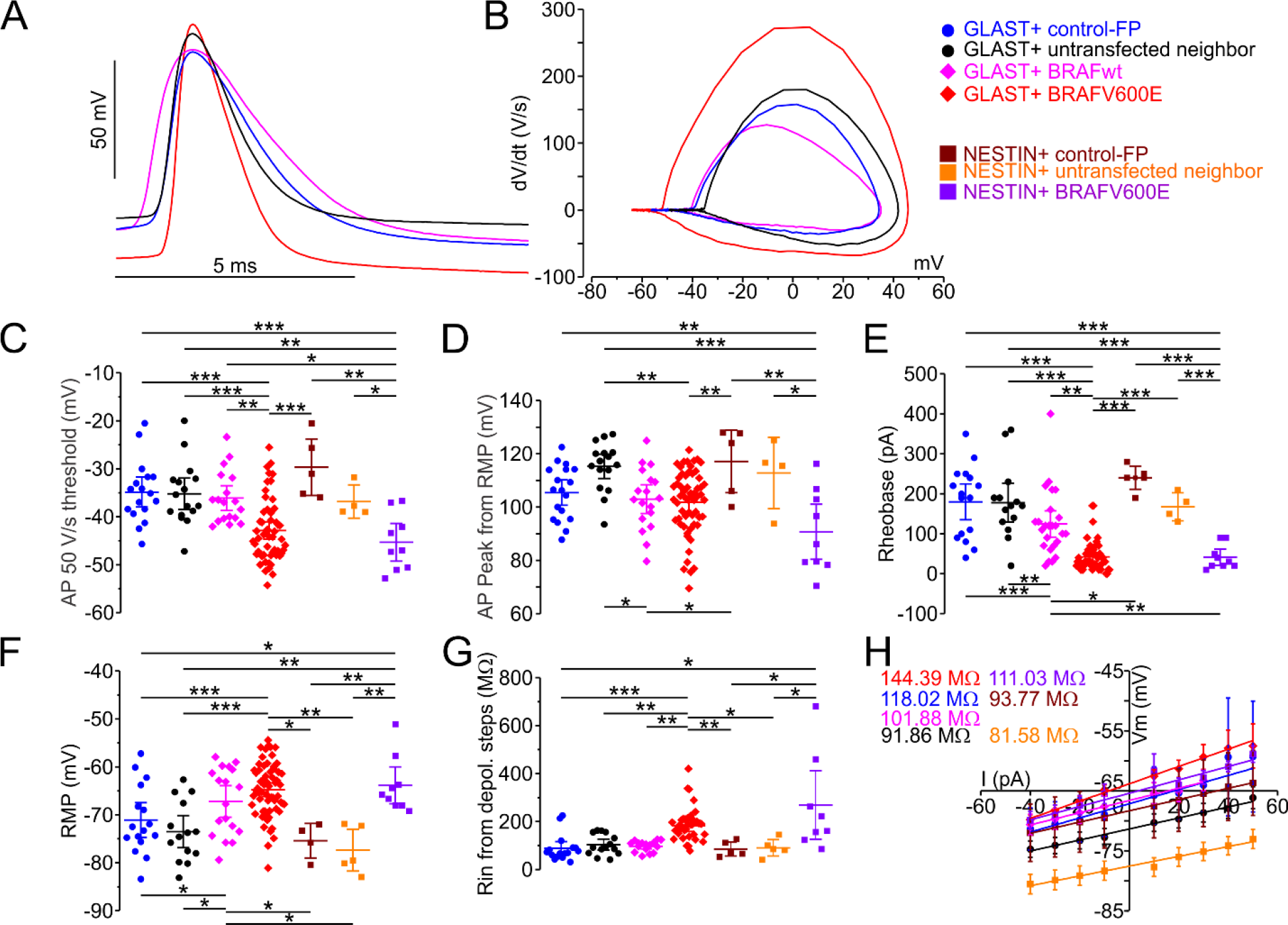
Properites of first action potential at rheobase are altered in GLAST+ and NESTIN+ BRAFV600E expressing neurons. **A.** Representative first APs at rheobase from GLAST+ BRAFV600E, control-FP, untransfected-neighbor, BRAFwt all the cells had similar RMPs (BRAFV600E - −72.87 mV, BRAFwt - −72.32 mV, control-FP - −72.48 mV, untransfected neighbor - −72.31 mV). **B.** First order derivative over time (dV/dt) of representative APs (phase-space plot) from A. showing hyperpolarised AP voltage threshold in BRAFV600E transfected neurons. **C.** One-way ANOVA F(6, 116)=9.72, p<0.001 with Tukey post-hoc correction showed that AP voltage threshold at 50 V/s was more hyperpolarized in GLAST+ BRAFV600E transfected neurons (n=54) compared to GLAST+ control-FP (n=18, p<0.001), and untransfected neighbor (n=16, p<0.001); GLAST+ BRAFwt (n=17, p=0.007) neurons; NESTIN+ control-FP (n=5, p<0.001); it was lower but not statistically significant compared to NESTIN+ BRAFV600E untransfected neighbor (n=4 T=1.81, p=0.076). AP voltage threshold at 50 V/s was significantly more hyperpolarized in NESTIN+ BRAFV600E neurons (n=9) compared to NESTIN+ control-FP neurons (n=5 p<0.001); to NESTIN+ BRAFV600E untransfected neighbor (n=4 T=2.63, p=0.023); to GLAST+ untransfected neighbor (p=0.001); to GLAST+ control-FP (p<0.001); to GLAST+ BRAFwt (p=0.012). **D.** One-way ANOVA F(6, 122) =6.29, p, 0.001 with Tukey post-hoc correction showed that AP peak measured from RMP was larger in GLAST+ untransfected neighbor neurons (n=16) compared to GLAST+ BRAFV600E (n=58, p=0.001) and to GLAST+ BRAFwt transfected neurons (n=18, p=0.04); to NESTIN+ BRAFV600E (n=9, p<0.001). GLAST+ control-FP (n=18) to NESTIN+ BRAFV600E (p=0.039). NESTIN+ BRAFV600E to NESTIN+ untransfected neighbor (n=4, p=0.033); to NESTIN+ control-FP (n=5, p=0.002). **E.** Due to non-homogenous variance – Levene test (6, 114) =5.55, p<0.001 Welch test with Games-Howell post-hoc correction was used to compare rheobase between experimental conditions, which showed lower current required to fire AP in GLAST+ BRAFV600E transfected neurons (n=49) Welch test (6, 20.94) =36.71, p<0.001, compared to control-FP (n=19, p<0.001), and untransfected neighbor neurons (n=17, p<0.001), to GLAST+ BRAFwt (n=18, p<0.001). Rheobase was significantly lower in GLAST+ BRAFV600E (n=49) compared to NESTIN+ control-FP (n=5 p<0.001); to NESTIN+ untransfected neighbor (n=4 T=6.46, p<0.001); NESTIN+ BRAFV600E neurons (n=9) to NESTIN+ control-FP (n=5 T=11.52, p<0.001); to their untransfected neighbor (n=4 T=6.66, p<0.001); to GLAST+ untransfected neighbor (p<0.001); to GLAST+ control-FP (p<0.001); to GLAST+ BRAFwt (p=0.004). **F.** GLAST+ BRAFV600E transfected neurons (n=61) had more depolarized resting membrane potential compared to GLAST+ control-FP neurons (n=20, p<0.001), and to their untransfected neighbor neurons (n=23, p<0.001); to NESTIN+ untransfected neighbor (n=5, p=0.001); to NESTIN+ control-FP (n=5, p=0.031). GLAST+ BRAFwt transfected neurons (n=18) had more depolarized RMP compared to untransfected neighbor neurons (p=0.044); to GLAST+ control-FP (p=0.050); to NESTIN+ untransfected neighbor (p=0.034); to NESTIN+ control-FP (T=2.25 p=0.036). NESTIN+ BRAFV600E to GLAST+ control-FP (p=0.033), and to their untransfected neighbor (p=0.005). One-way ANOVA F(6, 133) = 9.59, p<0.001 with Tukey post-hoc correction test was used for statistical comparison. In case of small n student t-test was used. RMP measured in current clamp before beginning of steps protocol. **G.** Due to nonhomogeneous variance (Levene test (6, 112) =5.58, p<0.001) Welch test (6, 24.56) =7.04, p<0.001 with Games-Howell post-hoc correction was used for input resistance comparison. Averaged input resistance (Rin) as a function of membrane potential response (Vm) to depolarizing current (I) steps was significantly larger in GLAST+ BRAFV600E transfected neurons (n=39) compared to GLAST+ control-FP (n=21, p<0.001), and their untransfected neighbor neurons (n=23, p=0.009); to GLAST+ BRAFwt (n=18, p=0.001). GLAST+ BRAFV600E to NESTIN+ untransfected neighbor (n=5, p=0.018); to NESTIN+ control-FP (n=5, p=0.003). Average Rin from depolarizing current steps was significantly larger in NESTIN+ BRAFV600E (n=8) compared to GLAST+ control-FP (n=21 T=2.496, p=0.040); to NESTIN+ untransfected neighbor (n=5 T=2.418, p=0.043); to NESTIN+ control-FP (n=5 T=2.512, p=0.038). **H.** Linear fit of averaged membrane potential responses to current step protocol from −40 to +50 pA 1 second pulse with 10 pA increment shows a different input resistance in between the recorded conditions. * - p<0.05, ** - p<0.01, *** - p<0.001. Error bars are ±SEM for A. and H, ±2SEM for C-G.

### 3.5 BRAFV600E decreases delayed rectifier potassium currents

Since the combined increased AP firing frequency, SAG ratio, rebound excitation, more depolarized resting membrane potential and higher input resistance were observed only in BRAFV600E expressing neurons, and not in any non-BRAFV600E neurons to test alterations in what ionic conductances make BRAFV600E transgenic neurons hyperexcitable we have performed whole-cell voltage clamp in GLAST+ BRAFV600E neurons and their untransfected neighbors only. Recorded calcium currents did not show any significant difference (data not shown). Recording of potassium currents showed a decreased sustained current sensitive to 25 mM TEA, a known potassium channel inhibitor, measured at the last 100 ms of 500 ms depolarizing voltage pulses across examined range of voltage steps in GLAST+ BRAFV600E neurons (n=18) compared to their untransfected neighbors (n=8, p<0.01; Table 1, 2), preserving kinetic properties of activation (Figure 8A, B, C, D, F).

**Table 1.**
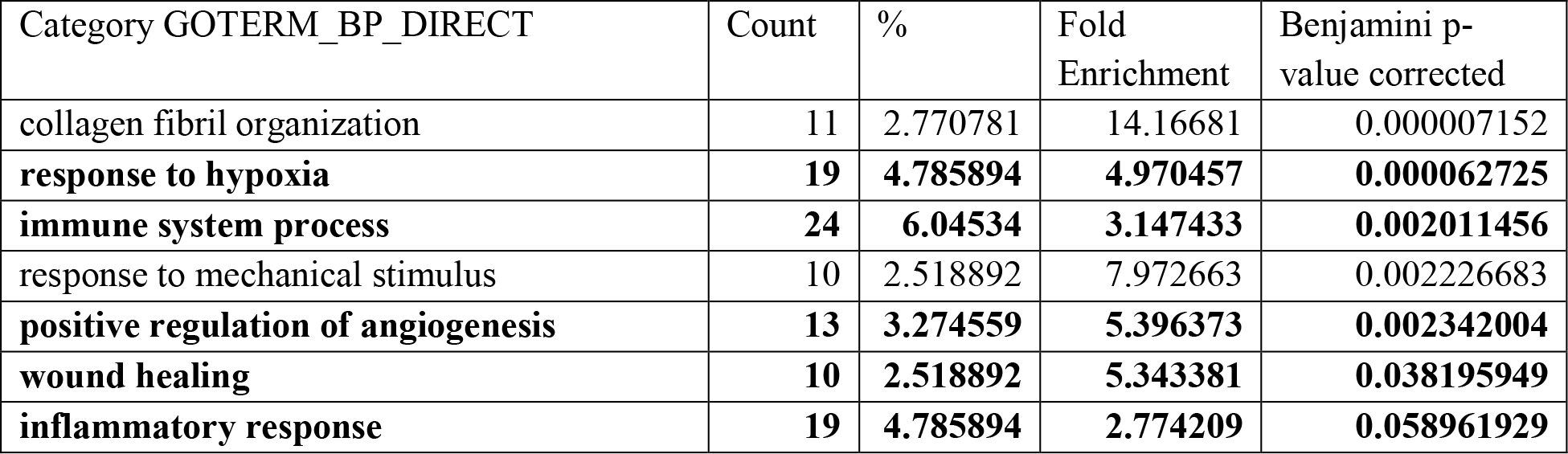
GOTERM Biological protein production pathways enrichment in GLAST+ BRAFV600E compared to control-FP and to GLAST+ BRAFwt from 402 upregulated genes at p<0.01

| Category GOTERM_BP_DIRECT | Count | % | Fold Enrichment | Benjamini p-value corrected |
| --- | --- | --- | --- | --- |
| collagen fibril organization | 11 | 2.770781 | 14.16681 | 0.000007152 |
| **response to hypoxia** | **19** | **4.785894** | **4.970457** | **0.000062725** |
| **immune system process** | **24** | **6.04534** | **3.147433** | **0.002011456** |
| response to mechanical stimulus | 10 | 2.518892 | 7.972663 | 0.002226683 |
| **positive regulation of angiogenesis** | **13** | **3.274559** | **5.396373** | **0.002342004** |
| **wound healing** | **10** | **2.518892** | **5.343381** | **0.038195949** |
| **inflammatory response** | **19** | **4.785894** | **2.774209** | **0.058961929** |

**Table 2.** GOTERM Biological protein production pathways enrichment in GLAST+ BRAFV600E compared to GLAST+ control-FP and to GLAST+ BRAFwt from 262 downregulated genes at p<0.01.

| Category GOTERM_BP_DIRECT | Count | % | Fold Enrichment | Benjamini p-value corrected |
| --- | --- | --- | --- | --- |
| **neuron projection development** | **10** | **3.802281** | **5.807425** | **0.023973** |
| **cytoskeleton organization** | **9** | **3.422053** | **6.985663** | **0.027481** |
| protein phosphorylation | 20 | 7.604563 | 2.802889 | 0.031236 |
| **potassium ion transport** | **10** | **3.802281** | **6.356159** | **0.035131** |
| ephrin receptor signaling pathway | 6 | 2.281369 | 11.2637 | 0.044375 |
| phosphorylation | 20 | 7.604563 | 2.638014 | 0.044999 |

**Figure 8.**
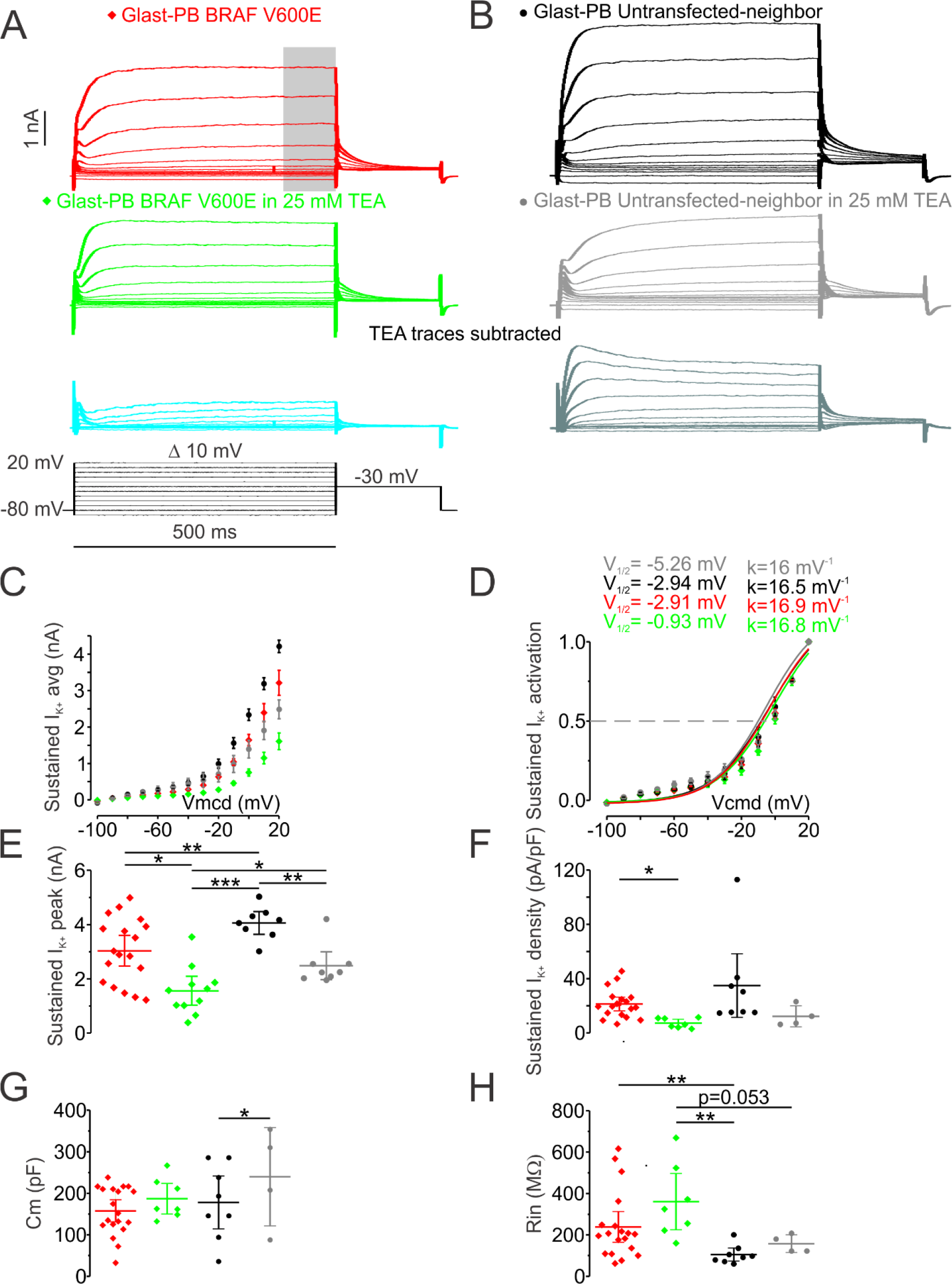
Sustained potassium currents are decreased in GLAST+ BRAFV600E neurons compared to their untransfected neighbors. **A.** Representative traces of potassium currents in GLAST+ BRAFV600E neuron recorded in the presence of 3 mM 4AP, 1μM TTX, 10μM NBQX, 50μM D-AP5, 10μM SR, 50μM ZD7288, and 1 mM Co^2+^ substitution for Ca^2+^ (5 min in, holding voltage is −80 mV, holding current −32.41 pA) with whole-cell capacitance compensated, grey bar indicate the region where the measurement was made in all conditions (upper panel); middle panel is showing the traces of the same neuron 9 min after application of 25 mM TEA with previous inhibitors cocktail (holding current −40.25 pA); lower panel is showing subtracted traces before and after 25 mM TEA with voltage step protocol. **B.** Representative traces of potassium currents in GLAST+ untransfected neighbor recorded in the presence of the same inhibitors cocktail as for A (6 min in, holding voltage is −80 mV, holding current is −48.04 pA, upper panel) with whole-cell capacitance compensated; middle panel is showing traces from the same neuron 9 min after application of 25 mM TEA with previous inhibitors cocktail (holding current −71.67 pA); lower panel is showing subtracted traces before and after 25 mM TEA. **C.** Average sustained current activation curve showing decreased peaks at all tested voltages in GLAST+ BRAFV600E neurons compared to their untransfected neighbor before and after application of 25 mM TEA. **D.** Normalized to the maximum current and averaged sustained current activation curve have similar kinetics between two conditions before and after application of 25 mM TEA. **E.** Maximal sustained current measured at +20 mV voltage step showing lower values in GLAST+ BRAFV600E neurons (n=18) compared to their untransfected neighbors (n=8, T=2.92, p=0.008), as well as after application of 25 mM TEA (T=2.411, p=0.028). It was also decreased in the same neurons when comparing before and after 25 mM TEA – GLAST+ BRAFV600E (paired sample T=2.474, p=0.035); and their untransfected neighbor (paired sample T=3.827, p=0.006); and comparing untransfected neighbor neurons to GLAST+ BRAFV600E after application of 25 mM TEA (T=6.919, p<0.001). **F.** Current density was not different in untransfected neighbors’ comparison. **G.** Capacitance was measured from −5 mV steps at the beginning of each trace recording using built-in procedure in Axograph acquisition software before application of whole-cell capacitance compensation. There was no statistically significant difference in capacitance measurements compared to untransfected neighbors condition. **J.** Input resistance was significantly increased in GLAST+ BRAFV600E (n=18) compared to their untransfected neighbors (n=8, T=3.293, p=0.003), it was increased in GLAST+ BRAFV600E neurons (n=7) compared to their untransfected neighbors (n=4), but not statistically significant (T=2.223, p=0.053); it was significantly increased in GLAST+ BRAFV600E neurons (n=7) after application of 25 mM TEA compared to their untransfected neighbors before application of TEA (n=8, T=3.664, p=0.009). * - p<0.05, ** - p<0.01, *** - p<0.001. Error bars are ±SEM for C. and D, ±2SEM for E-H.

### 3.6 Elevated I_H_ in BRAFV600E neurons

In response to hyperpolarizing current pulses in whole-cell current clamp mode BRAFV600E neurons in either GLAST+ or NESTIN+ condition displayed an initial deflection, SAG ratio that was absent in untransfected neighbor neurons, control-FP neurons, and in BRAFwt neurons (Figure 5A lower panel, 5E). SAG ratio was calculated as 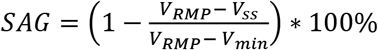, V_RMP_ – Resting Membrane Potential, V_ss_ – stable-state voltage in the last 100 ms of 1 second −40 pA pulse, V_min_ – minimal initial voltage deflection in response to 1 second −40 pA pulse. GLAST+ BRAFV600E neurons (n=57) had average SAG ratio of 23.41 ± 1.33% that was significantly larger than in their untransfected neighbor neurons of 4.46 ± 1.48% (n=20, p=0.001); and in GLAST+ control-FP neurons 6.82 ± 1.86% (n=15, p<0.001); and in GLAST+ BRAFwt neurons 6.59 ± 1.94% (n=17, p<0.001) (Figure 5E). In NESTIN+ BRAFV600E average SAG was 24.48 ± 2.27 % (n=8) and it was significantly increased compared to all non-BRAF V600E conditions (p<0.001, Figure 5E); in NESTIN+ control-FP the average SAG ratio was 10.03 ± 1.77% (n=5); and in NESTIN+ untransfected neighbor the average SAG ratio was 6.59 ± 2.41% (n=4).

Average rebound excitation measured as an overshoot above RMP at the end of 1 second −40 pA current step was also larger in GLAST+ BRAFV600E neurons (n=42) 2.28 ± 0.24 mV compared to their untransfected neighbor neurons 0.69 ± 0.18 mV (n=22, p<0.01); to GLAST+ control-FP neurons 0.37 ± 0.11 mV (n=14, p<0.01); to GLAST+ BRAFwt neurons 0.34 ± 0.06 mV (n=17, p<0.01); to NESTIN+ untransfected neighbor 0.18 ± 0.27 mV (n=4, p<0.05); to NESTIN+ control-FP 0.47 ± 0.13 mV (n=5, p<0.05); in NESTIN+ BRAFV600E rebound excitation was increased (n=7) 1.82 ± 0.17 mV compared to non-BRAFV600E conditions (to GLAST+ BRAFwt - p<0.001; to GLAST+ untransfected neighbor – p=0.02; to GLAST+ control-FP – p<0.001; to NESTIN+ untransfected neighbor – p<0.001; to NESTIN+ control-FP – p<0.001) (Figure 5F). In 20.34% of GLAST+ BRAFV600E neurons (12/59) and in 11.11% of NESTIN+ BRAFV600E (1/9) rebound excitation resulted in AP firing (Figure 5G upper right panel).

Increased SAG ratio and rebound excitation has been previously shown in layer 5 cortical, hippocampal and non-cortical neurons in mice, rats and cats to be associated with hyperpolarization activated conductances ^132–136^. To test whether BRAFV600E expressing neurons have increased hyperpolarization activated conductance we recorded cells in whole-cell voltage clamp configuration and show that BRAFV600E neurons have Ih that is absent in all other conditions and have half activation voltage of V_1/2_ = −82.79 mV and the slope factor k = 11.58^−1^ mV using recording protocol that hold the cell at −50 mV and the first voltage step is at −120 mV with 5 mV increase for 1.5 seconds (Figure 9; Table 3).

**Figure 9.**
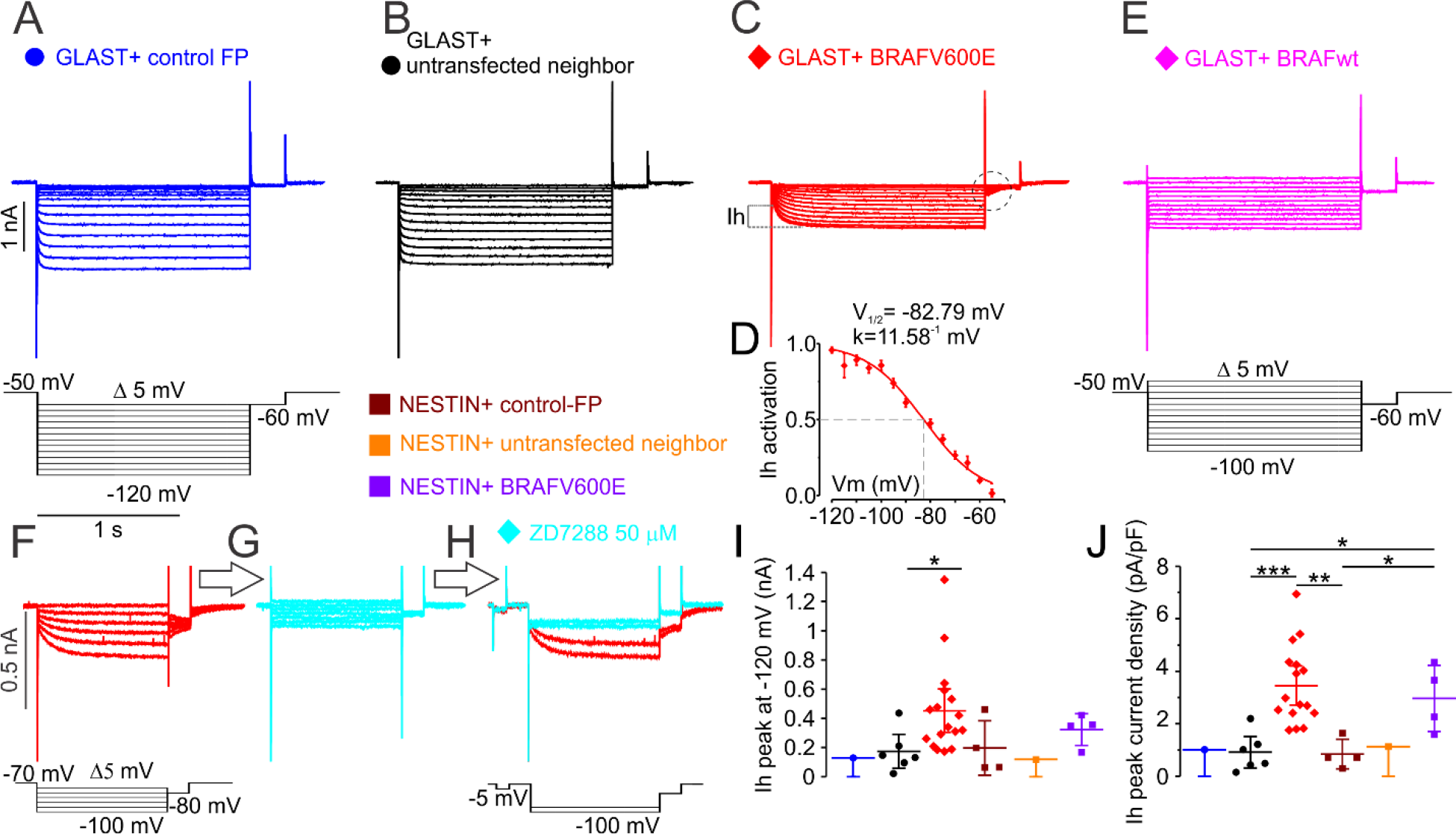
Hyperpolarization activated depolarizing current (Ih) recorded in whole-cell voltage-clamp configuration is increased in BRAFV600E expressing cortical neurons of layers 2/3. **A., B., C., E.** Representative traces of currents in response to hyperpolarizing voltage step protocols shown in the lower panels. Note that GLAST+ BRAFwt was recorded at the same holding potential as all other conditions with the first hyperpolarizing voltage steps been −100 mV and not −120mV. **D.** Ih activation curve from the voltage steps protocol shown in C. (tail currents, dashed circle) with maximal activation around −120 mV half activation −82.79 mV and k slope factor of 11.58^−1^ mV, which are averaged and fit with Boltzmann curve (n=18). **F., G., H.** Application of 50 μM ZD7288, a known Ih inhibitor in perfusion system for 5 minutes blocked Ih**. I.** Ih peak current measured as shown in C., recorded with protocol shown in A. lower panel. The significant increase was only found in GLAST+ BRAFV600E neurons (n=17) compared to their untransfected neighbors (n=6, T=2.117, p=0.046); **J.** right panel – Ih peak density was increased in GLAST+ BRAFV600E neurons (n=16) compared to their untransfected neighbor (n=6, T=3.918, p<0.001), to NESTIN+ control-FP (n=4, T=5.546, p<0.001); it was also significantly increased in NESTIN+ BRAFV600E neurons (n=4) compared to GLAST+ untransfected neighbors (n=6, T=3.275, p=0.011), and to NESTIN+ control-FP (n=4, T=3.066, p=0.022). Error bars are ±SEM and for H and I; ±2SEM for J.

**Table 3.** Sustained K^+^ current average of maximal values and current density.

| Condition | $I_{K-Sustained}$ maximal (pA) last 100 ms of 500 ms +20 mV pulse | $I_{K-Sustained}$ maximal density (pA/pF) | $C_m$ (pF) | n |
| --- | --- | --- | --- | --- |
| **GLAST+ untransfected neighbor** | **4062.66 ± 210.81** | **34.94 ± 11.76** | **178.04 ± 31.74** | **8** |
| **GLAST+ BRAFV600E** | **3034.19 ± 282.49\*\*** | **21.30 ± 2.53** | **157.76 ± 13.29** | **18** |

**\*\* - $t(23.5)=2.92$ , $P=0.008$**

**Table 4.** Sustained K^+^ current average of maximal values and current density – TEA sensitive

| Condition | I<sub>K</sub>-Sustained peak (pA) last 100 ms of 500 ms +20 mV pulse TEA sensitive | n |  | I<sub>K</sub>-Sustained density TEA sensitive (pA/pF) | C<sub>m</sub> (pF) | n |
| --- | --- | --- | --- | --- | --- | --- |
| GLAST+ untransfected neighbor | 2484.56 ± 258.29 | 8 |  | 12.23 ± 3.85 | 239.88 ± 59.23 | 4 |
| GLAST+ BRAFV600E | 1561.06 ± 266.57* | 11 |  | 7.28 ± 1.36 | 187.18 ± 18.45 | 7 |

\* - T(17)=2.41, P=0.028

**Table 5.** Ih peak and current density.

| Condition | I<sub>h</sub> peak -50 -120 mV (pA) | I<sub>h</sub> peak density -50 -120 mV (pA/pF) | C<sub>m</sub> (pF) | n |
| --- | --- | --- | --- | --- |
| GLAST+ control-FP | -129.22 | -1.02 | 126.94 | 1 |
| GLAST+ untransfected neighbor | -174.17 ± 58.24 | -0.92 ± 0.30 | 193.03 ± 24.99 | 6 |
| GLAST+ BRAFwt | -66.22 ± 13.03* | -0.36 ± 0.08 | 253.49 ± 41.27 | 12 |
| GLAST+ BRAFV600E | - 446.58 ± 58.24 | -3.45 ± 0.37 | 135.95 ± 14.18 | 18 |
| NESTIN+ control-FP | -197.17 ± 93.19 | -0.85 ± 0.28 | 214.29 ± 44.06 | 4 |
| NESTIN+ untransfected neighbor | -117.75 | -1.13 | 104.00 | 1 |
| NESTIN+ BRAFV600E | -322.76 ± 54.45 | -2.97 ± 0.63 | 84.34 ± 32.33 | 4 |

This current was blocked with application of 50 μM ZD7288, a known Ih inhibitor in perfusion system and recorded at least 5 minutes later. Ih peak was only significantly increased in GLAST+ BRAFV600E neurons (n=17) compared to their untransfected neighbors (n=6, p<0.05), however when normalized to cell capacitance the Ih peak density was significantly increased in GLAST+ BRAFV600E neurons compared to their untransfected neighbors (p<0.001), and to NESTIN+ control-FP (n=4, p<0.001). In NESTIN+ BRAFV600E neurons (n=4) the Ih peak density was significantly increased compared to GLAST+ untransfected neighbors (p<0.05), and to NESTIN+ control-FP (p<0.05). Consistent with that application of ZD7288 decreased SAG and rebound excitation in BRAFV600E neurons (data not shown). Hyperpolarization activated conductance is generated through ion channels with subunit composition of HCN1-4 ^137, 138^. Koh et al. ^14^ finding that HCN1 is downregulated in GG patients and in mouse model suggest that HCN channels subunit composition may change and, possibly, the expression may be redistributed across different cellular compartments.

## 4. Discussion

Here we showed that introduction of human BRAFV600E, an LNETs associated mutation that constitutively activate BRAF in a RAS-independent manner, into radial glia progenitors using different driver promoters - GLAST and NESTIN, increased astrogenesis in the first case and neurogenesis in the second case. The results from GLAST experiments consistent with previous studies that showed increased astrogenesis when constitutive MEK1 a downstream target of BRAF was expressed in hGFAPCre/CAG-loxpSTOPloxp-Mek1^S218, S222E^ mouse line ^128^ and also in tamoxifen induced knockdown of NF1, a RAS-GTPase activating protein in hGFAPCre driven mouse line ^129^ and in GG patients and BRAFV637E transgenic mouse line that were electroporated with episomal Cre plasmid ^14^. Additionally, Gronych et al. ^139^ showed that introducing truncated BRAFV600E containing kinase domain using retroviral vector into neonatal Ntv mice under promoter derived from the human NESTIN gene was sufficient to model tumor induced astrogenesis observed in pilocytic astrocytoma, another LNET entity. However full length BRAFV600E did not have such an effect suggesting that there is an increased negative regulation of BRAF activity through possible phosphorylation of inhibiting residues on C-terminus domain in later progenitors pool available at birth ^119, 140–143^. It may also reflect possible increased requirement of Hsp90 stabilizing binding in the full length BRAFV600E protein compared to truncated version when introduced in postnatal animals ^144, 145^. Those experimental studies mostly targeted progenitor population that may already have switched to glial fate. Together with our work this suggest that there are, probably, at least two separate populations of progenitors that may overlap at some developmental stage ^116–118, 146^, this notion is also supported by the previous work in which the effects of overactivation of RAS-RAF-ERK pathway was examined in Neurog2 driven and in Ascl1 driven radial glia progenitors, that were proposed as a progenitor molecular fate switch. Neurog2 driving the excitatory neuronal differentiation, and RAS-RAF-ERK pathway activation cause switching off Neurog2 and turning on Ascl1, through direct phosphorylation by ERK, subsequently driving inhibitory interneuronal differentiation at the low levels of RAS-RAF-ERK pathway activation, oligodendrogenesis and astrogenesis at the high levels of RAS-RAF-ERK pathway activation. Which was proposed as a probable explanation for different LNETs histopathology ^127^.

### 4.1 BRAFV600E LNETs, MCD histopathology and inflammation

Increased number of mislocalized neurons in lower cortical layers in both our GLAST+ and NESTIN+ BRAFV600E transgenic slices, together with balloon-like cells and clusters, increased astrogenesis in GLAST+ suggest that we partially recaptured histopathology of LNETs. Further, NESTIN+ BRAFV600E slices also showed increased soma size of the transgenic cells in the subventricular area compared to neurons in the upper cortical layers. Membrane blebbing, and DNA fragmentation, signs of apoptosis and pyroptosis were not observed in those balloon-like cells, additional examination with anti-caspase-3 immunostaining in the selected slices did not show immunoreactivity.

Increased inflammatory immune response and activation of classic complement pathway in current work is consistent with microarray study in GG resected tissue ^69^ and recent publication by Koh and colleagues ^14^, it was also reported in cortical tubers resected tissue ^63, 68^. However inflammatory response in those studies may result from seizure activity. In current work video-ECoG recording did not show any behavioral manifestations of seizures in the seven recorded animals with BRAFV600E under GLAST promoter, also presence of electrocorticographic seizures was rarely observed suggesting that seizures cannot account for activation of inflammatory pathways using current experimental design with GLAST+ driving promoter. This is supported by RNA sequencing results that did not show increase of IL-1R1, which mediates biological response to IL-1β and is increased in neurons and subsequently in astrocytes after seizures ^147–149^. However, this possibility cannot be completely excluded. Activation of inflammatory innate immunity has been shown to precipitate seizures in mouse model of kainate-induced seizures ^150, 151^ through phosphorylation of NR2B subunits of NMDA channels by Src serine/threonine kinase. Sequential injection of complement complex components was sufficient to induce seizures in rats’ hippocampus ^152^. In case our manipulation have similar results, but requires a longer time to induce seizures or dependent on the presence of “threshold” number of affected neuronal component (GLAST+ vs. NESTIN+), inhibition of inflammatory pathways may be used to reduce seizures ^150, 153^.

### 4.2 BRAFV600E and neuronal hyperexcitability

Increased excitability properties in BRAFV600E neurons observed in current work are described for the first time. Previous studies that concentrated on cortical tissue resected from FCD patients and TSC patients ^51, 54, 74–76^ did not observe significant increase in action potential firing properties in examined malformed components and in mouse model of synapsin-driven TSC1 KO ^77^. Those studies suggested that the difference in synaptic circuit excitability account for seizures observed in FCD and TSC patients. In current work BRAFV600E neurons with depolarized resting membrane potential, increased input resistance, low capacitance fired three times more action potentials then untransfected neighbor neurons, or control neurons transfected with fluorophore only. In addition, neurons transgenic for wild type BRAF displayed similar properties to control conditions. Significant increase in hyperpolarization activated depolarizing conductance (I_H_) observed in BRAFV600E neurons may explain more depolarized resting membrane potential. Indeed, I_H_ inhibition with ZD7288 hyperpolarized membrane potential in neurons, but it did not alter significantly action potential firing frequency. The reducing effects of IH on neuronal excitability suggest that increased IH conductance may be compensatory and counteract the hyperexcitability changes observed in BRAFV600E neurons, thus working to reduce input resistance ^138, 154^. I_H_ in dendrites has been previously shown to decrease amplitudes of propagating EPSPs and to dampen temporal dendritic summation ^155–157^. In addition to increased I_H_ conductance BRAFV600E neurons had decreased sustained potassium currents, which may contribute to action potential adaptation ^158–160^. Indeed, when retigabine ^161^ a Kv7.2, Kv7.3 and Kv7.4 activator was acutely applied to BRAFV600E neurons it decreased action potential firing frequency by 30%. This was consistent with previous studies in hippocampal pyramidal neurons ^162, 163^, that also showed opposite effect of blocking Kv7 channels with XE-991, which increased action potential firing frequency. Opening of multiple different voltage sensitive potassium channels may contribute to sustained, non-inactivating potassium currents, one of this channels is Kv1, global inhibition of which with α-dendrotoxin in avian nucleus magnocellularis neurons has been shown to depolarize membrane potential by about 5 mV, increase input resistance two-fold, hyperpolarized action potential voltage threshold by 8-10 mV and decreased rheobase ^164^. This suggest that retigabine in case of BRAFV600E associated epilepsy may be used to reduce seizures.

In a small number of BRAFV600E neurons we have also observed post-action potential depolarization waves, that were increased in potassium currents recording, when we substituted Ca^2+^ ions with Co^2+^ ions in the extracellular solution. One of the possible explanations to post-action potential depolarizing waves may be presence of gap junctions ^165^, or pannexin hemichannels as has been shown in severe inflammation with epileptic seizures Rasmussen encephalitis ^166^.

Reflecting on previous studies in FCD and TSC resected cortical tissue ^51, 54, 74–77^ that showed increased action potential dependent glutamatergic synaptic events frequencies in TSC compared to FCD tissue and the opposite effect on GABAergic events frequencies. In addition, they showed large amplitude GABAergic pacemaker rhythmic events in immature looking pyramidal neurons and decreased frequencies of action potential dependent mixed glutamatergic and GABAergic events in normal looking neurons from severe FCD cortical areas. This led us to examine the action potential dependent mixed glutamatergic and GABAergic events in our preparation as an initial step. Interestingly, in current work action potential dependent mixed synaptic events frequencies were increased in untransfected neighbor neurons compared to BRAFV600E neurons, and was comparable to PIK3CAE545K neurons, TSC1 KD neurons, their untransfected neighbors, and TSC2 untransfected neighbors. sPSCs frequencies in BRAFV600E neurons were higher compared to control-FP and to BRAFwt neurons, additionally sPSCs amplitudes, although significantly different did look similar in all conditions, which shows that increased I_H_ is not sufficient to counteract hyperexcitability changes in synaptic activity.

Depolarized membrane potential due to I_H_, lower rheobase, increased input resistance, hyperpolarized action potential voltage threshold, probably due to decreased Kv1 mediated I_K_^+^ currents allows BRAFV600E to fire action potentials in response to smaller depolarizing current input. Increased action potential dependent synaptic activity, which is largely mediated by miniature Post-Synaptic Currents (mPSCs) suggest that the neuronal network is more excitable than in control-FP and in BRAFwt conditions and that increased frequencies may increase probability of action potential firing. Since those neurons have a low action potential voltage threshold the may fire more action potentials in response to similar synaptic inputs as in control-FP. This was not tested yet, and further experiments to elucidate it may need to be performed.

### 4.3 Caveats in current work

In current work we have tested the effect of acute BRAFV600E inhibition with specific blocker Vemurafenib (PLX4032, PLX4720) on excitability in BRAFV600E neurons. This FDA approved cancer medication was developed for unresectable or metastatic melanoma treatment ^167, 168^. Preincubation of BRAFV600E transfected cortical slices in 10-50 μM of Vemurafenib for 1-5h did decrease action potential firing frequency, but this effect was indistinguishable in slices preincubated in comparable amount of solvent (DMSO). This was also consistent with previous work examining the effect of DMSO on neuronal excitability in layer 2 of perirhinal cortex ^169^. Similar results were obtained for Rapamycin experiments, which was also dissolved in DMSO.

Histopathological examination of immunopositivity to CD34, a hematopoietic stem cell marker previously shown to label extensively LNETs ^35^, showed only few immunopositive neuronal cells.

Electrocorticographic recordings did not show any behavioral manifestation of seizures in GLAST+ BRAFV600E transgenic mice. The increased astrogenesis and decrease neuronal content suggest that this manipulation may produce a different malformation which is not Ictogenic in mice. However, electrocorticographic recordings were not performed in NESTIN+ BRAFV600E mice, which has more neurons compared to GLAST+ BRAFV600E and may have sufficient number of BRAFV600E transgenic neurons that are as hyperexcitable as GLAST+ BRAFV600E neurons to initiate seizures. This would be addressed in the next set of experiments in ex-vivo cortical slices with ChR2 added to the plasmid mix. Stimulation of BRAFV600E transgenic neurons with ChR2 in NESTIN+ condition with high power blue laser under high extracellular potassium concentration may differentiate BRAFV600E condition from control-FP based on the amount of stimulation required to initiate ictal activity (threshold)^170^. Also electrocorticographic recording of NESTIN+ BRAFV600E freely moving mice to examine whether seizures are manifested in those animals.

### 4.4 Short summary

In summary, our study proposes that constitutively active RAS-RAF-ERK pathway resulting from BRAFV600E mutated protein switch cell fate dependent on the driver promoter, have histopathological features akin to LNETs, alters gene signature in the affected cortical tissue, increasing inflammation, and, increase excitability in neurons with BRAFV600E mutation that may contribute to seizure generation.

**Figure 10.**
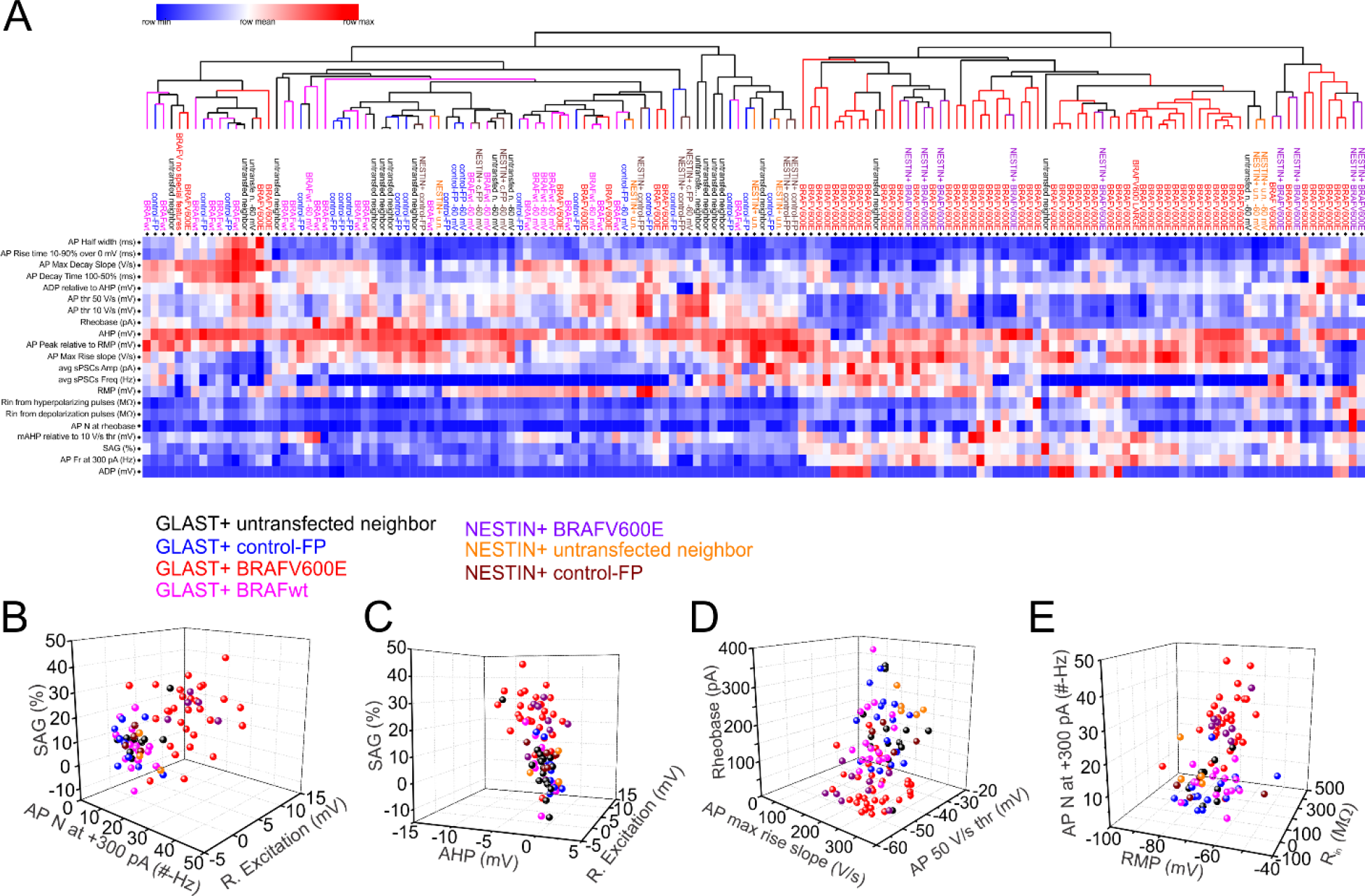
GLAST+ and NESTIN+ BRAF V600E expressing neurons segregate to separate clusters in HCA analysis of electrophysiological properties. **A.** Unsupervised Hierarchical Cluster Analysis was performed on 20 recorded electrophysiological parameters and showing that most of the BRAFV600E neurons segregate together by electrophysiological parameters recorded. The parameters are AP width at 50% height from RMP in ms, AP maximal decay slope in V/s, AP decay time from 100% to 50% height in ms, AP rise time from 10% to 90% height in ms, AP 50 V/s voltage threshold in mV, AP 10 V/s voltage threshold in mV, rheobase in pA, AHP measured at the end of +300 pA 1 second current step in mV, AP peak relative to RMP in mV, AP maximal rise slope in V/s, RMP in mV, Rin from hyperpolarizing pulses in MΩ, Rin from depolarizing pulses in MΩ, average sPSCs amplitude in pA, average sPSCs instantaneous frequency in Hz, mAHP measured relative to 10 V/s AP voltage threshold for rheobase APs in mV, SAG ratio in %, AP frequency at +300 pA 1 second current step in Hz, number of APs at rheobase, rebound excitation measured as an overshoot above RMP (mV). **B.** Most contributing electrophysiological parameters to the variability in PCA shown in 3D plots – upper left panel SAG ratio on the Z axis, AP number at +300 pA 1 second pulse is on the X axis and rebound excitation is on the Y axis. **C.** SAG ratio on the Z axis, AHP at the end of +300 pA 1 second pulse on the X axis and rebound excitation on the Y axis. **D.** Rheobase on the Z axis, AP maximal rise slope is on the X axis, and AP 50V/s voltage threshold on the Y axis. **E.** AP number at +300 pA 1 second pulse on the Z axis, resting membrane potential (RMP) is on the X axis, Input resistance from depolarizing current pulses (Rin) is on the Y axis.

## Supporting information

Sup. Table 1. DE genes in BRAF V600E to control-FP

Sup. Table 2. DE genes in BRAF V600E to BRAFwt

Sup. Table 3. DE genes in BRAFwt to control-FP

Sup. Table 6. Venn diagram genes

Gene Analytics 262 downregulated genes

Gene Analytics 402 upregulated genes

Gene Analytics 500 random mouse genes

Gene Analytics 825 upregulated genes in BV6 to Control

Gene Analytics 1000 random mouse genes

Gene Analytics for Venn diagram genes

Gene Analytics 1800 genes

Gene Analytics 825 upregulated genes in BV6 to Bw and Contorl

## Acknowledgements

We want to thank Dr. Bo Reese from the Center for Genome Innovation, Institute for Systems Genomics, University of Connecticut, Storrs, CT for helping with RNA-sequencing.

Current work was supported by NIH/NICHD grant #

And, NIH Akiko Nishiyama grant #S10OD016435 for acquisition of Leica SP8 microscope.

Author contribution
R.U.G. performed IUE, electrophysiological *ex-vivo* patch-clamp recordings, analysis, IHC, image acquisition, and final figures. R.U.G., A.S., S.B. counted the cells with ImageJ-Fiji and analyzed it. R.U.G. and J.J.L. performed RNA extraction, alignment and DE gene quantification and functional enrichment analysis. R.U.G. and J.J.L. wrote the paper. J.J.L. wrote the grant, acquired funding and provided the equipment and biomolecular research tools.

