## Supplementary material for "BRAFV600E Expression in Mouse Neuroglial Progenitors Increase Neuronal Excitability, Cause Appearance of Balloon-like cells, Neuronal Mislocalization, and Inflammatory Immune response": Sup. Table 1. DE genes in BRAF V600E to control-FP

| UP p<0.05 | Gene ID | Fold change (BRAF V600E vs. control-FP) | log-10 P-values | P-value | FDR step up |
|---|---|---|---|---|---|
| | C3 | 681.23 | 3.731428262 | 0.00018559734 | 0.014121412 |
| | H2-Aa | 258.75 | 2.595908848 | 0.00253566077 | 0.027051642 |
| | Cd74 | 231.76 | 2.458336483 | 0.00348067535 | 0.031057117 |
| | Serpinf2 | 147.73 | 2.841645191 | 0.00143997453 | 0.021719747 |
| | H2-Ab1 | 134.29 | 2.513260092 | 0.00306718455 | 0.029428615 |
| | Tgtp2 | 91.56 | 1.674117163 | 0.02117789726 | 0.082135518 |
| | Iigp1 | 63.64 | 1.763845253 | 0.01722482218 | 0.073534751 |
| | Ifi47 | 47.38 | 1.652430124 | 0.02226229204 | 0.084644758 |
| | Igtp | 43.56 | 1.993677334 | 0.01014664967 | 0.054019804 |
| | C4b | 37.75 | 3.240955692 | 0.00057417504 | 0.016426119 |
| | C1s1 | 35.31 | 2.535149615 | 0.00291642213 | 0.028906123 |
| | A2m | 35.22 | 4.462844067 | 0.00003444736 | 0.014121412 |
| | Psmb9 | 32.73 | 2.427532753 | 0.00373651945 | 0.032301194 |
| | Serpina3i | 26.34 | 3.158271888 | 0.00069458934 | 0.016949337 |
| | Slc43a3 | 26.15 | 3.187761376 | 0.00064899093 | 0.016529429 |
| | H2-Q7 | 24.83 | 2.385956172 | 0.00411191216 | 0.033646417 |
| | Gbp6 | 23.95 | 2.580855164 | 0.00262509386 | 0.027662128 |
| | Cybb | 21.82 | 2.478019788 | 0.00332644397 | 0.030310787 |
| | Psmb8 | 21.02 | 2.016139153 | 0.00963520252 | 0.052705939 |
| | H2-DMb1 | 20.92 | 1.847541134 | 0.01420557659 | 0.065568971 |
| | Gfap | 20.78 | 3.704824579 | 0.00019732196 | 0.014121412 |
| | Gbp2 | 19.87 | 3.033430809 | 0.00092591089 | 0.018883167 |

| | | | | | |
|---|---|---|---|---|---|
| | Serpina3h | 18.63 | 3.323259827 | 0.00047505093 | 0.015871978 |
| | Tap1 | 18.51 | 2.117532813 | 0.00762899248 | 0.047051161 |
| | Irgm2 | 18.11 | 1.416965245 | 0.03828553804 | 0.118593683 |
| | H2-Q4 | 16.04 | 1.771538941 | 0.01692236505 | 0.072949681 |
| | S100a4 | 15.85 | 4.818196229 | 0.00001519861 | 0.014121412 |
| | Fgl2 | 15.72 | 3.156356994 | 0.00069765869 | 0.016949337 |
| | Oasl2 | 15.54 | 1.867518471 | 0.01356692828 | 0.063647375 |
| | Cxcl16 | 15.43 | 2.574907014 | 0.00266129480 | 0.027727676 |
| | Cd109 | 15.20 | 3.553161661 | 0.00027979396 | 0.014292375 |
| | Lyz2 | 14.81 | 2.091855323 | 0.00809365480 | 0.048396393 |
| | Cfb | 14.52 | 1.308359241 | 0.04916326969 | 0.139519308 |
| | H2-K1 | 13.78 | 2.105436906 | 0.00784446074 | 0.047709242 |
| | Cd52 | 13.51 | 1.875985783 | 0.01330497971 | 0.062912500 |
| | H2-T-ps | 12.45 | 1.308850372 | 0.04910770384 | 0.139463975 |
| | Serpina3n | 12.25 | 3.204361752 | 0.00062465216 | 0.016529429 |
| | Serping1 | 12.12 | 3.193664709 | 0.00064022892 | 0.016529429 |
| | Gbp5 | 12.09 | 2.167075095 | 0.00680651655 | 0.043839765 |
| | Slfn8 | 11.46 | 1.992547905 | 0.01017307146 | 0.054076663 |
| | Bst2 | 11.38 | 1.455348586 | 0.03504704566 | 0.112477329 |
| | Gbp4 | 11.26 | 1.503425055 | 0.03137436504 | 0.105227032 |
| | Gbp3 | 11.23 | 2.174372857 | 0.00669309736 | 0.043422970 |
| | Vim | 11.07 | 4.094881289 | 0.00008037458 | 0.014121412 |
| | Irf4 | 10.48 | 1.49164982 | 0.03223667046 | 0.106815551 |

| | | | | | |
|---|---|---|---|---|---|
| | Irgm1 | 10.36 | 1.90517737 | 0.01244006443 | 0.060483225 |
| | Cybrd1 | 10.34 | 3.16281069 | 0.00068736800 | 0.016866292 |
| | Mmp19 | 10.16 | 2.870342996 | 0.00134789792 | 0.021189830 |
| | Parp14 | 10.04 | 1.981750281 | 0.01042916933 | 0.054655689 |
| | Casp12 | 9.80 | 2.225545493 | 0.00594914435 | 0.040833454 |
| | Uba7 | 9.71 | 2.616357086 | 0.00241903924 | 0.026631344 |
| | Il18bp | 9.42 | 1.548404803 | 0.02828754105 | 0.098706250 |
| | Csf2rb | 9.40 | 1.983594802 | 0.01038496884 | 0.054655689 |
| | Cnmd | 9.39 | 2.56153842 | 0.00274448954 | 0.028260938 |
| | Ggta1 | 9.36 | 3.846628748 | 0.00014235452 | 0.014121412 |
| | Gpnmb | 9.24 | 2.290116445 | 0.00512723891 | 0.037595192 |
| | H2-D1 | 9.23 | 2.104446321 | 0.00786237365 | 0.047709242 |
| | Rtp4 | 9.22 | 1.81013016 | 0.01548352499 | 0.069438354 |
| | Fam167a | 9.16 | 3.577734267 | 0.00026440261 | 0.014200135 |
| | Gm8909 | 9.11 | 1.820045433 | 0.01513402917 | 0.068470085 |
| | Ifitm3 | 8.93 | 3.087642926 | 0.00081725404 | 0.017954237 |
| | Ptprc | 8.72 | 2.781933236 | 0.00165221578 | 0.022978843 |
| | Capg | 8.54 | 3.457175839 | 0.00034899898 | 0.014883417 |
| | B2m | 8.51 | 2.031675874 | 0.00929659959 | 0.051957065 |
| | Cd274 | 8.24 | 1.591468026 | 0.02561721858 | 0.092926663 |
| | Tnfaip2 | 8.24 | 2.014790305 | 0.00966517440 | 0.052785835 |
| | Cdkn2b | 8.12 | 1.854916584 | 0.01396636591 | 0.064835079 |
| | Ly6a | 8.00 | 1.703344689 | 0.01979954961 | 0.078980679 |

| | | | | | |
|---|---|---|---|---|---|
| | Mamdc2 | 7.96 | 2.607668992 | 0.00246791961 | 0.026852745 |
| | Mgst1 | 7.94 | 4.030278898 | 0.00009326552 | 0.014121412 |
| | Csf2rb2 | 7.93 | 1.713824049 | 0.01932751200 | 0.077928211 |
| | Flnc | 7.93 | 3.641673186 | 0.00022820587 | 0.014200135 |
| | Lbp | 7.93 | 2.927588941 | 0.00118143834 | 0.020343539 |
| | Angpt1 | 7.87 | 3.520169617 | 0.00030187725 | 0.014504487 |
| | 1500015O10Rik | 7.84 | 1.925238053 | 0.01187850943 | 0.058967702 |
| | 2410004P03Rik | 7.84 | 2.210236709 | 0.00616259023 | 0.041540673 |
| | Tmem140 | 7.80 | 1.9589124 | 0.01099227540 | 0.056423181 |
| | Osmr | 7.71 | 3.48235624 | 0.00032933945 | 0.014875020 |
| | Prrx2 | 7.68 | 3.693179232 | 0.00020268461 | 0.014121412 |
| | Irf7 | 7.65 | 1.76029744 | 0.01736611049 | 0.073718790 |
| | Stat1 | 7.48 | 1.382738948 | 0.04142486026 | 0.124856110 |
| | Tlr2 | 7.46 | 2.301887902 | 0.00499013274 | 0.037064548 |
| | Gbp7 | 7.38 | 2.035539542 | 0.00921425991 | 0.051721056 |
| | Ms4a6b | 7.31 | 2.597141738 | 0.00252847266 | 0.027050644 |
| | Cd44 | 7.31 | 3.346211498 | 0.00045059721 | 0.015871978 |
| | Lgals3bp | 7.31 | 2.487088427 | 0.00325770364 | 0.030074830 |
| | Ifit3b | 7.31 | 2.30156521 | 0.00499384191 | 0.037072059 |
| | Ccr1 | 7.28 | 3.220253389 | 0.00060220813 | 0.016529429 |
| | Ms4a6c | 7.19 | 2.590123716 | 0.00256966367 | 0.027215913 |
| | Cd48 | 7.18 | 2.885522169 | 0.00130160087 | 0.020991075 |
| | Pycard | 7.14 | 2.999623854 | 0.00100086648 | 0.019234834 |

| | | | | | |
|---|---|---|---|---|---|
| | Lpo | 7.04 | 2.575952105 | 0.00265489833 | 0.027727676 |
| | Bcl2a1d | 7.02 | 2.312801733 | 0.00486629314 | 0.036440182 |
| | Gimap4 | 6.81 | 2.065039767 | 0.00860914917 | 0.049726069 |
| | C1ra | 6.80 | 2.12366584 | 0.00752201439 | 0.046516652 |
| | Plscr4 | 6.79 | 3.450583406 | 0.00035433707 | 0.014935416 |
| | Tekt1 | 6.78 | 3.130850861 | 0.00073985930 | 0.017360193 |
| | Ifit1 | 6.76 | 2.625188884 | 0.00237034257 | 0.026509970 |
| | Trim12c | 6.75 | 3.23040251 | 0.00058829816 | 0.016487783 |
| | Rsph4a | 6.70 | 2.373846023 | 0.00422818496 | 0.034075549 |
| | Irf1 | 6.61 | 1.995637435 | 0.01010095798 | 0.053901850 |
| | Cntf | 6.59 | 3.52014992 | 0.00030189094 | 0.014504487 |
| | Gm11992 | 6.51 | 3.711497756 | 0.00019431317 | 0.014121412 |
| | Icam1 | 6.41 | 2.413621788 | 0.00385814204 | 0.032725150 |
| | Dlk1 | 6.39 | 3.061772755 | 0.00086741563 | 0.018396503 |
| | Trim30a | 6.32 | 1.82424483 | 0.01498839640 | 0.068079032 |
| | Ctsc | 6.31 | 2.686947101 | 0.00205614103 | 0.025047370 |
| | Rac2 | 6.26 | 1.913283193 | 0.01221003213 | 0.059813922 |
| | Tap2 | 6.22 | 2.433080902 | 0.00368908871 | 0.032144431 |
| | Sp110 | 6.12 | 1.814912275 | 0.01531396763 | 0.068880271 |
| | Cd24a | 6.11 | 2.829113355 | 0.00148213118 | 0.021922459 |
| | Slc14a1 | 6.06 | 3.983646055 | 0.00010383743 | 0.014121412 |
| | Foxj1 | 6.06 | 2.735506712 | 0.00183862554 | 0.023856991 |
| | Chil1 | 6.01 | 3.248833886 | 0.00056385328 | 0.016356417 |

| | | | | | |
|---|---|---|---|---|---|
| | Lag3 | 5.95 | 1.706688245 | 0.01964770166 | 0.078665230 |
| | Sfrp5 | 5.94 | 2.603925567 | 0.00248928392 | 0.026933268 |
| | Aspg | 5.90 | 3.097846791 | 0.00079827625 | 0.017835958 |
| | Apobec1 | 5.90 | 3.312096635 | 0.00048742002 | 0.015871978 |
| | Fxyd1 | 5.88 | 3.581303256 | 0.00026223868 | 0.014200135 |
| | Isg15 | 5.84 | 1.517902339 | 0.03034573500 | 0.103034530 |
| | Parp9 | 5.83 | 2.303591316 | 0.00497059850 | 0.036999455 |
| | Plce1 | 5.70 | 3.443972354 | 0.00035977224 | 0.015028221 |
| | Igsf1 | 5.67 | 4.39228504 | 0.00004052425 | 0.014121412 |
| | Apobec3 | 5.64 | 3.667628066 | 0.00021496707 | 0.014121412 |
| | Ncf4 | 5.50 | 2.051045128 | 0.00889108726 | 0.050693954 |
| | C3ar1 | 5.45 | 2.59343027 | 0.00255017351 | 0.027143249 |
| | Glipr2 | 5.42 | 3.130554178 | 0.00074036490 | 0.017360193 |
| | Togaram2 | 5.38 | 2.296999287 | 0.00504662126 | 0.037264945 |
| | Hck | 5.37 | 2.693264699 | 0.00202644724 | 0.024906282 |
| | Ccl6 | 5.35 | 2.08979619 | 0.00813212059 | 0.048541907 |
| | Iqub | 5.32 | 2.029541232 | 0.00934240667 | 0.052099842 |
| | Tspo | 5.31 | 2.689147597 | 0.00204574926 | 0.024975124 |
| | Pik3ap1 | 5.25 | 1.751493397 | 0.01772174989 | 0.074537669 |
| | C2 | 5.25 | 2.277876526 | 0.00527379779 | 0.038362761 |
| | Plscr2 | 5.24 | 2.292869703 | 0.00509483703 | 0.037457547 |
| | Irf8 | 5.21 | 2.311486553 | 0.00488105214 | 0.036510908 |
| | Fyb | 5.19 | 2.488831829 | 0.00324465235 | 0.030008970 |

| | | | | | |
|---|---|---|---|---|---|
| | Parp10 | 5.12 | 1.4467245 41 | 0.03574995 164 | 0.1137595 56 |
| | Fas | 5.12 | 3.2717856 12 | 0.00053482 831 | 0.0160558 68 |
| | Mpeg1 | 5.05 | 1.4725007 42 | 0.03368986 403 | 0.1096996 26 |
| | Sdc1 | 5.01 | 2.3025365 32 | 0.00498268 541 | 0.0370292 48 |
| | Gm4070 | 5.01 | 1.4016585 71 | 0.03965896 988 | 0.1211639 44 |
| | Gvin1 | 5.01 | 1.4010559 1 | 0.03971404 195 | 0.1212420 91 |
| | Tmem106a | 4.97 | 1.6671811 1 | 0.02151884 165 | 0.0828264 43 |
| | Ifi30 | 4.96 | 1.7226535 62 | 0.01893853 748 | 0.0771065 02 |
| | Ctss | 4.95 | 2.7689163 84 | 0.00170248 626 | 0.0232543 38 |
| | Itgb2 | 4.94 | 2.2842373 63 | 0.00519711 871 | 0.0379657 67 |
| | Gadl1 | 4.92 | 2.1956675 51 | 0.00637283 170 | 0.0422776 38 |
| | Cd180 | 4.90 | 2.6370819 97 | 0.00230631 171 | 0.0261909 33 |
| | S1pr3 | 4.88 | 3.0890600 37 | 0.00081459 167 | 0.0179456 97 |
| | Ms4a6d | 4.88 | 2.1411383 97 | 0.00722539 514 | 0.0454284 20 |
| | Ubxn10 | 4.85 | 3.29346 | 0.00050879 168 | 0.0158719 78 |
| | Elf4 | 4.85 | 1.6174327 | 0.02413055 440 | 0.0891338 57 |
| | Ppp1r36 | 4.82 | 2.2626677 85 | 0.00546175 500 | 0.0390275 48 |
| | Ly86 | 4.81 | 2.5311687 66 | 0.00294327 766 | 0.0290542 95 |
| | Fcgr1 | 4.79 | 2.3842219 41 | 0.00412836 473 | 0.0336464 17 |
| | Naip2 | 4.78 | 1.6810973 32 | 0.02084023 769 | 0.0814853 26 |
| | Gbp9 | 4.75 | 1.3744551 77 | 0.04222258 538 | 0.1262359 76 |
| | Fcgr2b | 4.75 | 2.5427078 24 | 0.00286610 552 | 0.0287258 61 |
| | Adgrg6 | 4.73 | 3.7620186 7 | 0.00017297 420 | 0.0141214 12 |

| | | | | | |
|---|---|---|---|---|---|
| | B3gnt5 | 4.72 | 2.647734045 | 0.00225043231 | 0.025964275 |
| | Hfe | 4.71 | 2.696492477 | 0.00201144204 | 0.024775488 |
| | S100a6 | 4.71 | 3.800397592 | 0.00015834429 | 0.014121412 |
| | Slc15a3 | 4.71 | 2.326838165 | 0.00471152864 | 0.035960485 |
| | Cd14 | 4.71 | 3.720220171 | 0.00019044950 | 0.014121412 |
| | Hk3 | 4.69 | 1.965912528 | 0.01081651788 | 0.055960005 |
| | Top2a | 4.64 | 2.064599211 | 0.00861788687 | 0.049734726 |
| | Rcn3 | 4.63 | 4.206767318 | 0.00006212018 | 0.014121412 |
| | Cd84 | 4.62 | 2.04731494 | 0.00896778236 | 0.050973497 |
| | Hvcn1 | 4.61 | 1.819643886 | 0.01514802850 | 0.068470085 |
| | Cyba | 4.59 | 2.51688403 | 0.00304169715 | 0.029392377 |
| | Zc3hav1 | 4.56 | 2.66278031 | 0.00217380053 | 0.025519026 |
| | Slc11a1 | 4.56 | 1.80427247 | 0.01569377892 | 0.070083918 |
| | Hs3st3b1 | 4.55 | 1.794934822 | 0.01603486022 | 0.070884150 |
| | Cyp1b1 | 4.54 | 2.490062213 | 0.00323547305 | 0.029999079 |
| | Cd38 | 4.54 | 3.401602222 | 0.00039664116 | 0.015470257 |
| | Pipox | 4.52 | 2.396142624 | 0.00401658883 | 0.033368771 |
| | Ifi35 | 4.51 | 1.982065388 | 0.01042160508 | 0.054655689 |
| | Emp3 | 4.49 | 2.168629531 | 0.00678219808 | 0.043793278 |
| | Rbp1 | 4.48 | 3.724875253 | 0.00018841902 | 0.014121412 |
| | Cd86 | 4.47 | 2.080520053 | 0.00830768359 | 0.049062156 |
| | Ptpn6 | 4.46 | 1.677331477 | 0.02102173336 | 0.081806751 |
| | Col5a2 | 4.46 | 2.885034629 | 0.00130306287 | 0.020991075 |

| | | | | | |
|---|---|---|---|---|---|
| | Slc2a10 | 4.46 | 3.198685162 | 0.00063287048 | 0.016529429 |
| | Il7r | 4.44 | 1.875597428 | 0.01331688263 | 0.062925476 |
| | Dok1 | 4.43 | 3.427148986 | 0.00037398227 | 0.015193137 |
| | Tmem252 | 4.41 | 2.380129327 | 0.00416745264 | 0.033824552 |
| | Slc22a4 | 4.40 | 3.886735424 | 0.00012979698 | 0.014121412 |
| | Ccl9 | 4.38 | 2.774765553 | 0.00167971054 | 0.023195029 |
| | Serpinf1 | 4.38 | 3.129456035 | 0.00074223933 | 0.017360193 |
| | Mki67 | 4.37 | 1.501111819 | 0.03154192401 | 0.105620703 |
| | Srgn | 4.36 | 2.526802216 | 0.00297301968 | 0.029162245 |
| | Igfbp3 | 4.35 | 2.676246551 | 0.00210743141 | 0.025224926 |
| | C1qa | 4.34 | 2.196718382 | 0.00635743045 | 0.042262918 |
| | Kcne1l | 4.32 | 2.500463228 | 0.00315890650 | 0.029738034 |
| | BC026585 | 4.30 | 3.731336252 | 0.00018563666 | 0.014121412 |
| | Rhoh | 4.26 | 1.858157866 | 0.01386251835 | 0.064492408 |
| | C1qb | 4.25 | 2.260816918 | 0.00548508147 | 0.039133180 |
| | Cebpd | 4.25 | 2.425322514 | 0.00375558405 | 0.032339304 |
| | Btk | 4.24 | 2.356996535 | 0.00439545122 | 0.034711434 |
| | Sash3 | 4.24 | 1.978323677 | 0.01051178144 | 0.054875911 |
| | Ppm1j | 4.22 | 1.334205811 | 0.04632273458 | 0.134477885 |
| | Serpinb6b | 4.20 | 1.400736098 | 0.03974329787 | 0.121304455 |
| | Il13ra1 | 4.20 | 3.999307682 | 0.00010015954 | 0.014121412 |
| | Slc1a5 | 4.20 | 3.939420079 | 0.00011496878 | 0.014121412 |
| | Cpxm2 | 4.20 | 3.107116731 | 0.00078141774 | 0.017776366 |

| | | | | | |
|---|---|---|---|---|---|
| | Dnase1l1 | 4.19 | 3.063643562 | 0.00086368711 | 0.018396503 |
| | Pdlim4 | 4.19 | 3.580792244 | 0.00026254742 | 0.014200135 |
| | Hcls1 | 4.18 | 2.512717024 | 0.00307102234 | 0.029428615 |
| | Crybg1 | 4.18 | 1.983201096 | 0.01039438753 | 0.054655689 |
| | Trim21 | 4.17 | 1.666418413 | 0.02155666572 | 0.082925572 |
| | Myo1f | 4.17 | 1.945282211 | 0.01134273509 | 0.057386054 |
| | Fbln1 | 4.15 | 1.803425905 | 0.01572440043 | 0.070084005 |
| | Dapp1 | 4.13 | 4.206815825 | 0.00006211324 | 0.014121412 |
| | Meox1 | 4.12 | 2.472754433 | 0.00336701900 | 0.030579120 |
| | Prokr2 | 4.12 | 1.6368012 | 0.02307803355 | 0.086619847 |
| | Txlnb | 4.11 | 2.820991446 | 0.00151010990 | 0.022098424 |
| | Ccdc114 | 4.10 | 2.019696629 | 0.00955659918 | 0.052550236 |
| | Rsad2 | 4.09 | 1.542822008 | 0.02865352068 | 0.099439356 |
| | Gpsm3 | 4.08 | 2.198513854 | 0.00633120165 | 0.042129318 |
| | Lcp1 | 4.08 | 2.010624875 | 0.00975832156 | 0.053138496 |
| | Esyt3 | 4.07 | 3.968076594 | 0.00010762754 | 0.014121412 |
| | Clic1 | 4.06 | 3.167096173 | 0.00068061862 | 0.016820828 |
| | Col16a1 | 4.06 | 3.298973437 | 0.00050237332 | 0.015871978 |
| | Tgif1 | 4.05 | 2.386430775 | 0.00410742105 | 0.033646417 |
| | Cp | 4.05 | 2.878776349 | 0.00132197625 | 0.021138199 |
| | Gsdmd | 4.03 | 1.614308203 | 0.02430478573 | 0.089536745 |
| | Dock2 | 4.01 | 2.427001583 | 0.00374109224 | 0.032301194 |
| | Birc3 | 3.98 | 2.194941499 | 0.00638349469 | 0.042277638 |

| | | | | | |
|---|---|---|---|---|---|
| | C1qc | 3.97 | 1.901334421 | 0.01255063152 | 0.060788942 |
| | Tnfrsf1a | 3.96 | 3.578376416 | 0.00026401195 | 0.014200135 |
| | Ccna2 | 3.96 | 3.466435304 | 0.00034163684 | 0.014875020 |
| | Nfkb2 | 3.95 | 1.70737421 | 0.01961669274 | 0.078609791 |
| | Zc3h12a | 3.95 | 3.7207135 | 0.00019023328 | 0.014121412 |
| | Lat2 | 3.94 | 2.592840735 | 0.00255363761 | 0.027151188 |
| | Baiap3 | 3.93 | 1.575881929 | 0.02655327361 | 0.094965121 |
| | Clec5a | 3.90 | 2.24144365 | 0.00573530277 | 0.040147326 |
| | Xlr | 3.89 | 2.972256167 | 0.00106596718 | 0.019783855 |
| | Tes | 3.88 | 2.638712666 | 0.00229766831 | 0.026149651 |
| | Klhl6 | 3.88 | 2.45030596 | 0.00354563511 | 0.031513953 |
| | Fam183b | 3.87 | 1.772340681 | 0.01689115389 | 0.072896151 |
| | Ctsh | 3.83 | 2.895949754 | 0.00127072111 | 0.020822928 |
| | Gramd1c | 3.83 | 2.585362731 | 0.00259798877 | 0.027439634 |
| | Gm20743 | 3.82 | 2.059191206 | 0.00872587111 | 0.050078346 |
| | Dbi | 3.82 | 3.571848203 | 0.00026801049 | 0.014200135 |
| | Aqp4 | 3.81 | 3.660419072 | 0.00021856516 | 0.014121412 |
| | Ccdc40 | 3.80 | 1.508269273 | 0.03102635284 | 0.104436713 |
| | St8sia2 | 3.80 | 1.882208532 | 0.01311569980 | 0.062468919 |
| | Dnah9 | 3.80 | 2.870069323 | 0.00134874757 | 0.021189830 |
| | Lox | 3.79 | 2.930524155 | 0.00117348041 | 0.020335620 |
| | Rarres2 | 3.79 | 3.432867621 | 0.00036909009 | 0.015152753 |
| | Col4a6 | 3.78 | 3.080148014 | 0.00083148034 | 0.018142719 |

| | | | | | |
|---|---|---|---|---|---|
| | Paqr6 | 3.77 | 2.649053749 | 0.00224360423 | 0.025964275 |
| | Cnn3 | 3.77 | 3.574484869 | 0.00026638829 | 0.014200135 |
| | Parp12 | 3.77 | 2.579661757 | 0.00263231733 | 0.027675522 |
| | Adamtsl3 | 3.76 | 2.071732404 | 0.00847749604 | 0.049351543 |
| | Asap3 | 3.76 | 2.869347811 | 0.00135099017 | 0.021191731 |
| | Rhbdf2 | 3.75 | 1.523882207 | 0.02993076333 | 0.101991199 |
| | Hspb8 | 3.74 | 3.098111061 | 0.00079779064 | 0.017835958 |
| | Icosl | 3.74 | 2.114686186 | 0.00767916173 | 0.047254528 |
| | Adora3 | 3.74 | 1.808392758 | 0.01554559116 | 0.069693954 |
| | Lgals1 | 3.74 | 4.73837208 | 0.00001826535 | 0.014121412 |
| | Nnat | 3.74 | 2.418889026 | 0.00381163209 | 0.032543773 |
| | Nmi | 3.73 | 2.690050469 | 0.00204150069 | 0.024967673 |
| | Slc12a7 | 3.72 | 2.778448216 | 0.00166552741 | 0.023117184 |
| | S100a11 | 3.72 | 2.70118768 | 0.00198981326 | 0.024654306 |
| | Bgn | 3.71 | 3.659760606 | 0.00021889679 | 0.014121412 |
| | Pon3 | 3.71 | 2.001220522 | 0.00997193588 | 0.053713983 |
| | Pgf | 3.71 | 3.004640316 | 0.00098937216 | 0.019174842 |
| | Padi2 | 3.70 | 3.171266734 | 0.00067411387 | 0.016724165 |
| | Il1a | 3.69 | 1.881831765 | 0.01312708310 | 0.062484266 |
| | Dnaic2 | 3.68 | 1.924200931 | 0.01190690995 | 0.059087342 |
| | Gldn | 3.67 | 2.328899142 | 0.00468922270 | 0.035876731 |
| | Ifit3 | 3.66 | 3.05199897 | 0.00088715812 | 0.018531284 |
| | Naprt | 3.66 | 2.994824397 | 0.00101198856 | 0.019276474 |

| | | | | | |
|---|---|---|---|---|---|
| | Igsf10 | 3.64 | 2.29673818 | 0.00504965631 | 0.037264945 |
| | Klk6 | 3.64 | 1.468791756 | 0.03397881622 | 0.110300712 |
| | Mxra8 | 3.64 | 3.705254869 | 0.00019712655 | 0.014121412 |
| | Pld4 | 3.64 | 2.212480656 | 0.00613083099 | 0.041438145 |
| | Tlr4 | 3.62 | 2.369519159 | 0.00427052079 | 0.034181414 |
| | Ncf1 | 3.62 | 2.551583647 | 0.00280812447 | 0.028624954 |
| | Arhgap30 | 3.62 | 1.931763446 | 0.01170136573 | 0.058426042 |
| | Clic6 | 3.61 | 1.463994732 | 0.03435621153 | 0.111134503 |
| | Myoc | 3.58 | 1.455782529 | 0.03501204443 | 0.112477329 |
| | Igfbp5 | 3.58 | 2.910087993 | 0.00123001953 | 0.020485825 |
| | Ddx58 | 3.58 | 2.315225506 | 0.00483921028 | 0.036396053 |
| | C4a | 3.56 | 1.451553621 | 0.03535463669 | 0.112975240 |
| | Bin2 | 3.55 | 3.249675399 | 0.00056276179 | 0.016356417 |
| | Tmie | 3.54 | 3.60717544 | 0.00024707259 | 0.014200135 |
| | Spp1 | 3.54 | 2.12767323 | 0.00745292532 | 0.046318701 |
| | Irf5 | 3.54 | 2.440134306 | 0.00362965790 | 0.031869092 |
| | Zfp185 | 3.52 | 3.98444546 | 0.00010364648 | 0.014121412 |
| | Adgre1 | 3.52 | 1.932086692 | 0.01169265964 | 0.058426042 |
| | Cfap44 | 3.52 | 1.941806114 | 0.01143388674 | 0.057635010 |
| | Pros1 | 3.51 | 3.697832228 | 0.00020052465 | 0.014121412 |
| | Pdgfrl | 3.51 | 3.187366739 | 0.00064958092 | 0.016529429 |
| | Tlr1 | 3.51 | 3.153312554 | 0.00070256651 | 0.016949337 |
| | Islr | 3.49 | 2.123301752 | 0.00752832306 | 0.046534722 |

| | | | | | |
|---|---|---|---|---|---|
| | Sulf1 | 3.49 | 2.2042623<br>71 | 0.00624795<br>121 | 0.0418142<br>24 |
| | Sla | 3.48 | 2.9291980<br>13 | 0.00117706<br>918 | 0.0203356<br>20 |
| | Cmtm7 | 3.48 | 2.0948429<br>88 | 0.00803816<br>676 | 0.0481694<br>07 |
| | Samd9l | 3.48 | 1.7418505<br>13 | 0.01811963<br>675 | 0.0755406<br>34 |
| | Ccdc88b | 3.47 | 2.0212323<br>48 | 0.00952286<br>553 | 0.0524884<br>46 |
| | Vcam1 | 3.46 | 3.6831495<br>79 | 0.00020741<br>990 | 0.0141214<br>12 |
| | Gpx8 | 3.46 | 3.7199876<br>02 | 0.00019055<br>151 | 0.0141214<br>12 |
| | Tnc | 3.45 | 2.0726030<br>21 | 0.00846051<br>850 | 0.0493515<br>43 |
| | Mns1 | 3.44 | 2.5559917<br>04 | 0.00277976<br>637 | 0.0284413<br>77 |
| | Cd300c2 | 3.44 | 1.4668694<br>03 | 0.03412955<br>275 | 0.1106332<br>12 |
| | Ucp2 | 3.43 | 2.6745449<br>85 | 0.00211570<br>452 | 0.0252579<br>46 |
| | Tbxas1 | 3.42 | 2.2549275<br>77 | 0.00555996<br>967 | 0.0394644<br>88 |
| | Emilin1 | 3.42 | 1.5115043<br>81 | 0.03079609<br>278 | 0.1040750<br>40 |
| | Angpt2 | 3.42 | 3.5370385<br>66 | 0.00029037<br>648 | 0.0143061<br>46 |
| | Glis3 | 3.41 | 2.4926659<br>06 | 0.00321613<br>370 | 0.0299655<br>46 |
| | Vav1 | 3.40 | 2.1418569<br>68 | 0.00721345<br>010 | 0.0454284<br>20 |
| | Slc2a12 | 3.39 | 2.9696925<br>26 | 0.00107227<br>819 | 0.0197838<br>55 |
| | 2810459M11<br>Rik | 3.39 | 3.2001474<br>69 | 0.00063074<br>313 | 0.0165294<br>29 |
| | Smoc1 | 3.39 | 3.7152777<br>44 | 0.00019262<br>926 | 0.0141214<br>12 |
| | Pdpn | 3.38 | 3.6740388<br>99 | 0.00021181<br>714 | 0.0141214<br>12 |
| | Aldh1l2 | 3.37 | 3.0492548<br>46 | 0.00089278<br>144 | 0.0185312<br>84 |
| | Tnfrsf1b | 3.37 | 1.3131403<br>29 | 0.04862500<br>629 | 0.1386503<br>86 |
| | Tnfaip8l2 | 3.36 | 1.8768620<br>72 | 0.01327816<br>093 | 0.0628073<br>01 |

| | | | | | |
|---|---|---|---|---|---|
| | Syngr2 | 3.36 | 3.68028747 | 0.00020879136 | 0.014121412 |
| | Steap3 | 3.35 | 3.366469957 | 0.00043006098 | 0.015871978 |
| | Fabp7 | 3.35 | 2.814011405 | 0.00153457668 | 0.022290294 |
| | Was | 3.34 | 1.553863617 | 0.02793420932 | 0.098119624 |
| | Fermt3 | 3.33 | 1.67793267 | 0.02099265312 | 0.081788822 |
| | Anxa1 | 3.33 | 2.752630981 | 0.00176753906 | 0.023565836 |
| | Ikzf1 | 3.33 | 2.405239917 | 0.00393332727 | 0.033006104 |
| | Slc7a7 | 3.33 | 2.009716219 | 0.00977875985 | 0.053165568 |
| | Fcer1g | 3.32 | 2.108493848 | 0.00778943848 | 0.047698161 |
| | Rasal3 | 3.32 | 1.447734406 | 0.03566691884 | 0.113633001 |
| | Ttc12 | 3.32 | 3.049834071 | 0.00089159152 | 0.018531284 |
| | Lsp1 | 3.31 | 1.341632415 | 0.04553733231 | 0.132794670 |
| | Tlr9 | 3.31 | 1.422414185 | 0.03780818379 | 0.117778792 |
| | Cd37 | 3.31 | 1.919542246 | 0.01203532310 | 0.059360149 |
| | Samsn1 | 3.30 | 1.7945878 | 0.01604767793 | 0.070884150 |
| | Runx1 | 3.30 | 2.221591286 | 0.00600355804 | 0.040986626 |
| | Ube2l6 | 3.29 | 2.097846288 | 0.00798277175 | 0.048025949 |
| | Pgm5 | 3.29 | 3.992944257 | 0.00010163791 | 0.014121412 |
| | Hpse | 3.29 | 1.313907002 | 0.04853924285 | 0.138520817 |
| | Plpp4 | 3.28 | 3.48646626 | 0.00032623739 | 0.014875020 |
| | Maff | 3.28 | 2.066463797 | 0.00858096644 | 0.049625867 |
| | Ncf2 | 3.27 | 3.33566839 | 0.00046166995 | 0.015871978 |
| | Klk8 | 3.27 | 1.99917102 | 0.01001910619 | 0.053750484 |

| | | | | | |
|---|---|---|---|---|---|
| | P4ha3 | 3.26 | 3.227962666 | 0.00059161249 | 0.016487783 |
| | Stom | 3.25 | 2.787862624 | 0.00162981149 | 0.022891833 |
| | Mr1 | 3.24 | 3.580227347 | 0.00026288914 | 0.014200135 |
| | Tlr6 | 3.24 | 1.956224608 | 0.01106051610 | 0.056583229 |
| | Arhgap45 | 3.24 | 2.020397628 | 0.00954118622 | 0.052547270 |
| | Bmp6 | 3.23 | 3.927933247 | 0.00011805021 | 0.014121412 |
| | Hk2 | 3.23 | 1.797708327 | 0.01593278418 | 0.070600576 |
| | Stk32a | 3.22 | 1.709049144 | 0.01954118320 | 0.078444463 |
| | Plscr1 | 3.22 | 1.933719225 | 0.01164878889 | 0.058335681 |
| | Irf9 | 3.20 | 1.724780119 | 0.01884603014 | 0.076924296 |
| | Akna | 3.20 | 2.164967854 | 0.00683962271 | 0.043904822 |
| | Pqlc3 | 3.20 | 2.042844089 | 0.00906057816 | 0.051319623 |
| | Dcn | 3.18 | 2.265234956 | 0.00542956509 | 0.039000342 |
| | Col3a1 | 3.18 | 2.30313844 | 0.00497578446 | 0.037018004 |
| | Klc3 | 3.18 | 2.766437516 | 0.00171223150 | 0.023271537 |
| | Capn3 | 3.16 | 1.995135984 | 0.01011262764 | 0.053922242 |
| | Erap1 | 3.16 | 2.958296152 | 0.00110078841 | 0.019984262 |
| | P2ry6 | 3.15 | 2.107177202 | 0.00781308948 | 0.047709242 |
| | Dpyd | 3.15 | 1.588871011 | 0.02577086460 | 0.093262431 |
| | Lpxn | 3.15 | 1.455370437 | 0.03504528229 | 0.112477329 |
| | Parp3 | 3.14 | 3.303139359 | 0.00049757739 | 0.015871978 |
| | Ifitm1 | 3.14 | 1.64288694 | 0.02275689785 | 0.085883695 |
| | Casp8 | 3.13 | 2.597764136 | 0.00252485164 | 0.027050644 |

| | | | | | |
|---|---|---|---|---|---|
| | Havcr2 | 3.12 | 3.023803291 | 0.00094666585 | 0.018940288 |
| | Lrrc9 | 3.11 | 2.794021908 | 0.00160686019 | 0.022762748 |
| | Lcp2 | 3.11 | 1.301796117 | 0.04991187482 | 0.141028613 |
| | Mdfic | 3.08 | 2.670287681 | 0.00213654635 | 0.025440454 |
| | Zic4 | 3.08 | 2.254682249 | 0.00556311132 | 0.039464488 |
| | Id3 | 3.07 | 3.326583276 | 0.00047142947 | 0.015871978 |
| | Cfh | 3.06 | 3.418836002 | 0.00038120975 | 0.015193137 |
| | Anxa2 | 3.06 | 2.635433703 | 0.00231508157 | 0.026221089 |
| | Tmem176b | 3.05 | 3.459962531 | 0.00034676677 | 0.014875020 |
| | Gstt3 | 3.04 | 2.37787988 | 0.00418909414 | 0.033920060 |
| | Cd53 | 3.03 | 3.018710201 | 0.00095783301 | 0.018992061 |
| | Lgals9 | 3.03 | 1.547699484 | 0.02833351896 | 0.098813884 |
| | Gm973 | 3.03 | 2.83041708 | 0.00147768859 | 0.021914537 |
| | Slc13a4 | 3.03 | 2.925970633 | 0.00118584893 | 0.020344995 |
| | Pik3cg | 3.02 | 3.190210016 | 0.00064534208 | 0.016529429 |
| | Plin2 | 3.02 | 3.282698364 | 0.00052155683 | 0.015979254 |
| | F2r | 3.00 | 2.054096622 | 0.00882883455 | 0.050401751 |
| | Tmem176a | 2.99 | 2.593848832 | 0.00254771690 | 0.027138122 |
| | Lrrk1 | 2.99 | 2.448430996 | 0.00356097565 | 0.031568623 |
| | Plcg2 | 2.98 | 2.392543889 | 0.00405001014 | 0.033464335 |
| | Ptgdr | 2.97 | 1.623379454 | 0.02380238888 | 0.088468657 |
| | Ccl17 | 2.97 | 1.877951575 | 0.01324489212 | 0.062784854 |
| | Casp1 | 2.97 | 1.524846864 | 0.02986435480 | 0.101830650 |

| | | | | | |
|---|---|---|---|---|---|
| | Sema3b | 2.97 | 2.430386884 | 0.00371204400 | 0.032221855 |
| | Ccdc96 | 2.97 | 2.250887996 | 0.00561192689 | 0.039626664 |
| | Aldh1a2 | 2.97 | 2.529384649 | 0.00295539375 | 0.029132041 |
| | Tor4a | 2.96 | 1.650841295 | 0.02234388591 | 0.084814181 |
| | Unc93b1 | 2.96 | 1.696694717 | 0.02010505581 | 0.079660776 |
| | Sntb1 | 2.96 | 2.614349142 | 0.00243024947 | 0.026651584 |
| | Pcsk9 | 2.96 | 1.436341554 | 0.03661495003 | 0.115335181 |
| | Ptges | 2.96 | 3.307815289 | 0.00049224885 | 0.015871978 |
| | Anxa4 | 2.96 | 1.999708578 | 0.01000671250 | 0.053745583 |
| | Hopx | 2.95 | 4.066247807 | 0.00008585235 | 0.014121412 |
| | Col4a5 | 2.95 | 3.224832988 | 0.00059589126 | 0.016487783 |
| | Msn | 2.93 | 3.198142899 | 0.00063366118 | 0.016529429 |
| | Tyrobp | 2.92 | 2.632202741 | 0.00233236899 | 0.026229430 |
| | Wdfy4 | 2.92 | 1.871885757 | 0.01343118229 | 0.063226405 |
| | Dtx3l | 2.91 | 1.556927745 | 0.02773781552 | 0.097704518 |
| | Lst1 | 2.91 | 2.201068267 | 0.00629407238 | 0.041966210 |
| | Ddo | 2.90 | 2.819191971 | 0.00151637993 | 0.022166571 |
| | Iqck | 2.90 | 4.335954296 | 0.00004613661 | 0.014121412 |
| | Trp63 | 2.89 | 1.527627414 | 0.02967376039 | 0.101497148 |
| | Ccdc102a | 2.89 | 1.954170579 | 0.01111295155 | 0.056745844 |
| | S1pr2 | 2.88 | 2.033675415 | 0.00925389538 | 0.051837659 |
| | Arpc1b | 2.88 | 2.459963009 | 0.00346766385 | 0.030987986 |
| | Tagln2 | 2.88 | 2.800944324 | 0.00158145076 | 0.022636161 |

| | | | | | |
|---|---|---|---|---|---|
| | Pla2g4a | 2.88 | 2.967128646 | 0.00107862717 | 0.019788272 |
| | Cnn2 | 2.88 | 2.818638496 | 0.00151831368 | 0.022171252 |
| | Tmem123 | 2.87 | 2.543355835 | 0.00286183219 | 0.028725861 |
| | Gem | 2.86 | 3.066850131 | 0.00085733365 | 0.018362491 |
| | Fbln7 | 2.86 | 2.766840185 | 0.00171064470 | 0.023271537 |
| | Ccr5 | 2.86 | 2.97007139 | 0.00107134318 | 0.019783855 |
| | Arhgdib | 2.85 | 1.630195143 | 0.02343175714 | 0.087623433 |
| | Ncaph | 2.84 | 2.344547908 | 0.00452326562 | 0.035174982 |
| | Fgd2 | 2.84 | 2.177627772 | 0.00664312200 | 0.043244179 |
| | Naaa | 2.84 | 3.591953673 | 0.00025588588 | 0.014200135 |
| | Fam111a | 2.84 | 1.384173875 | 0.04128821663 | 0.124580893 |
| | Vstm4 | 2.84 | 2.889244565 | 0.00129049235 | 0.020978042 |
| | Irak4 | 2.84 | 1.653412589 | 0.02221198701 | 0.084547067 |
| | Hacd4 | 2.83 | 1.791330337 | 0.01616849745 | 0.071163140 |
| | Sparc | 2.83 | 3.316446824 | 0.00048256206 | 0.015871978 |
| | Cyth4 | 2.82 | 2.488748441 | 0.00324527541 | 0.030008970 |
| | Pax6 | 2.82 | 3.508955202 | 0.00030977388 | 0.014819035 |
| | Agt | 2.82 | 2.224648977 | 0.00596143789 | 0.040876306 |
| | Fcgr3 | 2.82 | 2.088031939 | 0.00816522321 | 0.048633867 |
| | Sema3d | 2.82 | 3.416816741 | 0.00038298632 | 0.015193137 |
| | Slc25a45 | 2.82 | 2.925356402 | 0.00118752729 | 0.020344995 |
| | Crip1 | 2.82 | 2.520311469 | 0.00301778663 | 0.029347135 |
| | Gm17750 | 2.80 | 3.234630968 | 0.00058259806 | 0.016472181 |

| | | | | | |
|---|---|---|---|---|---|
| | Catip | 2.80 | 2.349903213 | 0.00446783151 | 0.034842493 |
| | Rbl1 | 2.79 | 1.622037573 | 0.02387604709 | 0.088574720 |
| | Col1a2 | 2.79 | 2.519696188 | 0.00302206507 | 0.029347135 |
| | Aif1 | 2.79 | 2.043507166 | 0.00904675510 | 0.051262458 |
| | Lair1 | 2.79 | 3.154920344 | 0.00069997037 | 0.016949337 |
| | Apod | 2.78 | 3.495352412 | 0.00031963004 | 0.014837961 |
| | Trem2 | 2.78 | 2.499416647 | 0.00316652815 | 0.029738034 |
| | Micall2 | 2.78 | 1.985595498 | 0.01033723769 | 0.054548381 |
| | Mt2 | 2.78 | 4.101385356 | 0.00007917984 | 0.014121412 |
| | Spi1 | 2.78 | 1.386400216 | 0.04107710080 | 0.124144806 |
| | Gjb2 | 2.77 | 2.561350508 | 0.00274567729 | 0.028260938 |
| | Synpo2 | 2.77 | 2.427598863 | 0.00373595070 | 0.032301194 |
| | Aebp1 | 2.77 | 3.293466635 | 0.00050878391 | 0.015871978 |
| | Hs3st3a1 | 2.77 | 2.446868849 | 0.00357380746 | 0.031575132 |
| | Efcab1 | 2.77 | 2.919039877 | 0.00120492530 | 0.020382559 |
| | Plcb2 | 2.77 | 1.429038999 | 0.03723582677 | 0.116593001 |
| | Trim25 | 2.77 | 1.734274188 | 0.01843850951 | 0.076150386 |
| | Ptafr | 2.76 | 1.484131822 | 0.03279957214 | 0.108003576 |
| | Aim2 | 2.76 | 1.748045577 | 0.01786300101 | 0.074893683 |
| | Gsap | 2.76 | 2.758210884 | 0.00174497463 | 0.023409261 |
| | Endou | 2.76 | 1.706446349 | 0.01965864821 | 0.078686130 |
| | Cyp4v3 | 2.76 | 3.125815866 | 0.00074848678 | 0.017398965 |
| | Myof | 2.76 | 2.614080758 | 0.00243175178 | 0.026651584 |

| | | | | | |
|---|---|---|---|---|---|
| | Col1a1 | 2.76 | 2.416383593 | 0.00383368483 | 0.032658812 |
| | Cpq | 2.75 | 3.098581357 | 0.00079692719 | 0.017835958 |
| | Ly75 | 2.75 | 1.45392578 | 0.03516205267 | 0.112572639 |
| | Plin4 | 2.74 | 1.610749559 | 0.02450475931 | 0.090137355 |
| | Adam12 | 2.74 | 1.654900243 | 0.02213603112 | 0.084351415 |
| | Ifitm2 | 2.74 | 2.645190115 | 0.00226365316 | 0.026080691 |
| | Wfikkn2 | 2.74 | 2.880966362 | 0.00131532671 | 0.021114374 |
| | Ptgfr | 2.73 | 2.448613894 | 0.00355947630 | 0.031568623 |
| | Iqcg | 2.73 | 1.827729297 | 0.01486862141 | 0.067764420 |
| | Pde1c | 2.73 | 2.141337635 | 0.00722208116 | 0.045428420 |
| | Dock8 | 2.73 | 1.670306799 | 0.02136452301 | 0.082468256 |
| | Ptgr1 | 2.72 | 1.785665026 | 0.01638079495 | 0.071638607 |
| | Upp1 | 2.71 | 2.630444226 | 0.00234183220 | 0.026264441 |
| | Trp53inp1 | 2.71 | 2.224814952 | 0.00595916002 | 0.040876306 |
| | Postn | 2.71 | 2.144495608 | 0.00716975627 | 0.045273177 |
| | Eya2 | 2.70 | 1.831072903 | 0.01475458835 | 0.067500994 |
| | Clec2d | 2.70 | 1.47756574 | 0.03329923526 | 0.109125874 |
| | Cd63 | 2.70 | 3.491322903 | 0.00032260946 | 0.014875020 |
| | Efemp1 | 2.70 | 2.647950693 | 0.00224930997 | 0.025964275 |
| | Sh3bp2 | 2.69 | 1.725984277 | 0.01879384857 | 0.076836142 |
| | Pawr | 2.69 | 2.316380279 | 0.00482636009 | 0.036359109 |
| | Itih3 | 2.69 | 2.973765208 | 0.00106226970 | 0.019778656 |
| | Nckap1l | 2.68 | 2.357995098 | 0.00438535648 | 0.034705459 |

| | | | | | |
|---|---|---|---|---|---|
| | Itih2 | 2.68 | 2.536677024 | 0.00290618312 | 0.028906123 |
| | Ogn | 2.67 | 2.43310683 | 0.00368886847 | 0.032144431 |
| | Lgi4 | 2.67 | 3.504921085 | 0.00031266475 | 0.014837961 |
| | Csf3r | 2.66 | 2.329594027 | 0.00468172578 | 0.035847206 |
| | Fgfbp1 | 2.66 | 2.406484504 | 0.00392207140 | 0.033001988 |
| | Mx2 | 2.65 | 1.716387554 | 0.01921376375 | 0.077697566 |
| | Cdca7l | 2.64 | 1.79388006 | 0.01607385107 | 0.070951104 |
| | Olfml3 | 2.64 | 2.718950832 | 0.00191006949 | 0.024310539 |
| | Tgm2 | 2.63 | 1.742018358 | 0.01811263529 | 0.075534362 |
| | Samhd1 | 2.63 | 1.834281532 | 0.01464598106 | 0.067150626 |
| | Afap1l2 | 2.63 | 2.616554652 | 0.00241793904 | 0.026631344 |
| | Dera | 2.63 | 2.503654935 | 0.00313577624 | 0.029668607 |
| | Prelp | 2.63 | 2.948575685 | 0.00112570427 | 0.020012642 |
| | Xaf1 | 2.63 | 1.568782602 | 0.02699090196 | 0.096108314 |
| | Rassf9 | 2.63 | 1.751584496 | 0.01771803289 | 0.074537669 |
| | Npl | 2.63 | 3.246728988 | 0.00056659275 | 0.016356417 |
| | Mmp14 | 2.63 | 1.786204816 | 0.01636044771 | 0.071626989 |
| | Prkcd | 2.62 | 2.077555176 | 0.00836459320 | 0.049245461 |
| | Fam180a | 2.62 | 2.250960405 | 0.00561099130 | 0.039626664 |
| | Ormdl2 | 2.62 | 3.963075963 | 0.00010887396 | 0.014121412 |
| | Gcnt1 | 2.62 | 2.769273532 | 0.00170108677 | 0.023254338 |
| | Rarres1 | 2.61 | 1.575886243 | 0.02655300989 | 0.094965121 |
| | Nid2 | 2.60 | 2.281603135 | 0.00522873781 | 0.038135927 |

| | | | | | |
|---|---|---|---|---|---|
| | Vwf | 2.60 | 2.4462921 18 | 0.00357855 653 | 0.0315818 53 |
| | 1700113A16 Rik | 2.60 | 2.6635071 53 | 0.00217016 546 | 0.0255190 26 |
| | Efemp2 | 2.60 | 2.4204471 52 | 0.00379798 153 | 0.0325152 71 |
| | Crabp2 | 2.60 | 1.6215051 64 | 0.02390533 510 | 0.0885904 57 |
| | A330048O09 Rik | 2.60 | 1.6061381 02 | 0.02476634 384 | 0.0906806 77 |
| | Epb41l4a | 2.59 | 1.6652866 76 | 0.02161291 395 | 0.0830489 52 |
| | Cmklr1 | 2.59 | 1.4556580 04 | 0.03502208 485 | 0.1124773 29 |
| | Rab13 | 2.59 | 3.4425968 26 | 0.00036091 354 | 0.0150282 21 |
| | Tapbp | 2.59 | 1.7680383 68 | 0.01705931 672 | 0.0732089 56 |
| | Mvp | 2.58 | 1.6224359 91 | 0.02385415 349 | 0.0885413 08 |
| | Rsph1 | 2.58 | 3.6276496 39 | 0.00023569 500 | 0.0142001 35 |
| | Itpripl1 | 2.58 | 2.0340987 12 | 0.00924488 021 | 0.0518082 79 |
| | E130114P18 Rik | 2.58 | 4.1923231 08 | 0.00006422 097 | 0.0141214 12 |
| | Txnip | 2.57 | 2.3669766 47 | 0.00429559 525 | 0.0342851 07 |
| | Cdh1 | 2.56 | 1.8915908 61 | 0.01283539 205 | 0.0615820 96 |
| | Colec12 | 2.56 | 4.1863824 94 | 0.00006510 547 | 0.0141214 12 |
| | Nfkbie | 2.56 | 2.0166542 02 | 0.00962378 246 | 0.0526854 16 |
| | Svep1 | 2.56 | 2.1443189 36 | 0.00717267 353 | 0.0452731 77 |
| | Loxl1 | 2.56 | 2.5223905 59 | 0.00300337 417 | 0.0293471 35 |
| | Id4 | 2.56 | 2.9572001 | 0.00110357 003 | 0.0199842 62 |
| | Tlr13 | 2.56 | 2.4765952 73 | 0.00333737 284 | 0.0303700 93 |
| | Serpind1 | 2.56 | 3.3740527 72 | 0.00042261 726 | 0.0157846 12 |
| | Ednrb | 2.55 | 3.4232838 92 | 0.00037732 546 | 0.0151931 37 |

| | | | | | |
|---|---|---|---|---|---|
| | Pik3r5 | 2.55 | 1.48851952 | 0.03246986478 | 0.107277810 |
| | Gm13293 | 2.55 | 1.898198915 | 0.01264157208 | 0.061057238 |
| | Plek | 2.55 | 2.022127917 | 0.00950324844 | 0.052479510 |
| | Tmem173 | 2.55 | 1.642555549 | 0.02277426928 | 0.085925655 |
| | Nfe2l2 | 2.54 | 2.647730374 | 0.00225045134 | 0.025964275 |
| | Slc6a13 | 2.54 | 2.401717101 | 0.00396536253 | 0.033139740 |
| | Ccdc80 | 2.54 | 3.334441509 | 0.00046297601 | 0.015871978 |
| | Vamp8 | 2.53 | 2.777307731 | 0.00166990694 | 0.023152817 |
| | Nrp2 | 2.52 | 1.824357403 | 0.01498451177 | 0.068079032 |
| | Gypc | 2.52 | 1.448678526 | 0.03558946623 | 0.113465164 |
| | Itpripl2 | 2.52 | 2.32949857 | 0.00468275493 | 0.035847206 |
| | Tfap2b | 2.51 | 1.603395391 | 0.02492324630 | 0.091082534 |
| | 1110017D15Rik | 2.51 | 2.338963164 | 0.00458180747 | 0.035409796 |
| | Thbs2 | 2.51 | 3.625745788 | 0.00023673050 | 0.014200135 |
| | Rab7b | 2.51 | 2.770147514 | 0.00169766692 | 0.023241590 |
| | Zfp36l1 | 2.50 | 2.921713892 | 0.00119752919 | 0.020344995 |
| | Il10ra | 2.50 | 1.485514115 | 0.03269534203 | 0.107813298 |
| | Cd9 | 2.50 | 3.283262899 | 0.00052087930 | 0.015979254 |
| | Gna15 | 2.50 | 2.513547481 | 0.00306515555 | 0.029428615 |
| | Tifab | 2.49 | 2.168245781 | 0.00678819359 | 0.043793278 |
| | Stat5a | 2.49 | 1.508326739 | 0.03102224766 | 0.104436713 |
| | Entpd2 | 2.49 | 2.712984014 | 0.00193649324 | 0.024345246 |
| | Hn1l | 2.48 | 1.678075415 | 0.02098575437 | 0.081788822 |

| | | | | | |
|---|---|---|---|---|---|
| | Nt5e | 2.48 | 1.857991121 | 0.01386784181 | 0.064492408 |
| | Hist1h2bq | 2.48 | 2.27361662 | 0.00532578193 | 0.038557202 |
| | Hist1h2br | 2.48 | 2.27361662 | 0.00532578193 | 0.038557202 |
| | Uhrf1 | 2.48 | 1.539789091 | 0.02885432433 | 0.099745225 |
| | Ikzf2 | 2.48 | 2.252871204 | 0.00558635841 | 0.039547734 |
| | Gimap6 | 2.48 | 2.325278181 | 0.00472848286 | 0.036016676 |
| | Bmpr1b | 2.47 | 3.295843468 | 0.00050600701 | 0.015871978 |
| | Hmox1 | 2.47 | 2.916058584 | 0.00121322518 | 0.020405052 |
| | Creb3l1 | 2.47 | 2.140841921 | 0.00723032933 | 0.045428420 |
| | Fli1 | 2.47 | 2.632624064 | 0.00233010739 | 0.026229430 |
| | Vasp | 2.47 | 2.000989432 | 0.00997724342 | 0.053718891 |
| | Zfp36 | 2.47 | 2.462064021 | 0.00345092864 | 0.030987986 |
| | Clec4a2 | 2.47 | 2.369968683 | 0.00426610280 | 0.034181414 |
| | Adamts9 | 2.46 | 2.700248926 | 0.00199411901 | 0.024654306 |
| | 4932438H23Rik | 2.46 | 1.678527877 | 0.02096390212 | 0.081788822 |
| | Nt5dc1 | 2.46 | 3.080731326 | 0.00083036431 | 0.018142719 |
| | Pla1a | 2.45 | 1.729165625 | 0.01865668055 | 0.076571520 |
| | Tmem40 | 2.45 | 2.037525814 | 0.00917221415 | 0.051569310 |
| | Mcm3 | 2.45 | 1.829892266 | 0.01479475352 | 0.067539770 |
| | Matn4 | 2.45 | 1.466330123 | 0.03417195903 | 0.110742372 |
| | Phyhd1 | 2.45 | 2.66259735 | 0.00217471650 | 0.025519026 |
| | Nek3 | 2.45 | 2.718982061 | 0.00190993215 | 0.024310539 |
| | Plod2 | 2.45 | 2.665974592 | 0.00215787065 | 0.025517470 |

| | | | | | |
|---|---|---|---|---|---|
| | Tead3 | 2.44 | 2.538972851 | 0.00289086059 | 0.028872829 |
| | Nfam1 | 2.44 | 1.657726637 | 0.02199243731 | 0.083967236 |
| | Col6a3 | 2.44 | 1.318594165 | 0.04801819544 | 0.137605427 |
| | Emid1 | 2.44 | 1.841555572 | 0.01440271701 | 0.066234182 |
| | Rnaset2a | 2.44 | 2.937289831 | 0.00115534096 | 0.020327196 |
| | Zic1 | 2.44 | 1.631358885 | 0.02336905306 | 0.087520893 |
| | Tor3a | 2.43 | 1.511926353 | 0.03076618497 | 0.104025135 |
| | Sdc4 | 2.43 | 4.160736601 | 0.00006906586 | 0.014121412 |
| | Nr1h3 | 2.43 | 2.73559896 | 0.00183823504 | 0.023856991 |
| | Tlr7 | 2.43 | 3.141595878 | 0.00072177880 | 0.017129469 |
| | Prrx1 | 2.43 | 3.416041411 | 0.00038367066 | 0.015193137 |
| | Casp6 | 2.42 | 2.622800855 | 0.00238341213 | 0.026569497 |
| | Ecm2 | 2.42 | 3.606634755 | 0.00024738038 | 0.014200135 |
| | Psme1 | 2.42 | 1.729462205 | 0.01864394421 | 0.076546219 |
| | B4galt1 | 2.42 | 2.160160225 | 0.00691575780 | 0.044097182 |
| | Igf2 | 2.42 | 3.15793783 | 0.00069512382 | 0.016949337 |
| | Ddr2 | 2.42 | 1.952602819 | 0.01115314071 | 0.056883184 |
| | Slc7a11 | 2.41 | 3.214894005 | 0.00060968568 | 0.016529429 |
| | Il1r1 | 2.41 | 1.747483268 | 0.01788614439 | 0.074953800 |
| | Kctd14 | 2.40 | 2.465739353 | 0.00342184747 | 0.030894838 |
| | Ifi27 | 2.40 | 1.366282658 | 0.04302464957 | 0.127878166 |
| | Clec3b | 2.40 | 1.743877642 | 0.01803525794 | 0.075355839 |
| | Celsr1 | 2.40 | 3.005983582 | 0.00098631677 | 0.019169701 |

| | | | | | |
|---|---|---|---|---|---|
| | Kctd11 | 2.39 | 2.872359998 | 0.00134165237 | 0.021189830 |
| | Mlc1 | 2.39 | 3.887235248 | 0.00012964768 | 0.014121412 |
| | Trim56 | 2.39 | 3.275145638 | 0.00053070644 | 0.016055868 |
| | Vwa5a | 2.39 | 2.922078512 | 0.00119652420 | 0.020344995 |
| | Pygl | 2.39 | 2.423960023 | 0.00376738476 | 0.032351180 |
| | Tmem100 | 2.38 | 2.645018535 | 0.00226454766 | 0.026080691 |
| | C1qtnf1 | 2.38 | 2.124860568 | 0.00750135004 | 0.046430654 |
| | Pcolce | 2.38 | 1.873546335 | 0.01337992458 | 0.063074807 |
| | Cebpa | 2.37 | 2.928847858 | 0.00117801858 | 0.020335620 |
| | Clu | 2.37 | 3.619193371 | 0.00024032925 | 0.014200135 |
| | Mrc1 | 2.37 | 1.733026301 | 0.01849156631 | 0.076189689 |
| | Arhgef26 | 2.37 | 4.011788156 | 0.00009732218 | 0.014121412 |
| | Itpkb | 2.37 | 2.558917702 | 0.00276110103 | 0.028377180 |
| | Rinl | 2.37 | 1.533122719 | 0.02930065177 | 0.100750619 |
| | Vdr | 2.37 | 1.58758786 | 0.02584711886 | 0.093419852 |
| | Notch2 | 2.37 | 3.263759064 | 0.00054480481 | 0.016152730 |
| | Eif2ak2 | 2.37 | 2.740617789 | 0.00181711415 | 0.023799215 |
| | Hhex | 2.36 | 2.00244247 | 0.00994391790 | 0.053668254 |
| | Lhfpl3 | 2.36 | 1.311216667 | 0.04884086348 | 0.139063884 |
| | Ip6k3 | 2.36 | 1.39956235 | 0.03985085557 | 0.121578731 |
| | Ifih1 | 2.36 | 2.600707985 | 0.00250779490 | 0.026984816 |
| | Hmgb2 | 2.36 | 1.632290659 | 0.02331896881 | 0.087404787 |
| | Sfrp1 | 2.35 | 2.479934769 | 0.00331180861 | 0.030221111 |

| | | | | | |
|---|---|---|---|---|---|
| | Enkur | 2.35 | 1.375455843 | 0.04212541156 | 0.126027712 |
| | Cd82 | 2.35 | 2.580321508 | 0.00262832152 | 0.027674917 |
| | Lyn | 2.33 | 1.330546556 | 0.04671468709 | 0.135274217 |
| | Hcar1 | 2.33 | 2.508922205 | 0.00309797419 | 0.029582532 |
| | Apobr | 2.33 | 2.00413233 | 0.00990530084 | 0.053543957 |
| | Fkbp10 | 2.33 | 2.221334846 | 0.00600710404 | 0.040986626 |
| | Ahnak | 2.33 | 3.228562379 | 0.00059079610 | 0.016487783 |
| | Rdh5 | 2.32 | 2.435539433 | 0.00366826386 | 0.032023897 |
| | Ampd3 | 2.31 | 2.247485805 | 0.00565606243 | 0.039835958 |
| | Ccdc3 | 2.31 | 1.890074042 | 0.01288029939 | 0.061732889 |
| | Smc4 | 2.31 | 3.11023698 | 0.00077582366 | 0.017679259 |
| | Tspan4 | 2.31 | 3.911891012 | 0.00012249236 | 0.014121412 |
| | Gulp1 | 2.30 | 2.700414568 | 0.00199335859 | 0.024654306 |
| | Spata13 | 2.30 | 1.429790754 | 0.03717142805 | 0.116508347 |
| | Laptm5 | 2.30 | 1.908603689 | 0.01234230604 | 0.060268524 |
| | Ripk1 | 2.29 | 2.952692238 | 0.00111508446 | 0.020012642 |
| | Mmp2 | 2.29 | 1.900233156 | 0.01258249725 | 0.060878907 |
| | Ednra | 2.29 | 2.57376194 | 0.00266832091 | 0.027727676 |
| | Rida | 2.28 | 2.867571677 | 0.00135652663 | 0.021242026 |
| | Pm20d1 | 2.28 | 1.899573985 | 0.01260160942 | 0.060928471 |
| | Bicc1 | 2.28 | 2.484231712 | 0.00327920289 | 0.030148823 |
| | Stat3 | 2.28 | 2.954926868 | 0.00110936161 | 0.020012642 |
| | Prkch | 2.28 | 1.81858116 | 0.01518514141 | 0.068615267 |

| | | | | | |
|---|---|---|---|---|---|
| | Renbp | 2.28 | 2.677237515 | 0.00210262820 | 0.025189376 |
| | Mtbp | 2.27 | 2.147717636 | 0.00711676071 | 0.045052207 |
| | Fam117a | 2.27 | 1.426415879 | 0.03746141013 | 0.117123376 |
| | Hey2 | 2.27 | 2.225962905 | 0.00594342921 | 0.040833454 |
| | Lrig1 | 2.27 | 2.619831691 | 0.00239976275 | 0.026629713 |
| | Psme2 | 2.27 | 1.645772957 | 0.02260617281 | 0.085432184 |
| | Cela1 | 2.26 | 2.323104647 | 0.00475220704 | 0.036097334 |
| | 1700007K13Rik | 2.26 | 2.3886219 3 | 0.00408674999 | 0.033646417 |
| | Mapkapk3 | 2.26 | 1.5779813 5 | 0.02642522235 | 0.094658233 |
| | Fmod | 2.26 | 3.396387471 | 0.00040143250 | 0.015538265 |
| | Tspan12 | 2.26 | 5.2361313 5 | 0.00000580589 | 0.014121412 |
| | F5 | 2.26 | 1.592347211 | 0.02556541162 | 0.092787723 |
| | Nfatc1 | 2.26 | 1.918921565 | 0.01205253593 | 0.059384969 |
| | Pxdc1 | 2.26 | 1.782747153 | 0.01649122235 | 0.071915546 |
| | Rhoj | 2.25 | 3.201298601 | 0.00062907351 | 0.016529429 |
| | S100a13 | 2.25 | 3.954385998 | 0.00011107441 | 0.014121412 |
| | Cetn4 | 2.25 | 2.080829075 | 0.00830177436 | 0.049062156 |
| | Ssc5d | 2.25 | 1.427781759 | 0.03734377702 | 0.116861954 |
| | Mrc2 | 2.25 | 1.750101331 | 0.01777864543 | 0.074726009 |
| | Aox3 | 2.25 | 1.581232753 | 0.02622812512 | 0.094247036 |
| | 1700094D03Rik | 2.25 | 2.168027432 | 0.00679160733 | 0.043793278 |
| | Thbs1 | 2.25 | 1.3852777 3 | 0.04118340687 | 0.124346560 |
| | Alox5ap | 2.24 | 2.404979897 | 0.00393568293 | 0.033006104 |

| | | | | | |
|---|---|---|---|---|---|
| | Tspan18 | 2.24 | 2.037007615 | 0.00918316495 | 0.051609763 |
| | Meis1 | 2.24 | 1.731748596 | 0.01854604906 | 0.076322630 |
| | AI467606 | 2.24 | 1.935421632 | 0.01160321576 | 0.058210948 |
| | Nod1 | 2.24 | 1.9504314 | 0.01120904467 | 0.057036957 |
| | Col8a2 | 2.23 | 1.80599371 | 0.01563170282 | 0.069920322 |
| | Prdx6 | 2.23 | 3.774660377 | 0.00016801174 | 0.014121412 |
| | P2rx7 | 2.23 | 1.614892421 | 0.02427211265 | 0.089464351 |
| | Fam107b | 2.23 | 2.940232926 | 0.00114753799 | 0.020215794 |
| | Galnt4 | 2.23 | 3.040320684 | 0.00091133766 | 0.018746543 |
| | Aldh3b1 | 2.23 | 2.67868287 | 0.00209564218 | 0.025169139 |
| | Dab2 | 2.23 | 2.637284123 | 0.00230523857 | 0.026190933 |
| | Antxr2 | 2.22 | 1.698189728 | 0.02003596535 | 0.079501646 |
| | Lyl1 | 2.22 | 2.503485744 | 0.00313699810 | 0.029668607 |
| | Serhl | 2.22 | 2.836759668 | 0.00145626473 | 0.021867902 |
| | Fam114a1 | 2.22 | 3.175434534 | 0.00066767554 | 0.016682000 |
| | Hmgcs2 | 2.22 | 1.377130189 | 0.04196331716 | 0.125761819 |
| | Apbb1ip | 2.22 | 2.220899741 | 0.00601312538 | 0.041005636 |
| | Slc13a3 | 2.22 | 2.354578084 | 0.00441999641 | 0.034726255 |
| | Cenpj | 2.22 | 2.14152834 | 0.00721891054 | 0.045428420 |
| | Dap | 2.22 | 2.056295686 | 0.00878424245 | 0.050272501 |
| | Rcsd1 | 2.22 | 2.000157843 | 0.00999636619 | 0.053745583 |
| | Nxn | 2.22 | 2.4405088 | 0.00362652938 | 0.031861982 |
| | Acer2 | 2.22 | 2.257022348 | 0.00553321636 | 0.039415203 |

| | | | | | |
|---|---|---|---|---|---|
| | Mdk | 2.21 | 2.555238647 | 0.00278459060 | 0.028469538 |
| | Foxo1 | 2.21 | 2.764416702 | 0.00172021725 | 0.023294207 |
| | Plekhd1 | 2.20 | 1.902551175 | 0.01251551789 | 0.060731728 |
| | Alx4 | 2.20 | 1.365322688 | 0.04311985696 | 0.128054885 |
| | Wnt5a | 2.20 | 2.022223244 | 0.00950116273 | 0.052479510 |
| | Elf1 | 2.20 | 1.746664766 | 0.01791988566 | 0.075072301 |
| | Crispld2 | 2.20 | 1.52014986 | 0.03018909818 | 0.102680297 |
| | Skap2 | 2.19 | 3.101423926 | 0.00079172813 | 0.017776366 |
| | Ccdc122 | 2.19 | 1.811837527 | 0.01542277324 | 0.069188484 |
| | Casp7 | 2.19 | 1.457889923 | 0.03484256167 | 0.112177048 |
| | Filip1l | 2.19 | 1.870437113 | 0.01347605848 | 0.063415931 |
| | Frem2 | 2.19 | 1.866367207 | 0.01360294032 | 0.063754300 |
| | Tst | 2.19 | 2.717887531 | 0.00191475172 | 0.024310539 |
| | Itgam | 2.19 | 2.214581493 | 0.00610124559 | 0.041339850 |
| | Vamp5 | 2.18 | 1.736736835 | 0.01833425067 | 0.075974348 |
| | Rrad | 2.18 | 1.847250163 | 0.01421509730 | 0.065590884 |
| | Papss2 | 2.18 | 2.220423274 | 0.00601972603 | 0.041010978 |
| | Dnase2a | 2.18 | 2.894896141 | 0.00127380767 | 0.020822928 |
| | Bag3 | 2.18 | 2.579184344 | 0.00263521258 | 0.027675522 |
| | Gdpd2 | 2.18 | 3.240338297 | 0.00057499187 | 0.016426119 |
| | Tmem47 | 2.18 | 4.266321103 | 0.00005416003 | 0.014121412 |
| | Antxr1 | 2.18 | 3.312888355 | 0.00048653226 | 0.015871978 |
| | Rnase4 | 2.18 | 3.12204675 | 0.00075501095 | 0.017413556 |

| | | | | | |
|---|---|---|---|---|---|
| | Spidr | 2.17 | 1.964333543 | 0.01085591557 | 0.056058300 |
| | Tgfbr2 | 2.17 | 2.574944469 | 0.00266106529 | 0.027727676 |
| | Bambi | 2.17 | 2.184998063 | 0.00653133466 | 0.042797840 |
| | Csf1 | 2.17 | 2.359146059 | 0.00437374986 | 0.034674128 |
| | Cd68 | 2.17 | 1.700545973 | 0.01992755549 | 0.079323447 |
| | Rcn1 | 2.16 | 2.078802073 | 0.00834061215 | 0.049167032 |
| | Dynlt1c | 2.16 | 1.71823324 | 0.01913228140 | 0.077504917 |
| | Emp2 | 2.16 | 3.138227761 | 0.00072739823 | 0.017232921 |
| | Col9a3 | 2.16 | 3.148939049 | 0.00070967736 | 0.017019400 |
| | Gja1 | 2.16 | 3.948153021 | 0.00011268004 | 0.014121412 |
| | Fam129a | 2.15 | 1.665302192 | 0.02161214181 | 0.083048952 |
| | Sox9 | 2.15 | 2.66873097 | 0.00214421846 | 0.025453666 |
| | Palld | 2.15 | 1.772500798 | 0.01688492758 | 0.072892174 |
| | Hsd3b7 | 2.15 | 2.876462126 | 0.00132903946 | 0.021189830 |
| | Trip6 | 2.15 | 2.125410225 | 0.00749186211 | 0.046429600 |
| | Slc16a9 | 2.15 | 1.873523853 | 0.01338061722 | 0.063074807 |
| | C1ql1 | 2.15 | 1.705428384 | 0.01970478114 | 0.078744570 |
| | Abca1 | 2.15 | 3.331420468 | 0.00046620780 | 0.015871978 |
| | Foxc1 | 2.15 | 1.923227975 | 0.01193361506 | 0.059134441 |
| | Srebf1 | 2.14 | 3.260758054 | 0.00054858250 | 0.016187922 |
| | Hexb | 2.14 | 2.931823218 | 0.00116997554 | 0.020335620 |
| | Ston1 | 2.14 | 2.221316707 | 0.00600735494 | 0.040986626 |
| | Ctsz | 2.14 | 1.735818533 | 0.01837305890 | 0.076066346 |

| | | | | | |
|---|---|---|---|---|---|
| | Rab29 | 2.14 | 2.785024208 | 0.00164049833 | 0.022932214 |
| | Loxl3 | 2.14 | 2.601209327 | 0.00250490161 | 0.026984816 |
| | Rnf135 | 2.13 | 2.437148035 | 0.00365470195 | 0.031988693 |
| | Sqor | 2.13 | 2.905226078 | 0.00124386693 | 0.020617582 |
| | Inpp4b | 2.13 | 2.466483655 | 0.00341598807 | 0.030880982 |
| | Ccr6 | 2.13 | 2.387360043 | 0.00409864173 | 0.033646417 |
| | Spata6 | 2.13 | 2.307667438 | 0.00492416460 | 0.036793336 |
| | Cyp26b1 | 2.13 | 2.508144585 | 0.00310352619 | 0.029594416 |
| | Nfatc4 | 2.13 | 1.438891612 | 0.03640058705 | 0.115024120 |
| | Arhgap18 | 2.13 | 3.579497114 | 0.00026333154 | 0.014200135 |
| | Tmem220 | 2.13 | 2.131998985 | 0.00737905955 | 0.046026172 |
| | Cd59a | 2.12 | 2.521678639 | 0.00300830150 | 0.029347135 |
| | Mboat1 | 2.12 | 1.804751491 | 0.01567647843 | 0.070064091 |
| | Prc1 | 2.12 | 1.422651942 | 0.03778749108 | 0.117741024 |
| | Pbxip1 | 2.12 | 3.729441924 | 0.00018644815 | 0.014121412 |
| | Wipf1 | 2.12 | 3.008630001 | 0.00098032482 | 0.019169701 |
| | Ctso | 2.12 | 3.73288963 | 0.00018497386 | 0.014121412 |
| | Rfx2 | 2.12 | 1.568973771 | 0.02697902364 | 0.096107424 |
| | Plin3 | 2.11 | 2.379717995 | 0.00417140162 | 0.033836617 |
| | Hpgds | 2.11 | 3.401371175 | 0.00039685223 | 0.015470257 |
| | Adgrv1 | 2.11 | 2.620399905 | 0.00239662505 | 0.026629713 |
| | Sncaip | 2.11 | 2.128567173 | 0.00743760016 | 0.046297362 |
| | Rassf4 | 2.11 | 1.615536823 | 0.02423612464 | 0.089383015 |

| | | | | | |
|---|---|---|---|---|---|
| | Trim47 | 2.11 | 2.0595044<br>26 | 0.00871958<br>015 | 0.0500783<br>46 |
| | Itprip | 2.11 | 1.3832836<br>07 | 0.04137294<br>087 | 0.1247269<br>81 |
| | Wnt5b | 2.11 | 2.1410277<br>65 | 0.00722723<br>597 | 0.0454284<br>20 |
| | Cst3 | 2.11 | 3.4629870<br>58 | 0.00034436<br>019 | 0.0148750<br>20 |
| | Gm10790 | 2.11 | 1.6847960<br>67 | 0.02066350<br>230 | 0.0810554<br>34 |
| | Arl5c | 2.11 | 1.3158420<br>38 | 0.04832345<br>324 | 0.1381920<br>02 |
| | Klhl13 | 2.11 | 1.3757650<br>84 | 0.04209542<br>673 | 0.1259928<br>85 |
| | Dna2 | 2.10 | 1.4419741<br>97 | 0.03614313<br>359 | 0.1145129<br>81 |
| | Plekhg2 | 2.10 | 2.4948538<br>9 | 0.00319997<br>150 | 0.0299528<br>67 |
| | Prdm5 | 2.10 | 2.6384855<br>02 | 0.00229887<br>045 | 0.0261496<br>51 |
| | Map3k19 | 2.10 | 1.7645575<br>65 | 0.01719659<br>390 | 0.0734766<br>43 |
| | Rab3il1 | 2.10 | 1.8297004<br>79 | 0.01480128<br>841 | 0.0675459<br>91 |
| | Plcd4 | 2.10 | 2.8859043<br>66 | 0.00130045<br>592 | 0.0209910<br>75 |
| | Smo | 2.09 | 2.9730457<br>89 | 0.00106403<br>083 | 0.0197838<br>55 |
| | Serpinh1 | 2.09 | 2.2483640<br>06 | 0.00564463<br>669 | 0.0397758<br>73 |
| | Zfp217 | 2.09 | 1.7565022<br>94 | 0.01751853<br>179 | 0.0740683<br>52 |
| | Gng5 | 2.09 | 3.6985470<br>22 | 0.00020019<br>489 | 0.0141214<br>12 |
| | Ppp1r3c | 2.09 | 4.2612139<br>25 | 0.00005480<br>070 | 0.0141214<br>12 |
| | Btd | 2.09 | 3.8897212<br>41 | 0.00012890<br>767 | 0.0141214<br>12 |
| | Cox6b2 | 2.09 | 1.5515049<br>3 | 0.02808633<br>493 | 0.0984024<br>29 |
| | C1qtnf6 | 2.08 | 1.9039493<br>86 | 0.01247528<br>896 | 0.0606163<br>17 |
| | Car5b | 2.08 | 2.8337233<br>11 | 0.00146648<br>184 | 0.0219145<br>37 |
| | Spa17 | 2.07 | 1.7539264<br>94 | 0.01762274<br>293 | 0.0742427<br>89 |

| | | | | | |
|---|---|---|---|---|---|
| | Noxo1 | 2.07 | 2.921612237 | 0.00119780953 | 0.020344995 |
| | Clec14a | 2.07 | 2.012520624 | 0.00971581811 | 0.052978197 |
| | Agtrap | 2.07 | 3.283340464 | 0.00052078628 | 0.015979254 |
| | Vangl1 | 2.06 | 1.986908979 | 0.01030602096 | 0.054483415 |
| | Pdgfd | 2.06 | 1.914675065 | 0.01217096280 | 0.059710775 |
| | Mcm6 | 2.06 | 2.624696389 | 0.00237303209 | 0.026510434 |
| | Slc6a20a | 2.06 | 2.537416188 | 0.00290124104 | 0.028888372 |
| | Mrgprf | 2.06 | 1.339818194 | 0.04572795779 | 0.133211388 |
| | Fam181a | 2.06 | 1.815315171 | 0.01529976743 | 0.068870774 |
| | Ltbp1 | 2.06 | 3.352963887 | 0.00044364553 | 0.015871978 |
| | Cdc14a | 2.05 | 2.269901414 | 0.00537153717 | 0.038765910 |
| | Efs | 2.05 | 2.010384402 | 0.00976372634 | 0.053138496 |
| | Btg1 | 2.05 | 2.413885689 | 0.00385579833 | 0.032725150 |
| | Itgb5 | 2.05 | 2.856981023 | 0.00139001337 | 0.021509205 |
| | Ppp1r18 | 2.05 | 2.102703234 | 0.00789399353 | 0.047742678 |
| | Arap1 | 2.05 | 3.019140188 | 0.00095688514 | 0.018992061 |
| | Slc16a1 | 2.05 | 3.59474673 | 0.00025424550 | 0.014200135 |
| | Htra3 | 2.05 | 2.288603048 | 0.00514513711 | 0.037706309 |
| | Tm4sf1 | 2.05 | 2.660001586 | 0.00218775363 | 0.025606408 |
| | Lrrc2 | 2.05 | 2.403404643 | 0.00394998418 | 0.033035139 |
| | Sall3 | 2.05 | 1.770133963 | 0.01697719891 | 0.073032554 |
| | Rlbp1 | 2.04 | 2.157893721 | 0.00695194423 | 0.044286818 |
| | Sorbs1 | 2.04 | 1.854508098 | 0.01397950849 | 0.064852271 |

| | | | | | |
|---|---|---|---|---|---|
| | Eva1b | 2.04 | 1.659207691 | 0.02191756531 | 0.083757684 |
| | Snhg18 | 2.04 | 1.819756723 | 0.01514409331 | 0.068470085 |
| | Hhatl | 2.04 | 4.181532621 | 0.00006583660 | 0.014121412 |
| | Rnaset2b | 2.04 | 1.371720249 | 0.04248931708 | 0.126922980 |
| | Rab27a | 2.04 | 1.380179242 | 0.04166973691 | 0.125382267 |
| | Gatsl3 | 2.03 | 1.962210004 | 0.01090912696 | 0.056248523 |
| | Hist2h2aa1 | 2.03 | 1.794243596 | 0.01606040172 | 0.070914518 |
| | Ostf1 | 2.03 | 1.731943541 | 0.01853772602 | 0.076316810 |
| | Tfcp2l1 | 2.03 | 1.705239283 | 0.01971336292 | 0.078744570 |
| | Tcf7l2 | 2.03 | 1.657122559 | 0.02202304879 | 0.084014079 |
| | Zfp36l2 | 2.03 | 3.196754645 | 0.00063568996 | 0.016529429 |
| | Sbno2 | 2.03 | 1.651890287 | 0.02228998177 | 0.084703164 |
| | Lpar4 | 2.03 | 2.222845244 | 0.00598624871 | 0.040984885 |
| | Foxd1 | 2.03 | 1.388306442 | 0.04089719833 | 0.123997555 |
| | Six5 | 2.02 | 1.658065195 | 0.02197529961 | 0.083925123 |
| | Calcrl | 2.02 | 2.602316808 | 0.00249852208 | 0.026972730 |
| | Add3 | 2.02 | 2.859230067 | 0.00138283363 | 0.021470641 |
| | Thap6 | 2.02 | 1.944244978 | 0.01136985751 | 0.057417572 |
| | Fn1 | 2.02 | 1.911092941 | 0.01227176581 | 0.059988023 |
| | Aif1l | 2.02 | 2.013624939 | 0.00969114431 | 0.052864634 |
| | Slc8b1 | 2.02 | 2.459815105 | 0.00346884500 | 0.030987986 |
| | Prdm16 | 2.02 | 3.311785982 | 0.00048776880 | 0.015871978 |
| | Ifngr1 | 2.02 | 2.664122998 | 0.00216709027 | 0.025519026 |

| | | | | | |
|---|---|---|---|---|---|
| | Cyp4f14 | 2.01 | 4.505132983 | 0.00003125122 | 0.014121412 |
| | Cpne3 | 2.01 | 2.954715199 | 0.00110990243 | 0.020012642 |
| | Cdk5rap2 | 2.01 | 2.002071442 | 0.00995241685 | 0.053682259 |
| | Atp6v0e | 2.01 | 3.241862661 | 0.00057297720 | 0.016426119 |
| | Kif20a | 2.01 | 1.510379082 | 0.03087599187 | 0.104191308 |
| | Hist1h2bc | 2.01 | 2.786427143 | 0.00163520745 | 0.022904572 |
| | Npas3 | 2.01 | 3.582871115 | 0.00026129367 | 0.014200135 |
| | Abhd4 | 2.01 | 3.46379722 | 0.00034371840 | 0.014875020 |
| | Tgfb1 | 2.01 | 1.37567044 | 0.04210460140 | 0.125992885 |
| | Fam161a | 2.01 | 2.018353565 | 0.00958619889 | 0.052584415 |
| | Gal3st4 | 2.01 | 2.63649829 | 0.00230941355 | 0.026204502 |
| | Zfp521 | 2.01 | 2.570434092 | 0.00268884587 | 0.027863824 |
| | Pard3b | 2.00 | 2.46408219 | 0.00343492936 | 0.030950403 |
| | Phkg1 | 2.00 | 3.584834757 | 0.00026011491 | 0.014200135 |
| | Phldb2 | 2.00 | 2.535907562 | 0.00291133672 | 0.028906123 |
| | Bmf | 2.00 | 1.691484973 | 0.02034768598 | 0.080274922 |
| | Ttc41 | 2.00 | 1.380119607 | 0.04167545917 | 0.125382267 |
| | Fzd5 | 2.00 | 1.416460738 | 0.03833003911 | 0.118654387 |
| | Csf1r | 2.00 | 2.633429311 | 0.00232579102 | 0.026229430 |
| | Erbb2 | 1.99 | 2.606083384 | 0.00247694644 | 0.026884456 |
| | Gimap8 | 1.99 | 1.455045903 | 0.03507148027 | 0.112492346 |
| | Pxmp2 | 1.99 | 2.164646228 | 0.00684468982 | 0.043904822 |
| | Nid1 | 1.99 | 2.319367121 | 0.00479328088 | 0.036288966 |

| | | | | | |
|---|---|---|---|---|---|
| | Lamb2 | 1.99 | 3.3071691 | 0.00049298182 | 0.015871978 |
| | Npc2 | 1.98 | 2.901348883 | 0.00125502136 | 0.020677756 |
| | Mt1 | 1.98 | 3.592540808 | 0.00025554018 | 0.014200135 |
| | Pi4k2b | 1.98 | 2.35936924 | 0.00437150279 | 0.034674128 |
| | Tifa | 1.98 | 3.179017061 | 0.00066219049 | 0.016682000 |
| | Cald1 | 1.98 | 3.206973552 | 0.00062090685 | 0.016529429 |
| | Ptgds | 1.98 | 2.316455713 | 0.00482552186 | 0.036359109 |
| | Dse | 1.97 | 1.921587214 | 0.01197878544 | 0.059251437 |
| | Mfsd7b | 1.97 | 2.206912415 | 0.00620994258 | 0.041706169 |
| | Rras | 1.97 | 2.993309885 | 0.00101552382 | 0.019300571 |
| | Sall1 | 1.97 | 2.685391615 | 0.00206351858 | 0.025063632 |
| | Lsm5 | 1.97 | 4.094881518 | 0.00008037454 | 0.014121412 |
| | Cdca7 | 1.97 | 2.277125097 | 0.00528293057 | 0.038401868 |
| | Gusb | 1.97 | 2.248721096 | 0.00563999740 | 0.039763573 |
| | Sec24d | 1.97 | 1.477885401 | 0.03327473450 | 0.109083415 |
| | F3 | 1.97 | 4.037279499 | 0.00009177418 | 0.014121412 |
| | Abca6 | 1.97 | 1.711905289 | 0.01941309193 | 0.078158598 |
| | Npr1 | 1.97 | 1.341986279 | 0.04550024356 | 0.132742855 |
| | Anxa3 | 1.97 | 1.336457672 | 0.04608316808 | 0.133904194 |
| | Galnt12 | 1.96 | 1.960915276 | 0.01094169802 | 0.056340181 |
| | S100a16 | 1.96 | 3.29294871 | 0.00050939103 | 0.015871978 |
| | Lamp2 | 1.96 | 4.222982535 | 0.00005984357 | 0.014121412 |
| | Cmtm6 | 1.96 | 3.739484076 | 0.00018218639 | 0.014121412 |

| | | | | | |
|---|---|---|---|---|---|
| | Mfng | 1.96 | 1.598330016 | 0.02521563936 | 0.091809248 |
| | Cd83 | 1.96 | 1.35181541 | 0.04448202914 | 0.130632093 |
| | Kctd12b | 1.96 | 2.725982969 | 0.00187939052 | 0.024067759 |
| | Def6 | 1.96 | 2.162888021 | 0.00687245618 | 0.043964593 |
| | Cyp4f16 | 1.96 | 1.662522463 | 0.02175091528 | 0.083376207 |
| | Lum | 1.95 | 2.946155965 | 0.00113199377 | 0.020044750 |
| | Aox1 | 1.95 | 2.619580545 | 0.00240115091 | 0.026629713 |
| | Dock11 | 1.95 | 2.107161457 | 0.00781337274 | 0.047709242 |
| | Cd151 | 1.95 | 3.16510172 | 0.00068375148 | 0.016842314 |
| | Kif11 | 1.95 | 1.514416785 | 0.03059026327 | 0.103557725 |
| | Scara3 | 1.95 | 3.485500156 | 0.00032696393 | 0.014875020 |
| | Gpam | 1.95 | 3.784971125 | 0.00016406989 | 0.014121412 |
| | Hacl1 | 1.94 | 1.524727024 | 0.02987259670 | 0.101830650 |
| | Card6 | 1.94 | 2.311602598 | 0.00487974809 | 0.036510908 |
| | Rps27l | 1.94 | 2.531340238 | 0.00294211580 | 0.029054295 |
| | Nedd9 | 1.94 | 2.442201593 | 0.00361242141 | 0.031799028 |
| | Sned1 | 1.94 | 1.592227181 | 0.02557247831 | 0.092788863 |
| | Gli2 | 1.93 | 1.673436204 | 0.02121112953 | 0.082194622 |
| | Tmem119 | 1.93 | 2.29879287 | 0.00502582232 | 0.037208957 |
| | Ltbr | 1.93 | 1.550624935 | 0.02814330293 | 0.098514616 |
| | Pla2g16 | 1.93 | 3.6465431 | 0.00022566120 | 0.014200135 |
| | Adcyap1 | 1.93 | 1.437567064 | 0.03651177415 | 0.115202822 |
| | Tubb6 | 1.93 | 1.438972033 | 0.03639384716 | 0.115024120 |

| | | | | | |
|---|---|---|---|---|---|
| | Ptpn14 | 1.93 | 2.005643115 | 0.00987090302 | 0.053379016 |
| | Rnls | 1.93 | 2.109586593 | 0.00776986382 | 0.047641990 |
| | Pou3f4 | 1.93 | 1.982198497 | 0.01041841140 | 0.054655689 |
| | Wwtr1 | 1.92 | 3.779261223 | 0.00016624124 | 0.014121412 |
| | Crb2 | 1.92 | 1.887047199 | 0.01297038303 | 0.062056418 |
| | BC064078 | 1.92 | 1.785491957 | 0.01638732409 | 0.071644359 |
| | 1700029J07Rik | 1.92 | 2.716189379 | 0.00192225333 | 0.024311630 |
| | Qpct | 1.92 | 2.482502313 | 0.00329228700 | 0.030148823 |
| | Cep112 | 1.92 | 2.550002927 | 0.00281836394 | 0.028627616 |
| | Cd33 | 1.92 | 2.003850191 | 0.00991173790 | 0.053557684 |
| | Ddah2 | 1.92 | 2.643697705 | 0.00227144536 | 0.026096932 |
| | Vcan | 1.91 | 2.074189125 | 0.00842967585 | 0.049351543 |
| | Tbx18 | 1.91 | 1.527093499 | 0.02971026330 | 0.101497148 |
| | Fgfr1 | 1.91 | 3.002248454 | 0.00099483612 | 0.019226502 |
| | 10-Sep | 1.91 | 2.622319844 | 0.00238605338 | 0.026569497 |
| | Fanci | 1.91 | 1.853886857 | 0.01399951993 | 0.064901283 |
| | Ttc23 | 1.91 | 1.61475523 | 0.02427978127 | 0.089468617 |
| | Myh15 | 1.91 | 2.530236099 | 0.00294960527 | 0.029095855 |
| | Gsn | 1.91 | 3.121884618 | 0.00075529287 | 0.017413556 |
| | Gm16576 | 1.91 | 1.489080034 | 0.03242798519 | 0.107216782 |
| | Ccdc190 | 1.91 | 3.276514832 | 0.00052903593 | 0.016055868 |
| | Snap23 | 1.91 | 2.862832792 | 0.00137140967 | 0.021348116 |
| | Rnf43 | 1.90 | 2.053392832 | 0.00884315359 | 0.050462531 |

| | | | | | |
|---|---|---|---|---|---|
| | Nek6 | 1.90 | 2.931623296 | 0.00117051425 | 0.020335620 |
| | Cmtm3 | 1.90 | 2.177774441 | 0.00664087888 | 0.043244179 |
| | Mfap4 | 1.90 | 1.793363242 | 0.01609299060 | 0.070989979 |
| | LOC101056073 | 1.90 | 1.769897046 | 0.01698646285 | 0.073032554 |
| | Hells | 1.90 | 1.769362511 | 0.01700738289 | 0.073053594 |
| | Nupr1 | 1.90 | 2.874694392 | 0.00133446015 | 0.021189830 |
| | S100a1 | 1.90 | 3.536891497 | 0.00029047483 | 0.014306146 |
| | Isoc1 | 1.90 | 1.822925932 | 0.01503398345 | 0.068201376 |
| | 4933407L21Rik | 1.90 | 1.784456196 | 0.01642645331 | 0.071769760 |
| | Cx3cr1 | 1.90 | 2.218082238 | 0.00605226259 | 0.041149995 |
| | Dbx2 | 1.89 | 3.607640292 | 0.00024680827 | 0.014200135 |
| | Tmco4 | 1.89 | 1.473326321 | 0.03362588157 | 0.109569182 |
| | Cpxm1 | 1.89 | 1.388571298 | 0.04087226466 | 0.123986471 |
| | Plcb3 | 1.89 | 2.758386531 | 0.00174426903 | 0.023409261 |
| | Pacrg | 1.89 | 2.734412787 | 0.00184326261 | 0.023894596 |
| | Emp1 | 1.89 | 1.651614025 | 0.02230416530 | 0.084733629 |
| | Ccdc74a | 1.89 | 3.495920399 | 0.00031921229 | 0.014837961 |
| | Angptl7 | 1.89 | 1.664482254 | 0.02165298357 | 0.083156413 |
| | Lrig3 | 1.89 | 2.076180028 | 0.00839112077 | 0.049245461 |
| | Copz2 | 1.89 | 2.038607536 | 0.00914939681 | 0.051569310 |
| | Axl | 1.88 | 3.101733944 | 0.00079116316 | 0.017776366 |
| | Serpinb9 | 1.88 | 1.583719919 | 0.02607834829 | 0.093847003 |
| | Tmem255a | 1.88 | 1.907544541 | 0.01237244293 | 0.060329928 |

| | | | | | |
|---|---|---|---|---|---|
| | Selplg | 1.88 | 2.460880763 | 0.00346034370 | 0.030987986 |
| | Apoe | 1.88 | 2.950180017 | 0.00112155347 | 0.020012642 |
| | Tgfbr1 | 1.88 | 3.797738621 | 0.00015931673 | 0.014121412 |
| | Ppic | 1.88 | 2.116407414 | 0.00764878733 | 0.047130936 |
| | Fkbp14 | 1.88 | 2.642177778 | 0.00227940881 | 0.026149651 |
| | Enpp1 | 1.88 | 3.063696971 | 0.00086358090 | 0.018396503 |
| | Olfml1 | 1.88 | 2.740751768 | 0.00181655366 | 0.023799215 |
| | Abca9 | 1.88 | 2.596768883 | 0.00253064436 | 0.027050644 |
| | Maml2 | 1.88 | 2.106293918 | 0.00782899619 | 0.047709242 |
| | Cep72 | 1.87 | 1.579473065 | 0.02633461264 | 0.094506114 |
| | Fgf1 | 1.87 | 1.683406127 | 0.02072974086 | 0.081199364 |
| | Oat | 1.87 | 4.384635884 | 0.00004124432 | 0.014121412 |
| | Inpp5d | 1.87 | 1.634186144 | 0.02321741455 | 0.087095412 |
| | Cdk2 | 1.87 | 1.758866173 | 0.01742343692 | 0.073848071 |
| | Rfx4 | 1.87 | 3.78890272 | 0.00016259129 | 0.014121412 |
| | St6gal1 | 1.87 | 2.38675624 | 0.00410434406 | 0.033646417 |
| | P3h2 | 1.87 | 2.319903497 | 0.00478736459 | 0.036264155 |
| | Maob | 1.86 | 3.255934724 | 0.00055470908 | 0.016252148 |
| | Etfbkmt | 1.86 | 1.983974742 | 0.01037588760 | 0.054655689 |
| | Als2cl | 1.86 | 2.67967055 | 0.00209088164 | 0.025136312 |
| | Col5a3 | 1.86 | 1.992314833 | 0.01017853248 | 0.054084770 |
| | Tax1bp3 | 1.86 | 2.066653687 | 0.00857721533 | 0.049625059 |
| | Cyp7b1 | 1.85 | 4.219555325 | 0.00006031769 | 0.014121412 |

| | | | | | |
|---|---|---|---|---|---|
| | Shc4 | 1.85 | 1.743941115 | 0.01803262224 | 0.075355839 |
| | Prkd3 | 1.85 | 3.562422079 | 0.00027389110 | 0.014206378 |
| | Art3 | 1.85 | 1.603753404 | 0.02490270912 | 0.091079758 |
| | Fmo5 | 1.85 | 2.378684858 | 0.00418133671 | 0.033877210 |
| | Sox2 | 1.85 | 3.367546339 | 0.00042899641 | 0.015871978 |
| | Limd1 | 1.85 | 2.431436815 | 0.00370308077 | 0.032184714 |
| | Zfp229 | 1.85 | 2.751938138 | 0.00177036112 | 0.023565836 |
| | Gatm | 1.85 | 2.985919456 | 0.00103295296 | 0.019577664 |
| | Fibin | 1.85 | 1.994334116 | 0.01013131653 | 0.053959078 |
| | Siglech | 1.85 | 3.102652584 | 0.00078949142 | 0.017776366 |
| | Pgghg | 1.85 | 2.551764391 | 0.00280695602 | 0.028624954 |
| | Tlr3 | 1.84 | 2.76393318 | 0.00172213352 | 0.023294207 |
| | Cmbl | 1.84 | 1.988005624 | 0.01028002986 | 0.054434640 |
| | Gas1 | 1.84 | 2.327874192 | 0.00470030249 | 0.035927864 |
| | Gpr34 | 1.84 | 4.146704565 | 0.00007133381 | 0.014121412 |
| | Fgfrl1 | 1.84 | 3.643416725 | 0.00022729154 | 0.014200135 |
| | Stk17b | 1.84 | 2.658481131 | 0.00219542633 | 0.025674343 |
| | Loxl2 | 1.84 | 2.168950534 | 0.00677718696 | 0.043782476 |
| | Hspa2 | 1.84 | 2.261465761 | 0.00547689279 | 0.039095057 |
| | Tcn2 | 1.83 | 2.39971978 | 0.00398364125 | 0.033175281 |
| | Lama1 | 1.83 | 2.169713835 | 0.00676528606 | 0.043746727 |
| | Vsig10 | 1.83 | 1.699115723 | 0.01999329051 | 0.079434091 |
| | Lima1 | 1.83 | 2.163004343 | 0.00687061569 | 0.043964593 |

| | | | | | |
|---|---|---|---|---|---|
| | Galnt10 | 1.83 | 1.413623574 | 0.03858126166 | 0.119080215 |
| | Kif19a | 1.83 | 1.608430462 | 0.02463596268 | 0.090393261 |
| | Gm11266 | 1.83 | 2.096894697 | 0.00800028213 | 0.048026575 |
| | Rapgef3 | 1.83 | 2.944001344 | 0.00113762377 | 0.020092658 |
| | Rhbdd1 | 1.83 | 3.36506018 | 0.00043145929 | 0.015871978 |
| | Parvg | 1.82 | 1.741317037 | 0.01814190810 | 0.075610543 |
| | Idh1 | 1.82 | 4.199106372 | 0.00006322570 | 0.014121412 |
| | Parp4 | 1.82 | 2.619895882 | 0.00239940808 | 0.026629713 |
| | Elovl5 | 1.82 | 4.083165563 | 0.00008257231 | 0.014121412 |
| | Gadd45g | 1.82 | 2.461665049 | 0.00345410035 | 0.030987986 |
| | Tirap | 1.82 | 2.059062843 | 0.00872845058 | 0.050078346 |
| | Krcc1 | 1.82 | 3.07705595 | 0.00083742139 | 0.018142719 |
| | Grn | 1.82 | 1.659434487 | 0.02190612256 | 0.083754043 |
| | Ezh2 | 1.82 | 1.802316691 | 0.01576461284 | 0.070194447 |
| | Il10rb | 1.82 | 2.18727151 | 0.00649723373 | 0.042757897 |
| | Rin3 | 1.82 | 1.319128168 | 0.04795918914 | 0.137493682 |
| | Chd7 | 1.82 | 2.042426061 | 0.00906930356 | 0.051326730 |
| | Plxnb2 | 1.82 | 2.101096632 | 0.00792325016 | 0.047790729 |
| | Ngfr | 1.81 | 1.565176806 | 0.02721593093 | 0.096609173 |
| | Vwa1 | 1.81 | 3.302630742 | 0.00049816046 | 0.015871978 |
| | Sptssa | 1.81 | 3.210131478 | 0.00061640836 | 0.016529429 |
| | Pdgfra | 1.81 | 1.503823665 | 0.03134558178 | 0.105162280 |
| | P2ry13 | 1.81 | 2.764658098 | 0.00171926136 | 0.023294207 |

| | | | | | |
|---|---|---|---|---|---|
| | Itga6 | 1.81 | 3.216008196 | 0.00060812352 | 0.016529429 |
| | Fbn1 | 1.80 | 1.557589749 | 0.02769556636 | 0.097680898 |
| | Erbin | 1.80 | 3.394454902 | 0.00040322282 | 0.015548260 |
| | Sspn | 1.80 | 2.128108836 | 0.00744545365 | 0.046297362 |
| | Map3k8 | 1.80 | 1.575756772 | 0.02656092702 | 0.094965121 |
| | Nckap5 | 1.80 | 1.792284977 | 0.01613299587 | 0.071097978 |
| | Slc39a1 | 1.80 | 3.644519816 | 0.00022671496 | 0.014200135 |
| | Cldn10 | 1.80 | 2.573556674 | 0.00266958237 | 0.027727676 |
| | Rp2 | 1.80 | 2.828141634 | 0.00148545112 | 0.021947940 |
| | Fcgrt | 1.80 | 2.567628228 | 0.00270627404 | 0.027939077 |
| | Trim16 | 1.80 | 1.860976461 | 0.01377284115 | 0.064211317 |
| | Scpep1 | 1.80 | 1.500196874 | 0.03160844464 | 0.105676798 |
| | Neat1 | 1.79 | 2.350658453 | 0.00446006869 | 0.034801706 |
| | Slc22a6 | 1.79 | 1.638731931 | 0.02297566388 | 0.086353555 |
| | Bmp4 | 1.79 | 2.066244574 | 0.00858529901 | 0.049630035 |
| | Anpep | 1.79 | 1.484373714 | 0.03278130861 | 0.107993935 |
| | Cdc42bpg | 1.79 | 2.188513575 | 0.00647867844 | 0.042676664 |
| | Slc25a24 | 1.79 | 1.606460541 | 0.02474796306 | 0.090658960 |
| | Tspan6 | 1.79 | 2.490668346 | 0.00323096054 | 0.029977467 |
| | Lfng | 1.79 | 2.597680211 | 0.00252533960 | 0.027050644 |
| | Slc1a3 | 1.79 | 3.346434707 | 0.00045036569 | 0.015871978 |
| | Fbln5 | 1.79 | 2.112239416 | 0.00772254742 | 0.047446521 |
| | Kl | 1.79 | 1.362467819 | 0.04340424252 | 0.128621457 |

| | | | | | |
|---|---|---|---|---|---|
| | Crybg3 | 1.79 | 2.39995193 | 0.00398151238 | 0.033175281 |
| | Rbpms2 | 1.79 | 1.947699951 | 0.01127976493 | 0.057214932 |
| | F11r | 1.79 | 2.055392308 | 0.00880253361 | 0.050314315 |
| | Sat1 | 1.79 | 3.671768439 | 0.00021292740 | 0.014121412 |
| | Slfn5 | 1.79 | 1.543606589 | 0.02860180302 | 0.099357349 |
| | Zic2 | 1.79 | 2.644694923 | 0.00226623570 | 0.026080691 |
| | Entpd1 | 1.78 | 1.676170372 | 0.02107801107 | 0.081956126 |
| | Akr1c14 | 1.78 | 1.424136817 | 0.03765851435 | 0.117495157 |
| | 0610040J01Rik | 1.78 | 2.060825642 | 0.00869309364 | 0.050059592 |
| | Ptbp1 | 1.78 | 1.655719653 | 0.02209430511 | 0.084215769 |
| | Ets1 | 1.78 | 1.320607548 | 0.04779609896 | 0.137226535 |
| | Notch1 | 1.78 | 2.885359677 | 0.00130208796 | 0.020991075 |
| | Acot1 | 1.78 | 1.40160163 | 0.03966416990 | 0.121163944 |
| | Slc31a2 | 1.78 | 1.942210793 | 0.01142323753 | 0.057602461 |
| | Tgfb2 | 1.78 | 3.187821722 | 0.00064890075 | 0.016529429 |
| | Id1 | 1.78 | 1.952472692 | 0.01115648300 | 0.056883184 |
| | Cachd1 | 1.78 | 2.436845655 | 0.00365724744 | 0.031988693 |
| | Scamp2 | 1.78 | 2.355271882 | 0.00441294097 | 0.034726255 |
| | F13a1 | 1.77 | 1.661802775 | 0.02178698955 | 0.083484390 |
| | Cyp2d22 | 1.77 | 4.235098175 | 0.00005819716 | 0.014121412 |
| | Bmp2k | 1.77 | 3.678905389 | 0.00020945687 | 0.014121412 |
| | Cped1 | 1.77 | 1.925748793 | 0.01186454825 | 0.058919681 |
| | Plekha2 | 1.77 | 1.443503827 | 0.03601605765 | 0.114215705 |

| | | | | | |
|---|---|---|---|---|---|
| | Gpr183 | 1.77 | 1.587122332 | 0.02587483972 | 0.093463806 |
| | Pdia4 | 1.77 | 1.573531371 | 0.02669737912 | 0.095359679 |
| | Zbtb20 | 1.77 | 1.586837769 | 0.02589179922 | 0.093474221 |
| | Plpp2 | 1.77 | 1.786019799 | 0.01636741902 | 0.071626989 |
| | Accs | 1.77 | 2.750396613 | 0.00177665616 | 0.023565836 |
| | Apcdd1 | 1.76 | 3.463379137 | 0.00034404945 | 0.014875020 |
| | Dcxr | 1.76 | 2.212815536 | 0.00612610539 | 0.041426582 |
| | Slc16a2 | 1.76 | 2.714589517 | 0.00192934762 | 0.024344505 |
| | Slco2b1 | 1.76 | 2.331110944 | 0.00466540184 | 0.035814127 |
| | Katnal2 | 1.76 | 1.529810192 | 0.02952499334 | 0.101097168 |
| | Grap | 1.76 | 2.910952377 | 0.00122757384 | 0.020485825 |
| | Ugdh | 1.76 | 2.912577793 | 0.00122298803 | 0.020485825 |
| | Fzd7 | 1.76 | 1.315660854 | 0.04834361766 | 0.138220901 |
| | Mavs | 1.76 | 1.541888713 | 0.02871516308 | 0.099496673 |
| | Ikbip | 1.76 | 2.910604277 | 0.00122855817 | 0.020485825 |
| | Lipo3 | 1.76 | 1.725612066 | 0.01880996267 | 0.076879148 |
| | Gm5607 | 1.76 | 1.616898994 | 0.02416022674 | 0.089195507 |
| | Myo10 | 1.76 | 2.70133433 | 0.00198914146 | 0.024654306 |
| | Ntsr2 | 1.75 | 3.146898705 | 0.00071301932 | 0.017019400 |
| | Gna13 | 1.75 | 3.554928592 | 0.00027865793 | 0.014292375 |
| | Tspan15 | 1.75 | 2.259834444 | 0.00549750403 | 0.039201455 |
| | Lrp5 | 1.75 | 2.471347358 | 0.00337794553 | 0.030637854 |
| | Lhfpl2 | 1.75 | 2.896626552 | 0.00126874238 | 0.020822928 |

| | | | | | |
|---|---|---|---|---|---|
| | Lrp10 | 1.75 | 2.620497559 | 0.00239608622 | 0.026629713 |
| | Rps6ka1 | 1.75 | 1.990660144 | 0.01021738729 | 0.054192943 |
| | Mrvi1 | 1.75 | 2.233215911 | 0.00584499425 | 0.040584167 |
| | Ggh | 1.74 | 3.272826681 | 0.00053354778 | 0.016055868 |
| | Tcf7l1 | 1.74 | 1.421260077 | 0.03790879003 | 0.117887608 |
| | Cyp20a1 | 1.74 | 4.119431442 | 0.00007595713 | 0.014121412 |
| | Plpp3 | 1.74 | 3.726733958 | 0.00018761435 | 0.014121412 |
| | P2ry12 | 1.74 | 3.983350092 | 0.00010390822 | 0.014121412 |
| | Fstl1 | 1.74 | 2.444860244 | 0.00359037455 | 0.031665813 |
| | Crtap | 1.74 | 1.594947457 | 0.02541280146 | 0.092331387 |
| | Tmem98 | 1.74 | 2.384365795 | 0.00412699749 | 0.033646417 |
| | Arrdc4 | 1.74 | 1.592770455 | 0.02554050879 | 0.092721831 |
| | Prdx1 | 1.73 | 4.331880399 | 0.00004657143 | 0.014121412 |
| | Rest | 1.73 | 1.330902356 | 0.04667643127 | 0.135226827 |
| | Naga | 1.73 | 2.294635263 | 0.00507416677 | 0.037389130 |
| | Ctbs | 1.73 | 2.790745277 | 0.00161902936 | 0.022854666 |
| | Nde1 | 1.73 | 2.400071528 | 0.00398041608 | 0.033175281 |
| | Hspb6 | 1.73 | 2.707490482 | 0.00196114416 | 0.024453795 |
| | Decr1 | 1.73 | 4.349989721 | 0.00004466942 | 0.014121412 |
| | Fkbp9 | 1.73 | 3.38526221 | 0.00041184879 | 0.015676494 |
| | Fam126a | 1.73 | 1.553339358 | 0.02796795047 | 0.098119926 |
| | Necap2 | 1.73 | 3.05184361 | 0.00088747554 | 0.018531284 |
| | Fbxo36 | 1.72 | 1.831471954 | 0.01474103734 | 0.067473882 |

| | | | | | |
|---|---|---|---|---|---|
| | Hist2h2be | 1.72 | 1.766455945 | 0.01712158849 | 0.073317904 |
| | Proca1 | 1.72 | 1.454891074 | 0.03508398571 | 0.112506195 |
| | Ginm1 | 1.72 | 3.061708051 | 0.00086754488 | 0.018396503 |
| | Serp1 | 1.72 | 2.977987272 | 0.00105199270 | 0.019683538 |
| | Prorsd1 | 1.72 | 1.97031161 | 0.01070750756 | 0.055557879 |
| | Ttc32 | 1.72 | 1.927714309 | 0.01181097340 | 0.058845027 |
| | Prdx4 | 1.72 | 2.474652304 | 0.00335233720 | 0.030465916 |
| | Sowahc | 1.72 | 2.857798775 | 0.00138739851 | 0.021492946 |
| | Tns3 | 1.72 | 3.03826695 | 0.00091565749 | 0.018807249 |
| | Mcm4 | 1.72 | 2.665475399 | 0.00216035241 | 0.025519026 |
| | Itgb3bp | 1.71 | 1.344787919 | 0.04520766556 | 0.132198028 |
| | Pdlim5 | 1.71 | 2.776902827 | 0.00167146456 | 0.023152817 |
| | Rin2 | 1.71 | 2.968590944 | 0.00107500146 | 0.019788272 |
| | Szrd1 | 1.71 | 2.564319017 | 0.00272697391 | 0.028110539 |
| | Nkain4 | 1.71 | 3.907195965 | 0.00012382377 | 0.014121412 |
| | Ctdsp1 | 1.71 | 2.49532049 | 0.00319653534 | 0.029952867 |
| | Sgpl1 | 1.71 | 1.941344347 | 0.01144605038 | 0.057654024 |
| | Gpld1 | 1.71 | 1.802886358 | 0.01574394784 | 0.070144914 |
| | Lap3 | 1.71 | 2.965552474 | 0.00108254891 | 0.019833739 |
| | Plat | 1.71 | 2.193335908 | 0.00640713820 | 0.042388294 |
| | Slc26a2 | 1.71 | 1.317827519 | 0.04810303530 | 0.137791079 |
| | Ppm1m | 1.71 | 2.572846903 | 0.00267394886 | 0.027746394 |
| | Ndp | 1.71 | 2.867327896 | 0.00135728829 | 0.021242026 |

| | | | | | |
|---|---|---|---|---|---|
| | Pth1r | 1.71 | 1.9097393<br>86 | 0.01231007<br>259 | 0.0601324<br>95 |
| | Tnfaip8 | 1.71 | 2.2204123<br>12 | 0.00601987<br>798 | 0.0410109<br>78 |
| | Igsf11 | 1.71 | 2.5840499<br>56 | 0.00260585<br>378 | 0.0275015<br>64 |
| | Adap2 | 1.71 | 1.3613789<br>47 | 0.04351320<br>301 | 0.1288332<br>09 |
| | Soat1 | 1.71 | 2.3261970<br>69 | 0.00471848<br>882 | 0.0359604<br>85 |
| | Bmpr1a | 1.70 | 3.3480168<br>89 | 0.00044872<br>794 | 0.0158719<br>78 |
| | Cnbd2 | 1.70 | 2.7875550<br>8 | 0.00163096<br>604 | 0.0228918<br>33 |
| | Hmgn1 | 1.70 | 2.8648822<br>61 | 0.00136495<br>313 | 0.0213076<br>10 |
| | Adam17 | 1.70 | 2.6186309<br>13 | 0.00240640<br>702 | 0.0266313<br>44 |
| | Ephx1 | 1.70 | 2.0110817<br>84 | 0.00974806<br>050 | 0.0531118<br>55 |
| | Pgpep1 | 1.70 | 2.2552451<br>79 | 0.00555590<br>513 | 0.0394644<br>88 |
| | St5 | 1.70 | 3.0777649<br>6 | 0.00083605<br>537 | 0.0181427<br>19 |
| | Zfp182 | 1.70 | 1.8650087<br>64 | 0.01364555<br>600 | 0.0638577<br>60 |
| | Selenbp1 | 1.70 | 3.2250241<br>78 | 0.00059562<br>898 | 0.0164877<br>83 |
| | Lix1l | 1.69 | 2.1628810<br>08 | 0.00687256<br>715 | 0.0439645<br>93 |
| | Cotl1 | 1.69 | 1.5617391<br>73 | 0.02743221<br>192 | 0.0972009<br>24 |
| | Ccdc8 | 1.69 | 1.3525081<br>49 | 0.04441113<br>278 | 0.1305986<br>06 |
| | Iqgap1 | 1.69 | 1.5896149<br>68 | 0.02572675<br>629 | 0.0932009<br>91 |
| | Ttf2 | 1.69 | 2.2214166<br>74 | 0.00600597<br>232 | 0.0409866<br>26 |
| | Pttg1ip | 1.69 | 2.5179950<br>6 | 0.00303392<br>569 | 0.0293923<br>77 |
| | Adora2b | 1.69 | 2.5197372<br>25 | 0.00302177<br>953 | 0.0293471<br>35 |
| | Tead1 | 1.69 | 2.2002285<br>82 | 0.00630625<br>340 | 0.0420243<br>59 |
| | Magt1 | 1.69 | 3.9105688<br>19 | 0.00012286<br>585 | 0.0141214<br>12 |

| | | | | | |
|---|---|---|---|---|---|
| | Nedd1 | 1.69 | 2.505664277 | 0.00312130152 | 0.029668607 |
| | Smim1 | 1.69 | 2.846500749 | 0.00142396479 | 0.021677178 |
| | Fam69c | 1.69 | 2.347551684 | 0.00449208862 | 0.034991944 |
| | Sugct | 1.69 | 1.972952531 | 0.01064259337 | 0.055351959 |
| | 2-Sep | 1.69 | 3.36427551 | 0.00043223954 | 0.015871978 |
| | Mob3a | 1.69 | 1.44146186 | 0.03618579675 | 0.114595306 |
| | Wdr78 | 1.69 | 1.309672354 | 0.04901484639 | 0.139350193 |
| | Carhsp1 | 1.68 | 1.869551387 | 0.01350357040 | 0.063501903 |
| | Chdh | 1.68 | 1.91719754 | 0.01210047614 | 0.059531916 |
| | Rnd3 | 1.68 | 1.863412106 | 0.01369581540 | 0.064011632 |
| | Cdc42ep4 | 1.68 | 2.8228228 | 0.00150375540 | 0.022052404 |
| | Yap1 | 1.68 | 3.47247578 | 0.00033691800 | 0.014875020 |
| | Sox21 | 1.68 | 2.352606152 | 0.00444011121 | 0.034726255 |
| | Plp2 | 1.68 | 1.71612716 | 0.01922528738 | 0.077721293 |
| | Rftn2 | 1.68 | 2.4918981 | 0.00322182465 | 0.029973658 |
| | Il18 | 1.68 | 4.738363632 | 0.00001826570 | 0.014121412 |
| | Man2b1 | 1.68 | 1.711563666 | 0.01942836858 | 0.078197192 |
| | Mcm2 | 1.68 | 1.627704808 | 0.02356650566 | 0.087877903 |
| | Inhbb | 1.68 | 1.742404794 | 0.01809652577 | 0.075534362 |
| | Orai1 | 1.68 | 1.347068938 | 0.04497084649 | 0.131664294 |
| | Hebp1 | 1.68 | 2.544236761 | 0.00285603312 | 0.028725861 |
| | Pcdhgc3 | 1.67 | 2.915181964 | 0.00121567654 | 0.020421285 |
| | Kctd12 | 1.67 | 1.591014584 | 0.02564397918 | 0.092974649 |

| | | | | | |
|---|---|---|---|---|---|
| | Tpk1 | 1.67 | 2.208945178 | 0.00618094418 | 0.041613108 |
| | Syne3 | 1.67 | 1.595954397 | 0.02535394844 | 0.092190687 |
| | Serpine2 | 1.67 | 3.308165431 | 0.00049185214 | 0.015871978 |
| | Ctla2a | 1.67 | 2.622616764 | 0.00238442264 | 0.026569497 |
| | Rsu1 | 1.67 | 3.248359705 | 0.00056446926 | 0.016356417 |
| | Abcc4 | 1.67 | 1.95645151 | 0.01105473892 | 0.056574737 |
| | Slc22a8 | 1.67 | 1.980679675 | 0.01045491064 | 0.054700813 |
| | 1700008J07Rik | 1.66 | 1.499481831 | 0.03166052912 | 0.105748014 |
| | Ptgs1 | 1.66 | 2.096886237 | 0.00800043797 | 0.048026575 |
| | Homer3 | 1.66 | 1.937908415 | 0.01153696527 | 0.057984433 |
| | Lpp | 1.66 | 1.971816276 | 0.01067047430 | 0.055413072 |
| | Primpol | 1.66 | 2.35208972 | 0.00444539422 | 0.034726641 |
| | Tep1 | 1.66 | 1.474871975 | 0.03350641978 | 0.109335482 |
| | Sox1 | 1.66 | 1.824136042 | 0.01499215135 | 0.068079032 |
| | Laptm4a | 1.66 | 3.308445741 | 0.00049153479 | 0.015871978 |
| | Zfp449 | 1.66 | 2.426930006 | 0.00374170888 | 0.032301194 |
| | Hmgn5 | 1.66 | 3.922147951 | 0.00011963329 | 0.014121412 |
| | Col27a1 | 1.66 | 1.533226875 | 0.02929362550 | 0.100750619 |
| | Plgrkt | 1.66 | 3.131833478 | 0.00073818722 | 0.017360193 |
| | Prrg1 | 1.66 | 1.638834734 | 0.02297022589 | 0.086353555 |
| | Ptpn13 | 1.66 | 2.339901142 | 0.00457192248 | 0.035409796 |
| | Nqo1 | 1.66 | 1.330677848 | 0.04670056687 | 0.135268231 |
| | Gm6644 | 1.65 | 2.332765722 | 0.00464765924 | 0.035717833 |

| | | | | | |
|---|---|---|---|---|---|
| | Arhgef19 | 1.65 | 1.670037612 | 0.02137776937 | 0.082468256 |
| | Plekhf2 | 1.65 | 2.708283268 | 0.00195756744 | 0.024453795 |
| | Dhrs1 | 1.65 | 3.417003941 | 0.00038282127 | 0.015193137 |
| | Fzd6 | 1.65 | 2.677462285 | 0.00210154026 | 0.025189376 |
| | Pld2 | 1.65 | 3.82935218 | 0.00014813164 | 0.014121412 |
| | Elk3 | 1.65 | 1.733855772 | 0.01845628246 | 0.076158492 |
| | Cflar | 1.65 | 1.690231182 | 0.02040651383 | 0.080410454 |
| | Uaca | 1.65 | 2.067704965 | 0.00855647793 | 0.049556285 |
| | Hspg2 | 1.65 | 2.071691119 | 0.00847830197 | 0.049351543 |
| | Nfkbia | 1.65 | 2.077227549 | 0.00837090573 | 0.049245461 |
| | Lpcat3 | 1.65 | 2.21672667 | 0.00607118309 | 0.041217454 |
| | Sh3pxd2b | 1.64 | 1.548857408 | 0.02825807619 | 0.098706250 |
| | Slc16a4 | 1.64 | 2.686454841 | 0.00205847293 | 0.025053566 |
| | Mthfs | 1.64 | 1.536041504 | 0.02910438963 | 0.100281700 |
| | Slc16a12 | 1.64 | 1.631234589 | 0.02337574229 | 0.087522091 |
| | Gng12 | 1.64 | 2.992019853 | 0.00101854483 | 0.019331249 |
| | Acsf2 | 1.64 | 3.313543271 | 0.00048579913 | 0.015871978 |
| | Edn3 | 1.64 | 2.711886481 | 0.00194139327 | 0.024384538 |
| | Itgb1 | 1.64 | 3.212176977 | 0.00061351195 | 0.016529429 |
| | Adrb2 | 1.64 | 1.443276323 | 0.03603492954 | 0.114249185 |
| | Ccnd3 | 1.64 | 1.801163315 | 0.01580653529 | 0.070290486 |
| | Bche | 1.64 | 1.559018305 | 0.02760461502 | 0.097510287 |
| | Rhoc | 1.64 | 1.621336431 | 0.02391462466 | 0.088590457 |

| | | | | | |
|---|---|---|---|---|---|
| | Trim59 | 1.64 | 1.956917879 | 0.01104287410 | 0.056535072 |
| | Zfp808 | 1.64 | 1.495763479 | 0.03193276471 | 0.106218378 |
| | Tmem198b | 1.64 | 2.557262167 | 0.00277164646 | 0.028393015 |
| | BC028528 | 1.64 | 1.981403891 | 0.01043749089 | 0.054673525 |
| | Etohd2 | 1.64 | 1.833497485 | 0.01467244582 | 0.067249526 |
| | 2310022B05Rik | 1.64 | 2.663534612 | 0.00217002825 | 0.025519026 |
| | Lrrc51 | 1.63 | 1.858786517 | 0.01384246657 | 0.064477740 |
| | Il6st | 1.63 | 2.713575464 | 0.00193385780 | 0.024345246 |
| | Nbl1 | 1.63 | 1.787241115 | 0.01632145550 | 0.071626989 |
| | Neu4 | 1.63 | 1.462313727 | 0.03448945032 | 0.111305199 |
| | Snx33 | 1.63 | 2.82168868 | 0.00150768745 | 0.022086496 |
| | Zhx2 | 1.63 | 3.300501387 | 0.00050060895 | 0.015871978 |
| | Cd2ap | 1.63 | 3.547687259 | 0.00028334316 | 0.014306146 |
| | Scrg1 | 1.63 | 2.383887396 | 0.00413154611 | 0.033652386 |
| | Fhod1 | 1.63 | 1.326434848 | 0.04715906133 | 0.136231759 |
| | Tex9 | 1.63 | 2.11149247 | 0.00773584092 | 0.047496957 |
| | Sh3bgrl | 1.63 | 4.402747313 | 0.00003955967 | 0.014121412 |
| | Arsk | 1.63 | 2.427932581 | 0.00373308105 | 0.032301194 |
| | Dnaic1 | 1.63 | 1.580896711 | 0.02624842740 | 0.094295331 |
| | Ddr1 | 1.63 | 2.575098071 | 0.00266012429 | 0.027727676 |
| | Nit2 | 1.63 | 2.978581852 | 0.00105055344 | 0.019683538 |
| | Nat1 | 1.63 | 1.618892606 | 0.02404957432 | 0.088992445 |
| | Man1a | 1.62 | 2.789672195 | 0.00162303470 | 0.022854666 |

| | | | | | |
|---|---|---|---|---|---|
| | Lmcd1 | 1.62 | 1.690181839 | 0.02040883248 | 0.080410454 |
| | Etfrf1 | 1.62 | 4.170863792 | 0.00006747396 | 0.014121412 |
| | Ttc30b | 1.62 | 2.43701662 | 0.00365580801 | 0.031988693 |
| | Fzd1 | 1.62 | 1.45013019 | 0.03547070414 | 0.113191581 |
| | Rpl39 | 1.62 | 3.480742987 | 0.00033056511 | 0.014875020 |
| | Mospd2 | 1.62 | 2.805688327 | 0.00156426984 | 0.022531990 |
| | Slc29a3 | 1.61 | 2.096978824 | 0.00799873256 | 0.048026575 |
| | P2ry14 | 1.61 | 1.390381447 | 0.04070226269 | 0.123709310 |
| | Vsir | 1.61 | 1.656795525 | 0.02203963891 | 0.084054032 |
| | C1qtnf5 | 1.61 | 1.494814249 | 0.03200263592 | 0.106368279 |
| | Zfp953 | 1.61 | 1.54385616 | 0.02858537145 | 0.099357349 |
| | Tsc22d4 | 1.61 | 3.117349947 | 0.00076322054 | 0.017537481 |
| | Gnai3 | 1.61 | 2.542059442 | 0.00287038769 | 0.028747811 |
| | Mmd2 | 1.61 | 3.693617154 | 0.00020248033 | 0.014121412 |
| | Pcsk6 | 1.61 | 2.409874287 | 0.00389157776 | 0.032814001 |
| | Gng11 | 1.61 | 2.103903422 | 0.00787220831 | 0.047709242 |
| | Fadd | 1.61 | 1.587020218 | 0.02588092425 | 0.093463806 |
| | Itpr2 | 1.61 | 2.244526661 | 0.00569473264 | 0.039944523 |
| | Col22a1 | 1.61 | 1.543820728 | 0.02858770369 | 0.099357349 |
| | P2rx6 | 1.61 | 2.35757077 | 0.00438964329 | 0.034705459 |
| | Zeb2os | 1.61 | 2.959547824 | 0.00109762042 | 0.019976692 |
| | Xkr8 | 1.61 | 2.17522797 | 0.00667993182 | 0.043399028 |
| | Tppp3 | 1.61 | 1.418361511 | 0.03816264690 | 0.118407848 |

| | | | | | |
|---|---|---|---|---|---|
| | Pde3b | 1.61 | 1.949855306 | 0.01122392339 | 0.057036957 |
| | Hdac1 | 1.60 | 1.813832743 | 0.01535208114 | 0.069029106 |
| | Psph | 1.60 | 2.97856146 | 0.00105060277 | 0.019683538 |
| | Pla2g7 | 1.60 | 2.340674665 | 0.00456378666 | 0.035369990 |
| | Tprkb | 1.60 | 3.034328409 | 0.00092399919 | 0.018883167 |
| | Ssfa2 | 1.60 | 3.055369151 | 0.00088030030 | 0.018524053 |
| | Mtmr11 | 1.60 | 1.767453023 | 0.01708232486 | 0.073238136 |
| | Smad5 | 1.60 | 2.32991853 | 0.00467822893 | 0.035847206 |
| | Zdhhc12 | 1.60 | 1.714045615 | 0.01931765407 | 0.077911325 |
| | Pls1 | 1.60 | 1.723935335 | 0.01888272484 | 0.076989910 |
| | Ptprz1 | 1.60 | 3.074098818 | 0.00084314289 | 0.018187797 |
| | Rasgrp3 | 1.60 | 1.981277432 | 0.01044053054 | 0.054673525 |
| | Gfpt2 | 1.60 | 1.736287644 | 0.01835322361 | 0.076030041 |
| | Rilp | 1.60 | 1.310210068 | 0.04895419706 | 0.139295768 |
| | Gpr37l1 | 1.60 | 3.467548649 | 0.00034076215 | 0.014875020 |
| | Ddah1 | 1.60 | 2.755861331 | 0.00175444060 | 0.023473971 |
| | Pnrc2 | 1.59 | 3.588047712 | 0.00025819765 | 0.014200135 |
| | Col23a1 | 1.59 | 1.38096988 | 0.04159394564 | 0.125255842 |
| | Mif4gd | 1.59 | 1.407143799 | 0.03916121891 | 0.120195289 |
| | Aff1 | 1.59 | 1.648086251 | 0.02248607986 | 0.085212693 |
| | 9430091E24Rik | 1.59 | 1.476560902 | 0.03337636977 | 0.109189632 |
| | Ccdc173 | 1.59 | 2.022144935 | 0.00950287606 | 0.052479510 |
| | Utp14b | 1.59 | 2.164203811 | 0.00685166608 | 0.043912660 |

| | | | | | |
|---|---|---|---|---|---|
| | Pld1 | 1.59 | 2.753583089 | 0.00176366831 | 0.023565836 |
| | Fam198a | 1.59 | 2.807612646 | 0.00155735404 | 0.022478573 |
| | Chrnb1 | 1.59 | 1.557849751 | 0.02767899065 | 0.097652405 |
| | Pold4 | 1.59 | 1.77442603 | 0.01681024217 | 0.072729703 |
| | Ccdc113 | 1.59 | 1.546315017 | 0.02842398616 | 0.098982057 |
| | Cpt2 | 1.59 | 2.236392534 | 0.00580239735 | 0.040431411 |
| | Tjp2 | 1.59 | 2.589920746 | 0.00257086489 | 0.027215913 |
| | Tmod3 | 1.59 | 2.708734817 | 0.00195553315 | 0.024453795 |
| | Cyp2j6 | 1.59 | 2.92454692 | 0.00118974279 | 0.020344995 |
| | Ctsd | 1.59 | 2.384775164 | 0.00412310919 | 0.033646417 |
| | Phactr4 | 1.59 | 2.638615545 | 0.00229818219 | 0.026149651 |
| | Gm3414 | 1.58 | 1.508530518 | 0.03100769492 | 0.104436713 |
| | Rhpn2 | 1.58 | 1.418126313 | 0.03818332001 | 0.118437246 |
| | Eci1 | 1.58 | 4.10979783 | 0.00007766086 | 0.014121412 |
| | Cyp2j9 | 1.58 | 3.392044688 | 0.00040546681 | 0.015562904 |
| | Tpm4 | 1.58 | 1.881218924 | 0.01314562006 | 0.062524737 |
| | Sfxn2 | 1.58 | 1.553962353 | 0.02792785925 | 0.098119624 |
| | Tln1 | 1.58 | 2.107278514 | 0.00781126706 | 0.047709242 |
| | Gnai2 | 1.58 | 2.383060319 | 0.00413942179 | 0.033676610 |
| | Cpe | 1.58 | 2.422305885 | 0.00378176131 | 0.032417456 |
| | Brca1 | 1.58 | 1.926207178 | 0.01185203218 | 0.058902112 |
| | Rrbp1 | 1.58 | 2.461214362 | 0.00345768669 | 0.030987986 |
| | Ranbp3l | 1.58 | 2.057339468 | 0.00876315578 | 0.050214564 |

| | | | | | |
|---|---|---|---|---|---|
| | Timp3 | 1.58 | 3.871237029 | 0.00013451260 | 0.014121412 |
| | Mcm7 | 1.58 | 1.630875948 | 0.02339505398 | 0.087546688 |
| | Heatr5a | 1.58 | 1.847883927 | 0.01419436840 | 0.065568971 |
| | Nup35 | 1.58 | 1.875004185 | 0.01333508581 | 0.062949311 |
| | Clic4 | 1.58 | 2.522360054 | 0.00300358513 | 0.029347135 |
| | Cd81 | 1.58 | 2.977050997 | 0.00105426309 | 0.019683538 |
| | Lxn | 1.57 | 1.652854396 | 0.02224055415 | 0.084608930 |
| | Myo6 | 1.57 | 3.682555523 | 0.00020770382 | 0.014121412 |
| | Fads2 | 1.57 | 2.672016001 | 0.00212806064 | 0.025383404 |
| | Prkd1 | 1.57 | 2.4477434 | 0.00356661802 | 0.031575132 |
| | Anxa5 | 1.57 | 2.503156774 | 0.00313937522 | 0.029668607 |
| | Zcchc24 | 1.57 | 2.72724983 | 0.00187391622 | 0.024052594 |
| | Stard13 | 1.57 | 2.911582796 | 0.00122579319 | 0.020485825 |
| | Litaf | 1.57 | 1.699369467 | 0.01998161252 | 0.079434091 |
| | Zfp51 | 1.57 | 3.314094024 | 0.00048518345 | 0.015871978 |
| | Wnt7b | 1.57 | 2.4268567 | 0.00374234051 | 0.032301194 |
| | Idh2 | 1.57 | 2.654583987 | 0.00221521566 | 0.025856593 |
| | Rsph9 | 1.57 | 1.352362805 | 0.04442599828 | 0.130598606 |
| | Rom1 | 1.57 | 2.267736541 | 0.00539838009 | 0.038894788 |
| | Nsmce1 | 1.57 | 2.179373908 | 0.00661646610 | 0.043190908 |
| | Hacd1 | 1.57 | 2.708821288 | 0.00195514383 | 0.024453795 |
| | Mob1a | 1.57 | 1.414967594 | 0.03846204801 | 0.118850526 |
| | Sox2ot | 1.57 | 1.671468229 | 0.02130746437 | 0.082400951 |

| | | | | | |
|---|---|---|---|---|---|
| | Pard6g | 1.56 | 2.047749233 | 0.00895881910 | 0.050973497 |
| | Prex2 | 1.56 | 1.86759578 | 0.01356451342 | 0.063647375 |
| | 1600020E01Rik | 1.56 | 1.476619008 | 0.03337190448 | 0.109189632 |
| | Ppp1r14b | 1.56 | 1.511319676 | 0.03080919308 | 0.104093711 |
| | Pign | 1.56 | 2.235672248 | 0.00581202873 | 0.040477996 |
| | Pgap2 | 1.56 | 1.636871556 | 0.02307429518 | 0.086619847 |
| | Slc15a2 | 1.56 | 3.538097406 | 0.00028966938 | 0.014306146 |
| | Shc1 | 1.56 | 1.876917086 | 0.01327647902 | 0.062807301 |
| | Gm3055 | 1.56 | 1.453753685 | 0.03517598887 | 0.112591023 |
| | Leprot | 1.56 | 2.685786514 | 0.00206164310 | 0.025063632 |
| | Cd93 | 1.56 | 1.306754356 | 0.04934528304 | 0.139920251 |
| | Zfp937 | 1.56 | 2.195225834 | 0.00637931674 | 0.042277638 |
| | Tfeb | 1.56 | 1.875136499 | 0.01333102370 | 0.062949311 |
| | 1190005I06Rik | 1.56 | 1.788298443 | 0.01628176779 | 0.071528245 |
| | Perp | 1.56 | 1.49707804 | 0.03183625394 | 0.106025682 |
| | Mfap3l | 1.56 | 3.021988308 | 0.00095063039 | 0.018986355 |
| | Ehd2 | 1.56 | 3.188091705 | 0.00064849748 | 0.016529429 |
| | Lbh | 1.56 | 1.872355895 | 0.01341665046 | 0.063195952 |
| | Igfbp7 | 1.56 | 1.812985151 | 0.01538207233 | 0.069073548 |
| | Sorbs3 | 1.56 | 4.664055992 | 0.00002167425 | 0.014121412 |
| | Ldlrad3 | 1.55 | 1.998356616 | 0.01003791201 | 0.053768732 |
| | Mad2l2 | 1.55 | 2.680950136 | 0.00208473023 | 0.025136312 |
| | Cav1 | 1.55 | 2.835134917 | 0.00146172301 | 0.021903529 |

| | | | | | |
|---|---|---|---|---|---|
| | Znrd1as | 1.55 | 1.462195251 | 0.03449886038 | 0.111305199 |
| | Ninj1 | 1.55 | 1.56062816 | 0.02750247892 | 0.097349707 |
| | Ss18 | 1.55 | 2.293668602 | 0.00508547353 | 0.037430923 |
| | Ppib | 1.55 | 2.600818177 | 0.00250715869 | 0.026984816 |
| | Stxbp3 | 1.55 | 3.387994372 | 0.00040926596 | 0.015621454 |
| | Mro | 1.55 | 2.068734155 | 0.00853622484 | 0.049512987 |
| | Acads | 1.55 | 1.351896551 | 0.04447371915 | 0.130632093 |
| | Lix1 | 1.55 | 2.104005727 | 0.00787035411 | 0.047709242 |
| | Gpc4 | 1.55 | 1.747964202 | 0.01786634837 | 0.074893683 |
| | Gch1 | 1.55 | 1.322551057 | 0.04758268482 | 0.136907288 |
| | Fhl1 | 1.55 | 2.745569303 | 0.00179651438 | 0.023614609 |
| | Sfxn5 | 1.55 | 3.48816073 | 0.00032496701 | 0.014875020 |
| | Rps6ka6 | 1.55 | 2.161584379 | 0.00689311654 | 0.044055257 |
| | Kank2 | 1.54 | 1.952314554 | 0.01116054610 | 0.056883184 |
| | Slc7a2 | 1.54 | 2.68058554 | 0.00208648112 | 0.025136312 |
| | Cep192 | 1.54 | 1.91140057 | 0.01226307629 | 0.059966879 |
| | Phgdh | 1.54 | 2.594483483 | 0.00254399654 | 0.027119516 |
| | Lgmn | 1.54 | 1.629364341 | 0.02347662480 | 0.087708619 |
| | Slc38a3 | 1.54 | 3.399464425 | 0.00039859842 | 0.015472149 |
| | Anapc10 | 1.54 | 3.74377955 | 0.00018039332 | 0.014121412 |
| | Dgka | 1.54 | 1.405899168 | 0.03927361077 | 0.120459528 |
| | Aldh4a1 | 1.54 | 2.490669564 | 0.00323095148 | 0.029977467 |
| | Il11ra1 | 1.54 | 2.377271061 | 0.00419497077 | 0.033927659 |

| | | | | | |
|---|---|---|---|---|---|
| | Pnpla7 | 1.54 | 2.662174117 | 0.00217683686 | 0.025522112 |
| | Gjb6 | 1.54 | 1.962391796 | 0.01090456143 | 0.056246088 |
| | Tulp3 | 1.54 | 1.77716152 | 0.01670469229 | 0.072478427 |
| | Lmo2 | 1.54 | 1.459321137 | 0.03472792728 | 0.111886623 |
| | Thbd | 1.54 | 1.388094705 | 0.04091714233 | 0.123997555 |
| | Ankrd44 | 1.54 | 1.719824412 | 0.01906231262 | 0.077379379 |
| | Tubb2b | 1.54 | 2.793106207 | 0.00161025180 | 0.022787302 |
| | Poc1b | 1.54 | 1.727383337 | 0.01873340246 | 0.076748862 |
| | Reep3 | 1.54 | 3.214320176 | 0.00061049178 | 0.016529429 |
| | Ifnar2 | 1.54 | 1.593206633 | 0.02551487043 | 0.092653233 |
| | Slc39a12 | 1.53 | 3.375387834 | 0.00042132009 | 0.015784612 |
| | Ccdc13 | 1.53 | 1.617063789 | 0.02415106079 | 0.089185629 |
| | Katna1 | 1.53 | 1.455301078 | 0.03505087971 | 0.112477329 |
| | Snhg3 | 1.53 | 2.023127663 | 0.00948139712 | 0.052449226 |
| | Pms1 | 1.53 | 1.524800841 | 0.02986751971 | 0.101830650 |
| | Chic2 | 1.53 | 2.873909737 | 0.00133687334 | 0.021189830 |
| | Oplah | 1.53 | 3.731509718 | 0.00018556253 | 0.014121412 |
| | Mthfsl | 1.53 | 1.707960594 | 0.01959022420 | 0.078526625 |
| | Peli2 | 1.53 | 1.75670343 | 0.01751042029 | 0.074064721 |
| | Ogfod3 | 1.53 | 2.082565853 | 0.00826864120 | 0.048973879 |
| | Pald1 | 1.53 | 1.344388752 | 0.04524923571 | 0.132257650 |
| | Icam2 | 1.53 | 1.427355343 | 0.03738046137 | 0.116929898 |
| | Tmem51 | 1.53 | 1.478136051 | 0.03325553580 | 0.109060696 |

| | | | | | |
|---|---|---|---|---|---|
| | Stab1 | 1.53 | 1.616467698 | 0.02418423208 | 0.089260149 |
| | Rdh10 | 1.53 | 1.776065593 | 0.01674689923 | 0.072592789 |
| | Sft2d2 | 1.53 | 1.879752876 | 0.01319007073 | 0.062627768 |
| | Rnf180 | 1.52 | 1.799525391 | 0.01586626155 | 0.070396610 |
| | Lcat | 1.52 | 1.361566665 | 0.04349439913 | 0.128805288 |
| | Gm2a | 1.52 | 2.823920981 | 0.00149995773 | 0.022050044 |
| | Acadm | 1.52 | 3.308947138 | 0.00049096763 | 0.015871978 |
| | Mtmr10 | 1.52 | 2.550422817 | 0.00281564037 | 0.028627616 |
| | Sgk3 | 1.52 | 2.213959327 | 0.00610999244 | 0.041358328 |
| | Glb1l | 1.52 | 1.527160424 | 0.02970568530 | 0.101497148 |
| | Itgb4 | 1.52 | 1.496952112 | 0.03184548648 | 0.106030732 |
| | Itih5 | 1.52 | 1.803525407 | 0.01572079821 | 0.070084005 |
| | Frmd4b | 1.52 | 1.916799759 | 0.01211156435 | 0.059565142 |
| | Scarb2 | 1.52 | 3.430267952 | 0.00037130607 | 0.015152753 |
| | Nynrin | 1.52 | 2.341898326 | 0.00455094591 | 0.035290377 |
| | Sppl2a | 1.52 | 2.625442304 | 0.00236895983 | 0.026509970 |
| | Kansl1l | 1.52 | 2.365080893 | 0.00431438709 | 0.034367532 |
| | Txnrd3 | 1.52 | 1.453937183 | 0.03516112944 | 0.112572639 |
| | Rplp0 | 1.52 | 1.849306237 | 0.01414795803 | 0.065412884 |
| | Tmed1 | 1.52 | 3.293266984 | 0.00050901785 | 0.015871978 |
| | Arhgap17 | 1.51 | 1.444726404 | 0.03591481188 | 0.114029026 |
| | Nadk2 | 1.51 | 3.869841263 | 0.00013494560 | 0.014121412 |
| | Man2b2 | 1.51 | 1.827533378 | 0.01487533044 | 0.067772518 |

| | | | | | |
|---|---|---|---|---|---|
| | Oard1 | 1.51 | 2.037670502 | 0.00916915887 | 0.051569310 |
| | Slc1a4 | 1.51 | 3.075858842 | 0.00083973288 | 0.018142719 |
| | Znrf2 | 1.51 | 2.386111472 | 0.00411044203 | 0.033646417 |
| | Adgrg1 | 1.51 | 1.64038883 | 0.02288817526 | 0.086118953 |
| | Lims1 | 1.51 | 2.748596104 | 0.00178403716 | 0.023565836 |
| | Selenop | 1.51 | 3.049604264 | 0.00089206343 | 0.018531284 |
| | Dnajc3 | 1.51 | 2.269607896 | 0.00537516876 | 0.038771755 |
| | Pnp | 1.51 | 1.506792414 | 0.03113204043 | 0.104644170 |
| | Rnf182 | 1.51 | 2.245560515 | 0.00568119224 | 0.039912161 |
| | Hexa | 1.51 | 1.763351232 | 0.01724442699 | 0.073543039 |
| | Vkorc1 | 1.51 | 1.874985156 | 0.01333567012 | 0.062949311 |
| | Atp7a | 1.51 | 1.569213985 | 0.02696410531 | 0.096107424 |
| | Slc12a4 | 1.51 | 2.431128025 | 0.00370571466 | 0.032187247 |
| | Acss1 | 1.50 | 2.702094264 | 0.00198566388 | 0.024654306 |
| | Pdk4 | 1.50 | 3.123534884 | 0.00075242829 | 0.017405921 |
| | Rock1 | 1.50 | 3.104100144 | 0.00078686433 | 0.017776366 |
| | Ghr | 1.50 | 2.079450368 | 0.00832817097 | 0.049115165 |
| | Suclg2 | 1.50 | 3.473829685 | 0.00033586930 | 0.014875020 |
| | Gpx7 | 1.50 | 1.865127954 | 0.01364181155 | 0.063857760 |
| | Csk | 1.50 | 1.868750108 | 0.01352850765 | 0.063534657 |
| | Stt3a | 1.50 | 1.846732202 | 0.01423206106 | 0.065647113 |
| | Mfsd1 | 1.50 | 2.413476728 | 0.00385943092 | 0.032725150 |
| | H3f3a | 1.50 | 3.330767335 | 0.00046690945 | 0.015871978 |

| | | | | | |
|---|---|---|---|---|---|
| | Atp11c | 1.50 | 2.631083832 | 0.00233838581 | 0.026264441 |
| | Dhrs4 | 1.50 | 1.607809039 | 0.02467123903 | 0.090498531 |
| | Smpdl3a | 1.50 | 3.190089635 | 0.00064552098 | 0.016529429 |
| | Gdf11 | 1.50 | 2.373656045 | 0.00423003494 | 0.034075549 |
| | Alg14 | 1.50 | 2.435841879 | 0.00366571014 | 0.032021947 |
| | Mettl25 | 1.50 | 2.404315491 | 0.00394170854 | 0.033006104 |
| | Arfip1 | 1.50 | 1.926004522 | 0.01185756402 | 0.058906286 |
| | Cd55 | 1.50 | 1.37010986 | 0.04264716240 | 0.127228541 |
| | Adra2a | 1.50 | 1.452999153 | 0.03523715579 | 0.112708044 |
| | Twsg1 | 1.50 | 2.076260849 | 0.00838955935 | 0.049245461 |
| | Tmsb4x | 1.49 | 2.573545032 | 0.00266965393 | 0.027727676 |
| | Ccdc90b | 1.49 | 1.991694569 | 0.01019307995 | 0.054141133 |
| | Ssr4 | 1.49 | 2.071413941 | 0.00848371477 | 0.049351543 |
| | B4galt4 | 1.49 | 2.798290202 | 0.00159114515 | 0.022676450 |
| | Dock1 | 1.49 | 2.854349221 | 0.00139846235 | 0.021544296 |
| | Cntrl | 1.49 | 1.475264523 | 0.03347614786 | 0.109262648 |
| | F8 | 1.49 | 1.703835455 | 0.01977718813 | 0.078969608 |
| | Pard3 | 1.49 | 2.043930813 | 0.00903793444 | 0.051262458 |
| | Hsd17b11 | 1.49 | 3.713866994 | 0.00019325601 | 0.014121412 |
| | Xrcc4 | 1.49 | 2.177932253 | 0.00663846618 | 0.043244179 |
| | Slc9a3r1 | 1.49 | 2.21704345 | 0.00606675631 | 0.041207760 |
| | Cnpy2 | 1.49 | 2.705212298 | 0.00197145879 | 0.024560123 |
| | AI987944 | 1.49 | 2.547277477 | 0.00283610642 | 0.028655621 |

| | | | | | |
|---|---|---|---|---|---|
| | As3mt | 1.49 | 1.492080431 | 0.03220472305 | 0.106747469 |
| | Cmtm5 | 1.49 | 2.215265135 | 0.00609164890 | 0.041315571 |
| | Cpt1a | 1.49 | 1.766303269 | 0.01712760863 | 0.073317904 |
| | Zfp959 | 1.49 | 2.953194179 | 0.00111379643 | 0.020012642 |
| | Vamp3 | 1.48 | 3.542678748 | 0.00028662974 | 0.014306146 |
| | Ttc7 | 1.48 | 1.761690226 | 0.01731050644 | 0.073619210 |
| | Reck | 1.48 | 1.692866357 | 0.02028306781 | 0.080124081 |
| | Lama4 | 1.48 | 1.479146759 | 0.03317823212 | 0.108885142 |
| | Gprc5b | 1.48 | 3.02727234 | 0.00093913421 | 0.018931026 |
| | Zfp429 | 1.48 | 1.564056892 | 0.02728620317 | 0.096758637 |
| | Plau | 1.48 | 1.648464171 | 0.02246652115 | 0.085185559 |
| | Fbxo2 | 1.48 | 2.640176631 | 0.00228993613 | 0.026149651 |
| | Cwf19l2 | 1.48 | 2.513839169 | 0.00306309757 | 0.029428615 |
| | Pcdhb7 | 1.48 | 1.915614233 | 0.01214467137 | 0.059642577 |
| | Rcbtb2 | 1.48 | 1.47644882 | 0.03338498459 | 0.109189632 |
| | Gpr146 | 1.48 | 2.059793986 | 0.00871376843 | 0.050078346 |
| | Acadl | 1.48 | 2.607606728 | 0.00246827345 | 0.026852745 |
| | Nr2e1 | 1.48 | 2.153473613 | 0.00702306014 | 0.044636387 |
| | Gm5141 | 1.48 | 2.156904037 | 0.00696780459 | 0.044317675 |
| | Plekhg1 | 1.48 | 2.83734374 | 0.00145430755 | 0.021867902 |
| | Gm12191 | 1.48 | 2.843610668 | 0.00143347239 | 0.021677178 |
| | Yes1 | 1.48 | 2.195885407 | 0.00636963569 | 0.042277638 |
| | Zfp53 | 1.48 | 1.878385516 | 0.01323166460 | 0.062781873 |

| | | | | | |
|---|---|---|---|---|---|
| | Rgs10 | 1.48 | 2.238203591 | 0.00577825108 | 0.040324504 |
| | Cecr2 | 1.48 | 1.740472481 | 0.01817722234 | 0.075643008 |
| | Eif4ebp1 | 1.48 | 1.37895343 | 0.04178751735 | 0.125532802 |
| | Cd164 | 1.48 | 3.443776554 | 0.00035993447 | 0.015028221 |
| | Zfp960 | 1.48 | 1.450483592 | 0.03544185199 | 0.113125781 |
| | Slc19a3 | 1.47 | 1.679033716 | 0.02093949887 | 0.081758078 |
| | Sec61b | 1.47 | 1.398478056 | 0.03995047474 | 0.121774506 |
| | Slc22a5 | 1.47 | 1.642401847 | 0.02278233075 | 0.085932475 |
| | Rnh1 | 1.47 | 2.450608277 | 0.00354316782 | 0.031512407 |
| | Pygb | 1.47 | 3.126720355 | 0.00074692956 | 0.017398965 |
| | Cyb5r3 | 1.47 | 2.728558814 | 0.00186827665 | 0.024052594 |
| | Gpr137b | 1.47 | 1.866994798 | 0.01358329716 | 0.063702419 |
| | Nipsnap3b | 1.47 | 2.419166834 | 0.00380919465 | 0.032543773 |
| | C330018D20Rik | 1.47 | 2.423832 | 0.00376849549 | 0.032351180 |
| | Arhgap12 | 1.47 | 2.449311843 | 0.00355376051 | 0.031565755 |
| | 9130023H24Rik | 1.47 | 1.523719748 | 0.02994196180 | 0.101991199 |
| | Chaf1a | 1.47 | 1.546304786 | 0.02842465578 | 0.098982057 |
| | Mtm1 | 1.47 | 1.732707085 | 0.01850516301 | 0.076222855 |
| | Gng10 | 1.47 | 3.124321109 | 0.00075106736 | 0.017403738 |
| | Igdcc4 | 1.47 | 2.576001795 | 0.00265459459 | 0.027727676 |
| | Haus7 | 1.47 | 1.724453797 | 0.01886019605 | 0.076924296 |
| | Lhfp | 1.47 | 2.032001118 | 0.00928963995 | 0.051957065 |
| | Sardh | 1.47 | 2.441119897 | 0.00362143007 | 0.031837537 |

| | | | | | |
|---|---|---|---|---|---|
| | Rab31 | 1.47 | 2.63015971 | 0.00234336689 | 0.026264441 |
| | Cavin3 | 1.47 | 2.192808922 | 0.00641491754 | 0.042419337 |
| | Brip1os | 1.47 | 1.972542442 | 0.01065264755 | 0.055383288 |
| | Lamp1 | 1.46 | 3.220353884 | 0.00060206879 | 0.016529429 |
| | Plekhb1 | 1.46 | 1.994431327 | 0.01012904902 | 0.053959078 |
| | Sema6d | 1.46 | 1.674706786 | 0.02114916443 | 0.082128920 |
| | Ctnna1 | 1.46 | 2.1744992 | 0.00669115052 | 0.043422970 |
| | Helb | 1.46 | 1.427198314 | 0.03739397951 | 0.116939161 |
| | Itm2a | 1.46 | 3.666710948 | 0.00021542150 | 0.014121412 |
| | Ticam1 | 1.46 | 1.555092475 | 0.02785527976 | 0.097951238 |
| | Ctnnal1 | 1.46 | 2.426202944 | 0.00374797820 | 0.032327888 |
| | Rcn2 | 1.46 | 3.526954536 | 0.00029719771 | 0.014430367 |
| | Rpl22l1 | 1.46 | 3.042693946 | 0.00090637111 | 0.018700369 |
| | S100b | 1.46 | 3.010830426 | 0.00097537041 | 0.019169701 |
| | Casp3 | 1.46 | 1.821280861 | 0.01509103891 | 0.068324865 |
| | Rpl13a | 1.46 | 3.379199563 | 0.00041763841 | 0.015765850 |
| | Cttnbp2nl | 1.46 | 1.649911585 | 0.02239176953 | 0.084949007 |
| | Scg3 | 1.46 | 2.715575362 | 0.00192497298 | 0.024311630 |
| | Maoa | 1.46 | 3.863257713 | 0.00013700685 | 0.014121412 |
| | Bcl10 | 1.46 | 2.104447396 | 0.00786235419 | 0.047709242 |
| | Bloc1s1 | 1.46 | 2.446663578 | 0.00357549703 | 0.031575132 |
| | Cyr61 | 1.46 | 1.318875753 | 0.04798707145 | 0.137544920 |
| | Arhgef40 | 1.46 | 1.389104248 | 0.04082213855 | 0.123925394 |

| | | | | | |
|---|---|---|---|---|---|
| | Cbs | 1.46 | 3.092038068 | 0.00080902498 | 0.017943901 |
| | Tmed5 | 1.45 | 3.537932134 | 0.00028977964 | 0.014306146 |
| | A930005H10Rik | 1.45 | 1.307783751 | 0.04922845990 | 0.139646628 |
| | Atp1b2 | 1.45 | 2.538606733 | 0.00289329866 | 0.028872829 |
| | Ech1 | 1.45 | 2.263336422 | 0.00545335258 | 0.039027548 |
| | Crlf3 | 1.45 | 1.601618504 | 0.02502542695 | 0.091237567 |
| | Tmem150a | 1.45 | 1.540008521 | 0.02883974917 | 0.099719928 |
| | Lbr | 1.45 | 1.705511102 | 0.01970102845 | 0.078744570 |
| | Jam3 | 1.45 | 1.412249918 | 0.03870348589 | 0.119269926 |
| | Egflam | 1.45 | 1.498877763 | 0.03170459695 | 0.105785453 |
| | Zfyve21 | 1.45 | 1.785977148 | 0.01636902649 | 0.071626989 |
| | Nqo2 | 1.45 | 2.073463992 | 0.00844376248 | 0.049351543 |
| | Trip10 | 1.45 | 1.817126413 | 0.01523609203 | 0.068777641 |
| | Lats2 | 1.45 | 2.148136098 | 0.00710990671 | 0.045042521 |
| | Lpar1 | 1.45 | 1.411979376 | 0.03872760354 | 0.119317489 |
| | Chpt1 | 1.45 | 2.805215848 | 0.00156597257 | 0.022531990 |
| | Afap1 | 1.45 | 1.919712416 | 0.01203060823 | 0.059358200 |
| | Cdo1 | 1.45 | 1.815799007 | 0.01528273186 | 0.068854277 |
| | Tmed3 | 1.44 | 2.99445354 | 0.00101285310 | 0.019276474 |
| | Gstk1 | 1.44 | 2.182510003 | 0.00656885986 | 0.043002717 |
| | Slc39a11 | 1.44 | 1.762171755 | 0.01729132388 | 0.073608873 |
| | Slc44a5 | 1.44 | 1.442276036 | 0.03611802246 | 0.114459812 |
| | Mplkip | 1.44 | 1.805415295 | 0.01565253578 | 0.069990724 |

| | | | | | |
|---|---|---|---|---|---|
| | Rps24 | 1.44 | 3.56701454 | 0.00027101009 | 0.014206378 |
| | Abhd3 | 1.44 | 3.57887525 | 0.00026370888 | 0.014200135 |
| | Golim4 | 1.44 | 2.276974465 | 0.00528476324 | 0.038401868 |
| | Sat2 | 1.44 | 1.412476627 | 0.03868328730 | 0.119269926 |
| | Sil1 | 1.44 | 2.521716296 | 0.00300804067 | 0.029347135 |
| | Egfr | 1.44 | 1.551657568 | 0.02807646534 | 0.098392938 |
| | Marcks | 1.44 | 1.815114344 | 0.01530684398 | 0.068870774 |
| | Ppfibp2 | 1.44 | 1.763534987 | 0.01723713221 | 0.073534751 |
| | Crtc3 | 1.44 | 1.897326068 | 0.01266700471 | 0.061158578 |
| | B9d1 | 1.44 | 1.885370932 | 0.01302054215 | 0.062253051 |
| | Dhx40 | 1.44 | 2.077268234 | 0.00837012157 | 0.049245461 |
| | Raver2 | 1.44 | 2.425952686 | 0.00375013856 | 0.032327888 |
| | Cdca4 | 1.44 | 1.791855535 | 0.01614895653 | 0.071109869 |
| | Pkd2 | 1.44 | 1.891965896 | 0.01282431284 | 0.061550431 |
| | Nfe2l3 | 1.44 | 1.336454847 | 0.04608346790 | 0.133904194 |
| | Rbm7 | 1.44 | 2.789593643 | 0.00162332829 | 0.022854666 |
| | Gab1 | 1.44 | 1.342760666 | 0.04541918474 | 0.132675386 |
| | Col11a2 | 1.44 | 1.853257812 | 0.01401981193 | 0.064949171 |
| | Asah1 | 1.44 | 3.030713807 | 0.00093172166 | 0.018883167 |
| | Ntrk2 | 1.44 | 1.844404897 | 0.01430853278 | 0.065933450 |
| | Thap3 | 1.44 | 1.990608187 | 0.01021860974 | 0.054192943 |
| | Ezr | 1.44 | 3.016858958 | 0.00096192462 | 0.018992061 |
| | Nfia | 1.44 | 2.090235543 | 0.00812389791 | 0.048541907 |

| | | | | | |
|---|---|---|---|---|---|
| | Tom1l1 | 1.44 | 3.274780072 | 0.00053115335 | 0.016055868 |
| | Eci2 | 1.44 | 3.360046943 | 0.00043646865 | 0.015871978 |
| | Sash1 | 1.44 | 2.072003965 | 0.00847219679 | 0.049351543 |
| | Gpt2 | 1.43 | 2.259414772 | 0.00550281900 | 0.039219002 |
| | Odc1 | 1.43 | 2.1521468 | 0.00704454909 | 0.044731585 |
| | Pcdhb8 | 1.43 | 1.394944606 | 0.04027684037 | 0.122578973 |
| | Kat2b | 1.43 | 2.348627109 | 0.00448097881 | 0.034925201 |
| | Vav3 | 1.43 | 1.734032433 | 0.01844877638 | 0.076150386 |
| | Dennd4c | 1.43 | 1.940291096 | 0.01147384303 | 0.057730530 |
| | Zfp934 | 1.43 | 1.327700888 | 0.04702178509 | 0.135912147 |
| | Edem2 | 1.43 | 1.945411383 | 0.01133936193 | 0.057386054 |
| | Prom1 | 1.43 | 2.022003672 | 0.00950596757 | 0.052479510 |
| | Rassf8 | 1.43 | 1.656478243 | 0.02205574628 | 0.084092123 |
| | Nat6 | 1.43 | 1.37050014 | 0.04260885466 | 0.127169477 |
| | 1810037I17Rik | 1.43 | 3.654180244 | 0.00022172760 | 0.014180233 |
| | Kcnn3 | 1.43 | 1.614008321 | 0.02432157408 | 0.089574578 |
| | Manba | 1.43 | 1.785873711 | 0.01637292564 | 0.071626989 |
| | Klf15 | 1.43 | 2.967630789 | 0.00107738075 | 0.019788272 |
| | Btg3 | 1.43 | 2.576787802 | 0.00264979452 | 0.027727676 |
| | Gm5069 | 1.43 | 1.768814378 | 0.01702886185 | 0.073122997 |
| | Rgcc | 1.43 | 1.714610699 | 0.01929253517 | 0.077878592 |
| | Mitd1 | 1.43 | 1.468586643 | 0.03399486784 | 0.110326755 |
| | Ccdc34 | 1.43 | 1.505134439 | 0.03125111820 | 0.104967396 |

| | | | | | |
|---|---|---|---|---|---|
| | Capza1 | 1.43 | 1.440365307 | 0.03627727802 | 0.114726370 |
| | Ptma | 1.43 | 2.484096078 | 0.00328022717 | 0.030148823 |
| | Adk | 1.43 | 2.996919663 | 0.00100711795 | 0.019247299 |
| | Naa16 | 1.43 | 1.971913878 | 0.01066807650 | 0.055413072 |
| | Rnf13 | 1.43 | 3.627646279 | 0.00023569682 | 0.014200135 |
| | Ston2 | 1.43 | 1.978278458 | 0.01051287599 | 0.054875911 |
| | Hacd2 | 1.42 | 2.29829557 | 0.00503158055 | 0.037231528 |
| | Pltp | 1.42 | 2.230991879 | 0.00587500338 | 0.040628295 |
| | Trps1 | 1.42 | 1.74986601 | 0.01778828134 | 0.074726009 |
| | Gpr137b-ps | 1.42 | 1.646673924 | 0.02255932372 | 0.085349027 |
| | Rhog | 1.42 | 1.617757688 | 0.02411250395 | 0.089115093 |
| | Hapln1 | 1.42 | 2.6814804 | 0.00208218637 | 0.025136312 |
| | Hmmr | 1.42 | 1.30871094 | 0.04912347259 | 0.139463975 |
| | Fuca1 | 1.42 | 1.866048746 | 0.01361291881 | 0.063754300 |
| | Pon2 | 1.42 | 2.395470914 | 0.00402280597 | 0.033380058 |
| | Txndc5 | 1.42 | 1.386824336 | 0.04103700566 | 0.124095399 |
| | Ldlr | 1.42 | 3.602365922 | 0.00024982395 | 0.014200135 |
| | Tcf3 | 1.42 | 1.697493437 | 0.02006811417 | 0.079583249 |
| | Rps4x | 1.42 | 2.547269612 | 0.00283615778 | 0.028655621 |
| | Sumf1 | 1.42 | 2.848534533 | 0.00141731201 | 0.021674489 |
| | Erlin2 | 1.42 | 2.438837814 | 0.00364050965 | 0.031923576 |
| | Rps20 | 1.42 | 2.423395401 | 0.00377228589 | 0.032356417 |
| | Fam181b | 1.42 | 1.95578842 | 0.01107163043 | 0.056619008 |

| | | | | | |
|---|---|---|---|---|---|
| | Kctd5 | 1.42 | 2.466558064 | 0.00341540285 | 0.030880982 |
| | Wasf2 | 1.42 | 1.553307558 | 0.02796999845 | 0.098119926 |
| | Lactb2 | 1.41 | 1.669456354 | 0.02140640047 | 0.082509214 |
| | Evc | 1.41 | 1.436975366 | 0.03656155298 | 0.115254026 |
| | Kyat3 | 1.41 | 1.618785546 | 0.02405550362 | 0.088992445 |
| | Abcg2 | 1.41 | 2.140222231 | 0.00724065357 | 0.045472496 |
| | Etfdh | 1.41 | 3.248084831 | 0.00056482664 | 0.016356417 |
| | Rps27rt | 1.41 | 2.49503515 | 0.00319863622 | 0.029952867 |
| | Wdr5b | 1.41 | 2.108951566 | 0.00778123325 | 0.047673920 |
| | 1810058I24Rik | 1.41 | 3.533708914 | 0.00029261129 | 0.014331583 |
| | Tiparp | 1.41 | 1.652989612 | 0.02223363071 | 0.084606015 |
| | Cbfb | 1.41 | 1.720667292 | 0.01902535232 | 0.077345375 |
| | Adh5 | 1.41 | 3.881797815 | 0.00013128109 | 0.014121412 |
| | Hltf | 1.41 | 1.602050233 | 0.02500056173 | 0.091195306 |
| | Tmem219 | 1.41 | 2.10123764 | 0.00792067803 | 0.047790729 |
| | Stx2 | 1.41 | 1.932004575 | 0.01169487071 | 0.058426042 |
| | Cr1l | 1.41 | 1.733122786 | 0.01848745856 | 0.076189689 |
| | Tor1aip1 | 1.41 | 1.75496958 | 0.01758046753 | 0.074124948 |
| | Fancl | 1.41 | 1.328912437 | 0.04689079149 | 0.135647656 |
| | Tmem205 | 1.41 | 1.671246513 | 0.02131834506 | 0.082400951 |
| | Nemp1 | 1.41 | 1.424269412 | 0.03764701857 | 0.117495157 |
| | Rack1 | 1.41 | 1.631003236 | 0.02338819810 | 0.087544873 |
| | Hepacam | 1.41 | 2.67583965 | 0.00210940684 | 0.025226597 |

| | | | | | |
|---|---|---|---|---|---|
| | Agbl3 | 1.41 | 1.3685531 24 | 0.04280030 623 | 0.1274916 56 |
| | Hps5 | 1.41 | 1.7223204 62 | 0.01895306 877 | 0.0771199 64 |
| | Cers2 | 1.41 | 1.7396300 75 | 0.01821251 509 | 0.0757210 80 |
| | Aga | 1.41 | 1.5501575 94 | 0.02817360 403 | 0.0985574 07 |
| | Atf5 | 1.41 | 1.6414936 6 | 0.02283002 253 | 0.0860651 14 |
| | Svbp | 1.40 | 1.5655780 99 | 0.02719079 469 | 0.0965844 30 |
| | Galnt1 | 1.40 | 2.9229986 32 | 0.00119399 187 | 0.0203449 95 |
| | Rps18 | 1.40 | 2.1797314 81 | 0.00661102 073 | 0.0431758 73 |
| | Nfatc3 | 1.40 | 1.6467000 77 | 0.02255796 521 | 0.0853490 27 |
| | Tmcc3 | 1.40 | 1.5654002 51 | 0.02720193 187 | 0.0965844 30 |
| | Uxt | 1.40 | 1.8412694 82 | 0.01441220 788 | 0.0662556 54 |
| | Rgma | 1.40 | 1.9817937 07 | 0.01042812 654 | 0.0546556 89 |
| | Bnip2 | 1.40 | 2.7374437 59 | 0.00183044 313 | 0.0238288 09 |
| | 1110038B12 Rik | 1.40 | 1.6471310 8 | 0.02253558 935 | 0.0853062 07 |
| | Sdc2 | 1.40 | 2.1774658 42 | 0.00664559 940 | 0.0432441 79 |
| | Scd1 | 1.40 | 2.6401528 39 | 0.00229006 158 | 0.0261496 51 |
| | Sec11a | 1.40 | 2.3307160 43 | 0.00466964 599 | 0.0358266 92 |
| | Hmbs | 1.40 | 2.4936651 61 | 0.00320874 230 | 0.0299655 46 |
| | Uvssa | 1.40 | 1.6871843 27 | 0.02055018 205 | 0.0808185 61 |
| | Slc16a13 | 1.40 | 1.6840899 81 | 0.02069712 485 | 0.0811409 96 |
| | Syne2 | 1.40 | 1.7561801 03 | 0.01753153 314 | 0.0740777 36 |
| | Nebl | 1.40 | 1.4487928 74 | 0.03558009 686 | 0.1134616 18 |
| | Taf13 | 1.40 | 2.0981941 37 | 0.00797638 049 | 0.0480085 17 |

| | | | | | |
|---|---|---|---|---|---|
| | Rps9 | 1.40 | 1.595984951 | 0.02535216479 | 0.092190687 |
| | Pecr | 1.40 | 1.622941498 | 0.02382640404 | 0.088510035 |
| | Rap1b | 1.40 | 2.463372493 | 0.00344054709 | 0.030960418 |
| | Pigx | 1.40 | 1.508393005 | 0.03101751461 | 0.104436713 |
| | Cdk5rap3 | 1.40 | 2.76638478 | 0.00171243943 | 0.023271537 |
| | Adal | 1.40 | 2.707940132 | 0.00195911472 | 0.024453795 |
| | Rps27 | 1.40 | 2.394094462 | 0.00403557607 | 0.033410655 |
| | Rbpms | 1.40 | 1.415269555 | 0.03843531505 | 0.118843309 |
| | Mut | 1.39 | 1.753880106 | 0.01762462534 | 0.074242789 |
| | Zfp41 | 1.39 | 1.331120056 | 0.04665303955 | 0.135216076 |
| | Ssr3 | 1.39 | 3.306781547 | 0.00049342194 | 0.015871978 |
| | Glud1 | 1.39 | 3.349511702 | 0.00044718610 | 0.015871978 |
| | Tram1 | 1.39 | 1.813070177 | 0.01537906113 | 0.069073548 |
| | Oaf | 1.39 | 1.650696872 | 0.02235131750 | 0.084818960 |
| | Qk | 1.39 | 2.922967308 | 0.00119407799 | 0.020344995 |
| | Rpl22 | 1.39 | 2.716794013 | 0.00191957899 | 0.024310539 |
| | Hnrnpf | 1.39 | 1.992910239 | 0.01016458756 | 0.054073402 |
| | 4931406C07Rik | 1.39 | 2.29130346 | 0.00511324426 | 0.037512594 |
| | Zfp119a | 1.39 | 1.316446808 | 0.04825620798 | 0.138057163 |
| | Myl12a | 1.39 | 1.677549681 | 0.02101117400 | 0.081788822 |
| | Mpzl1 | 1.39 | 1.484274674 | 0.03278878518 | 0.107993935 |
| | Prkcq | 1.39 | 1.498575352 | 0.03172668144 | 0.105788966 |
| | Mettl23 | 1.39 | 2.48454405 | 0.00327684539 | 0.030148823 |

| | | | | | |
|---|---|---|---|---|---|
| | Gltp | 1.39 | 1.435373858 | 0.03669662650 | 0.115520812 |
| | Mtus1 | 1.39 | 1.753221615 | 0.01765136862 | 0.074332656 |
| | Vtn | 1.39 | 1.912945689 | 0.01221952462 | 0.059817773 |
| | Idua | 1.39 | 1.848006571 | 0.01419036051 | 0.065568971 |
| | Klf3 | 1.39 | 1.437103264 | 0.03655078728 | 0.115254026 |
| | Zfp105 | 1.39 | 1.784166965 | 0.01643739662 | 0.071780756 |
| | Cdk4 | 1.39 | 1.353040264 | 0.04435675179 | 0.130571150 |
| | Hdac8 | 1.39 | 2.486053131 | 0.00326547881 | 0.030114728 |
| | Psat1 | 1.39 | 2.732922942 | 0.00184959677 | 0.023952830 |
| | Ost4 | 1.38 | 2.233393005 | 0.00584261131 | 0.040584167 |
| | Snx5 | 1.38 | 3.903035302 | 0.00012501574 | 0.014121412 |
| | Farp1 | 1.38 | 1.728146598 | 0.01870050789 | 0.076659809 |
| | Slc25a18 | 1.38 | 2.185291283 | 0.00652692642 | 0.042797840 |
| | Zfp110 | 1.38 | 1.696302938 | 0.02012320083 | 0.079686715 |
| | Bhlhe41 | 1.38 | 2.427955455 | 0.00373288443 | 0.032301194 |
| | Tvp23b | 1.38 | 2.477077409 | 0.00333366988 | 0.030356500 |
| | Sec22a | 1.38 | 2.638594592 | 0.00229829307 | 0.026149651 |
| | Itm2b | 1.38 | 2.736646028 | 0.00183380846 | 0.023839510 |
| | Etfa | 1.38 | 2.717720157 | 0.00191548980 | 0.024310539 |
| | Smc2 | 1.38 | 1.816724524 | 0.01525019777 | 0.068796115 |
| | Ttc30a1 | 1.38 | 1.990308943 | 0.01022565313 | 0.054209375 |
| | Notch3 | 1.38 | 2.366627553 | 0.00429904952 | 0.034285107 |
| | Slc31a1 | 1.38 | 1.982252791 | 0.01041710900 | 0.054655689 |

| | | | | | |
|---|---|---|---|---|---|
| | Nhsl1 | 1.38 | 1.3097785<br>1 | 0.04900286<br>699 | 0.1393501<br>93 |
| | Mrps6 | 1.38 | 1.7917946<br>94 | 0.01615121<br>901 | 0.0711098<br>69 |
| | Aldh9a1 | 1.38 | 2.9424123<br>5 | 0.00114179<br>372 | 0.0201404<br>20 |
| | Katnbl1 | 1.38 | 2.4431055<br>6 | 0.00360491<br>011 | 0.0317532<br>50 |
| | Hmgn2 | 1.38 | 1.8233151<br>92 | 0.01502051<br>448 | 0.0681627<br>77 |
| | Plxdc2 | 1.38 | 2.0221613<br>33 | 0.00950251<br>727 | 0.0524795<br>10 |
| | Rpl10a | 1.38 | 2.1919102 | 0.00642820<br>621 | 0.0424791<br>52 |
| | Fbxo30 | 1.38 | 1.7154004<br>53 | 0.01925748<br>408 | 0.0777599<br>44 |
| | Xrcc2 | 1.37 | 1.5875125<br>57 | 0.02585160<br>092 | 0.0934198<br>52 |
| | Fam213a | 1.37 | 4.2166218<br>45 | 0.00006072<br>649 | 0.0141214<br>12 |
| | Smim11 | 1.37 | 1.4826141<br>61 | 0.03291439<br>213 | 0.1082778<br>70 |
| | Entpd5 | 1.37 | 2.6158793<br>6 | 0.00242170<br>166 | 0.0266313<br>44 |
| | Arl13b | 1.37 | 2.0232250<br>38 | 0.00947927<br>149 | 0.0524492<br>26 |
| | Metrn | 1.37 | 1.9964245<br>03 | 0.01008266<br>870 | 0.0538460<br>75 |
| | Appl2 | 1.37 | 2.7372520<br>81 | 0.00183125<br>119 | 0.0238288<br>09 |
| | Slc41a1 | 1.37 | 2.0198629<br>29 | 0.00955294<br>045 | 0.0525502<br>36 |
| | E2f5 | 1.37 | 1.5616281<br>67 | 0.02743922<br>451 | 0.0972009<br>24 |
| | Pik3c2a | 1.37 | 2.7260829<br>03 | 0.00187895<br>811 | 0.0240677<br>59 |
| | Atad5 | 1.37 | 1.4381572<br>94 | 0.03646218<br>636 | 0.1150992<br>20 |
| | Hsd17b10 | 1.37 | 2.8824112<br>77 | 0.00131095<br>783 | 0.0210688<br>56 |
| | Ssbp1 | 1.37 | 2.7138039<br>66 | 0.00193284<br>058 | 0.0243452<br>46 |
| | Dbndd2 | 1.37 | 2.0148564<br>8 | 0.00966370<br>180 | 0.0527858<br>35 |
| | Rpl5 | 1.37 | 2.8505716<br>73 | 0.00141067<br>941 | 0.0216582<br>63 |

| | | | | | |
|---|---|---|---|---|---|
| | Tmbim6 | 1.37 | 2.084769417 | 0.00822679324 | 0.048915779 |
| | Zfp882 | 1.37 | 1.32893046 | 0.04688884556 | 0.135647656 |
| | Cobll1 | 1.37 | 1.395066448 | 0.04026554221 | 0.122571736 |
| | Tpmt | 1.37 | 1.524769175 | 0.02986969754 | 0.101830650 |
| | Utp11 | 1.37 | 2.481323641 | 0.00330123438 | 0.030161078 |
| | Ncapg2 | 1.37 | 1.610008843 | 0.02454658936 | 0.090209865 |
| | Calu | 1.37 | 2.615715249 | 0.00242261694 | 0.026631344 |
| | Cat | 1.37 | 2.810932457 | 0.00154549478 | 0.022354362 |
| | Nampt | 1.37 | 1.617580721 | 0.02412233136 | 0.089127442 |
| | Ftl1 | 1.37 | 1.46220496 | 0.03449808916 | 0.111305199 |
| | Sypl | 1.37 | 2.125419296 | 0.00749170563 | 0.046429600 |
| | Tmco1 | 1.37 | 2.52821985 | 0.00296333090 | 0.029147552 |
| | Twf1 | 1.37 | 2.611686711 | 0.00244519382 | 0.026751121 |
| | Pcbd2 | 1.36 | 1.339562729 | 0.04575486418 | 0.133231106 |
| | Kdelc2 | 1.36 | 1.996795941 | 0.01007404900 | 0.053841292 |
| | Mcl1 | 1.36 | 2.036516754 | 0.00919355008 | 0.051647004 |
| | Ptgr2 | 1.36 | 2.962322974 | 0.00109062896 | 0.019947158 |
| | Bckdha | 1.36 | 2.961853588 | 0.00109180835 | 0.019947158 |
| | Amdhd2 | 1.36 | 1.540326202 | 0.02881866097 | 0.099697185 |
| | Lrp4 | 1.36 | 1.728324509 | 0.01869284870 | 0.076659809 |
| | Rpl36a | 1.36 | 2.116432576 | 0.00764834419 | 0.047130936 |
| | Blvrb | 1.36 | 1.304905708 | 0.04955577722 | 0.140285524 |
| | Rps3 | 1.36 | 2.770722308 | 0.00169542152 | 0.023241590 |

| | | | | | |
|---|---|---|---|---|---|
| | Rnf138 | 1.36 | 2.543250687 | 0.00286252516 | 0.028725861 |
| | Gpr19 | 1.36 | 2.557254646 | 0.00277169446 | 0.028393015 |
| | Cep44 | 1.36 | 1.791010007 | 0.01618042754 | 0.071192845 |
| | Rpl34-ps1 | 1.36 | 1.734368873 | 0.01843449000 | 0.076150386 |
| | Gmds | 1.36 | 1.470093496 | 0.03387712166 | 0.110100645 |
| | Rps27a | 1.36 | 2.518587158 | 0.00302979219 | 0.029392377 |
| | Phtf1os | 1.36 | 1.442449707 | 0.03610358202 | 0.114440443 |
| | Cln5 | 1.36 | 1.462328838 | 0.03448825028 | 0.111305199 |
| | Tpt1 | 1.36 | 1.999903471 | 0.01000222292 | 0.053745583 |
| | 1810026B05Rik | 1.36 | 2.569801807 | 0.00269276339 | 0.027883392 |
| | Fmnl3 | 1.36 | 1.375970417 | 0.04207552878 | 0.125992885 |
| | Mpp5 | 1.36 | 3.359308073 | 0.00043721185 | 0.015871978 |
| | Eef1a1 | 1.36 | 1.389047538 | 0.04082746943 | 0.123925394 |
| | Anp32b | 1.36 | 1.749378149 | 0.01780827489 | 0.074754258 |
| | Eml3 | 1.36 | 1.492031454 | 0.03220835509 | 0.106747469 |
| | Pex2 | 1.36 | 2.431502436 | 0.00370252129 | 0.032184714 |
| | Lgals8 | 1.36 | 2.297296195 | 0.00504317228 | 0.037264945 |
| | Mt3 | 1.36 | 2.096701245 | 0.00800384656 | 0.048026575 |
| | Pabpc1 | 1.36 | 1.355637408 | 0.04409228364 | 0.129959689 |
| | Ugp2 | 1.36 | 3.116283124 | 0.00076509766 | 0.017551264 |
| | Rpl7 | 1.36 | 2.376107497 | 0.00420622503 | 0.033998669 |
| | 4930503L19Rik | 1.36 | 1.409750446 | 0.03892687624 | 0.119716698 |
| | Usp53 | 1.36 | 1.585789112 | 0.02595439370 | 0.093606122 |

| | | | | | |
|---|---|---|---|---|---|
| | Spcs2 | 1.36 | 1.9054264<br>07 | 0.01243293<br>301 | 0.0604832<br>25 |
| | Serpinb6a | 1.36 | 3.7007064<br>41 | 0.00019920<br>194 | 0.0141214<br>12 |
| | Cpne2 | 1.36 | 1.3043460<br>81 | 0.04961967<br>543 | 0.1403796<br>50 |
| | Sec61g | 1.36 | 2.0472176<br>18 | 0.00896979<br>221 | 0.0509734<br>97 |
| | Rps19 | 1.35 | 1.8390410<br>19 | 0.01448635<br>022 | 0.0664852<br>83 |
| | Npm1 | 1.35 | 2.1410755<br>86 | 0.00722644<br>022 | 0.0454284<br>20 |
| | Ybx1 | 1.35 | 1.5607423<br>59 | 0.02749524<br>801 | 0.0973491<br>89 |
| | Snrpg | 1.35 | 2.1464031<br>69 | 0.00713833<br>344 | 0.0451186<br>02 |
| | Sccpdh | 1.35 | 1.9589836<br>07 | 0.01099047<br>324 | 0.0564231<br>81 |
| | Pter | 1.35 | 1.7609691<br>37 | 0.01733927<br>214 | 0.0736503<br>67 |
| | Gstm1 | 1.35 | 2.1681878<br>27 | 0.00678909<br>949 | 0.0437932<br>78 |
| | Tpp1 | 1.35 | 2.1451615<br>87 | 0.00715877<br>006 | 0.0452269<br>70 |
| | Ppp1r2 | 1.35 | 1.8040445<br>66 | 0.01570201<br>667 | 0.0700840<br>05 |
| | Ryk | 1.35 | 2.6639995<br>3 | 0.00216770<br>645 | 0.0255190<br>26 |
| | Pik3ip1 | 1.35 | 1.6896097<br>8 | 0.02043573<br>303 | 0.0804605<br>75 |
| | Gcsh | 1.35 | 2.3605811<br>61 | 0.00435932<br>090 | 0.0346436<br>43 |
| | Tmpo | 1.35 | 1.5878889<br>92 | 0.02582920<br>312 | 0.0934198<br>52 |
| | Ormdl1 | 1.35 | 1.4921586<br>05 | 0.03219892<br>668 | 0.1067474<br>69 |
| | Trim26 | 1.35 | 1.5310632<br>04 | 0.02943993<br>156 | 0.1009489<br>14 |
| | Cmtr2 | 1.35 | 1.3088338<br>96 | 0.04910956<br>687 | 0.1394639<br>75 |
| | Mageh1 | 1.35 | 2.3735964<br>45 | 0.00423061<br>549 | 0.0340755<br>49 |
| | 1810009A15<br>Rik | 1.35 | 1.5220691<br>89 | 0.03005597<br>435 | 0.1023034<br>79 |
| | Dag1 | 1.35 | 2.1653306<br>14 | 0.00683391<br>206 | 0.0439012<br>56 |

| | | | | | |
|---|---|---|---|---|---|
| | S1pr1 | 1.35 | 3.114518616 | 0.00076821252 | 0.017593347 |
| | Rela | 1.35 | 1.493580106 | 0.03209370780 | 0.106495928 |
| | Pts | 1.35 | 1.786243113 | 0.01635900507 | 0.071626989 |
| | Plod1 | 1.34 | 1.737882126 | 0.01828596457 | 0.075819988 |
| | Ktn1 | 1.34 | 3.590252891 | 0.00025688995 | 0.014200135 |
| | Tab2 | 1.34 | 1.922274301 | 0.01195984907 | 0.059200391 |
| | Ephx2 | 1.34 | 2.38443342 | 0.00412635491 | 0.033646417 |
| | Naf1 | 1.34 | 2.08874655 | 0.00815179876 | 0.048574964 |
| | Ankrd13a | 1.34 | 1.800534641 | 0.01582943299 | 0.070291175 |
| | Bphl | 1.34 | 2.227962353 | 0.00591612917 | 0.040769073 |
| | Pdia3 | 1.34 | 2.263339476 | 0.00545331424 | 0.039027548 |
| | Alcam | 1.34 | 2.146748479 | 0.00713265997 | 0.045118602 |
| | Mapk12 | 1.34 | 1.322080102 | 0.04763431210 | 0.136962352 |
| | Haus5 | 1.34 | 1.346982799 | 0.04497976699 | 0.131664294 |
| | Dnajc17 | 1.34 | 1.581439077 | 0.02621566768 | 0.094226913 |
| | Nxt1 | 1.34 | 1.480165348 | 0.03310050746 | 0.108707952 |
| | Cnksr3 | 1.34 | 1.47725208 | 0.03332329367 | 0.109126639 |
| | Fgd4 | 1.34 | 1.997418151 | 0.01005962634 | 0.053827531 |
| | Trmt2b | 1.34 | 1.630140732 | 0.02343469299 | 0.087623433 |
| | Rpl23 | 1.34 | 2.037817765 | 0.00916605028 | 0.051569310 |
| | Nsa2 | 1.34 | 2.189532542 | 0.00646349561 | 0.042597071 |
| | Zmpste24 | 1.34 | 2.493558586 | 0.00320952981 | 0.029965546 |
| | Pdgfrb | 1.34 | 2.668062304 | 0.00214752237 | 0.025460833 |

| | | | | | |
|---|---|---|---|---|---|
| | H3f3b | 1.34 | 1.927357334 | 0.01182068559 | 0.058855750 |
| | Suco | 1.34 | 2.632281791 | 0.00233194449 | 0.026229430 |
| | Rhoa | 1.34 | 2.845978742 | 0.00142567738 | 0.021677178 |
| | Kcnj10 | 1.34 | 1.916234453 | 0.01212733982 | 0.059603303 |
| | Jam2 | 1.34 | 1.860879284 | 0.01377592330 | 0.064211317 |
| | 2610001J05Rik | 1.34 | 2.072505193 | 0.00846242451 | 0.049351543 |
| | Desi2 | 1.34 | 2.484899776 | 0.00327416245 | 0.030148823 |
| | Slc6a9 | 1.34 | 2.352531982 | 0.00444086957 | 0.034726255 |
| | Fra10ac1 | 1.34 | 1.847830585 | 0.01419611194 | 0.065568971 |
| | Me1 | 1.34 | 2.48276625 | 0.00329028676 | 0.030148823 |
| | Pofut1 | 1.34 | 1.508262924 | 0.03102680641 | 0.104436713 |
| | Ltbp3 | 1.34 | 1.559101192 | 0.02759934707 | 0.097510287 |
| | Prps1l3 | 1.34 | 1.53429525 | 0.02922165105 | 0.100610049 |
| | Cep41 | 1.34 | 1.416225013 | 0.03835084942 | 0.118654387 |
| | Dpy19l4 | 1.34 | 1.680499348 | 0.02086895255 | 0.081558668 |
| | Nab1 | 1.34 | 2.263221292 | 0.00545479844 | 0.039027548 |
| | Flna | 1.34 | 1.451465624 | 0.03536180101 | 0.112975240 |
| | Gm14325 | 1.34 | 2.46361684 | 0.00343861189 | 0.030960418 |
| | Fbxo8 | 1.34 | 2.854662783 | 0.00139745302 | 0.021544296 |
| | Hadh | 1.34 | 2.228169686 | 0.00591330546 | 0.040769073 |
| | Wscd1 | 1.34 | 1.346625816 | 0.04501675477 | 0.131723856 |
| | Bet1 | 1.33 | 2.270434182 | 0.00536495171 | 0.038759096 |
| | Slc35b2 | 1.33 | 2.742429235 | 0.00180955074 | 0.023748841 |

| | | | | | |
|---|---|---|---|---|---|
| | Srbd1 | 1.33 | 1.5706920<br>28 | 0.02687249<br>383 | 0.0959103<br>73 |
| | Rps3a1 | 1.33 | 2.9144917<br>96 | 0.00121760<br>999 | 0.0204287<br>90 |
| | Nudcd2 | 1.33 | 2.6922075<br>19 | 0.00203138<br>612 | 0.0249225<br>68 |
| | Chst2 | 1.33 | 1.9338218<br>56 | 0.01164603<br>642 | 0.0583356<br>81 |
| | Scd2 | 1.33 | 2.1327180<br>7 | 0.00736685<br>175 | 0.0459708<br>95 |
| | Arxes2 | 1.33 | 3.0913348<br>86 | 0.00081033<br>596 | 0.0179439<br>01 |
| | Arhgap29 | 1.33 | 1.4171627<br>08 | 0.03826813<br>451 | 0.1185665<br>02 |
| | Rhobtb3 | 1.33 | 1.7822607<br>39 | 0.01650970<br>303 | 0.0719520<br>69 |
| | Adcyap1r1 | 1.33 | 1.8217519<br>75 | 0.01507467<br>734 | 0.0682927<br>59 |
| | Snrpe | 1.33 | 2.8147000<br>11 | 0.00153214<br>543 | 0.0222792<br>97 |
| | Ebpl | 1.33 | 1.8983821<br>15 | 0.01263624<br>056 | 0.0610529<br>47 |
| | Asrgl1 | 1.33 | 2.4317102<br>54 | 0.00370074<br>998 | 0.0321847<br>14 |
| | P2rx4 | 1.33 | 1.9136751<br>53 | 0.01219901<br>729 | 0.0597812<br>76 |
| | Slc38a1 | 1.33 | 1.9015722<br>92 | 0.01254375<br>918 | 0.0607770<br>79 |
| | Pex12 | 1.33 | 1.6102599<br>62 | 0.02453240<br>004 | 0.0902059<br>70 |
| | Wdr92 | 1.33 | 1.4475187<br>81 | 0.03568463<br>170 | 0.1136367<br>38 |
| | Snx6 | 1.33 | 1.9292132<br>76 | 0.01177027<br>809 | 0.0586635<br>44 |
| | Slc4a4 | 1.33 | 1.9689948<br>73 | 0.01074002<br>090 | 0.0556270<br>74 |
| | Prkx | 1.33 | 1.6824594<br>33 | 0.02077497<br>771 | 0.0813070<br>26 |
| | Gpc6 | 1.33 | 2.5269964<br>91 | 0.00297169<br>004 | 0.0291622<br>45 |
| | Fmnl2 | 1.33 | 2.2948135<br>98 | 0.00507208<br>358 | 0.0373891<br>30 |
| | Dph6 | 1.33 | 1.3900656<br>41 | 0.04073187<br>095 | 0.1237445<br>59 |
| | Acat3 | 1.33 | 1.9602972<br>87 | 0.01095727<br>881 | 0.0563487<br>90 |

| | | | | | |
|---|---|---|---|---|---|
| | Wbp1 | 1.33 | 1.720274612 | 0.01904256241 | 0.077372960 |
| | Dazap2 | 1.32 | 2.823448842 | 0.00150158928 | 0.022050044 |
| | Tmem168 | 1.32 | 1.763566689 | 0.01723587403 | 0.073534751 |
| | Tor1aip2 | 1.32 | 2.344037159 | 0.00452858831 | 0.035196455 |
| | Rpl27 | 1.32 | 2.135331942 | 0.00732264631 | 0.045840767 |
| | Sgce | 1.32 | 1.842225109 | 0.01438052996 | 0.066176444 |
| | Fcho2 | 1.32 | 2.544160021 | 0.00285653782 | 0.028725861 |
| | Zfp120 | 1.32 | 2.293172935 | 0.00509128097 | 0.037451441 |
| | Med7 | 1.32 | 2.392789352 | 0.00404772173 | 0.033464335 |
| | Rbl2 | 1.32 | 2.314423897 | 0.00484815061 | 0.036412040 |
| | Gabrg1 | 1.32 | 1.781865031 | 0.01652475271 | 0.071976233 |
| | Acadvl | 1.32 | 2.90195428 | 0.00125327311 | 0.020677756 |
| | Selenos | 1.32 | 3.632848322 | 0.00023289045 | 0.014200135 |
| | 2700060E02Rik | 1.32 | 2.366634048 | 0.00429898522 | 0.034285107 |
| | Rfx3 | 1.32 | 1.378776601 | 0.04180453520 | 0.125532802 |
| | Myl9 | 1.32 | 1.672891828 | 0.02123773377 | 0.082251325 |
| | Ocln | 1.32 | 1.631992873 | 0.02333496357 | 0.087440887 |
| | Aldh6a1 | 1.32 | 3.016213476 | 0.00096335537 | 0.018992061 |
| | Jun | 1.32 | 1.39590688 | 0.04018769706 | 0.122416126 |
| | Sparcl1 | 1.32 | 1.888293908 | 0.01293320293 | 0.061948638 |
| | Fdx1 | 1.32 | 1.520844887 | 0.03014082341 | 0.102541484 |
| | Snx18 | 1.32 | 1.317071691 | 0.04818682473 | 0.137944825 |
| | Cenpx | 1.32 | 2.134926837 | 0.00732947998 | 0.045862652 |

| | | | | | |
|---|---|---|---|---|---|
| | Tcf12 | 1.32 | 1.416243482 | 0.03834921848 | 0.118654387 |
| | Psd2 | 1.32 | 2.067438883 | 0.00856172190 | 0.049556285 |
| | Trnau1ap | 1.32 | 2.404630312 | 0.00393885223 | 0.033006104 |
| | Fuca2 | 1.32 | 2.957373749 | 0.00110312887 | 0.019984262 |
| | Dek | 1.32 | 2.574016075 | 0.00266675995 | 0.027727676 |
| | Vmac | 1.32 | 1.390182617 | 0.04072090139 | 0.123738590 |
| | 2010315B03 Rik | 1.32 | 2.274889377 | 0.00531019687 | 0.038484924 |
| | Sox6 | 1.32 | 1.508185765 | 0.03103231926 | 0.104436713 |
| | Cyfip1 | 1.32 | 2.418724566 | 0.00381307576 | 0.032543773 |
| | Sumf2 | 1.32 | 1.717452856 | 0.01916669112 | 0.077560752 |
| | Asf1a | 1.32 | 1.512721584 | 0.03070990098 | 0.103911536 |
| | Zmat1 | 1.31 | 1.652046163 | 0.02228198290 | 0.084696190 |
| | Ctsa | 1.31 | 1.824416805 | 0.01498246236 | 0.068079032 |
| | Ctsl | 1.31 | 1.985656644 | 0.01033578236 | 0.054548381 |
| | Idi1 | 1.31 | 2.505776813 | 0.00312049281 | 0.029668607 |
| | Tmed10 | 1.31 | 2.756582927 | 0.00175152796 | 0.023457842 |
| | 1810022K09 Rik | 1.31 | 2.684744425 | 0.00206659596 | 0.025063632 |
| | Hbp1 | 1.31 | 2.060496709 | 0.00869968026 | 0.050059592 |
| | Plod3 | 1.31 | 1.403202435 | 0.03951823733 | 0.120966830 |
| | Golph3l | 1.31 | 1.558279044 | 0.02765164397 | 0.097651308 |
| | Gkap1 | 1.31 | 2.071860702 | 0.00847499202 | 0.049351543 |
| | Mid1ip1 | 1.31 | 2.469451789 | 0.00339272150 | 0.030751574 |
| | Hadhb | 1.31 | 2.19364761 | 0.00640254133 | 0.042378285 |

| | | | | | |
|---|---|---|---|---|---|
| | Zfp944 | 1.31 | 1.905344423 | 0.01243528023 | 0.060483225 |
| | Fam204a | 1.31 | 2.394871068 | 0.00402836609 | 0.033406022 |
| | Elf2 | 1.31 | 1.471248894 | 0.03378711466 | 0.109902050 |
| | Zbtb41 | 1.31 | 2.46468653 | 0.00343015283 | 0.030940003 |
| | Arl6ip6 | 1.31 | 1.567212417 | 0.02708866377 | 0.096406457 |
| | Mtfr1 | 1.31 | 2.048580744 | 0.00894168273 | 0.050940159 |
| | Gtf2e1 | 1.31 | 1.590175912 | 0.02569354848 | 0.093129794 |
| | Rwdd3 | 1.31 | 1.35448891 | 0.04420904059 | 0.130247947 |
| | Enpp4 | 1.31 | 1.58026847 | 0.02628642530 | 0.094407154 |
| | Mccc2 | 1.31 | 1.351232319 | 0.04454179152 | 0.130723784 |
| | Ostc | 1.31 | 2.081549581 | 0.00828801292 | 0.049025220 |
| | Gabpa | 1.31 | 1.388190647 | 0.04090810413 | 0.123997555 |
| | Smad1 | 1.31 | 2.024240961 | 0.00945712303 | 0.052408546 |
| | Zfp943 | 1.31 | 2.521258353 | 0.00301121418 | 0.029347135 |
| | Uimc1 | 1.31 | 1.405203284 | 0.03933659064 | 0.120520647 |
| | Cetn2 | 1.31 | 1.699697774 | 0.01996651303 | 0.079409510 |
| | Rps5 | 1.31 | 1.648236597 | 0.02247829689 | 0.085206697 |
| | Rassf2 | 1.31 | 1.862293628 | 0.01373113293 | 0.064089503 |
| | Mettl5 | 1.31 | 1.558031248 | 0.02766742566 | 0.097652405 |
| | N4bp2l2 | 1.31 | 1.756235329 | 0.01752930391 | 0.074077736 |
| | Cryz | 1.31 | 1.70991246 | 0.01950237667 | 0.078382862 |
| | Rpl15 | 1.31 | 3.052428764 | 0.00088628058 | 0.018531284 |
| | Zfp606 | 1.31 | 1.803466908 | 0.01572291591 | 0.070084005 |

| | | | | | |
|---|---|---|---|---|---|
| | Rps6 | 1.31 | 2.405716464 | 0.00392901364 | 0.033001988 |
| | St8sia4 | 1.31 | 1.662517673 | 0.02175115517 | 0.083376207 |
| | Cetn3 | 1.31 | 3.772077764 | 0.00016901383 | 0.014121412 |
| | Dnajc1 | 1.30 | 1.389340039 | 0.04079998102 | 0.123896694 |
| | Hus1 | 1.30 | 1.829563715 | 0.01480595021 | 0.067545991 |
| | Pigp | 1.30 | 1.911541119 | 0.01225910828 | 0.059966879 |
| | Slc25a17 | 1.30 | 1.518818083 | 0.03028181603 | 0.102880883 |
| | Rps7 | 1.30 | 2.481919615 | 0.00329670726 | 0.030148823 |
| | Selenof | 1.30 | 2.126332788 | 0.00747596419 | 0.046398927 |
| | Cox16 | 1.30 | 2.234499478 | 0.00582774475 | 0.040525830 |
| | Eef1b2 | 1.30 | 2.110201008 | 0.00775887924 | 0.047598730 |
| | Rpn2 | 1.30 | 1.708808659 | 0.01955200686 | 0.078450131 |
| | Slc3a2 | 1.30 | 1.883013547 | 0.01309141086 | 0.062404912 |
| | Kdelr2 | 1.30 | 1.68609802 | 0.02060164884 | 0.080928351 |
| | Pak2 | 1.30 | 1.986099409 | 0.01032525036 | 0.054527004 |
| | Gabarap | 1.30 | 2.251151838 | 0.00560851857 | 0.039626664 |
| | 1810030O07Rik | 1.30 | 1.480617657 | 0.03306605188 | 0.108620755 |
| | Fam162a | 1.30 | 2.678344495 | 0.00209727560 | 0.025169139 |
| | Sf3b6 | 1.30 | 3.192489749 | 0.00064196337 | 0.016529429 |
| | Hadha | 1.30 | 2.062621174 | 0.00865722742 | 0.049898893 |
| | Coa6 | 1.30 | 1.317924134 | 0.04809233523 | 0.137789153 |
| | Rap1a | 1.30 | 1.757349467 | 0.01748439193 | 0.073992310 |
| | Usp1 | 1.30 | 2.390044174 | 0.00407338843 | 0.033596897 |

| | | | | | |
|---|---|---|---|---|---|
| | Socs6 | 1.30 | 1.482990648 | 0.03288587122 | 0.108209951 |
| | Rps14 | 1.30 | 2.245322275 | 0.00568430961 | 0.039912161 |
| | Commd1 | 1.30 | 2.336143335 | 0.00461165346 | 0.035520912 |
| | Rps15a | 1.30 | 2.001356738 | 0.00996880870 | 0.053713983 |
| | P4hb | 1.30 | 1.754779003 | 0.01758818388 | 0.074134735 |
| | Eif3e | 1.30 | 2.718304021 | 0.00191291635 | 0.024310539 |
| | Cc2d2a | 1.30 | 2.635148935 | 0.00231660007 | 0.026221089 |
| | Pnpla2 | 1.30 | 1.425848671 | 0.03751036839 | 0.117223100 |
| | Dars | 1.30 | 2.423830434 | 0.00376850908 | 0.032351180 |
| | Abcd4 | 1.30 | 2.069783774 | 0.00851561908 | 0.049442186 |
| | Fads1 | 1.30 | 2.367626608 | 0.00428917131 | 0.034285107 |
| | Clcc1 | 1.30 | 2.610470888 | 0.00245204882 | 0.026781419 |
| | Crim1 | 1.30 | 1.502469503 | 0.03144347213 | 0.105433077 |
| | Aldh1l1 | 1.29 | 2.727066951 | 0.00187470548 | 0.024052594 |
| | Dscr3 | 1.29 | 2.907600675 | 0.00123708438 | 0.020529923 |
| | Rpf1 | 1.29 | 1.470945306 | 0.03381074138 | 0.109936914 |
| | Mapkapk2 | 1.29 | 1.562359965 | 0.02739302760 | 0.097087334 |
| | Fchsd2 | 1.29 | 1.418869657 | 0.03811802080 | 0.118341555 |
| | Elp4 | 1.29 | 1.306262327 | 0.04940121984 | 0.139979586 |
| | Gpm6b | 1.29 | 1.926879114 | 0.01183370903 | 0.058860516 |
| | Gyg | 1.29 | 1.948421895 | 0.01126102972 | 0.057183226 |
| | Stag2 | 1.29 | 2.613997775 | 0.00243221647 | 0.026651584 |
| | Fam114a2 | 1.29 | 2.20429383 | 0.00624749863 | 0.041814224 |

| | | | | | |
|---|---|---|---|---|---|
| | Slc6a11 | 1.29 | 1.731222787 | 0.01856851670 | 0.076392212 |
| | Scfd1 | 1.29 | 2.268581268 | 0.00538789015 | 0.038843126 |
| | Lsm3 | 1.29 | 1.979558606 | 0.01048193336 | 0.054806791 |
| | Dram2 | 1.29 | 1.623119323 | 0.02381665011 | 0.088497726 |
| | Msmo1 | 1.29 | 1.720928609 | 0.01901390813 | 0.077344616 |
| | Parvb | 1.29 | 1.991171992 | 0.01020535244 | 0.054164445 |
| | Chmp5 | 1.29 | 2.843131907 | 0.00143505350 | 0.021677178 |
| | Mmgt2 | 1.29 | 1.413791203 | 0.03856637295 | 0.119061004 |
| | Mcur1 | 1.29 | 1.500342644 | 0.03159783709 | 0.105676798 |
| | Fam76b | 1.28 | 1.67768316 | 0.02100471726 | 0.081788822 |
| | Alg5 | 1.28 | 2.263060068 | 0.00545682382 | 0.039027548 |
| | Rpl18a | 1.28 | 1.451721556 | 0.03534096827 | 0.112975240 |
| | St3gal4 | 1.28 | 2.254764844 | 0.00556205342 | 0.039464488 |
| | Rpe | 1.28 | 1.659652149 | 0.02189514627 | 0.083735376 |
| | Rps23 | 1.28 | 2.041805648 | 0.00908226882 | 0.051357801 |
| | Abcb4 | 1.28 | 1.511955625 | 0.03076411141 | 0.104025135 |
| | Syf2 | 1.28 | 2.799351439 | 0.00158726179 | 0.022676450 |
| | Mettl21a | 1.28 | 1.690010465 | 0.02041688748 | 0.080410454 |
| | B230118H07Rik | 1.28 | 2.03298059 | 0.00926871248 | 0.051899502 |
| | Itgav | 1.28 | 1.447194169 | 0.03571131406 | 0.113695358 |
| | Rps12 | 1.28 | 1.421697407 | 0.03787063553 | 0.117866456 |
| | Fgfr2 | 1.28 | 1.670278774 | 0.02136590170 | 0.082468256 |
| | Rps17 | 1.28 | 2.738398816 | 0.00182642223 | 0.023828809 |

| | | | | | |
|---|---|---|---|---|---|
| | Megf10 | 1.28 | 1.661423642 | 0.02180601761 | 0.083534008 |
| | H2afv | 1.28 | 1.657212261 | 0.02201850046 | 0.084014079 |
| | A130010J15Rik | 1.28 | 1.559818386 | 0.02755380714 | 0.097430497 |
| | Gorab | 1.28 | 1.6544105 | 0.02216100745 | 0.084399779 |
| | Ppie | 1.28 | 1.397959653 | 0.03999819069 | 0.121892912 |
| | Wls | 1.28 | 1.960851531 | 0.01094330413 | 0.056340181 |
| | Gtf3c6 | 1.28 | 1.699060847 | 0.01999581697 | 0.079434091 |
| | Rnft1 | 1.28 | 1.893410446 | 0.01278172748 | 0.061431870 |
| | Zfp277 | 1.28 | 2.923013293 | 0.00119395156 | 0.020344995 |
| | Commd6 | 1.28 | 1.761078948 | 0.01733488846 | 0.073650367 |
| | Rab33b | 1.28 | 1.947785433 | 0.01127754496 | 0.057214932 |
| | Mrpl24 | 1.28 | 2.730948471 | 0.00185802490 | 0.023995414 |
| | Slc25a1 | 1.28 | 1.447550392 | 0.03568203447 | 0.113636738 |
| | Zfp958 | 1.28 | 1.333776406 | 0.04636855844 | 0.134546086 |
| | Epb41l2 | 1.28 | 2.045610922 | 0.00900303789 | 0.051099027 |
| | Phpt1 | 1.28 | 1.783217849 | 0.01647335855 | 0.071883271 |
| | Tmem38b | 1.28 | 1.950055712 | 0.01121874529 | 0.057036957 |
| | Rnf139 | 1.28 | 2.836830928 | 0.00145602581 | 0.021867902 |
| | Slc25a20 | 1.28 | 1.667031658 | 0.02152624812 | 0.082831749 |
| | Rpl17 | 1.28 | 2.149872524 | 0.00708153614 | 0.044945676 |
| | Mpp6 | 1.28 | 2.71012106 | 0.00194930115 | 0.024439185 |
| | Rhoq | 1.28 | 1.684256507 | 0.02068919026 | 0.081133037 |
| | Smim15 | 1.28 | 2.384608641 | 0.00412469043 | 0.033646417 |

| | | | | | |
|---|---|---|---|---|---|
| | Tmem164 | 1.28 | 1.438800049 | 0.03640826223 | 0.115024120 |
| | Abhd5 | 1.27 | 2.038137077 | 0.00915931348 | 0.051569310 |
| | Mesd | 1.27 | 2.541658446 | 0.00287303921 | 0.028753410 |
| | Ppp4r3b | 1.27 | 2.737257784 | 0.00183122714 | 0.023828809 |
| | Ap1s2 | 1.27 | 1.342496334 | 0.04544683733 | 0.132680639 |
| | Arl6 | 1.27 | 2.089804897 | 0.00813195756 | 0.048541907 |
| | Smurf2 | 1.27 | 1.735427848 | 0.01838959445 | 0.076088954 |
| | Slc7a10 | 1.27 | 1.864867009 | 0.01365001067 | 0.063857760 |
| | Hmgcl | 1.27 | 1.945620186 | 0.01133391141 | 0.057383669 |
| | Usp40 | 1.27 | 1.306226852 | 0.04940525529 | 0.139979586 |
| | Tmbim1 | 1.27 | 2.17135768 | 0.00673972723 | 0.043684241 |
| | Zadh2 | 1.27 | 1.674641665 | 0.02115233591 | 0.082128920 |
| | Nek9 | 1.27 | 2.288004055 | 0.00515223834 | 0.037738223 |
| | Ddx21 | 1.27 | 1.375786871 | 0.04209331494 | 0.125992885 |
| | Sp1 | 1.27 | 1.504164182 | 0.03132101429 | 0.105125075 |
| | 1110032A03Rik | 1.27 | 2.086003256 | 0.00820345393 | 0.048798121 |
| | Mrpl33 | 1.27 | 1.821628103 | 0.01507897765 | 0.068292759 |
| | Ece2 | 1.27 | 1.319922558 | 0.04787154474 | 0.137318801 |
| | Acsl3 | 1.27 | 1.634511855 | 0.02320000855 | 0.087053882 |
| | Adipor2 | 1.27 | 2.8429742 | 0.00143557471 | 0.021677178 |
| | Rft1 | 1.27 | 1.505560462 | 0.03122047726 | 0.104890117 |
| | Lman1 | 1.27 | 1.78225108 | 0.01651007019 | 0.071952069 |
| | Wdr43 | 1.27 | 1.742177535 | 0.01810599789 | 0.075534362 |

| | | | | | |
|---|---|---|---|---|---|
| | Vezf1 | 1.27 | 1.931803513 | 0.01170028624 | 0.058426042 |
| | Pigc | 1.27 | 1.705939307 | 0.01968161323 | 0.078715332 |
| | Abracl | 1.27 | 1.517463535 | 0.03037641134 | 0.103087742 |
| | Nxt2 | 1.27 | 2.517466685 | 0.00303761910 | 0.029392377 |
| | Taf9b | 1.27 | 1.729973571 | 0.01862200456 | 0.076520623 |
| | Rpl31 | 1.27 | 2.632200879 | 0.00233237899 | 0.026229430 |
| | Snapin | 1.27 | 2.738030777 | 0.00182797067 | 0.023828809 |
| | Gnpnat1 | 1.27 | 1.880359972 | 0.01317164534 | 0.062564375 |
| | Mtdh | 1.27 | 2.969551147 | 0.00107262732 | 0.019783855 |
| | Ccdc191 | 1.27 | 1.341466435 | 0.04555473919 | 0.132817244 |
| | Smim20 | 1.27 | 1.857658423 | 0.01387846956 | 0.064492408 |
| | Fundc1 | 1.27 | 2.520111624 | 0.00301917562 | 0.029347135 |
| | Gt(ROSA)26Sor | 1.27 | 1.698277515 | 0.02003191577 | 0.079501646 |
| | Atp1b3 | 1.27 | 3.00031371 | 0.00099927792 | 0.019232744 |
| | Tgs1 | 1.26 | 1.862502667 | 0.01372452533 | 0.064080429 |
| | Rpl7a | 1.26 | 1.773438883 | 0.01684849515 | 0.072853387 |
| | B3glct | 1.26 | 1.947122956 | 0.01129476096 | 0.057227622 |
| | Stag1 | 1.26 | 1.488763623 | 0.03245161961 | 0.107243315 |
| | Derl1 | 1.26 | 2.354094606 | 0.00442491971 | 0.034726255 |
| | Dhrs7 | 1.26 | 1.431527825 | 0.03702304845 | 0.116175773 |
| | Sar1b | 1.26 | 2.479882944 | 0.00331220383 | 0.030221111 |
| | Sp3 | 1.26 | 2.26345751 | 0.00545183232 | 0.039027548 |
| | Tmed7 | 1.26 | 2.314510747 | 0.00484718117 | 0.036412040 |

| | | | | | |
|---|---|---|---|---|---|
| | Atr | 1.26 | 2.027379912 | 0.00938901618 | 0.052190320 |
| | Vcl | 1.26 | 2.499871008 | 0.00316321705 | 0.029738034 |
| | Zfp386 | 1.26 | 1.974611852 | 0.01060200851 | 0.055245430 |
| | Zbtb8os | 1.26 | 1.94399653 | 0.01137636376 | 0.057429322 |
| | 2010320M18Rik | 1.26 | 1.31038306 | 0.04893470105 | 0.139273349 |
| | Tmed2 | 1.26 | 3.294374221 | 0.00050772176 | 0.015871978 |
| | Hsdl2 | 1.26 | 1.370705045 | 0.04258875601 | 0.127164732 |
| | Golga7 | 1.26 | 2.390912569 | 0.00406525161 | 0.033549923 |
| | Ptp4a2 | 1.26 | 2.499119723 | 0.00316869382 | 0.029738034 |
| | Stk3 | 1.26 | 1.511110674 | 0.03082402339 | 0.104113940 |
| | Suox | 1.26 | 1.474180405 | 0.03355981788 | 0.109404072 |
| | Garem2 | 1.26 | 1.603701327 | 0.02490569543 | 0.091079758 |
| | Wsb1 | 1.26 | 1.676761789 | 0.02104932680 | 0.081881837 |
| | Rdx | 1.26 | 2.222428158 | 0.00599200052 | 0.040984885 |
| | Aco1 | 1.26 | 2.139072472 | 0.00725984800 | 0.045563515 |
| | Snx10 | 1.26 | 1.906986637 | 0.01238834705 | 0.060350637 |
| | Syap1 | 1.26 | 2.306307493 | 0.00493960824 | 0.036864758 |
| | Rpl12 | 1.26 | 2.776356027 | 0.00167357035 | 0.023158641 |
| | Gramd3 | 1.26 | 1.49955747 | 0.03165501549 | 0.105748014 |
| | Ak3 | 1.26 | 2.909527083 | 0.00123160918 | 0.020485825 |
| | Actr6 | 1.26 | 1.629792211 | 0.02345350683 | 0.087669923 |
| | Hsd17b4 | 1.26 | 2.056069582 | 0.00878881693 | 0.050277741 |
| | 9530068E07Rik | 1.26 | 1.965596181 | 0.01082439967 | 0.055979705 |

| | | | | | |
|---|---|---|---|---|---|
| | Mettl9 | 1.26 | 2.381467546 | 0.00415463097 | 0.033753568 |
| | Glt8d1 | 1.26 | 1.777186367 | 0.01670373661 | 0.072478427 |
| | Med30 | 1.26 | 1.694466035 | 0.02020849469 | 0.079909331 |
| | Arf4 | 1.25 | 1.958674002 | 0.01099831104 | 0.056433081 |
| | C1d | 1.25 | 2.313636974 | 0.00485694322 | 0.036412040 |
| | Atp1a2 | 1.25 | 1.541941307 | 0.02871168578 | 0.099496673 |
| | Smc3 | 1.25 | 1.918893294 | 0.01205332054 | 0.059384969 |
| | Phka1 | 1.25 | 1.624096922 | 0.02376309903 | 0.088394354 |
| | Tceal8 | 1.25 | 2.231985356 | 0.00586157929 | 0.040628295 |
| | A830080D01Rik | 1.25 | 1.538639751 | 0.02893078710 | 0.099883906 |
| | Pdcd10 | 1.25 | 2.179831842 | 0.00660949316 | 0.043175873 |
| | Zfp518a | 1.25 | 1.654628663 | 0.02214987792 | 0.084380780 |
| | Rplp1 | 1.25 | 1.379959183 | 0.04169085644 | 0.125382810 |
| | Slc12a9 | 1.25 | 1.406759735 | 0.03919586602 | 0.120256685 |
| | Smpd2 | 1.25 | 1.970062021 | 0.01071366294 | 0.055557879 |
| | Ppt1 | 1.25 | 2.668777665 | 0.00214398792 | 0.025453666 |
| | Slco1a4 | 1.25 | 1.494355336 | 0.03203647052 | 0.106368279 |
| | Pkn2 | 1.25 | 1.579810459 | 0.02631416181 | 0.094482074 |
| | Slc35f5 | 1.25 | 1.767464187 | 0.01708188575 | 0.073238136 |
| | Smarca1 | 1.25 | 1.376337148 | 0.04204001401 | 0.125936741 |
| | Maged2 | 1.25 | 2.070896496 | 0.00849382882 | 0.049351543 |
| | Pea15a | 1.25 | 1.894264471 | 0.01275661737 | 0.061397392 |
| | Rab8b | 1.25 | 1.327288955 | 0.04706640694 | 0.136012513 |

| | | | | | |
|---|---|---|---|---|---|
| | Ofd1 | 1.25 | 1.371121596 | 0.04254792685 | 0.127070433 |
| | Magoh | 1.25 | 2.214342376 | 0.00610460578 | 0.041342232 |
| | Prpf40a | 1.25 | 1.726667397 | 0.01876431015 | 0.076806788 |
| | Kctd18 | 1.25 | 1.522263203 | 0.03004255029 | 0.102303479 |
| | Rpl23a | 1.25 | 1.894684955 | 0.01274427238 | 0.061394633 |
| | Itm2c | 1.25 | 1.32061094 | 0.04779572571 | 0.137226535 |
| | Txndc9 | 1.25 | 2.597480599 | 0.00252650057 | 0.027050644 |
| | Echdc1 | 1.25 | 1.312067042 | 0.04874532365 | 0.138878186 |
| | Fam96a | 1.25 | 1.353255229 | 0.04433480175 | 0.130534500 |
| | Cib1 | 1.24 | 1.674407718 | 0.02116373342 | 0.082135518 |
| | Fnta | 1.24 | 3.067444285 | 0.00085616154 | 0.018362491 |
| | Scp2 | 1.24 | 2.321855313 | 0.00476589738 | 0.036141388 |
| | Atraid | 1.24 | 2.231405094 | 0.00586941619 | 0.040628295 |
| | Rpl41 | 1.24 | 1.519362646 | 0.03024386939 | 0.102841131 |
| | G6pdx | 1.24 | 1.734161509 | 0.01844329408 | 0.076150386 |
| | Slc30a10 | 1.24 | 1.584834596 | 0.02601150043 | 0.093713694 |
| | M6pr | 1.24 | 1.885537956 | 0.01301553558 | 0.062250774 |
| | Mbnl1 | 1.24 | 1.931964386 | 0.01169595298 | 0.058426042 |
| | Pnpt1 | 1.24 | 1.454548333 | 0.03511168466 | 0.112568749 |
| | Lrch2 | 1.24 | 1.304477412 | 0.04960467270 | 0.140379650 |
| | 2810001G20Rik | 1.24 | 1.624111345 | 0.02376230988 | 0.088394354 |
| | Tril | 1.24 | 2.183159842 | 0.00655903817 | 0.042958886 |
| | Pdcl3 | 1.24 | 1.335645836 | 0.04616939289 | 0.134097152 |

| | | | | | |
|---|---|---|---|---|---|
| | Trip4 | 1.24 | 1.391625105 | 0.04058587324 | 0.123390593 |
| | Cyb5a | 1.24 | 2.668974352 | 0.00214301716 | 0.025453666 |
| | Nras | 1.24 | 2.121391297 | 0.00756151300 | 0.046718862 |
| | Amot | 1.24 | 1.906933394 | 0.01238986591 | 0.060350637 |
| | Asnsd1 | 1.24 | 1.755969191 | 0.01754004929 | 0.074090937 |
| | Erlec1 | 1.24 | 1.905737681 | 0.01242402508 | 0.060474151 |
| | Pcca | 1.24 | 2.047832729 | 0.00895709687 | 0.050973497 |
| | Haus2 | 1.24 | 2.064703124 | 0.00861582514 | 0.049734726 |
| | Rpl19 | 1.24 | 1.671321208 | 0.02131467878 | 0.082400951 |
| | Nudt13 | 1.24 | 1.405194438 | 0.03933739186 | 0.120520647 |
| | Abcd3 | 1.24 | 1.69778902 | 0.02005446033 | 0.079552061 |
| | Papd4 | 1.24 | 1.598617652 | 0.02519894436 | 0.091772779 |
| | Riok1 | 1.24 | 1.738596925 | 0.01825589272 | 0.075775370 |
| | Papss1 | 1.24 | 1.731076554 | 0.01857477003 | 0.076395066 |
| | Igbp1 | 1.24 | 2.222414679 | 0.00599218648 | 0.040984885 |
| | Rhou | 1.24 | 1.332776843 | 0.04647540218 | 0.134796950 |
| | Macrod1 | 1.24 | 1.455275817 | 0.03505291845 | 0.112477329 |
| | Immp1l | 1.24 | 1.589708872 | 0.02572119418 | 0.093200991 |
| | Nln | 1.23 | 1.458653111 | 0.03478138635 | 0.112032590 |
| | Thoc2 | 1.23 | 1.735664186 | 0.01837958980 | 0.076070465 |
| | Eif2s2 | 1.23 | 3.688013524 | 0.00020510983 | 0.014121412 |
| | Snrpd1 | 1.23 | 2.409771549 | 0.00389249847 | 0.032814001 |
| | Ndrg2 | 1.23 | 1.419462895 | 0.03806598789 | 0.118206721 |

| | | | | | |
|---|---|---|---|---|---|
| | Smndc1 | 1.23 | 2.186292198 | 0.00651190119 | 0.042792938 |
| | Slc25a53 | 1.23 | 1.338602784 | 0.04585611061 | 0.133384593 |
| | Amn1 | 1.23 | 1.368344425 | 0.04282087873 | 0.127525291 |
| | Lrrcc1 | 1.23 | 1.589258967 | 0.02574785370 | 0.093252835 |
| | Sall2 | 1.23 | 1.547593935 | 0.02834040580 | 0.098813884 |
| | Lrrc75b | 1.23 | 1.535709259 | 0.02912666364 | 0.100308142 |
| | Hspe1 | 1.23 | 1.728195355 | 0.01869840859 | 0.076659809 |
| | Plxnb1 | 1.23 | 1.75517456 | 0.01757217178 | 0.074112711 |
| | Selenon | 1.23 | 1.691277965 | 0.02035738709 | 0.080290142 |
| | Ccdc141 | 1.23 | 1.479522795 | 0.03314951702 | 0.108826469 |
| | Vhl | 1.23 | 2.231881169 | 0.00586298564 | 0.040628295 |
| | Rps25 | 1.23 | 1.957609881 | 0.01102529248 | 0.056523731 |
| | Tmem59 | 1.23 | 2.103772768 | 0.00787457696 | 0.047709242 |
| | Glrx3 | 1.23 | 2.083319637 | 0.00825430216 | 0.048953126 |
| | Ilvbl | 1.23 | 1.695516571 | 0.02015967045 | 0.079762174 |
| | BC017643 | 1.23 | 1.324678952 | 0.04735011613 | 0.136545214 |
| | Rfk | 1.23 | 1.681303445 | 0.02083034941 | 0.081485326 |
| | Atp6v0a2 | 1.23 | 1.607487405 | 0.02468951706 | 0.090517250 |
| | Kif1c | 1.23 | 2.030372461 | 0.00932454263 | 0.052042462 |
| | Psenen | 1.23 | 1.982202619 | 0.01041831251 | 0.054655689 |
| | Chchd1 | 1.23 | 1.555053857 | 0.02785775683 | 0.097951238 |
| | Elmod2 | 1.23 | 1.696525352 | 0.02011289785 | 0.079668876 |
| | Tbc1d12 | 1.23 | 1.552506274 | 0.02802165143 | 0.098254067 |

| | | | | | |
|---|---|---|---|---|---|
| | Slirp | 1.23 | 1.363703572 | 0.04328091442 | 0.128394440 |
| | Aldoc | 1.23 | 1.427331564 | 0.03738250806 | 0.116929898 |
| | Spg21 | 1.23 | 1.421545427 | 0.03788389049 | 0.117881010 |
| | Ppil3 | 1.23 | 1.577789547 | 0.02643689544 | 0.094675366 |
| | Pxmp4 | 1.23 | 1.712261256 | 0.01939718663 | 0.078117451 |
| | Tmem43 | 1.23 | 1.449137368 | 0.03555188497 | 0.113397969 |
| | Cdc42se1 | 1.23 | 1.807547268 | 0.01557588502 | 0.069784231 |
| | Slc30a1 | 1.22 | 1.508977546 | 0.03097579448 | 0.104436713 |
| | Ddx52 | 1.22 | 1.804677727 | 0.01567914127 | 0.070064091 |
| | Vrk3 | 1.22 | 1.455758268 | 0.03501400034 | 0.112477329 |
| | Lin7c | 1.22 | 2.003403218 | 0.00992194423 | 0.053591759 |
| | Zfp37 | 1.22 | 1.730753907 | 0.01858857477 | 0.076406104 |
| | Cnot6 | 1.22 | 1.708764088 | 0.01955401355 | 0.078450131 |
| | Mgat2 | 1.22 | 1.362864826 | 0.04336458292 | 0.128567685 |
| | Washc3 | 1.22 | 1.403279505 | 0.03951122504 | 0.120966830 |
| | Tcp11l2 | 1.22 | 1.349761518 | 0.04469289443 | 0.131055284 |
| | Rpl35 | 1.22 | 2.02135653 | 0.00952014296 | 0.052488446 |
| | Vps54 | 1.22 | 1.71949859 | 0.01907661918 | 0.077379379 |
| | Lrif1 | 1.22 | 1.357007364 | 0.04395341625 | 0.129717331 |
| | Rdh11 | 1.22 | 1.988384885 | 0.01027105642 | 0.054429073 |
| | Rpl11 | 1.22 | 1.527497824 | 0.02968261619 | 0.101497148 |
| | Cebpg | 1.22 | 1.719321532 | 0.01908439813 | 0.077379379 |
| | Phip | 1.22 | 2.000753448 | 0.00998266627 | 0.053718891 |

| | | | | | |
|---|---|---|---|---|---|
| | Hibadh | 1.22 | 2.025168668 | 0.00943694300 | 0.052393145 |
| | Rpl21 | 1.22 | 1.7096769 | 0.01951295758 | 0.078393478 |
| | Csrp1 | 1.22 | 1.734873568 | 0.01841307965 | 0.076140273 |
| | Eva1a | 1.22 | 1.456327524 | 0.03496813546 | 0.112423292 |
| | Rab9 | 1.22 | 1.603090301 | 0.02494076092 | 0.091098244 |
| | Galc | 1.22 | 1.337886838 | 0.04593176795 | 0.133576386 |
| | Dcaf13 | 1.22 | 2.076065243 | 0.00839333886 | 0.049245461 |
| | Osgep | 1.22 | 1.79837122 | 0.01590848343 | 0.070515636 |
| | Zfp760 | 1.22 | 1.333735577 | 0.04637291783 | 0.134546086 |
| | 1110059E24Rik | 1.22 | 2.322556493 | 0.00475820892 | 0.036113508 |
| | Fgfr3 | 1.22 | 1.532927813 | 0.02931380446 | 0.100750619 |
| | Nudt10 | 1.22 | 1.477074767 | 0.03333690161 | 0.109145191 |
| | Rbbp9 | 1.22 | 1.726134164 | 0.01878736340 | 0.076836142 |
| | Tmco3 | 1.22 | 1.579537501 | 0.02633070569 | 0.094506114 |
| | Uchl3 | 1.22 | 1.618658454 | 0.02406254428 | 0.088992445 |
| | Tjp1 | 1.22 | 1.50462312 | 0.03128793350 | 0.105048396 |
| | Med21 | 1.22 | 1.716442916 | 0.01921131460 | 0.077697566 |
| | Agpat5 | 1.22 | 1.997220424 | 0.01006420736 | 0.053827531 |
| | Mpst | 1.22 | 1.30960412 | 0.04902254792 | 0.139350193 |
| | Sarnp | 1.21 | 2.267510047 | 0.00540119619 | 0.038894788 |
| | Zfand6 | 1.21 | 2.622810259 | 0.00238336052 | 0.026569497 |
| | Cbx3 | 1.21 | 2.886500318 | 0.00129867261 | 0.020991075 |
| | Sh3glb1 | 1.21 | 1.565412005 | 0.02720119572 | 0.096584430 |

| | | | | | |
|---|---|---|---|---|---|
| | Myg1 | 1.21 | 1.533127867 | 0.02930030447 | 0.100750619 |
| | Prpf38a | 1.21 | 1.460877097 | 0.03460372908 | 0.111584362 |
| | Nme7 | 1.21 | 1.30372667 | 0.04969049585 | 0.140551071 |
| | 8-Mar | 1.21 | 1.341667223 | 0.04553368268 | 0.132794670 |
| | Cmc1 | 1.21 | 1.501223894 | 0.03153378528 | 0.105620703 |
| | Pccb | 1.21 | 2.681094485 | 0.00208403743 | 0.025136312 |
| | Ankfy1 | 1.21 | 1.570368191 | 0.02689253910 | 0.095956993 |
| | Pgrmc1 | 1.21 | 1.690004857 | 0.02041715109 | 0.080410454 |
| | Pcna | 1.21 | 1.74486101 | 0.01799446713 | 0.075292927 |
| | Ptges3 | 1.21 | 2.446726591 | 0.00357497830 | 0.031575132 |
| | Tpcn1 | 1.21 | 1.323013872 | 0.04753200430 | 0.136839990 |
| | Topbp1 | 1.21 | 1.549643035 | 0.02820700431 | 0.098624032 |
| | 2410015M20Rik | 1.21 | 2.060636289 | 0.00869688468 | 0.050059592 |
| | Btf3 | 1.21 | 1.97635452 | 0.01055955169 | 0.055065958 |
| | Churc1 | 1.21 | 1.545403263 | 0.02848372189 | 0.099112389 |
| | Cav2 | 1.21 | 1.358394818 | 0.04381322100 | 0.129470424 |
| | Rad50 | 1.21 | 1.366696609 | 0.04298365991 | 0.127788505 |
| | Gmfb | 1.21 | 2.502532275 | 0.00314389277 | 0.029690880 |
| | Ptpmt1 | 1.21 | 1.787071758 | 0.01632782142 | 0.071626989 |
| | Pet100 | 1.21 | 1.541257079 | 0.02875695655 | 0.099584007 |
| | Hmgb1 | 1.21 | 1.615489685 | 0.02423875535 | 0.089383015 |
| | Tra2b | 1.21 | 1.57868549 | 0.02638241274 | 0.094554182 |
| | Degs1 | 1.21 | 1.540784177 | 0.02878828693 | 0.099667385 |

| | | | | | |
|---|---|---|---|---|---|
| | Commd3 | 1.21 | 2.309424795 | 0.00490427941 | 0.036664692 |
| | Wdr44 | 1.21 | 1.960326596 | 0.01095653938 | 0.056348790 |
| | AU019823 | 1.21 | 1.421087127 | 0.03792388958 | 0.117894224 |
| | Ssb | 1.21 | 2.575262835 | 0.00265911528 | 0.027727676 |
| | Prps2 | 1.21 | 2.23934288 | 0.00576311279 | 0.040285418 |
| | Mrps14 | 1.20 | 2.579078013 | 0.00263585786 | 0.027675522 |
| | Sox8 | 1.20 | 1.326406823 | 0.04716210454 | 0.136231759 |
| | Ubxn4 | 1.20 | 1.557348609 | 0.02771094850 | 0.097685003 |
| | Ercc6l2 | 1.20 | 1.332417342 | 0.04651388959 | 0.134869668 |
| | Ergic3 | 1.20 | 1.548999221 | 0.02824885042 | 0.098706250 |
| | Srsf10 | 1.20 | 2.112231813 | 0.00772268262 | 0.047446521 |
| | Suz12 | 1.20 | 1.715406823 | 0.01925720161 | 0.077759944 |
| | Abhd17c | 1.20 | 1.774620928 | 0.01680269993 | 0.072729703 |
| | Smarcad1 | 1.20 | 1.445113411 | 0.03588282189 | 0.114003666 |
| | Vps25 | 1.20 | 1.803511219 | 0.01572131178 | 0.070084005 |
| | Nacc2 | 1.20 | 1.509146256 | 0.03096376365 | 0.104436713 |
| | Fam118b | 1.20 | 1.678162493 | 0.02098154705 | 0.081788822 |
| | Rnf2 | 1.20 | 1.724759915 | 0.01884690691 | 0.076924296 |
| | Zfp24 | 1.20 | 1.628020065 | 0.02354940479 | 0.087861084 |
| | Taldo1 | 1.20 | 1.356641343 | 0.04399047560 | 0.129783849 |
| | Mrpl39 | 1.20 | 2.506244839 | 0.00311713176 | 0.029668607 |
| | Psma2 | 1.20 | 1.669877158 | 0.02138566908 | 0.082475576 |
| | Mipep | 1.20 | 2.015710871 | 0.00964470901 | 0.052736947 |

| | | | | | |
|---|---|---|---|---|---|
| | Fam208a | 1.20 | 1.395620175 | 0.04021423625 | 0.122442681 |
| | Txndc12 | 1.20 | 1.567950668 | 0.02704265529 | 0.096267649 |
| | Sdhaf1 | 1.20 | 1.377728605 | 0.04190553554 | 0.125670878 |
| | Efcab14 | 1.20 | 1.759762075 | 0.01738753129 | 0.073741379 |
| | 2810025M15Rik | 1.20 | 1.424093856 | 0.03766223978 | 0.117495157 |
| | Rnf130 | 1.20 | 1.508648043 | 0.03099930497 | 0.104436713 |
| | C1galt1c1 | 1.20 | 1.44100807 | 0.03622362670 | 0.114662256 |
| | Zfp932 | 1.20 | 1.831008243 | 0.01475678526 | 0.067500994 |
| | Mettl6 | 1.20 | 1.401528945 | 0.03967080881 | 0.121163944 |
| | Ufl1 | 1.20 | 1.531923223 | 0.02938169029 | 0.100890549 |
| | Thoc1 | 1.20 | 1.611068953 | 0.02448674437 | 0.090110432 |
| | Selenot | 1.20 | 1.825583526 | 0.01494226637 | 0.067987312 |
| | Vbp1 | 1.20 | 2.061444335 | 0.00868071833 | 0.050013313 |
| | Wnk1 | 1.20 | 2.169001414 | 0.00677639301 | 0.043782476 |
| | Frg1 | 1.20 | 1.381041711 | 0.04158706669 | 0.125255842 |
| | Cspg5 | 1.20 | 1.560159245 | 0.02753218983 | 0.097429776 |
| | Aph1a | 1.20 | 1.498800596 | 0.03171023088 | 0.105785453 |
| | Yipf5 | 1.20 | 2.179109425 | 0.00662049673 | 0.043196698 |
| | Ptprg | 1.20 | 1.986762269 | 0.01030950304 | 0.054483415 |
| | Hps3 | 1.19 | 1.313993654 | 0.04852955912 | 0.138520817 |
| | Rpl3 | 1.19 | 1.724688663 | 0.01884999926 | 0.076924296 |
| | Chchd3 | 1.19 | 2.082845738 | 0.00826331411 | 0.048969915 |
| | Sgpp1 | 1.19 | 1.9179323 | 0.01208002131 | 0.059452569 |

| | | | | | |
|---|---|---|---|---|---|
| | Fabp5 | 1.19 | 2.038001069 | 0.00916218235 | 0.051569310 |
| | Ccnc | 1.19 | 1.710881096 | 0.01945892767 | 0.078274334 |
| | Hsd17b12 | 1.19 | 1.683650675 | 0.02071807137 | 0.081176795 |
| | Dtna | 1.19 | 1.637746408 | 0.02302786061 | 0.086526069 |
| | Tmem68 | 1.19 | 1.312155762 | 0.04873536665 | 0.138878186 |
| | Tbca | 1.19 | 1.461041123 | 0.03459066229 | 0.111575186 |
| | Aldh7a1 | 1.19 | 2.191570423 | 0.00643323738 | 0.042479152 |
| | Fam174b | 1.19 | 1.377543189 | 0.04192343030 | 0.125697110 |
| | Snx4 | 1.19 | 1.826974742 | 0.01489447701 | 0.067814781 |
| | Rpl37a | 1.19 | 1.628462458 | 0.02352542845 | 0.087819319 |
| | Wtap | 1.19 | 1.900626532 | 0.01257110542 | 0.060845213 |
| | Usp25 | 1.19 | 2.35362966 | 0.00442965946 | 0.034726255 |
| | Rchy1 | 1.19 | 2.021557855 | 0.00951573075 | 0.052488446 |
| | Psmb3 | 1.19 | 2.026743793 | 0.00940277853 | 0.052233414 |
| | Sephs1 | 1.19 | 1.720686025 | 0.01902453171 | 0.077345375 |
| | Tgoln1 | 1.19 | 1.84529488 | 0.01427924090 | 0.065820547 |
| | Ap3m1 | 1.19 | 1.982488157 | 0.01041146499 | 0.054655689 |
| | Lsm14a | 1.19 | 1.902509453 | 0.01251672027 | 0.060731728 |
| | Rpf2 | 1.19 | 1.5757437 | 0.02656172647 | 0.094965121 |
| | Nsdhl | 1.19 | 2.196249942 | 0.00636429143 | 0.042267631 |
| | Galnt16 | 1.19 | 1.927197704 | 0.01182503121 | 0.058855750 |
| | Dnttip2 | 1.19 | 1.92267974 | 0.01194868906 | 0.059166464 |
| | Mrps33 | 1.19 | 1.862822157 | 0.01371443255 | 0.064055071 |

| | | | | | |
|---|---|---|---|---|---|
| | Ptpn12 | 1.19 | 1.738240618 | 0.01827087651 | 0.075794131 |
| | Hif1a | 1.19 | 1.462591341 | 0.03446741067 | 0.111305199 |
| | Ctsb | 1.19 | 1.423917129 | 0.03767756873 | 0.117496086 |
| | Zhx1 | 1.19 | 1.786181481 | 0.01636132677 | 0.071626989 |
| | Rnf128 | 1.19 | 1.61610887 | 0.02420422214 | 0.089309940 |
| | Lrp6 | 1.19 | 1.531458377 | 0.02941315578 | 0.100928405 |
| | Capns1 | 1.19 | 1.830577808 | 0.01477141814 | 0.067539770 |
| | 4930430F08Rik | 1.19 | 1.565567071 | 0.02719148518 | 0.096584430 |
| | Gsta4 | 1.19 | 1.797343608 | 0.01594617008 | 0.070637113 |
| | Snrpd2 | 1.19 | 1.77147059 | 0.01692502856 | 0.072949681 |
| | Fyco1 | 1.18 | 1.375720479 | 0.04209975043 | 0.125992885 |
| | Slc20a2 | 1.18 | 1.772689919 | 0.01687757634 | 0.072883336 |
| | Timp4 | 1.18 | 1.357421337 | 0.04391153951 | 0.129649434 |
| | Txndc17 | 1.18 | 1.724601289 | 0.01885379197 | 0.076924296 |
| | Dbnl | 1.18 | 1.352481241 | 0.04441388455 | 0.130598606 |
| | Zfp617 | 1.18 | 1.626806814 | 0.02361528469 | 0.087987426 |
| | Rpl24 | 1.18 | 1.548343184 | 0.02829155483 | 0.098706250 |
| | Atg5 | 1.18 | 1.498874883 | 0.03170480722 | 0.105785453 |
| | Nsmce4a | 1.18 | 1.526759702 | 0.02973310732 | 0.101506243 |
| | Cxcl14 | 1.18 | 1.660140057 | 0.02187056202 | 0.083711251 |
| | Arsb | 1.18 | 1.541401751 | 0.02874737860 | 0.099575934 |
| | C330007P06Rik | 1.18 | 1.651153951 | 0.02232780595 | 0.084776563 |
| | Trmt10c | 1.18 | 1.460801559 | 0.03460974830 | 0.111584362 |

| | | | | | |
|---|---|---|---|---|---|
| | Pcyt1a | 1.18 | 1.380626258 | 0.04162686860 | 0.125327520 |
| | Lin52 | 1.18 | 1.682562635 | 0.02077004154 | 0.081307026 |
| | Snrnp27 | 1.18 | 1.575670833 | 0.02656618343 | 0.094965121 |
| | 1600012H06Rik | 1.18 | 1.757975587 | 0.01745920293 | 0.073936737 |
| | Metap2 | 1.18 | 1.700057486 | 0.01994998228 | 0.079366736 |
| | Uba52 | 1.18 | 2.423736887 | 0.00376932090 | 0.032351180 |
| | Dnajb9 | 1.18 | 1.478404375 | 0.03323499552 | 0.109045385 |
| | Rps29 | 1.18 | 1.342260001 | 0.04547157518 | 0.132687389 |
| | Rpl9 | 1.18 | 2.509369565 | 0.00309478465 | 0.029572626 |
| | Pbrm1 | 1.18 | 1.621544468 | 0.02390317177 | 0.088590457 |
| | Nudt9 | 1.18 | 1.501013097 | 0.03154909479 | 0.105620703 |
| | Rbm39 | 1.18 | 2.186620301 | 0.00650698340 | 0.042781081 |
| | Adam10 | 1.18 | 1.489823415 | 0.03237252576 | 0.107110734 |
| | Ddrgk1 | 1.18 | 2.391629817 | 0.00405854329 | 0.033514690 |
| | Tmx1 | 1.18 | 1.31358695 | 0.04857502695 | 0.138590741 |
| | Pdrg1 | 1.18 | 1.527316339 | 0.02969502263 | 0.101497148 |
| | Psma6 | 1.18 | 2.201455151 | 0.00628846790 | 0.041966210 |
| | Gtf2b | 1.18 | 1.759600173 | 0.01739401446 | 0.073746113 |
| | Adgrl4 | 1.18 | 1.595824187 | 0.02536155123 | 0.092193935 |
| | Gdi2 | 1.18 | 2.503591935 | 0.00313623115 | 0.029668607 |
| | Tle1 | 1.18 | 1.953267416 | 0.01113608621 | 0.056842853 |
| | Stx8 | 1.18 | 1.856708264 | 0.01390886640 | 0.064611810 |
| | Arl6ip1 | 1.18 | 2.464513026 | 0.00343152347 | 0.030940003 |

| | | | | | |
|---|---|---|---|---|---|
| | Rpl7l1 | 1.18 | 1.6489303 18 | 0.02244241 979 | 0.0851176 62 |
| | Rb1 | 1.18 | 2.0245977 09 | 0.00944935 774 | 0.0524085 46 |
| | Bpgm | 1.17 | 2.2262470 63 | 0.00593954 172 | 0.0408334 54 |
| | Capn2 | 1.17 | 1.5188121 9 | 0.03028222 699 | 0.1028808 83 |
| | Qars | 1.17 | 2.3823279 92 | 0.00414640 775 | 0.0337134 85 |
| | Vma21 | 1.17 | 1.7480003 8 | 0.01786486 010 | 0.0748936 83 |
| | Sptlc1 | 1.17 | 1.4348166 53 | 0.03674373 897 | 0.1156321 32 |
| | Ptch1 | 1.17 | 1.4996098 61 | 0.03165119 696 | 0.1057480 14 |
| | Cant1 | 1.17 | 1.4707678 91 | 0.03382455 632 | 0.1099558 15 |
| | Hsf2 | 1.17 | 1.5155699 95 | 0.03050914 275 | 0.1034104 91 |
| | Picalm | 1.17 | 1.9233106 59 | 0.01193134 326 | 0.0591344 41 |
| | Atox1 | 1.17 | 1.3044096 57 | 0.04961241 220 | 0.1403796 50 |
| | Htra1 | 1.17 | 1.8947895 38 | 0.01274120 378 | 0.0613946 33 |
| | Arfgap3 | 1.17 | 1.4690277 31 | 0.03396035 871 | 0.1102831 33 |
| | Rrm2b | 1.17 | 1.5357291 61 | 0.02912532 894 | 0.1003081 42 |
| | Cept1 | 1.17 | 2.0591077 11 | 0.00872754 887 | 0.0500783 46 |
| | Psma1 | 1.17 | 1.9160065 97 | 0.01213370 419 | 0.0596100 21 |
| | Lrrc58 | 1.17 | 1.4938111 | 0.03207664 222 | 0.1064650 10 |
| | Rab5c | 1.17 | 1.3532958 47 | 0.04433065 551 | 0.1305345 00 |
| | Cdc16 | 1.17 | 1.3456800 62 | 0.04511489 370 | 0.1319829 16 |
| | Ammecr1l | 1.17 | 1.3019965 33 | 0.04988884 697 | 0.1409960 19 |
| | Mcfd2 | 1.17 | 1.3201954 59 | 0.04784147 282 | 0.1372994 31 |
| | Cln8 | 1.17 | 1.7193339 68 | 0.01908385 166 | 0.0773793 79 |

| | | | | | |
|---|---|---|---|---|---|
| | Emc2 | 1.17 | 1.7713404<br>2 | 0.01693010<br>219 | 0.0729496<br>81 |
| | Tma7 | 1.17 | 2.2866058<br>82 | 0.00516885<br>225 | 0.0378202<br>99 |
| | Chmp2a | 1.17 | 1.9114515<br>97 | 0.01226163<br>554 | 0.0599668<br>79 |
| | Osbpl9 | 1.16 | 1.6590599<br>29 | 0.02192502<br>368 | 0.0837576<br>84 |
| | Acad11 | 1.16 | 1.7608227<br>75 | 0.01734511<br>666 | 0.0736524<br>25 |
| | Ccdc50 | 1.16 | 1.3126578<br>78 | 0.04867905<br>314 | 0.1387756<br>99 |
| | Srsf3 | 1.16 | 1.5661754<br>17 | 0.02715342<br>287 | 0.0965619<br>01 |
| | Rwdd1 | 1.16 | 1.4744221<br>5 | 0.03354114<br>235 | 0.1093708<br>68 |
| | Nol7 | 1.16 | 1.5037983<br>98 | 0.03134740<br>552 | 0.1051622<br>80 |
| | Btbd1 | 1.16 | 1.9696991<br>65 | 0.01072261<br>801 | 0.0555578<br>79 |
| | Zdhhc20 | 1.16 | 1.4637804<br>96 | 0.03437316<br>352 | 0.1111345<br>03 |
| | Paip2 | 1.16 | 1.7656781<br>34 | 0.01715228<br>034 | 0.0733760<br>00 |
| | Bbip1 | 1.16 | 1.4153061<br>9 | 0.03843207<br>298 | 0.1188433<br>09 |
| | Atg3 | 1.16 | 2.1043242<br>35 | 0.00786458<br>417 | 0.0477092<br>42 |
| | Ndufc2 | 1.16 | 1.3586588<br>43 | 0.04378659<br>330 | 0.1294195<br>69 |
| | Canx | 1.16 | 1.4443776<br>78 | 0.03594366<br>203 | 0.1140914<br>44 |
| | Tm9sf3 | 1.16 | 2.0712335<br>34 | 0.00848723<br>967 | 0.0493515<br>43 |
| | Mrpl9 | 1.16 | 1.4511466<br>88 | 0.03538777<br>945 | 0.1130056<br>88 |
| | Tom1 | 1.16 | 1.5533086<br>03 | 0.02796993<br>115 | 0.0981199<br>26 |
| | Mex3c | 1.16 | 1.3115173<br>5 | 0.04880706<br>020 | 0.1390252<br>52 |
| | Rab18 | 1.16 | 1.7552730<br>32 | 0.01756818<br>792 | 0.0741127<br>11 |
| | Cwc15 | 1.16 | 1.7332910<br>8 | 0.01848029<br>587 | 0.0761896<br>89 |
| | Hnrnpc | 1.16 | 1.3255648<br>82 | 0.04725362<br>362 | 0.1363528<br>02 |

| | | | | | |
|---|---|---|---|---|---|
| | Tspan3 | 1.16 | 1.745638071 | 0.01796229931 | 0.075181223 |
| | Tmem129 | 1.15 | 1.569862436 | 0.02692387492 | 0.096043864 |
| | Cdc42 | 1.15 | 2.245868026 | 0.00567717098 | 0.039912161 |
| | Commd8 | 1.15 | 1.497209392 | 0.03182662654 | 0.106025682 |
| | Psma4 | 1.15 | 1.607522907 | 0.02468749884 | 0.090517250 |
| | Mapre1 | 1.15 | 1.316643555 | 0.04823435168 | 0.138023371 |
| | Rnf181 | 1.15 | 1.645881197 | 0.02260053934 | 0.085432184 |
| | Brox | 1.15 | 1.421335569 | 0.03790220106 | 0.117887608 |
| | Rps15 | 1.15 | 1.561268751 | 0.02746194224 | 0.097256327 |
| | Cacybp | 1.15 | 1.877355626 | 0.01326307954 | 0.062785576 |
| | 0610037L13Rik | 1.15 | 1.449787552 | 0.03549869989 | 0.113254617 |
| | Ube2a | 1.15 | 1.429656316 | 0.03718293641 | 0.116517840 |
| | Dnajc10 | 1.15 | 1.302426002 | 0.04983953684 | 0.140933116 |
| | Nubp2 | 1.15 | 1.44537836 | 0.03586093760 | 0.113976303 |
| | Tmem87a | 1.15 | 1.771854114 | 0.01691008872 | 0.072949681 |
| | Cldnd1 | 1.15 | 1.556089538 | 0.02779140239 | 0.097843110 |
| | Ntan1 | 1.15 | 2.125066098 | 0.00749780087 | 0.046429600 |
| | Actr3 | 1.15 | 1.439389548 | 0.03635887625 | 0.114957966 |
| | Psmb1 | 1.15 | 1.305961536 | 0.04943544683 | 0.140031432 |
| | Spcs1 | 1.14 | 1.393589804 | 0.04040268210 | 0.122934733 |
| | Rbm45 | 1.14 | 1.346914729 | 0.04498681750 | 0.131664294 |
| | Rap2c | 1.14 | 1.476225464 | 0.03340215874 | 0.109189632 |
| | Ankra2 | 1.14 | 1.445582736 | 0.03584406565 | 0.113959580 |

| | | | | | |
|---|---|---|---|---|---|
| | Ubr7 | 1.14 | 1.999844686 | 0.01000357688 | 0.053745583 |
| | Srp9 | 1.14 | 1.710011825 | 0.01949791512 | 0.078382862 |
| | Glod4 | 1.14 | 1.405765943 | 0.03928566028 | 0.120469596 |
| | Bag1 | 1.14 | 2.10547744 | 0.00784372863 | 0.047709242 |
| | Ppig | 1.14 | 1.407940868 | 0.03908941149 | 0.120055343 |
| | Paics | 1.14 | 1.6294025 | 0.02347456213 | 0.087708619 |
| | Naa50 | 1.14 | 1.440447982 | 0.03627037264 | 0.114726370 |
| | Fam45a | 1.14 | 1.90370843 | 0.01248221243 | 0.060628519 |
| | Cox6c | 1.14 | 1.400300564 | 0.03978317466 | 0.121399201 |
| | Sdcbp | 1.14 | 1.344364951 | 0.04525171558 | 0.132257650 |
| | Svip | 1.14 | 1.481072279 | 0.03303145624 | 0.108562745 |
| | Srp19 | 1.14 | 1.438249709 | 0.03645442820 | 0.115099220 |
| | Eif2s1 | 1.13 | 1.695661511 | 0.02015294357 | 0.079758525 |
| | 4933434E20Rik | 1.13 | 1.602562753 | 0.02497107545 | 0.091160347 |
| | Ireb2 | 1.13 | 1.673105874 | 0.02122726911 | 0.082233974 |
| | Heatr3 | 1.13 | 1.45415568 | 0.03514344406 | 0.112572639 |
| | Amfr | 1.13 | 1.647833255 | 0.02249918286 | 0.085238840 |
| | Sec62 | 1.13 | 1.978185737 | 0.01051512072 | 0.054875911 |
| | Mrpl3 | 1.13 | 1.444811548 | 0.03590777144 | 0.114029026 |
| | Pex7 | 1.13 | 1.325682842 | 0.04724079067 | 0.136344403 |
| | Cnbp | 1.13 | 1.759859327 | 0.01738363812 | 0.073741379 |
| | Phf20l1 | 1.13 | 1.323088359 | 0.04752385265 | 0.136839990 |
| | Mpv17 | 1.13 | 1.54705229 | 0.02837577355 | 0.098895281 |

| | | | | | |
|---|---|---|---|---|---|
| | Ctsf | 1.13 | 1.305174304 | 0.04952513814 | 0.140227678 |
| | Seh1l | 1.13 | 1.441099802 | 0.03621597635 | 0.114662256 |
| | Gpx4 | 1.12 | 1.378672366 | 0.04181456992 | 0.125532802 |
| | Acp2 | 1.12 | 1.45424522 | 0.03513619917 | 0.112572639 |
| | Thap12 | 1.12 | 1.742142036 | 0.01810747791 | 0.075534362 |
| | Hnrnpk | 1.12 | 1.352338459 | 0.04442848877 | 0.130598606 |
| | Uqcrh | 1.12 | 1.629067527 | 0.02349267510 | 0.087744726 |
| | Gosr2 | 1.12 | 1.337741711 | 0.04594711943 | 0.133592757 |
| | Acox1 | 1.12 | 1.425526728 | 0.03753818516 | 0.117283357 |
| | Psma3 | 1.12 | 1.52961811 | 0.02953805469 | 0.101116694 |
| | Tcea1 | 1.11 | 1.528590979 | 0.02960799649 | 0.101305647 |
| | Rsl1d1 | 1.11 | 1.313926464 | 0.04853706777 | 0.138520817 |
| | Vps4b | 1.11 | 1.414270221 | 0.03852385851 | 0.118956481 |
| | Dync1i2 | 1.11 | 1.34101149 | 0.04560248504 | 0.132928245 |
| | Myl6 | 1.11 | 1.434756615 | 0.03674881888 | 0.115632132 |
| | Uri1 | 1.10 | 1.339682993 | 0.04574219566 | 0.133222448 |
| | Washc5 | 1.10 | 1.308389117 | 0.04915988781 | 0.139519308 |
| | Slc30a5 | 1.10 | 1.308075423 | 0.04919540919 | 0.139581689 |
| | Arf2 | 1.10 | 1.324153897 | 0.04740739617 | 0.136595729 |
| | Srp54c | 1.09 | 1.306658752 | 0.04935614694 | 0.139922182 |

| DOWN p<0.05 | Gene ID | Fold change (BRAF V600E vs. control-FP) | log-10 P-values | P-value | FDR step up |
|---|---|---|---|---|---|
| | Ube2v1 | -1.09 | 1.436743048 | 0.03658111610 | 1.15E-01 |
| | Eif4g1 | -1.09 | 1.324157494 | 0.04740700362 | 1.37E-01 |
| | Atp6v0b | -1.09 | 1.352228307 | 0.04443975880 | 1.31E-01 |
| | Sqstm1 | -1.10 | 1.356310631 | 0.04402398675 | 1.30E-01 |
| | Wdr82 | -1.10 | 1.688667803 | 0.02048010585 | 8.06E-02 |
| | Vps39 | -1.10 | 1.582742338 | 0.02613711578 | 9.40E-02 |
| | Ankrd13c | -1.10 | 1.474550006 | 0.03353126936 | 1.09E-01 |
| | Ciapin1 | -1.10 | 1.484824211 | 0.03274732191 | 1.08E-01 |
| | Csnk1g3 | -1.10 | 1.367741263 | 0.04288039100 | 1.28E-01 |
| | Nrd1 | -1.10 | 1.407357226 | 0.03914197846 | 1.20E-01 |
| | Snx19 | -1.10 | 1.326176829 | 0.04718708729 | 1.36E-01 |
| | Tmem184b | -1.10 | 1.421005296 | 0.03793103598 | 1.18E-01 |
| | Eif2ak1 | -1.10 | 1.481057358 | 0.03303259116 | 1.09E-01 |
| | Hdlbp | -1.11 | 1.543513831 | 0.02860791252 | 9.94E-02 |
| | Rtf1 | -1.11 | 1.448042477 | 0.03564162715 | 1.14E-01 |
| | Rab3gap1 | -1.11 | 1.347499915 | 0.04492624123 | 1.32E-01 |
| | Samm50 | -1.11 | 1.508213079 | 0.03103036760 | 1.04E-01 |
| | Rnf10 | -1.11 | 1.44383939 | 0.03598824014 | 1.14E-01 |
| | Ipo5 | -1.11 | 1.689189305 | 0.02045552807 | 8.05E-02 |
| | Vps4a | -1.11 | 1.46971511 | 0.03390665054 | 1.10E-01 |
| | Glyr1 | -1.11 | 1.317378163 | 0.04815283233 | 1.38E-01 |
| | Abcf1 | -1.11 | 1.501127143 | 0.03154081106 | 1.06E-01 |

| | | | | | |
|---|---|---|---|---|---|
| | 6-Mar | -1.11 | 1.487130637 | 0.03257387030 | 1.08E-01 |
| | Otud5 | -1.11 | 1.500975164 | 0.03155185051 | 1.06E-01 |
| | Mtmr3 | -1.11 | 1.430237001 | 0.03713325324 | 1.16E-01 |
| | Akt1 | -1.11 | 1.387884861 | 0.04093691766 | 1.24E-01 |
| | Nsd3 | -1.11 | 1.406728051 | 0.03919872574 | 1.20E-01 |
| | Arl8a | -1.11 | 1.453643735 | 0.03518489542 | 1.13E-01 |
| | Atg4b | -1.11 | 1.55045677 | 0.02815420248 | 9.85E-02 |
| | Rnf6 | -1.11 | 1.358710552 | 0.04378138016 | 1.29E-01 |
| | Ap1g1 | -1.11 | 1.434428927 | 0.03677655743 | 1.16E-01 |
| | Fbxo18 | -1.11 | 1.521551262 | 0.03009183955 | 1.02E-01 |
| | Yars | -1.11 | 1.578815503 | 0.02637451589 | 9.46E-02 |
| | Dgcr2 | -1.11 | 1.537824302 | 0.02898515970 | 1.00E-01 |
| | Pdap1 | -1.11 | 1.322394727 | 0.04759981593 | 1.37E-01 |
| | Dcaf6 | -1.11 | 1.374654992 | 0.04220316361 | 1.26E-01 |
| | Hectd1 | -1.11 | 1.322454621 | 0.04759325182 | 1.37E-01 |
| | Letmd1 | -1.11 | 1.575079376 | 0.02660238806 | 9.51E-02 |
| | Sppl3 | -1.11 | 1.472698747 | 0.03367450751 | 1.10E-01 |
| | Xpo7 | -1.11 | 1.432633849 | 0.03692888125 | 1.16E-01 |
| | Neu1 | -1.11 | 1.309021386 | 0.04908837032 | 1.39E-01 |
| | Vdac1 | -1.11 | 1.403453954 | 0.03949535718 | 1.21E-01 |
| | Leprotl1 | -1.11 | 1.412328153 | 0.03869651432 | 1.19E-01 |
| | Ubl4a | -1.12 | 1.61070666 | 0.02450717998 | 9.01E-02 |
| | Cs | -1.12 | 1.630241302 | 0.02342926683 | 8.76E-02 |

| | | | | | |
|---|---|---|---|---|---|
| | Tex264 | -1.12 | 1.487398303 | 0.03255380047 | 1.07E-01 |
| | Togaram1 | -1.12 | 1.429380642 | 0.03720654623 | 1.17E-01 |
| | Rhot1 | -1.12 | 1.500538179 | 0.03158361380 | 1.06E-01 |
| | Poldip2 | -1.12 | 1.323607625 | 0.04746706453 | 1.37E-01 |
| | Gas6 | -1.12 | 1.516342344 | 0.03045493352 | 1.03E-01 |
| | Pafah1b2 | -1.12 | 1.402601511 | 0.03957295572 | 1.21E-01 |
| | Ndufa10 | -1.12 | 1.538944182 | 0.02891051430 | 9.99E-02 |
| | Hars | -1.12 | 1.496233717 | 0.03189820778 | 1.06E-01 |
| | Prpf19 | -1.12 | 1.786248118 | 0.01635881653 | 7.16E-02 |
| | Elk1 | -1.12 | 1.367470373 | 0.04290714586 | 1.28E-01 |
| | Zdhhc5 | -1.12 | 1.412521005 | 0.03867933470 | 1.19E-01 |
| | Vapb | -1.12 | 1.637295356 | 0.02305178943 | 8.66E-02 |
| | Tex2 | -1.12 | 1.597792516 | 0.02524686654 | 9.18E-02 |
| | Tmed8 | -1.12 | 1.325685932 | 0.04724045458 | 1.36E-01 |
| | Urgcp | -1.12 | 1.677572057 | 0.02101009146 | 8.18E-02 |
| | Trim41 | -1.12 | 1.487030582 | 0.03258137575 | 1.08E-01 |
| | Tacc2 | -1.12 | 1.378585328 | 0.04182295086 | 1.26E-01 |
| | Ikbkap | -1.12 | 1.433435837 | 0.03686074962 | 1.16E-01 |
| | Eif4h | -1.12 | 1.778251201 | 0.01666283137 | 7.24E-02 |
| | Fam20b | -1.12 | 1.888054476 | 0.01294033514 | 6.20E-02 |
| | Smap1 | -1.12 | 1.44076935 | 0.03624354339 | 1.15E-01 |
| | Med9 | -1.12 | 1.339139854 | 0.04579943767 | 1.33E-01 |
| | Ywhab | -1.12 | 1.405450882 | 0.03931417060 | 1.21E-01 |

| | | | | | |
|---|---|---|---|---|---|
| | Ttc17 | -1.12 | 1.410994038 | 0.03881556947 | 1.19E-01 |
| | Actl6b | -1.12 | 1.501173353 | 0.03153745522 | 1.06E-01 |
| | Ppp2r2b | -1.12 | 1.330042604 | 0.04676892588 | 1.35E-01 |
| | Srr | -1.12 | 1.880386977 | 0.01317082631 | 6.26E-02 |
| | Fam53c | -1.12 | 1.351623473 | 0.04450169237 | 1.31E-01 |
| | Rnf145 | -1.12 | 1.418451487 | 0.03815474124 | 1.18E-01 |
| | Hnrnpul2 | -1.12 | 1.664335728 | 0.02166029026 | 8.32E-02 |
| | Sec31a | -1.12 | 1.636965504 | 0.02306930422 | 8.66E-02 |
| | Bscl2 | -1.12 | 1.347628707 | 0.04491292020 | 1.32E-01 |
| | Setd3 | -1.12 | 1.316677309 | 0.04823060289 | 1.38E-01 |
| | Luc7l | -1.12 | 1.375295901 | 0.04214092841 | 1.26E-01 |
| | Zfp318 | -1.12 | 1.331183491 | 0.04664622567 | 1.35E-01 |
| | Cyth1 | -1.12 | 1.454264109 | 0.03513467101 | 1.13E-01 |
| | Gpatch2l | -1.12 | 1.387658209 | 0.04095828756 | 1.24E-01 |
| | Ranbp3 | -1.12 | 1.402948242 | 0.03954137415 | 1.21E-01 |
| | Ociad1 | -1.12 | 1.537983289 | 0.02897455077 | 1.00E-01 |
| | Supt5 | -1.12 | 1.77282442 | 0.01687235015 | 7.29E-02 |
| | Kif3b | -1.12 | 1.729450688 | 0.01864443863 | 7.65E-02 |
| | Kctd20 | -1.12 | 1.463036513 | 0.03443209810 | 1.11E-01 |
| | Bptf | -1.12 | 1.409507122 | 0.03894869204 | 1.20E-01 |
| | Add1 | -1.12 | 1.802766981 | 0.01574827605 | 7.01E-02 |
| | Trappc11 | -1.12 | 1.857713534 | 0.01387670853 | 6.45E-02 |
| | Tmem55b | -1.12 | 1.409299539 | 0.03896731309 | 1.20E-01 |

| | | | | | |
|---|---|---|---|---|---|
| | Sms | -1.12 | 1.659742219 | 0.02189060584 | 8.37E-02 |
| | Ank | -1.13 | 1.511021769 | 0.03083033411 | 1.04E-01 |
| | Pum1 | -1.13 | 1.69523264 | 0.02017285468 | 7.98E-02 |
| | Fkbp2 | -1.13 | 1.853578925 | 0.01400944965 | 6.49E-02 |
| | Dcaf7 | -1.13 | 1.546905866 | 0.02838534216 | 9.89E-02 |
| | Hs2st1 | -1.13 | 1.421744467 | 0.03786653209 | 1.18E-01 |
| | Upf2 | -1.13 | 1.350578138 | 0.04460893567 | 1.31E-01 |
| | Qrich1 | -1.13 | 1.930525912 | 0.01173475665 | 5.86E-02 |
| | Cpsf2 | -1.13 | 1.538069813 | 0.02896877873 | 1.00E-01 |
| | Foxj3 | -1.13 | 1.982209938 | 0.01041813694 | 5.47E-02 |
| | Polr2c | -1.13 | 1.619814646 | 0.02399856943 | 8.89E-02 |
| | Kat14 | -1.13 | 1.326209066 | 0.04718358483 | 1.36E-01 |
| | Vezt | -1.13 | 1.555085158 | 0.02785574910 | 9.80E-02 |
| | Dcun1d4 | -1.13 | 1.815645207 | 0.01528814499 | 6.89E-02 |
| | Ppp2r1a | -1.13 | 1.665725749 | 0.02159107425 | 8.30E-02 |
| | Ogdh | -1.13 | 1.388517141 | 0.04087736179 | 1.24E-01 |
| | Klhdc10 | -1.13 | 1.565495322 | 0.02719597781 | 9.66E-02 |
| | Atxn3 | -1.13 | 1.429464742 | 0.03719934197 | 1.17E-01 |
| | Elp3 | -1.13 | 1.564108811 | 0.02728294132 | 9.68E-02 |
| | Aars | -1.13 | 2.00943006 | 0.00978520526 | 5.32E-02 |
| | Rrp1 | -1.13 | 1.541879629 | 0.02871576370 | 9.95E-02 |
| | Atp6v1a | -1.13 | 1.342311859 | 0.04546614589 | 1.33E-01 |
| | B4galt6 | -1.13 | 1.779089606 | 0.01663069482 | 7.23E-02 |

| | | | | | |
|---|---|---|---|---|---|
| | Dhdds | -1.13 | 1.613643062 | 0.02434203813 | 8.96E-02 |
| | Atp6v1e1 | -1.13 | 1.476142776 | 0.03340851900 | 1.09E-01 |
| | Ube2j1 | -1.13 | 1.542764851 | 0.02865729198 | 9.94E-02 |
| | Supt6 | -1.13 | 1.920085506 | 0.01202027750 | 5.93E-02 |
| | Rab11b | -1.13 | 1.890613961 | 0.01286429644 | 6.17E-02 |
| | Ubqln1 | -1.13 | 1.531896731 | 0.02938348269 | 1.01E-01 |
| | Tm9sf4 | -1.13 | 1.676687124 | 0.02105294598 | 8.19E-02 |
| | Iars | -1.13 | 1.627583324 | 0.02357309879 | 8.79E-02 |
| | Slc35e1 | -1.13 | 1.86475287 | 0.01365359856 | 6.39E-02 |
| | Sik3 | -1.13 | 1.556518095 | 0.02776399167 | 9.78E-02 |
| | Kbtbd11 | -1.13 | 1.549734691 | 0.02820105198 | 9.86E-02 |
| | Psmd3 | -1.13 | 1.9229532 | 0.01194116776 | 5.92E-02 |
| | Hmg20a | -1.13 | 1.875634275 | 0.01331575283 | 6.29E-02 |
| | Rabl6 | -1.13 | 1.792456455 | 0.01612662715 | 7.11E-02 |
| | Slc25a36 | -1.13 | 1.387645034 | 0.04095953009 | 1.24E-01 |
| | Ankmy2 | -1.13 | 1.446582214 | 0.03576166958 | 1.14E-01 |
| | Ap1b1 | -1.13 | 1.883261227 | 0.01308394691 | 6.24E-02 |
| | Pik3c3 | -1.13 | 1.385050911 | 0.04120492130 | 1.24E-01 |
| | Msl1 | -1.13 | 1.662487046 | 0.02175268916 | 8.34E-02 |
| | Tmx2 | -1.13 | 1.506864608 | 0.03112686570 | 1.05E-01 |
| | Tspyl1 | -1.13 | 1.726196128 | 0.01878468306 | 7.68E-02 |
| | Ehd3 | -1.13 | 1.824788768 | 0.01496963571 | 6.81E-02 |
| | Larp4b | -1.13 | 1.852389711 | 0.01404786383 | 6.50E-02 |

| | | | | | |
|---|---|---|---|---|---|
| | Fto | -1.13 | 1.883539474 | 0.01307556687 | 6.24E-02 |
| | Rnf214 | -1.13 | 1.739020447 | 0.01823809834 | 7.58E-02 |
| | Hcfc1 | -1.13 | 1.496047941 | 0.03191185567 | 1.06E-01 |
| | D230025D16Rik | -1.13 | 1.618589493 | 0.02406636543 | 8.90E-02 |
| | Clock | -1.13 | 1.402535706 | 0.03957895235 | 1.21E-01 |
| | Kbtbd2 | -1.13 | 1.549211695 | 0.02823503337 | 9.87E-02 |
| | 1700025G04Rik | -1.13 | 1.418332171 | 0.03816522513 | 1.18E-01 |
| | Cdip1 | -1.13 | 1.940709595 | 0.01146279181 | 5.77E-02 |
| | Psmc5 | -1.13 | 1.843668211 | 0.01433282470 | 6.60E-02 |
| | Nudt3 | -1.13 | 1.576661864 | 0.02650563027 | 9.49E-02 |
| | Dlg1 | -1.13 | 1.633652416 | 0.02324596523 | 8.72E-02 |
| | Arl6ip5 | -1.13 | 1.939301457 | 0.01150001859 | 5.78E-02 |
| | Lemd3 | -1.13 | 1.729438884 | 0.01864494539 | 7.65E-02 |
| | St6galnac6 | -1.13 | 1.30162393 | 0.04993166750 | 1.41E-01 |
| | Ltn1 | -1.13 | 1.369793853 | 0.04267820519 | 1.27E-01 |
| | Csnk2b | -1.13 | 1.697274889 | 0.02007821550 | 7.96E-02 |
| | Miga1 | -1.13 | 1.58323974 | 0.02610719782 | 9.39E-02 |
| | Myo9a | -1.13 | 1.321876547 | 0.04765664366 | 1.37E-01 |
| | Vps53 | -1.13 | 1.831865586 | 0.01472768254 | 6.74E-02 |
| | Isca1 | -1.13 | 1.386857952 | 0.04103382934 | 1.24E-01 |
| | Praf2 | -1.14 | 1.416383611 | 0.03833684681 | 1.19E-01 |
| | Nf1 | -1.14 | 1.50717997 | 0.03110427122 | 1.05E-01 |
| | Srcap | -1.14 | 1.731911798 | 0.01853908099 | 7.63E-02 |

| | | | | | |
|---|---|---|---|---|---|
| | Gpr137 | -1.14 | 1.347177648 | 0.04495959099 | 1.32E-01 |
| | Akt3 | -1.14 | 1.366204115 | 0.04303243141 | 1.28E-01 |
| | Fastk | -1.14 | 1.356252851 | 0.04402984425 | 1.30E-01 |
| | Cdc37l1 | -1.14 | 1.376429062 | 0.04203111763 | 1.26E-01 |
| | Eif4enif1 | -1.14 | 1.950285076 | 0.01121282190 | 5.70E-02 |
| | Rhbdd2 | -1.14 | 1.454007536 | 0.03515543404 | 1.13E-01 |
| | Arrb1 | -1.14 | 1.557827935 | 0.02768038109 | 9.77E-02 |
| | Gle1 | -1.14 | 1.74853071 | 0.01784305811 | 7.49E-02 |
| | Parp1 | -1.14 | 1.401318095 | 0.03969007365 | 1.21E-01 |
| | Parp6 | -1.14 | 1.696133435 | 0.02013105635 | 7.97E-02 |
| | Slc25a51 | -1.14 | 1.531402409 | 0.02941694650 | 1.01E-01 |
| | Pkm | -1.14 | 1.451550129 | 0.03535492101 | 1.13E-01 |
| | Gdpd1 | -1.14 | 1.746148786 | 0.01794118870 | 7.51E-02 |
| | Asns | -1.14 | 1.51057592 | 0.03086200090 | 1.04E-01 |
| | Kansl2 | -1.14 | 1.579112218 | 0.02635650271 | 9.45E-02 |
| | Usp5 | -1.14 | 1.490582774 | 0.03231597221 | 1.07E-01 |
| | Gdi1 | -1.14 | 1.815126945 | 0.01530639988 | 6.89E-02 |
| | Ak1 | -1.14 | 2.156945364 | 0.00696714158 | 4.43E-02 |
| | Ppp2r5c | -1.14 | 1.94492788 | 0.01135199313 | 5.74E-02 |
| | Umad1 | -1.14 | 1.463810196 | 0.03437081296 | 1.11E-01 |
| | Rprd2 | -1.14 | 1.569519958 | 0.02694511504 | 9.61E-02 |
| | Dnajb2 | -1.14 | 1.431637725 | 0.03701368079 | 1.16E-01 |
| | Lztr1 | -1.14 | 1.485418299 | 0.03270255621 | 1.08E-01 |

| | | | | | |
|---|---|---|---|---|---|
| | Bcl7b | -1.14 | 1.477838341 | 0.03327834032 | 1.09E-01 |
| | Prpf8 | -1.14 | 1.968112151 | 0.01076187265 | 5.57E-02 |
| | Stoml1 | -1.14 | 1.38844998 | 0.04088368371 | 1.24E-01 |
| | Cxx1c | -1.14 | 2.083600374 | 0.00824896814 | 4.90E-02 |
| | Senp2 | -1.14 | 1.368868127 | 0.04276927348 | 1.27E-01 |
| | Clasp2 | -1.14 | 1.380056127 | 0.04168155116 | 1.25E-01 |
| | Chst10 | -1.14 | 1.513737762 | 0.03063812884 | 1.04E-01 |
| | Pdk2 | -1.14 | 1.480934795 | 0.03304191463 | 1.09E-01 |
| | Asxl1 | -1.14 | 1.599193793 | 0.02516553729 | 9.17E-02 |
| | Gopc | -1.14 | 1.663174824 | 0.02171826741 | 8.33E-02 |
| | Hipk3 | -1.14 | 1.332743217 | 0.04647900076 | 1.35E-01 |
| | Tbpl1 | -1.14 | 1.586487548 | 0.02591268715 | 9.35E-02 |
| | Dctn2 | -1.14 | 1.71114861 | 0.01944694518 | 7.82E-02 |
| | Rbsn | -1.14 | 2.279919542 | 0.00524904696 | 3.82E-02 |
| | Dym | -1.14 | 1.681195705 | 0.02083551766 | 8.15E-02 |
| | Nfx1 | -1.14 | 1.735196127 | 0.01839940899 | 7.61E-02 |
| | Tmub2 | -1.14 | 1.998667534 | 0.01003072828 | 5.38E-02 |
| | Exoc6b | -1.14 | 1.569029688 | 0.02697555023 | 9.61E-02 |
| | Prrc2c | -1.14 | 1.640635229 | 0.02287519319 | 8.61E-02 |
| | Aes | -1.14 | 1.310106858 | 0.04896583243 | 1.39E-01 |
| | Cand1 | -1.14 | 2.198252581 | 0.00633501166 | 4.21E-02 |
| | Dnaja3 | -1.14 | 2.089030301 | 0.00814647443 | 4.86E-02 |
| | Slc35b4 | -1.14 | 1.362836776 | 0.04336738384 | 1.29E-01 |

| | | | | | |
|---|---|---|---|---|---|
| | Ap3m2 | -1.14 | 1.766432321 | 0.01712251989 | 7.33E-02 |
| | Dlst | -1.14 | 1.92018044 | 0.01201765025 | 5.93E-02 |
| | Ndfip1 | -1.14 | 1.545801786 | 0.02845759627 | 9.90E-02 |
| | Usp15 | -1.14 | 1.557083908 | 0.02772784339 | 9.77E-02 |
| | 2410089E03Rik | -1.14 | 1.3392272 | 0.04579022733 | 1.33E-01 |
| | Oxr1 | -1.14 | 1.306211895 | 0.04940695688 | 1.40E-01 |
| | Gga1 | -1.14 | 1.671111069 | 0.02132499465 | 8.24E-02 |
| | Ppp2r5e | -1.14 | 1.531824745 | 0.02938835348 | 1.01E-01 |
| | Ank2 | -1.14 | 1.471653124 | 0.03375568119 | 1.10E-01 |
| | Arfip2 | -1.14 | 1.663290028 | 0.02171250704 | 8.33E-02 |
| | Kcmf1 | -1.14 | 1.597854033 | 0.02524329063 | 9.18E-02 |
| | Wasl | -1.14 | 1.868837467 | 0.01352578665 | 6.35E-02 |
| | 6030458C11Rik | -1.14 | 1.918315084 | 0.01206937874 | 5.94E-02 |
| | Pafah1b1 | -1.14 | 1.357720673 | 0.04388128406 | 1.30E-01 |
| | Ndufa11 | -1.14 | 1.769512525 | 0.01700150921 | 7.31E-02 |
| | Fzr1 | -1.14 | 1.553533483 | 0.02795545187 | 9.81E-02 |
| | Scmh1 | -1.14 | 1.345241277 | 0.04516049805 | 1.32E-01 |
| | Dpcd | -1.14 | 1.552381737 | 0.02802968798 | 9.83E-02 |
| | Nicn1 | -1.14 | 1.715683201 | 0.01924495051 | 7.78E-02 |
| | Fam149b | -1.15 | 1.433183784 | 0.03688214885 | 1.16E-01 |
| | Tars2 | -1.15 | 1.411664721 | 0.03875567269 | 1.19E-01 |
| | Cmtr1 | -1.15 | 1.723519843 | 0.01890079873 | 7.70E-02 |
| | Maneal | -1.15 | 1.432407101 | 0.03694816711 | 1.16E-01 |

| | | | | | |
|---|---|---|---|---|---|
| | Ywhaz | -1.15 | 1.5630339 | 0.02735055224 | 9.70E-02 |
| | Lrrc59 | -1.15 | 2.79757869 | 0.00159375408 | 2.27E-02 |
| | Heatr5b | -1.15 | 1.934271645 | 0.01163398112 | 5.83E-02 |
| | Wipi2 | -1.15 | 2.306117331 | 0.00494177160 | 3.69E-02 |
| | Mtfp1 | -1.15 | 1.421226183 | 0.03791174874 | 1.18E-01 |
| | Prkci | -1.15 | 2.089549953 | 0.00813673265 | 4.85E-02 |
| | Usp7 | -1.15 | 2.418050127 | 0.00381900189 | 3.26E-02 |
| | Slc6a8 | -1.15 | 1.498229674 | 0.03175194445 | 1.06E-01 |
| | Ppp1r12a | -1.15 | 1.539597803 | 0.02886703618 | 9.98E-02 |
| | Crmp1 | -1.15 | 1.432510875 | 0.03693933946 | 1.16E-01 |
| | Ip6k1 | -1.15 | 1.907090162 | 0.01238539431 | 6.04E-02 |
| | Socs5 | -1.15 | 1.781700153 | 0.01653102745 | 7.20E-02 |
| | Fam13b | -1.15 | 1.466236246 | 0.03417934646 | 1.11E-01 |
| | Ddx46 | -1.15 | 1.551027512 | 0.02811722709 | 9.85E-02 |
| | Ctps | -1.15 | 1.667556457 | 0.02150025164 | 8.28E-02 |
| | Pmm1 | -1.15 | 1.719622406 | 0.01907118127 | 7.74E-02 |
| | Snx32 | -1.15 | 1.882528911 | 0.01310602792 | 6.24E-02 |
| | Rad23b | -1.15 | 2.352871903 | 0.00443739507 | 3.47E-02 |
| | Ncaph2 | -1.15 | 2.292331113 | 0.00510115931 | 3.75E-02 |
| | Tpgs2 | -1.15 | 1.776906813 | 0.01671449219 | 7.25E-02 |
| | Cpeb4 | -1.15 | 1.313507476 | 0.04858391677 | 1.39E-01 |
| | Rrp7a | -1.15 | 2.262875613 | 0.00545914194 | 3.90E-02 |
| | Fbxo45 | -1.15 | 1.383929505 | 0.04131145533 | 1.25E-01 |

| | | | | | |
|---|---|---|---|---|---|
| | Atp6v1c1 | -1.15 | 1.515742236 | 0.03049704521 | 1.03E-01 |
| | Prkab2 | -1.15 | 1.417396624 | 0.03824752843 | 1.19E-01 |
| | Akap11 | -1.15 | 1.335305976 | 0.04620553721 | 1.34E-01 |
| | Tufm | -1.15 | 1.60210462 | 0.02499743108 | 9.12E-02 |
| | Nrxn3 | -1.15 | 1.425909312 | 0.03750513108 | 1.17E-01 |
| | Slc25a23 | -1.15 | 1.489715756 | 0.03238055168 | 1.07E-01 |
| | Stk25 | -1.15 | 1.85029085 | 0.01411591875 | 6.53E-02 |
| | Brap | -1.15 | 2.317002064 | 0.00481945507 | 3.64E-02 |
| | Zfp148 | -1.15 | 1.367999248 | 0.04285492627 | 1.28E-01 |
| | Srgap2 | -1.15 | 1.763569854 | 0.01723574840 | 7.35E-02 |
| | Kifap3 | -1.15 | 1.492523231 | 0.03217190434 | 1.07E-01 |
| | Ubr2 | -1.15 | 1.836450793 | 0.01457300812 | 6.68E-02 |
| | Kat7 | -1.15 | 1.687754089 | 0.02052323941 | 8.07E-02 |
| | Nt5m | -1.15 | 1.719336164 | 0.01908375518 | 7.74E-02 |
| | Sestd1 | -1.15 | 1.370374401 | 0.04262119270 | 1.27E-01 |
| | Ap2m1 | -1.15 | 1.932715528 | 0.01167574153 | 5.84E-02 |
| | Hectd3 | -1.15 | 1.987141222 | 0.01030051118 | 5.45E-02 |
| | Brd4 | -1.15 | 1.86396292 | 0.01367845607 | 6.40E-02 |
| | Man1a2 | -1.15 | 1.807124338 | 0.01559106068 | 6.98E-02 |
| | Tmub1 | -1.15 | 1.324505835 | 0.04736899442 | 1.37E-01 |
| | Rnasek | -1.15 | 1.978508155 | 0.01050731724 | 5.49E-02 |
| | Lrsam1 | -1.15 | 1.803435784 | 0.01572404276 | 7.01E-02 |
| | Bysl | -1.15 | 1.543010067 | 0.02864111579 | 9.94E-02 |

| | | | | | |
|---|---|---|---|---|---|
| | Fam49a | -1.15 | 1.459473037 | 0.03471578282 | 1.12E-01 |
| | Ankrd50 | -1.15 | 1.361057678 | 0.04354540374 | 1.29E-01 |
| | Zdhhc3 | -1.15 | 1.705905853 | 0.01968312938 | 7.87E-02 |
| | Rnf123 | -1.15 | 1.699236096 | 0.01998774979 | 7.94E-02 |
| | Pef1 | -1.15 | 1.778941386 | 0.01663637166 | 7.23E-02 |
| | Trappc12 | -1.15 | 1.361770402 | 0.04347399965 | 1.29E-01 |
| | Ocrl | -1.15 | 1.946151809 | 0.01132004599 | 5.73E-02 |
| | Mtmr9 | -1.15 | 1.841646768 | 0.01439969295 | 6.62E-02 |
| | Pja1 | -1.15 | 1.801801285 | 0.01578333285 | 7.02E-02 |
| | Ubtf | -1.15 | 1.690808597 | 0.02037940046 | 8.04E-02 |
| | Adam15 | -1.15 | 1.380479263 | 0.04164096037 | 1.25E-01 |
| | Man2a2 | -1.15 | 2.148775872 | 0.00709944057 | 4.50E-02 |
| | Trim8 | -1.16 | 1.770444926 | 0.01696504726 | 7.30E-02 |
| | Nfe2l1 | -1.16 | 2.248763204 | 0.00563945059 | 3.98E-02 |
| | Zfp651 | -1.16 | 1.362304568 | 0.04342056116 | 1.29E-01 |
| | Zfand2a | -1.16 | 2.500018237 | 0.00316214487 | 2.97E-02 |
| | Zc3h7b | -1.16 | 1.377267139 | 0.04195008649 | 1.26E-01 |
| | Dync1li2 | -1.16 | 1.584303785 | 0.02604331207 | 9.38E-02 |
| | Ncoa7 | -1.16 | 1.417379317 | 0.03824905262 | 1.19E-01 |
| | Ap1ar | -1.16 | 1.483449167 | 0.03285116936 | 1.08E-01 |
| | Xpo6 | -1.16 | 1.530988974 | 0.02944496386 | 1.01E-01 |
| | Gprasp1 | -1.16 | 1.79913053 | 0.01588069372 | 7.04E-02 |
| | Pam | -1.16 | 1.816173545 | 0.01526955763 | 6.88E-02 |

| | | | | | |
|---|---|---|---|---|---|
| | Nsd1 | -1.16 | 2.1667225 41 | 0.00681204 423 | 4.38E-02 |
| | Smcr8 | -1.16 | 2.0167411 6 | 0.00962185 570 | 5.27E-02 |
| | Jakmip2 | -1.16 | 1.5220707 98 | 0.03005586 295 | 1.02E-01 |
| | Ctnna2 | -1.16 | 1.4659662 02 | 0.03420060 571 | 1.11E-01 |
| | Rufy3 | -1.16 | 1.7141472 77 | 0.01931313 263 | 7.79E-02 |
| | Dnajc18 | -1.16 | 2.4091128 8 | 0.00389840 648 | 3.28E-02 |
| | Ddx54 | -1.16 | 1.3087598 69 | 0.04911793 848 | 1.39E-01 |
| | Zwint | -1.16 | 1.3574264 25 | 0.04391102 501 | 1.30E-01 |
| | Dennd4b | -1.16 | 1.3854685 95 | 0.04116531 139 | 1.24E-01 |
| | Ift172 | -1.16 | 1.9877571 05 | 0.01028591 414 | 5.44E-02 |
| | Farsa | -1.16 | 1.4565158 11 | 0.03495297 847 | 1.12E-01 |
| | Ttc19 | -1.16 | 1.8270415 83 | 0.01489218 482 | 6.78E-02 |
| | Rtn2 | -1.16 | 1.6852963 52 | 0.02063971 270 | 8.10E-02 |
| | Idh3b | -1.16 | 2.0656297 11 | 0.00859746 247 | 4.97E-02 |
| | Casp9 | -1.16 | 2.0193299 76 | 0.00956467 075 | 5.26E-02 |
| | Dennd1a | -1.16 | 1.8675605 93 | 0.01356561 250 | 6.36E-02 |
| | Dctn3 | -1.16 | 1.6789450 02 | 0.02094377 667 | 8.18E-02 |
| | Dip2c | -1.16 | 2.0080778 16 | 0.00981572 050 | 5.32E-02 |
| | Mcu | -1.16 | 1.3143159 36 | 0.04849355 958 | 1.39E-01 |
| | Itsn1 | -1.16 | 2.0737605 33 | 0.00843799 895 | 4.94E-02 |
| | Mlf2 | -1.16 | 1.4684432 09 | 0.03400609 712 | 1.10E-01 |
| | B4gat1 | -1.16 | 1.9611125 89 | 0.01093672 798 | 5.63E-02 |
| | Bap1 | -1.16 | 2.0183814 | 0.00958558 450 | 5.26E-02 |

| | | | | | |
|---|---|---|---|---|---|
| | Btbd2 | -1.16 | 2.649382399 | 0.00224190704 | 2.60E-02 |
| | Pex5 | -1.16 | 2.228855957 | 0.00590396865 | 4.08E-02 |
| | Gars | -1.16 | 2.035320692 | 0.00921890432 | 5.17E-02 |
| | Birc6 | -1.16 | 1.698778284 | 0.02000883098 | 7.95E-02 |
| | Tmem199 | -1.16 | 2.037594665 | 0.00917076014 | 5.16E-02 |
| | Nrxn2 | -1.16 | 1.323943749 | 0.04743034148 | 1.37E-01 |
| | Tmem63b | -1.16 | 1.42412702 | 0.03765936391 | 1.17E-01 |
| | Snx15 | -1.16 | 1.434174312 | 0.03679812476 | 1.16E-01 |
| | St3gal2 | -1.16 | 1.796705885 | 0.01596960283 | 7.07E-02 |
| | Carm1 | -1.16 | 1.383825174 | 0.04132138084 | 1.25E-01 |
| | Klhl17 | -1.16 | 1.588574408 | 0.02578847091 | 9.33E-02 |
| | Rere | -1.16 | 1.365919725 | 0.04306061965 | 1.28E-01 |
| | Fntb | -1.16 | 1.500956126 | 0.03155323373 | 1.06E-01 |
| | Mtmr4 | -1.16 | 2.501618616 | 0.00315051378 | 2.97E-02 |
| | Inpp5a | -1.16 | 1.799636721 | 0.01586219479 | 7.04E-02 |
| | Rap2a | -1.16 | 1.672361795 | 0.02126366909 | 8.23E-02 |
| | Ezh1 | -1.16 | 2.011297334 | 0.00974322353 | 5.31E-02 |
| | Ldb1 | -1.16 | 1.309353194 | 0.04905088032 | 1.39E-01 |
| | Emc1 | -1.16 | 2.592491235 | 0.00255569348 | 2.72E-02 |
| | Pim2 | -1.16 | 1.497662209 | 0.03179345983 | 1.06E-01 |
| | Fam155a | -1.16 | 1.478195501 | 0.03325098377 | 1.09E-01 |
| | Gpat4 | -1.16 | 2.551813358 | 0.00280663956 | 2.86E-02 |
| | Tspyl2 | -1.16 | 1.615403945 | 0.02424354111 | 8.94E-02 |

| | | | | | |
|---|---|---|---|---|---|
| | Tbck | -1.16 | 1.840565049 | 0.01443560371 | 6.63E-02 |
| | Sgta | -1.16 | 2.251400975 | 0.00560530212 | 3.96E-02 |
| | Prrc2a | -1.16 | 1.55369363 | 0.02794514515 | 9.81E-02 |
| | Ncoa6 | -1.16 | 1.986864024 | 0.01030708781 | 5.45E-02 |
| | Ptp4a3 | -1.16 | 1.410645207 | 0.03884675919 | 1.19E-01 |
| | Fbxo42 | -1.16 | 1.991206613 | 0.01020453892 | 5.42E-02 |
| | Lsm14b | -1.16 | 2.253364134 | 0.00558002142 | 3.95E-02 |
| | Usp36 | -1.16 | 1.532819212 | 0.02932113572 | 1.01E-01 |
| | Emc4 | -1.16 | 1.744552874 | 0.01800723887 | 7.53E-02 |
| | Wars | -1.16 | 1.78162414 | 0.01653392107 | 7.20E-02 |
| | Arfgef1 | -1.16 | 1.771531048 | 0.01692267260 | 7.29E-02 |
| | Nsg1 | -1.16 | 1.999312052 | 0.01001585315 | 5.38E-02 |
| | Zc3h18 | -1.16 | 1.738531091 | 0.01825866030 | 7.58E-02 |
| | Cramp1l | -1.16 | 1.327774562 | 0.04701380898 | 1.36E-01 |
| | Porcn | -1.16 | 1.857981892 | 0.01386813652 | 6.45E-02 |
| | Stmn1 | -1.16 | 1.606806221 | 0.02472827258 | 9.06E-02 |
| | Hif1an | -1.16 | 1.772871425 | 0.01687052411 | 7.29E-02 |
| | Acot7 | -1.16 | 2.039305597 | 0.00913470240 | 5.16E-02 |
| | Iffo2 | -1.16 | 1.47198592 | 0.03372982441 | 1.10E-01 |
| | Pdxk | -1.16 | 1.815776633 | 0.01528351921 | 6.89E-02 |
| | 8030462N17Rik | -1.16 | 1.486468844 | 0.03262354537 | 1.08E-01 |
| | Bbs1 | -1.17 | 1.364998248 | 0.04315208175 | 1.28E-01 |
| | Wsb2 | -1.17 | 1.384294683 | 0.04127673304 | 1.25E-01 |

| | | | | | |
|---|---|---|---|---|---|
| | Zmynd19 | -1.17 | 1.530590441 | 0.02947199667 | 1.01E-01 |
| | Mtch1 | -1.17 | 1.36070421 | 0.04358085943 | 1.29E-01 |
| | Tmem250-ps | -1.17 | 1.641193032 | 0.02284583144 | 8.61E-02 |
| | Gda | -1.17 | 1.508365562 | 0.03101947463 | 1.04E-01 |
| | Cacna1d | -1.17 | 1.325198076 | 0.04729355100 | 1.36E-01 |
| | Napg | -1.17 | 1.6444316 | 0.02267610191 | 8.56E-02 |
| | Reep1 | -1.17 | 2.009267357 | 0.00978887186 | 5.32E-02 |
| | Dalrd3 | -1.17 | 1.659769634 | 0.02188922401 | 8.37E-02 |
| | Ranbp10 | -1.17 | 1.370588203 | 0.04260021563 | 1.27E-01 |
| | Ttpal | -1.17 | 1.713158499 | 0.01935715384 | 7.80E-02 |
| | Otub1 | -1.17 | 2.420185426 | 0.00380027106 | 3.25E-02 |
| | B3galnt1 | -1.17 | 2.342499292 | 0.00454465278 | 3.53E-02 |
| | 11-Sep | -1.17 | 1.578361406 | 0.02640210744 | 9.46E-02 |
| | Dopey1 | -1.17 | 1.869931712 | 0.01349175008 | 6.35E-02 |
| | Slc35c1 | -1.17 | 1.436271326 | 0.03662087138 | 1.15E-01 |
| | Vps8 | -1.17 | 1.940948861 | 0.01145647836 | 5.77E-02 |
| | Klhl18 | -1.17 | 1.824940694 | 0.01496439991 | 6.81E-02 |
| | Cul9 | -1.17 | 1.905057591 | 0.01244349590 | 6.05E-02 |
| | Bcl2l1 | -1.17 | 1.900830909 | 0.01256519090 | 6.08E-02 |
| | Prcc | -1.17 | 1.476485396 | 0.03338217302 | 1.09E-01 |
| | Pik3cb | -1.17 | 1.624591212 | 0.02373606855 | 8.84E-02 |
| | Ddx19b | -1.17 | 1.486248745 | 0.03264008304 | 1.08E-01 |
| | Trim44 | -1.17 | 2.244228862 | 0.00569863889 | 4.00E-02 |

| | | | | | |
|---|---|---|---|---|---|
| | Bag4 | -1.17 | 2.323381197 | 0.00474918189 | 3.61E-02 |
| | Ankrd17 | -1.17 | 2.664990957 | 0.00216276356 | 2.55E-02 |
| | Ipo4 | -1.17 | 1.765553581 | 0.01715720019 | 7.34E-02 |
| | Smg1 | -1.17 | 1.99346853 | 0.01015152924 | 5.40E-02 |
| | Wdr47 | -1.17 | 1.667322027 | 0.02151186049 | 8.28E-02 |
| | Synm | -1.17 | 1.319951057 | 0.04786840349 | 1.37E-01 |
| | Nacc1 | -1.17 | 2.101074577 | 0.00792365253 | 4.78E-02 |
| | Map1lc3a | -1.17 | 1.618407711 | 0.02407644095 | 8.90E-02 |
| | Ncoa2 | -1.17 | 2.177396769 | 0.00664665644 | 4.32E-02 |
| | Rab35 | -1.17 | 2.491397019 | 0.00322554408 | 3.00E-02 |
| | Ccdc97 | -1.17 | 1.782775417 | 0.01649014913 | 7.19E-02 |
| | Golgb1 | -1.17 | 1.906530382 | 0.01240136867 | 6.04E-02 |
| | Nisch | -1.17 | 1.907678072 | 0.01236863939 | 6.03E-02 |
| | Inpp5e | -1.17 | 1.531782264 | 0.02939122830 | 1.01E-01 |
| | Clk3 | -1.17 | 1.327756159 | 0.04701580118 | 1.36E-01 |
| | Tagln3 | -1.17 | 1.386364832 | 0.04108044774 | 1.24E-01 |
| | Sart3 | -1.17 | 1.738534936 | 0.01825849867 | 7.58E-02 |
| | Nsd2 | -1.17 | 2.095178679 | 0.00803195601 | 4.82E-02 |
| | Mecp2 | -1.17 | 1.853127184 | 0.01402402948 | 6.49E-02 |
| | Nr2c2 | -1.17 | 1.310043381 | 0.04897298987 | 1.39E-01 |
| | Sec61a2 | -1.17 | 1.387704925 | 0.04095388204 | 1.24E-01 |
| | Dock4 | -1.17 | 2.264393623 | 0.00544009366 | 3.90E-02 |
| | Vps33a | -1.17 | 1.877311423 | 0.01326442955 | 6.28E-02 |

| | | | | | |
|---|---|---|---|---|---|
| | Ldoc1l | -1.17 | 1.490269551 | 0.03233928762 | 1.07E-01 |
| | Map2k7 | -1.17 | 2.339555458 | 0.00457556303 | 3.54E-02 |
| | Ppm1a | -1.17 | 2.512985025 | 0.00306912782 | 2.94E-02 |
| | Cers6 | -1.17 | 1.538708489 | 0.02892620840 | 9.99E-02 |
| | Dpp6 | -1.17 | 1.7947 | 0.01604353256 | 7.09E-02 |
| | Dlg2 | -1.17 | 1.540390296 | 0.02881440817 | 9.97E-02 |
| | Polr2l | -1.17 | 1.559849747 | 0.02755181752 | 9.74E-02 |
| | Nudcd3 | -1.17 | 2.110174953 | 0.00775934473 | 4.76E-02 |
| | Gria2 | -1.17 | 1.738161325 | 0.01827421267 | 7.58E-02 |
| | Repin1 | -1.17 | 1.505686577 | 0.03121141247 | 1.05E-01 |
| | Cdk9 | -1.17 | 1.477349854 | 0.03331579233 | 1.09E-01 |
| | Pex1 | -1.17 | 1.52524023 | 0.02983731710 | 1.02E-01 |
| | Ubap2l | -1.17 | 2.384906554 | 0.00412186199 | 3.36E-02 |
| | Zfp385a | -1.17 | 1.640534839 | 0.02288048156 | 8.61E-02 |
| | Ddx55 | -1.17 | 1.348112382 | 0.04486292837 | 1.32E-01 |
| | Zscan26 | -1.17 | 2.399322207 | 0.00398728972 | 3.32E-02 |
| | Bag6 | -1.17 | 2.007238345 | 0.00983471218 | 5.32E-02 |
| | Zfp523 | -1.17 | 1.663792719 | 0.02168738958 | 8.32E-02 |
| | Snn | -1.17 | 1.601803503 | 0.02501476903 | 9.12E-02 |
| | Cxx1b | -1.17 | 2.680240033 | 0.00208814170 | 2.51E-02 |
| | Mtor | -1.17 | 2.238377513 | 0.00577593752 | 4.03E-02 |
| | Elavl4 | -1.17 | 1.459347892 | 0.03472578790 | 1.12E-01 |
| | Letm1 | -1.17 | 2.360608211 | 0.00435904938 | 3.46E-02 |

| | | | | | |
|---|---|---|---|---|---|
| | Sars | -1.17 | 2.217085847 | 0.00606616407 | 4.12E-02 |
| | Rbbp5 | -1.17 | 2.483149605 | 0.00328738368 | 3.01E-02 |
| | Gucy1b3 | -1.17 | 1.383416248 | 0.04136030682 | 1.25E-01 |
| | Smim12 | -1.17 | 1.424168948 | 0.03765572830 | 1.17E-01 |
| | Tmem246 | -1.17 | 1.423253732 | 0.03773516623 | 1.18E-01 |
| | Mex3b | -1.17 | 1.367906514 | 0.04286407792 | 1.28E-01 |
| | Klhl26 | -1.17 | 2.204691404 | 0.00624178199 | 4.18E-02 |
| | Rab3gap2 | -1.17 | 2.871129604 | 0.00134545878 | 2.12E-02 |
| | Slc39a3 | -1.18 | 1.756657237 | 0.01751228283 | 7.41E-02 |
| | Rubcn | -1.18 | 2.545886289 | 0.00284520597 | 2.87E-02 |
| | Mrpl19 | -1.18 | 1.90771092 | 0.01236770392 | 6.03E-02 |
| | Tnrc6c | -1.18 | 1.793564017 | 0.01608555252 | 7.10E-02 |
| | Rapgef1 | -1.18 | 1.717498991 | 0.01916465517 | 7.76E-02 |
| | Actr8 | -1.18 | 2.406031034 | 0.00392616879 | 3.30E-02 |
| | Rtn3 | -1.18 | 1.579107272 | 0.02635680283 | 9.45E-02 |
| | Atg4d | -1.18 | 1.672261257 | 0.02126859213 | 8.23E-02 |
| | Tspan7 | -1.18 | 1.769930996 | 0.01698513505 | 7.30E-02 |
| | Golga2 | -1.18 | 2.35355468 | 0.00443042430 | 3.47E-02 |
| | Gabrb3 | -1.18 | 1.787036086 | 0.01632916260 | 7.16E-02 |
| | Zfp13 | -1.18 | 1.311024246 | 0.04886250795 | 1.39E-01 |
| | Cic | -1.18 | 1.481933227 | 0.03296603940 | 1.08E-01 |
| | Tln2 | -1.18 | 2.429242204 | 0.00372184083 | 3.23E-02 |
| | Aldoa | -1.18 | 1.692800852 | 0.02028612738 | 8.01E-02 |

| | | | | | |
|---|---|---|---|---|---|
| | Vps51 | -1.18 | 1.395754731 | 0.04020177870 | 1.22E-01 |
| | Clasp1 | -1.18 | 1.62266419 | 0.02384162265 | 8.85E-02 |
| | Atp8b2 | -1.18 | 2.009334099 | 0.00978736763 | 5.32E-02 |
| | Dmwd | -1.18 | 1.974282348 | 0.01061005542 | 5.53E-02 |
| | Retreg2 | -1.18 | 2.118227496 | 0.00761679916 | 4.70E-02 |
| | Stx6 | -1.18 | 2.522234136 | 0.00300445611 | 2.93E-02 |
| | Ext1 | -1.18 | 2.01912459 | 0.00956919511 | 5.26E-02 |
| | Nomo1 | -1.18 | 2.219064683 | 0.00603858685 | 4.11E-02 |
| | Agpat1 | -1.18 | 2.083311873 | 0.00825444974 | 4.90E-02 |
| | Slc45a4 | -1.18 | 2.517306928 | 0.00303873671 | 2.94E-02 |
| | Ercc6 | -1.18 | 1.652707269 | 0.02224808992 | 8.46E-02 |
| | Zbtb11 | -1.18 | 1.640505502 | 0.02288202720 | 8.61E-02 |
| | Jazf1 | -1.18 | 1.621300171 | 0.02391662143 | 8.86E-02 |
| | Skiv2l | -1.18 | 2.514780589 | 0.00305646489 | 2.94E-02 |
| | Fbxo44 | -1.18 | 1.385428739 | 0.04116908943 | 1.24E-01 |
| | Fam168a | -1.18 | 1.72680499 | 0.01875836621 | 7.68E-02 |
| | Zswim1 | -1.18 | 1.610030978 | 0.02454533826 | 9.02E-02 |
| | Lyrm9 | -1.18 | 1.678959502 | 0.02094307739 | 8.18E-02 |
| | Narf | -1.18 | 1.622543244 | 0.02384826318 | 8.85E-02 |
| | Selenoi | -1.18 | 2.210346772 | 0.00616102864 | 4.15E-02 |
| | Rapgef2 | -1.18 | 1.884190779 | 0.01305597233 | 6.24E-02 |
| | Tti2 | -1.18 | 1.458187493 | 0.03481869637 | 1.12E-01 |
| | Polr2e | -1.18 | 1.550979774 | 0.02812031790 | 9.85E-02 |

| | | | | | |
|---|---|---|---|---|---|
| | Rab4b | -1.18 | 1.558041516 | 0.02766677152 | 9.77E-02 |
| | Phf8 | -1.18 | 1.322345627 | 0.04760519765 | 1.37E-01 |
| | Rnf41 | -1.18 | 2.253624155 | 0.00557668155 | 3.95E-02 |
| | Bicral | -1.18 | 1.611076311 | 0.02448632948 | 9.01E-02 |
| | Plekha5 | -1.18 | 1.420836481 | 0.03794578304 | 1.18E-01 |
| | Gnao1 | -1.18 | 2.195236311 | 0.00637916285 | 4.23E-02 |
| | Med12l | -1.18 | 1.739970116 | 0.01819826079 | 7.57E-02 |
| | Supv3l1 | -1.18 | 1.666232038 | 0.02156591862 | 8.29E-02 |
| | Ppp1r13b | -1.18 | 2.136568456 | 0.00730182708 | 4.57E-02 |
| | Atp6v0e2 | -1.18 | 2.313610869 | 0.00485723518 | 3.64E-02 |
| | Zbtb4 | -1.18 | 1.763903811 | 0.01722249981 | 7.35E-02 |
| | Scrn1 | -1.18 | 1.391597852 | 0.04058842020 | 1.23E-01 |
| | Srxn1 | -1.18 | 1.720033648 | 0.01905313096 | 7.74E-02 |
| | Dido1 | -1.18 | 1.902164056 | 0.01252667886 | 6.07E-02 |
| | Srrm1 | -1.18 | 1.530745541 | 0.02946147317 | 1.01E-01 |
| | Rnf187 | -1.18 | 2.514728644 | 0.00305683049 | 2.94E-02 |
| | Cdk7 | -1.18 | 1.465210647 | 0.03426015730 | 1.11E-01 |
| | Tspan33 | -1.18 | 1.654252615 | 0.02216906541 | 8.44E-02 |
| | Bicd2 | -1.18 | 2.161125647 | 0.00690040138 | 4.41E-02 |
| | Plxnd1 | -1.18 | 1.416285631 | 0.03834549683 | 1.19E-01 |
| | Klhl22 | -1.18 | 2.539196928 | 0.00288936942 | 2.89E-02 |
| | Tmem109 | -1.18 | 1.386055365 | 0.04110973103 | 1.24E-01 |
| | Arhgef7 | -1.18 | 1.770653303 | 0.01695690930 | 7.30E-02 |

| | | | | | |
|---|---|---|---|---|---|
| | Necap1 | -1.18 | 3.049718492 | 0.00089182883 | 1.85E-02 |
| | Nfyb | -1.18 | 1.903540749 | 0.01248703277 | 6.06E-02 |
| | Ubr3 | -1.18 | 1.315484523 | 0.04836324992 | 1.38E-01 |
| | Flot2 | -1.18 | 1.627693824 | 0.02356710171 | 8.79E-02 |
| | Ylpm1 | -1.18 | 1.899758877 | 0.01259624569 | 6.09E-02 |
| | Mtmr12 | -1.18 | 1.411173393 | 0.03879954271 | 1.19E-01 |
| | Csnk1g2 | -1.18 | 1.358093718 | 0.04384360763 | 1.30E-01 |
| | Snapc3 | -1.18 | 1.455428766 | 0.03504057579 | 1.12E-01 |
| | Gabra3 | -1.18 | 1.417862356 | 0.03820653427 | 1.18E-01 |
| | Dhx30 | -1.18 | 2.557379046 | 0.00277090064 | 2.84E-02 |
| | Cyth2 | -1.18 | 2.019583338 | 0.00955909245 | 5.26E-02 |
| | Zbtb43 | -1.18 | 1.490603083 | 0.03231446108 | 1.07E-01 |
| | Evi5l | -1.18 | 2.231407209 | 0.00586938761 | 4.06E-02 |
| | Rictor | -1.18 | 1.840536443 | 0.01443655457 | 6.63E-02 |
| | Armcx1 | -1.18 | 1.540031488 | 0.02883822408 | 9.97E-02 |
| | Syngr3 | -1.18 | 1.880743496 | 0.01316001863 | 6.26E-02 |
| | Ddx24 | -1.18 | 2.983210171 | 0.00103941703 | 1.97E-02 |
| | Gcc2 | -1.18 | 2.104307121 | 0.00786489409 | 4.77E-02 |
| | Trappc10 | -1.18 | 2.797551799 | 0.00159385277 | 2.27E-02 |
| | Huwe1 | -1.18 | 2.079890465 | 0.00831973579 | 4.91E-02 |
| | Bcl2l2 | -1.18 | 2.081186474 | 0.00829494529 | 4.90E-02 |
| | Ica1 | -1.18 | 1.386871534 | 0.04103254606 | 1.24E-01 |
| | Gpr89 | -1.18 | 1.453028499 | 0.03523477490 | 1.13E-01 |

| | | | | | |
|---|---|---|---|---|---|
| | Larp1 | -1.18 | 2.4890244 98 | 0.00324321 322 | 3.00E-02 |
| | Gnas | -1.18 | 2.2133808 61 | 0.00611813 617 | 4.14E-02 |
| | Tnik | -1.18 | 2.3603486 78 | 0.00436165 512 | 3.46E-02 |
| | Mllt11 | -1.18 | 1.7342695 68 | 0.01843870 566 | 7.62E-02 |
| | Sh3bp5 | -1.18 | 2.0278714 03 | 0.00937839 665 | 5.22E-02 |
| | Slitrk4 | -1.18 | 1.4107612 54 | 0.03883638 041 | 1.19E-01 |
| | Inafm2 | -1.18 | 1.5002184 09 | 0.03160687 733 | 1.06E-01 |
| | Tollip | -1.18 | 2.7075685 73 | 0.00196079 155 | 2.45E-02 |
| | Tmem259 | -1.18 | 1.3150683 69 | 0.04840961 525 | 1.38E-01 |
| | Erc2 | -1.18 | 2.1526069 11 | 0.00703708 974 | 4.47E-02 |
| | Tecr | -1.18 | 2.3140621 85 | 0.00485219 019 | 3.64E-02 |
| | Usp22 | -1.19 | 2.5207797 48 | 0.00301453 446 | 2.93E-02 |
| | Spin1 | -1.19 | 2.8121893 73 | 0.00154102 834 | 2.23E-02 |
| | Rufy2 | -1.19 | 1.6061250 22 | 0.02476708 975 | 9.07E-02 |
| | Gria3 | -1.19 | 2.3598248 22 | 0.00436691 941 | 3.47E-02 |
| | Pomgnt2 | -1.19 | 1.3470018 68 | 0.04497779 205 | 1.32E-01 |
| | Fam169a | -1.19 | 1.7388187 44 | 0.01824657 077 | 7.58E-02 |
| | Kpna6 | -1.19 | 2.8858693 07 | 0.00130056 090 | 2.10E-02 |
| | Stim2 | -1.19 | 1.7431041 68 | 0.01806740 718 | 7.54E-02 |
| | Chst12 | -1.19 | 1.5291579 6 | 0.02956936 783 | 1.01E-01 |
| | Smarcal1 | -1.19 | 2.1858011 63 | 0.00651926 804 | 4.28E-02 |
| | Ap2b1 | -1.19 | 2.3987605 37 | 0.00399244 979 | 3.32E-02 |
| | Mcm3ap | -1.19 | 1.7733638 31 | 0.01685140 706 | 7.29E-02 |

| | | | | | |
|---|---|---|---|---|---|
| | Abhd17a | -1.19 | 1.325044359 | 0.04731029338 | 1.36E-01 |
| | Pdlim7 | -1.19 | 1.417502742 | 0.03823818391 | 1.19E-01 |
| | Gipc1 | -1.19 | 1.410899386 | 0.03882402998 | 1.19E-01 |
| | Smarca4 | -1.19 | 3.016236886 | 0.00096330345 | 1.90E-02 |
| | Cdipt | -1.19 | 2.688972295 | 0.00204657519 | 2.50E-02 |
| | Itga3 | -1.19 | 1.410770717 | 0.03883553412 | 1.19E-01 |
| | Slc6a1 | -1.19 | 1.76753648 | 0.01707904254 | 7.32E-02 |
| | Ralgps1 | -1.19 | 1.364256958 | 0.04322580019 | 1.28E-01 |
| | Phtf2 | -1.19 | 1.774448633 | 0.01680936730 | 7.27E-02 |
| | Sptbn1 | -1.19 | 1.621588459 | 0.02390075063 | 8.86E-02 |
| | Golga4 | -1.19 | 2.057911564 | 0.00875161967 | 5.02E-02 |
| | Klhl12 | -1.19 | 1.96167954 | 0.01092245992 | 5.63E-02 |
| | Sympk | -1.19 | 1.467074916 | 0.03411340605 | 1.11E-01 |
| | Nlgn3 | -1.19 | 1.566510438 | 0.02713248439 | 9.65E-02 |
| | 4932438A13Rik | -1.19 | 1.387029411 | 0.04101763244 | 1.24E-01 |
| | Eya3 | -1.19 | 1.67467446 | 0.02115073870 | 8.21E-02 |
| | Naa25 | -1.19 | 2.075421473 | 0.00840578983 | 4.93E-02 |
| | Slc9a8 | -1.19 | 2.223459597 | 0.00597778654 | 4.09E-02 |
| | Nfasc | -1.19 | 2.335896016 | 0.00461428042 | 3.55E-02 |
| | Zfp280b | -1.19 | 1.367368409 | 0.04291722089 | 1.28E-01 |
| | Lrrc47 | -1.19 | 1.73885601 | 0.01824500516 | 7.58E-02 |
| | Trim32 | -1.19 | 2.499434999 | 0.00316639435 | 2.97E-02 |
| | Foxk2 | -1.19 | 2.378746309 | 0.00418074512 | 3.39E-02 |

| | | | | | |
|---|---|---|---|---|---|
| | Mink1 | -1.19 | 1.955273417 | 0.01108476735 | 5.67E-02 |
| | Agk | -1.19 | 2.124605167 | 0.00750576275 | 4.64E-02 |
| | Zfp574 | -1.19 | 2.166004481 | 0.00682331654 | 4.39E-02 |
| | Smap2 | -1.19 | 2.024313576 | 0.00945554191 | 5.24E-02 |
| | Cep85 | -1.19 | 1.472584453 | 0.03368337082 | 1.10E-01 |
| | Gbf1 | -1.19 | 3.382084954 | 0.00041487288 | 1.57E-02 |
| | Shc2 | -1.19 | 1.392400822 | 0.04051344536 | 1.23E-01 |
| | Ddx56 | -1.19 | 1.56469066 | 0.02724641327 | 9.67E-02 |
| | Got2 | -1.19 | 2.226876884 | 0.00593093433 | 4.08E-02 |
| | Rmnd5a | -1.19 | 1.717408645 | 0.01916864237 | 7.76E-02 |
| | Nanp | -1.19 | 1.588903813 | 0.02576891820 | 9.33E-02 |
| | Ncor1 | -1.19 | 2.723227872 | 0.00189135098 | 2.42E-02 |
| | Hirip3 | -1.19 | 1.30228482 | 0.04985574152 | 1.41E-01 |
| | Gnl1 | -1.19 | 1.709552635 | 0.01951854164 | 7.84E-02 |
| | Fam234b | -1.19 | 2.471356318 | 0.00337787584 | 3.06E-02 |
| | Marf1 | -1.19 | 1.877872536 | 0.01324730283 | 6.28E-02 |
| | Brwd1 | -1.19 | 2.862698073 | 0.00137183515 | 2.13E-02 |
| | Slc4a10 | -1.19 | 1.583510822 | 0.02609090707 | 9.39E-02 |
| | Vps13d | -1.19 | 1.830467427 | 0.01477517296 | 6.75E-02 |
| | Chd8 | -1.19 | 3.184375025 | 0.00065407112 | 1.66E-02 |
| | Abca2 | -1.19 | 1.854745953 | 0.01397185426 | 6.48E-02 |
| | Ube3b | -1.19 | 2.266887923 | 0.00540893892 | 3.89E-02 |
| | Slc6a15 | -1.19 | 1.58674361 | 0.02589741341 | 9.35E-02 |

| | | | | | |
|---|---|---|---|---|---|
| | Atg16l1 | -1.19 | 1.70374749 | 0.01978119434 | 7.90E-02 |
| | Fam212b | -1.19 | 1.740663442 | 0.01816923146 | 7.56E-02 |
| | Ciz1 | -1.19 | 1.339811111 | 0.04572870355 | 1.33E-01 |
| | Crebl2 | -1.19 | 1.762986071 | 0.01725893246 | 7.36E-02 |
| | Hdgfl2 | -1.19 | 1.378789402 | 0.04180330303 | 1.26E-01 |
| | Elac2 | -1.19 | 1.880342794 | 0.01317216632 | 6.26E-02 |
| | Cog1 | -1.19 | 1.70607046 | 0.01967567047 | 7.87E-02 |
| | 3-Sep | -1.19 | 1.995651006 | 0.01010064237 | 5.39E-02 |
| | Stub1 | -1.19 | 1.744127178 | 0.01802489826 | 7.54E-02 |
| | Cdk5 | -1.19 | 1.929747638 | 0.01175580469 | 5.86E-02 |
| | Serac1 | -1.19 | 1.624527764 | 0.02373953653 | 8.84E-02 |
| | Sipa1l3 | -1.19 | 1.313354647 | 0.04860101648 | 1.39E-01 |
| | Ddx51 | -1.19 | 1.330475577 | 0.04672232249 | 1.35E-01 |
| | Tbc1d22b | -1.19 | 2.108908199 | 0.00778201029 | 4.77E-02 |
| | Sec14l1 | -1.19 | 2.216205056 | 0.00607847932 | 4.12E-02 |
| | Ptk2b | -1.19 | 1.957284867 | 0.01103354658 | 5.65E-02 |
| | Tbc1d10b | -1.19 | 1.878904415 | 0.01321586474 | 6.27E-02 |
| | Zbtb34 | -1.19 | 1.408358678 | 0.03905182391 | 1.20E-01 |
| | Prkcz | -1.19 | 1.852302657 | 0.01405068001 | 6.50E-02 |
| | Ppp1r9b | -1.19 | 1.49466139 | 0.03201390188 | 1.06E-01 |
| | Atg2a | -1.19 | 1.884854697 | 0.01303602856 | 6.23E-02 |
| | Pwp2 | -1.19 | 1.714463946 | 0.01929905547 | 7.79E-02 |
| | Irf2bp1 | -1.19 | 1.492899763 | 0.03214402353 | 1.07E-01 |

| | | | | | |
|---|---|---|---|---|---|
| | Pagr1a | -1.19 | 1.516684636 | 0.03043093974 | 1.03E-01 |
| | Jph4 | -1.19 | 1.673926325 | 0.02118720531 | 8.21E-02 |
| | Brpf1 | -1.19 | 1.837056989 | 0.01455268104 | 6.68E-02 |
| | Ankrd34b | -1.19 | 1.483809097 | 0.03282395455 | 1.08E-01 |
| | Smyd5 | -1.19 | 2.016311335 | 0.00963138326 | 5.27E-02 |
| | Zfp507 | -1.19 | 1.543452292 | 0.02861196650 | 9.94E-02 |
| | Tsc2 | -1.19 | 2.239022349 | 0.00576736784 | 4.03E-02 |
| | Dusp3 | -1.19 | 1.468000889 | 0.03404074928 | 1.10E-01 |
| | Adarb1 | -1.19 | 1.660771819 | 0.02183877032 | 8.36E-02 |
| | Lrp11 | -1.20 | 1.74344891 | 0.01805307099 | 7.54E-02 |
| | Slc12a6 | -1.20 | 2.119790125 | 0.00758944250 | 4.69E-02 |
| | Jmjd4 | -1.20 | 1.475863517 | 0.03343000816 | 1.09E-01 |
| | Cables1 | -1.20 | 1.342466586 | 0.04544995046 | 1.33E-01 |
| | Polr3a | -1.20 | 1.80068132 | 0.01582408764 | 7.03E-02 |
| | Cyb561d1 | -1.20 | 2.161229687 | 0.00689874851 | 4.41E-02 |
| | Cds2 | -1.20 | 2.648585702 | 0.00224602351 | 2.60E-02 |
| | Senp5 | -1.20 | 2.16158214 | 0.00689315207 | 4.41E-02 |
| | Cep104 | -1.20 | 1.78063722 | 0.01657153659 | 7.21E-02 |
| | Nudt18 | -1.20 | 1.54855212 | 0.02827794726 | 9.87E-02 |
| | Myl12b | -1.20 | 2.910755111 | 0.00122813155 | 2.05E-02 |
| | Sgsm3 | -1.20 | 2.550461016 | 0.00281539273 | 2.86E-02 |
| | Large1 | -1.20 | 1.657500344 | 0.02200389964 | 8.40E-02 |
| | Gpd2 | -1.20 | 1.927163035 | 0.01182597523 | 5.89E-02 |

| | | | | | |
|---|---|---|---|---|---|
| | Cdkl2 | -1.20 | 1.964524175 | 0.01085115146 | 5.61E-02 |
| | Napb | -1.20 | 2.175278552 | 0.00667915386 | 4.34E-02 |
| | Car11 | -1.20 | 1.472318408 | 0.03370401131 | 1.10E-01 |
| | Nup214 | -1.20 | 2.103856314 | 0.00787306227 | 4.77E-02 |
| | Srpk2 | -1.20 | 1.872243792 | 0.01342011410 | 6.32E-02 |
| | Atp6v0d1 | -1.20 | 3.016430188 | 0.00096287478 | 1.90E-02 |
| | Ankrd11 | -1.20 | 2.487574231 | 0.00325406159 | 3.01E-02 |
| | Kctd2 | -1.20 | 1.6090518 | 0.02460074163 | 9.03E-02 |
| | Zfp609 | -1.20 | 2.001931547 | 0.00995562325 | 5.37E-02 |
| | Cdan1 | -1.20 | 2.008322966 | 0.00981018130 | 5.32E-02 |
| | Tmem11 | -1.20 | 1.89485209 | 0.01273936878 | 6.14E-02 |
| | Trak1 | -1.20 | 2.032523494 | 0.00927847295 | 5.19E-02 |
| | Hsph1 | -1.20 | 2.005973524 | 0.00986339614 | 5.34E-02 |
| | Grm7 | -1.20 | 1.950409351 | 0.01120961377 | 5.70E-02 |
| | Phactr1 | -1.20 | 1.82367273 | 0.01500815375 | 6.81E-02 |
| | Ogfrl1 | -1.20 | 1.373240813 | 0.04234081245 | 1.27E-01 |
| | Pdxp | -1.20 | 2.498868455 | 0.00317052765 | 2.97E-02 |
| | Cplx2 | -1.20 | 1.628249976 | 0.02353694127 | 8.78E-02 |
| | Gpatch8 | -1.20 | 1.804281196 | 0.01569346358 | 7.01E-02 |
| | Mtmr7 | -1.20 | 1.713218985 | 0.01935445806 | 7.80E-02 |
| | Kansl1 | -1.20 | 2.933345302 | 0.00116588227 | 2.03E-02 |
| | Sh2b1 | -1.20 | 2.203837429 | 0.00625406760 | 4.18E-02 |
| | Podxl2 | -1.20 | 1.536585976 | 0.02906792455 | 1.00E-01 |

| | | | | | |
|---|---|---|---|---|---|
| | Hdac6 | -1.20 | 1.481119181 | 0.03302788917 | 1.09E-01 |
| | Slc7a6 | -1.20 | 1.712564908 | 0.01938362912 | 7.81E-02 |
| | Kpna1 | -1.20 | 2.352747879 | 0.00443866247 | 3.47E-02 |
| | Cluh | -1.20 | 2.975274489 | 0.00105858445 | 1.97E-02 |
| | Gtf3c1 | -1.20 | 2.207686575 | 0.00619888279 | 4.17E-02 |
| | 2210016L21Rik | -1.20 | 2.008096801 | 0.00981529143 | 5.32E-02 |
| | Dnmt3a | -1.20 | 2.225864411 | 0.00594477729 | 4.08E-02 |
| | Glce | -1.20 | 1.7068723 | 0.01963937667 | 7.87E-02 |
| | Lancl1 | -1.20 | 1.643947015 | 0.02270141798 | 8.57E-02 |
| | Cops7a | -1.20 | 2.830444889 | 0.00147759397 | 2.19E-02 |
| | Mcf2l | -1.20 | 2.784099361 | 0.00164399555 | 2.30E-02 |
| | Adcy3 | -1.20 | 2.263939799 | 0.00544578136 | 3.90E-02 |
| | Ubfd1 | -1.20 | 3.030779026 | 0.00093158175 | 1.89E-02 |
| | Dync1i1 | -1.20 | 1.881654896 | 0.01313243028 | 6.25E-02 |
| | Dapk1 | -1.20 | 1.633023466 | 0.02327965467 | 8.73E-02 |
| | Klf13 | -1.20 | 2.039702893 | 0.00912634972 | 5.16E-02 |
| | Tmem201 | -1.20 | 1.595239538 | 0.02539571602 | 9.23E-02 |
| | Copg1 | -1.20 | 2.618868815 | 0.00240508918 | 2.66E-02 |
| | Mxd1 | -1.20 | 1.807309052 | 0.01558443090 | 6.98E-02 |
| | Eml2 | -1.20 | 1.536682038 | 0.02906149568 | 1.00E-01 |
| | Exoc1 | -1.20 | 2.587019757 | 0.00258809518 | 2.74E-02 |
| | Sun1 | -1.20 | 1.355826413 | 0.04407309880 | 1.30E-01 |
| | Rbm15b | -1.20 | 1.857773629 | 0.01387478848 | 6.45E-02 |

| | | | | | |
|---|---|---|---|---|---|
| | Larp6 | -1.20 | 1.36543706 | 0.04310850280 | 1.28E-01 |
| | Syvn1 | -1.20 | 1.496738082 | 0.03186118454 | 1.06E-01 |
| | Sarm1 | -1.20 | 2.01811901 | 0.00959137762 | 5.26E-02 |
| | Ncs1 | -1.20 | 1.402286962 | 0.03960162779 | 1.21E-01 |
| | Rundc1 | -1.20 | 1.918159571 | 0.01207370136 | 5.94E-02 |
| | Stmn2 | -1.20 | 1.546945783 | 0.02838273333 | 9.89E-02 |
| | 1810055G02Rik | -1.20 | 1.715441405 | 0.01925566825 | 7.78E-02 |
| | Fam160b1 | -1.20 | 2.034712739 | 0.00923181858 | 5.18E-02 |
| | Hgsnat | -1.20 | 1.352205297 | 0.04444211334 | 1.31E-01 |
| | Kif1b | -1.20 | 1.73607183 | 0.01836234615 | 7.60E-02 |
| | Gdap1l1 | -1.20 | 2.164483943 | 0.00684724799 | 4.39E-02 |
| | Gtpbp2 | -1.20 | 2.062678288 | 0.00865608898 | 4.99E-02 |
| | Iglon5 | -1.20 | 1.701647042 | 0.01987709702 | 7.92E-02 |
| | Tmem38a | -1.20 | 2.296832298 | 0.00504856209 | 3.73E-02 |
| | Slc25a44 | -1.20 | 2.632724861 | 0.00232956664 | 2.62E-02 |
| | Fkrp | -1.20 | 2.138524735 | 0.00726900997 | 4.56E-02 |
| | Rnf220 | -1.20 | 2.372312331 | 0.00424314301 | 3.41E-02 |
| | Nup133 | -1.20 | 1.789721785 | 0.01622849385 | 7.13E-02 |
| | Tuba4a | -1.20 | 1.583648273 | 0.02608265079 | 9.38E-02 |
| | Gak | -1.20 | 2.275820874 | 0.00529881950 | 3.84E-02 |
| | Pi4k2a | -1.20 | 3.233221157 | 0.00058449236 | 1.65E-02 |
| | Pkd1 | -1.20 | 1.414852454 | 0.03847224641 | 1.19E-01 |
| | Trappc9 | -1.20 | 2.549614496 | 0.00282088580 | 2.86E-02 |

| | | | | | |
|---|---|---|---|---|---|
| | Snca | -1.20 | 1.405619254 | 0.03929893177 | 1.20E-01 |
| | Dusp8 | -1.20 | 2.52647402 | 0.00297526724 | 2.92E-02 |
| | Znfx1 | -1.21 | 1.749303727 | 0.01781132682 | 7.48E-02 |
| | Gramd1a | -1.21 | 2.020830634 | 0.00953167807 | 5.25E-02 |
| | Nacad | -1.21 | 1.800465236 | 0.01583196293 | 7.03E-02 |
| | Smarca2 | -1.21 | 1.717619229 | 0.01915934998 | 7.76E-02 |
| | Eml5 | -1.21 | 1.347527897 | 0.04492334669 | 1.32E-01 |
| | Dpp8 | -1.21 | 1.822102336 | 0.01506252096 | 6.83E-02 |
| | Ube3c | -1.21 | 2.918282338 | 0.00120702888 | 2.04E-02 |
| | Hspa4l | -1.21 | 1.833336456 | 0.01467788715 | 6.73E-02 |
| | Ptgfrn | -1.21 | 1.358737813 | 0.04377863200 | 1.29E-01 |
| | Gabarapl1 | -1.21 | 2.461673714 | 0.00345403144 | 3.10E-02 |
| | Chp1 | -1.21 | 2.801516377 | 0.00157936905 | 2.26E-02 |
| | Slc35e2 | -1.21 | 2.385949385 | 0.00411197642 | 3.36E-02 |
| | Fn3krp | -1.21 | 1.412391497 | 0.03869087065 | 1.19E-01 |
| | Pikfyve | -1.21 | 1.578689699 | 0.02638215703 | 9.46E-02 |
| | Tbrg4 | -1.21 | 1.523817205 | 0.02993524348 | 1.02E-01 |
| | Tspyl3 | -1.21 | 2.067634224 | 0.00855787179 | 4.96E-02 |
| | Dclk1 | -1.21 | 2.32390728 | 0.00474343245 | 3.61E-02 |
| | Malat1 | -1.21 | 1.507457106 | 0.03108442901 | 1.05E-01 |
| | Khdrbs3 | -1.21 | 1.430728637 | 0.03709124087 | 1.16E-01 |
| | Dpysl4 | -1.21 | 1.91074636 | 0.01228156299 | 6.00E-02 |
| | Tomm34 | -1.21 | 1.995438996 | 0.01010557440 | 5.39E-02 |

| | | | | | |
|---|---|---|---|---|---|
| | Pip5k1a | -1.21 | 2.410124696 | 0.00388933457 | 3.28E-02 |
| | Vps52 | -1.21 | 2.160389351 | 0.00691211013 | 4.41E-02 |
| | Jcad | -1.21 | 1.673583029 | 0.02120395976 | 8.22E-02 |
| | Spns1 | -1.21 | 1.722674922 | 0.01893760606 | 7.71E-02 |
| | Camsap2 | -1.21 | 2.64199112 | 0.00228038870 | 2.61E-02 |
| | 1700003E16Rik | -1.21 | 1.423851889 | 0.03768322914 | 1.17E-01 |
| | Dync1li1 | -1.21 | 2.554907123 | 0.00278671706 | 2.85E-02 |
| | Ablim2 | -1.21 | 1.414890256 | 0.03846889785 | 1.19E-01 |
| | Nkrf | -1.21 | 2.314193796 | 0.00485071998 | 3.64E-02 |
| | Actn4 | -1.21 | 1.432129556 | 0.03697178718 | 1.16E-01 |
| | Trim9 | -1.21 | 1.585885385 | 0.02594864082 | 9.36E-02 |
| | Pitpna | -1.21 | 2.549845983 | 0.00281938261 | 2.86E-02 |
| | Wasf1 | -1.21 | 2.758689541 | 0.00174305246 | 2.34E-02 |
| | Ing2 | -1.21 | 1.538493833 | 0.02894050915 | 9.99E-02 |
| | Rxrb | -1.21 | 2.007489098 | 0.00982903546 | 5.32E-02 |
| | Snph | -1.21 | 1.420039858 | 0.03801545054 | 1.18E-01 |
| | Slc2a3 | -1.21 | 2.035084653 | 0.00922391617 | 5.17E-02 |
| | Diras1 | -1.21 | 2.194889526 | 0.00638425865 | 4.23E-02 |
| | Dzank1 | -1.21 | 2.319993691 | 0.00478637045 | 3.63E-02 |
| | Atp1b1 | -1.21 | 1.542121147 | 0.02869979886 | 9.95E-02 |
| | Exd2 | -1.21 | 2.418137083 | 0.00381823731 | 3.26E-02 |
| | Rpgrip1l | -1.21 | 1.639738634 | 0.02292246751 | 8.62E-02 |
| | Smarcd1 | -1.21 | 1.980706386 | 0.01045426764 | 5.47E-02 |

| | | | | | |
|---|---|---|---|---|---|
| | Rabgef1 | -1.21 | 2.134036642 | 0.00734451899 | 4.59E-02 |
| | Clip3 | -1.21 | 2.534141559 | 0.00292319940 | 2.89E-02 |
| | Cbfa2t2 | -1.21 | 3.142789113 | 0.00071979842 | 1.71E-02 |
| | Slc24a3 | -1.21 | 2.48692046 | 0.00325896383 | 3.01E-02 |
| | Sh2d3c | -1.21 | 1.36343981 | 0.04330720845 | 1.28E-01 |
| | Fam81a | -1.21 | 1.877724836 | 0.01325180887 | 6.28E-02 |
| | Dip2a | -1.21 | 1.79022431 | 0.01620972661 | 7.13E-02 |
| | Rnf169 | -1.21 | 1.839721728 | 0.01446366226 | 6.64E-02 |
| | Katnal1 | -1.21 | 2.730080802 | 0.00186174072 | 2.40E-02 |
| | Prr12 | -1.21 | 2.067606631 | 0.00855841553 | 4.96E-02 |
| | Ythdc2 | -1.21 | 1.70989812 | 0.01950302063 | 7.84E-02 |
| | Tmem8b | -1.21 | 2.671035833 | 0.00213286893 | 2.54E-02 |
| | Gnb1 | -1.21 | 3.102238079 | 0.00079024530 | 1.78E-02 |
| | Map7d1 | -1.21 | 1.883403181 | 0.01307967097 | 6.24E-02 |
| | Magee1 | -1.21 | 2.978223685 | 0.00105142020 | 1.97E-02 |
| | Schip1 | -1.21 | 1.554735089 | 0.02787821166 | 9.80E-02 |
| | Safb | -1.21 | 1.627052429 | 0.02360193288 | 8.80E-02 |
| | Srf | -1.21 | 1.368962162 | 0.04276001388 | 1.27E-01 |
| | Tnks | -1.21 | 2.10488413 | 0.00785445163 | 4.77E-02 |
| | Pcnx3 | -1.21 | 1.37942061 | 0.04174258979 | 1.25E-01 |
| | Exoc7 | -1.21 | 2.700020767 | 0.00199516691 | 2.47E-02 |
| | Dnajc27 | -1.21 | 2.106004566 | 0.00783421407 | 4.77E-02 |
| | Ldlrad4 | -1.21 | 1.737689779 | 0.01829406511 | 7.58E-02 |

| | | | | | |
|---|---|---|---|---|---|
| | Asb1 | -1.21 | 1.722342397 | 0.01895211150 | 7.71E-02 |
| | Lgi3 | -1.21 | 1.312211527 | 0.04872910931 | 1.39E-01 |
| | Arhgef18 | -1.22 | 2.374952769 | 0.00421742367 | 3.40E-02 |
| | Cxxc5 | -1.22 | 2.833738095 | 0.00146643192 | 2.19E-02 |
| | Pigo | -1.22 | 1.674214056 | 0.02117317292 | 8.21E-02 |
| | Tro | -1.22 | 2.61624705 | 0.00241965223 | 2.66E-02 |
| | Pcnx | -1.22 | 3.191901314 | 0.00064283377 | 1.65E-02 |
| | 4933427D14Rik | -1.22 | 1.998687682 | 0.01003026295 | 5.38E-02 |
| | Usp19 | -1.22 | 2.125813819 | 0.00748490308 | 4.64E-02 |
| | Cxx1a | -1.22 | 1.683709929 | 0.02071524487 | 8.12E-02 |
| | Rnf44 | -1.22 | 2.630262821 | 0.00234281059 | 2.63E-02 |
| | Tet3 | -1.22 | 1.559790925 | 0.02755554943 | 9.74E-02 |
| | Slc27a4 | -1.22 | 2.389305263 | 0.00408032482 | 3.36E-02 |
| | Ap3d1 | -1.22 | 2.713280855 | 0.00193517010 | 2.43E-02 |
| | Ap2a2 | -1.22 | 2.176555099 | 0.00665955027 | 4.33E-02 |
| | Lrrc20 | -1.22 | 1.311224036 | 0.04884003474 | 1.39E-01 |
| | Magi3 | -1.22 | 1.64151069 | 0.02282912733 | 8.61E-02 |
| | 2900011O08Rik | -1.22 | 2.618352374 | 0.00240795090 | 2.66E-02 |
| | Pnma2 | -1.22 | 1.702622425 | 0.01983250513 | 7.91E-02 |
| | Mier2 | -1.22 | 1.463823114 | 0.03436979060 | 1.11E-01 |
| | Camk2b | -1.22 | 2.648998604 | 0.00224388914 | 2.60E-02 |
| | Chchd10 | -1.22 | 1.504587168 | 0.03129052372 | 1.05E-01 |
| | Amph | -1.22 | 1.960637617 | 0.01094869564 | 5.63E-02 |

| | | | | | |
|---|---|---|---|---|---|
| | Dohh | -1.22 | 1.54185707 | 0.02871725533 | 9.95E-02 |
| | Pik3r2 | -1.22 | 1.693838546 | 0.02023771397 | 8.00E-02 |
| | Ica1l | -1.22 | 2.219952121 | 0.00602626019 | 4.10E-02 |
| | Serp2 | -1.22 | 1.378806586 | 0.04180164902 | 1.26E-01 |
| | Mpp3 | -1.22 | 2.054862852 | 0.00881327149 | 5.03E-02 |
| | Icmt | -1.22 | 2.73164707 | 0.00185503851 | 2.40E-02 |
| | Ogdhl | -1.22 | 1.462700593 | 0.03445874110 | 1.11E-01 |
| | Vopp1 | -1.22 | 2.508307061 | 0.00310236533 | 2.96E-02 |
| | Galt | -1.22 | 2.082373216 | 0.00827230967 | 4.90E-02 |
| | Extl3 | -1.22 | 2.607129005 | 0.00247099004 | 2.69E-02 |
| | Meaf6 | -1.22 | 1.752212926 | 0.01769241320 | 7.45E-02 |
| | Oprl1 | -1.22 | 1.498654807 | 0.03172087752 | 1.06E-01 |
| | Rnf26 | -1.22 | 2.199427246 | 0.00631790009 | 4.21E-02 |
| | Pdpk1 | -1.22 | 3.209128902 | 0.00061783299 | 1.65E-02 |
| | Slc23a2 | -1.22 | 2.094334589 | 0.00804758201 | 4.82E-02 |
| | Lgi1 | -1.22 | 1.539059801 | 0.02890281872 | 9.99E-02 |
| | Upf1 | -1.22 | 2.048368007 | 0.00894606384 | 5.09E-02 |
| | Fam69b | -1.22 | 2.316595792 | 0.00482396566 | 3.64E-02 |
| | Rnft2 | -1.22 | 3.463796776 | 0.00034371875 | 1.49E-02 |
| | Zfp385b | -1.22 | 2.072440722 | 0.00846368083 | 4.94E-02 |
| | Rrp12 | -1.22 | 1.980564487 | 0.01045768396 | 5.47E-02 |
| | Rab6a | -1.22 | 1.602596245 | 0.02496914978 | 9.12E-02 |
| | Zfp251 | -1.22 | 2.808021943 | 0.00155588702 | 2.25E-02 |

| | | | | | |
|---|---|---|---|---|---|
| | Appbp2 | -1.22 | 2.0974255 57 | 0.00799050 896 | 4.80E-02 |
| | Ube4b | -1.22 | 3.5783180 4 | 0.00026404 744 | 1.42E-02 |
| | Prkaca | -1.22 | 4.0594128 69 | 0.00008721 419 | 1.41E-02 |
| | Prkce | -1.22 | 2.4319286 23 | 0.00369888 967 | 3.22E-02 |
| | Neurl4 | -1.22 | 2.3941854 37 | 0.00403473 080 | 3.34E-02 |
| | Scaf1 | -1.22 | 1.8299401 57 | 0.01479312 213 | 6.75E-02 |
| | Elovl4 | -1.22 | 2.0241587 28 | 0.00945891 389 | 5.24E-02 |
| | Csrnp3 | -1.22 | 1.8441571 15 | 0.01431669 868 | 6.59E-02 |
| | Cntn4 | -1.22 | 1.3896359 88 | 0.04077218 747 | 1.24E-01 |
| | Gsk3b | -1.22 | 2.4056949 33 | 0.00392920 842 | 3.30E-02 |
| | Plcg1 | -1.22 | 2.8601015 04 | 0.00138006 168 | 2.15E-02 |
| | Mark2 | -1.22 | 2.7496623 52 | 0.00177966 250 | 2.36E-02 |
| | Nol4l | -1.22 | 1.7052829 99 | 0.01971137 869 | 7.87E-02 |
| | Igf1r | -1.22 | 2.6053162 78 | 0.00248132 541 | 2.69E-02 |
| | Slc20a1 | -1.22 | 2.8052154 19 | 0.00156597 412 | 2.25E-02 |
| | Atxn1 | -1.22 | 1.9138442 55 | 0.01219426 826 | 5.98E-02 |
| | Hgs | -1.22 | 2.2994308 46 | 0.00501844 483 | 3.72E-02 |
| | Abtb1 | -1.22 | 1.9210530 43 | 0.01199352 809 | 5.93E-02 |
| | Kctd1 | -1.22 | 2.5456160 61 | 0.00284697 687 | 2.87E-02 |
| | Cntnap2 | -1.22 | 1.6221834 76 | 0.02386802 721 | 8.86E-02 |
| | Fxr2 | -1.22 | 2.8441155 44 | 0.00143180 692 | 2.17E-02 |
| | Rptor | -1.22 | 3.2392873 99 | 0.00057638 491 | 1.64E-02 |
| | Flrt2 | -1.22 | 1.7898150 31 | 0.01622500 985 | 7.13E-02 |

| | | | | | |
|---|---|---|---|---|---|
| | Smarcc2 | -1.22 | 2.785018838 | 0.00164051861 | 2.29E-02 |
| | L3mbtl2 | -1.22 | 2.774006006 | 0.00168265079 | 2.32E-02 |
| | Fam168b | -1.22 | 2.316798567 | 0.00482171385 | 3.64E-02 |
| | 6430548M08Rik | -1.22 | 2.369458019 | 0.00427112204 | 3.42E-02 |
| | Stmn3 | -1.22 | 2.369883919 | 0.00426693553 | 3.42E-02 |
| | Pip4k2b | -1.22 | 3.61493681 | 0.00024269632 | 1.42E-02 |
| | Zfyve9 | -1.22 | 1.829419631 | 0.01481086315 | 6.75E-02 |
| | Katnb1 | -1.22 | 3.305273734 | 0.00049513801 | 1.59E-02 |
| | Clcn6 | -1.22 | 2.604960956 | 0.00248335636 | 2.69E-02 |
| | Tenm4 | -1.22 | 1.796930367 | 0.01596135043 | 7.07E-02 |
| | Arid1b | -1.23 | 3.540359072 | 0.00028816480 | 1.43E-02 |
| | Epb41l3 | -1.23 | 2.165632804 | 0.00682915856 | 4.39E-02 |
| | Depdc5 | -1.23 | 2.239946344 | 0.00575511036 | 4.03E-02 |
| | Cadps | -1.23 | 1.741103596 | 0.01815082646 | 7.56E-02 |
| | Lrrtm3 | -1.23 | 2.056935461 | 0.00877131158 | 5.02E-02 |
| | Rasal2 | -1.23 | 2.41046547 | 0.00388628396 | 3.28E-02 |
| | Tppp | -1.23 | 1.860284788 | 0.01379479376 | 6.43E-02 |
| | Abr | -1.23 | 2.384831868 | 0.00412257089 | 3.36E-02 |
| | Elavl3 | -1.23 | 2.926702157 | 0.00118385317 | 2.03E-02 |
| | Adar | -1.23 | 3.240542123 | 0.00057472207 | 1.64E-02 |
| | Pja2 | -1.23 | 1.982692146 | 0.01040657585 | 5.47E-02 |
| | Fbxo21 | -1.23 | 2.228814838 | 0.00590452767 | 4.08E-02 |
| | Tmem169 | -1.23 | 1.942786873 | 0.01140809494 | 5.75E-02 |

| | | | | | |
|---|---|---|---|---|---|
| | Zfp598 | -1.23 | 1.475891077 | 0.03342788685 | 1.09E-01 |
| | Zmat3 | -1.23 | 2.144113148 | 0.00717607307 | 4.53E-02 |
| | Oaz1-ps | -1.23 | 1.476292655 | 0.03339699145 | 1.09E-01 |
| | Bahd1 | -1.23 | 2.156786603 | 0.00696968896 | 4.43E-02 |
| | Epn1 | -1.23 | 1.749882077 | 0.01778762326 | 7.47E-02 |
| | Jakmip3 | -1.23 | 1.95139496 | 0.01118420298 | 5.70E-02 |
| | Hs6st1 | -1.23 | 2.089936834 | 0.00812948747 | 4.85E-02 |
| | Pcgf3 | -1.23 | 2.44871981 | 0.00355860832 | 3.16E-02 |
| | 8-Sep | -1.23 | 2.072723898 | 0.00845816401 | 4.94E-02 |
| | Afg3l2 | -1.23 | 3.497907483 | 0.00031775509 | 1.48E-02 |
| | Chchd4 | -1.23 | 1.47603842 | 0.03341654769 | 1.09E-01 |
| | Ipo9 | -1.23 | 2.371565347 | 0.00425044747 | 3.41E-02 |
| | Pde4dip | -1.23 | 3.018359462 | 0.00095860687 | 1.90E-02 |
| | Bace1 | -1.23 | 3.024087333 | 0.00094604690 | 1.89E-02 |
| | Med1 | -1.23 | 2.102337713 | 0.00790064025 | 4.78E-02 |
| | Ugcg | -1.23 | 2.750201061 | 0.00177745633 | 2.36E-02 |
| | Chrnb2 | -1.23 | 2.751962791 | 0.00177026062 | 2.36E-02 |
| | Glcci1 | -1.23 | 2.031194364 | 0.00930691260 | 5.20E-02 |
| | Ttbk2 | -1.23 | 1.446953279 | 0.03573112752 | 1.14E-01 |
| | Ccdc149 | -1.23 | 1.645941118 | 0.02259742128 | 8.54E-02 |
| | Fam8a1 | -1.23 | 2.209463013 | 0.00617357867 | 4.16E-02 |
| | Ambra1 | -1.23 | 2.968267667 | 0.00107580196 | 1.98E-02 |
| | Pcdhga12 | -1.23 | 1.399316198 | 0.03987344892 | 1.22E-01 |

| | | | | | |
|---|---|---|---|---|---|
| | Opa3 | -1.23 | 2.322430209 | 0.00475959271 | 3.61E-02 |
| | Hrasls | -1.23 | 1.67586271 | 0.02109294837 | 8.20E-02 |
| | Bcas3 | -1.23 | 2.953562677 | 0.00111285178 | 2.00E-02 |
| | Atp6v0c | -1.23 | 2.782720755 | 0.00164922248 | 2.30E-02 |
| | Dyrk1b | -1.23 | 1.416155986 | 0.03835694537 | 1.19E-01 |
| | Creld1 | -1.23 | 2.683386287 | 0.00207306879 | 2.51E-02 |
| | Hira | -1.23 | 2.818084646 | 0.00152025120 | 2.22E-02 |
| | Dexi | -1.23 | 2.435130478 | 0.00367171972 | 3.20E-02 |
| | Mgat3 | -1.23 | 3.641246071 | 0.00022843042 | 1.42E-02 |
| | Tcf25 | -1.23 | 2.333292993 | 0.00464202000 | 3.57E-02 |
| | Kctd13 | -1.23 | 1.661135098 | 0.02182051024 | 8.36E-02 |
| | A430033K04Rik | -1.23 | 1.456913294 | 0.03492100276 | 1.12E-01 |
| | Map2k4 | -1.23 | 2.010347695 | 0.00976455160 | 5.31E-02 |
| | Dlg3 | -1.23 | 3.263254435 | 0.00054543822 | 1.62E-02 |
| | Nol4 | -1.23 | 1.725987604 | 0.01879370461 | 7.68E-02 |
| | Plk2 | -1.23 | 1.964393721 | 0.01085441141 | 5.61E-02 |
| | Snx30 | -1.23 | 2.146445293 | 0.00713764111 | 4.51E-02 |
| | Dnajc5 | -1.23 | 3.570752513 | 0.00026868751 | 1.42E-02 |
| | Cbx6 | -1.23 | 1.832597442 | 0.01470288492 | 6.73E-02 |
| | Gnaq | -1.23 | 2.462719758 | 0.00344572205 | 3.10E-02 |
| | Med15 | -1.23 | 2.170665778 | 0.00675047327 | 4.37E-02 |
| | Atp9a | -1.23 | 2.491002612 | 0.00322847470 | 3.00E-02 |
| | Uhrf1bp1l | -1.23 | 2.10628285 | 0.00782919573 | 4.77E-02 |

| | | | | | |
|---|---|---|---|---|---|
| | Mturn | -1.23 | 1.564744951 | 0.02724300741 | 9.67E-02 |
| | Fam160a2 | -1.23 | 2.07115359 | 0.00848880212 | 4.94E-02 |
| | Dock9 | -1.23 | 2.949390584 | 0.00112359401 | 2.00E-02 |
| | Anxa11 | -1.23 | 1.494431708 | 0.03203083733 | 1.06E-01 |
| | Frmd5 | -1.23 | 1.468963584 | 0.03396537515 | 1.10E-01 |
| | Slc25a12 | -1.23 | 2.946819696 | 0.00113026506 | 2.00E-02 |
| | Map4k3 | -1.23 | 1.983015997 | 0.01039881861 | 5.47E-02 |
| | Taok2 | -1.23 | 3.148067505 | 0.00071110297 | 1.70E-02 |
| | Ptprd | -1.23 | 1.443613231 | 0.03600698593 | 1.14E-01 |
| | Apbb1 | -1.23 | 2.12811709 | 0.00744531213 | 4.63E-02 |
| | Arf3 | -1.23 | 2.101553533 | 0.00791491886 | 4.78E-02 |
| | Dixdc1 | -1.23 | 2.202875789 | 0.00626793105 | 4.19E-02 |
| | Atp6v1g2 | -1.23 | 1.819734426 | 0.01514487083 | 6.85E-02 |
| | Zfpl1 | -1.23 | 1.783975588 | 0.01644464159 | 7.18E-02 |
| | Cdc42bpa | -1.23 | 1.675312696 | 0.02111967856 | 8.21E-02 |
| | Tbc1d17 | -1.23 | 1.621257486 | 0.02391897218 | 8.86E-02 |
| | Atp13a2 | -1.24 | 2.949162326 | 0.00112418471 | 2.00E-02 |
| | Cers1 | -1.24 | 1.720255523 | 0.01904339946 | 7.74E-02 |
| | Gria4 | -1.24 | 1.726953888 | 0.01875193599 | 7.68E-02 |
| | Ubr4 | -1.24 | 2.234650653 | 0.00582571651 | 4.05E-02 |
| | Ppp5c | -1.24 | 2.494491417 | 0.00320264339 | 3.00E-02 |
| | Clip2 | -1.24 | 3.133605433 | 0.00073518149 | 1.73E-02 |
| | 2010107G23Rik | -1.24 | 2.008794361 | 0.00979953884 | 5.32E-02 |

| | | | | | |
|---|---|---|---|---|---|
| | Syt16 | -1.24 | 1.72471108 | 0.01884902630 | 7.69E-02 |
| | Lancl2 | -1.24 | 2.537834715 | 0.00289844647 | 2.89E-02 |
| | Arhgap10 | -1.24 | 1.527084167 | 0.02971090177 | 1.01E-01 |
| | Wbp2 | -1.24 | 2.548114259 | 0.00283064718 | 2.87E-02 |
| | Strn4 | -1.24 | 2.377417452 | 0.00419355697 | 3.39E-02 |
| | Asphd2 | -1.24 | 1.775191078 | 0.01678065553 | 7.27E-02 |
| | Lhfpl4 | -1.24 | 3.798331667 | 0.00015909932 | 1.41E-02 |
| | Zfp282 | -1.24 | 1.305844152 | 0.04944881035 | 1.40E-01 |
| | Lrrn2 | -1.24 | 1.681013601 | 0.02084425605 | 8.15E-02 |
| | Slc11a2 | -1.24 | 2.342920219 | 0.00454025015 | 3.52E-02 |
| | Fgf12 | -1.24 | 1.446040654 | 0.03580629173 | 1.14E-01 |
| | Tspan5 | -1.24 | 1.800456885 | 0.01583226735 | 7.03E-02 |
| | Spryd3 | -1.24 | 2.783558622 | 0.00164604377 | 2.30E-02 |
| | Ckmt1 | -1.24 | 2.18698622 | 0.00650150318 | 4.28E-02 |
| | AI837181 | -1.24 | 3.237663621 | 0.00057854398 | 1.64E-02 |
| | Mapk1ip1 | -1.24 | 1.703207058 | 0.01980582522 | 7.90E-02 |
| | Tom1l2 | -1.24 | 2.578982877 | 0.00263643533 | 2.77E-02 |
| | Sppl2b | -1.24 | 1.740478675 | 0.01817696306 | 7.56E-02 |
| | Slc7a1 | -1.24 | 2.027805964 | 0.00937980989 | 5.22E-02 |
| | Mgrn1 | -1.24 | 3.04074173 | 0.00091045455 | 1.87E-02 |
| | Bcl11a | -1.24 | 2.413904321 | 0.00385563291 | 3.27E-02 |
| | P4htm | -1.24 | 2.230030454 | 0.00588802365 | 4.07E-02 |
| | Tagap1 | -1.24 | 1.77040247 | 0.01696670584 | 7.30E-02 |

| | | | | | |
|---|---|---|---|---|---|
| | Brsk1 | -1.24 | 1.477395674 | 0.03331227754 | 1.09E-01 |
| | Sox5 | -1.24 | 1.367058164 | 0.04294789040 | 1.28E-01 |
| | Rassf5 | -1.24 | 1.792068386 | 0.01614104372 | 7.11E-02 |
| | Armcx4 | -1.24 | 2.058553922 | 0.00873868486 | 5.01E-02 |
| | Cnih3 | -1.24 | 1.530069767 | 0.02950735169 | 1.01E-01 |
| | Snap91 | -1.24 | 2.609626977 | 0.00245681822 | 2.68E-02 |
| | Ap3b2 | -1.24 | 3.325912252 | 0.00047215843 | 1.59E-02 |
| | Dvl1 | -1.24 | 1.466948883 | 0.03412330728 | 1.11E-01 |
| | Rasgrp1 | -1.24 | 2.419553638 | 0.00380580351 | 3.25E-02 |
| | Map7 | -1.24 | 1.377854489 | 0.04189339060 | 1.26E-01 |
| | Cables2 | -1.24 | 1.692147349 | 0.02031667585 | 8.02E-02 |
| | Mrm1 | -1.24 | 1.769856572 | 0.01698804598 | 7.30E-02 |
| | Abca8b | -1.24 | 1.359520642 | 0.04369979082 | 1.29E-01 |
| | Ttc3 | -1.24 | 2.606939131 | 0.00247207059 | 2.69E-02 |
| | Ergic1 | -1.24 | 3.315385762 | 0.00048374249 | 1.59E-02 |
| | 9330151L19Rik | -1.24 | 1.757948598 | 0.01746028798 | 7.39E-02 |
| | Zyg11b | -1.24 | 2.845570191 | 0.00142701918 | 2.17E-02 |
| | Celf3 | -1.24 | 1.387442313 | 0.04097865376 | 1.24E-01 |
| | Snap47 | -1.24 | 2.520296699 | 0.00301788927 | 2.93E-02 |
| | Man2c1 | -1.24 | 1.471185832 | 0.03379202115 | 1.10E-01 |
| | Atxn7l3 | -1.24 | 1.937612016 | 0.01154484176 | 5.80E-02 |
| | Mbd5 | -1.24 | 3.02543643 | 0.00094311265 | 1.89E-02 |
| | Psd3 | -1.24 | 1.926813387 | 0.01183550008 | 5.89E-02 |

| | | | | | |
|---|---|---|---|---|---|
| | Ankrd27 | -1.24 | 2.016747123 | 0.00962172360 | 5.27E-02 |
| | Arhgef9 | -1.24 | 2.365220249 | 0.00431300291 | 3.44E-02 |
| | Gfra4 | -1.24 | 1.356598206 | 0.04399484524 | 1.30E-01 |
| | Lrrc4c | -1.24 | 1.761885652 | 0.01730271874 | 7.36E-02 |
| | Asb6 | -1.24 | 2.105439884 | 0.00784440695 | 4.77E-02 |
| | Ubqln4 | -1.24 | 2.014886757 | 0.00966302812 | 5.28E-02 |
| | Dgkq | -1.24 | 2.771043349 | 0.00169416869 | 2.32E-02 |
| | Mark4 | -1.24 | 2.046668988 | 0.00898113060 | 5.10E-02 |
| | Glg1 | -1.24 | 2.41154776 | 0.00387661114 | 3.28E-02 |
| | Gripap1 | -1.24 | 2.783027284 | 0.00164805885 | 2.30E-02 |
| | Osbpl5 | -1.24 | 2.452283856 | 0.00352952404 | 3.14E-02 |
| | Pitpnm1 | -1.24 | 1.670146399 | 0.02137241515 | 8.25E-02 |
| | Klc1 | -1.24 | 2.879869078 | 0.00131865420 | 2.11E-02 |
| | Cyb561 | -1.24 | 1.866193276 | 0.01360838929 | 6.38E-02 |
| | Kcnn1 | -1.24 | 1.59999285 | 0.02511927787 | 9.15E-02 |
| | Jph1 | -1.24 | 1.316023603 | 0.04830325497 | 1.38E-01 |
| | Usp46 | -1.24 | 2.293643488 | 0.00508576762 | 3.74E-02 |
| | Rpap1 | -1.24 | 2.119664922 | 0.00759163077 | 4.69E-02 |
| | Abi2 | -1.24 | 2.831909207 | 0.00147262034 | 2.19E-02 |
| | Zfr2 | -1.24 | 1.402062243 | 0.03962212437 | 1.21E-01 |
| | Lonrf2 | -1.24 | 2.639043767 | 0.00229591726 | 2.61E-02 |
| | Spsb3 | -1.24 | 2.256033469 | 0.00554582972 | 3.94E-02 |
| | Gar1 | -1.24 | 1.541908124 | 0.02871387965 | 9.95E-02 |

| | | | | | |
|---|---|---|---|---|---|
| | Ptprk | -1.24 | 2.338968426 | 0.00458175195 | 3.54E-02 |
| | Mfsd6 | -1.24 | 3.562289086 | 0.00027397499 | 1.42E-02 |
| | Hk1 | -1.24 | 3.006467831 | 0.00098521762 | 1.92E-02 |
| | Zc3h13 | -1.24 | 3.057326983 | 0.00087634077 | 1.85E-02 |
| | Spns2 | -1.24 | 1.734041205 | 0.01844840375 | 7.62E-02 |
| | Ogfod2 | -1.24 | 2.009911513 | 0.00977436353 | 5.32E-02 |
| | Opcml | -1.24 | 2.059466558 | 0.00872034049 | 5.01E-02 |
| | Ndrg3 | -1.25 | 2.460652886 | 0.00346215983 | 3.10E-02 |
| | Sv2b | -1.25 | 1.576301602 | 0.02652762672 | 9.50E-02 |
| | Atf6 | -1.25 | 1.948479415 | 0.01125953837 | 5.72E-02 |
| | Map1b | -1.25 | 1.997080756 | 0.01006744450 | 5.38E-02 |
| | Zfp142 | -1.25 | 2.070886455 | 0.00849402519 | 4.94E-02 |
| | Ash1l | -1.25 | 2.590414999 | 0.00256794076 | 2.72E-02 |
| | Ccdc124 | -1.25 | 1.526995003 | 0.02971700223 | 1.01E-01 |
| | Myadm | -1.25 | 2.459807444 | 0.00346890620 | 3.10E-02 |
| | Rundc3a | -1.25 | 1.752888999 | 0.01766489259 | 7.44E-02 |
| | D17Wsu92e | -1.25 | 2.132784658 | 0.00736572231 | 4.60E-02 |
| | Josd1 | -1.25 | 2.684994221 | 0.00206540764 | 2.51E-02 |
| | Ptk2 | -1.25 | 1.647493739 | 0.02251677879 | 8.53E-02 |
| | Map4k2 | -1.25 | 1.975053503 | 0.01059123238 | 5.52E-02 |
| | Rfng | -1.25 | 1.964559143 | 0.01085027779 | 5.61E-02 |
| | Myl6b | -1.25 | 1.317480544 | 0.04814148203 | 1.38E-01 |
| | Entpd4 | -1.25 | 2.617648869 | 0.00241185464 | 2.66E-02 |

| | | | | | |
|---|---|---|---|---|---|
| | Necab3 | -1.25 | 2.0678536 97 | 0.00855354 812 | 4.96E-02 |
| | Arhgap35 | -1.25 | 2.5995004 79 | 0.00251477 724 | 2.70E-02 |
| | Pde8b | -1.25 | 1.8921859 45 | 0.01281781 666 | 6.15E-02 |
| | Rnf150 | -1.25 | 1.6848258 04 | 0.02066208 748 | 8.11E-02 |
| | Ctxn1 | -1.25 | 1.5654875 35 | 0.02719646 546 | 9.66E-02 |
| | Grip2 | -1.25 | 2.7974661 5 | 0.00159416 713 | 2.27E-02 |
| | Slc22a17 | -1.25 | 2.4934052 43 | 0.00321066 325 | 3.00E-02 |
| | Rtn1 | -1.25 | 2.8410809 57 | 0.00144184 655 | 2.17E-02 |
| | Ppp6r2 | -1.25 | 2.7623054 46 | 0.00172860 018 | 2.34E-02 |
| | Prkaa2 | -1.25 | 1.7460441 7 | 0.01794551 100 | 7.51E-02 |
| | Ice1 | -1.25 | 2.5034047 | 0.00313758 355 | 2.97E-02 |
| | Slc38a7 | -1.25 | 2.1676375 07 | 0.00679770 780 | 4.38E-02 |
| | Srpk1 | -1.25 | 2.4819699 27 | 0.00329632 537 | 3.01E-02 |
| | Plcb1 | -1.25 | 1.7006218 6 | 0.01992407 373 | 7.93E-02 |
| | Gnl3l | -1.25 | 2.5922952 31 | 0.00255684 716 | 2.72E-02 |
| | Vars2 | -1.25 | 2.1345197 55 | 0.00733635 343 | 4.59E-02 |
| | Fam160b2 | -1.25 | 2.2548573 57 | 0.00556086 873 | 3.95E-02 |
| | Ablim1 | -1.25 | 2.9176020 22 | 0.00120892 116 | 2.04E-02 |
| | Arhgef2 | -1.25 | 2.1073078 52 | 0.00781073 941 | 4.77E-02 |
| | Lsamp | -1.25 | 1.7665674 98 | 0.01711719 120 | 7.33E-02 |
| | Atp6v1b2 | -1.25 | 3.0002788 66 | 0.00099935 809 | 1.92E-02 |
| | Lrrc73 | -1.25 | 1.3778454 05 | 0.04189426 687 | 1.26E-01 |
| | Lrrtm1 | -1.25 | 1.5542969 23 | 0.02790635 259 | 9.81E-02 |

| | | | | | |
|---|---|---|---|---|---|
| | Chga | -1.25 | 2.5972747<br>9 | 0.00252769<br>815 | 2.71E-<br>02 |
| | Zfp827 | -1.25 | 2.8242509<br>77 | 0.00149881<br>843 | 2.21E-<br>02 |
| | Btrc | -1.25 | 2.6399505<br>67 | 0.00229112<br>842 | 2.61E-<br>02 |
| | Dnajc30 | -1.25 | 2.0314386<br>23 | 0.00930167<br>962 | 5.20E-<br>02 |
| | App | -1.25 | 2.0283606<br>51 | 0.00936783<br>751 | 5.22E-<br>02 |
| | Slc25a42 | -1.25 | 1.4438972 | 0.03598345<br>002 | 1.14E-<br>01 |
| | Fam131a | -1.25 | 1.3197715<br>27 | 0.04788819<br>552 | 1.37E-<br>01 |
| | Mpg | -1.25 | 2.0072783<br>14 | 0.00983380<br>712 | 5.32E-<br>02 |
| | Miga2 | -1.25 | 2.9316381<br>18 | 0.00117047<br>430 | 2.03E-<br>02 |
| | Gtpbp3 | -1.25 | 1.9990568<br>39 | 0.01002174<br>069 | 5.38E-<br>02 |
| | Slc6a7 | -1.25 | 1.6192338<br>51 | 0.02403068<br>491 | 8.90E-<br>02 |
| | Zbtb40 | -1.25 | 1.3286246<br>4 | 0.04692187<br>517 | 1.36E-<br>01 |
| | Dgke | -1.25 | 2.4468091<br>51 | 0.00357429<br>875 | 3.16E-<br>02 |
| | Smpd4 | -1.25 | 1.5595971<br>63 | 0.02756784<br>617 | 9.74E-<br>02 |
| | Cmc2 | -1.25 | 1.8476495<br>06 | 0.01420203<br>223 | 6.56E-<br>02 |
| | Wasf3 | -1.25 | 2.4991808<br>33 | 0.00316824<br>798 | 2.97E-<br>02 |
| | Gpr61 | -1.25 | 1.4622560<br>94 | 0.03449402<br>756 | 1.11E-<br>01 |
| | Mapk14 | -1.25 | 2.6659824<br>39 | 0.00215783<br>166 | 2.55E-<br>02 |
| | Scamp5 | -1.25 | 2.7583569<br>08 | 0.00174438<br>801 | 2.34E-<br>02 |
| | Ncdn | -1.25 | 2.2642073<br>98 | 0.00544242<br>686 | 3.90E-<br>02 |
| | 9530082P21<br>Rik | -1.25 | 1.6090916<br>24 | 0.02459848<br>590 | 9.03E-<br>02 |
| | Pnkd | -1.25 | 2.6250572<br>23 | 0.00237106<br>127 | 2.65E-<br>02 |
| | Lrrc4b | -1.25 | 2.0837767<br>68 | 0.00824561<br>839 | 4.90E-<br>02 |

| | | | | | |
|---|---|---|---|---|---|
| | Cdr2l | -1.25 | 1.428980277 | 0.03724086182 | 1.17E-01 |
| | Fabp3 | -1.25 | 1.762946754 | 0.01726049497 | 7.36E-02 |
| | Dcaf5 | -1.25 | 3.479165525 | 0.00033176798 | 1.49E-02 |
| | Tbc1d24 | -1.25 | 1.969925426 | 0.01071703315 | 5.56E-02 |
| | Pacs1 | -1.25 | 2.092713278 | 0.00807768145 | 4.83E-02 |
| | Nova2 | -1.25 | 2.634131275 | 0.00232203480 | 2.62E-02 |
| | Lhx2 | -1.25 | 1.465650801 | 0.03422545256 | 1.11E-01 |
| | Dcbld2 | -1.25 | 1.518838711 | 0.03028037776 | 1.03E-01 |
| | Grik5 | -1.25 | 2.786769363 | 0.00163391943 | 2.29E-02 |
| | Esrra | -1.25 | 1.957878622 | 0.01101847214 | 5.65E-02 |
| | Elavl2 | -1.25 | 1.686097841 | 0.02060165732 | 8.09E-02 |
| | B4galnt1 | -1.25 | 2.400872663 | 0.00397308024 | 3.32E-02 |
| | Vamp2 | -1.25 | 2.957088456 | 0.00110385377 | 2.00E-02 |
| | Gon4l | -1.25 | 1.338611355 | 0.04585520562 | 1.33E-01 |
| | Plekhm3 | -1.25 | 2.22868363 | 0.00590631180 | 4.08E-02 |
| | Tceal5 | -1.25 | 2.590970642 | 0.00256465740 | 2.72E-02 |
| | Slc4a3 | -1.26 | 2.775434021 | 0.00167712711 | 2.32E-02 |
| | Ppp2r5b | -1.26 | 3.672155447 | 0.00021273775 | 1.41E-02 |
| | Kdm7a | -1.26 | 1.894736888 | 0.01274274850 | 6.14E-02 |
| | Zfyve27 | -1.26 | 2.700358523 | 0.00199361585 | 2.47E-02 |
| | Jade1 | -1.26 | 2.700971148 | 0.00199080559 | 2.47E-02 |
| | Prrc2b | -1.26 | 3.29490201 | 0.00050710511 | 1.59E-02 |
| | Ak5 | -1.26 | 1.912996591 | 0.01221809252 | 5.98E-02 |

| | | | | | |
|---|---|---|---|---|---|
| | Usp32 | -1.26 | 2.3243482<br>14 | 0.00473861<br>894 | 3.61E-<br>02 |
| | Rundc3b | -1.26 | 2.6354465<br>37 | 0.00231501<br>315 | 2.62E-<br>02 |
| | Dnm3 | -1.26 | 1.6068387<br>65 | 0.02472641<br>963 | 9.06E-<br>02 |
| | Drosha | -1.26 | 3.2208023<br>75 | 0.00060144<br>736 | 1.65E-<br>02 |
| | Iffo1 | -1.26 | 1.8009830<br>89 | 0.01581309<br>611 | 7.03E-<br>02 |
| | Chn1 | -1.26 | 1.6460577<br>13 | 0.02259135<br>536 | 8.54E-<br>02 |
| | Ccdc136 | -1.26 | 1.7070948<br>79 | 0.01962931<br>396 | 7.86E-<br>02 |
| | Usf3 | -1.26 | 3.1921108<br>64 | 0.00064252<br>368 | 1.65E-<br>02 |
| | Vstm5 | -1.26 | 1.4745794<br>01 | 0.03352899<br>988 | 1.09E-<br>01 |
| | Ncoa1 | -1.26 | 3.2671368<br>19 | 0.00054058<br>399 | 1.61E-<br>02 |
| | Pianp | -1.26 | 1.8426997<br>36 | 0.01436482<br>249 | 6.61E-<br>02 |
| | Krba1 | -1.26 | 1.9624310<br>41 | 0.01090357<br>609 | 5.62E-<br>02 |
| | Impdh1 | -1.26 | 2.6928793<br>45 | 0.00202824<br>612 | 2.49E-<br>02 |
| | Trp53bp1 | -1.26 | 2.8723608<br>77 | 0.00134164<br>965 | 2.12E-<br>02 |
| | Rell2 | -1.26 | 1.3693095<br>63 | 0.04272582<br>304 | 1.27E-<br>01 |
| | Foxk1 | -1.26 | 2.2447236<br>74 | 0.00569214<br>987 | 3.99E-<br>02 |
| | Pcnt | -1.26 | 2.0356669<br>02 | 0.00921155<br>814 | 5.17E-<br>02 |
| | Jakmip1 | -1.26 | 2.5170220<br>73 | 0.00304073<br>047 | 2.94E-<br>02 |
| | Sertm1 | -1.26 | 1.4575456<br>52 | 0.03487019<br>274 | 1.12E-<br>01 |
| | Sgip1 | -1.26 | 3.0493348<br>65 | 0.00089261<br>696 | 1.85E-<br>02 |
| | AI854703 | -1.26 | 1.4387401<br>64 | 0.03641328<br>293 | 1.15E-<br>01 |
| | Tspyl4 | -1.26 | 2.8437630<br>24 | 0.00143296<br>960 | 2.17E-<br>02 |
| | Map3k10 | -1.26 | 2.5963727<br>12 | 0.00253295<br>391 | 2.71E-<br>02 |

| | | | | | |
|---|---|---|---|---|---|
| | Syp | -1.26 | 2.164685403 | 0.00684407243 | 4.39E-02 |
| | Rhot2 | -1.26 | 2.043625887 | 0.00904428236 | 5.13E-02 |
| | Mapre2 | -1.26 | 3.454770685 | 0.00035093713 | 1.49E-02 |
| | Pknox2 | -1.26 | 2.016737194 | 0.00962194358 | 5.27E-02 |
| | Drp2 | -1.26 | 1.868733325 | 0.01352903046 | 6.35E-02 |
| | Zfp612 | -1.26 | 2.071796534 | 0.00847624431 | 4.94E-02 |
| | Napa | -1.26 | 2.106503931 | 0.00782521122 | 4.77E-02 |
| | Arhgap33 | -1.26 | 1.302279663 | 0.04985633350 | 1.41E-01 |
| | Gabbr1 | -1.26 | 2.874816956 | 0.00133408360 | 2.12E-02 |
| | Eno2 | -1.26 | 2.457281812 | 0.00348913833 | 3.11E-02 |
| | Reps2 | -1.26 | 1.708128294 | 0.01958266102 | 7.85E-02 |
| | St3gal3 | -1.26 | 2.503991091 | 0.00313335000 | 2.97E-02 |
| | Gdf1 | -1.26 | 2.20409386 | 0.00625037594 | 4.18E-02 |
| | Tcaf1 | -1.26 | 3.904958365 | 0.00012446339 | 1.41E-02 |
| | Sprn | -1.26 | 2.299084895 | 0.00502244402 | 3.72E-02 |
| | Zbtb38 | -1.26 | 2.283635739 | 0.00520432322 | 3.80E-02 |
| | Sorcs1 | -1.26 | 2.043675275 | 0.00904325392 | 5.13E-02 |
| | Olfm1 | -1.26 | 1.813425913 | 0.01536646911 | 6.90E-02 |
| | Ppp1r9a | -1.26 | 2.436412519 | 0.00366089675 | 3.20E-02 |
| | Vti1a | -1.26 | 3.245811788 | 0.00056779062 | 1.64E-02 |
| | Mycbp2 | -1.26 | 2.370097354 | 0.00426483905 | 3.42E-02 |
| | Bcorl1 | -1.26 | 2.550396329 | 0.00281581210 | 2.86E-02 |
| | Prpf40b | -1.26 | 2.420291519 | 0.00379934281 | 3.25E-02 |

| | | | | | |
|---|---|---|---|---|---|
| | Kat2a | -1.26 | 3.964953344 | 0.00010840434 | 1.41E-02 |
| | Nap1l2 | -1.26 | 1.70017885 | 0.01994440800 | 7.94E-02 |
| | Cdkl1 | -1.26 | 1.434561281 | 0.03676535122 | 1.16E-01 |
| | Ece1 | -1.26 | 2.36342076 | 0.00433091081 | 3.45E-02 |
| | Fscn1 | -1.26 | 1.764552991 | 0.01719677500 | 7.35E-02 |
| | Zer1 | -1.26 | 2.437908663 | 0.00364830667 | 3.20E-02 |
| | Zfp335 | -1.26 | 1.9473022 | 0.01129010030 | 5.72E-02 |
| | Setd1a | -1.26 | 2.317714793 | 0.00481155225 | 3.64E-02 |
| | Ywhah | -1.26 | 1.822574735 | 0.01504614576 | 6.82E-02 |
| | Ypel3 | -1.26 | 2.80334601 | 0.00157272935 | 2.26E-02 |
| | Lin37 | -1.26 | 1.830012947 | 0.01479064293 | 6.75E-02 |
| | Sntg1 | -1.26 | 1.37330083 | 0.04233496158 | 1.27E-01 |
| | Cpeb1 | -1.26 | 1.926192339 | 0.01185243715 | 5.89E-02 |
| | Mta3 | -1.26 | 3.314611534 | 0.00048460564 | 1.59E-02 |
| | Tesk1 | -1.26 | 2.724952447 | 0.00188385535 | 2.41E-02 |
| | Rasgef1c | -1.26 | 1.53120364 | 0.02943041323 | 1.01E-01 |
| | Cyp46a1 | -1.26 | 2.266238894 | 0.00541702832 | 3.90E-02 |
| | Ubox5 | -1.26 | 2.23471817 | 0.00582481089 | 4.05E-02 |
| | Lrfn3 | -1.26 | 1.973525761 | 0.01062855536 | 5.53E-02 |
| | Trrap | -1.26 | 2.780597137 | 0.00165730661 | 2.30E-02 |
| | Wipf2 | -1.26 | 1.853947278 | 0.01399757238 | 6.49E-02 |
| | Abhd8 | -1.26 | 2.204928389 | 0.00623837692 | 4.18E-02 |
| | Cry2 | -1.26 | 2.055771323 | 0.00879485487 | 5.03E-02 |

| | | | | | |
|---|---|---|---|---|---|
| | Rap1gap | -1.26 | 1.761461354 | 0.01731963143 | 7.36E-02 |
| | Syndig1 | -1.26 | 1.515060406 | 0.03054496233 | 1.03E-01 |
| | Nol6 | -1.26 | 3.256539267 | 0.00055393746 | 1.63E-02 |
| | Adcy6 | -1.27 | 1.77496719 | 0.01678930854 | 7.27E-02 |
| | Zcchc2 | -1.27 | 2.136750963 | 0.00729875921 | 4.57E-02 |
| | Agap3 | -1.27 | 2.027749437 | 0.00938103083 | 5.22E-02 |
| | Brpf3 | -1.27 | 2.601902325 | 0.00250090776 | 2.70E-02 |
| | Slc25a37 | -1.27 | 1.802178952 | 0.01576961344 | 7.02E-02 |
| | Mcoln1 | -1.27 | 3.147731448 | 0.00071165344 | 1.70E-02 |
| | Chpf | -1.27 | 2.20793242 | 0.00619537473 | 4.16E-02 |
| | Nlk | -1.27 | 1.895878708 | 0.01270929007 | 6.13E-02 |
| | Gpr176 | -1.27 | 2.112147982 | 0.00772417345 | 4.74E-02 |
| | Ralgapa1 | -1.27 | 2.304358882 | 0.00496182129 | 3.70E-02 |
| | Soga3 | -1.27 | 2.788706543 | 0.00162664753 | 2.29E-02 |
| | Ptpn23 | -1.27 | 2.108113165 | 0.00779626935 | 4.77E-02 |
| | Dusp26 | -1.27 | 2.190772256 | 0.00644507157 | 4.25E-02 |
| | Prkacb | -1.27 | 2.439844943 | 0.00363207709 | 3.19E-02 |
| | Amigo1 | -1.27 | 2.996934922 | 0.00100708257 | 1.92E-02 |
| | Tspyl5 | -1.27 | 2.148877799 | 0.00709777457 | 4.50E-02 |
| | Ncam2 | -1.27 | 1.874082975 | 0.01336340176 | 6.31E-02 |
| | Hook1 | -1.27 | 2.163570405 | 0.00686166633 | 4.40E-02 |
| | Ss18l1 | -1.27 | 2.795961928 | 0.00159969826 | 2.27E-02 |
| | Lin7a | -1.27 | 1.484864122 | 0.03274431264 | 1.08E-01 |

| | | | | | |
|---|---|---|---|---|---|
| | Rnf144b | -1.27 | 1.923963118 | 0.01191343179 | 5.91E-02 |
| | Erc1 | -1.27 | 2.089120825 | 0.00814477657 | 4.86E-02 |
| | Numb | -1.27 | 3.307194832 | 0.00049295261 | 1.59E-02 |
| | Zswim8 | -1.27 | 2.766935024 | 0.00171027118 | 2.33E-02 |
| | Rbm33 | -1.27 | 2.086658743 | 0.00819108168 | 4.88E-02 |
| | Socs7 | -1.27 | 2.105837797 | 0.00783722298 | 4.77E-02 |
| | Plxna2 | -1.27 | 3.451810471 | 0.00035333733 | 1.49E-02 |
| | Erf | -1.27 | 2.071159314 | 0.00848869024 | 4.94E-02 |
| | Trub1 | -1.27 | 1.769759213 | 0.01699185476 | 7.30E-02 |
| | Gigyf1 | -1.27 | 1.728935522 | 0.01866656804 | 7.66E-02 |
| | Slc1a1 | -1.27 | 3.283020663 | 0.00052116991 | 1.60E-02 |
| | Dapk3 | -1.27 | 2.237782447 | 0.00578385708 | 4.03E-02 |
| | Baiap2 | -1.27 | 1.419853812 | 0.03803173932 | 1.18E-01 |
| | Arhgap21 | -1.27 | 3.090311102 | 0.00081224846 | 1.79E-02 |
| | Sfswap | -1.27 | 1.415011165 | 0.03845818947 | 1.19E-01 |
| | Mnt | -1.27 | 1.320926574 | 0.04776100158 | 1.37E-01 |
| | Smurf1 | -1.27 | 2.245736178 | 0.00567889478 | 3.99E-02 |
| | Map6d1 | -1.27 | 1.51876513 | 0.03028550848 | 1.03E-01 |
| | Spata2l | -1.27 | 1.404505361 | 0.03939985637 | 1.21E-01 |
| | Sh3bp5l | -1.27 | 3.271001489 | 0.00053579482 | 1.61E-02 |
| | Sugp2 | -1.27 | 2.482587348 | 0.00329164243 | 3.01E-02 |
| | Limk1 | -1.27 | 1.977953824 | 0.01052073729 | 5.49E-02 |
| | Asphd1 | -1.27 | 1.455205271 | 0.03505861288 | 1.12E-01 |

| | | | | | |
|---|---|---|---|---|---|
| | Atrnl1 | -1.27 | 1.984855315 | 0.01035487081 | 5.46E-02 |
| | Zfp239 | -1.27 | 1.893564931 | 0.01277718166 | 6.14E-02 |
| | Arfgef2 | -1.27 | 3.316404259 | 0.00048260936 | 1.59E-02 |
| | Gng3 | -1.27 | 2.727209097 | 0.00187409198 | 2.41E-02 |
| | Gsk3a | -1.27 | 2.537866164 | 0.00289823659 | 2.89E-02 |
| | Cdc42bpb | -1.27 | 2.384771585 | 0.00412314316 | 3.36E-02 |
| | Stxbp5 | -1.27 | 2.849180125 | 0.00141520670 | 2.17E-02 |
| | Wdr7 | -1.27 | 3.111877866 | 0.00077289791 | 1.76E-02 |
| | Cacng8 | -1.27 | 1.651163995 | 0.02232728955 | 8.48E-02 |
| | Cacnb3 | -1.27 | 2.101588177 | 0.00791428750 | 4.78E-02 |
| | Tceal6 | -1.27 | 2.648795908 | 0.00224493666 | 2.60E-02 |
| | B4galt2 | -1.27 | 1.792718076 | 0.01611691531 | 7.11E-02 |
| | Tsc1 | -1.27 | 3.247073077 | 0.00056614402 | 1.64E-02 |
| | Rasl11b | -1.27 | 1.447970456 | 0.03564753825 | 1.14E-01 |
| | Tnpo2 | -1.27 | 3.45905574 | 0.00034749156 | 1.49E-02 |
| | Snrpn | -1.27 | 2.272147412 | 0.00534382944 | 3.86E-02 |
| | Rhof | -1.27 | 2.53559495 | 0.00291343310 | 2.89E-02 |
| | Edc4 | -1.27 | 2.690655118 | 0.00203866038 | 2.50E-02 |
| | Dusp14 | -1.27 | 1.936423933 | 0.01157646776 | 5.81E-02 |
| | Got1 | -1.27 | 2.690609416 | 0.00203887492 | 2.50E-02 |
| | Lin7b | -1.27 | 1.861593605 | 0.01375328349 | 6.41E-02 |
| | Gmeb2 | -1.27 | 1.588953731 | 0.02576595648 | 9.33E-02 |
| | Kcnip2 | -1.27 | 1.545830438 | 0.02845571887 | 9.90E-02 |

| | | | | | |
|---|---|---|---|---|---|
| | Lrrc3b | -1.28 | 1.584169246 | 0.02605138123 | 9.38E-02 |
| | Dbn1 | -1.28 | 2.789654476 | 0.00162310092 | 2.29E-02 |
| | Sbno1 | -1.28 | 3.874339397 | 0.00013355514 | 1.41E-02 |
| | Pdzrn3 | -1.28 | 1.360492219 | 0.04360213763 | 1.29E-01 |
| | Tmem44 | -1.28 | 2.533458212 | 0.00292780257 | 2.90E-02 |
| | Pde4a | -1.28 | 2.80188843 | 0.00157801661 | 2.26E-02 |
| | Snurf | -1.28 | 2.252396917 | 0.00559246253 | 3.96E-02 |
| | Agtpbp1 | -1.28 | 1.76132134 | 0.01732521607 | 7.36E-02 |
| | Dip2b | -1.28 | 3.74844612 | 0.00017846534 | 1.41E-02 |
| | Zfp697 | -1.28 | 1.628935051 | 0.02349984238 | 8.77E-02 |
| | Dennd5b | -1.28 | 2.094437851 | 0.00804566876 | 4.82E-02 |
| | Rab15 | -1.28 | 2.076423489 | 0.00838641812 | 4.92E-02 |
| | Mfn2 | -1.28 | 3.227416682 | 0.00059235672 | 1.65E-02 |
| | Gpd1 | -1.28 | 2.00309483 | 0.00992899222 | 5.36E-02 |
| | Otud3 | -1.28 | 1.540522747 | 0.02880562171 | 9.97E-02 |
| | Atxn2 | -1.28 | 2.878473394 | 0.00132289875 | 2.11E-02 |
| | Rgs7bp | -1.28 | 1.474084252 | 0.03356724882 | 1.09E-01 |
| | Habp4 | -1.28 | 2.210127582 | 0.00616413892 | 4.15E-02 |
| | Rangap1 | -1.28 | 3.577043331 | 0.00026482359 | 1.42E-02 |
| | Ets2 | -1.28 | 1.644480359 | 0.02267355616 | 8.56E-02 |
| | Pde5a | -1.28 | 1.369307002 | 0.04272607491 | 1.27E-01 |
| | Ralgapb | -1.28 | 2.615172675 | 0.00242564547 | 2.66E-02 |
| | Btbd10 | -1.28 | 2.210299772 | 0.00616169543 | 4.15E-02 |

| | | | | | |
|---|---|---|---|---|---|
| | Atxn2l | -1.28 | 1.852028583 | 0.01405954988 | 6.50E-02 |
| | Panx2 | -1.28 | 1.722778981 | 0.01893306906 | 7.71E-02 |
| | Jade2 | -1.28 | 2.050675395 | 0.00889865983 | 5.07E-02 |
| | Mllt6 | -1.28 | 2.763876707 | 0.00172235747 | 2.33E-02 |
| | Fbxl16 | -1.28 | 1.740665751 | 0.01816913487 | 7.56E-02 |
| | Amer2 | -1.28 | 1.764999388 | 0.01717910806 | 7.34E-02 |
| | Plcl2 | -1.28 | 1.555155886 | 0.02785121295 | 9.80E-02 |
| | Sfxn3 | -1.28 | 3.213602925 | 0.00061150086 | 1.65E-02 |
| | Zc3h3 | -1.28 | 2.211985077 | 0.00613783095 | 4.15E-02 |
| | Dffa | -1.28 | 2.499726884 | 0.00316426695 | 2.97E-02 |
| | Tmem108 | -1.28 | 1.7399566 | 0.01819882713 | 7.57E-02 |
| | Nedd4l | -1.28 | 3.182178937 | 0.00065738693 | 1.66E-02 |
| | Mdn1 | -1.28 | 2.413604094 | 0.00385829923 | 3.27E-02 |
| | Camk1d | -1.28 | 2.854321178 | 0.00139855266 | 2.15E-02 |
| | Spats2 | -1.28 | 2.255968399 | 0.00554666072 | 3.94E-02 |
| | Slc36a1 | -1.28 | 2.572595911 | 0.00267549466 | 2.77E-02 |
| | Caskin1 | -1.28 | 2.385373167 | 0.00411743577 | 3.36E-02 |
| | Lpgat1 | -1.28 | 2.001817724 | 0.00995823283 | 5.37E-02 |
| | Susd4 | -1.28 | 2.641127565 | 0.00228492755 | 2.61E-02 |
| | Kctd17 | -1.28 | 2.029778039 | 0.00933731394 | 5.21E-02 |
| | Mkl1 | -1.28 | 2.441120174 | 0.00362142775 | 3.18E-02 |
| | Srgap3 | -1.28 | 3.031300886 | 0.00093046301 | 1.89E-02 |
| | Adap1 | -1.28 | 1.957166848 | 0.01103654536 | 5.65E-02 |

| | | | | | |
|---|---|---|---|---|---|
| | Gdpgp1 | -1.28 | 1.557173653 | 0.02772211414 | 9.77E-02 |
| | Fsd1 | -1.28 | 2.901618889 | 0.00125424134 | 2.07E-02 |
| | Rab3ip | -1.28 | 2.639349441 | 0.00229430187 | 2.61E-02 |
| | Epb41l1 | -1.28 | 2.133964154 | 0.00734574497 | 4.59E-02 |
| | Fam131b | -1.28 | 2.890129655 | 0.00128786501 | 2.10E-02 |
| | Helq | -1.28 | 1.894586158 | 0.01274717190 | 6.14E-02 |
| | Creg2 | -1.28 | 2.304142816 | 0.00496429046 | 3.70E-02 |
| | Vps9d1 | -1.28 | 2.169779321 | 0.00676426602 | 4.37E-02 |
| | Dnajc16 | -1.28 | 2.978969845 | 0.00104961531 | 1.97E-02 |
| | Cntrob | -1.28 | 1.779939963 | 0.01659816347 | 7.22E-02 |
| | Setbp1 | -1.28 | 2.056621752 | 0.00877764975 | 5.03E-02 |
| | Coro2a | -1.28 | 2.51296776 | 0.00306924983 | 2.94E-02 |
| | Map9 | -1.28 | 2.073750207 | 0.00843819958 | 4.94E-02 |
| | A230050P20Rik | -1.28 | 2.280658061 | 0.00524012853 | 3.82E-02 |
| | Nsf | -1.29 | 3.31694747 | 0.00048200610 | 1.59E-02 |
| | Napepld | -1.29 | 1.87392402 | 0.01336829374 | 6.31E-02 |
| | Atg4c | -1.29 | 1.978878422 | 0.01049836282 | 5.49E-02 |
| | Crebbp | -1.29 | 2.727941132 | 0.00187093573 | 2.41E-02 |
| | Otub2 | -1.29 | 2.060227716 | 0.00870507033 | 5.01E-02 |
| | Foxp1 | -1.29 | 1.514607048 | 0.03057686470 | 1.04E-01 |
| | Dnajc6 | -1.29 | 2.829131918 | 0.00148206783 | 2.19E-02 |
| | Herc1 | -1.29 | 2.326345398 | 0.00471687755 | 3.60E-02 |
| | Rcor2 | -1.29 | 1.511693829 | 0.03078266179 | 1.04E-01 |

| | | | | | |
|---|---|---|---|---|---|
| | Zgpat | -1.29 | 1.730878874 | 0.01858322676 | 7.64E-02 |
| | Mthfsd | -1.29 | 2.313223691 | 0.00486156737 | 3.64E-02 |
| | Atp2a2 | -1.29 | 2.112414729 | 0.00771943067 | 4.74E-02 |
| | Atp1a3 | -1.29 | 2.64399819 | 0.00226987431 | 2.61E-02 |
| | Wee1 | -1.29 | 1.497100644 | 0.03183459694 | 1.06E-01 |
| | Nlgn2 | -1.29 | 3.285160718 | 0.00051860808 | 1.60E-02 |
| | Mapk9 | -1.29 | 2.356002592 | 0.00440552234 | 3.47E-02 |
| | Zfp668 | -1.29 | 1.678436214 | 0.02096832726 | 8.18E-02 |
| | Clip1 | -1.29 | 3.59386271 | 0.00025476355 | 1.42E-02 |
| | Particl | -1.29 | 1.566973025 | 0.02710359973 | 9.64E-02 |
| | Mapt | -1.29 | 3.217783702 | 0.00060564244 | 1.65E-02 |
| | Ppp1r26 | -1.29 | 1.68312974 | 0.02074293757 | 8.12E-02 |
| | Bri3bp | -1.29 | 2.459586374 | 0.00347067243 | 3.10E-02 |
| | Matk | -1.29 | 1.743837555 | 0.01803692275 | 7.54E-02 |
| | B3gat1 | -1.29 | 1.866273422 | 0.01360587819 | 6.38E-02 |
| | Sorbs2 | -1.29 | 1.700884116 | 0.01991204587 | 7.93E-02 |
| | Kifc2 | -1.29 | 1.355272882 | 0.04412930806 | 1.30E-01 |
| | Celsr2 | -1.29 | 2.538595064 | 0.00289337641 | 2.89E-02 |
| | R3hdm4 | -1.29 | 2.134123383 | 0.00734305222 | 4.59E-02 |
| | Cnnm4 | -1.29 | 1.733324512 | 0.01847887332 | 7.62E-02 |
| | Ttll1 | -1.29 | 2.533017037 | 0.00293077827 | 2.90E-02 |
| | Med24 | -1.29 | 2.605105812 | 0.00248252819 | 2.69E-02 |
| | Ap5z1 | -1.29 | 2.751342643 | 0.00177279026 | 2.36E-02 |

| | | | | | |
|---|---|---|---|---|---|
| | Usp20 | -1.29 | 3.2974886<br>2 | 0.00050409<br>383 | 1.59E-<br>02 |
| | Pdp1 | -1.29 | 2.0377350<br>55 | 0.00916779<br>609 | 5.16E-<br>02 |
| | Pip4k2c | -1.29 | 2.8946805<br>47 | 0.00127444<br>017 | 2.08E-<br>02 |
| | Dlg4 | -1.29 | 2.1570932<br>14 | 0.00696477<br>011 | 4.43E-<br>02 |
| | Zfp94 | -1.29 | 1.6923280<br>79 | 0.02030822<br>287 | 8.02E-<br>02 |
| | Tacc1 | -1.29 | 3.2775229<br>5 | 0.00052780<br>931 | 1.61E-<br>02 |
| | Megf8 | -1.29 | 3.9573036<br>64 | 0.00011033<br>069 | 1.41E-<br>02 |
| | 2310057M21<br>Rik | -1.29 | 2.2865977<br>89 | 0.00516894<br>856 | 3.78E-<br>02 |
| | Ppp2r2c | -1.29 | 2.3009261<br>11 | 0.00500119<br>616 | 3.71E-<br>02 |
| | 4931428F04<br>Rik | -1.29 | 1.7815546<br>45 | 0.01653656<br>703 | 7.20E-<br>02 |
| | Acd | -1.29 | 2.1703506<br>5 | 0.00675537<br>326 | 4.37E-<br>02 |
| | 6-Sep | -1.29 | 1.9590061<br>93 | 0.01098990<br>167 | 5.64E-<br>02 |
| | Kif1a | -1.29 | 1.9336999<br>59 | 0.01164930<br>566 | 5.83E-<br>02 |
| | Zzef1 | -1.29 | 4.4683455<br>13 | 0.00003401<br>375 | 1.41E-<br>02 |
| | Mast2 | -1.29 | 2.4606223<br>8 | 0.00346240<br>303 | 3.10E-<br>02 |
| | Mgat5 | -1.29 | 1.6092700<br>54 | 0.02458838<br>171 | 9.03E-<br>02 |
| | Csnk1e | -1.29 | 2.7701286<br>86 | 0.00169774<br>052 | 2.32E-<br>02 |
| | Gm5113 | -1.29 | 2.6632442<br>49 | 0.00217147<br>958 | 2.55E-<br>02 |
| | Atg2b | -1.29 | 2.7232511<br>6 | 0.00189124<br>956 | 2.42E-<br>02 |
| | Cox10 | -1.29 | 3.1753394<br>89 | 0.00066782<br>168 | 1.67E-<br>02 |
| | Gsg1l | -1.29 | 1.7794330<br>78 | 0.01661754<br>722 | 7.22E-<br>02 |
| | Arhgap44 | -1.29 | 3.5036526<br>94 | 0.00031357<br>924 | 1.48E-<br>02 |
| | 5-Sep | -1.29 | 2.2414414<br>16 | 0.00573533<br>228 | 4.01E-<br>02 |

| | | | | | |
|---|---|---|---|---|---|
| | Sptan1 | -1.29 | 2.888369978 | 0.00129309378 | 2.10E-02 |
| | Plxna4 | -1.29 | 2.892121927 | 0.00128197062 | 2.09E-02 |
| | Vps13a | -1.29 | 2.019970626 | 0.00955057180 | 5.26E-02 |
| | Chgb | -1.29 | 2.919320728 | 0.00120414635 | 2.04E-02 |
| | Tecpr2 | -1.29 | 3.019875985 | 0.00095526533 | 1.90E-02 |
| | Ptpn4 | -1.29 | 1.354312359 | 0.04422701627 | 1.30E-01 |
| | Fam219a | -1.29 | 3.461091703 | 0.00034586634 | 1.49E-02 |
| | Prickle1 | -1.29 | 1.457045615 | 0.03491036466 | 1.12E-01 |
| | Ncald | -1.29 | 1.806643146 | 0.01560834495 | 6.99E-02 |
| | Trpc1 | -1.29 | 3.16498237 | 0.00068393941 | 1.68E-02 |
| | Syt3 | -1.29 | 1.87761572 | 0.01325513881 | 6.28E-02 |
| | Map1s | -1.30 | 1.568899182 | 0.02698365767 | 9.61E-02 |
| | Mlxip | -1.30 | 2.528578337 | 0.00296088584 | 2.91E-02 |
| | Mib2 | -1.30 | 2.661790425 | 0.00217876091 | 2.55E-02 |
| | Plch2 | -1.30 | 2.481789 | 0.00329769891 | 3.01E-02 |
| | Akap6 | -1.30 | 1.785174368 | 0.01639931214 | 7.17E-02 |
| | Cend1 | -1.30 | 2.900543177 | 0.00125735184 | 2.07E-02 |
| | Pak1 | -1.30 | 1.755517386 | 0.01755830603 | 7.41E-02 |
| | Fbxw7 | -1.30 | 1.773278062 | 0.01685473536 | 7.29E-02 |
| | Trim46 | -1.30 | 2.67457384 | 0.00211556396 | 2.53E-02 |
| | Adam23 | -1.30 | 1.349789868 | 0.04468997712 | 1.31E-01 |
| | Tbc1d9 | -1.30 | 3.027099204 | 0.00093950868 | 1.89E-02 |
| | Mfsd4a | -1.30 | 1.545056637 | 0.02850646486 | 9.91E-02 |

| | | | | | |
|---|---|---|---|---|---|
| | Igsf8 | -1.30 | 3.153521735 | 0.00070222820 | 1.69E-02 |
| | Kcnh2 | -1.30 | 1.82666267 | 0.01490518361 | 6.78E-02 |
| | Mef2d | -1.30 | 3.175547802 | 0.00066750143 | 1.67E-02 |
| | Atg13 | -1.30 | 4.523939028 | 0.00002992685 | 1.41E-02 |
| | Gga3 | -1.30 | 3.846102525 | 0.00014252711 | 1.41E-02 |
| | Nmnat2 | -1.30 | 5.033582674 | 0.00000925587 | 1.41E-02 |
| | Hmga1 | -1.30 | 1.813665477 | 0.01535799505 | 6.90E-02 |
| | B230217C12Rik | -1.30 | 1.845442208 | 0.01427439767 | 6.58E-02 |
| | Asic1 | -1.30 | 3.967098125 | 0.00010787030 | 1.41E-02 |
| | Sobp | -1.30 | 2.717080564 | 0.00191831285 | 2.43E-02 |
| | Dynll2 | -1.30 | 2.667323264 | 0.00215117992 | 2.55E-02 |
| | Ajap1 | -1.30 | 1.585089661 | 0.02599622811 | 9.37E-02 |
| | Ttl | -1.30 | 1.997224235 | 0.01006411904 | 5.38E-02 |
| | Lyst | -1.30 | 2.950392507 | 0.00112100485 | 2.00E-02 |
| | Fbxo34 | -1.30 | 2.71590988 | 0.00192349083 | 2.43E-02 |
| | Fam219aos | -1.30 | 2.534952026 | 0.00291774930 | 2.89E-02 |
| | Ccser1 | -1.30 | 1.322590044 | 0.04757841344 | 1.37E-01 |
| | Wdfy3 | -1.30 | 3.146651613 | 0.00071342510 | 1.70E-02 |
| | Frat2 | -1.30 | 1.36249053 | 0.04340197276 | 1.29E-01 |
| | Slitrk3 | -1.30 | 3.572797501 | 0.00026742530 | 1.42E-02 |
| | Clec16a | -1.30 | 3.721672761 | 0.00018981356 | 1.41E-02 |
| | 9330102E08Rik | -1.30 | 1.552477267 | 0.02802352306 | 9.83E-02 |
| | Cds1 | -1.30 | 1.475761207 | 0.03343788446 | 1.09E-01 |

| | | | | | |
|---|---|---|---|---|---|
| | Gatad2b | -1.30 | 1.685382499 | 0.02063561901 | 8.10E-02 |
| | Mib1 | -1.30 | 3.070025975 | 0.00085108713 | 1.83E-02 |
| | Svop | -1.30 | 2.221720637 | 0.00600177021 | 4.10E-02 |
| | Cdr1 | -1.30 | 1.446546704 | 0.03576459374 | 1.14E-01 |
| | Klhl34 | -1.30 | 2.174896183 | 0.00668503702 | 4.34E-02 |
| | Dopey2 | -1.30 | 3.328063778 | 0.00046982511 | 1.59E-02 |
| | Kif21a | -1.30 | 2.632158281 | 0.00233260778 | 2.62E-02 |
| | Camk2g | -1.30 | 1.379515426 | 0.04173347743 | 1.25E-01 |
| | Trim37 | -1.30 | 1.677715569 | 0.02100314985 | 8.18E-02 |
| | Gse1 | -1.30 | 2.276147726 | 0.00529483308 | 3.84E-02 |
| | Ankrd13b | -1.30 | 2.094098829 | 0.00805195189 | 4.82E-02 |
| | Trim3 | -1.30 | 3.062929336 | 0.00086510867 | 1.84E-02 |
| | Adora1 | -1.30 | 2.131671878 | 0.00738461948 | 4.60E-02 |
| | Lnx1 | -1.30 | 3.155214536 | 0.00069949637 | 1.69E-02 |
| | Ripor1 | -1.30 | 2.930784989 | 0.00117277584 | 2.03E-02 |
| | Dus3l | -1.30 | 2.179986905 | 0.00660713369 | 4.32E-02 |
| | Efr3b | -1.30 | 4.132516426 | 0.00007370273 | 1.41E-02 |
| | Lrrc4 | -1.30 | 2.129196285 | 0.00742683397 | 4.63E-02 |
| | Arhgap23 | -1.30 | 2.528818438 | 0.00295924936 | 2.91E-02 |
| | Lrrk2 | -1.30 | 2.521042324 | 0.00301271241 | 2.93E-02 |
| | Fbxl2 | -1.30 | 2.249337132 | 0.00563200288 | 3.97E-02 |
| | Ipcef1 | -1.30 | 1.526906529 | 0.02972305677 | 1.01E-01 |
| | Tmem56 | -1.30 | 1.445318087 | 0.03586591488 | 1.14E-01 |

| | | | | | |
|---|---|---|---|---|---|
| | Sstr3 | -1.30 | 2.518330601 | 0.00303158255 | 2.94E-02 |
| | Adgrl1 | -1.30 | 2.527614087 | 0.00296746710 | 2.92E-02 |
| | Ttc7b | -1.30 | 2.415312882 | 0.00384314807 | 3.27E-02 |
| | Ap2a1 | -1.30 | 3.681685591 | 0.00020812028 | 1.41E-02 |
| | Atrn | -1.30 | 3.152990781 | 0.00070308725 | 1.69E-02 |
| | 6530402F18Rik | -1.31 | 1.329947438 | 0.04677917539 | 1.35E-01 |
| | Akap8l | -1.31 | 1.881650481 | 0.01313256379 | 6.25E-02 |
| | Tnfrsf21 | -1.31 | 3.927771913 | 0.00011809407 | 1.41E-02 |
| | C1qtnf4 | -1.31 | 1.437911663 | 0.03648281464 | 1.15E-01 |
| | Atn1 | -1.31 | 2.237744745 | 0.00578435922 | 4.03E-02 |
| | Mark1 | -1.31 | 2.870647355 | 0.00134695363 | 2.12E-02 |
| | Madd | -1.31 | 2.225695728 | 0.00594708673 | 4.08E-02 |
| | Slc8a1 | -1.31 | 1.515014334 | 0.03054820285 | 1.03E-01 |
| | Mapkbp1 | -1.31 | 2.668559446 | 0.00214506548 | 2.55E-02 |
| | Atp6v0a1 | -1.31 | 3.576973264 | 0.00026486632 | 1.42E-02 |
| | Kif3c | -1.31 | 3.54304856 | 0.00028638577 | 1.43E-02 |
| | Zfp106 | -1.31 | 3.926595813 | 0.00011841431 | 1.41E-02 |
| | Pou6f1 | -1.31 | 1.570880433 | 0.02686083857 | 9.59E-02 |
| | Rbfox2 | -1.31 | 3.228823949 | 0.00059044038 | 1.65E-02 |
| | Lrrtm2 | -1.31 | 2.902139732 | 0.00125273805 | 2.07E-02 |
| | Zbtb17 | -1.31 | 2.612272361 | 0.00244189867 | 2.67E-02 |
| | Frmpd3 | -1.31 | 1.623771182 | 0.02378092911 | 8.84E-02 |
| | Rnf157 | -1.31 | 3.180830805 | 0.00065943075 | 1.67E-02 |

| | | | | | |
|---|---|---|---|---|---|
| | Myh10 | -1.31 | 3.707241697 | 0.00019622679 | 1.41E-02 |
| | Mfsd12 | -1.31 | 1.895977462 | 0.01270640045 | 6.13E-02 |
| | Ulk1 | -1.31 | 3.649031098 | 0.00022437213 | 1.42E-02 |
| | Gls | -1.31 | 3.058891877 | 0.00087318873 | 1.85E-02 |
| | Pclo | -1.31 | 1.450996364 | 0.03540003045 | 1.13E-01 |
| | Raph1 | -1.31 | 1.90773443 | 0.01236703442 | 6.03E-02 |
| | Unc80 | -1.31 | 2.29199047 | 0.00510516203 | 3.75E-02 |
| | Hlf | -1.31 | 1.334139216 | 0.04632983822 | 1.34E-01 |
| | Scn2a | -1.31 | 2.372012765 | 0.00424607084 | 3.41E-02 |
| | Mapre3 | -1.31 | 3.42519096 | 0.00037567218 | 1.52E-02 |
| | Ppme1 | -1.31 | 2.214828827 | 0.00609777188 | 4.13E-02 |
| | Brinp2 | -1.31 | 2.76601276 | 0.00171390695 | 2.33E-02 |
| | Fam189b | -1.31 | 3.226341888 | 0.00059382450 | 1.65E-02 |
| | Bcl9 | -1.31 | 3.003519848 | 0.00099192800 | 1.92E-02 |
| | Ttll5 | -1.31 | 2.334870679 | 0.00462518727 | 3.56E-02 |
| | Sbf1 | -1.31 | 2.56840157 | 0.00270145931 | 2.79E-02 |
| | Dlgap1 | -1.31 | 2.892651349 | 0.00128040880 | 2.09E-02 |
| | Rnf208 | -1.31 | 1.820357099 | 0.01512317232 | 6.84E-02 |
| | Pex14 | -1.31 | 3.697517158 | 0.00020067018 | 1.41E-02 |
| | Shisa7 | -1.31 | 2.509452339 | 0.00309419487 | 2.96E-02 |
| | Dctn1 | -1.31 | 4.406042565 | 0.00003926065 | 1.41E-02 |
| | 5730409E04Rik | -1.31 | 2.208280362 | 0.00619041320 | 4.16E-02 |
| | Mansc1 | -1.31 | 1.464915149 | 0.03428347618 | 1.11E-01 |

| | | | | | |
|---|---|---|---|---|---|
| | Lmbrd2 | -1.31 | 2.243319279 | 0.00571058659 | 4.00E-02 |
| | St3gal5 | -1.31 | 3.543403377 | 0.00028615189 | 1.43E-02 |
| | Slx4 | -1.31 | 2.306924128 | 0.00493259969 | 3.68E-02 |
| | Nrg3 | -1.31 | 1.915448562 | 0.01214930509 | 5.96E-02 |
| | Caln1 | -1.31 | 2.103034005 | 0.00788798352 | 4.77E-02 |
| | Rbfox1 | -1.31 | 1.92384192 | 0.01191675691 | 5.91E-02 |
| | Tigar | -1.31 | 1.988110923 | 0.01027753767 | 5.44E-02 |
| | Crtc1 | -1.31 | 2.888641864 | 0.00129228450 | 2.10E-02 |
| | Ubtd2 | -1.31 | 1.959721234 | 0.01097182231 | 5.64E-02 |
| | Capn15 | -1.31 | 2.374516554 | 0.00422166187 | 3.41E-02 |
| | 3110035E14Rik | -1.31 | 1.724510451 | 0.01885773588 | 7.69E-02 |
| | Smg7 | -1.31 | 3.430431123 | 0.00037116659 | 1.52E-02 |
| | Mroh1 | -1.31 | 2.355870635 | 0.00440686113 | 3.47E-02 |
| | Rap1gds1 | -1.31 | 2.512396622 | 0.00307328884 | 2.94E-02 |
| | Slc39a10 | -1.31 | 2.752396198 | 0.00176849486 | 2.36E-02 |
| | Bcl7a | -1.31 | 3.43534639 | 0.00036698948 | 1.52E-02 |
| | Sipa1l1 | -1.31 | 2.748232967 | 0.00178552951 | 2.36E-02 |
| | Chst1 | -1.31 | 3.088873413 | 0.00081494179 | 1.79E-02 |
| | Rock2 | -1.31 | 3.373936611 | 0.00042273031 | 1.58E-02 |
| | Uhmk1 | -1.31 | 2.492146991 | 0.00321997878 | 3.00E-02 |
| | Abcc5 | -1.31 | 3.259133107 | 0.00055063891 | 1.62E-02 |
| | Slc16a7 | -1.31 | 1.727824326 | 0.01871438992 | 7.67E-02 |
| | Csrnp2 | -1.31 | 2.565060974 | 0.00272231907 | 2.81E-02 |

| | | | | | |
|---|---|---|---|---|---|
| | Reep2 | -1.31 | 2.526449373 | 0.00297543609 | 2.92E-02 |
| | Lrrc7 | -1.31 | 2.836315742 | 0.00145775406 | 2.19E-02 |
| | Il1rapl1 | -1.31 | 1.352598126 | 0.04440193269 | 1.31E-01 |
| | Herc2 | -1.31 | 3.084418886 | 0.00082334360 | 1.80E-02 |
| | Mapk8ip2 | -1.32 | 3.917450691 | 0.00012093425 | 1.41E-02 |
| | L1cam | -1.32 | 2.31567546 | 0.00483419918 | 3.64E-02 |
| | Tuba8 | -1.32 | 1.877890543 | 0.01324675358 | 6.28E-02 |
| | Arfgap1 | -1.32 | 2.574171663 | 0.00266580474 | 2.77E-02 |
| | Ankrd33b | -1.32 | 1.314068395 | 0.04852120800 | 1.39E-01 |
| | Ppp1r12b | -1.32 | 2.086506144 | 0.00819396031 | 4.88E-02 |
| | Rab11fip3 | -1.32 | 3.564835818 | 0.00027237308 | 1.42E-02 |
| | Sort1 | -1.32 | 2.385447856 | 0.00411672772 | 3.36E-02 |
| | Ttll11 | -1.32 | 1.545213436 | 0.02849617465 | 9.91E-02 |
| | Tubgcp6 | -1.32 | 2.98274214 | 0.00104053780 | 1.97E-02 |
| | Mlip | -1.32 | 1.315434905 | 0.04836877575 | 1.38E-01 |
| | Rgag4 | -1.32 | 2.338017023 | 0.00459180014 | 3.54E-02 |
| | Med14 | -1.32 | 3.994193901 | 0.00010134588 | 1.41E-02 |
| | Kcnab2 | -1.32 | 1.948226502 | 0.01126609731 | 5.72E-02 |
| | Mb21d2 | -1.32 | 1.686158001 | 0.02059880371 | 8.09E-02 |
| | Zfp526 | -1.32 | 1.344314914 | 0.04525692950 | 1.32E-01 |
| | Agap2 | -1.32 | 2.479406982 | 0.00331583581 | 3.02E-02 |
| | Elk4 | -1.32 | 1.944971982 | 0.01135084041 | 5.74E-02 |
| | Ankrd13d | -1.32 | 2.18987202 | 0.00645844522 | 4.26E-02 |

| | | | | | |
|---|---|---|---|---|---|
| | Spata2 | -1.32 | 3.8349969 1 | 0.00014621 876 | 1.41E-02 |
| | Zfyve28 | -1.32 | 4.0647613 78 | 0.00008614 670 | 1.41E-02 |
| | Iqsec2 | -1.32 | 2.2762802 61 | 0.00529321 749 | 3.84E-02 |
| | Mapk8ip3 | -1.32 | 2.7475174 85 | 0.00178847 352 | 2.36E-02 |
| | 1700020I14Rik | -1.32 | 2.3172466 32 | 0.00481674 182 | 3.64E-02 |
| | Btbd9 | -1.32 | 1.3866655 | 0.04105201 700 | 1.24E-01 |
| | Add2 | -1.32 | 3.8171602 65 | 0.00015234 904 | 1.41E-02 |
| | Lsm11 | -1.32 | 1.3017175 39 | 0.04992090 624 | 1.41E-01 |
| | Ubl7 | -1.32 | 2.5477104 38 | 0.00283328 043 | 2.87E-02 |
| | Lrfn4 | -1.32 | 3.0792062 2 | 0.00083328 541 | 1.81E-02 |
| | Slc36a4 | -1.32 | 1.7950514 26 | 0.01603055 558 | 7.09E-02 |
| | Zfp369 | -1.32 | 1.3823239 63 | 0.04146446 223 | 1.25E-01 |
| | Kcnq5 | -1.32 | 2.0783426 96 | 0.00834943 915 | 4.92E-02 |
| | Stk32c | -1.32 | 2.0184852 9 | 0.00958329 176 | 5.26E-02 |
| | Rap1gap2 | -1.32 | 1.9571726 26 | 0.01103639 851 | 5.65E-02 |
| | Hcn1 | -1.32 | 1.3356976 78 | 0.04616388 198 | 1.34E-01 |
| | Fam193b | -1.32 | 1.4894905 76 | 0.03239734 522 | 1.07E-01 |
| | Camta1 | -1.32 | 1.8167711 1 | 0.01524856 202 | 6.88E-02 |
| | Kcnh3 | -1.32 | 1.9996052 74 | 0.01000909 303 | 5.37E-02 |
| | Myo5a | -1.32 | 1.8839291 87 | 0.01306383 879 | 6.24E-02 |
| | Rpusd1 | -1.32 | 1.8064020 3 | 0.01561701 293 | 6.99E-02 |
| | Ttc39b | -1.32 | 2.0266702 34 | 0.00940437 126 | 5.22E-02 |
| | St8sia5 | -1.32 | 1.4020464 57 | 0.03962356 461 | 1.21E-01 |

| | | | | | |
|---|---|---|---|---|---|
| | Ankrd52 | -1.32 | 3.008499412 | 0.00098061964 | 1.92E-02 |
| | Calm3 | -1.32 | 2.94871532 | 0.00112534239 | 2.00E-02 |
| | Fam189a1 | -1.32 | 2.013775597 | 0.00968778301 | 5.29E-02 |
| | Dync1h1 | -1.32 | 3.191931607 | 0.00064278894 | 1.65E-02 |
| | Usp31 | -1.32 | 3.092789085 | 0.00080762716 | 1.79E-02 |
| | Inha | -1.32 | 1.969771993 | 0.01072082007 | 5.56E-02 |
| | Unc13b | -1.32 | 2.843513455 | 0.00143379329 | 2.17E-02 |
| | Magi2 | -1.32 | 2.456309075 | 0.00349696210 | 3.11E-02 |
| | Crispld1 | -1.32 | 1.351541676 | 0.04451007482 | 1.31E-01 |
| | Nck2 | -1.32 | 2.375001405 | 0.00421695139 | 3.40E-02 |
| | Dvl3 | -1.32 | 2.457050076 | 0.00349100060 | 3.11E-02 |
| | Rexo1 | -1.32 | 3.023675023 | 0.00094694548 | 1.89E-02 |
| | Diexf | -1.32 | 2.69542877 | 0.00201637466 | 2.48E-02 |
| | Thy1 | -1.33 | 2.185187564 | 0.00652848538 | 4.28E-02 |
| | Camsap1 | -1.33 | 4.945116497 | 0.00001134706 | 1.41E-02 |
| | Eif4g3 | -1.33 | 2.652208394 | 0.00222736610 | 2.59E-02 |
| | Pcdh9 | -1.33 | 2.075293282 | 0.00840827134 | 4.93E-02 |
| | Nckipsd | -1.33 | 2.546111926 | 0.00284372813 | 2.87E-02 |
| | Ccdc9 | -1.33 | 3.046654725 | 0.00089814256 | 1.86E-02 |
| | Braf | -1.33 | 2.18043435 | 0.00660033000 | 4.32E-02 |
| | Ano8 | -1.33 | 3.854133539 | 0.00013991570 | 1.41E-02 |
| | Prkcb | -1.33 | 1.888256028 | 0.01293433104 | 6.19E-02 |
| | Zbed6 | -1.33 | 1.884371313 | 0.01305054615 | 6.24E-02 |

| | | | | | |
|---|---|---|---|---|---|
| | Klf9 | -1.33 | 3.076435854 | 0.00083861794 | 1.81E-02 |
| | Acvr1b | -1.33 | 3.949289637 | 0.00011238552 | 1.41E-02 |
| | Thrb | -1.33 | 2.281296876 | 0.00523242634 | 3.81E-02 |
| | Rab11fip5 | -1.33 | 2.355010485 | 0.00441559787 | 3.47E-02 |
| | Tecpr1 | -1.33 | 3.775550762 | 0.00016766763 | 1.41E-02 |
| | Tnk2 | -1.33 | 2.503998541 | 0.00313329625 | 2.97E-02 |
| | Tmem191c | -1.33 | 1.762202129 | 0.01729011457 | 7.36E-02 |
| | Ago2 | -1.33 | 1.733126597 | 0.01848729635 | 7.62E-02 |
| | Plbd2 | -1.33 | 2.369618642 | 0.00426954267 | 3.42E-02 |
| | Nfic | -1.33 | 1.639155118 | 0.02295328675 | 8.63E-02 |
| | Prepl | -1.33 | 2.878451899 | 0.00132296423 | 2.11E-02 |
| | Ints1 | -1.33 | 2.935779133 | 0.00115936682 | 2.03E-02 |
| | Dmxl2 | -1.33 | 2.651736563 | 0.00222978730 | 2.59E-02 |
| | Atcay | -1.33 | 3.698069085 | 0.00020041532 | 1.41E-02 |
| | Vps18 | -1.33 | 4.095714711 | 0.00008022049 | 1.41E-02 |
| | Ptprs | -1.33 | 4.013974444 | 0.00009683348 | 1.41E-02 |
| | Mchr1 | -1.33 | 1.669751648 | 0.02139185036 | 8.25E-02 |
| | Igsf9 | -1.33 | 1.499175338 | 0.03168288071 | 1.06E-01 |
| | Itpr1 | -1.33 | 2.228585428 | 0.00590764747 | 4.08E-02 |
| | Myt1l | -1.33 | 4.515068899 | 0.00003054437 | 1.41E-02 |
| | Osbpl6 | -1.33 | 1.930197333 | 0.01174363831 | 5.86E-02 |
| | Nefl | -1.33 | 2.326673481 | 0.00471331558 | 3.60E-02 |
| | Rita1 | -1.33 | 2.929052842 | 0.00117746270 | 2.03E-02 |

| | | | | | |
|---|---|---|---|---|---|
| | Ppip5k1 | -1.33 | 3.282215499 | 0.00052213704 | 1.60E-02 |
| | Prr36 | -1.33 | 1.801470638 | 0.01579535394 | 7.03E-02 |
| | Lrp3 | -1.33 | 2.832267161 | 0.00147140707 | 2.19E-02 |
| | Herc3 | -1.33 | 1.70861784 | 0.01956059945 | 7.85E-02 |
| | Kcnq2 | -1.33 | 2.696331936 | 0.00201218573 | 2.48E-02 |
| | Dok6 | -1.33 | 1.465312225 | 0.03425214507 | 1.11E-01 |
| | Syn1 | -1.33 | 2.41354611 | 0.00385881440 | 3.27E-02 |
| | Bmp3 | -1.33 | 1.56923242 | 0.02696296079 | 9.61E-02 |
| | Acap3 | -1.33 | 2.218225311 | 0.00605026906 | 4.11E-02 |
| | Rsad1 | -1.33 | 1.62563095 | 0.02367931039 | 8.82E-02 |
| | Lhx6 | -1.33 | 1.858216583 | 0.01386064423 | 6.45E-02 |
| | Aak1 | -1.33 | 2.850780879 | 0.00141000003 | 2.17E-02 |
| | Ppp1r37 | -1.33 | 2.69912451 | 0.00199928860 | 2.47E-02 |
| | Slc25a27 | -1.33 | 2.851427573 | 0.00140790200 | 2.17E-02 |
| | Rab3a | -1.33 | 3.455392285 | 0.00035043519 | 1.49E-02 |
| | Elfn2 | -1.34 | 2.210729198 | 0.00615560582 | 4.15E-02 |
| | Rims2 | -1.34 | 1.602441231 | 0.02497806369 | 9.12E-02 |
| | Cntnap5a | -1.34 | 1.718983971 | 0.01909923749 | 7.74E-02 |
| | Zmiz2 | -1.34 | 2.602265031 | 0.00249881997 | 2.70E-02 |
| | Syna | -1.34 | 1.583990257 | 0.02606212015 | 9.38E-02 |
| | Lzts1 | -1.34 | 1.755489952 | 0.01755941518 | 7.41E-02 |
| | Rhov | -1.34 | 1.402368246 | 0.03959421659 | 1.21E-01 |
| | Aifm3 | -1.34 | 1.373307297 | 0.04233433121 | 1.27E-01 |

| | | | | | |
|---|---|---|---|---|---|
| | Dscam | -1.34 | 3.270571865 | 0.00053632512 | 1.61E-02 |
| | Sptbn2 | -1.34 | 1.81743752 | 0.01522518155 | 6.88E-02 |
| | Cpeb3 | -1.34 | 3.092381922 | 0.00080838469 | 1.79E-02 |
| | Galntl6 | -1.34 | 1.761888114 | 0.01730262065 | 7.36E-02 |
| | Prrg2 | -1.34 | 1.428880568 | 0.03724941290 | 1.17E-01 |
| | Cdk5r1 | -1.34 | 3.708991049 | 0.00019543797 | 1.41E-02 |
| | Map1a | -1.34 | 2.125119575 | 0.00749687769 | 4.64E-02 |
| | Hint3 | -1.34 | 2.492604117 | 0.00321659130 | 3.00E-02 |
| | Dpp9 | -1.34 | 4.489896724 | 0.00003236706 | 1.41E-02 |
| | Smg9 | -1.34 | 3.113151125 | 0.00077063526 | 1.76E-02 |
| | Prkar1b | -1.34 | 2.270604856 | 0.00536284375 | 3.88E-02 |
| | Dnajb5 | -1.34 | 1.944703031 | 0.01135787197 | 5.74E-02 |
| | Ddhd2 | -1.34 | 3.381647799 | 0.00041529070 | 1.57E-02 |
| | Cntn2 | -1.34 | 1.921124621 | 0.01199155156 | 5.93E-02 |
| | Mast4 | -1.34 | 2.079446807 | 0.00832823925 | 4.91E-02 |
| | Il17ra | -1.34 | 1.677672491 | 0.02100523329 | 8.18E-02 |
| | Nfix | -1.34 | 1.557416371 | 0.02770662513 | 9.77E-02 |
| | Trio | -1.34 | 4.24025902 | 0.00005750968 | 1.41E-02 |
| | H1fx | -1.34 | 1.437042986 | 0.03655586073 | 1.15E-01 |
| | Sh3gl2 | -1.34 | 2.424824658 | 0.00375989176 | 3.24E-02 |
| | Pcdhgc4 | -1.34 | 1.548410902 | 0.02828714380 | 9.87E-02 |
| | Shank3 | -1.34 | 2.358713308 | 0.00437811023 | 3.47E-02 |
| | Mpp2 | -1.34 | 2.483124051 | 0.00328757712 | 3.01E-02 |

| | | | | | |
|---|---|---|---|---|---|
| | Slc25a22 | -1.34 | 3.056917934 | 0.00087716656 | 1.85E-02 |
| | Nphp4 | -1.34 | 1.765700861 | 0.01715138274 | 7.34E-02 |
| | Fam43b | -1.34 | 1.640931223 | 0.02285960791 | 8.61E-02 |
| | Mafg | -1.34 | 2.223518061 | 0.00597698187 | 4.09E-02 |
| | Hivep3 | -1.34 | 1.593701954 | 0.02548578683 | 9.26E-02 |
| | Map3k12 | -1.34 | 3.400746104 | 0.00039742382 | 1.55E-02 |
| | Dlc1 | -1.34 | 2.326263427 | 0.00471776792 | 3.60E-02 |
| | Dab2ip | -1.34 | 2.267321005 | 0.00540354777 | 3.89E-02 |
| | Zfp653 | -1.34 | 2.031611181 | 0.00929798453 | 5.20E-02 |
| | Cdh7 | -1.34 | 1.78403765 | 0.01644229176 | 7.18E-02 |
| | Rph3a | -1.34 | 1.430399666 | 0.03711934756 | 1.16E-01 |
| | Spock2 | -1.34 | 4.723657324 | 0.00001889482 | 1.41E-02 |
| | Syt1 | -1.34 | 2.239286956 | 0.00576385496 | 4.03E-02 |
| | Srrm2 | -1.34 | 2.102970352 | 0.00788913973 | 4.77E-02 |
| | Ywhag | -1.34 | 3.41624072 | 0.00038349462 | 1.52E-02 |
| | Slc35f3 | -1.34 | 2.739450903 | 0.00182200304 | 2.38E-02 |
| | Reep6 | -1.34 | 1.306864913 | 0.04933272291 | 1.40E-01 |
| | Scn2b | -1.34 | 2.981632862 | 0.00104319894 | 1.97E-02 |
| | Ccsap | -1.34 | 2.557850116 | 0.00276789674 | 2.84E-02 |
| | Irgq | -1.34 | 3.047264312 | 0.00089688278 | 1.86E-02 |
| | Pcdhgc5 | -1.34 | 3.034469579 | 0.00092369889 | 1.89E-02 |
| | Snap25 | -1.34 | 1.451257935 | 0.03537871586 | 1.13E-01 |
| | Faah | -1.34 | 2.147642447 | 0.00711799294 | 4.51E-02 |

| | | | | | |
|---|---|---|---|---|---|
| | Coro2b | -1.34 | 3.0081826<br>21 | 0.00098133<br>520 | 1.92E-<br>02 |
| | Zfp169 | -1.34 | 1.4569486<br>41 | 0.03491816<br>069 | 1.12E-<br>01 |
| | Mapk6 | -1.34 | 1.6706153<br>85 | 0.02134934<br>795 | 8.25E-<br>02 |
| | Auts2 | -1.34 | 3.1264189<br>76 | 0.00074744<br>807 | 1.74E-<br>02 |
| | Atg9a | -1.35 | 4.1291255<br>67 | 0.00007428<br>043 | 1.41E-<br>02 |
| | Grik4 | -1.35 | 2.0976376<br>43 | 0.00798660<br>778 | 4.80E-<br>02 |
| | Rapgef5 | -1.35 | 3.5330178<br>5 | 0.00029307<br>728 | 1.43E-<br>02 |
| | Mmp17 | -1.35 | 1.9475268<br>62 | 0.01128426<br>139 | 5.72E-<br>02 |
| | Syne1 | -1.35 | 2.5862283<br>01 | 0.00259281<br>601 | 2.74E-<br>02 |
| | Tmem59l | -1.35 | 1.4366483<br>14 | 0.03658909<br>652 | 1.15E-<br>01 |
| | Fzd3 | -1.35 | 1.5165708<br>1 | 0.03043891<br>658 | 1.03E-<br>01 |
| | Map6 | -1.35 | 4.0127251<br>07 | 0.00009711<br>245 | 1.41E-<br>02 |
| | Htt | -1.35 | 3.9411048<br>56 | 0.00011452<br>364 | 1.41E-<br>02 |
| | Rasgrf2 | -1.35 | 1.4447156<br>73 | 0.03591569<br>932 | 1.14E-<br>01 |
| | Syngr1 | -1.35 | 3.3767194<br>01 | 0.00042003<br>028 | 1.58E-<br>02 |
| | Mpped1 | -1.35 | 2.2311330<br>03 | 0.00587309<br>461 | 4.06E-<br>02 |
| | 9430015G10<br>Rik | -1.35 | 1.6853740<br>42 | 0.02063602<br>086 | 8.10E-<br>02 |
| | Dlgap4 | -1.35 | 2.1154304<br>63 | 0.00766601<br>275 | 4.72E-<br>02 |
| | Zdhhc14 | -1.35 | 1.9508412<br>44 | 0.01119847<br>167 | 5.70E-<br>02 |
| | Aatk | -1.35 | 3.1208010<br>7 | 0.00075717<br>964 | 1.74E-<br>02 |
| | Alpl | -1.35 | 1.4753849<br>19 | 0.03346686<br>881 | 1.09E-<br>01 |
| | Fgf9 | -1.35 | 1.7620296<br>28 | 0.01729698<br>355 | 7.36E-<br>02 |
| | Gpr162 | -1.35 | 2.8026255<br>13 | 0.00157534<br>068 | 2.26E-<br>02 |

| | | | | | |
|---|---|---|---|---|---|
| | Adgra1 | -1.35 | 2.5736505 82 | 0.00266900 519 | 2.77E-02 |
| | Sema6b | -1.35 | 3.8121123 05 | 0.00015413 018 | 1.41E-02 |
| | Bean1 | -1.35 | 1.6055129 23 | 0.02480202 132 | 9.08E-02 |
| | Sv2a | -1.35 | 3.7905489 64 | 0.00016197 614 | 1.41E-02 |
| | Actr1b | -1.35 | 2.7591687 65 | 0.00174113 014 | 2.34E-02 |
| | Tmem240 | -1.35 | 2.2043635 69 | 0.00624649 549 | 4.18E-02 |
| | Map7d2 | -1.35 | 2.7205326 55 | 0.00190312 513 | 2.43E-02 |
| | Phlpp2 | -1.35 | 3.2993731 12 | 0.00050191 120 | 1.59E-02 |
| | Fry | -1.35 | 3.4968642 07 | 0.00031851 933 | 1.48E-02 |
| | Iqsec1 | -1.35 | 2.3534706 62 | 0.00443128 149 | 3.47E-02 |
| | Hspa12a | -1.35 | 2.0457161 81 | 0.00900085 612 | 5.11E-02 |
| | Wnt4 | -1.35 | 1.4943107 82 | 0.03203975 730 | 1.06E-01 |
| | Zfp871 | -1.35 | 1.8619942 14 | 0.01374060 282 | 6.41E-02 |
| | Ccdc177 | -1.35 | 2.0216655 26 | 0.00951337 188 | 5.25E-02 |
| | Ubald1 | -1.35 | 2.6797528 39 | 0.00209048 551 | 2.51E-02 |
| | Slc30a3 | -1.35 | 1.3972625 49 | 0.04006244 501 | 1.22E-01 |
| | Ercc8 | -1.35 | 1.9499260 13 | 0.01122209 619 | 5.70E-02 |
| | Rab6b | -1.35 | 2.6543017 42 | 0.00221665 578 | 2.59E-02 |
| | Pi4ka | -1.36 | 3.7662939 26 | 0.00017127 977 | 1.41E-02 |
| | Fgf14 | -1.36 | 2.2454250 2 | 0.00568296 498 | 3.99E-02 |
| | Zfp692 | -1.36 | 1.3386522 9 | 0.04585088 368 | 1.33E-01 |
| | Sult4a1 | -1.36 | 2.6553546 49 | 0.00221128 821 | 2.58E-02 |
| | R3hdm2 | -1.36 | 2.2970612 31 | 0.00504590 151 | 3.73E-02 |

| | | | | | |
|---|---|---|---|---|---|
| | Fam217b | -1.36 | 2.578085744 | 0.00264188711 | 2.77E-02 |
| | Cabp1 | -1.36 | 1.500390533 | 0.03159435309 | 1.06E-01 |
| | Ccdc92b | -1.36 | 3.309720658 | 0.00049009395 | 1.59E-02 |
| | Inpp4a | -1.36 | 2.870652977 | 0.00134693619 | 2.12E-02 |
| | Fbxo31 | -1.36 | 2.383087151 | 0.00413916605 | 3.37E-02 |
| | Eef1a2 | -1.36 | 2.796173336 | 0.00159891974 | 2.27E-02 |
| | Hmgxb3 | -1.36 | 4.550039215 | 0.00002818128 | 1.41E-02 |
| | Rhobtb2 | -1.36 | 3.390629859 | 0.00040678988 | 1.56E-02 |
| | Hrh3 | -1.36 | 1.996631903 | 0.01007785481 | 5.38E-02 |
| | Slitrk1 | -1.36 | 3.156352074 | 0.00069766659 | 1.69E-02 |
| | Osbpl10 | -1.36 | 2.357094513 | 0.00439445971 | 3.47E-02 |
| | Nrip1 | -1.36 | 1.817781952 | 0.01521311150 | 6.87E-02 |
| | Pde4d | -1.36 | 2.047539094 | 0.00896315499 | 5.10E-02 |
| | Amer3 | -1.36 | 3.311379683 | 0.00048822534 | 1.59E-02 |
| | Prickle2 | -1.36 | 3.34123892 | 0.00045578610 | 1.59E-02 |
| | Kcnv1 | -1.36 | 2.282602107 | 0.00521672439 | 3.81E-02 |
| | Klhdc3 | -1.36 | 3.264884859 | 0.00054339438 | 1.62E-02 |
| | Nbea | -1.36 | 2.94618777 | 0.00113191087 | 2.00E-02 |
| | Kcnt1 | -1.36 | 2.996200736 | 0.00100878650 | 1.93E-02 |
| | Prrt1 | -1.36 | 1.670510918 | 0.02135448400 | 8.25E-02 |
| | Chrm4 | -1.36 | 2.650913857 | 0.00223401530 | 2.59E-02 |
| | Plxna1 | -1.36 | 1.787057529 | 0.01632835639 | 7.16E-02 |
| | Ccm2 | -1.36 | 1.340064187 | 0.04570206389 | 1.33E-01 |

| | | | | | |
|---|---|---|---|---|---|
| | Epb41l4b | -1.36 | 1.42466642 | 0.03761261948 | 1.17E-01 |
| | Hid1 | -1.36 | 3.208245699 | 0.00061909073 | 1.65E-02 |
| | Zkscan16 | -1.36 | 2.055036887 | 0.00880974043 | 5.03E-02 |
| | Gm15506 | -1.36 | 1.559663743 | 0.02756362018 | 9.74E-02 |
| | 5330417C22Rik | -1.36 | 3.205584286 | 0.00062289625 | 1.65E-02 |
| | Cacng3 | -1.36 | 1.798694227 | 0.01589665588 | 7.05E-02 |
| | Dkkl1 | -1.36 | 1.351922205 | 0.04447109214 | 1.31E-01 |
| | Gls2 | -1.36 | 1.49949588 | 0.03165950501 | 1.06E-01 |
| | Basp1 | -1.36 | 3.585432063 | 0.00025975741 | 1.42E-02 |
| | Cecr6 | -1.36 | 2.569315574 | 0.00269577987 | 2.79E-02 |
| | Scn1b | -1.36 | 1.423594012 | 0.03770561145 | 1.18E-01 |
| | Etv5 | -1.36 | 3.311960502 | 0.00048757283 | 1.59E-02 |
| | Synj1 | -1.36 | 4.047758119 | 0.00008958636 | 1.41E-02 |
| | Rab11fip4 | -1.37 | 3.602233771 | 0.00024989998 | 1.42E-02 |
| | Astn2 | -1.37 | 1.749616001 | 0.01779852444 | 7.47E-02 |
| | Dlgap3 | -1.37 | 2.082788236 | 0.00826440827 | 4.90E-02 |
| | Cntnap1 | -1.37 | 2.581492595 | 0.00262124373 | 2.76E-02 |
| | Arhgap32 | -1.37 | 2.932109242 | 0.00116920525 | 2.03E-02 |
| | Dnajb14 | -1.37 | 2.717031259 | 0.00191853065 | 2.43E-02 |
| | Cap2 | -1.37 | 3.018780033 | 0.00095767901 | 1.90E-02 |
| | Nell2 | -1.37 | 3.279176933 | 0.00052580301 | 1.61E-02 |
| | Kcnc2 | -1.37 | 2.023992946 | 0.00946252531 | 5.24E-02 |
| | Pgbd5 | -1.37 | 3.554433059 | 0.00027897606 | 1.43E-02 |

| | | | | | |
|---|---|---|---|---|---|
| | Pitpnm2 | -1.37 | 3.030988696 | 0.00093113211 | 1.89E-02 |
| | Gatsl2 | -1.37 | 2.9455024 | 0.00113369857 | 2.00E-02 |
| | Npcd | -1.37 | 1.631497327 | 0.02336160482 | 8.75E-02 |
| | Anks6 | -1.37 | 1.507327085 | 0.03109373657 | 1.05E-01 |
| | Adam11 | -1.37 | 2.191644224 | 0.00643214426 | 4.25E-02 |
| | Mkl2 | -1.37 | 3.523698575 | 0.00029943422 | 1.45E-02 |
| | Ube2ql1 | -1.37 | 2.740266411 | 0.00181858493 | 2.38E-02 |
| | Tanc2 | -1.37 | 2.403471411 | 0.00394937697 | 3.30E-02 |
| | Chsy3 | -1.37 | 1.757820553 | 0.01746543660 | 7.39E-02 |
| | Rai1 | -1.37 | 3.465978075 | 0.00034199671 | 1.49E-02 |
| | Unc79 | -1.37 | 3.814710317 | 0.00015321091 | 1.41E-02 |
| | Arhgap26 | -1.37 | 2.393934181 | 0.00403706571 | 3.34E-02 |
| | Rps6kc1 | -1.37 | 4.796111004 | 0.00001599149 | 1.41E-02 |
| | Slc7a8 | -1.37 | 3.198926827 | 0.00063251841 | 1.65E-02 |
| | Lrfn5 | -1.37 | 3.030961385 | 0.00093119067 | 1.89E-02 |
| | Rusc2 | -1.37 | 4.366696358 | 0.00004298368 | 1.41E-02 |
| | Sez6l2 | -1.37 | 2.413353681 | 0.00386052456 | 3.27E-02 |
| | Luzp1 | -1.37 | 2.617569938 | 0.00241229303 | 2.66E-02 |
| | Epha4 | -1.37 | 2.185404548 | 0.00652522441 | 4.28E-02 |
| | Fam78b | -1.37 | 3.056725954 | 0.00087755440 | 1.85E-02 |
| | Dgkz | -1.37 | 1.933054133 | 0.01166664189 | 5.84E-02 |
| | Plppr2 | -1.37 | 2.610040889 | 0.00245447782 | 2.68E-02 |
| | Fbxl19 | -1.37 | 3.529760867 | 0.00029528347 | 1.44E-02 |

| | | | | | |
|---|---|---|---|---|---|
| | Tnip1 | -1.38 | 2.56815603 | 0.00270298708 | 2.79E-02 |
| | Nrn1 | -1.38 | 1.491293046 | 0.03226316391 | 1.07E-01 |
| | Mn1 | -1.38 | 3.194748261 | 0.00063863356 | 1.65E-02 |
| | Slc7a14 | -1.38 | 4.020302497 | 0.00009543276 | 1.41E-02 |
| | Slc4a8 | -1.38 | 4.315805676 | 0.00004832750 | 1.41E-02 |
| | Abcb9 | -1.38 | 2.46571777 | 0.00342201753 | 3.09E-02 |
| | Slc7a4 | -1.38 | 2.279117597 | 0.00525874852 | 3.83E-02 |
| | Speg | -1.38 | 3.405375545 | 0.00039320991 | 1.54E-02 |
| | Sigmar1 | -1.38 | 2.747153607 | 0.00178997264 | 2.36E-02 |
| | Celf5 | -1.38 | 2.959624929 | 0.00109742556 | 2.00E-02 |
| | Gpr45 | -1.38 | 1.760102204 | 0.01737391916 | 7.37E-02 |
| | Camk2n2 | -1.38 | 1.816515466 | 0.01525754060 | 6.88E-02 |
| | Sytl2 | -1.38 | 1.488871959 | 0.03244352547 | 1.07E-01 |
| | Tle2 | -1.38 | 1.336571933 | 0.04607104545 | 1.34E-01 |
| | Ephb3 | -1.38 | 1.780141451 | 0.01659046465 | 7.22E-02 |
| | Pom121 | -1.38 | 4.353555494 | 0.00004430416 | 1.41E-02 |
| | Ctif | -1.38 | 3.260441317 | 0.00054898273 | 1.62E-02 |
| | Flywch1 | -1.38 | 2.106515913 | 0.00782499533 | 4.77E-02 |
| | Trmt44 | -1.38 | 1.320836483 | 0.04777091030 | 1.37E-01 |
| | Atxn7l2 | -1.38 | 1.533808013 | 0.02925445333 | 1.01E-01 |
| | Scrt2 | -1.38 | 1.8942623 | 0.01275668112 | 6.14E-02 |
| | Csmd2 | -1.38 | 3.206857412 | 0.00062107291 | 1.65E-02 |
| | Cacna1b | -1.38 | 2.33697441 | 0.00460283694 | 3.55E-02 |

| | | | | | |
|---|---|---|---|---|---|
| | Ciart | -1.38 | 1.812413128 | 0.01540234587 | 6.91E-02 |
| | Ttc9 | -1.38 | 3.429896891 | 0.00037162345 | 1.52E-02 |
| | Abcg4 | -1.38 | 4.298488302 | 0.00005029348 | 1.41E-02 |
| | Hdac4 | -1.38 | 3.205699051 | 0.00062273166 | 1.65E-02 |
| | Prr14l | -1.38 | 2.895398271 | 0.00127233575 | 2.08E-02 |
| | Tnks1bp1 | -1.38 | 2.126581682 | 0.00747168093 | 4.64E-02 |
| | Ppm1l | -1.38 | 2.304303457 | 0.00496245456 | 3.70E-02 |
| | Rnf165 | -1.38 | 1.469946154 | 0.03388861706 | 1.10E-01 |
| | Cadm3 | -1.38 | 4.091355618 | 0.00008102973 | 1.41E-02 |
| | Zmat4 | -1.38 | 1.352252627 | 0.04443727033 | 1.31E-01 |
| | Gas7 | -1.38 | 3.187379717 | 0.00064956151 | 1.65E-02 |
| | Slc17a7 | -1.38 | 2.772404144 | 0.00168886858 | 2.32E-02 |
| | Ank1 | -1.38 | 1.399076516 | 0.03989546068 | 1.22E-01 |
| | Adgrb2 | -1.38 | 2.687929991 | 0.00205149286 | 2.50E-02 |
| | Syt7 | -1.38 | 3.190906428 | 0.00064430807 | 1.65E-02 |
| | Zmynd8 | -1.39 | 3.957208385 | 0.00011035490 | 1.41E-02 |
| | Pitpnm3 | -1.39 | 1.983988794 | 0.01037555188 | 5.47E-02 |
| | Pld3 | -1.39 | 3.066551725 | 0.00085792293 | 1.84E-02 |
| | Arhgef17 | -1.39 | 2.949824015 | 0.00112247321 | 2.00E-02 |
| | Atp8a2 | -1.39 | 2.903225471 | 0.00124961011 | 2.07E-02 |
| | Pcdha12 | -1.39 | 1.698685415 | 0.02001311013 | 7.95E-02 |
| | A830010M20Rik | -1.39 | 1.659053242 | 0.02192536127 | 8.38E-02 |
| | Git1 | -1.39 | 2.300773372 | 0.00500295536 | 3.71E-02 |

| | | | | | |
|---|---|---|---|---|---|
| | Grin2d | -1.39 | 1.598184842 | 0.02522406974 | 9.18E-02 |
| | Rgs4 | -1.39 | 1.436060482 | 0.03663865463 | 1.15E-01 |
| | R3hdm1 | -1.39 | 2.04203796 | 0.00907741185 | 5.14E-02 |
| | Med13l | -1.39 | 3.21293807 | 0.00061243772 | 1.65E-02 |
| | Srcin1 | -1.39 | 3.010229345 | 0.00097672129 | 1.92E-02 |
| | Efhd2 | -1.39 | 2.535439129 | 0.00291447860 | 2.89E-02 |
| | Rgs8 | -1.39 | 3.177067051 | 0.00066517045 | 1.67E-02 |
| | Rasgrf1 | -1.39 | 3.794933936 | 0.00016034893 | 1.41E-02 |
| | Asap1 | -1.39 | 1.921871968 | 0.01197093388 | 5.92E-02 |
| | Snapc4 | -1.39 | 1.971828323 | 0.01067017830 | 5.54E-02 |
| | Gpr158 | -1.39 | 2.02804366 | 0.00937467758 | 5.22E-02 |
| | Ptprn | -1.39 | 3.174618627 | 0.00066893108 | 1.67E-02 |
| | Aff2 | -1.39 | 2.201167808 | 0.00629262994 | 4.20E-02 |
| | Nectin1 | -1.39 | 1.488260142 | 0.03248926287 | 1.07E-01 |
| | Shf | -1.39 | 2.352341519 | 0.00444281758 | 3.47E-02 |
| | Gng2 | -1.39 | 2.680235095 | 0.00208816544 | 2.51E-02 |
| | Dock3 | -1.39 | 3.845110319 | 0.00014285310 | 1.41E-02 |
| | Klhl29 | -1.39 | 3.296072023 | 0.00050574078 | 1.59E-02 |
| | Plk3 | -1.39 | 2.512477814 | 0.00307271433 | 2.94E-02 |
| | Tonsl | -1.39 | 1.412323763 | 0.03869690547 | 1.19E-01 |
| | Trpc3 | -1.39 | 2.232484956 | 0.00585484017 | 4.06E-02 |
| | Cnksr2 | -1.39 | 2.748932006 | 0.00178265784 | 2.36E-02 |
| | Dpf1 | -1.39 | 1.986613667 | 0.01031303123 | 5.45E-02 |

| | | | | | |
|---|---|---|---|---|---|
| | Arnt2 | -1.39 | 1.914239389 | 0.01218317861 | 5.97E-02 |
| | Ppp1r3f | -1.39 | 3.46138373 | 0.00034563385 | 1.49E-02 |
| | Carmil2 | -1.39 | 1.890836945 | 0.01285769310 | 6.17E-02 |
| | Lingo2 | -1.39 | 2.071508896 | 0.00848186008 | 4.94E-02 |
| | Gfra2 | -1.40 | 1.510394899 | 0.03087486738 | 1.04E-01 |
| | Gpr12 | -1.40 | 1.701769882 | 0.01987147562 | 7.92E-02 |
| | Hivep2 | -1.40 | 2.294594138 | 0.00507464728 | 3.74E-02 |
| | Stxbp1 | -1.40 | 3.341676131 | 0.00045532749 | 1.59E-02 |
| | Homer1 | -1.40 | 2.083961543 | 0.00824211097 | 4.90E-02 |
| | Pgr | -1.40 | 1.936352786 | 0.01157836440 | 5.81E-02 |
| | Sco2 | -1.40 | 1.378062233 | 0.04187335576 | 1.26E-01 |
| | Apba1 | -1.40 | 2.918113462 | 0.00120749833 | 2.04E-02 |
| | Prrg3 | -1.40 | 2.728879097 | 0.00186689934 | 2.41E-02 |
| | Mapk10 | -1.40 | 2.265492238 | 0.00542634950 | 3.90E-02 |
| | Mypop | -1.40 | 3.028438171 | 0.00093661655 | 1.89E-02 |
| | Nrsn1 | -1.40 | 1.321615392 | 0.04768530975 | 1.37E-01 |
| | Ahdc1 | -1.40 | 2.969829244 | 0.00107194069 | 1.98E-02 |
| | Kcnh1 | -1.40 | 2.54927072 | 0.00282311962 | 2.86E-02 |
| | Ggt7 | -1.40 | 3.290424559 | 0.00051236026 | 1.59E-02 |
| | Cbarp | -1.40 | 2.130011932 | 0.00741289874 | 4.62E-02 |
| | Camta2 | -1.40 | 2.178248546 | 0.00663363320 | 4.32E-02 |
| | Fam57b | -1.40 | 2.351182884 | 0.00445468619 | 3.48E-02 |
| | Slc8a2 | -1.40 | 2.396929612 | 0.00400931693 | 3.33E-02 |

| | | | | | |
|---|---|---|---|---|---|
| | Adam19 | -1.40 | 2.360553768 | 0.00435959586 | 3.46E-02 |
| | Fam196a | -1.40 | 1.674116186 | 0.02117794491 | 8.21E-02 |
| | A230077H06Rik | -1.40 | 1.35996341 | 0.04365526106 | 1.29E-01 |
| | Parm1 | -1.40 | 3.036188063 | 0.00092005108 | 1.89E-02 |
| | Ptprj | -1.40 | 5.076956325 | 0.00000837614 | 1.41E-02 |
| | Mirg | -1.40 | 1.412817551 | 0.03865293255 | 1.19E-01 |
| | E330009J07Rik | -1.40 | 2.19638861 | 0.00636225966 | 4.23E-02 |
| | Zfp831 | -1.40 | 2.410739882 | 0.00388382916 | 3.28E-02 |
| | Clvs1 | -1.40 | 3.046965549 | 0.00089749999 | 1.86E-02 |
| | Camsap3 | -1.40 | 3.076112382 | 0.00083924279 | 1.81E-02 |
| | Dot1l | -1.40 | 1.664718663 | 0.02164119990 | 8.31E-02 |
| | Glt1d1 | -1.40 | 3.194311462 | 0.00063927620 | 1.65E-02 |
| | Grasp | -1.41 | 2.029288881 | 0.00934783675 | 5.21E-02 |
| | Tubg2 | -1.41 | 3.484082603 | 0.00032803290 | 1.49E-02 |
| | Fcho1 | -1.41 | 2.246885221 | 0.00566388959 | 3.99E-02 |
| | Rgs11 | -1.41 | 2.126736451 | 0.00746901873 | 4.64E-02 |
| | Peg13 | -1.41 | 3.922692253 | 0.00011948345 | 1.41E-02 |
| | Fbxo41 | -1.41 | 3.267413349 | 0.00054023989 | 1.61E-02 |
| | Nuak1 | -1.41 | 1.587708851 | 0.02583991905 | 9.34E-02 |
| | Pcsk2 | -1.41 | 1.954404702 | 0.01110696231 | 5.67E-02 |
| | Ccdc92 | -1.41 | 2.831628745 | 0.00147357164 | 2.19E-02 |
| | Cnnm1 | -1.41 | 3.507805928 | 0.00031059472 | 1.48E-02 |
| | Tmem132a | -1.41 | 2.799102077 | 0.00158817342 | 2.27E-02 |

| | | | | | |
|---|---|---|---|---|---|
| | Frmpd4 | -1.41 | 1.974050752 | 0.01061571495 | 5.53E-02 |
| | Fam43a | -1.41 | 1.919096122 | 0.01204769261 | 5.94E-02 |
| | Nr6a1 | -1.41 | 2.339336453 | 0.00457787097 | 3.54E-02 |
| | Chrna7 | -1.41 | 2.006583676 | 0.00984954852 | 5.33E-02 |
| | Jph3 | -1.41 | 2.344811203 | 0.00452052418 | 3.52E-02 |
| | Galnt14 | -1.41 | 1.916210845 | 0.01212799907 | 5.96E-02 |
| | Shd | -1.41 | 2.042495183 | 0.00906786021 | 5.13E-02 |
| | Nyap1 | -1.41 | 3.836279827 | 0.00014578746 | 1.41E-02 |
| | Clstn1 | -1.41 | 2.609984887 | 0.00245479434 | 2.68E-02 |
| | Dusp7 | -1.41 | 3.106084711 | 0.00078327685 | 1.78E-02 |
| | Trerf1 | -1.41 | 3.75370612 | 0.00017631688 | 1.41E-02 |
| | Pex5l | -1.41 | 1.788274024 | 0.01628268330 | 7.15E-02 |
| | Disp2 | -1.41 | 4.082116111 | 0.00008277208 | 1.41E-02 |
| | Hs3st4 | -1.41 | 2.285095305 | 0.00518686202 | 3.79E-02 |
| | Ube2o | -1.41 | 3.590261642 | 0.00025688477 | 1.42E-02 |
| | Wnk2 | -1.41 | 3.419808338 | 0.00038035722 | 1.52E-02 |
| | Atp1a1 | -1.41 | 2.100930363 | 0.00792628414 | 4.78E-02 |
| | Apba2 | -1.41 | 3.886914297 | 0.00012974353 | 1.41E-02 |
| | Apc2 | -1.41 | 3.830506351 | 0.00014773849 | 1.41E-02 |
| | Gm996 | -1.41 | 2.790069032 | 0.00162155233 | 2.29E-02 |
| | Mapk4 | -1.41 | 2.844380515 | 0.00143093361 | 2.17E-02 |
| | Ypel4 | -1.42 | 2.285429297 | 0.00518287462 | 3.79E-02 |
| | Klhl11 | -1.42 | 2.107696852 | 0.00780374641 | 4.77E-02 |

| | | | | | |
|---|---|---|---|---|---|
| | Epg5 | -1.42 | 5.104137341 | 0.00000786797 | 1.41E-02 |
| | Prrt2 | -1.42 | 2.338103698 | 0.00459088382 | 3.54E-02 |
| | Ncor2 | -1.42 | 2.747034996 | 0.00179046157 | 2.36E-02 |
| | Ltbp4 | -1.42 | 1.608987749 | 0.02460437010 | 9.03E-02 |
| | Zc3h12b | -1.42 | 1.766364271 | 0.01712520303 | 7.33E-02 |
| | Cx3cl1 | -1.42 | 3.63348093 | 0.00023255146 | 1.42E-02 |
| | 2900026A02Rik | -1.42 | 2.467822031 | 0.00340547713 | 3.08E-02 |
| | Sgtb | -1.42 | 3.151188924 | 0.00070601036 | 1.70E-02 |
| | Nat8l | -1.42 | 2.031619232 | 0.00929781215 | 5.20E-02 |
| | Wdr90 | -1.42 | 1.555541355 | 0.02782650387 | 9.79E-02 |
| | Snhg11 | -1.42 | 1.585556617 | 0.02596829182 | 9.36E-02 |
| | Ptpru | -1.42 | 1.693454924 | 0.02025559832 | 8.00E-02 |
| | Arhgef11 | -1.42 | 4.357107868 | 0.00004394325 | 1.41E-02 |
| | Npy5r | -1.42 | 1.882954919 | 0.01309317827 | 6.24E-02 |
| | Peli3 | -1.42 | 2.492740978 | 0.00321557780 | 3.00E-02 |
| | Ryr2 | -1.42 | 2.080475332 | 0.00830853911 | 4.91E-02 |
| | Ccdc159 | -1.42 | 2.019356903 | 0.00956407773 | 5.26E-02 |
| | Zfp771 | -1.42 | 1.797088001 | 0.01595555806 | 7.07E-02 |
| | Tmem63c | -1.42 | 2.315837606 | 0.00483239464 | 3.64E-02 |
| | Gpr3 | -1.42 | 1.42405246 | 0.03766582982 | 1.17E-01 |
| | Cdh12 | -1.42 | 1.723825773 | 0.01888748909 | 7.70E-02 |
| | Bicdl1 | -1.42 | 2.597123023 | 0.00252858162 | 2.71E-02 |
| | Arhgap20 | -1.42 | 3.318754479 | 0.00048000473 | 1.59E-02 |

| | | | | | |
|---|---|---|---|---|---|
| | Phf24 | -1.42 | 2.5570543<br>1 | 0.00277297<br>332 | 2.84E-<br>02 |
| | Cyfip2 | -1.42 | 2.7599497<br>61 | 0.00173800<br>187 | 2.34E-<br>02 |
| | Ptprn2 | -1.42 | 3.6097953<br>64 | 0.00024558<br>658 | 1.42E-<br>02 |
| | Spock1 | -1.42 | 2.0239894<br>67 | 0.00946260<br>110 | 5.24E-<br>02 |
| | Enc1 | -1.42 | 2.8478269<br>68 | 0.00141962<br>302 | 2.17E-<br>02 |
| | C130074G19<br>Rik | -1.42 | 2.2730555<br>42 | 0.00533266<br>692 | 3.86E-<br>02 |
| | Faim2 | -1.42 | 2.9090593<br>61 | 0.00123293<br>630 | 2.05E-<br>02 |
| | Agbl4 | -1.42 | 1.5740063<br>39 | 0.02666819<br>737 | 9.53E-<br>02 |
| | Epop | -1.43 | 1.5177167<br>1 | 0.03035870<br>837 | 1.03E-<br>01 |
| | Peg3 | -1.43 | 3.6657874<br>18 | 0.00021588<br>009 | 1.41E-<br>02 |
| | Kcnc4 | -1.43 | 3.3178313<br>34 | 0.00048102<br>613 | 1.59E-<br>02 |
| | Doc2a | -1.43 | 1.8724467<br>14 | 0.01341384<br>507 | 6.32E-<br>02 |
| | Usp13 | -1.43 | 2.6898739<br>68 | 0.00204233<br>054 | 2.50E-<br>02 |
| | Slc24a4 | -1.43 | 1.7779963<br>56 | 0.01667261<br>202 | 7.24E-<br>02 |
| | Plcxd2 | -1.43 | 1.3878224<br>82 | 0.04094279<br>789 | 1.24E-<br>01 |
| | Fbxo27 | -1.43 | 1.6471725<br>36 | 0.02253343<br>828 | 8.53E-<br>02 |
| | Chrna4 | -1.43 | 3.4464996<br>27 | 0.00035768<br>471 | 1.50E-<br>02 |
| | Pcsk1n | -1.43 | 1.5124170<br>08 | 0.03073144<br>573 | 1.04E-<br>01 |
| | Asic2 | -1.43 | 2.4894118<br>11 | 0.00324032<br>214 | 3.00E-<br>02 |
| | Synpo | -1.43 | 2.8712167<br>51 | 0.00134518<br>882 | 2.12E-<br>02 |
| | Dnajc21 | -1.43 | 1.5733508<br>51 | 0.02670847<br>856 | 9.54E-<br>02 |
| | Sncb | -1.43 | 3.2369773<br>51 | 0.00057945<br>891 | 1.64E-<br>02 |
| | Cacna1a | -1.43 | 2.2056329<br>79 | 0.00622826<br>411 | 4.18E-<br>02 |

| | | | | | |
|---|---|---|---|---|---|
| | A330023F24Rik | -1.43 | 1.822629573 | 0.01504424601 | 6.82E-02 |
| | Phyhip | -1.43 | 3.163329152 | 0.00068654791 | 1.69E-02 |
| | Cmip | -1.43 | 4.175503632 | 0.00006675693 | 1.41E-02 |
| | Syngap1 | -1.43 | 4.234877342 | 0.00005822676 | 1.41E-02 |
| | Hrh2 | -1.43 | 1.441542408 | 0.03617908608 | 1.15E-01 |
| | Lrfn1 | -1.43 | 2.823493811 | 0.00150143380 | 2.21E-02 |
| | Pcdhga3 | -1.43 | 1.43857247 | 0.03642734589 | 1.15E-01 |
| | Mccc1os | -1.43 | 1.408953682 | 0.03899835768 | 1.20E-01 |
| | Frrs1l | -1.43 | 2.166955703 | 0.00680838799 | 4.38E-02 |
| | Rims4 | -1.43 | 2.029035694 | 0.00935328798 | 5.21E-02 |
| | Tmem25 | -1.43 | 2.205263058 | 0.00623357145 | 4.18E-02 |
| | Pdgfb | -1.43 | 1.530064851 | 0.02950768572 | 1.01E-01 |
| | Grik3 | -1.43 | 4.017863309 | 0.00009597026 | 1.41E-02 |
| | Nckap1 | -1.43 | 3.102016848 | 0.00079064796 | 1.78E-02 |
| | Cep170b | -1.43 | 2.832037176 | 0.00147218648 | 2.19E-02 |
| | Tub | -1.43 | 3.168997306 | 0.00067764571 | 1.68E-02 |
| | Jdp2 | -1.43 | 2.013886793 | 0.00968530289 | 5.29E-02 |
| | Prmt8 | -1.43 | 3.890207288 | 0.00012876348 | 1.41E-02 |
| | Grin1 | -1.43 | 2.996929335 | 0.00100709552 | 1.92E-02 |
| | Tmem151a | -1.43 | 2.900025487 | 0.00125885153 | 2.07E-02 |
| | Olfm2 | -1.44 | 2.185220126 | 0.00652799592 | 4.28E-02 |
| | Synj2 | -1.44 | 1.920915512 | 0.01199732679 | 5.93E-02 |
| | Cdk5r2 | -1.44 | 3.32237159 | 0.00047602352 | 1.59E-02 |

| | | | | | |
|---|---|---|---|---|---|
| | Sptb | -1.44 | 3.0530276<br>03 | 0.00088505<br>936 | 1.85E-<br>02 |
| | Klc2 | -1.44 | 2.3886237<br>27 | 0.00408673<br>307 | 3.36E-<br>02 |
| | Nr1d1 | -1.44 | 1.9331974<br>66 | 0.01166279<br>209 | 5.84E-<br>02 |
| | Syn3 | -1.44 | 2.2339081<br>81 | 0.00583568<br>470 | 4.06E-<br>02 |
| | Camkk1 | -1.44 | 2.0525462<br>1 | 0.00886040<br>941 | 5.05E-<br>02 |
| | Rtn4rl1 | -1.44 | 2.5606425<br>87 | 0.00275015<br>653 | 2.83E-<br>02 |
| | Atmin | -1.44 | 3.3527371<br>69 | 0.00044387<br>719 | 1.59E-<br>02 |
| | Gramd1b | -1.44 | 3.8093311<br>24 | 0.00015512<br>039 | 1.41E-<br>02 |
| | Scube1 | -1.44 | 3.8681147<br>55 | 0.00013548<br>314 | 1.41E-<br>02 |
| | Lrrc8b | -1.44 | 2.7504911<br>37 | 0.00177626<br>952 | 2.36E-<br>02 |
| | Camkk2 | -1.44 | 2.3670159<br>11 | 0.00429520<br>691 | 3.43E-<br>02 |
| | Pip5k1c | -1.44 | 2.5486984<br>25 | 0.00282684<br>226 | 2.86E-<br>02 |
| | Syt12 | -1.44 | 2.8762516<br>6 | 0.00132968<br>369 | 2.12E-<br>02 |
| | Ptpn3 | -1.44 | 2.0006704<br>1 | 0.00998457<br>514 | 5.37E-<br>02 |
| | Slc45a1 | -1.44 | 3.9095821<br>06 | 0.00012314<br>532 | 1.41E-<br>02 |
| | 1700030J22R<br>ik | -1.44 | 2.7672945<br>84 | 0.00170885<br>580 | 2.33E-<br>02 |
| | Kcna2 | -1.44 | 1.8927005<br>39 | 0.01280263<br>785 | 6.15E-<br>02 |
| | B230209E15<br>Rik | -1.44 | 3.6634886<br>95 | 0.00021702<br>577 | 1.41E-<br>02 |
| | Numbl | -1.44 | 2.3301068<br>74 | 0.00467620<br>052 | 3.58E-<br>02 |
| | Epha10 | -1.44 | 2.6094696<br>7 | 0.00245770<br>826 | 2.68E-<br>02 |
| | Lypd6 | -1.44 | 2.2699093<br>42 | 0.00537143<br>913 | 3.88E-<br>02 |
| | Prdm8 | -1.44 | 1.3425169<br>96 | 0.04544467<br>519 | 1.33E-<br>01 |
| | Arhgap39 | -1.44 | 2.5111080<br>77 | 0.00308242<br>077 | 2.95E-<br>02 |

| | | | | | |
|---|---|---|---|---|---|
| | Plekha6 | -1.44 | 3.212438374 | 0.00061314279 | 1.65E-02 |
| | Golga7b | -1.44 | 1.856145795 | 0.01392689192 | 6.47E-02 |
| | Cacnb4 | -1.44 | 1.887904173 | 0.01294481437 | 6.20E-02 |
| | Sh3rf3 | -1.44 | 2.26231341 | 0.00546621350 | 3.90E-02 |
| | Slit1 | -1.44 | 1.626099668 | 0.02365376796 | 8.81E-02 |
| | Srrm4 | -1.44 | 2.115557965 | 0.00766376246 | 4.72E-02 |
| | Slc6a17 | -1.45 | 3.723304671 | 0.00018910165 | 1.41E-02 |
| | Ccdc184 | -1.45 | 2.546441122 | 0.00284157339 | 2.87E-02 |
| | Pacsin1 | -1.45 | 4.004824116 | 0.00009889535 | 1.41E-02 |
| | Btbd11 | -1.45 | 1.984748532 | 0.01035741716 | 5.46E-02 |
| | Slc2a13 | -1.45 | 2.929744356 | 0.00117558935 | 2.03E-02 |
| | Dgki | -1.45 | 3.136736462 | 0.00072990029 | 1.72E-02 |
| | Tchh | -1.45 | 2.336115563 | 0.00461194837 | 3.55E-02 |
| | Car7 | -1.45 | 2.316625977 | 0.00482363040 | 3.64E-02 |
| | Far2 | -1.45 | 2.869882813 | 0.00134932693 | 2.12E-02 |
| | Zbtb7a | -1.45 | 4.34671509 | 0.00004500750 | 1.41E-02 |
| | Rbfox3 | -1.45 | 2.919019222 | 0.00120498261 | 2.04E-02 |
| | Dnm1 | -1.45 | 3.316813096 | 0.00048215525 | 1.59E-02 |
| | Lmln | -1.45 | 2.137778717 | 0.00728150719 | 4.56E-02 |
| | Chd5 | -1.45 | 2.814685049 | 0.00153219821 | 2.23E-02 |
| | 2010111I01Rik | -1.45 | 2.71118405 | 0.00194453583 | 2.44E-02 |
| | Trhde | -1.45 | 1.364586845 | 0.04319297869 | 1.28E-01 |
| | Sema6c | -1.45 | 2.336878516 | 0.00460385338 | 3.55E-02 |

| | | | | | |
|---|---|---|---|---|---|
| | Diras2 | -1.45 | 2.140887785 | 0.00722956580 | 4.54E-02 |
| | Whrn | -1.45 | 1.350242683 | 0.04464340561 | 1.31E-01 |
| | Lrrc24 | -1.45 | 3.986313048 | 0.00010320172 | 1.41E-02 |
| | Ccl27a | -1.45 | 1.952897842 | 0.01114556676 | 5.69E-02 |
| | Mical2 | -1.45 | 1.437420033 | 0.03652413729 | 1.15E-01 |
| | Ppm1h | -1.45 | 3.331828276 | 0.00046577023 | 1.59E-02 |
| | D630045J12Rik | -1.45 | 3.370625765 | 0.00042596531 | 1.59E-02 |
| | Stx1b | -1.45 | 3.757621036 | 0.00017473462 | 1.41E-02 |
| | Sorl1 | -1.45 | 2.773928257 | 0.00168295205 | 2.32E-02 |
| | Cacna1i | -1.45 | 2.971105705 | 0.00106879471 | 1.98E-02 |
| | Nbas | -1.45 | 2.503447073 | 0.00313727744 | 2.97E-02 |
| | Slit3 | -1.46 | 1.808204465 | 0.01555233259 | 6.97E-02 |
| | Rnf126 | -1.46 | 3.070955073 | 0.00084926833 | 1.83E-02 |
| | B4galnt4 | -1.46 | 3.272757828 | 0.00053363238 | 1.61E-02 |
| | Cit | -1.46 | 4.407248616 | 0.00003915177 | 1.41E-02 |
| | Pcdh1 | -1.46 | 3.236156012 | 0.00058055583 | 1.64E-02 |
| | Hs3st2 | -1.46 | 2.411430909 | 0.00387765433 | 3.28E-02 |
| | Pcsk1 | -1.46 | 1.971475487 | 0.01067885067 | 5.54E-02 |
| | Rusc1 | -1.46 | 3.861351394 | 0.00013760956 | 1.41E-02 |
| | Slit2 | -1.46 | 1.440868122 | 0.03623530143 | 1.15E-01 |
| | Galnt9 | -1.46 | 1.381464601 | 0.04154659138 | 1.25E-01 |
| | Adnp | -1.46 | 1.49539271 | 0.03196003824 | 1.06E-01 |
| | Gm715 | -1.46 | 1.550553508 | 0.02814793190 | 9.85E-02 |

| | | | | | |
|---|---|---|---|---|---|
| | Rasgef1b | -1.46 | 2.357705141 | 0.00438828534 | 3.47E-02 |
| | A230103J11Rik | -1.46 | 1.407252988 | 0.03915137437 | 1.20E-01 |
| | Vwa5b2 | -1.46 | 1.534163879 | 0.02923049172 | 1.01E-01 |
| | Satb1 | -1.46 | 1.433797456 | 0.03683006998 | 1.16E-01 |
| | Car10 | -1.46 | 1.796357326 | 0.01598242497 | 7.07E-02 |
| | Sptbn4 | -1.46 | 4.231062157 | 0.00005874053 | 1.41E-02 |
| | Cacnb1 | -1.46 | 3.609984291 | 0.00024547977 | 1.42E-02 |
| | Pip5k1b | -1.46 | 2.356372536 | 0.00440177119 | 3.47E-02 |
| | Nt5dc3 | -1.47 | 2.140928843 | 0.00722888236 | 4.54E-02 |
| | Fam171a2 | -1.47 | 3.878699099 | 0.00013222114 | 1.41E-02 |
| | C77080 | -1.47 | 1.77622396 | 0.01674079352 | 7.26E-02 |
| | Paqr9 | -1.47 | 3.334198843 | 0.00046323478 | 1.59E-02 |
| | Lrrc14b | -1.47 | 2.353417263 | 0.00443182637 | 3.47E-02 |
| | Entpd7 | -1.47 | 2.256294415 | 0.00554249851 | 3.94E-02 |
| | Mical3 | -1.47 | 4.541690219 | 0.00002872829 | 1.41E-02 |
| | Brsk2 | -1.47 | 2.354878734 | 0.00441693762 | 3.47E-02 |
| | Ngef | -1.47 | 2.935590951 | 0.00115986929 | 2.03E-02 |
| | Npas2 | -1.47 | 2.823333027 | 0.00150198976 | 2.21E-02 |
| | Fxyd7 | -1.47 | 1.902283291 | 0.01252324016 | 6.07E-02 |
| | Car4 | -1.47 | 1.548598018 | 0.02827495890 | 9.87E-02 |
| | Extl1 | -1.47 | 2.410383385 | 0.00388701857 | 3.28E-02 |
| | Rapgefl1 | -1.47 | 3.423616636 | 0.00037703647 | 1.52E-02 |
| | Tmem74 | -1.47 | 1.321285944 | 0.04772149669 | 1.37E-01 |

| | | | | | |
|---|---|---|---|---|---|
| | Ntng2 | -1.47 | 2.098387174 | 0.00797283592 | 4.80E-02 |
| | Hectd4 | -1.47 | 4.709881195 | 0.00001950378 | 1.41E-02 |
| | Cbfa2t3 | -1.47 | 2.252933919 | 0.00558555177 | 3.95E-02 |
| | Gal3st3 | -1.47 | 2.814940116 | 0.00153129859 | 2.23E-02 |
| | Dyrk2 | -1.47 | 2.543129198 | 0.00286332603 | 2.87E-02 |
| | Adamts17 | -1.47 | 1.840276147 | 0.01444520978 | 6.63E-02 |
| | Sidt1 | -1.47 | 2.864507698 | 0.00136613086 | 2.13E-02 |
| | Scrt1 | -1.47 | 1.697219568 | 0.02008077322 | 7.96E-02 |
| | Zfhx2 | -1.48 | 3.714078938 | 0.00019316172 | 1.41E-02 |
| | Map3k5 | -1.48 | 3.090976441 | 0.00081100505 | 1.79E-02 |
| | Mtus2 | -1.48 | 2.354228769 | 0.00442355296 | 3.47E-02 |
| | Cyp4x1 | -1.48 | 2.227085827 | 0.00592808160 | 4.08E-02 |
| | Chrm1 | -1.48 | 3.763677301 | 0.00017231485 | 1.41E-02 |
| | Hic2 | -1.48 | 1.936697534 | 0.01156917701 | 5.81E-02 |
| | Ccnf | -1.48 | 1.587452841 | 0.02585515580 | 9.34E-02 |
| | Slc12a5 | -1.48 | 3.463715649 | 0.00034378296 | 1.49E-02 |
| | Fhod3 | -1.48 | 2.759633768 | 0.00173926690 | 2.34E-02 |
| | Brinp1 | -1.48 | 2.399997262 | 0.00398109680 | 3.32E-02 |
| | Dmtn | -1.48 | 3.207082876 | 0.00062075057 | 1.65E-02 |
| | Kndc1 | -1.48 | 2.425254788 | 0.00375616976 | 3.23E-02 |
| | Hecw1 | -1.48 | 1.679988583 | 0.02089351054 | 8.16E-02 |
| | Flrt1 | -1.49 | 3.825205995 | 0.00014955261 | 1.41E-02 |
| | Ankrd9 | -1.49 | 1.893805132 | 0.01277011675 | 6.14E-02 |

| | | | | | |
|---|---|---|---|---|---|
| | Otud7a | -1.49 | 2.916214198 | 0.00121279054 | 2.04E-02 |
| | Pcnx2 | -1.49 | 2.950855221 | 0.00111981113 | 2.00E-02 |
| | Zdhhc8 | -1.49 | 2.603457194 | 0.00249196998 | 2.69E-02 |
| | Unc5a | -1.49 | 3.718774584 | 0.00019108448 | 1.41E-02 |
| | Tshz3 | -1.49 | 1.703613868 | 0.01978728148 | 7.90E-02 |
| | Tmppe | -1.49 | 2.128069816 | 0.00744612262 | 4.63E-02 |
| | Has3 | -1.49 | 1.585436588 | 0.02597546983 | 9.36E-02 |
| | Chrd | -1.49 | 1.543690605 | 0.02859627042 | 9.94E-02 |
| | Faxc | -1.49 | 3.312449014 | 0.00048702470 | 1.59E-02 |
| | Cobl | -1.49 | 2.404488069 | 0.00394014252 | 3.30E-02 |
| | Fam135b | -1.49 | 2.770018784 | 0.00169817020 | 2.32E-02 |
| | Adgrb1 | -1.49 | 3.563184984 | 0.00027341039 | 1.42E-02 |
| | Hspb3 | -1.49 | 1.479484576 | 0.03315243435 | 1.09E-01 |
| | Adrb1 | -1.49 | 2.747964155 | 0.00178663503 | 2.36E-02 |
| | Trank1 | -1.50 | 3.300508215 | 0.00050060108 | 1.59E-02 |
| | Cacna1e | -1.50 | 3.548525741 | 0.00028279665 | 1.43E-02 |
| | Etl4 | -1.50 | 2.732536984 | 0.00185124124 | 2.40E-02 |
| | Osbpl3 | -1.50 | 2.19860898 | 0.00632981504 | 4.21E-02 |
| | Wbscr17 | -1.50 | 3.621965576 | 0.00023880006 | 1.42E-02 |
| | Dact2 | -1.50 | 1.6431739 | 0.02274186618 | 8.59E-02 |
| | Clstn3 | -1.50 | 3.705125899 | 0.00019718510 | 1.41E-02 |
| | Ina | -1.50 | 4.648018305 | 0.00002248960 | 1.41E-02 |
| | Mtcl1 | -1.50 | 3.499018199 | 0.00031694346 | 1.48E-02 |

| | | | | | |
|---|---|---|---|---|---|
| | Pcdha5 | -1.50 | 1.938141572 | 0.01153077316 | 5.80E-02 |
| | Ttbk1 | -1.50 | 3.460742998 | 0.00034614415 | 1.49E-02 |
| | Scn8a | -1.50 | 2.292391139 | 0.00510045431 | 3.75E-02 |
| | Acta1 | -1.50 | 1.391807587 | 0.04056882345 | 1.23E-01 |
| | Slc25a34 | -1.50 | 1.515143645 | 0.03053910852 | 1.03E-01 |
| | Syt13 | -1.50 | 2.825785624 | 0.00149353146 | 2.20E-02 |
| | Ankrd34a | -1.50 | 3.32507372 | 0.00047307095 | 1.59E-02 |
| | Klhl3 | -1.51 | 2.366873174 | 0.00429661881 | 3.43E-02 |
| | Tbc1d30 | -1.51 | 2.409286405 | 0.00389684915 | 3.28E-02 |
| | Osbp2 | -1.51 | 1.943693839 | 0.01138429552 | 5.74E-02 |
| | Sez6l | -1.51 | 2.848078026 | 0.00141880259 | 2.17E-02 |
| | Ephb6 | -1.51 | 1.7189398 | 0.01910118012 | 7.74E-02 |
| | Pcdhac2 | -1.51 | 3.136965702 | 0.00072951512 | 1.72E-02 |
| | Gm9899 | -1.51 | 2.201040598 | 0.00629447339 | 4.20E-02 |
| | Gm11549 | -1.51 | 1.604881737 | 0.02483809381 | 9.09E-02 |
| | Cacng2 | -1.51 | 2.372433466 | 0.00424195966 | 3.41E-02 |
| | Adcy9 | -1.52 | 3.393668331 | 0.00040395377 | 1.55E-02 |
| | Serpinb8 | -1.52 | 1.930138438 | 0.01174523097 | 5.86E-02 |
| | Jag2 | -1.52 | 2.979190183 | 0.00104908292 | 1.97E-02 |
| | Dlgap2 | -1.52 | 3.31629374 | 0.00048273219 | 1.59E-02 |
| | 2410018L13Rik | -1.52 | 1.67181717 | 0.02129035144 | 8.24E-02 |
| | Stx1a | -1.52 | 2.358784874 | 0.00437738884 | 3.47E-02 |
| | Sowahb | -1.52 | 1.599977734 | 0.02512015218 | 9.15E-02 |

| | | | | | |
|---|---|---|---|---|---|
| | Foxo3 | -1.52 | 3.001155117 | 0.00099734378 | 1.92E-02 |
| | Ramp3 | -1.52 | 1.959171029 | 0.01098573126 | 5.64E-02 |
| | Clstn2 | -1.52 | 2.083826559 | 0.00824467311 | 4.90E-02 |
| | Sema7a | -1.52 | 1.623882576 | 0.02377483022 | 8.84E-02 |
| | Srrm3 | -1.52 | 3.468052515 | 0.00034036703 | 1.49E-02 |
| | Efna5 | -1.53 | 1.490954073 | 0.03228835552 | 1.07E-01 |
| | Dlk2 | -1.53 | 1.983577816 | 0.01038537504 | 5.47E-02 |
| | Lpcat4 | -1.53 | 3.466366323 | 0.00034169111 | 1.49E-02 |
| | Kcnj9 | -1.53 | 1.944414719 | 0.01136541454 | 5.74E-02 |
| | Serinc2 | -1.53 | 1.830113006 | 0.01478723564 | 6.75E-02 |
| | Lmtk2 | -1.53 | 3.092840277 | 0.00080753197 | 1.79E-02 |
| | Irs2 | -1.53 | 1.667510319 | 0.02150253587 | 8.28E-02 |
| | Gfod1 | -1.53 | 2.70383774 | 0.00197770841 | 2.46E-02 |
| | Sstr4 | -1.53 | 2.072941778 | 0.00845392173 | 4.94E-02 |
| | Lmtk3 | -1.53 | 3.986172246 | 0.00010323519 | 1.41E-02 |
| | Krt80 | -1.53 | 1.331028428 | 0.04666288346 | 1.35E-01 |
| | Tmem198 | -1.53 | 2.848584524 | 0.00141714887 | 2.17E-02 |
| | Xkr4 | -1.54 | 2.59626218 | 0.00253359866 | 2.71E-02 |
| | Trim17 | -1.54 | 1.91465252 | 0.01217159463 | 5.97E-02 |
| | Kcnj11 | -1.54 | 1.369303786 | 0.04272639131 | 1.27E-01 |
| | Vipr1 | -1.54 | 2.653653146 | 0.00221996871 | 2.59E-02 |
| | Shc3 | -1.54 | 1.992706271 | 0.01016936250 | 5.41E-02 |
| | Ier5 | -1.54 | 3.28589603 | 0.00051773076 | 1.60E-02 |

| | | | | | |
|---|---|---|---|---|---|
| | Kcnt2 | -1.54 | 2.148723431 | 0.00710029787 | 4.50E-02 |
| | Tspoap1 | -1.54 | 2.343725149 | 0.00453184295 | 3.52E-02 |
| | Slc9a5 | -1.54 | 2.138957003 | 0.00726177848 | 4.56E-02 |
| | Mast3 | -1.55 | 2.616624359 | 0.00241755098 | 2.66E-02 |
| | Zdhhc22 | -1.55 | 1.367984641 | 0.04285636762 | 1.28E-01 |
| | Sik2 | -1.55 | 1.973984736 | 0.01061732874 | 5.53E-02 |
| | Kcnk3 | -1.55 | 3.433223289 | 0.00036878794 | 1.52E-02 |
| | Palm2 | -1.55 | 1.955154106 | 0.01108781304 | 5.67E-02 |
| | Neurod2 | -1.55 | 2.883371424 | 0.00130806274 | 2.10E-02 |
| | Plcb4 | -1.55 | 2.323735439 | 0.00474530969 | 3.61E-02 |
| | Rnf112 | -1.55 | 3.010421364 | 0.00097628954 | 1.92E-02 |
| | Kalrn | -1.55 | 2.427268999 | 0.00373878939 | 3.23E-02 |
| | Gnaz | -1.55 | 3.895461682 | 0.00012721500 | 1.41E-02 |
| | Oprd1 | -1.55 | 3.224499683 | 0.00059634876 | 1.65E-02 |
| | Lzts3 | -1.56 | 4.23413724 | 0.00005832608 | 1.41E-02 |
| | Gpr26 | -1.56 | 3.702582639 | 0.00019834322 | 1.41E-02 |
| | Mast1 | -1.56 | 3.748689608 | 0.00017836531 | 1.41E-02 |
| | Kcnc1 | -1.56 | 3.079387241 | 0.00083293816 | 1.81E-02 |
| | Tbr1 | -1.56 | 2.998368841 | 0.00100376294 | 1.92E-02 |
| | Grm2 | -1.57 | 2.866049213 | 0.00136129042 | 2.13E-02 |
| | Phospho1 | -1.57 | 1.641374478 | 0.02283628854 | 8.61E-02 |
| | Ankrd24 | -1.57 | 2.653447094 | 0.00222102223 | 2.59E-02 |
| | Igfbp6 | -1.57 | 2.039538682 | 0.00912980114 | 5.16E-02 |

| | | | | | |
|---|---|---|---|---|---|
| | Vamp1 | -1.57 | 1.388816019 | 0.04084924003 | 1.24E-01 |
| | Pcdha3 | -1.57 | 2.171822394 | 0.00673251928 | 4.37E-02 |
| | Mapk11 | -1.57 | 2.008224045 | 0.00981241608 | 5.32E-02 |
| | Cdh24 | -1.57 | 2.076932018 | 0.00837660395 | 4.92E-02 |
| | Celsr3 | -1.58 | 3.217120074 | 0.00060656860 | 1.65E-02 |
| | Tcp11l1 | -1.58 | 2.170262907 | 0.00675673822 | 4.37E-02 |
| | Nptx1 | -1.58 | 3.141597476 | 0.00072177615 | 1.71E-02 |
| | Cckbr | -1.58 | 2.20322254 | 0.00626292860 | 4.19E-02 |
| | A230001M10Rik | -1.58 | 1.378693848 | 0.04181250164 | 1.26E-01 |
| | Bsn | -1.58 | 3.438233804 | 0.00036455763 | 1.51E-02 |
| | Kcnma1 | -1.58 | 3.007044791 | 0.00098390962 | 1.92E-02 |
| | Fbxl18 | -1.58 | 3.653896502 | 0.00022187251 | 1.42E-02 |
| | Nxph3 | -1.59 | 2.160406096 | 0.00691184362 | 4.41E-02 |
| | Hipk4 | -1.59 | 2.381298296 | 0.00415625040 | 3.38E-02 |
| | Trpv6 | -1.59 | 1.603089544 | 0.02494080436 | 9.11E-02 |
| | Dgkh | -1.59 | 1.982740595 | 0.01040541498 | 5.47E-02 |
| | Rimbp2 | -1.59 | 3.183606686 | 0.00065522931 | 1.66E-02 |
| | Nyap2 | -1.59 | 1.768032799 | 0.01705953546 | 7.32E-02 |
| | D030047H15Rik | -1.59 | 1.414376635 | 0.03851442027 | 1.19E-01 |
| | Ddi2 | -1.59 | 3.245951313 | 0.00056760823 | 1.64E-02 |
| | D130017N08Rik | -1.60 | 2.772491788 | 0.00168852779 | 2.32E-02 |
| | Lrfn2 | -1.60 | 1.812270374 | 0.01540740952 | 6.91E-02 |
| | Crhr1 | -1.60 | 2.076398142 | 0.00838690760 | 4.92E-02 |

| | | | | | |
|---|---|---|---|---|---|
| | Pakap | -1.60 | 2.977037358 | 0.00105429620 | 1.97E-02 |
| | Bdnf | -1.60 | 1.691829352 | 0.02033155744 | 8.02E-02 |
| | Chrm3 | -1.60 | 3.027059808 | 0.00093959391 | 1.89E-02 |
| | Gabrb2 | -1.60 | 3.200883582 | 0.00062967495 | 1.65E-02 |
| | Sstr1 | -1.60 | 3.171164137 | 0.00067427314 | 1.67E-02 |
| | Epha8 | -1.60 | 2.063088751 | 0.00864791175 | 4.99E-02 |
| | D430041D05Rik | -1.61 | 2.683932672 | 0.00207046230 | 2.51E-02 |
| | Gm12522 | -1.61 | 1.320207855 | 0.04784010732 | 1.37E-01 |
| | Qrfpr | -1.61 | 1.892912446 | 0.01279639255 | 6.15E-02 |
| | Efna3 | -1.61 | 3.087277043 | 0.00081794284 | 1.80E-02 |
| | Shank2 | -1.61 | 3.361455089 | 0.00043505575 | 1.59E-02 |
| | Kcnq3 | -1.61 | 3.502230298 | 0.00031460796 | 1.48E-02 |
| | Atp2b2 | -1.61 | 2.346078919 | 0.00450734791 | 3.51E-02 |
| | Sema4f | -1.61 | 4.481309899 | 0.00003301339 | 1.41E-02 |
| | Prrt3 | -1.62 | 3.129086465 | 0.00074287122 | 1.74E-02 |
| | Kcnh4 | -1.62 | 1.901670051 | 0.01254093593 | 6.08E-02 |
| | Tfr2 | -1.62 | 1.31981691 | 0.04788319162 | 1.37E-01 |
| | Grin2b | -1.62 | 2.512529268 | 0.00307235030 | 2.94E-02 |
| | Dusp1 | -1.62 | 1.603472028 | 0.02491884866 | 9.11E-02 |
| | Kcnb1 | -1.62 | 4.269145326 | 0.00005380897 | 1.41E-02 |
| | Tmem178b | -1.62 | 3.006256582 | 0.00098569696 | 1.92E-02 |
| | Adgrd1 | -1.62 | 1.357248248 | 0.04392904402 | 1.30E-01 |
| | Gm20063 | -1.62 | 1.314681223 | 0.04845278858 | 1.38E-01 |

| | | | | | |
|---|---|---|---|---|---|
| | Map3k9 | -1.62 | 3.172422808 | 0.00067232180 | 1.67E-02 |
| | AF529169 | -1.62 | 1.430524302 | 0.03710869637 | 1.16E-01 |
| | Sstr2 | -1.62 | 3.357412701 | 0.00043912413 | 1.59E-02 |
| | Abcc8 | -1.63 | 2.505776297 | 0.00312049652 | 2.97E-02 |
| | Eps8l2 | -1.63 | 1.526200434 | 0.02977142111 | 1.02E-01 |
| | Cdkl5 | -1.63 | 4.40621565 | 0.00003924500 | 1.41E-02 |
| | Mybpc3 | -1.63 | 1.536290214 | 0.02908772705 | 1.00E-01 |
| | Csmd1 | -1.63 | 3.700428543 | 0.00019932944 | 1.41E-02 |
| | Mamld1 | -1.63 | 2.181190369 | 0.00658885015 | 4.31E-02 |
| | Nrgn | -1.64 | 1.962545907 | 0.01090069260 | 5.62E-02 |
| | Tmem200b | -1.64 | 2.040518031 | 0.00910923632 | 5.15E-02 |
| | Kcnk4 | -1.64 | 1.686670253 | 0.02057452166 | 8.09E-02 |
| | Fmnl1 | -1.64 | 3.158411571 | 0.00069436597 | 1.69E-02 |
| | Ptges3l | -1.64 | 2.276062319 | 0.00529587445 | 3.84E-02 |
| | Prr7 | -1.64 | 2.190047218 | 0.00645584035 | 4.26E-02 |
| | Kcnmb4os2 | -1.64 | 1.78613676 | 0.01636301168 | 7.16E-02 |
| | Myh2 | -1.65 | 1.422085166 | 0.03783683791 | 1.18E-01 |
| | Hsf4 | -1.65 | 3.00525124 | 0.00098798138 | 1.92E-02 |
| | Kcna1 | -1.65 | 2.411184551 | 0.00387985459 | 3.28E-02 |
| | Stxbp5l | -1.65 | 3.314026782 | 0.00048525857 | 1.59E-02 |
| | Unc13a | -1.65 | 3.874248478 | 0.00013358310 | 1.41E-02 |
| | Ltk | -1.65 | 1.399084484 | 0.03989472866 | 1.22E-01 |
| | Emx1 | -1.65 | 1.99449583 | 0.01012754473 | 5.40E-02 |

| | | | | | |
|---|---|---|---|---|---|
| | Wnt10a | -1.65 | 1.757809542 | 0.01746587944 | 7.39E-02 |
| | Igsf9b | -1.66 | 3.175508428 | 0.00066756195 | 1.67E-02 |
| | Tenm1 | -1.66 | 1.703182203 | 0.01980695873 | 7.90E-02 |
| | Ephb2 | -1.66 | 3.682421292 | 0.00020776802 | 1.41E-02 |
| | Htr1a | -1.66 | 3.41469354 | 0.00038486327 | 1.52E-02 |
| | Nkx3-1 | -1.67 | 1.48740139 | 0.03255356904 | 1.07E-01 |
| | Fezf2 | -1.67 | 1.495472043 | 0.03195420062 | 1.06E-01 |
| | Tmem178 | -1.67 | 1.967372175 | 0.01078022498 | 5.58E-02 |
| | D130043K22Rik | -1.67 | 3.667544171 | 0.00021500860 | 1.41E-02 |
| | L3mbtl1 | -1.67 | 1.869320395 | 0.01351075456 | 6.35E-02 |
| | Sdk2 | -1.67 | 1.862869737 | 0.01371293012 | 6.41E-02 |
| | Adra1d | -1.67 | 2.098453679 | 0.00797161509 | 4.80E-02 |
| | Zfp57 | -1.68 | 1.672153063 | 0.02127389137 | 8.23E-02 |
| | Cdh6 | -1.68 | 2.165368798 | 0.00683331124 | 4.39E-02 |
| | Nav3 | -1.69 | 2.812400878 | 0.00154027803 | 2.23E-02 |
| | Npc1l1 | -1.69 | 1.548287706 | 0.02829516911 | 9.87E-02 |
| | Atp2b3 | -1.69 | 5.182961315 | 0.00000656204 | 1.41E-02 |
| | Zbtb46 | -1.69 | 2.302872708 | 0.00497882994 | 3.70E-02 |
| | Kcnj14 | -1.69 | 2.158069206 | 0.00694913573 | 4.43E-02 |
| | Kcnj6 | -1.69 | 2.830206297 | 0.00147840596 | 2.19E-02 |
| | Gabbr2 | -1.69 | 3.231673201 | 0.00058657939 | 1.65E-02 |
| | BC030499 | -1.70 | 1.499238591 | 0.03167826658 | 1.06E-01 |
| | Pnpla1 | -1.70 | 1.494336851 | 0.03203783414 | 1.06E-01 |

| | | | | | |
|---|---|---|---|---|---|
| | Siah3 | -1.71 | 1.78591022 | 0.01637154931 | 7.16E-02 |
| | Trim66 | -1.71 | 3.556858956 | 0.00027742209 | 1.43E-02 |
| | Rtn4r | -1.71 | 2.332174055 | 0.00465399535 | 3.57E-02 |
| | Syt2 | -1.71 | 1.750085739 | 0.01777928375 | 7.47E-02 |
| | Pla2g4e | -1.71 | 2.476251991 | 0.00334001186 | 3.04E-02 |
| | Tmem132d | -1.71 | 2.948534411 | 0.00112581126 | 2.00E-02 |
| | 9430037G07Rik | -1.71 | 1.609349361 | 0.02458389200 | 9.03E-02 |
| | Lingo1 | -1.71 | 2.858405238 | 0.00138546246 | 2.15E-02 |
| | Fstl4 | -1.72 | 2.856469303 | 0.00139165216 | 2.15E-02 |
| | Hs6st3 | -1.72 | 3.062921265 | 0.00086512475 | 1.84E-02 |
| | Meg3 | -1.72 | 2.039218427 | 0.00913653607 | 5.16E-02 |
| | Cdsn | -1.72 | 1.343002855 | 0.04539386326 | 1.33E-01 |
| | Plekhg5 | -1.72 | 2.870981908 | 0.00134591642 | 2.12E-02 |
| | Kcnc3 | -1.72 | 2.401503346 | 0.00396731472 | 3.31E-02 |
| | Dagla | -1.74 | 3.767524365 | 0.00017079519 | 1.41E-02 |
| | Tmem145 | -1.74 | 2.038424235 | 0.00915325928 | 5.16E-02 |
| | Phf21b | -1.74 | 1.755665463 | 0.01755232037 | 7.41E-02 |
| | Myl4 | -1.75 | 1.668409743 | 0.02145805018 | 8.27E-02 |
| | Sgsm1 | -1.76 | 2.100378361 | 0.00793636511 | 4.78E-02 |
| | Tmem181b-ps | -1.76 | 1.828272905 | 0.01485002192 | 6.77E-02 |
| | Rtn4rl2 | -1.76 | 2.923672983 | 0.00119213933 | 2.03E-02 |
| | Gm11762 | -1.76 | 1.603639694 | 0.02490923021 | 9.11E-02 |
| | Klhl33 | -1.77 | 2.443871119 | 0.00359856110 | 3.17E-02 |

| | | | | | |
|---|---|---|---|---|---|
| | Shank1 | -1.77 | 2.961342344 | 0.00109309437 | 1.99E-02 |
| | Nos1ap | -1.79 | 3.025812236 | 0.00094229690 | 1.89E-02 |
| | Cux2 | -1.80 | 3.693029758 | 0.00020275438 | 1.41E-02 |
| | Miat | -1.80 | 2.069722418 | 0.00851682221 | 4.94E-02 |
| | Kctd16 | -1.80 | 2.542779899 | 0.00286562991 | 2.87E-02 |
| | Coro6 | -1.80 | 2.373289566 | 0.00423360596 | 3.41E-02 |
| | Arc | -1.81 | 1.920694389 | 0.01200343683 | 5.93E-02 |
| | Kcng4 | -1.81 | 1.421826422 | 0.03785938699 | 1.18E-01 |
| | Cpne9 | -1.82 | 2.23170105 | 0.00586541776 | 4.06E-02 |
| | Arfgef3 | -1.82 | 2.932100755 | 0.00116922810 | 2.03E-02 |
| | Myo5c | -1.82 | 2.103436911 | 0.00788066905 | 4.77E-02 |
| | Tspan11 | -1.83 | 1.893750148 | 0.01277173362 | 6.14E-02 |
| | Npbwr1 | -1.83 | 1.408067999 | 0.03907797051 | 1.20E-01 |
| | Fgf5 | -1.83 | 2.069455347 | 0.00852206128 | 4.95E-02 |
| | Robo3 | -1.83 | 1.53288517 | 0.02931668290 | 1.01E-01 |
| | Nxf7 | -1.83 | 1.324473807 | 0.04737248785 | 1.37E-01 |
| | Grin2a | -1.83 | 2.796400408 | 0.00159808396 | 2.27E-02 |
| | Proser2 | -1.83 | 1.451980118 | 0.03531993391 | 1.13E-01 |
| | Mir124a-1hg | -1.83 | 2.831461941 | 0.00147413772 | 2.19E-02 |
| | Cdyl2 | -1.84 | 3.406789242 | 0.00039193203 | 1.54E-02 |
| | Tekt2 | -1.84 | 1.364882932 | 0.04316354127 | 1.28E-01 |
| | Slc26a8 | -1.86 | 1.7094584 | 0.01952277727 | 7.84E-02 |
| | Kcnh5 | -1.87 | 2.225538148 | 0.00594924498 | 4.08E-02 |

| | | | | | |
|---|---|---|---|---|---|
| | Rims3 | -1.87 | 2.095307014 | 0.00802958290 | 4.82E-02 |
| | Ksr2 | -1.88 | 4.613005182 | 0.00002437782 | 1.41E-02 |
| | Gm13308 | -1.88 | 1.794569192 | 0.01604836556 | 7.09E-02 |
| | Col19a1 | -1.89 | 2.514739101 | 0.00305675688 | 2.94E-02 |
| | Pygm | -1.90 | 2.536472476 | 0.00290755223 | 2.89E-02 |
| | Pcdha10 | -1.90 | 2.393763719 | 0.00403865059 | 3.34E-02 |
| | Barx2 | -1.91 | 2.077047193 | 0.00837438277 | 4.92E-02 |
| | Hkdc1 | -1.91 | 1.998231896 | 0.01004079509 | 5.38E-02 |
| | Nptxr | -1.93 | 2.468648677 | 0.00339900124 | 3.08E-02 |
| | Kcnk9 | -1.93 | 3.125173207 | 0.00074959519 | 1.74E-02 |
| | Chrm2 | -1.94 | 3.104355491 | 0.00078640182 | 1.78E-02 |
| | P2rx3 | -1.94 | 1.591306711 | 0.02562673566 | 9.29E-02 |
| | Satb2 | -1.96 | 2.20861719 | 0.00618561393 | 4.16E-02 |
| | Rxfp1 | -1.97 | 2.929833841 | 0.00117534715 | 2.03E-02 |
| | Zbtb16 | -1.98 | 2.082070255 | 0.00827808239 | 4.90E-02 |
| | Kirrel2 | -1.99 | 2.503990214 | 0.00313335633 | 2.97E-02 |
| | Gm21949 | -2.00 | 1.941601606 | 0.01143927219 | 5.76E-02 |
| | Kcnk12 | -2.00 | 2.745305894 | 0.00179760433 | 2.36E-02 |
| | Nphs1 | -2.01 | 1.7756151 | 0.01676427976 | 7.26E-02 |
| | Smyd1 | -2.01 | 1.462764956 | 0.03445363467 | 1.11E-01 |
| | Kcnh7 | -2.02 | 2.697751835 | 0.00200561775 | 2.47E-02 |
| | Sacs | -2.02 | 2.318736957 | 0.00480024101 | 3.63E-02 |
| | Col24a1 | -2.05 | 2.327797257 | 0.00470113522 | 3.59E-02 |

| | | | | | |
|---|---|---|---|---|---|
| | Egr3 | -2.07 | 2.447403048 | 0.00356941424 | 3.16E-02 |
| | Plec | -2.08 | 2.482845464 | 0.00328968668 | 3.01E-02 |
| | B230217O12Rik | -2.16 | 1.762134605 | 0.01729280303 | 7.36E-02 |
| | Trhr2 | -2.24 | 2.92807938 | 0.00118010492 | 2.03E-02 |
| | Nr4a2 | -2.25 | 1.898575027 | 0.01263062884 | 6.10E-02 |
| | H60b | -2.27 | 1.517990469 | 0.03033957765 | 1.03E-01 |
| | Tcap | -2.27 | 2.514680224 | 0.00305717131 | 2.94E-02 |
| | Gm13830 | -2.32 | 2.395667345 | 0.00402098687 | 3.34E-02 |
| | Grin1os | -2.38 | 3.320345547 | 0.00047824942 | 1.59E-02 |
| | Six4 | -2.39 | 2.187917767 | 0.00648757264 | 4.27E-02 |
| | 2900055J20Rik | -2.42 | 2.81200159 | 0.00154169481 | 2.23E-02 |
| | Igfn1 | -2.42 | 2.757906768 | 0.00174619698 | 2.34E-02 |
| | Pla2g4b | -2.58 | 2.385467016 | 0.00411654611 | 3.36E-02 |
| | Alox12b | -2.67 | 2.023293005 | 0.00947778810 | 5.24E-02 |
| | Xkr7 | -2.74 | 3.315684328 | 0.00048341005 | 1.59E-02 |
| | Fam227b | -3.42 | 1.934500643 | 0.01162784831 | 5.83E-02 |
| | Lyrm7 | -3.62 | 4.047512996 | 0.00008963694 | 1.41E-02 |
