## Supplementary material for "BRAFV600E Expression in Mouse Neuroglial Progenitors Increase Neuronal Excitability, Cause Appearance of Balloon-like cells, Neuronal Mislocalization, and Inflammatory Immune response": Sup. Table 2. DE genes in BRAF V600E to BRAFwt

| UP p<0.05 | Gene ID | Fold change (BRAF V600E vs. BRAFwt) | log-10 P-values | P-value | FDR step up |
|---|---|---|---|---|---|
| | Cd74 | 154.39 | 2.129027835 | 0.007429715176258620 | 0.136287102 |
| | H2-Aa | 128.49 | 1.970840156 | 0.010694484221932000 | 0.141501604 |
| | Iigp1 | 98.64 | 2.116487396 | 0.007647378816291930 | 0.136287102 |
| | H2-Ab1 | 82.69 | 2.215954024 | 0.006081993847299390 | 0.131469154 |
| | Tgtp1 | 75.67 | 1.550884453 | 0.028126490583718600 | 0.179433875 |
| | Tgtp2 | 62.35 | 1.323598339 | 0.047468079541589400 | 0.217347178 |
| | Ifi47 | 41.34 | 1.523480644 | 0.029958451114424400 | 0.183233156 |
| | Igtp | 35.83 | 1.801909789 | 0.015779390016188800 | 0.152122332 |
| | Psmb9 | 22.62 | 1.849074405 | 0.014155512401210900 | 0.149025511 |
| | H2-DMb1 | 21.46 | 1.897466434 | 0.012662911343761400 | 0.145739007 |
| | Psmb8 | 20.39 | 1.996262256 | 0.010086436150717500 | 0.141095284 |
| | H2-Q7 | 19.38 | 1.742528158 | 0.018091386086750800 | 0.158982987 |
| | Cybb | 16.39 | 2.056375475 | 0.008782628749564740 | 0.140488765 |
| | Gbp6 | 16.20 | 1.918148096 | 0.012074020357045400 | 0.143647440 |
| | Tap1 | 16.16 | 1.949724686 | 0.011227299657927200 | 0.141933868 |
| | Irgm2 | 16.01 | 1.307068107 | 0.049309646992239900 | 0.221281469 |
| | C1s1 | 13.93 | 1.826014215 | 0.014927455508828200 | 0.151437765 |
| | Oasl2 | 13.81 | 1.740757233 | 0.018165308052048600 | 0.159035359 |
| | Casp12 | 13.74 | 2.947910117 | 0.001127430768989640 | 0.127020757 |
| | H2-K1 | 13.20 | 2.055161454 | 0.008807213943254330 | 0.140488765 |
| | H2-Q4 | 13.14 | 1.522985831 | 0.029992603719333100 | 0.183233156 |
| | Lyz2 | 12.50 | 1.865144387 | 0.013641295395169500 | 0.147964655 |

| | | | | | |
|---|---|---|---|---|---|
| | Gbp2 | 12.41 | 2.476246209 | 0.003340056331844350 | 0.128703904 |
| | Bst2 | 11.82 | 1.549823925 | 0.028195258155532900 | 0.179474072 |
| | Cxcl16 | 11.65 | 2.178667925 | 0.006627230479406520 | 0.133240732 |
| | Cd274 | 11.33 | 2.118888111 | 0.007605221880561080 | 0.136287102 |
| | Cd52 | 10.95 | 1.660382022 | 0.021858380329383200 | 0.164554124 |
| | Lbp | 10.50 | 2.767198295 | 0.001709234717186100 | 0.127020757 |
| | Gbp4 | 10.20 | 1.397747496 | 0.040017734998871100 | 0.202460860 |
| | Slfn8 | 9.64 | 1.695509342 | 0.020160006056668500 | 0.160920515 |
| | Lck | 8.89 | 1.330855973 | 0.046681416593040700 | 0.215957866 |
| | Fgl2 | 8.73 | 2.453827435 | 0.003517001589945120 | 0.129362173 |
| | Irf7 | 8.64 | 1.947854133 | 0.011275761123204600 | 0.141933868 |
| | Uba7 | 8.50 | 2.408118964 | 0.003907338498925050 | 0.131162125 |
| | Gbp3 | 8.29 | 1.731869177 | 0.018540900481683800 | 0.159967493 |
| | Rsph4a | 8.23 | 3.017199741 | 0.000961170115160377 | 0.127020757 |
| | C3 | 8.23 | 1.812223224 | 0.015409082352506200 | 0.151749058 |
| | Gbp5 | 8.18 | 1.684108566 | 0.020696239154862600 | 0.161859432 |
| | Csf2rb | 7.95 | 1.694487973 | 0.020207473920083800 | 0.160920515 |
| | Irgm1 | 7.73 | 1.442376631 | 0.036109657518406300 | 0.196819835 |
| | Csf2rb2 | 7.46 | 1.608597547 | 0.024626486396572700 | 0.170079334 |
| | B2m | 7.36 | 1.823910239 | 0.014999948268888100 | 0.151437765 |
| | Sp110 | 6.97 | 1.916759158 | 0.012112696675159400 | 0.143647440 |
| | Irf1 | 6.76 | 2.049439349 | 0.008924022381159370 | 0.140488765 |
| | Top2a | 6.74 | 2.519777151 | 0.003021501741149040 | 0.128703904 |

| | | | | | |
|---|---|---|---|---|---|
| | Parp14 | 6.72 | 1.365778331 | 0.043074641199089500 | 0.208705446 |
| | H2-D1 | 6.63 | 1.535897795 | 0.029114021968780700 | 0.180612088 |
| | Cnmd | 6.62 | 1.627108139 | 0.023598905480916800 | 0.167979916 |
| | 1500015O10Rik | 6.55 | 1.728103869 | 0.018702347876902200 | 0.159967493 |
| | Tnfaip2 | 6.54 | 1.621255577 | 0.023919077349924000 | 0.168887506 |
| | Serping1 | 6.45 | 2.489399668 | 0.003240412748766060 | 0.128703904 |
| | Ptprc | 6.20 | 2.219356958 | 0.006034524318852680 | 0.131469154 |
| | 2410004P03Rik | 6.10 | 1.926226129 | 0.011851515010624900 | 0.143647440 |
| | C1ra | 6.05 | 1.843464768 | 0.014339540395061400 | 0.150411927 |
| | Isg15 | 5.92 | 1.513367015 | 0.030664295069532100 | 0.184968428 |
| | Cd48 | 5.87 | 2.41860838 | 0.003814096000649400 | 0.130257845 |
| | Dlk1 | 5.81 | 2.968931309 | 0.001074159294807140 | 0.127020757 |
| | Cd180 | 5.77 | 2.929372398 | 0.001176596633733830 | 0.127020757 |
| | Bcl2a1d | 5.69 | 2.040626816 | 0.009106954855961410 | 0.140488765 |
| | Rac2 | 5.60 | 1.694327036 | 0.020214963629601800 | 0.160920515 |
| | Mki67 | 5.55 | 1.829701474 | 0.014801254503517500 | 0.151437765 |
| | Tlr2 | 5.53 | 1.840508123 | 0.014437495997797600 | 0.150441568 |
| | Lag3 | 5.47 | 1.608967055 | 0.024605542502215000 | 0.170072817 |
| | B3gnt5 | 5.31 | 2.776595336 | 0.001672648419135390 | 0.127020757 |
| | Ifit3b | 5.21 | 1.747399368 | 0.017889600065961600 | 0.158185968 |
| | Ms4a6c | 5.21 | 1.886576077 | 0.012984460895565200 | 0.146280491 |
| | Gbp7 | 5.11 | 1.43439167 | 0.036779712502892000 | 0.198192168 |
| | Gimap4 | 5.11 | 1.58654818 | 0.025909069745827300 | 0.174261638 |

| | | | | | |
|---|---|---|---|---|---|
| | Irf8 | 5.05 | 2.248100838 | 0.005648058183937650 | 0.131469154 |
| | Serpina3h | 5.04 | 1.929156167 | 0.011771825966976300 | 0.143647440 |
| | Fam46c | 5.03 | 1.358007724 | 0.043852289857182700 | 0.210611848 |
| | Ifit1 | 5.01 | 2.068849992 | 0.008533948306108860 | 0.140434588 |
| | Melk | 4.92 | 2.087344067 | 0.008178166210824740 | 0.138564959 |
| | Ifitm3 | 4.91 | 2.216257954 | 0.006077739005348350 | 0.131469154 |
| | Trim30a | 4.84 | 1.448871933 | 0.035573620427615900 | 0.196075860 |
| | Prss22 | 4.70 | 2.152691184 | 0.007035724353205700 | 0.134660365 |
| | Ms4a6b | 4.66 | 1.791687592 | 0.016155202557577100 | 0.152370846 |
| | Lgals3bp | 4.66 | 1.670403784 | 0.021359752483285700 | 0.163057111 |
| | Col5a2 | 4.61 | 2.872459612 | 0.001341344669496120 | 0.127020757 |
| | Togaram2 | 4.60 | 2.943382536 | 0.001139245873027230 | 0.127020757 |
| | Fam183b | 4.60 | 2.03377361 | 0.009251803292751690 | 0.140488765 |
| | Pycard | 4.58 | 2.003555535 | 0.009918464993094220 | 0.141095284 |
| | Slc11a1 | 4.57 | 1.866959705 | 0.013584394810660900 | 0.147964655 |
| | Ctsc | 4.51 | 2.047648827 | 0.008960890555719000 | 0.140488765 |
| | Samsn1 | 4.51 | 2.476665688 | 0.003336831774469920 | 0.128703904 |
| | Tnc | 4.44 | 2.431956719 | 0.003698650384426510 | 0.129856967 |
| | Slc15a3 | 4.43 | 2.209196169 | 0.006177373079910500 | 0.131469154 |
| | Zcchc12 | 4.42 | 3.018517715 | 0.000958257628594116 | 0.127020757 |
| | H2-DMa | 4.39 | 1.432573059 | 0.036934050712112300 | 0.198531875 |
| | Hvcn1 | 4.36 | 1.542799895 | 0.028654979701113400 | 0.179884640 |
| | Baiap3 | 4.35 | 1.305259328 | 0.049515443384727000 | 0.221308908 |

| | | | | | |
|---|---|---|---|---|---|
| | Cd24a | 4.33 | 2.250030502 | 0.005623018308991990 | 0.131469154 |
| | Icam1 | 4.30 | 1.67296204 | 0.021234300551018100 | 0.162736895 |
| | Tap2 | 4.30 | 1.673571787 | 0.021204508641529800 | 0.162736895 |
| | Iqub | 4.29 | 1.782209056 | 0.016511667844219700 | 0.153363089 |
| | Rhoh | 4.28 | 1.807487171 | 0.015578040497284900 | 0.151848760 |
| | Naip2 | 4.24 | 1.478706869 | 0.033211854825937800 | 0.190867460 |
| | Capg | 4.23 | 2.272319335 | 0.005341714405261250 | 0.131469154 |
| | Pik3ap1 | 4.20 | 1.352377982 | 0.044424445706099800 | 0.211327864 |
| | Sp100 | 4.20 | 1.475580724 | 0.033451783390253100 | 0.191129135 |
| | Cyp1b1 | 4.20 | 2.458137274 | 0.003482272279541990 | 0.129362173 |
| | Ncf4 | 4.18 | 1.647125124 | 0.022535898382249400 | 0.166458646 |
| | Cd86 | 4.18 | 2.008441073 | 0.009807513776485340 | 0.140783376 |
| | Parp9 | 4.16 | 1.709294867 | 0.019530129953121800 | 0.160482390 |
| | Trim12c | 4.11 | 2.179659021 | 0.006612123834528280 | 0.133240732 |
| | Cd44 | 4.10 | 2.514482119 | 0.003058566171442040 | 0.128703904 |
| | Emp3 | 4.09 | 2.251774043 | 0.005600489113103380 | 0.131469154 |
| | Serpina3i | 4.06 | 1.466856036 | 0.034130603201420000 | 0.192208450 |
| | Itgb2 | 4.06 | 1.877433125 | 0.013260712964715200 | 0.146863953 |
| | A2m | 4.06 | 1.97556721 | 0.010578711909163300 | 0.141402802 |
| | Ubxn10 | 4.04 | 2.71697447 | 0.001918781534629820 | 0.127020757 |
| | Ly86 | 4.01 | 2.058262262 | 0.008744555498861910 | 0.140488765 |
| | S100a4 | 4.01 | 2.459372267 | 0.003472383891694720 | 0.129362173 |
| | Ccl6 | 4.00 | 1.597101459 | 0.025287071733030100 | 0.171929566 |

| | | | | | |
|---|---|---|---|---|---|
| | Myo1f | 3.99 | 1.87556449 | 0.013317892659391200 | 0.146863953 |
| | Ptpn6 | 3.98 | 1.437691581 | 0.036501307323998000 | 0.197466324 |
| | Lcp1 | 3.94 | 1.956607463 | 0.011050769935195600 | 0.141933868 |
| | Shisa8 | 3.93 | 1.594580233 | 0.025434298699792800 | 0.172537713 |
| | Ifi30 | 3.92 | 1.360735966 | 0.043577672798006500 | 0.209958205 |
| | Tmem106a | 3.92 | 1.408951812 | 0.038998525565102900 | 0.201744197 |
| | Foxj1 | 3.92 | 2.105447083 | 0.007844276926399070 | 0.137039524 |
| | Lsp1 | 3.91 | 1.716220168 | 0.019221170529104800 | 0.160482390 |
| | Osr1 | 3.91 | 2.381727409 | 0.004152145765099100 | 0.131469154 |
| | Sash3 | 3.91 | 1.797900712 | 0.015925727778343800 | 0.152122332 |
| | Crybg1 | 3.87 | 1.787402094 | 0.016315406790421600 | 0.152555604 |
| | Ifi203 | 3.86 | 1.646134884 | 0.022587341374629100 | 0.166458646 |
| | Cxcr4 | 3.85 | 1.563110639 | 0.027345719913564500 | 0.178337702 |
| | Col3a1 | 3.83 | 2.663994509 | 0.002167731512793850 | 0.127020757 |
| | Fam167a | 3.83 | 2.333406609 | 0.004640805756748350 | 0.131469154 |
| | Hck | 3.82 | 1.987870693 | 0.010283224253785200 | 0.141095284 |
| | Mamdc2 | 3.82 | 1.686039248 | 0.020604436965838400 | 0.161859432 |
| | Gsdmd | 3.81 | 1.487684021 | 0.032532390645587700 | 0.188778539 |
| | Apobec1 | 3.79 | 2.425714671 | 0.003752194382853000 | 0.129856967 |
| | Ctss | 3.79 | 2.192770626 | 0.006415483227783410 | 0.132629528 |
| | Glipr2 | 3.76 | 2.239609696 | 0.005759573221877200 | 0.131469154 |
| | C3ar1 | 3.75 | 1.836218516 | 0.014580804402237000 | 0.150869603 |
| | Ccdc40 | 3.73 | 1.795008892 | 0.016032125639386500 | 0.152122332 |

| | | | | | |
|---|---|---|---|---|---|
| | Fcgr1 | 3.73 | 1.928471095 | 0.011790409940162600 | 0.143647440 |
| | Gm11992 | 3.73 | 2.593240754 | 0.002551286585738150 | 0.127020757 |
| | C2 | 3.69 | 1.450247864 | 0.035461094526782500 | 0.195808417 |
| | Tspo | 3.68 | 1.988850667 | 0.010260046587569500 | 0.141095284 |
| | Prrx2 | 3.65 | 2.042331325 | 0.009071282152788980 | 0.140488765 |
| | Aldh1a2 | 3.64 | 3.218119129 | 0.000605174849574600 | 0.123542711 |
| | Fyb | 3.64 | 1.676431188 | 0.021065356421869500 | 0.162610003 |
| | Vim | 3.63 | 2.375004555 | 0.004216920805384960 | 0.131469154 |
| | P2ry6 | 3.63 | 2.432319301 | 0.003695563756653320 | 0.129856967 |
| | Plscr2 | 3.61 | 1.927343138 | 0.011821071972102900 | 0.143647440 |
| | Klk6 | 3.60 | 1.620466413 | 0.023962580630668100 | 0.168887506 |
| | C1qb | 3.60 | 1.864897376 | 0.013649056260540900 | 0.147964655 |
| | Dcn | 3.60 | 2.477506622 | 0.003330376842967380 | 0.128703904 |
| | Cpxm2 | 3.59 | 2.746940616 | 0.001790850712633610 | 0.127020757 |
| | Hspb8 | 3.59 | 3.131933219 | 0.000738017704943420 | 0.123542711 |
| | C1qa | 3.57 | 1.750762962 | 0.017751580963217300 | 0.157554283 |
| | Birc3 | 3.56 | 1.881024567 | 0.013151504348848500 | 0.146825043 |
| | Islr | 3.54 | 2.087717616 | 0.008171134965332360 | 0.138564959 |
| | Mns1 | 3.51 | 2.739362153 | 0.001822375409670420 | 0.127020757 |
| | Il16 | 3.49 | 1.391473676 | 0.040600027169264700 | 0.203309852 |
| | Nfkb2 | 3.47 | 1.495803899 | 0.031929792858115100 | 0.187100141 |
| | Igsf1 | 3.47 | 3.194136871 | 0.000639533250644370 | 0.123542711 |
| | Cyba | 3.46 | 1.877032949 | 0.013272937538299500 | 0.146863953 |

| | | | | | |
|---|---|---|---|---|---|
| | Tfap2b | 3.45 | 2.663582312 | 0.002169789922646010 | 0.127020757 |
| | Fbln1 | 3.44 | 1.578796873 | 0.026375647331059000 | 0.175170503 |
| | Apobec3 | 3.42 | 2.611601633 | 0.002445672873815330 | 0.127020757 |
| | Trim21 | 3.42 | 1.304931904 | 0.049552788179571800 | 0.221308908 |
| | Ifi35 | 3.42 | 1.304991582 | 0.049545979483359000 | 0.221308908 |
| | Parp12 | 3.40 | 2.365070429 | 0.004314491037348110 | 0.131469154 |
| | Clec5a | 3.40 | 1.925835877 | 0.011862169435554900 | 0.143647440 |
| | Thsd4 | 3.39 | 2.269389757 | 0.005377869299054900 | 0.131469154 |
| | Cd84 | 3.38 | 1.498809694 | 0.031709566560837700 | 0.186764318 |
| | Hk3 | 3.38 | 1.336832402 | 0.046043422490918800 | 0.214520487 |
| | Cd109 | 3.37 | 2.107514282 | 0.007807027651718860 | 0.137039524 |
| | Lgals9 | 3.37 | 1.862589775 | 0.013721772818634900 | 0.147964655 |
| | Rbp1 | 3.36 | 2.728024636 | 0.001870576027959680 | 0.127020757 |
| | Lox | 3.35 | 2.602225894 | 0.002499045170806890 | 0.127020757 |
| | Hpse | 3.34 | 1.303842515 | 0.049677243017781200 | 0.221317530 |
| | C1qc | 3.34 | 1.565534818 | 0.027193504639612400 | 0.177766864 |
| | Slc13a4 | 3.33 | 3.318344189 | 0.000480458423009719 | 0.123542711 |
| | Akr1c14 | 3.33 | 2.712904802 | 0.001936846477541770 | 0.127020757 |
| | Samd9l | 3.32 | 1.684894269 | 0.020658830439718900 | 0.161859432 |
| | Tlr9 | 3.31 | 1.421622575 | 0.037877161458036000 | 0.199763957 |
| | Ncf1 | 3.30 | 2.331486743 | 0.004661366572136180 | 0.131469154 |
| | Emilin1 | 3.28 | 1.53830962 | 0.028952787289059000 | 0.180151698 |
| | Cmtm7 | 3.26 | 1.947173252 | 0.011293452970021200 | 0.141933868 |

| | | | | | |
|---|---|---|---|---|---|
| | Tekt1 | 3.25 | 2.017945433 | 0.009595211825471850 | 0.140783376 |
| | Tbxas1 | 3.25 | 2.107913354 | 0.007799857093884260 | 0.137039524 |
| | Cfap44 | 3.25 | 1.919633528 | 0.012032793719183200 | 0.143647440 |
| | Zc3hav1 | 3.23 | 1.811444954 | 0.015436720684597300 | 0.151749058 |
| | Unc93b1 | 3.22 | 1.949318089 | 0.011237815848519300 | 0.141933868 |
| | Fcgr2b | 3.22 | 1.639556905 | 0.022932061362949900 | 0.166831290 |
| | Hcls1 | 3.22 | 1.808542161 | 0.015540244198404500 | 0.151768654 |
| | Mmp19 | 3.20 | 1.352558839 | 0.044405949564143200 | 0.211327864 |
| | S1pr3 | 3.20 | 2.42842702 | 0.003728833401675250 | 0.129856967 |
| | C4b | 3.20 | 1.400379127 | 0.039775978589059600 | 0.202412377 |
| | Nnat | 3.19 | 2.411358709 | 0.003878299028749380 | 0.130937855 |
| | Ddx58 | 3.17 | 1.985979462 | 0.010328102455479100 | 0.141095284 |
| | Pld4 | 3.17 | 1.879426179 | 0.013199996652754600 | 0.146825043 |
| | Hcn4 | 3.16 | 1.653312222 | 0.022217120906783000 | 0.165260680 |
| | Sdc1 | 3.15 | 1.397341606 | 0.040055152949498600 | 0.202575582 |
| | Tgif1 | 3.15 | 1.979592046 | 0.010481126287551600 | 0.141095284 |
| | Klhl6 | 3.15 | 1.867887921 | 0.013555391916945900 | 0.147964655 |
| | Kcne1l | 3.14 | 2.252474653 | 0.005591461602805550 | 0.131469154 |
| | Dnah5 | 3.14 | 1.95265524 | 0.011151794564460800 | 0.141933868 |
| | Lst1 | 3.14 | 2.33899357 | 0.004581486700071940 | 0.131469154 |
| | Cd37 | 3.14 | 1.817723328 | 0.015215165212243500 | 0.151522906 |
| | Adgre1 | 3.13 | 1.713866946 | 0.019325603022009400 | 0.160482390 |
| | Vit | 3.12 | 2.124936737 | 0.007500034536829760 | 0.136287102 |

| | | | | | |
|---|---|---|---|---|---|
| | Dock2 | 3.12 | 1.81889582 | 0.015174143276281100 | 0.151522906 |
| | Btk | 3.10 | 1.400523561 | 0.039762752380621000 | 0.202412377 |
| | Svep1 | 3.09 | 2.568068375 | 0.002703532690293400 | 0.128544092 |
| | Fgfbp1 | 3.09 | 3.001314949 | 0.000996976796424005 | 0.127020757 |
| | Flnc | 3.06 | 2.144837874 | 0.007164108025552050 | 0.135222539 |
| | Ppp1r36 | 3.05 | 1.869894304 | 0.013492912245437000 | 0.147616327 |
| | Ccl9 | 3.05 | 2.019200225 | 0.009567528720379330 | 0.140783376 |
| | Slc7a7 | 3.04 | 1.703501744 | 0.019792390718017900 | 0.160671218 |
| | Uhrf1 | 3.03 | 1.680965037 | 0.020846587012180700 | 0.162388295 |
| | Lpo | 3.03 | 1.563410371 | 0.027326853501144800 | 0.178328856 |
| | Dok1 | 3.03 | 2.302232968 | 0.004986169434808630 | 0.131469154 |
| | Slc43a3 | 3.01 | 1.47922507 | 0.033172250007692200 | 0.190719618 |
| | Ogn | 3.01 | 2.814951449 | 0.001531258636869200 | 0.127020757 |
| | Klk8 | 3.01 | 2.138493605 | 0.007269531035428250 | 0.135646175 |
| | Hfe | 3.01 | 1.768255837 | 0.017050776533161400 | 0.155633632 |
| | Runx1 | 3.00 | 1.908453807 | 0.012346566303777300 | 0.145127603 |
| | Ccl17 | 2.99 | 1.782327723 | 0.016507156827314200 | 0.153363089 |
| | Cdh1 | 2.99 | 2.023904721 | 0.009464447782043420 | 0.140783376 |
| | Cdca7l | 2.97 | 2.210828206 | 0.006154202669740830 | 0.131469154 |
| | Pon3 | 2.97 | 1.49281911 | 0.032149993563519600 | 0.187746849 |
| | S100a6 | 2.97 | 2.700390234 | 0.001993470281591060 | 0.127020757 |
| | Esr1 | 2.95 | 1.86497332 | 0.013646669705151800 | 0.147964655 |
| | Enpp1 | 2.95 | 5.310632126 | 0.000004890664526575 | 0.048608941 |

| | | | | | |
|---|---|---|---|---|---|
| | Col1a1 | 2.94 | 2.633841044 | 0.002323587093738530 | 0.127020757 |
| | Arhgap30 | 2.94 | 1.509142488 | 0.030964032305856600 | 0.185473744 |
| | Ccdc88b | 2.93 | 1.583108958 | 0.026115060824779800 | 0.174737719 |
| | Lat2 | 2.93 | 1.828258943 | 0.014850499351852800 | 0.151437765 |
| | Serpind1 | 2.92 | 3.984230024 | 0.000103697903546761 | 0.112761169 |
| | Irf5 | 2.92 | 1.886759735 | 0.012978971090989900 | 0.146280491 |
| | Rhbdf2 | 2.91 | 1.321123675 | 0.047739330628218700 | 0.217912197 |
| | Vav1 | 2.91 | 1.760624012 | 0.017353056791478900 | 0.156718571 |
| | Ms4a6d | 2.91 | 1.378691203 | 0.041812756299386300 | 0.205721435 |
| | Zc3h12a | 2.89 | 2.739544647 | 0.001821609794297160 | 0.127020757 |
| | Col1a2 | 2.89 | 2.602006928 | 0.002500305476200180 | 0.127020757 |
| | Bmp6 | 2.87 | 3.566131144 | 0.000271561911028221 | 0.123542711 |
| | Hacd4 | 2.87 | 1.694218202 | 0.020220030132184700 | 0.160920515 |
| | Col18a1 | 2.86 | 1.620442107 | 0.023963921751553200 | 0.168887506 |
| | Dsg2 | 2.86 | 2.488054032 | 0.003250468550397970 | 0.128703904 |
| | Adamtsl3 | 2.85 | 1.647269119 | 0.022528427632665700 | 0.166458646 |
| | Mgst1 | 2.85 | 2.160795773 | 0.006905644652359300 | 0.134513024 |
| | Antxr2 | 2.85 | 2.488358942 | 0.003248187255393460 | 0.128703904 |
| | S100a11 | 2.85 | 2.007551743 | 0.009827617783023280 | 0.140783376 |
| | Pik3r5 | 2.84 | 1.702051739 | 0.019858583233783300 | 0.160735761 |
| | Clic1 | 2.84 | 2.160685008 | 0.006907406139936510 | 0.134513024 |
| | Gypc | 2.84 | 1.840465649 | 0.014438908075662300 | 0.150441568 |
| | Sulf1 | 2.83 | 1.894396607 | 0.012752736695406900 | 0.145878492 |

| | | | | | |
|---|---|---|---|---|---|
| | Pqlc3 | 2.83 | 1.843086726 | 0.014352028032466500 | 0.150428083 |
| | Srgn | 2.83 | 1.631787653 | 0.023345992801709400 | 0.167518166 |
| | Zic1 | 2.83 | 2.039572456 | 0.009129091171389220 | 0.140488765 |
| | Ctsh | 2.81 | 2.034106138 | 0.009244722139413710 | 0.140488765 |
| | Serpinf1 | 2.81 | 2.164302475 | 0.006850109673949790 | 0.134513024 |
| | Ccrl2 | 2.81 | 1.52907304 | 0.029575150298140400 | 0.182155577 |
| | Cacng5 | 2.81 | 1.684176344 | 0.020693009441098000 | 0.161859432 |
| | Col8a2 | 2.80 | 2.596900994 | 0.002529874666188530 | 0.127020757 |
| | Dtx3l | 2.80 | 1.432349389 | 0.036953077315633600 | 0.198531875 |
| | Ccdc96 | 2.80 | 2.409120791 | 0.003898335463541810 | 0.131162125 |
| | Bgn | 2.79 | 2.940808183 | 0.001146018998847760 | 0.127020757 |
| | Wnt6 | 2.79 | 2.012772676 | 0.009710180967467860 | 0.140783376 |
| | Cntf | 2.79 | 2.073739045 | 0.008438416453478520 | 0.140434588 |
| | Bin2 | 2.79 | 2.415702618 | 0.003839700783841550 | 0.130257845 |
| | Vcam1 | 2.79 | 3.025943066 | 0.000942013083054663 | 0.127020757 |
| | 1700007K13Rik | 2.78 | 2.879877308 | 0.001318629210056410 | 0.127020757 |
| | Aox3 | 2.78 | 2.23056645 | 0.005880761280022420 | 0.131469154 |
| | Syngr2 | 2.76 | 2.930644538 | 0.001173155177252020 | 0.127020757 |
| | Rcn3 | 2.76 | 2.742131535 | 0.001810791575602980 | 0.127020757 |
| | Col4a6 | 2.76 | 2.265660198 | 0.005424251305268280 | 0.131469154 |
| | Cp | 2.75 | 1.964502291 | 0.010851698248624400 | 0.141933868 |
| | Nrp2 | 2.75 | 2.1254173 | 0.007491740054532330 | 0.136287102 |
| | Trem2 | 2.75 | 2.485489448 | 0.003269719913162840 | 0.128703904 |

| | | | | | |
|---|---|---|---|---|---|
| | Efcab1 | 2.75 | 2.836125036 | 0.001458394318637970 | 0.127020757 |
| | Slc2a12 | 2.75 | 2.554135679 | 0.002791671551571050 | 0.128703904 |
| | Psme1 | 2.74 | 2.152312479 | 0.007041862182712220 | 0.134660365 |
| | Rarres2 | 2.73 | 2.484362856 | 0.003278212820503840 | 0.128703904 |
| | Plcg2 | 2.73 | 2.106679648 | 0.007822045757374510 | 0.137039524 |
| | Thbs1 | 2.73 | 1.71001982 | 0.019497556149192700 | 0.160482390 |
| | Asap3 | 2.73 | 2.150490189 | 0.007071471746250140 | 0.134660365 |
| | Fam111a | 2.72 | 1.301201101 | 0.049980304694250600 | 0.221634296 |
| | Adcy7 | 2.72 | 2.188191298 | 0.006483487858981540 | 0.133060702 |
| | Loxl1 | 2.71 | 2.779134245 | 0.001662898550819340 | 0.127020757 |
| | Aif1 | 2.70 | 1.942878327 | 0.011405692884273600 | 0.141961618 |
| | Fbln7 | 2.70 | 2.475461199 | 0.003346099116581830 | 0.128703904 |
| | Pgf | 2.70 | 2.285609752 | 0.005180721513838360 | 0.131469154 |
| | Nek2 | 2.70 | 1.996269265 | 0.010086273371001400 | 0.141095284 |
| | Clec2d | 2.69 | 1.429429204 | 0.037202386100848300 | 0.198909723 |
| | Mmp2 | 2.69 | 2.443703455 | 0.003599950634695670 | 0.129856967 |
| | Cd14 | 2.69 | 2.20649128 | 0.006215967280011220 | 0.131469154 |
| | Slc12a7 | 2.68 | 1.830429842 | 0.014776451675664100 | 0.151437765 |
| | Aebp1 | 2.68 | 3.364465007 | 0.000432050978844704 | 0.123542711 |
| | Pdlim4 | 2.68 | 2.436160986 | 0.003663017670313300 | 0.129856967 |
| | Tnfaip8l2 | 2.68 | 1.349904756 | 0.044678156405140700 | 0.211647507 |
| | Gfap | 2.67 | 1.513454197 | 0.030658139980834000 | 0.184968428 |
| | Icosl | 2.66 | 1.570010853 | 0.026914675461247200 | 0.176630258 |

| | | | | | |
|---|---|---|---|---|---|
| | Mrgprf | 2.65 | 2.048311427 | 0.008947229413622170 | 0.140488765 |
| | Mxra8 | 2.65 | 2.712584725 | 0.001938274466572110 | 0.127020757 |
| | Rbl1 | 2.65 | 1.686401022 | 0.020587280306784700 | 0.161859432 |
| | Anxa1 | 2.65 | 2.069573096 | 0.008519751029518700 | 0.140434588 |
| | Fcer1g | 2.65 | 1.519213725 | 0.030254241918674400 | 0.183623471 |
| | Rasa4 | 2.65 | 2.035478381 | 0.009215557614740350 | 0.140488765 |
| | Arhgap45 | 2.64 | 1.462997474 | 0.034435193346989000 | 0.192834679 |
| | Zic4 | 2.64 | 1.8713005 | 0.013449294396626600 | 0.147515895 |
| | Hk2 | 2.64 | 1.379856476 | 0.041700717121320300 | 0.205721435 |
| | Rnaset2b | 2.63 | 1.920104859 | 0.012019741868103700 | 0.143647440 |
| | Tyrobp | 2.63 | 2.290321269 | 0.005124821351131570 | 0.131469154 |
| | Anxa2 | 2.62 | 2.243810735 | 0.005704128036289460 | 0.131469154 |
| | Pla2g4a | 2.62 | 2.711351871 | 0.001943784565728390 | 0.127020757 |
| | Csf3r | 2.61 | 2.256811218 | 0.005535906952131110 | 0.131469154 |
| | Pgm5 | 2.61 | 3.155067052 | 0.000699733953353644 | 0.123542711 |
| | Plscr1 | 2.61 | 1.509532788 | 0.030936217487278000 | 0.185473744 |
| | Mtbp | 2.61 | 2.789054264 | 0.001625345660728820 | 0.127020757 |
| | Slc1a5 | 2.60 | 2.377527294 | 0.004192496471285540 | 0.131469154 |
| | Hcar1 | 2.60 | 2.861604352 | 0.001375294316994670 | 0.127020757 |
| | Gpsm3 | 2.59 | 1.306784713 | 0.049341833888357800 | 0.221308908 |
| | Ifit2 | 2.59 | 1.709024623 | 0.019542286542008300 | 0.160482390 |
| | Gpx8 | 2.59 | 2.897195898 | 0.001267080193524980 | 0.127020757 |
| | Dpyd | 2.59 | 1.43523775 | 0.036708129031318900 | 0.197920025 |

| | | | | | |
|---|---|---|---|---|---|
| | Alx4 | 2.59 | 1.805946251 | 0.015633411119684300 | 0.151922703 |
| | Postn | 2.59 | 2.183598586 | 0.006552415266433370 | 0.133060702 |
| | Ube2l6 | 2.58 | 1.407542209 | 0.039125309973862400 | 0.201809641 |
| | Ptgfr | 2.58 | 2.162634698 | 0.006876466028993840 | 0.134513024 |
| | Tmem173 | 2.57 | 1.707020086 | 0.019632694755480800 | 0.160483557 |
| | Tes | 2.57 | 1.639295877 | 0.022945848536794500 | 0.166831290 |
| | Tlr13 | 2.56 | 2.336583401 | 0.004606982883597720 | 0.131469154 |
| | Mcm3 | 2.56 | 1.931317828 | 0.011713378365436000 | 0.143647440 |
| | Sla | 2.56 | 2.155398939 | 0.006991994230496810 | 0.134660365 |
| | Ncf2 | 2.55 | 2.503163658 | 0.003139325457633960 | 0.128703904 |
| | Trpc7 | 2.55 | 1.799414378 | 0.015870317757450500 | 0.152122332 |
| | Pdgfrl | 2.53 | 2.148427509 | 0.007105137567782100 | 0.134660365 |
| | Cfap206 | 2.52 | 1.826655176 | 0.014905440811630500 | 0.151437765 |
| | Ifi27 | 2.52 | 1.511895155 | 0.030768395224057700 | 0.185189890 |
| | Nfkbie | 2.52 | 2.00983511 | 0.009776083227925230 | 0.140783376 |
| | Wdfy4 | 2.52 | 1.469407866 | 0.033930646444523800 | 0.191843625 |
| | Crabp2 | 2.51 | 1.543296432 | 0.028622236658786800 | 0.179884640 |
| | Tor4a | 2.51 | 1.355459619 | 0.044110337482936300 | 0.210751094 |
| | Nmi | 2.51 | 1.62155306 | 0.023902698840835200 | 0.168887506 |
| | Ptgr1 | 2.51 | 1.604851985 | 0.024839795423215500 | 0.170576526 |
| | Adgrg6 | 2.51 | 2.384845239 | 0.004122443967968770 | 0.131469154 |
| | Tapbp | 2.51 | 1.690622142 | 0.020388151794089200 | 0.161483332 |
| | Rinl | 2.51 | 1.66942315 | 0.021408037145463800 | 0.163245193 |

| | | | | | |
|---|---|---|---|---|---|
| | Dnah9 | 2.51 | 1.892672705 | 0.012803458411266800 | 0.145881393 |
| | Kcnj13 | 2.51 | 1.391771863 | 0.040572160688149100 | 0.203309852 |
| | Cd53 | 2.51 | 2.360899569 | 0.004356125983321010 | 0.131469154 |
| | Samhd1 | 2.50 | 1.686937765 | 0.020561852274751700 | 0.161859432 |
| | Cd83 | 2.50 | 2.088124555 | 0.008163482109376110 | 0.138564959 |
| | Mmp14 | 2.50 | 1.568542577 | 0.027005823400931800 | 0.176876558 |
| | Vstm4 | 2.49 | 2.663282565 | 0.002171288013789970 | 0.127020757 |
| | 4933407L21Rik | 2.49 | 2.50647165 | 0.003115504262704090 | 0.128703904 |
| | Eya2 | 2.49 | 1.599315677 | 0.025158475636297000 | 0.171564573 |
| | Ikzf1 | 2.49 | 1.548204576 | 0.028300585746872800 | 0.179474072 |
| | Rrm2 | 2.49 | 1.82362783 | 0.015009705464521100 | 0.151437765 |
| | F5 | 2.49 | 1.547012113 | 0.028378398778647500 | 0.179474072 |
| | Gstt3 | 2.49 | 1.720875107 | 0.019016250648727500 | 0.160480708 |
| | Angpt1 | 2.48 | 1.61400428 | 0.024321800387948700 | 0.169664195 |
| | Pla2g5 | 2.48 | 1.678614047 | 0.020959742986566800 | 0.162532635 |
| | Enkur | 2.48 | 1.580660552 | 0.026262704534173500 | 0.174878392 |
| | Ccna2 | 2.48 | 1.994324494 | 0.010131540997670800 | 0.141095284 |
| | Prkcd | 2.47 | 1.959110503 | 0.010987262410353600 | 0.141933868 |
| | Gstm2 | 2.47 | 1.477593667 | 0.033297094069798500 | 0.191129135 |
| | Efemp1 | 2.47 | 2.436204074 | 0.003662654269161990 | 0.129856967 |
| | Ctxn3 | 2.47 | 1.679122805 | 0.020935203883754800 | 0.162468873 |
| | Chil1 | 2.47 | 1.745557012 | 0.017965652213317900 | 0.158552362 |
| | Bicc1 | 2.47 | 2.843228018 | 0.001434735954712120 | 0.127020757 |

| | | | | | |
|---|---|---|---|---|---|
| | Plpp4 | 2.47 | 2.533254164 | 0.002929178488240010 | 0.128703904 |
| | Sema3b | 2.46 | 1.863917634 | 0.013679882472825400 | 0.147964655 |
| | Txnip | 2.45 | 2.192556471 | 0.006418647547543720 | 0.132629528 |
| | Erap1 | 2.45 | 2.00497308 | 0.009886143731328440 | 0.141064902 |
| | Rsph1 | 2.45 | 3.400181029 | 0.000397941259896742 | 0.123542711 |
| | Wnt5a | 2.45 | 2.474543594 | 0.003353176439855610 | 0.128703904 |
| | Mrc2 | 2.45 | 1.988799118 | 0.010261264488065900 | 0.141095284 |
| | Clec4a2 | 2.44 | 2.2372917 | 0.005790396461477830 | 0.131469154 |
| | Tlr6 | 2.44 | 1.527770325 | 0.029663997472002600 | 0.182340276 |
| | Tmem123 | 2.44 | 2.039985258 | 0.009120417978180110 | 0.140488765 |
| | Dynlt1f | 2.44 | 2.117252918 | 0.007633910824817950 | 0.136287102 |
| | Mrc1 | 2.44 | 1.918478197 | 0.012064846572356500 | 0.143647440 |
| | Micall2 | 2.43 | 1.495908739 | 0.031922085806008200 | 0.187100141 |
| | Plin2 | 2.42 | 2.471814747 | 0.003374312127261190 | 0.129154381 |
| | Stom | 2.41 | 1.728109815 | 0.018702091835383200 | 0.159967493 |
| | Pawr | 2.40 | 2.023329961 | 0.009476981633290870 | 0.140783376 |
| | Osmr | 2.40 | 1.592229566 | 0.025572337858444700 | 0.173183585 |
| | Lrrk1 | 2.40 | 1.774510355 | 0.016806978522800800 | 0.154749105 |
| | Il13ra1 | 2.39 | 2.307817924 | 0.004922458638917100 | 0.131469154 |
| | Adam12 | 2.39 | 1.359039235 | 0.043748258056815700 | 0.210484879 |
| | Crip1 | 2.39 | 1.946914979 | 0.011300171138743300 | 0.141933868 |
| | Slc22a6 | 2.39 | 2.821075065 | 0.001509819167821890 | 0.127020757 |
| | Aldh3b1 | 2.39 | 2.98570271 | 0.001033468610808570 | 0.127020757 |

| | | | | | |
|---|---|---|---|---|---|
| | Hs3st3a1 | 2.39 | 2.116547712 | 0.007646316798928870 | 0.136287102 |
| | Cd82 | 2.39 | 2.636418558 | 0.002309837575532770 | 0.127020757 |
| | Dynlt1c | 2.38 | 1.861290238 | 0.013762893915359700 | 0.147977908 |
| | Serpina3n | 2.38 | 1.393308291 | 0.040428879897605000 | 0.203278828 |
| | Parp3 | 2.37 | 2.285445782 | 0.005182677889568480 | 0.131469154 |
| | Tbx15 | 2.37 | 1.897540414 | 0.012660754452663900 | 0.145739007 |
| | Igf2 | 2.37 | 3.126534531 | 0.000747249217104334 | 0.123542711 |
| | Bche | 2.36 | 2.544783079 | 0.002852442648134860 | 0.128703904 |
| | BC026585 | 2.36 | 2.076746244 | 0.008380187884990410 | 0.140384356 |
| | Slc6a13 | 2.36 | 2.151693812 | 0.007051900696404040 | 0.134660365 |
| | Slc6a20a | 2.35 | 3.166791179 | 0.000681096769945875 | 0.123542711 |
| | Capn3 | 2.35 | 1.475271228 | 0.033475631001229000 | 0.191129135 |
| | Rassf9 | 2.35 | 1.799510647 | 0.015866800196375700 | 0.152122332 |
| | Arhgap6 | 2.34 | 1.65165963 | 0.022301823283246900 | 0.165648299 |
| | Igsf10 | 2.34 | 1.498338888 | 0.031743960671635300 | 0.186886788 |
| | Tnfrsf1a | 2.34 | 2.045668974 | 0.009001834545365380 | 0.140488765 |
| | Tagln2 | 2.34 | 2.172604076 | 0.006720412390599220 | 0.133677072 |
| | Mob3b | 2.33 | 3.081356683 | 0.000829169496354415 | 0.123542711 |
| | Ckap2l | 2.33 | 1.654791282 | 0.022141585569144700 | 0.165260680 |
| | Casp8 | 2.33 | 1.765496576 | 0.017159452396138000 | 0.156151017 |
| | Cdca7 | 2.33 | 2.920255518 | 0.001201557287573080 | 0.127020757 |
| | Plek | 2.33 | 1.728725242 | 0.018675608351591700 | 0.159967493 |
| | Bambi | 2.32 | 2.486217882 | 0.003264240268473740 | 0.128703904 |

| | | | | | |
|---|---|---|---|---|---|
| | Foxd1 | 2.32 | 1.415664838 | 0.038400348152368400 | 0.200478413 |
| | Fgd2 | 2.32 | 1.606337914 | 0.024754951877901600 | 0.170282757 |
| | Pdpn | 2.32 | 2.47860363 | 0.003321975078129390 | 0.128703904 |
| | Fam180a | 2.32 | 1.607926466 | 0.024664569193816600 | 0.170091091 |
| | Cped1 | 2.31 | 3.106831813 | 0.000781930560585841 | 0.123542711 |
| | Serpinb9 | 2.31 | 2.352653309 | 0.004439629120065720 | 0.131469154 |
| | Spata6 | 2.31 | 2.568564347 | 0.002700446971172030 | 0.128544092 |
| | Dapp1 | 2.31 | 2.699949274 | 0.001995495378920250 | 0.127020757 |
| | Slc25a45 | 2.31 | 2.042587568 | 0.009065931478833780 | 0.140488765 |
| | Col5a3 | 2.30 | 2.681698589 | 0.002081140550315170 | 0.127020757 |
| | Vasp | 2.29 | 1.77929274 | 0.016622917916969100 | 0.153711652 |
| | Tbx18 | 2.29 | 1.966481487 | 0.010802356666868500 | 0.141501604 |
| | Cnn3 | 2.29 | 2.121990577 | 0.007551086117602900 | 0.136287102 |
| | Kif11 | 2.29 | 2.020749632 | 0.009533456043689600 | 0.140783376 |
| | Cfh | 2.29 | 2.387056704 | 0.004101505473906130 | 0.131469154 |
| | Klc3 | 2.29 | 2.059771046 | 0.008714228700375000 | 0.140488765 |
| | AI467606 | 2.28 | 1.887456096 | 0.012958176898815000 | 0.146280491 |
| | Fcgr3 | 2.28 | 1.494388605 | 0.032034016523634600 | 0.187346622 |
| | Dbi | 2.28 | 2.092296576 | 0.008085435627812190 | 0.138221114 |
| | Lrrc9 | 2.28 | 2.11709499 | 0.007636687342159870 | 0.136287102 |
| | Cela1 | 2.28 | 2.40742604 | 0.003913577690689040 | 0.131162125 |
| | Clec3b | 2.28 | 1.384659375 | 0.041242086121260300 | 0.204948695 |
| | Wfikkn2 | 2.28 | 2.454253773 | 0.003513550715447710 | 0.129362173 |

| | | | | | |
|---|---|---|---|---|---|
| | Mcm6 | 2.27 | 3.096250656 | 0.000801215502760299 | 0.123542711 |
| | Kif6 | 2.27 | 1.330600507 | 0.046708884244469500 | 0.215957866 |
| | Ppic | 2.27 | 2.778971903 | 0.001663520269670600 | 0.127020757 |
| | Angpt2 | 2.26 | 2.04226083 | 0.009072754721165920 | 0.140488765 |
| | Ggta1 | 2.26 | 1.424408423 | 0.037634970208689100 | 0.199593499 |
| | Epb41l4a | 2.26 | 1.535537463 | 0.029138187740399900 | 0.180680432 |
| | Chek2 | 2.26 | 1.817965192 | 0.015206694032994600 | 0.151522906 |
| | Pygl | 2.26 | 2.221315532 | 0.006007371201838940 | 0.131469154 |
| | Nt5dc1 | 2.26 | 2.865941703 | 0.001361627446782820 | 0.127020757 |
| | BC055324 | 2.25 | 2.207735 | 0.006198191635533140 | 0.131469154 |
| | Cebpd | 2.25 | 1.404130076 | 0.039433917553379000 | 0.202168336 |
| | Arpc1b | 2.25 | 1.637632096 | 0.023033922656231700 | 0.166831290 |
| | Apod | 2.25 | 2.722342816 | 0.001895209321840240 | 0.127020757 |
| | Fabp7 | 2.25 | 1.533126243 | 0.029300414049566900 | 0.181277348 |
| | Myof | 2.25 | 1.943714864 | 0.011383744399364900 | 0.141961618 |
| | Ccdc122 | 2.24 | 1.696388411 | 0.020119240825966500 | 0.160920515 |
| | Tspan18 | 2.24 | 2.174438466 | 0.006692086320037540 | 0.133462929 |
| | Pcolce | 2.24 | 1.666447456 | 0.021555224166567000 | 0.163912748 |
| | Kl | 2.24 | 1.800652579 | 0.015825134915700300 | 0.152122332 |
| | Thbs2 | 2.24 | 3.28816434 | 0.000515033715648280 | 0.123542711 |
| | Mdfic | 2.24 | 1.614543734 | 0.024291608086224300 | 0.169664195 |
| | Ecscr | 2.23 | 2.29059917 | 0.005121543076203080 | 0.131469154 |
| | Aqp4 | 2.23 | 2.069211543 | 0.008526846738537930 | 0.140434588 |

| | | | | | |
|---|---|---|---|---|---|
| | P4ha3 | 2.23 | 2.264418497 | 0.005439782094179200 | 0.131469154 |
| | Ndnf | 2.23 | 1.522201267 | 0.030046835050228100 | 0.183250722 |
| | Cyth4 | 2.23 | 1.701968161 | 0.019862405255309000 | 0.160735761 |
| | Cpq | 2.23 | 2.289679541 | 0.005132399560876870 | 0.131469154 |
| | Fmod | 2.22 | 3.495848727 | 0.000319264972257602 | 0.123542711 |
| | S1pr2 | 2.22 | 1.387567154 | 0.040966875888124300 | 0.204032563 |
| | Lair1 | 2.21 | 2.250826002 | 0.005612728020322640 | 0.131469154 |
| | Id3 | 2.21 | 2.271771868 | 0.005348452364428480 | 0.131469154 |
| | Padi2 | 2.21 | 2.132811228 | 0.007365271691606810 | 0.136212918 |
| | Cdk2 | 2.21 | 2.428388779 | 0.003729161751207710 | 0.129856967 |
| | 1700094D03Rik | 2.21 | 1.957708856 | 0.011022780111375100 | 0.141933868 |
| | Bmp5 | 2.20 | 1.581553686 | 0.026208750345896500 | 0.174737719 |
| | Tsku | 2.20 | 1.603736987 | 0.024903650530995400 | 0.170773773 |
| | Catip | 2.20 | 1.819094549 | 0.015167201317104800 | 0.151522906 |
| | Casp7 | 2.20 | 1.455052012 | 0.035070986996816300 | 0.194379196 |
| | Hmgcs2 | 2.20 | 1.538997575 | 0.028906960257250600 | 0.180105205 |
| | Gldn | 2.20 | 1.611805671 | 0.024445241345887500 | 0.169826196 |
| | Slc22a4 | 2.20 | 2.017859575 | 0.009597108956792890 | 0.140783376 |
| | Slc2a10 | 2.19 | 1.60383594 | 0.024897976919826100 | 0.170773773 |
| | Prc1 | 2.19 | 1.638816853 | 0.022971171624440200 | 0.166831290 |
| | Snhg18 | 2.19 | 2.021127023 | 0.009525175299370510 | 0.140783376 |
| | Cnn2 | 2.19 | 1.828367393 | 0.014846791405819200 | 0.151437765 |
| | Esyt3 | 2.19 | 1.898272534 | 0.012639429326706800 | 0.145739007 |

| | | | | | |
|---|---|---|---|---|---|
| | Anxa4 | 2.19 | 1.392999158 | 0.040457667647326600 | 0.203278828 |
| | Steap3 | 2.18 | 2.068337978 | 0.008544015398287320 | 0.140434588 |
| | Ucp2 | 2.18 | 1.425302435 | 0.037557576886836700 | 0.199566382 |
| | Gjb2 | 2.18 | 1.966945933 | 0.010790810523099100 | 0.141501604 |
| | Rnaset2a | 2.17 | 2.435470164 | 0.003668848986766780 | 0.129856967 |
| | Itih2 | 2.17 | 1.676987707 | 0.021038379892842200 | 0.162610003 |
| | Psme2 | 2.17 | 1.503653114 | 0.031357893880266500 | 0.186051321 |
| | Cd38 | 2.16 | 1.489323121 | 0.032409839412162500 | 0.188385619 |
| | Itpripl2 | 2.16 | 1.921124825 | 0.011991545927416400 | 0.143647440 |
| | Tifab | 2.16 | 1.816090275 | 0.015272485640887200 | 0.151522906 |
| | Rab27a | 2.16 | 1.554636663 | 0.027884530517506800 | 0.178882497 |
| | 1700113A16Rik | 2.15 | 2.124638713 | 0.007505183019560490 | 0.136287102 |
| | Pik3cg | 2.15 | 2.082647555 | 0.008267085806720180 | 0.139726969 |
| | Loxl2 | 2.14 | 2.736373108 | 0.001834961228943460 | 0.127020757 |
| | Olfml2a | 2.14 | 1.661712794 | 0.021791504017333100 | 0.164435506 |
| | Rftn1 | 2.14 | 1.788882757 | 0.016259876520881700 | 0.152555604 |
| | Rab7b | 2.14 | 2.163293844 | 0.006866037254593220 | 0.134513024 |
| | Fanci | 2.14 | 2.233974747 | 0.005834790313460480 | 0.131469154 |
| | Hopx | 2.13 | 2.731871263 | 0.001854081141215580 | 0.127020757 |
| | Olfml3 | 2.13 | 2.043879594 | 0.009039000412255610 | 0.140488765 |
| | Sncaip | 2.13 | 2.275789795 | 0.005299198714652550 | 0.131469154 |
| | Gramd1c | 2.13 | 1.560240809 | 0.027527019508358800 | 0.178595240 |
| | Lgals1 | 2.13 | 2.715596118 | 0.001924880985134190 | 0.127020757 |

| | | | | | |
|---|---|---|---|---|---|
| | Msn | 2.13 | 2.074328154 | 0.008426977718708270 | 0.140434588 |
| | Plce1 | 2.12 | 1.500117247 | 0.031614240511339100 | 0.186755061 |
| | Lrriq1 | 2.12 | 1.624354533 | 0.023749007602114500 | 0.168509056 |
| | C1qtnf6 | 2.12 | 1.989618678 | 0.010241918642995300 | 0.141095284 |
| | Gcnt1 | 2.12 | 2.029977207 | 0.009333032814533400 | 0.140783376 |
| | Fam60a | 2.11 | 2.115946283 | 0.007656913079276390 | 0.136287102 |
| | Rab13 | 2.11 | 2.577143762 | 0.002647623566344280 | 0.127347630 |
| | Adamts9 | 2.11 | 2.201906603 | 0.006281934398559020 | 0.132111948 |
| | Tlr1 | 2.10 | 1.672846374 | 0.021239956625136500 | 0.162736895 |
| | Plscr4 | 2.10 | 1.443030158 | 0.036055360459641800 | 0.196819835 |
| | C1qtnf1 | 2.10 | 1.888997042 | 0.012912280681315100 | 0.146280491 |
| | Nckap1l | 2.10 | 1.575637147 | 0.026568244101765500 | 0.175939394 |
| | Trp53inp1 | 2.10 | 1.390844938 | 0.040658847267948100 | 0.203309852 |
| | Patj | 2.10 | 1.481231087 | 0.033019379893056000 | 0.190158969 |
| | Nkd2 | 2.10 | 2.049789867 | 0.008916822736996490 | 0.140488765 |
| | Pmf1 | 2.09 | 2.224415832 | 0.005964639051642170 | 0.131469154 |
| | Iqck | 2.09 | 3.013065209 | 0.000970364256884459 | 0.127020757 |
| | Eif4ebp1 | 2.09 | 2.796732175 | 0.001596863613868040 | 0.127020757 |
| | Ncaph | 2.09 | 1.504559352 | 0.031292527908239600 | 0.185821360 |
| | Clec14a | 2.09 | 2.151129013 | 0.007061077637108730 | 0.134660365 |
| | Fxyd1 | 2.09 | 1.482544951 | 0.032919637848029900 | 0.189849178 |
| | Rnf135 | 2.08 | 2.405759495 | 0.003928624359470170 | 0.131346052 |
| | Hmgb2 | 2.08 | 1.307385375 | 0.049273637546492900 | 0.221264397 |

| | | | | | |
|---|---|---|---|---|---|
| | Xlr | 2.08 | 1.327495784 | 0.047043997303900500 | 0.216606590 |
| | Rab29 | 2.08 | 2.643512519 | 0.002272414130166040 | 0.127020757 |
| | Zfp36 | 2.07 | 1.704040521 | 0.019767851933149500 | 0.160671218 |
| | Alox5ap | 2.07 | 2.101351997 | 0.007918592668169480 | 0.137039524 |
| | Smoc1 | 2.07 | 2.229818185 | 0.005890902227039970 | 0.131469154 |
| | Pi4k2b | 2.07 | 2.589576186 | 0.002572905372485080 | 0.127173715 |
| | Tgfb1 | 2.07 | 1.318659059 | 0.048011020909271300 | 0.218254040 |
| | Rab3il1 | 2.06 | 1.789473347 | 0.016237780014319300 | 0.152510824 |
| | Gm20743 | 2.06 | 1.394627517 | 0.040306258215997800 | 0.203107134 |
| | Foxc1 | 2.06 | 1.834579659 | 0.014635930598196700 | 0.151070715 |
| | Kif20a | 2.06 | 1.590291481 | 0.025686712156117800 | 0.173610331 |
| | B4galt1 | 2.06 | 1.522224625 | 0.030045219092699800 | 0.183250722 |
| | Ctsz | 2.05 | 1.546545091 | 0.028408932111284300 | 0.179474072 |
| | Fam181a | 2.05 | 1.54846267 | 0.028283772163479100 | 0.179474072 |
| | Elf1 | 2.05 | 1.52283176 | 0.030003245774088800 | 0.183233156 |
| | Cenpf | 2.05 | 1.51646093 | 0.030446618816692900 | 0.184263623 |
| | Slc14a1 | 2.05 | 1.666750721 | 0.021540177558262800 | 0.163889025 |
| | Col22a1 | 2.05 | 2.537978362 | 0.002897487946231260 | 0.128703904 |
| | Anpep | 2.05 | 2.033248547 | 0.009262995508130580 | 0.140488765 |
| | Cd55 | 2.04 | 2.607182363 | 0.002470686471757770 | 0.127020757 |
| | Slc13a3 | 2.04 | 2.006291893 | 0.009856168213198590 | 0.140783376 |
| | Zfp607a | 2.04 | 1.719411333 | 0.019080452382249000 | 0.160480708 |
| | Msx1 | 2.03 | 1.823421507 | 0.015016837905162500 | 0.151437765 |

| | | | | | |
|---|---|---|---|---|---|
| | Itpripl1 | 2.02 | 1.437199085 | 0.036542723759833000 | 0.197534842 |
| | Ccr5 | 2.02 | 1.713270807 | 0.019352148717746200 | 0.160482390 |
| | Ikzf2 | 2.02 | 1.684205542 | 0.020691618304937600 | 0.161859432 |
| | Cenpe | 2.02 | 1.984791819 | 0.010356384845428000 | 0.141095284 |
| | Kif18a | 2.01 | 1.309640722 | 0.049018416491267400 | 0.220509619 |
| | Col4a5 | 2.01 | 1.899144807 | 0.012614068738019900 | 0.145739007 |
| | Pard6g | 2.01 | 3.392292902 | 0.000405235140179762 | 0.123542711 |
| | Emp1 | 2.01 | 1.586335457 | 0.025921763469328400 | 0.174261718 |
| | Bmp7 | 2.01 | 1.657804751 | 0.021988482035621100 | 0.164780689 |
| | Ttc12 | 2.01 | 1.789932343 | 0.016220627730853700 | 0.152510824 |
| | Adcyap1 | 2.00 | 1.875648685 | 0.013315311015165800 | 0.146863953 |
| | Map3k19 | 2.00 | 1.814937772 | 0.015313068590749600 | 0.151597173 |
| | Chst14 | 2.00 | 1.665152942 | 0.021619570309648500 | 0.163994790 |
| | Creb5 | 2.00 | 1.887412891 | 0.012959466058326500 | 0.146280491 |
| | Scml4 | 2.00 | 2.266546124 | 0.005413197542506920 | 0.131469154 |
| | Card6 | 2.00 | 2.476496775 | 0.003338129836310830 | 0.128703904 |
| | Mfap4 | 2.00 | 1.888235012 | 0.012934956945832200 | 0.146280491 |
| | 1110017D15Rik | 1.99 | 1.512367925 | 0.030734919162436500 | 0.185102416 |
| | Spa17 | 1.98 | 1.817525113 | 0.015222111093370200 | 0.151522906 |
| | Sned1 | 1.98 | 2.009202767 | 0.009790327802511110 | 0.140783376 |
| | P2rx7 | 1.98 | 1.315530633 | 0.048358115346301200 | 0.218869849 |
| | Gal3st4 | 1.98 | 2.626904076 | 0.002360999657217900 | 0.127020757 |
| | Tmem176b | 1.97 | 1.927214025 | 0.011824586830026500 | 0.143647440 |

| | | | | | |
|---|---|---|---|---|---|
| | Vamp8 | 1.97 | 1.797285238 | 0.015948313427592100 | 0.152122332 |
| | Ccdc113 | 1.97 | 2.576661439 | 0.002650565621595000 | 0.127347630 |
| | Galm | 1.96 | 1.854031053 | 0.013994872527350300 | 0.148646776 |
| | Vwa5b1 | 1.96 | 1.553416213 | 0.027963001549299400 | 0.178882497 |
| | Cd33 | 1.96 | 2.018741623 | 0.009577637084651020 | 0.140783376 |
| | Tgfbr2 | 1.96 | 2.171700576 | 0.006734407991849130 | 0.133724711 |
| | Marcksl1 | 1.96 | 1.753457795 | 0.017641772009053000 | 0.157412720 |
| | Mcm2 | 1.96 | 2.115196987 | 0.007670135082522150 | 0.136345829 |
| | Ampd3 | 1.95 | 1.949585373 | 0.011230901716456100 | 0.141933868 |
| | Dna2 | 1.95 | 1.405106828 | 0.039345328173541100 | 0.202034437 |
| | Colec12 | 1.95 | 2.944246957 | 0.001136980569224200 | 0.127020757 |
| | Itpkb | 1.95 | 1.904237827 | 0.012467006118720500 | 0.145640913 |
| | Upp1 | 1.95 | 1.426597566 | 0.037445741409556600 | 0.199280377 |
| | Ifitm2 | 1.94 | 1.644995532 | 0.022646676038091500 | 0.166458646 |
| | Slc16a1 | 1.94 | 3.348281668 | 0.000448454444360335 | 0.123542711 |
| | Casp6 | 1.94 | 1.774624884 | 0.016802546900212700 | 0.154749105 |
| | C1ql1 | 1.94 | 1.313057957 | 0.048634229859590400 | 0.219541049 |
| | Zmynd10 | 1.94 | 1.603482788 | 0.024918231242225700 | 0.170773773 |
| | Slc37a1 | 1.93 | 1.43963415 | 0.036338404065489900 | 0.197393563 |
| | Csrp2 | 1.93 | 1.413642947 | 0.038579540713533500 | 0.200781458 |
| | Nid1 | 1.93 | 2.182713951 | 0.006565775796499470 | 0.133060702 |
| | Pros1 | 1.93 | 1.826375562 | 0.014915040551447500 | 0.151437765 |
| | Edem1 | 1.93 | 1.406008057 | 0.039263765122456500 | 0.201979414 |

| | | | | | |
|---|---|---|---|---|---|
| | Laptm5 | 1.93 | 1.30307647 | 0.049764945230214500 | 0.221444337 |
| | Fbn1 | 1.93 | 1.818953446 | 0.015172129956079400 | 0.151522906 |
| | Cenpj | 1.93 | 1.654354966 | 0.022163841423215100 | 0.165260680 |
| | Csf1 | 1.93 | 2.033548367 | 0.009256602896055850 | 0.140488765 |
| | Efs | 1.93 | 1.657198884 | 0.022019178693500500 | 0.164885850 |
| | Galnt12 | 1.93 | 1.980466132 | 0.010460052577513600 | 0.141095284 |
| | Vamp5 | 1.92 | 1.318474262 | 0.048031454502642000 | 0.218254040 |
| | Qpct | 1.92 | 2.455975972 | 0.003499645284905970 | 0.129362173 |
| | Hhex | 1.92 | 1.458075215 | 0.034827699189736900 | 0.194066267 |
| | Naprt | 1.91 | 1.37717446 | 0.041959039665337000 | 0.205913987 |
| | Slc26a2 | 1.91 | 1.653374788 | 0.022213920446285600 | 0.165260680 |
| | Dse | 1.91 | 1.763962959 | 0.017220154379292300 | 0.156474694 |
| | Dnase2a | 1.91 | 2.305686185 | 0.004946679980721590 | 0.131469154 |
| | Mr1 | 1.91 | 1.667118311 | 0.021521953521065800 | 0.163841088 |
| | Tmc6 | 1.91 | 1.71848399 | 0.019121238114719600 | 0.160480708 |
| | Apobr | 1.91 | 1.303124555 | 0.049759435514289600 | 0.221444337 |
| | Gadd45g | 1.91 | 2.558007428 | 0.002766894318015290 | 0.128703904 |
| | Fzd7 | 1.91 | 1.80014934 | 0.015843482934715500 | 0.152122332 |
| | Pax6 | 1.90 | 1.987030543 | 0.010303136575644000 | 0.141095284 |
| | Vwa5a | 1.90 | 2.053221738 | 0.008846638105299720 | 0.140488765 |
| | Apbb1ip | 1.90 | 1.937668123 | 0.011543350351859300 | 0.142769736 |
| | Col23a1 | 1.90 | 2.292454175 | 0.005099714049029510 | 0.131469154 |
| | Cpxm1 | 1.90 | 1.403594824 | 0.039482548406336200 | 0.202168336 |

| | | | | | |
|---|---|---|---|---|---|
| | Ddo | 1.89 | 1.556211283 | 0.027783612712679900 | 0.178882497 |
| | Tlr7 | 1.89 | 2.078681024 | 0.008342937223801420 | 0.140384356 |
| | Shroom3 | 1.89 | 1.553424935 | 0.027962439968574500 | 0.178882497 |
| | Angptl2 | 1.89 | 2.206659535 | 0.006213559547169810 | 0.131469154 |
| | Capn6 | 1.89 | 1.788111621 | 0.016288773298782100 | 0.152555604 |
| | Ednra | 1.89 | 2.013082904 | 0.009703247213767890 | 0.140783376 |
| | Adamts6 | 1.88 | 1.408825735 | 0.039009848637404600 | 0.201744197 |
| | Cyp4v3 | 1.88 | 1.698086703 | 0.020040718904436900 | 0.160920515 |
| | Ecm1 | 1.88 | 1.942312279 | 0.011420568439979300 | 0.142018128 |
| | Ptges | 1.88 | 1.643319739 | 0.022734230605557700 | 0.166786472 |
| | Cd63 | 1.88 | 2.09081017 | 0.008113156067046070 | 0.138316225 |
| | Lyl1 | 1.88 | 1.784994119 | 0.016406119875958700 | 0.153045820 |
| | Havcr2 | 1.88 | 1.329751993 | 0.046800232139281500 | 0.216162013 |
| | Tspan6 | 1.87 | 2.778978839 | 0.001663493703684440 | 0.127020757 |
| | Tmie | 1.87 | 1.676512387 | 0.021061418266851800 | 0.162610003 |
| | Rcsd1 | 1.87 | 1.409125662 | 0.038982917433662500 | 0.201744197 |
| | Cd9 | 1.87 | 2.233716295 | 0.005838263667233430 | 0.131469154 |
| | Plp2 | 1.87 | 2.267421877 | 0.005402292846906350 | 0.131469154 |
| | Col16a1 | 1.87 | 1.358019077 | 0.043851143521329600 | 0.210611848 |
| | Fam161a | 1.87 | 1.401034872 | 0.039715965759865400 | 0.202412377 |
| | Naaa | 1.86 | 2.018435201 | 0.009584397101316110 | 0.140783376 |
| | Loxl3 | 1.86 | 1.999009091 | 0.010022842577091200 | 0.141095284 |
| | Galnt4 | 1.86 | 2.397950027 | 0.003999907728817790 | 0.131469154 |

| | | | | | |
|---|---|---|---|---|---|
| | Zfp185 | 1.86 | 1.723831101 | 0.018887257412975900 | 0.160468942 |
| | Itgam | 1.86 | 1.607829474 | 0.024670078205483400 | 0.170091091 |
| | Entpd1 | 1.85 | 1.834423518 | 0.014641193548880500 | 0.151070715 |
| | Ranbp3l | 1.85 | 2.99842176 | 0.001003640643708230 | 0.127020757 |
| | Dcx | 1.85 | 1.602333067 | 0.024984285427086100 | 0.170961919 |
| | Rdh5 | 1.85 | 1.660006788 | 0.021877274285283100 | 0.164554124 |
| | Eif2ak2 | 1.85 | 1.734271444 | 0.018438626014876800 | 0.159751047 |
| | Igfbp5 | 1.85 | 1.571390782 | 0.026829292337253200 | 0.176398764 |
| | Relb | 1.85 | 1.38633919 | 0.041082873297461700 | 0.204388038 |
| | Dab2 | 1.84 | 1.872763849 | 0.013404053453062800 | 0.147466052 |
| | Aif1l | 1.84 | 1.761290975 | 0.017326427454774900 | 0.156718571 |
| | 5430405H02Rik | 1.84 | 1.710476621 | 0.019477058968412300 | 0.160482390 |
| | Katnal2 | 1.83 | 1.745153362 | 0.017982357950044900 | 0.158597136 |
| | Art3 | 1.83 | 1.490437139 | 0.032326810758209900 | 0.188347112 |
| | Plcxd3 | 1.83 | 1.759834749 | 0.017384621970646500 | 0.156718571 |
| | Efcab7 | 1.83 | 1.80106618 | 0.015810070994449300 | 0.152122332 |
| | Cyp26b1 | 1.83 | 1.958153456 | 0.011011501532610500 | 0.141933868 |
| | Scpep1 | 1.83 | 1.582589834 | 0.026146295532593200 | 0.174737719 |
| | Ecm2 | 1.82 | 2.267770793 | 0.005397954343218710 | 0.131469154 |
| | Tmem255a | 1.82 | 1.793351525 | 0.016093424786202500 | 0.152219810 |
| | Slc44a5 | 1.82 | 2.595677832 | 0.002537009935646040 | 0.127020757 |
| | Prelp | 1.82 | 1.69626052 | 0.020125166425184000 | 0.160920515 |
| | Smc4 | 1.82 | 1.999871079 | 0.010002968957460800 | 0.141095284 |

| | | | | | |
|---|---|---|---|---|---|
| | Tspan4 | 1.81 | 2.634256976 | 0.002321362818612460 | 0.127020757 |
| | Eva1c | 1.81 | 2.120946765 | 0.007569256722470410 | 0.136287102 |
| | Prrx1 | 1.81 | 2.172483367 | 0.006722280537348980 | 0.133677072 |
| | Notch2 | 1.81 | 2.273523396 | 0.005326925266537570 | 0.131469154 |
| | Bmf | 1.81 | 1.359837263 | 0.043667943223450000 | 0.210245693 |
| | Lhfpl2 | 1.81 | 3.187557745 | 0.000649295293845551 | 0.123542711 |
| | Ifih1 | 1.81 | 1.542313559 | 0.028687086370824300 | 0.179912941 |
| | Gem | 1.81 | 1.618433268 | 0.024075024168804700 | 0.169474850 |
| | Ptgds | 1.81 | 1.968554593 | 0.010750914471392600 | 0.141501604 |
| | E130114P18Rik | 1.81 | 2.741769992 | 0.001812299655068720 | 0.127020757 |
| | Htra3 | 1.81 | 1.779458087 | 0.016616590356742600 | 0.153711652 |
| | Slc25a24 | 1.81 | 1.720041219 | 0.019052798775448600 | 0.160480708 |
| | Sall1 | 1.81 | 2.207926374 | 0.006195460977209090 | 0.131469154 |
| | Ripk1 | 1.80 | 1.868604822 | 0.013533034167700100 | 0.147937488 |
| | Ptpn14 | 1.80 | 1.570852308 | 0.026862578181986300 | 0.176527349 |
| | Btg1 | 1.80 | 1.849685449 | 0.014135609876510500 | 0.149025511 |
| | Tifa | 1.80 | 2.653846007 | 0.002218983092571990 | 0.127020757 |
| | Limd1 | 1.80 | 2.276971626 | 0.005284797776248340 | 0.131469154 |
| | Jun | 1.80 | 3.679951094 | 0.000208953141972961 | 0.119849580 |
| | Tmem220 | 1.80 | 1.443240355 | 0.036037914011383600 | 0.196819835 |
| | Smoc2 | 1.80 | 1.439816057 | 0.036323186663251700 | 0.197393563 |
| | Syt9 | 1.80 | 1.542580143 | 0.028669482739105900 | 0.179884640 |
| | Tpk1 | 1.80 | 2.532321809 | 0.002935473681864620 | 0.128703904 |

| | | | | | |
|---|---|---|---|---|---|
| | Cmbl | 1.80 | 2.007829363 | 0.009821337545895950 | 0.140783376 |
| | Pard3b | 1.79 | 1.861251867 | 0.013764109938903700 | 0.147977908 |
| | Heyl | 1.79 | 1.774653215 | 0.016801450806324300 | 0.154749105 |
| | Cebpa | 1.79 | 1.893009488 | 0.012793533532900800 | 0.145881393 |
| | Arrdc4 | 1.79 | 1.702261454 | 0.019848996094891300 | 0.160735761 |
| | Fam107b | 1.79 | 1.901886825 | 0.012534677812408400 | 0.145640913 |
| | Hist1h2bq | 1.79 | 1.393862783 | 0.040377294680918500 | 0.203232383 |
| | Hist1h2br | 1.79 | 1.393862783 | 0.040377294680918500 | 0.203232383 |
| | Fn1 | 1.78 | 1.399904748 | 0.039819449537845700 | 0.202412377 |
| | Hmox1 | 1.78 | 1.560551829 | 0.027507313138223400 | 0.178595240 |
| | Agt | 1.78 | 1.302042735 | 0.049883539969118700 | 0.221613838 |
| | Mboat1 | 1.78 | 1.320202429 | 0.047840704961042900 | 0.218108536 |
| | Ednrb | 1.78 | 1.961923556 | 0.010916324679897200 | 0.141933868 |
| | Fibin | 1.78 | 1.768089915 | 0.017057292043767000 | 0.155633632 |
| | Etfbkmt | 1.78 | 1.694840061 | 0.020191098113413100 | 0.160920515 |
| | Smo | 1.77 | 2.221886593 | 0.005999477196696250 | 0.131469154 |
| | Ppp1r18 | 1.77 | 1.489075043 | 0.032428357887624600 | 0.188413558 |
| | Isoc1 | 1.77 | 1.5163616 | 0.030453583249448800 | 0.184263623 |
| | Akip1 | 1.77 | 1.642156228 | 0.022795219168671800 | 0.166831290 |
| | Ccdc3 | 1.77 | 1.363516524 | 0.043299559282175200 | 0.209205079 |
| | 2810459M11Rik | 1.77 | 1.378223607 | 0.041857799486918200 | 0.205721435 |
| | Pbld1 | 1.77 | 1.4924622 | 0.032176425765564500 | 0.187746849 |
| | Pms1 | 1.76 | 2.266650276 | 0.005411899513438450 | 0.131469154 |

| | | | | | |
|---|---|---|---|---|---|
| | Wdr89 | 1.76 | 1.580133498 | 0.026294595996113400 | 0.174878392 |
| | Nectin4 | 1.76 | 2.618699695 | 0.002406025938215880 | 0.127020757 |
| | Triobp | 1.76 | 1.679797415 | 0.020902709504263500 | 0.162468873 |
| | Rnf43 | 1.76 | 1.725677324 | 0.018807136478351400 | 0.160315671 |
| | Cmtm3 | 1.76 | 1.796859501 | 0.015963955141871700 | 0.152122332 |
| | Npas3 | 1.76 | 2.896635018 | 0.001268717650404800 | 0.127020757 |
| | Hexb | 1.75 | 2.012909675 | 0.009707118355229640 | 0.140783376 |
| | Slc8b1 | 1.75 | 1.841176768 | 0.014415284955144100 | 0.150441568 |
| | Agtrap | 1.75 | 2.502734907 | 0.003142426240883030 | 0.128703904 |
| | Tpcn2 | 1.75 | 1.713857621 | 0.019326017956965500 | 0.160482390 |
| | Cdon | 1.75 | 2.270303769 | 0.005366562984135260 | 0.131469154 |
| | Gusb | 1.75 | 1.74036074 | 0.018181899808508400 | 0.159035359 |
| | Tm4sf1 | 1.75 | 1.889031948 | 0.012911242921688200 | 0.146280491 |
| | Mdk | 1.74 | 1.497442741 | 0.031809530474795400 | 0.187032417 |
| | Sema3d | 1.74 | 1.64030582 | 0.022892550441681900 | 0.166831290 |
| | 1700096K18Rik | 1.74 | 2.198531359 | 0.006330946465834530 | 0.132111948 |
| | Fli1 | 1.74 | 1.349469606 | 0.044722944974736700 | 0.211647507 |
| | Rnase4 | 1.74 | 2.114392373 | 0.007684358665720050 | 0.136422187 |
| | Hells | 1.74 | 1.427366142 | 0.037379531894929400 | 0.199280377 |
| | Srebf1 | 1.73 | 2.247821233 | 0.005651695657340070 | 0.131469154 |
| | Wnt5b | 1.73 | 1.396481286 | 0.040134579199571500 | 0.202779537 |
| | Cx3cr1 | 1.73 | 1.789954257 | 0.016219809272364100 | 0.152510824 |
| | Wipf1 | 1.73 | 2.070297289 | 0.008505556057416240 | 0.140434588 |

| | | | | | |
|---|---|---|---|---|---|
| | Rnls | 1.73 | 1.70632173 | 0.019664290002894900 | 0.160646260 |
| | Crb2 | 1.73 | 1.564386583 | 0.027265496932374100 | 0.178068058 |
| | Nfe2l2 | 1.73 | 1.33897944 | 0.045816357626176600 | 0.213863298 |
| | Lama4 | 1.73 | 2.139233572 | 0.007257155492962200 | 0.135646175 |
| | Tmem119 | 1.72 | 1.757890116 | 0.017462639325999600 | 0.156808305 |
| | Rras | 1.72 | 2.287674849 | 0.005156145342869370 | 0.131469154 |
| | Cd59a | 1.72 | 1.516390894 | 0.030451529148550100 | 0.184263623 |
| | Inpp4b | 1.72 | 1.53372972 | 0.029259727672785500 | 0.181205864 |
| | Slc9a2 | 1.72 | 1.391590547 | 0.040589102950292200 | 0.203309852 |
| | Zfp808 | 1.72 | 1.635754424 | 0.023133725404241300 | 0.166831290 |
| | Mob3a | 1.72 | 1.472132457 | 0.033718445408804100 | 0.191535824 |
| | Erbb2 | 1.72 | 1.93970349 | 0.011489377783087400 | 0.142358467 |
| | Ldlrap1 | 1.72 | 1.413934628 | 0.038553638589415500 | 0.200781458 |
| | Hpgds | 1.72 | 2.249131441 | 0.005634670937872880 | 0.131469154 |
| | Ston1 | 1.72 | 1.506550906 | 0.031149357546650400 | 0.185693415 |
| | Trim56 | 1.71 | 1.853560319 | 0.014010049851701600 | 0.148646776 |
| | Slc7a11 | 1.71 | 1.787560149 | 0.016309470085670200 | 0.152555604 |
| | Bmp4 | 1.71 | 1.94326506 | 0.011395540784639100 | 0.141961618 |
| | Id4 | 1.70 | 1.506208999 | 0.031173890170544800 | 0.185759074 |
| | Nov | 1.70 | 1.828032853 | 0.014858232417684400 | 0.151437765 |
| | Cyr61 | 1.70 | 1.774039595 | 0.016825206573836300 | 0.154749105 |
| | 4930570G19Rik | 1.70 | 1.77319238 | 0.016858060943550300 | 0.154794652 |
| | Skap2 | 1.70 | 2.055683714 | 0.008796629200311120 | 0.140488765 |

| | | | | | |
|---|---|---|---|---|---|
| | Itgb5 | 1.70 | 1.917615352 | 0.012088840505203300 | 0.143647440 |
| | Dnah7a | 1.70 | 1.51941954 | 0.030239907621474800 | 0.183617574 |
| | Grn | 1.70 | 1.38298532 | 0.041401366899048000 | 0.205392575 |
| | Kif19a | 1.70 | 1.349346781 | 0.044735595082966200 | 0.211647507 |
| | Efcab10 | 1.70 | 1.628358857 | 0.023531041152297300 | 0.167968850 |
| | Ajuba | 1.70 | 2.132942009 | 0.007363054094350880 | 0.136212918 |
| | Mgp | 1.70 | 1.352314035 | 0.044430987441584900 | 0.211327864 |
| | Plekhg2 | 1.69 | 1.437314455 | 0.036533017500423600 | 0.197534842 |
| | Chaf1a | 1.69 | 2.207748838 | 0.006197994145029590 | 0.131469154 |
| | Glra2 | 1.69 | 1.345036474 | 0.045181799682208100 | 0.212544714 |
| | Nr1h3 | 1.69 | 1.317708542 | 0.048116215129428200 | 0.218422502 |
| | Lpar6 | 1.69 | 1.371381603 | 0.042522461565005000 | 0.207645692 |
| | Npl | 1.69 | 1.750256427 | 0.017772297436698300 | 0.157554283 |
| | Pcdhb9 | 1.69 | 1.456050085 | 0.034990481173777000 | 0.194290171 |
| | Bag3 | 1.69 | 1.546516787 | 0.028410783645119300 | 0.179474072 |
| | Itgb4 | 1.68 | 1.876353284 | 0.013293725790029600 | 0.146863953 |
| | Tfpi | 1.68 | 1.328743168 | 0.046909070976033600 | 0.216399537 |
| | 5930430L01Rik | 1.68 | 1.381329851 | 0.041559484221265500 | 0.205568349 |
| | Zfp558 | 1.68 | 1.820666322 | 0.015112408264007100 | 0.151522906 |
| | Csf1r | 1.67 | 1.795788698 | 0.016003364670109400 | 0.152122332 |
| | 1700016K19Rik | 1.67 | 1.540340511 | 0.028817711494402200 | 0.179952010 |
| | Tnfrsf11b | 1.67 | 1.369279268 | 0.042728803564305500 | 0.207844103 |
| | Zic2 | 1.67 | 2.228476097 | 0.005909134872021210 | 0.131469154 |

| | | | | | |
|---|---|---|---|---|---|
| | Phldb2 | 1.67 | 1.696223675 | 0.020126873863177900 | 0.160920515 |
| | Btd | 1.67 | 2.656563893 | 0.002205139693789530 | 0.127020757 |
| | Tmem100 | 1.67 | 1.509293024 | 0.030953301354018400 | 0.185473744 |
| | Cdk5rap2 | 1.67 | 1.343312243 | 0.045361536499266800 | 0.212880080 |
| | Maml2 | 1.67 | 1.659739869 | 0.021890724269121900 | 0.164554124 |
| | Fzd9 | 1.67 | 1.324560318 | 0.047363052284912800 | 0.217172036 |
| | Ifngr1 | 1.67 | 1.731837844 | 0.018542238199465000 | 0.159967493 |
| | Zfp953 | 1.67 | 1.735009709 | 0.018407308511703200 | 0.159580332 |
| | Emp2 | 1.66 | 2.073456653 | 0.008443905173155520 | 0.140434588 |
| | D16Ertd472e | 1.66 | 1.825522877 | 0.014944353205591200 | 0.151437765 |
| | Nckap5 | 1.66 | 1.416103505 | 0.038361580774460000 | 0.200478413 |
| | Stat3 | 1.66 | 1.557374445 | 0.027709300016750700 | 0.178673623 |
| | Plat | 1.66 | 2.068449416 | 0.008541823314437920 | 0.140434588 |
| | Herc6 | 1.66 | 1.494442571 | 0.032030036195698000 | 0.187346622 |
| | Ehhadh | 1.66 | 1.418310114 | 0.038167163537032000 | 0.200173662 |
| | Pdzrn4 | 1.65 | 1.571401312 | 0.026828641867249400 | 0.176398764 |
| | Sparc | 1.65 | 1.58237343 | 0.026159327197233700 | 0.174737719 |
| | Sqor | 1.65 | 1.809242079 | 0.015515219405544300 | 0.151749058 |
| | Zfp229 | 1.65 | 2.208575972 | 0.006186201015036250 | 0.131469154 |
| | Ptbp1 | 1.65 | 1.348268236 | 0.044846831441647100 | 0.211951088 |
| | Zfp758 | 1.65 | 2.306470702 | 0.004937752281831980 | 0.131469154 |
| | Il10rb | 1.65 | 1.730421143 | 0.018602823137947600 | 0.159967493 |
| | Zfp677 | 1.65 | 1.468154599 | 0.034028703388794000 | 0.192026086 |

| | | | | | |
|---|---|---|---|---|---|
| | Orai1 | 1.65 | 1.311148605 | 0.048848518335111500 | 0.220002455 |
| | Rhoj | 1.65 | 1.673604846 | 0.021202894561457000 | 0.162736895 |
| | Npc2 | 1.64 | 1.963393265 | 0.010879444858687000 | 0.141933868 |
| | S100a16 | 1.64 | 2.288445218 | 0.005147007276871520 | 0.131469154 |
| | Zfp36l1 | 1.64 | 1.492145272 | 0.032199915238561200 | 0.187800949 |
| | Crtap | 1.64 | 1.447431269 | 0.035691823047611500 | 0.196291813 |
| | Ezh2 | 1.64 | 1.365985678 | 0.043054080858847700 | 0.208705446 |
| | Pla2g16 | 1.64 | 2.642262871 | 0.002278962239930030 | 0.127020757 |
| | Edn3 | 1.64 | 2.671180263 | 0.002132159732234840 | 0.127020757 |
| | Gm13293 | 1.64 | 1.305710032 | 0.049464083604813400 | 0.221308908 |
| | P2rx6 | 1.64 | 2.420125443 | 0.003800795970242540 | 0.130257845 |
| | Ltbp1 | 1.64 | 2.255759158 | 0.005549333713423100 | 0.131469154 |
| | Fam126a | 1.64 | 1.438165585 | 0.036461490210046000 | 0.197424556 |
| | Arap1 | 1.63 | 1.918296896 | 0.012069884201954300 | 0.143647440 |
| | 1700029J07 Rik | 1.63 | 1.849994471 | 0.014125555271400500 | 0.149025511 |
| | Olfml1 | 1.63 | 2.066630204 | 0.008577679130816490 | 0.140483777 |
| | Lgi4 | 1.63 | 1.584223226 | 0.026048143387614000 | 0.174737719 |
| | Ldlrad3 | 1.63 | 2.294941941 | 0.005070584897441380 | 0.131469154 |
| | Fadd | 1.63 | 1.507063207 | 0.031112634952062500 | 0.185635570 |
| | Slc16a12 | 1.63 | 1.610335755 | 0.024528119008035500 | 0.169879477 |
| | Accs | 1.63 | 2.308011656 | 0.004920263304308780 | 0.131469154 |
| | Gpr39 | 1.63 | 1.398631732 | 0.039936340603754200 | 0.202412377 |
| | Slc35g1 | 1.63 | 1.652985121 | 0.022233860650046200 | 0.165260680 |

| | | | | | |
|---|---|---|---|---|---|
| | Mycl | 1.63 | 2.05714894 | 0.008767001081878240 | 0.140488765 |
| | 9430038I01Rik | 1.63 | 1.608513717 | 0.024631240394933400 | 0.170079334 |
| | Cald1 | 1.63 | 2.207673752 | 0.006199065830135430 | 0.131469154 |
| | Plod2 | 1.63 | 1.441269269 | 0.036201847171348800 | 0.197087790 |
| | Gas1 | 1.63 | 1.758255398 | 0.017447957779184600 | 0.156808305 |
| | Rp2 | 1.63 | 2.274926293 | 0.005309745516439500 | 0.131469154 |
| | Krt1 | 1.63 | 1.336133389 | 0.046117590720087700 | 0.214668636 |
| | Fancl | 1.62 | 2.192951912 | 0.006412805794295300 | 0.132629528 |
| | Man1a | 1.62 | 2.846935729 | 0.001422539291669620 | 0.127020757 |
| | Zfp92 | 1.62 | 1.330104263 | 0.046762286339074100 | 0.216132047 |
| | Pacrg | 1.62 | 2.049356492 | 0.008925725115798710 | 0.140488765 |
| | Nek6 | 1.62 | 2.01360507 | 0.009691587695302350 | 0.140783376 |
| | Nfatc3 | 1.62 | 2.624207368 | 0.002375705660019440 | 0.127020757 |
| | Fhod1 | 1.62 | 1.344572595 | 0.045230085114858000 | 0.212626274 |
| | Gatm | 1.62 | 2.254378984 | 0.005566997367524650 | 0.131469154 |
| | Selplg | 1.62 | 1.754036956 | 0.017618261189426700 | 0.157305086 |
| | Mettl25 | 1.62 | 2.848829447 | 0.001416349890651760 | 0.127020757 |
| | Dyx1c1 | 1.62 | 1.550608649 | 0.028144358290180600 | 0.179457832 |
| | Lmnb1 | 1.61 | 1.598725365 | 0.025192695371572200 | 0.171597570 |
| | Plpp2 | 1.61 | 1.431588484 | 0.037017877698585000 | 0.198531875 |
| | Ddah2 | 1.61 | 1.88224418 | 0.013114623283955600 | 0.146652067 |
| | Litaf | 1.61 | 1.603256885 | 0.024931196119803400 | 0.170774077 |
| | Kctd11 | 1.61 | 1.481695829 | 0.032984064516290200 | 0.190035233 |

| | | | | | |
|---|---|---|---|---|---|
| | Gng5 | 1.61 | 2.238733782 | 0.005771201237588970 | 0.131469154 |
| | Cep112 | 1.61 | 1.73338382 | 0.018476349985835000 | 0.159876275 |
| | Vsir | 1.61 | 1.636076845 | 0.023116557233107200 | 0.166831290 |
| | Slc16a2 | 1.61 | 2.281128937 | 0.005234450090840220 | 0.131469154 |
| | Noxo1 | 1.61 | 1.62945621 | 0.023471659157803000 | 0.167893841 |
| | Utp14b | 1.61 | 2.313044482 | 0.004863573883952750 | 0.131469154 |
| | Antxr1 | 1.61 | 1.87082569 | 0.013464006433666700 | 0.147515895 |
| | Lamp2 | 1.60 | 2.940804769 | 0.001146028006055690 | 0.127020757 |
| | St6gal1 | 1.60 | 1.602139018 | 0.024995451258784500 | 0.170961919 |
| | Parp4 | 1.60 | 1.902950576 | 0.012504013217649100 | 0.145640913 |
| | Gfpt2 | 1.60 | 1.598897739 | 0.025182698190358900 | 0.171597570 |
| | Tgfb2 | 1.60 | 2.529016498 | 0.002957900098415590 | 0.128703904 |
| | Tst | 1.60 | 1.508134267 | 0.031035999276970900 | 0.185581230 |
| | Ccdc74a | 1.60 | 2.540577541 | 0.002880198761202200 | 0.128703904 |
| | Wasf2 | 1.60 | 2.325569818 | 0.004725308659353950 | 0.131469154 |
| | Mcm4 | 1.60 | 2.277279817 | 0.005281048823423500 | 0.131469154 |
| | Hsd3b7 | 1.60 | 1.558418615 | 0.027642758819924000 | 0.178595240 |
| | Amz1 | 1.60 | 1.314376331 | 0.048486816333898700 | 0.219235717 |
| | Ccdc80 | 1.60 | 1.536694741 | 0.029060645639190800 | 0.180443891 |
| | Nfya | 1.60 | 1.559716659 | 0.027560261944180500 | 0.178595240 |
| | Ctdsp1 | 1.60 | 2.069949114 | 0.008512377708098460 | 0.140434588 |
| | Ddr1 | 1.59 | 2.502708861 | 0.003142614707975320 | 0.128703904 |
| | Scrn2 | 1.59 | 1.307471341 | 0.049263885170017700 | 0.221264397 |

| | | | | | |
|---|---|---|---|---|---|
| | Prdm16 | 1.59 | 2.037901177 | 0.009164289976010650 | 0.140488765 |
| | Copz2 | 1.59 | 1.341053005 | 0.045598126062966900 | 0.213460217 |
| | Hspb6 | 1.59 | 2.162541498 | 0.006877941880973280 | 0.134513024 |
| | Zfp521 | 1.59 | 1.554144974 | 0.027916118019378300 | 0.178882497 |
| | Lrp10 | 1.59 | 2.09164807 | 0.008097518148599070 | 0.138221114 |
| | Gm3414 | 1.59 | 1.633544513 | 0.023251741559401900 | 0.167340671 |
| | Ttc30b | 1.59 | 2.336295057 | 0.004610042649958020 | 0.131469154 |
| | Aspa | 1.59 | 1.713856347 | 0.019326074649864100 | 0.160482390 |
| | Pih1d2 | 1.59 | 1.87904744 | 0.013211513113729500 | 0.146825043 |
| | Celsr1 | 1.59 | 1.476959599 | 0.033345743159139700 | 0.191129135 |
| | Pgpep1 | 1.59 | 1.87607016 | 0.013302394998065600 | 0.146863953 |
| | P2ry13 | 1.59 | 2.042501759 | 0.009067722910961420 | 0.140488765 |
| | Bmp2k | 1.58 | 2.903303604 | 0.001249385313438080 | 0.127020757 |
| | Fbln5 | 1.58 | 1.715461712 | 0.019254767890894200 | 0.160482390 |
| | Ogfod3 | 1.58 | 2.237903576 | 0.005782244127395610 | 0.131469154 |
| | Tax1bp3 | 1.58 | 1.374564632 | 0.042211945462388900 | 0.206785862 |
| | Pola2 | 1.58 | 1.476709806 | 0.033364928153653800 | 0.191129135 |
| | Plxnb2 | 1.58 | 1.415717279 | 0.038395711609354600 | 0.200478413 |
| | Tfap4 | 1.58 | 1.378486169 | 0.041832501061077400 | 0.205721435 |
| | Pold4 | 1.58 | 1.740855926 | 0.018161180482620000 | 0.159035359 |
| | Rps6ka1 | 1.58 | 1.423634181 | 0.037702124103594800 | 0.199593499 |
| | Fcgrt | 1.58 | 1.830480971 | 0.014774712183702200 | 0.151437765 |
| | Tmem98 | 1.57 | 1.811030587 | 0.015451456115488000 | 0.151749058 |

| | | | | | |
|---|---|---|---|---|---|
| | Gsn | 1.57 | 2.157371995 | 0.006960300748156720 | 0.134660365 |
| | Psmg4 | 1.57 | 1.617555546 | 0.024123729694050300 | 0.169658425 |
| | Dusp16 | 1.57 | 2.893333833 | 0.001278398247479440 | 0.127020757 |
| | Fancf | 1.57 | 1.332308041 | 0.046525597528491400 | 0.215472948 |
| | Prdx4 | 1.57 | 1.970618669 | 0.010699939723222100 | 0.141501604 |
| | Col9a3 | 1.57 | 1.662822105 | 0.021735913391364200 | 0.164352442 |
| | Inhbb | 1.57 | 1.347240965 | 0.044953036684347100 | 0.211951088 |
| | Sgpl1 | 1.57 | 1.494843878 | 0.032000452655143300 | 0.187346622 |
| | Hspa2 | 1.56 | 1.612634501 | 0.024398633225649400 | 0.169826196 |
| | Rsu1 | 1.56 | 2.798569218 | 0.001590123227538530 | 0.127020757 |
| | Nbl1 | 1.56 | 1.896268147 | 0.012697898561177100 | 0.145765935 |
| | Txnrd3 | 1.56 | 1.737879251 | 0.018286085624547800 | 0.159232638 |
| | Abca1 | 1.56 | 1.709504495 | 0.019520705312340900 | 0.160482390 |
| | Pola1 | 1.56 | 1.98324903 | 0.010393240333879500 | 0.141095284 |
| | Clu | 1.56 | 1.8162009 | 0.015268595863394400 | 0.151522906 |
| | Gprc5c | 1.56 | 1.606196483 | 0.024763014789236600 | 0.170282757 |
| | Bbs10 | 1.56 | 1.968446008 | 0.010753602820717000 | 0.141501604 |
| | Mcm5 | 1.55 | 1.475815863 | 0.033433676579311500 | 0.191129135 |
| | Pth1r | 1.55 | 1.406001492 | 0.039264358693668700 | 0.201979414 |
| | Necap2 | 1.55 | 2.384979508 | 0.004121169638402680 | 0.131469154 |
| | Tmem159 | 1.55 | 2.655351308 | 0.002211305222137830 | 0.127020757 |
| | Tex9 | 1.55 | 1.975682779 | 0.010575897214136500 | 0.141402802 |
| | Rplp0 | 1.55 | 1.986371751 | 0.010318777519743700 | 0.141095284 |

| | | | | | |
|---|---|---|---|---|---|
| | Lama1 | 1.55 | 1.349531616 | 0.044716559761687600 | 0.211647507 |
| | Rfx4 | 1.55 | 2.581614654 | 0.002620507133304370 | 0.127347630 |
| | Syt17 | 1.55 | 1.429928683 | 0.037159624525620700 | 0.198835826 |
| | Cst3 | 1.55 | 1.930780962 | 0.011727867156792500 | 0.143647440 |
| | Lap3 | 1.55 | 2.27863773 | 0.005264562305543110 | 0.131469154 |
| | Cxadr | 1.55 | 1.517482126 | 0.030375111040319400 | 0.184100966 |
| | Slc22a8 | 1.55 | 1.571587537 | 0.026817140240672800 | 0.176398764 |
| | Fstl1 | 1.55 | 1.87612918 | 0.013300587353142600 | 0.146863953 |
| | Ctso | 1.55 | 2.030506412 | 0.009321667078711470 | 0.140783376 |
| | Hspg2 | 1.55 | 1.720408233 | 0.019036704437295500 | 0.160480708 |
| | Chd7 | 1.54 | 1.36238447 | 0.043412573315571400 | 0.209530091 |
| | Scamp2 | 1.54 | 1.632395141 | 0.023313359398400300 | 0.167460405 |
| | Kank2 | 1.54 | 2.011038435 | 0.009749033558498530 | 0.140783376 |
| | Zfp40 | 1.54 | 2.030697026 | 0.009317576639625380 | 0.140783376 |
| | Casp3 | 1.54 | 2.215751523 | 0.006084830391827260 | 0.131469154 |
| | Stt3a | 1.54 | 2.075948113 | 0.008395602856785540 | 0.140384356 |
| | Tppp3 | 1.54 | 1.431623057 | 0.037014930904617500 | 0.198531875 |
| | Ninj1 | 1.54 | 1.50060507 | 0.031578749621696600 | 0.186633806 |
| | Lima1 | 1.54 | 1.387971481 | 0.040928753609374300 | 0.204032563 |
| | Sec61b | 1.54 | 1.710184103 | 0.019490182100664300 | 0.160482390 |
| | Zc2hc1c | 1.54 | 1.46171636 | 0.034536922845889900 | 0.192993842 |
| | Hist1h2bc | 1.54 | 1.576486083 | 0.026516360635629500 | 0.175765225 |
| | Cdh4 | 1.54 | 1.718966891 | 0.019099988623955700 | 0.160480708 |

| | | | | | |
|---|---|---|---|---|---|
| | Zfp277 | 1.54 | 5.150272188 | 0.000007075022284362 | 0.048608941 |
| | Xkr8 | 1.54 | 1.879323012 | 0.013203132707340600 | 0.146825043 |
| | Abhd4 | 1.54 | 1.993747383 | 0.010145013219625100 | 0.141095284 |
| | Foxo1 | 1.54 | 1.439564833 | 0.036344204490638900 | 0.197393563 |
| | Scara3 | 1.54 | 2.264791566 | 0.005435111198188620 | 0.131469154 |
| | Bak1 | 1.54 | 1.704642053 | 0.019740490843608500 | 0.160671218 |
| | Tspan15 | 1.54 | 1.647963386 | 0.022492442263977000 | 0.166458646 |
| | Cd151 | 1.53 | 1.9677585 | 0.010770639740509900 | 0.141501604 |
| | Cecr2 | 1.53 | 2.105251646 | 0.007847807730699280 | 0.137039524 |
| | Naga | 1.53 | 1.733847956 | 0.018456614599635300 | 0.159806138 |
| | Fmo5 | 1.53 | 1.521441086 | 0.030099474507228600 | 0.183250722 |
| | 43353 | 1.53 | 1.508611139 | 0.031001939274842400 | 0.185581230 |
| | Sat1 | 1.53 | 2.658573911 | 0.002194957364781860 | 0.127020757 |
| | Ccdc13 | 1.53 | 1.796037737 | 0.015994190431880800 | 0.152122332 |
| | Cdc42bpg | 1.53 | 1.476947207 | 0.033346694659006900 | 0.191129135 |
| | Nynrin | 1.53 | 2.37991505 | 0.004169509327723660 | 0.131469154 |
| | Brca1 | 1.53 | 1.795628617 | 0.016009264628154400 | 0.152122332 |
| | Lum | 1.53 | 1.621868783 | 0.023885328410074300 | 0.168887506 |
| | Cavin3 | 1.53 | 2.417588948 | 0.003823059452541790 | 0.130257845 |
| | Cmtm6 | 1.53 | 2.196874783 | 0.006355141383258300 | 0.132111948 |
| | Nsmce1 | 1.53 | 2.008456919 | 0.009807155937173810 | 0.140783376 |
| | Tgfbr1 | 1.53 | 2.426087742 | 0.003748972527705340 | 0.129856967 |
| | Fam35a | 1.52 | 2.254614983 | 0.005563973039565230 | 0.131469154 |

| | | | | | |
|---|---|---|---|---|---|
| | Rps27l | 1.52 | 1.410727369 | 0.038839410584997100 | 0.201396247 |
| | Rcbtb2 | 1.52 | 1.663969628 | 0.021678557075522900 | 0.164132783 |
| | Slc39a1 | 1.52 | 2.509097036 | 0.003096727308332010 | 0.128703904 |
| | Lrp5 | 1.52 | 1.739795924 | 0.018205561403351200 | 0.159035359 |
| | Zkscan7 | 1.52 | 1.34102682 | 0.045600875415307400 | 0.213460217 |
| | Nedd1 | 1.52 | 1.91709013 | 0.012103469210308200 | 0.143647440 |
| | Calcrl | 1.52 | 1.356321721 | 0.044022862583212700 | 0.210699462 |
| | Frmd4b | 1.52 | 1.936819084 | 0.011565939481990700 | 0.142792070 |
| | Zfp992 | 1.52 | 1.673337118 | 0.021215969489884200 | 0.162736895 |
| | Bid | 1.51 | 1.606533816 | 0.024743787906812300 | 0.170282757 |
| | H3f3b | 1.51 | 2.971693279 | 0.001067349672412350 | 0.127020757 |
| | 1810058I24Rik | 1.51 | 4.279630347 | 0.000052525434479666 | 0.103107428 |
| | Ggh | 1.51 | 2.347265776 | 0.004495046863239180 | 0.131469154 |
| | Prrg1 | 1.51 | 1.351800532 | 0.044483553000682300 | 0.211431512 |
| | Spata24 | 1.51 | 1.335262038 | 0.046210212154891800 | 0.214941372 |
| | Pcdhb7 | 1.51 | 2.206285662 | 0.006218910955111730 | 0.131469154 |
| | Mlc1 | 1.51 | 1.728258809 | 0.018695676779631500 | 0.159967493 |
| | Rab40b | 1.51 | 2.375449571 | 0.004212601993424040 | 0.131469154 |
| | Homer3 | 1.51 | 1.318544183 | 0.048023722053167100 | 0.218254040 |
| | Haus8 | 1.51 | 2.087368205 | 0.008177711679758180 | 0.138564959 |
| | Hexa | 1.51 | 1.810968839 | 0.015453653180700900 | 0.151749058 |
| | Zfp41 | 1.51 | 1.794440811 | 0.016053110290629500 | 0.152122332 |
| | Heatr5a | 1.51 | 1.646047694 | 0.022591876549216100 | 0.166458646 |

| | | | | | |
|---|---|---|---|---|---|
| | Nupr1 | 1.51 | 1.7175911 14 | 0.0191605903590 64700 | 0.1604823 90 |
| | Ankrd44 | 1.51 | 1.5833746 58 | 0.0260990886242 30500 | 0.1747377 19 |
| | Zfp36l2 | 1.51 | 1.6902305 | 0.0204065458685 73400 | 0.1614833 32 |
| | Lfng | 1.51 | 1.6035635 47 | 0.0249135980476 22500 | 0.1707737 73 |
| | Zfp458 | 1.50 | 2.1979224 46 | 0.0063398291464 08930 | 0.1321119 48 |
| | Tnfaip8 | 1.50 | 1.5013602 45 | 0.0315238864586 02500 | 0.1865502 69 |
| | Ssr4 | 1.50 | 2.1084272 91 | 0.0077906323420 02740 | 0.1370395 24 |
| | Adgrg1 | 1.50 | 1.6279949 13 | 0.0235507687082 93100 | 0.1679799 16 |
| | Tmem198b | 1.50 | 2.0491500 54 | 0.0089299688957 92930 | 0.1404887 65 |
| | Mplkip | 1.50 | 1.9492316 69 | 0.0112400522825 93100 | 0.1419338 68 |
| | Fhl1 | 1.50 | 2.4991816 27 | 0.0031682421905 34590 | 0.1287039 04 |
| | Fut10 | 1.50 | 1.7185240 76 | 0.0191194733084 86000 | 0.1604807 08 |
| | Xrcc2 | 1.50 | 2.0674092 57 | 0.0085623059738 86430 | 0.1404728 27 |
| | B3gnt9 | 1.50 | 1.6597333 78 | 0.0218910514665 20000 | 0.1645541 24 |
| | Dock10 | 1.50 | 1.4099897 74 | 0.0389054305710 39000 | 0.2015091 96 |
| | Adam17 | 1.50 | 1.8590206 3 | 0.0138350065650 74400 | 0.1480582 75 |
| | Col6a2 | 1.50 | 1.7053210 48 | 0.0197096518147 47700 | 0.1606712 18 |
| | Pbxip1 | 1.50 | 1.8989381 1 | 0.0126200736784 07800 | 0.1457390 07 |
| | Sema6a | 1.50 | 2.2985713 95 | 0.0050283859590 90140 | 0.1314691 54 |
| | Twsg1 | 1.50 | 2.1096528 64 | 0.0077686782615 40680 | 0.1370339 00 |
| | Oat | 1.49 | 2.8024166 98 | 0.0015760983040 32710 | 0.1270207 57 |
| | Fam69c | 1.49 | 1.6634665 35 | 0.0217036843556 34400 | 0.1642237 48 |
| | Ifnar2 | 1.49 | 1.4481883 32 | 0.0356296592039 90900 | 0.1962273 13 |

| | | | | | |
|---|---|---|---|---|---|
| | Il6st | 1.49 | 2.183394258 | 0.006555498793371940 | 0.133060702 |
| | Lrrc51 | 1.49 | 1.422230339 | 0.037824192126434300 | 0.199763957 |
| | Adra2a | 1.49 | 1.592302558 | 0.025568040294791400 | 0.173183585 |
| | Cpne3 | 1.49 | 1.456722855 | 0.034936319066635200 | 0.194290171 |
| | Vmac | 1.49 | 2.274036307 | 0.005320637765646050 | 0.131469154 |
| | Atp6v0e | 1.49 | 1.674418439 | 0.021163210961161600 | 0.162736895 |
| | Dnmbp | 1.48 | 1.803217112 | 0.015731961971399400 | 0.152122332 |
| | Pde3b | 1.48 | 1.502948286 | 0.031408826741091200 | 0.186151134 |
| | Fancd2 | 1.48 | 1.447221401 | 0.035709074888006900 | 0.196291813 |
| | Sptssa | 1.48 | 2.06070484 | 0.008695512016522540 | 0.140488765 |
| | D130020L05Rik | 1.48 | 1.614931084 | 0.024269951931652300 | 0.169664195 |
| | Ispd | 1.48 | 2.49247802 | 0.003217525370497600 | 0.128703904 |
| | Cpne2 | 1.48 | 1.916962685 | 0.012107021540418800 | 0.143647440 |
| | Vangl2 | 1.48 | 2.254428555 | 0.005566361983608580 | 0.131469154 |
| | Gab1 | 1.48 | 1.353286293 | 0.044331630731609000 | 0.211220852 |
| | Fgfr1 | 1.48 | 1.616153236 | 0.024201749656575600 | 0.169664195 |
| | Cep192 | 1.48 | 1.64755526 | 0.022513589317950800 | 0.166458646 |
| | Txndc5 | 1.48 | 1.630461089 | 0.023417412782603500 | 0.167794957 |
| | Pskh1 | 1.48 | 2.453339435 | 0.003520955732538860 | 0.129362173 |
| | Sox2 | 1.47 | 1.977781975 | 0.010524901128478500 | 0.141095284 |
| | Uaca | 1.47 | 1.412482325 | 0.038682779767695000 | 0.200960331 |
| | Prdx1 | 1.47 | 3.080910512 | 0.000830021778728244 | 0.123542711 |
| | Wfs1 | 1.47 | 1.852891297 | 0.014031648696051100 | 0.148646776 |

| | | | | | |
|---|---|---|---|---|---|
| | A730017C20 Rik | 1.47 | 2.34307965 | 0.004538583712547440 | 0.131469154 |
| | Fkbp14 | 1.47 | 1.433077089 | 0.036891210976610900 | 0.198531875 |
| | Snhg1 | 1.47 | 1.702709529 | 0.019828527813240100 | 0.160735761 |
| | Plxdc2 | 1.47 | 2.61809387 | 0.002409384597883810 | 0.127020757 |
| | Frmd8 | 1.47 | 1.400903728 | 0.039727960613773200 | 0.202412377 |
| | Coa4 | 1.47 | 1.906217108 | 0.012410317488271100 | 0.145323755 |
| | Svbp | 1.47 | 1.800332639 | 0.015836797402671400 | 0.152122332 |
| | Cyp20a1 | 1.47 | 2.796644683 | 0.001597185345043120 | 0.127020757 |
| | Tmcc3 | 1.47 | 1.765850282 | 0.017145482752276000 | 0.156127289 |
| | Grem2 | 1.47 | 3.146721957 | 0.000713309556886860 | 0.123542711 |
| | Rhog | 1.47 | 1.852253693 | 0.014052264213763600 | 0.148646776 |
| | Gja1 | 1.47 | 1.892024017 | 0.012822596698543800 | 0.145881393 |
| | Leprot | 1.47 | 2.249365681 | 0.005631632655672410 | 0.131469154 |
| | Yap1 | 1.47 | 2.440614825 | 0.003625644139049720 | 0.129856967 |
| | Slc29a3 | 1.47 | 1.505097074 | 0.031253807004166800 | 0.185821360 |
| | Gnai2 | 1.47 | 1.888064505 | 0.012940036298069200 | 0.146280491 |
| | Nkain2 | 1.47 | 1.639968915 | 0.022910316306732300 | 0.166831290 |
| | Smim4 | 1.46 | 1.946325361 | 0.011315523178130500 | 0.141961618 |
| | Chadl | 1.46 | 1.559299146 | 0.027586770007720400 | 0.178595240 |
| | Tln1 | 1.46 | 1.615913961 | 0.024215087268502100 | 0.169664195 |
| | Lpcat3 | 1.46 | 1.640812814 | 0.022865841349062500 | 0.166831290 |
| | Ccdc190 | 1.46 | 1.849107258 | 0.014154441624120200 | 0.149025511 |
| | B230354K17 Rik | 1.46 | 2.038968191 | 0.009141801964011540 | 0.140488765 |

| | | | | | |
|---|---|---|---|---|---|
| | Gpc4 | 1.46 | 1.446577381 | 0.035762067534769300 | 0.196302663 |
| | Gng11 | 1.46 | 1.456080257 | 0.034988050370909600 | 0.194290171 |
| | BC028528 | 1.46 | 1.391006294 | 0.040643743907834900 | 0.203309852 |
| | Fzd6 | 1.46 | 1.937106729 | 0.011558281590925200 | 0.142792070 |
| | Stard13 | 1.46 | 2.352846992 | 0.004437649609449330 | 0.131469154 |
| | Gstcd | 1.46 | 1.340070038 | 0.045701448134167600 | 0.213599864 |
| | Cdo1 | 1.46 | 1.922210336 | 0.011961610699755200 | 0.143647440 |
| | Tmem47 | 1.46 | 2.076815672 | 0.008378848313498900 | 0.140384356 |
| | Fxyd6 | 1.46 | 1.473007024 | 0.033650612654314700 | 0.191453200 |
| | Xrcc4 | 1.46 | 1.986509331 | 0.010315509160038400 | 0.141095284 |
| | Add3 | 1.46 | 1.495185623 | 0.031975281529392100 | 0.187285739 |
| | Rimbp3 | 1.46 | 1.3013134 | 0.049967382547677800 | 0.221634296 |
| | 9130023H24Rik | 1.46 | 1.513983327 | 0.030620809911804200 | 0.184949692 |
| | Cyp7b1 | 1.46 | 2.461479118 | 0.003455579446541030 | 0.129362173 |
| | Chrac1 | 1.45 | 2.528599524 | 0.002960741396724300 | 0.128703904 |
| | Abca9 | 1.45 | 1.317898066 | 0.048095222068559200 | 0.218399354 |
| | Egflam | 1.45 | 1.53830108 | 0.028953356630060300 | 0.180151698 |
| | Pign | 1.45 | 1.739927199 | 0.018200059214360200 | 0.159035359 |
| | Marcks | 1.45 | 1.917976322 | 0.012078796863055300 | 0.143647440 |
| | Lamb2 | 1.45 | 1.555092982 | 0.027855247291313200 | 0.178882497 |
| | Nfkbia | 1.45 | 1.390843187 | 0.040659011209746500 | 0.203309852 |
| | AU041133 | 1.45 | 1.924962557 | 0.011886046998099100 | 0.143647440 |
| | Rela | 1.45 | 1.954027012 | 0.011116625813209300 | 0.141933868 |

| | | | | | |
|---|---|---|---|---|---|
| | Cav1 | 1.45 | 2.459448446 | 0.003471774860081510 | 0.129362173 |
| | Snap23 | 1.45 | 1.423655979 | 0.037700231870555900 | 0.199593499 |
| | Aldh4a1 | 1.45 | 2.112430926 | 0.007719142783638950 | 0.136821077 |
| | Siglech | 1.45 | 1.658615004 | 0.021947496895008100 | 0.164777465 |
| | Sh2b2 | 1.45 | 1.410721093 | 0.038839971905922800 | 0.201396247 |
| | Zfp449 | 1.45 | 1.589451746 | 0.025736427013013300 | 0.173610331 |
| | Lbh | 1.45 | 1.450593219 | 0.035432906725136000 | 0.195808417 |
| | Apoe | 1.45 | 1.50308041 | 0.031399272801841600 | 0.186151134 |
| | Mospd2 | 1.45 | 2.006373327 | 0.009854320266665870 | 0.140783376 |
| | Fem1c | 1.45 | 2.349921701 | 0.004467641321380190 | 0.131469154 |
| | Igfbp7 | 1.45 | 1.366687733 | 0.042984538385372900 | 0.208636716 |
| | Dsel | 1.45 | 1.769773563 | 0.016991293296955500 | 0.155547876 |
| | Brip1os | 1.45 | 1.855635432 | 0.013943267765564400 | 0.148627530 |
| | Arhgef26 | 1.45 | 1.541869003 | 0.028716466283106000 | 0.179932351 |
| | Laptm4a | 1.45 | 2.297942143 | 0.005035676902888980 | 0.131469154 |
| | Siva1 | 1.44 | 1.321221797 | 0.047728545902950900 | 0.217912197 |
| | Poc1b | 1.44 | 1.352799636 | 0.044381335219584500 | 0.211311132 |
| | Dhrs1 | 1.44 | 2.507586652 | 0.003107515816712600 | 0.128703904 |
| | Fzd8 | 1.44 | 1.548198408 | 0.028300987646373900 | 0.179474072 |
| | C330027C09Rik | 1.44 | 1.532416034 | 0.029348368637649900 | 0.181410676 |
| | Bmpr1b | 1.44 | 1.305210445 | 0.049521017037377100 | 0.221308908 |
| | Mipol1 | 1.44 | 1.312095612 | 0.048742116971050200 | 0.219795160 |
| | 2610524H06Rik | 1.44 | 2.308851197 | 0.004910761050935960 | 0.131469154 |

| | | | | | |
|---|---|---|---|---|---|
| | Slc39a11 | 1.44 | 1.754541842 | 0.017597791116958000 | 0.157224478 |
| | Gm6644 | 1.44 | 1.428643892 | 0.037269718122986600 | 0.198960061 |
| | Cttnbp2nl | 1.44 | 1.57290999 | 0.026735604591631800 | 0.176398764 |
| | St5 | 1.44 | 1.982465273 | 0.010412013605241800 | 0.141095284 |
| | Rnf180 | 1.44 | 1.445651068 | 0.035838426380056100 | 0.196510701 |
| | Ormdl1 | 1.43 | 1.969255119 | 0.010733587009599800 | 0.141501604 |
| | Cradd | 1.43 | 1.558100712 | 0.027663000690584600 | 0.178595240 |
| | Peli2 | 1.43 | 1.394362812 | 0.040330832666270100 | 0.203147350 |
| | Ptpn13 | 1.43 | 1.507543499 | 0.031078246081425800 | 0.185593785 |
| | Zfp580 | 1.43 | 1.679775478 | 0.020903765347669600 | 0.162468873 |
| | Tmem86a | 1.43 | 1.528229629 | 0.029632641811180600 | 0.182265949 |
| | Ticam1 | 1.43 | 1.444395775 | 0.035942164317979100 | 0.196696198 |
| | Riox2 | 1.43 | 1.671689223 | 0.021296624656752100 | 0.163028924 |
| | Man2b2 | 1.43 | 1.488411528 | 0.032477939758512700 | 0.188595618 |
| | Prkcq | 1.43 | 1.695653789 | 0.020153301918009200 | 0.160920515 |
| | Zfp110 | 1.43 | 1.821425867 | 0.015086001045690500 | 0.151522906 |
| | Wdr31 | 1.43 | 1.30839808 | 0.049158873207531100 | 0.220965678 |
| | Sec22a | 1.43 | 2.880052248 | 0.001318098155835820 | 0.127020757 |
| | Cnpy2 | 1.43 | 2.369355891 | 0.004272126547017900 | 0.131469154 |
| | Ttc30a1 | 1.43 | 2.198308844 | 0.006334191022214240 | 0.132111948 |
| | Zeb2os | 1.43 | 2.121385115 | 0.007561620633841150 | 0.136287102 |
| | Plk4 | 1.43 | 1.829799679 | 0.014797907921465200 | 0.151437765 |
| | Cpne8 | 1.43 | 2.562450216 | 0.002738733564402430 | 0.128703904 |

| | | | | | |
|---|---|---|---|---|---|
| | Psph | 1.43 | 2.095341066 | 0.008028953344709470 | 0.137641801 |
| | Cdca4 | 1.42 | 1.746049885 | 0.017945274859799800 | 0.158474307 |
| | Tcf3 | 1.42 | 1.654168786 | 0.022173344990723600 | 0.165260680 |
| | Ginm1 | 1.42 | 1.76352145 | 0.017237669498649900 | 0.156474694 |
| | Arl13b | 1.42 | 2.331477734 | 0.004661463265938560 | 0.131469154 |
| | Ncapg2 | 1.42 | 1.789518967 | 0.016236074412997500 | 0.152510824 |
| | Cdc42ep4 | 1.42 | 1.767493209 | 0.017080744284686100 | 0.155744199 |
| | Ctbs | 1.42 | 1.699891359 | 0.019957615033073400 | 0.160872834 |
| | Zfp429 | 1.42 | 1.351043454 | 0.044561165972943100 | 0.211585183 |
| | Krcc1 | 1.42 | 1.66615 | 0.021569992801824200 | 0.163934331 |
| | Fkbp9 | 1.42 | 2.060718352 | 0.008695241484680190 | 0.140488765 |
| | Gpc6 | 1.42 | 3.254034078 | 0.000557142029794589 | 0.123542711 |
| | Sowahc | 1.42 | 1.611129182 | 0.024483348718427900 | 0.169826196 |
| | Rpl39 | 1.42 | 2.340924084 | 0.004561166400724800 | 0.131469154 |
| | Plau | 1.42 | 1.325303819 | 0.047282037296556400 | 0.217136270 |
| | Pld2 | 1.42 | 2.545513538 | 0.002847649033126760 | 0.128703904 |
| | Rhbdd1 | 1.42 | 1.746981098 | 0.017906837896998800 | 0.158236566 |
| | Tmem150a | 1.41 | 1.324527694 | 0.047366610228671200 | 0.217172036 |
| | Plcb3 | 1.41 | 1.340797446 | 0.045624965989457100 | 0.213460217 |
| | Susd2 | 1.41 | 1.38822474 | 0.040904892908114600 | 0.204032563 |
| | Rps27rt | 1.41 | 2.480024807 | 0.003311122080394060 | 0.128703904 |
| | Htr2a | 1.41 | 1.907692224 | 0.012368236360358400 | 0.145185458 |
| | Zfp105 | 1.41 | 2.022660835 | 0.009491594252565190 | 0.140783376 |

| | | | | | |
|---|---|---|---|---|---|
| | Gpx7 | 1.41 | 1.501114675 | 0.031541716580233500 | 0.186575432 |
| | Ccdc173 | 1.41 | 1.325842893 | 0.047223384216248600 | 0.217063784 |
| | 1810037I17Rik | 1.41 | 3.494848421 | 0.000320001179735828 | 0.123542711 |
| | Vtn | 1.41 | 1.994358923 | 0.010130737841096500 | 0.141095284 |
| | Blvrb | 1.41 | 1.526882643 | 0.029724691543850900 | 0.182539181 |
| | Erbin | 1.41 | 1.837106215 | 0.014551031631155200 | 0.150675000 |
| | Smarca1 | 1.41 | 2.492763496 | 0.003215411077309840 | 0.128703904 |
| | Serp1 | 1.41 | 1.629835872 | 0.023451149087094500 | 0.167834500 |
| | Ntsr2 | 1.41 | 1.817759614 | 0.015213894003233900 | 0.151522906 |
| | Dtwd2 | 1.41 | 1.406772377 | 0.039194725136259600 | 0.201979414 |
| | Gpr34 | 1.41 | 2.143385921 | 0.007188099481752310 | 0.135303664 |
| | Rps20 | 1.41 | 2.319073932 | 0.004796517879085030 | 0.131469154 |
| | P2ry12 | 1.41 | 2.2384261 | 0.005775291367138380 | 0.131469154 |
| | Bmpr1a | 1.40 | 2.046786337 | 0.008978704173300360 | 0.140488765 |
| | Wnt7b | 1.40 | 1.724295363 | 0.018867077649350600 | 0.160468942 |
| | Prkd3 | 1.40 | 1.759343344 | 0.017404303833717700 | 0.156718571 |
| | Lzic | 1.40 | 1.726271305 | 0.018781431685555000 | 0.160215585 |
| | Prdx6 | 1.40 | 1.393009122 | 0.040456739363935400 | 0.203278828 |
| | Nit2 | 1.40 | 1.990825782 | 0.010213491168020700 | 0.141095284 |
| | Erlin2 | 1.40 | 2.355093168 | 0.004414757288412690 | 0.131469154 |
| | Chl1 | 1.40 | 3.151872318 | 0.000704900278113618 | 0.123542711 |
| | Gpam | 1.40 | 1.703423296 | 0.019795966211334000 | 0.160671218 |
| | Gltp | 1.40 | 1.381917879 | 0.041503251436698700 | 0.205392575 |

| | | | | | |
|---|---|---|---|---|---|
| | Itgb1 | 1.40 | 1.982850628 | 0.010402778984198000 | 0.141095284 |
| | Apcdd1 | 1.40 | 1.914463453 | 0.012176894602022000 | 0.143871633 |
| | Itih5 | 1.40 | 1.349259045 | 0.044744633448721100 | 0.211647507 |
| | Zfp518a | 1.40 | 2.716513199 | 0.001920820582676940 | 0.127020757 |
| | Aga | 1.40 | 1.580174623 | 0.026292106155580300 | 0.174878392 |
| | Myl12a | 1.40 | 1.729216197 | 0.018654508174838900 | 0.159967493 |
| | Timp2 | 1.40 | 1.74435474 | 0.018015456013295100 | 0.158663525 |
| | Rack1 | 1.40 | 1.557098003 | 0.027726943463983100 | 0.178703532 |
| | Wwtr1 | 1.40 | 1.828390655 | 0.014845996189574700 | 0.151437765 |
| | Sdc4 | 1.40 | 1.357126395 | 0.043941371243503600 | 0.210667443 |
| | Ss18 | 1.39 | 1.576002561 | 0.026545899092042300 | 0.175876181 |
| | 43345 | 1.39 | 2.029485159 | 0.009343612963960970 | 0.140783376 |
| | Rft1 | 1.39 | 2.293950725 | 0.005082171011219190 | 0.131469154 |
| | Snx33 | 1.39 | 1.703842348 | 0.019776874257833400 | 0.160671218 |
| | Odc1 | 1.39 | 1.892327081 | 0.012813651813454600 | 0.145881393 |
| | Clp1 | 1.39 | 1.679724799 | 0.020906204838376900 | 0.162468873 |
| | Wdr6 | 1.39 | 1.809626199 | 0.015501502710832700 | 0.151749058 |
| | Nipsnap3b | 1.39 | 1.897438208 | 0.012663734374025200 | 0.145739007 |
| | Tmsb4x | 1.39 | 1.985809476 | 0.010332145748444200 | 0.141095284 |
| | Prps1l3 | 1.39 | 1.825154411 | 0.014957037706943900 | 0.151437765 |
| | Vkorc1 | 1.39 | 1.341062037 | 0.045597177735336100 | 0.213460217 |
| | Zfp423 | 1.39 | 1.416563901 | 0.038320935214672900 | 0.200478413 |
| | Wdr5b | 1.39 | 1.987717078 | 0.010286862186033600 | 0.141095284 |

| | | | | | |
|---|---|---|---|---|---|
| | Kcns2 | 1.38 | 1.549155246 | 0.028238703527964600 | 0.179474072 |
| | Sccpdh | 1.38 | 2.147933706 | 0.007113220876369240 | 0.134660365 |
| | Tmed1 | 1.38 | 2.492791265 | 0.003215205490795370 | 0.128703904 |
| | Decr1 | 1.38 | 2.486987926 | 0.003258457600501810 | 0.128703904 |
| | Rps9 | 1.38 | 1.523256081 | 0.029973945883760400 | 0.183233156 |
| | Phgdh | 1.38 | 1.811297173 | 0.015441974360091000 | 0.151749058 |
| | Rpl22l1 | 1.38 | 2.491486282 | 0.003224881180233190 | 0.128703904 |
| | Nr3c2 | 1.38 | 2.504382192 | 0.003130529553875450 | 0.128703904 |
| | Rps6ka6 | 1.38 | 1.508310702 | 0.031023393260855200 | 0.185581230 |
| | Lsm5 | 1.38 | 1.683019569 | 0.020748200247458600 | 0.162078657 |
| | Rbm11 | 1.38 | 1.645352602 | 0.022628064001189400 | 0.166458646 |
| | Eef1a1 | 1.38 | 1.502664667 | 0.031429345174113400 | 0.186151134 |
| | Haus5 | 1.38 | 1.403786321 | 0.039465142866636800 | 0.202168336 |
| | Pold2 | 1.38 | 1.449204723 | 0.035546371594899900 | 0.196070699 |
| | Sorbs3 | 1.38 | 3.361922372 | 0.000434587897348989 | 0.123542711 |
| | Afap1 | 1.38 | 1.692807309 | 0.020285825781014900 | 0.161032659 |
| | Hltf | 1.38 | 1.438706814 | 0.036416079270336900 | 0.197424556 |
| | Dubr | 1.37 | 1.304522801 | 0.049599488622369700 | 0.221308908 |
| | Tom1l1 | 1.37 | 2.909586653 | 0.001231440256272330 | 0.127020757 |
| | Myo10 | 1.37 | 1.316624493 | 0.048236468724850500 | 0.218577458 |
| | Rnf113a2 | 1.37 | 1.577397344 | 0.026460780833096700 | 0.175566195 |
| | Snx7 | 1.37 | 1.391720184 | 0.040576988896180400 | 0.203309852 |
| | Reep3 | 1.37 | 2.270268717 | 0.005366996142150640 | 0.131469154 |

| | | | | | |
|---|---|---|---|---|---|
| | Kctd5 | 1.37 | 2.185776271 | 0.006519641709106200 | 0.133060702 |
| | Cd81 | 1.37 | 1.948893934 | 0.011248796660077300 | 0.141933868 |
| | Hdhd3 | 1.37 | 1.720674315 | 0.019025044677927000 | 0.160480708 |
| | Hibch | 1.37 | 1.301041792 | 0.049998641898001500 | 0.221634296 |
| | Hmgn2 | 1.37 | 1.800982745 | 0.015813108639958700 | 0.152122332 |
| | BC017643 | 1.37 | 2.347685269 | 0.004490707116382530 | 0.131469154 |
| | Mfsd1 | 1.37 | 1.743619418 | 0.018045984567719300 | 0.158751520 |
| | Rrbp1 | 1.37 | 1.485023545 | 0.032732294851830300 | 0.189458493 |
| | Ctsd | 1.37 | 1.388779287 | 0.040852695123147100 | 0.204032563 |
| | Egfr | 1.36 | 1.38325679 | 0.041375495702846700 | 0.205392575 |
| | Crim1 | 1.36 | 1.967058021 | 0.010788025867115300 | 0.141501604 |
| | Gng10 | 1.36 | 2.399966967 | 0.003981374518500150 | 0.131469154 |
| | Pgd | 1.36 | 1.581728295 | 0.026198215166653900 | 0.174737719 |
| | Hacd1 | 1.36 | 1.838432864 | 0.014506650098251400 | 0.150441568 |
| | Zdhhc15 | 1.36 | 1.347448066 | 0.044931605172207500 | 0.211951088 |
| | 2810001G20Rik | 1.36 | 2.582390806 | 0.002615828057044590 | 0.127347630 |
| | D11Wsu47e | 1.36 | 1.665880642 | 0.021583375059628600 | 0.163945360 |
| | Rnh1 | 1.36 | 1.795918402 | 0.015998585926760900 | 0.152122332 |
| | Crtc3 | 1.36 | 1.475713085 | 0.033441589754785400 | 0.191129135 |
| | Trim13 | 1.36 | 2.063853984 | 0.008632687421543970 | 0.140488765 |
| | Nampt | 1.36 | 1.645168966 | 0.022637634005042200 | 0.166458646 |
| | Mtmr10 | 1.36 | 1.795156748 | 0.016026668425938500 | 0.152122332 |
| | Tmed3 | 1.36 | 2.40118428 | 0.003970230483233340 | 0.131469154 |

| | | | | | |
|---|---|---|---|---|---|
| | Vat1 | 1.36 | 1.422878215 | 0.037767808484148100 | 0.199603556 |
| | Mtss1 | 1.36 | 1.716943809 | 0.019189170023477600 | 0.160482390 |
| | Dnajc24 | 1.36 | 1.54347561 | 0.028610430307115000 | 0.179884640 |
| | Ldlr | 1.36 | 3.087093147 | 0.000818289264367845 | 0.123542711 |
| | Gpr137b | 1.36 | 1.485876574 | 0.032668066117265700 | 0.189166412 |
| | Slc35b2 | 1.36 | 2.906017439 | 0.001241602451575600 | 0.127020757 |
| | Elovl5 | 1.36 | 1.928999551 | 0.011776071918031100 | 0.143647440 |
| | Trps1 | 1.36 | 1.526608147 | 0.029743484987893400 | 0.182539181 |
| | Rgs3 | 1.36 | 1.4766257 | 0.033371390269082800 | 0.191129135 |
| | Sh3bgrl | 1.36 | 2.662141384 | 0.002177000936922080 | 0.127020757 |
| | Pter | 1.36 | 1.860758184 | 0.013779765146801000 | 0.147977908 |
| | Disp1 | 1.36 | 1.305365949 | 0.049503288573330400 | 0.221308908 |
| | Rpp38 | 1.35 | 1.377448114 | 0.041932609164140400 | 0.205857800 |
| | 2210016F16Rik | 1.35 | 1.611653681 | 0.024453797920188600 | 0.169826196 |
| | Lims1 | 1.35 | 1.895569272 | 0.012718348702145700 | 0.145878492 |
| | Rnf141 | 1.35 | 2.850949506 | 0.001409452660739120 | 0.127020757 |
| | Clic4 | 1.35 | 1.421360176 | 0.037900053607010100 | 0.199763957 |
| | Kansl1l | 1.35 | 1.531697425 | 0.029396970421384500 | 0.181444365 |
| | Ikbip | 1.35 | 1.318837172 | 0.047991334710400900 | 0.218254040 |
| | Lrrcc1 | 1.35 | 2.536431112 | 0.002907829164227080 | 0.128703904 |
| | Rps27 | 1.35 | 2.091982263 | 0.008091289439227360 | 0.138221114 |
| | Bcl10 | 1.35 | 1.580280057 | 0.026285723958178800 | 0.174878392 |
| | 2310015A10Rik | 1.35 | 1.302041544 | 0.049883676667061200 | 0.221613838 |

| | | | | | |
|---|---|---|---|---|---|
| | Mmgt2 | 1.35 | 1.825510761 | 0.014944770122620300 | 0.151437765 |
| | Stxbp3 | 1.35 | 2.237175275 | 0.005791948950483310 | 0.131469154 |
| | Ndp | 1.35 | 1.44153359 | 0.036179820696073400 | 0.197087790 |
| | Ppp1r3c | 1.35 | 1.565580736 | 0.027190629634335200 | 0.177766864 |
| | H3f3a | 1.35 | 2.347214218 | 0.004495580536146880 | 0.131469154 |
| | Rgs14 | 1.35 | 1.589491048 | 0.025734098095260600 | 0.173610331 |
| | 1110038B12Rik | 1.35 | 1.396368615 | 0.040144992831472800 | 0.202779537 |
| | Prdm9 | 1.35 | 1.331652141 | 0.046595916533445800 | 0.215653247 |
| | Zfp810 | 1.35 | 2.096433719 | 0.008008778463951290 | 0.137560781 |
| | Frmd3 | 1.35 | 1.412775346 | 0.038656689048056600 | 0.200960331 |
| | Rbm7 | 1.35 | 2.292878739 | 0.005094731033956280 | 0.131469154 |
| | Ccdc90b | 1.35 | 1.462804741 | 0.034450478550238200 | 0.192834679 |
| | Dek | 1.35 | 2.796354991 | 0.001598251088713010 | 0.127020757 |
| | Mpp5 | 1.35 | 3.297777607 | 0.000503758506342762 | 0.123542711 |
| | Tjp2 | 1.35 | 1.499366439 | 0.031668942441328000 | 0.186764318 |
| | Rfx3 | 1.35 | 1.573381308 | 0.026706605525080800 | 0.176398764 |
| | Bphl | 1.35 | 2.264969225 | 0.005432888290622520 | 0.131469154 |
| | Heatr3 | 1.35 | 3.939730736 | 0.000114886570158549 | 0.112761169 |
| | Uvssa | 1.35 | 1.356133558 | 0.044041940159142400 | 0.210717375 |
| | Uxt | 1.34 | 1.541775274 | 0.028722664522664300 | 0.179932351 |
| | Nkain4 | 1.34 | 2.019176107 | 0.009568060073643090 | 0.140783376 |
| | Scarb2 | 1.34 | 2.347015607 | 0.004497636914846430 | 0.131469154 |
| | Gna13 | 1.34 | 1.731715405 | 0.018547466489638500 | 0.159967493 |

| | | | | | |
|---|---|---|---|---|---|
| | Sdc2 | 1.34 | 1.813488364 | 0.015364259597985800 | 0.151749058 |
| | Trim26 | 1.34 | 1.545810016 | 0.028457057009185500 | 0.179474072 |
| | Rad50 | 1.34 | 2.265206116 | 0.005429925660019540 | 0.131469154 |
| | Selenos | 1.34 | 3.846871105 | 0.000142275098578740 | 0.119849580 |
| | Gtf2e1 | 1.34 | 1.761549188 | 0.017316128995976500 | 0.156718571 |
| | Snx10 | 1.34 | 2.601421874 | 0.002503676000865720 | 0.127020757 |
| | Cwf19l2 | 1.34 | 1.801389436 | 0.015798307531618700 | 0.152122332 |
| | H2afv | 1.34 | 2.043074276 | 0.009055777098199860 | 0.140488765 |
| | Egln3 | 1.34 | 2.122299114 | 0.007545723474336960 | 0.136287102 |
| | Ugdh | 1.34 | 1.306634816 | 0.049358867159315600 | 0.221308908 |
| | Galk1 | 1.34 | 1.583708437 | 0.026079037732037600 | 0.174737719 |
| | Lactb2 | 1.34 | 1.352031719 | 0.044459879426492700 | 0.211392112 |
| | Pcsk6 | 1.34 | 1.346244078 | 0.045056341157664600 | 0.212245178 |
| | Il11ra1 | 1.34 | 1.399203652 | 0.039883783336756800 | 0.202412377 |
| | 1190002N15Rik | 1.34 | 1.68705445 | 0.020556328530244200 | 0.161859432 |
| | Dpagt1 | 1.34 | 1.821436167 | 0.015085643246511800 | 0.151522906 |
| | Wls | 1.34 | 2.39566063 | 0.004021049039330210 | 0.131469154 |
| | Zfp935 | 1.34 | 1.46080157 | 0.034609747421384000 | 0.193270786 |
| | Syne2 | 1.34 | 1.411866119 | 0.038737704409907400 | 0.201153343 |
| | Pltp | 1.33 | 1.71344977 | 0.019344175783180900 | 0.160482390 |
| | Magt1 | 1.33 | 1.908625205 | 0.012341694601916700 | 0.145127603 |
| | Wdr83os | 1.33 | 2.037491528 | 0.009172938298421880 | 0.140488765 |
| | Tprkb | 1.33 | 1.896987476 | 0.012676884220731300 | 0.145765935 |

| | | | | | |
|---|---|---|---|---|---|
| | Lrrc8d | 1.33 | 2.729598386 | 0.001863809898802660 | 0.127020757 |
| | Fam135a | 1.33 | 2.455501712 | 0.003503469071404330 | 0.129362173 |
| | Mrps6 | 1.33 | 1.538035996 | 0.028971034522669500 | 0.180151698 |
| | P2rx4 | 1.33 | 1.909452533 | 0.012318206137926700 | 0.145042391 |
| | Slc38a3 | 1.33 | 2.029051528 | 0.009352946973680660 | 0.140783376 |
| | Ech1 | 1.33 | 1.573050272 | 0.026726970142808100 | 0.176398764 |
| | Btg3 | 1.33 | 1.917303225 | 0.012097531869488900 | 0.143647440 |
| | Dpp7 | 1.33 | 1.531770777 | 0.029392005681633300 | 0.181444365 |
| | Edem2 | 1.33 | 1.37070365 | 0.042588892810384000 | 0.207669970 |
| | Smpdl3a | 1.33 | 2.115991005 | 0.007656124632320470 | 0.136287102 |
| | 2610035D17Rik | 1.33 | 1.315280543 | 0.048385970556315000 | 0.218923813 |
| | Kirrel3 | 1.33 | 1.350300478 | 0.044637464968458500 | 0.211647507 |
| | Rpl22 | 1.33 | 2.226545926 | 0.005935455779557870 | 0.131469154 |
| | Zfp51 | 1.33 | 1.870582454 | 0.013471549335327300 | 0.147515895 |
| | Tamm41 | 1.33 | 1.771471184 | 0.016925005405668800 | 0.155147765 |
| | Slc9a3r1 | 1.33 | 1.418521587 | 0.038148583156040600 | 0.200173662 |
| | C330018D20Rik | 1.33 | 1.658685614 | 0.021943928844564900 | 0.164777465 |
| | Stn1 | 1.33 | 2.139366972 | 0.007254926683088020 | 0.135646175 |
| | Rgs10 | 1.33 | 1.429575999 | 0.037189813547565800 | 0.198909723 |
| | 6330403K07Rik | 1.33 | 1.643359161 | 0.022732167052080300 | 0.166786472 |
| | Tpmt | 1.32 | 1.33799659 | 0.045920161821785300 | 0.214257706 |
| | Oplah | 1.32 | 2.424684084 | 0.003761108967201780 | 0.129856967 |
| | Snx4 | 1.32 | 3.287689926 | 0.000515596634288972 | 0.123542711 |

| | | | | | |
|---|---|---|---|---|---|
| | Etfrf1 | 1.32 | 2.238010334 | 0.005780822918044890 | 0.131469154 |
| | Tmem18 | 1.32 | 3.113970927 | 0.000769181929686484 | 0.123542711 |
| | Fance | 1.32 | 1.561243997 | 0.027463507521491900 | 0.178595240 |
| | Ssr3 | 1.32 | 2.749900487 | 0.001778686925340560 | 0.127020757 |
| | Lhfp | 1.32 | 1.403362985 | 0.039503630898160400 | 0.202168336 |
| | Tmpo | 1.32 | 1.42691839 | 0.037418089571766400 | 0.199280377 |
| | Rpl34-ps1 | 1.32 | 1.484032407 | 0.032807081157098200 | 0.189571952 |
| | Rps18 | 1.32 | 1.657835714 | 0.021986914404784400 | 0.164780689 |
| | Msl3l2 | 1.32 | 1.652911894 | 0.022237609829083100 | 0.165260680 |
| | 2610001J05Rik | 1.32 | 1.967158894 | 0.010785520440521500 | 0.141501604 |
| | Nlrx1 | 1.32 | 1.599717357 | 0.025135217249950000 | 0.171535493 |
| | Gorab | 1.32 | 1.894122607 | 0.012760785029781600 | 0.145878492 |
| | Acadm | 1.32 | 2.036290229 | 0.009198346625913580 | 0.140488765 |
| | Rpl13a | 1.32 | 2.33991535 | 0.004571772908824800 | 0.131469154 |
| | Hs6st2 | 1.32 | 1.301108795 | 0.049990928711823300 | 0.221634296 |
| | Rnf113a1 | 1.32 | 1.524610393 | 0.029880620174382400 | 0.183082275 |
| | Lamp1 | 1.32 | 2.228907481 | 0.005903268253497360 | 0.131469154 |
| | Rpl27 | 1.32 | 2.097017974 | 0.007998011529367570 | 0.137560781 |
| | Sumf2 | 1.32 | 1.728819255 | 0.018671566058292200 | 0.159967493 |
| | Fgfr2 | 1.32 | 1.903689094 | 0.012482768215623500 | 0.145640913 |
| | Tspan12 | 1.32 | 1.65299608 | 0.022233299582418900 | 0.165260680 |
| | Sp1 | 1.32 | 1.82843602 | 0.014844445524016300 | 0.151437765 |
| | Ltbp3 | 1.32 | 1.376693058 | 0.042005575747282500 | 0.206068767 |

| | | | | | |
|---|---|---|---|---|---|
| | F3 | 1.32 | 1.42144609 | 0.037892556807061900 | 0.199763957 |
| | Rps5 | 1.32 | 1.689068573 | 0.020461215391410400 | 0.161584805 |
| | Cenpt | 1.32 | 1.878848504 | 0.013217566249974900 | 0.146825043 |
| | Itm2c | 1.32 | 1.849733028 | 0.014134061332219100 | 0.149025511 |
| | Tuba1a | 1.32 | 2.034860329 | 0.009228681781518510 | 0.140488765 |
| | Tab2 | 1.31 | 1.761162006 | 0.017331573516023600 | 0.156718571 |
| | Cd2ap | 1.31 | 1.763448354 | 0.017240571042200700 | 0.156474694 |
| | Asl | 1.31 | 1.407847406 | 0.039097824579791400 | 0.201809641 |
| | Pcdh19 | 1.31 | 1.428068756 | 0.037319107111184700 | 0.199146350 |
| | Kctd18 | 1.31 | 1.89767238 | 0.012656907893607400 | 0.145739007 |
| | Eci1 | 1.31 | 2.323732044 | 0.004745346791654920 | 0.131469154 |
| | Scfd1 | 1.31 | 2.466146808 | 0.003418638598827060 | 0.129362173 |
| | Alcam | 1.31 | 1.940426678 | 0.011470261589020200 | 0.142249878 |
| | Mfsd9 | 1.31 | 1.316215129 | 0.048281957650353100 | 0.218669209 |
| | Dnah1 | 1.31 | 1.480228019 | 0.033095731196909500 | 0.190389012 |
| | Amdhd2 | 1.31 | 1.32869151 | 0.046914650992822100 | 0.216399537 |
| | Smyd4 | 1.31 | 2.059032624 | 0.008729057936699680 | 0.140488765 |
| | Rps3 | 1.31 | 2.359512982 | 0.004370056159559410 | 0.131469154 |
| | Hhatl | 1.31 | 1.358303576 | 0.043822426748331000 | 0.210611848 |
| | Selenbp1 | 1.31 | 1.456572096 | 0.034948448762596900 | 0.194290171 |
| | Spg20 | 1.31 | 1.788497479 | 0.016274307632903000 | 0.152555604 |
| | Pfn4 | 1.31 | 1.431493308 | 0.037025991038512100 | 0.198531875 |
| | Znrf2 | 1.31 | 1.394620286 | 0.040306929334424100 | 0.203107134 |

| | | | | | |
|---|---|---|---|---|---|
| | Adamtsl2 | 1.31 | 1.387834645 | 0.040941651268889400 | 0.204032563 |
| | Aldh9a1 | 1.31 | 2.386334098 | 0.004108335493242360 | 0.131469154 |
| | Nectin3 | 1.31 | 1.46733151 | 0.034093256859793500 | 0.192101954 |
| | Vps25 | 1.31 | 2.908719556 | 0.001233901362680910 | 0.127020757 |
| | Alg14 | 1.31 | 1.445350113 | 0.035863270090291500 | 0.196568486 |
| | Rev3l | 1.31 | 1.507672555 | 0.031069012175987600 | 0.185593785 |
| | Bloc1s1 | 1.31 | 1.634394368 | 0.023206285557186100 | 0.167214247 |
| | Dhx40 | 1.31 | 1.48450362 | 0.032771504538545900 | 0.189525776 |
| | Fam204a | 1.31 | 2.316848048 | 0.004821164518454680 | 0.131469154 |
| | Ascc1 | 1.31 | 1.304391639 | 0.049614470565163900 | 0.221308908 |
| | Rin2 | 1.31 | 1.322415887 | 0.047597496720434200 | 0.217555144 |
| | St3gal6 | 1.31 | 1.455641503 | 0.035023415566423500 | 0.194290171 |
| | Prmt6 | 1.31 | 1.301663245 | 0.049927147588913700 | 0.221634296 |
| | Dock5 | 1.31 | 1.499822039 | 0.031635737321992800 | 0.186755061 |
| | Nupl2 | 1.31 | 1.524884517 | 0.029861765676659700 | 0.183082275 |
| | Papss1 | 1.31 | 2.34634824 | 0.004504553612612350 | 0.131469154 |
| | Fads2 | 1.31 | 1.312305497 | 0.048718566704880000 | 0.219777356 |
| | Slc4a7 | 1.30 | 1.382105172 | 0.041485356680038600 | 0.205392575 |
| | Idh1 | 1.30 | 1.582313957 | 0.026162909727479500 | 0.174737719 |
| | Rpn2 | 1.30 | 1.719874343 | 0.019060121166383500 | 0.160480708 |
| | Pmp22 | 1.30 | 2.016603757 | 0.009624900382999550 | 0.140783376 |
| | Idua | 1.30 | 1.329600838 | 0.046816523679359600 | 0.216164601 |
| | Rdh11 | 1.30 | 2.783222861 | 0.001647316842576320 | 0.127020757 |

| | | | | | |
|---|---|---|---|---|---|
| | Tcp11l2 | 1.30 | 1.943112147 | 0.011399553817849000 | 0.141961618 |
| | Rbm12 | 1.30 | 1.668925949 | 0.021432560126603000 | 0.163341547 |
| | Rps12 | 1.30 | 1.533754621 | 0.029258050107863600 | 0.181205864 |
| | Zc3h6 | 1.30 | 1.613961913 | 0.024324173213254000 | 0.169664195 |
| | Cep44 | 1.30 | 1.4751587 | 0.033484305831371800 | 0.191129135 |
| | Kdelc2 | 1.30 | 1.615842731 | 0.024219059204987100 | 0.169664195 |
| | Tmem200c | 1.30 | 1.312553854 | 0.048690714363269200 | 0.219723844 |
| | Slc1a3 | 1.30 | 1.327516628 | 0.047041739454383300 | 0.216606590 |
| | Itm2b | 1.30 | 2.101960265 | 0.007907509723512460 | 0.137039524 |
| | Slc16a4 | 1.30 | 1.316665147 | 0.048231953629888700 | 0.218577458 |
| | Zhx2 | 1.30 | 1.505157697 | 0.031249444627561100 | 0.185821360 |
| | Sh2d3c | 1.30 | 1.885979324 | 0.013002314785430200 | 0.146312614 |
| | Tmem258 | 1.30 | 1.67322838 | 0.021221282150055500 | 0.162736895 |
| | Lrrc3 | 1.30 | 1.345925057 | 0.045089450578951700 | 0.212328355 |
| | Ilvbl | 1.30 | 2.302281782 | 0.004985609023525210 | 0.131469154 |
| | Ctnnal1 | 1.30 | 1.447544249 | 0.035682539128250500 | 0.196291813 |
| | Slc7a2 | 1.30 | 1.399098902 | 0.039893404281964600 | 0.202412377 |
| | Etfdh | 1.30 | 2.312426915 | 0.004870494816536940 | 0.131469154 |
| | Zfp90 | 1.30 | 1.601496661 | 0.025032448948622700 | 0.171129792 |
| | Ift74 | 1.30 | 2.121232148 | 0.007564284459662870 | 0.136287102 |
| | Ost4 | 1.30 | 1.718131758 | 0.019136752571260600 | 0.160480708 |
| | Dpy19l4 | 1.30 | 1.581813309 | 0.026193087311372400 | 0.174737719 |
| | Sppl2a | 1.30 | 1.42867637 | 0.037266931101256600 | 0.198960061 |

| | | | | | |
|---|---|---|---|---|---|
| | Pld1 | 1.30 | 1.356635605 | 0.043991056769154300 | 0.210694009 |
| | Ece2 | 1.30 | 1.474854646 | 0.033507756721600900 | 0.191129135 |
| | Calu | 1.29 | 2.112003612 | 0.007726741586144950 | 0.136821077 |
| | Sumf1 | 1.29 | 1.959346393 | 0.010981296218876600 | 0.141933868 |
| | Zfp606 | 1.29 | 1.714864343 | 0.019281270943421300 | 0.160482390 |
| | Sdhaf1 | 1.29 | 2.106505259 | 0.007825187299920290 | 0.137039524 |
| | Prrc1 | 1.29 | 1.946961762 | 0.011298953935043500 | 0.141933868 |
| | Tmbim6 | 1.29 | 1.628553178 | 0.023520514740441600 | 0.167968850 |
| | Rps27a | 1.29 | 1.983882799 | 0.010378084471653700 | 0.141095284 |
| | Pnrc2 | 1.29 | 1.735853979 | 0.018371559366658400 | 0.159370958 |
| | Plgrkt | 1.29 | 1.320356281 | 0.047823760076461700 | 0.218103647 |
| | Alg10b | 1.29 | 1.913577107 | 0.012201771648607300 | 0.144041705 |
| | Cav2 | 1.29 | 1.969614766 | 0.010724701999609800 | 0.141501604 |
| | Dag1 | 1.29 | 1.792279032 | 0.016133216726314300 | 0.152361877 |
| | Cyb5r3 | 1.29 | 1.595746103 | 0.025366111475018500 | 0.172334465 |
| | Parvb | 1.29 | 1.955500764 | 0.011078966160999500 | 0.141933868 |
| | Rnf13 | 1.29 | 2.48771975 | 0.003252971437418940 | 0.128703904 |
| | Maoa | 1.29 | 2.535108962 | 0.002916695140100040 | 0.128703904 |
| | Ehd2 | 1.29 | 1.553652219 | 0.027947809897849900 | 0.178882497 |
| | Fam136a | 1.29 | 2.23421744 | 0.005831530619880970 | 0.131469154 |
| | Yif1a | 1.29 | 1.553028792 | 0.027987957618600100 | 0.178958830 |
| | Gpr19 | 1.29 | 2.043450494 | 0.009047935714985440 | 0.140488765 |
| | Nphp1 | 1.29 | 1.661108342 | 0.021821854611241400 | 0.164554124 |

| | | | | | |
|---|---|---|---|---|---|
| | Spef1 | 1.29 | 1.4314394<br>9 | 0.0370305796069<br>14900 | 0.1985318<br>75 |
| | Rpl15 | 1.29 | 2.8613185<br>89 | 0.0013761995476<br>29940 | 0.1270207<br>57 |
| | Rps4x | 1.29 | 1.6500917<br>93 | 0.0223824801088<br>20800 | 0.1661575<br>68 |
| | Wdr43 | 1.29 | 1.8849551<br>51 | 0.0130330136066<br>37000 | 0.1463126<br>14 |
| | Arl4a | 1.29 | 3.4837559<br>57 | 0.0003282797117<br>27804 | 0.1235427<br>11 |
| | Smad1 | 1.29 | 1.8603260<br>9 | 0.0137934819271<br>03000 | 0.1479779<br>08 |
| | Tpt1 | 1.28 | 1.4988852<br>46 | 0.0317040506912<br>94600 | 0.1867643<br>18 |
| | Sec11c | 1.28 | 1.5511929<br>33 | 0.0281065193422<br>95800 | 0.1794338<br>75 |
| | Gpr137c | 1.28 | 1.5291934<br>47 | 0.0295669517955<br>68300 | 0.1821555<br>77 |
| | Pex2 | 1.28 | 1.8526358<br>71 | 0.0140399037024<br>82200 | 0.1486467<br>76 |
| | 2610301B20<br>Rik | 1.28 | 2.3910716<br>94 | 0.0040637623825<br>84000 | 0.1314691<br>54 |
| | Rps19 | 1.28 | 1.3611122<br>73 | 0.0435399300135<br>32200 | 0.2098624<br>94 |
| | Nucb2 | 1.28 | 1.6957280<br>18 | 0.0201498576189<br>07300 | 0.1609205<br>15 |
| | Fuz | 1.28 | 1.3422805<br>92 | 0.0454694193757<br>23700 | 0.2132407<br>14 |
| | Rps15a | 1.28 | 1.8864720<br>99 | 0.0129875699776<br>32200 | 0.1462804<br>91 |
| | Tubb2b | 1.28 | 1.3377972<br>13 | 0.0459412478533<br>86900 | 0.2142833<br>29 |
| | Fau | 1.28 | 1.3638381<br>61 | 0.0432675036691<br>16200 | 0.2091237<br>31 |
| | Zfp85 | 1.28 | 1.5065539<br>74 | 0.0311491374851<br>62900 | 0.1856934<br>15 |
| | Prps2 | 1.28 | 3.1033338<br>61 | 0.0007882539192<br>84104 | 0.1235427<br>11 |
| | Rpl7 | 1.28 | 1.8001018<br>95 | 0.0158452138700<br>61600 | 0.1521223<br>32 |
| | Serpine2 | 1.28 | 1.3327555<br>08 | 0.0464776854247<br>56100 | 0.2153236<br>26 |
| | Vamp3 | 1.28 | 1.9901892<br>63 | 0.0102284714449<br>42900 | 0.1410952<br>84 |
| | Itm2a | 1.28 | 2.2132904<br>43 | 0.0061194100715<br>78610 | 0.1314691<br>54 |

| | | | | | |
|---|---|---|---|---|---|
| | Sypl | 1.28 | 1.620823624 | 0.023942879263869000 | 0.168887506 |
| | Gm2a | 1.28 | 1.416923212 | 0.038289243677340300 | 0.200478413 |
| | Mgat2 | 1.28 | 1.777072646 | 0.016708111077479500 | 0.154188149 |
| | Macrod1 | 1.28 | 1.73074917 | 0.018588777522445200 | 0.159967493 |
| | Tmem209 | 1.28 | 1.751907955 | 0.017704841587474200 | 0.157464225 |
| | Pcna | 1.28 | 2.362904236 | 0.004336064802186570 | 0.131469154 |
| | Ppie | 1.28 | 1.365310724 | 0.043121044915242400 | 0.208758381 |
| | Pgrmc1 | 1.28 | 2.283253558 | 0.005208905064192000 | 0.131469154 |
| | Fam58b | 1.28 | 1.363301754 | 0.043320977336445100 | 0.209234991 |
| | Gabra5 | 1.28 | 1.420526992 | 0.037972833738036600 | 0.200069899 |
| | Amn1 | 1.28 | 1.731830134 | 0.018542567408416000 | 0.159967493 |
| | Gyg | 1.28 | 1.880353864 | 0.013171830573008800 | 0.146825043 |
| | Plekhg1 | 1.28 | 1.523823672 | 0.029934797737714800 | 0.183222297 |
| | Slc35f5 | 1.27 | 1.966556943 | 0.010800480002634100 | 0.141501604 |
| | Cln6 | 1.27 | 1.38185772 | 0.041509000900346800 | 0.205392575 |
| | Rap1b | 1.27 | 1.599604879 | 0.025141727876237700 | 0.171535493 |
| | Trmt6 | 1.27 | 2.378844531 | 0.004179799681519310 | 0.131469154 |
| | Mpst | 1.27 | 1.783779061 | 0.016452084795849300 | 0.153278292 |
| | Nudt21 | 1.27 | 1.820949858 | 0.015102545130924500 | 0.151522906 |
| | Gng12 | 1.27 | 1.305921569 | 0.049439996505234000 | 0.221308908 |
| | Cd164 | 1.27 | 1.969585296 | 0.010725429795148400 | 0.141501604 |
| | Rasgef1c | 1.27 | 1.428700425 | 0.037264866934856800 | 0.198960061 |
| | Dync2li1 | 1.27 | 1.558452784 | 0.027640584063897100 | 0.178595240 |

| | | | | | |
|---|---|---|---|---|---|
| | Galk2 | 1.27 | 1.917503715 | 0.012091948401675600 | 0.143647440 |
| | Abcg2 | 1.27 | 1.371034005 | 0.042556509056830400 | 0.207659443 |
| | Psmb1 | 1.27 | 2.625807294 | 0.002366969740194120 | 0.127020757 |
| | Ptma | 1.27 | 1.472858779 | 0.033662101140280700 | 0.191453200 |
| | Anapc10 | 1.27 | 1.860628648 | 0.013783875840108800 | 0.147977908 |
| | Tmed10 | 1.27 | 2.431536053 | 0.003702234700343040 | 0.129856967 |
| | Rpl36a | 1.27 | 1.50459015 | 0.031290308847105900 | 0.185821360 |
| | Rpsa | 1.27 | 1.787816232 | 0.016299856029658600 | 0.152555604 |
| | Ofd1 | 1.27 | 1.543205175 | 0.028628251553712400 | 0.179884640 |
| | Mtdh | 1.27 | 3.003459864 | 0.000992065019367351 | 0.127020757 |
| | Zadh2 | 1.27 | 1.696081854 | 0.020133447472486900 | 0.160920515 |
| | Bin3 | 1.27 | 1.713933257 | 0.019322652466323000 | 0.160482390 |
| | Cenpx | 1.27 | 1.764607244 | 0.017194626877167300 | 0.156367550 |
| | Sri | 1.27 | 1.645134559 | 0.022639427534161900 | 0.166458646 |
| | Lgals8 | 1.27 | 1.710727364 | 0.019465816987639000 | 0.160482390 |
| | Arfgap3 | 1.27 | 2.508587085 | 0.003100365637001090 | 0.128703904 |
| | Zfp687 | 1.27 | 2.109890385 | 0.007764430639743920 | 0.137033900 |
| | Naf1 | 1.27 | 1.654389319 | 0.022162088328960600 | 0.165260680 |
| | Mterf2 | 1.27 | 1.710389209 | 0.019480979580640200 | 0.160482390 |
| | Exo5 | 1.27 | 1.349599077 | 0.044709614249009500 | 0.211647507 |
| | Rps7 | 1.27 | 2.13696635 | 0.007295140319097450 | 0.135646175 |
| | Tmem216 | 1.27 | 1.356841768 | 0.043970178803955100 | 0.210667443 |
| | Ergic3 | 1.27 | 2.154672724 | 0.007003695817804620 | 0.134660365 |

| | | | | | |
|---|---|---|---|---|---|
| | B4galt4 | 1.27 | 1.409164217 | 0.038979456791216900 | 0.201744197 |
| | Nudt16 | 1.27 | 1.444411105 | 0.035940895598995300 | 0.196696198 |
| | Slc38a1 | 1.26 | 1.453557094 | 0.035191915482762700 | 0.194674763 |
| | Rpl23 | 1.26 | 1.496197257 | 0.031900885810564400 | 0.187100141 |
| | Dars2 | 1.26 | 2.150552177 | 0.007070462495557250 | 0.134660365 |
| | Naxe | 1.26 | 2.152997609 | 0.007030761906186160 | 0.134660365 |
| | M6pr | 1.26 | 2.069716367 | 0.008516940894609180 | 0.140434588 |
| | Adcyap1r1 | 1.26 | 1.382346946 | 0.041462268003425100 | 0.205392575 |
| | Prkab1 | 1.26 | 1.449057605 | 0.035558415083076300 | 0.196070699 |
| | Tctn2 | 1.26 | 2.179188057 | 0.006619298149422460 | 0.133240732 |
| | Zfp941 | 1.26 | 1.802859867 | 0.015744908211332000 | 0.152122332 |
| | Sec11a | 1.26 | 1.416678268 | 0.038310845087733200 | 0.200478413 |
| | Fam19a1 | 1.26 | 1.405957902 | 0.039268299763571800 | 0.201979414 |
| | Cat | 1.26 | 2.026093487 | 0.009416868654506550 | 0.140783376 |
| | Zfp212 | 1.26 | 1.71206774 | 0.019405831683855500 | 0.160482390 |
| | Grk5 | 1.26 | 1.882295379 | 0.013113077296197000 | 0.146652067 |
| | Cyp2d22 | 1.26 | 1.458986498 | 0.034754696652662900 | 0.193856046 |
| | A830080D01Rik | 1.26 | 1.546734158 | 0.028396567207153900 | 0.179474072 |
| | Pigp | 1.26 | 1.613484791 | 0.024350910733434000 | 0.169764518 |
| | Cyp2j6 | 1.26 | 1.313190692 | 0.048619367838306300 | 0.219541049 |
| | Nwd2 | 1.26 | 1.899050795 | 0.012616799587334300 | 0.145739007 |
| | Rab8b | 1.26 | 1.41043078 | 0.038865943921845900 | 0.201454898 |
| | Kdelr2 | 1.26 | 1.420073429 | 0.038012512043425200 | 0.200069899 |

| | | | | | |
|---|---|---|---|---|---|
| | Slc1a4 | 1.26 | 1.438322247 | 0.036448339982968500 | 0.197424556 |
| | Gprc5b | 1.26 | 1.573316577 | 0.026710586412661600 | 0.176398764 |
| | Zfp944 | 1.26 | 1.593519232 | 0.025496511780762200 | 0.172840438 |
| | Rps3a1 | 1.26 | 2.200578771 | 0.006301170463858810 | 0.132111948 |
| | Tgs1 | 1.26 | 1.80557734 | 0.015646696558553900 | 0.151944352 |
| | Smim20 | 1.26 | 1.722576501 | 0.018941898230708700 | 0.160468942 |
| | Notch3 | 1.26 | 1.567543547 | 0.027068017751897000 | 0.177189689 |
| | Rfc2 | 1.26 | 2.248760964 | 0.005639479676806140 | 0.131469154 |
| | Plpp3 | 1.26 | 1.324703571 | 0.047347431947182800 | 0.217172036 |
| | Fam76b | 1.26 | 1.526745247 | 0.029734096931967900 | 0.182539181 |
| | Dhps | 1.26 | 1.675087431 | 0.021130636026638000 | 0.162735056 |
| | Eef1b2 | 1.26 | 1.728661774 | 0.018678337821811100 | 0.159967493 |
| | Cetn2 | 1.25 | 1.419558932 | 0.038057571152854200 | 0.200069899 |
| | Lamc1 | 1.25 | 1.368590888 | 0.042796584661378400 | 0.207981123 |
| | Slc3a2 | 1.25 | 1.551225428 | 0.028104416420066300 | 0.179433875 |
| | Txn1 | 1.25 | 1.493380542 | 0.032108458645655700 | 0.187665815 |
| | Twf1 | 1.25 | 1.755208891 | 0.017570782740361400 | 0.157085313 |
| | Sav1 | 1.25 | 1.462604838 | 0.034466339528871300 | 0.192834679 |
| | Fbxl4 | 1.25 | 1.585999701 | 0.025941811510821200 | 0.174311214 |
| | Ctsl | 1.25 | 1.497775333 | 0.031785179418158000 | 0.187015633 |
| | Erlec1 | 1.25 | 2.047771648 | 0.008958356716394610 | 0.140488765 |
| | Mapkapk2 | 1.25 | 1.335390265 | 0.046196570387070500 | 0.214941372 |
| | Wars2 | 1.25 | 1.419316057 | 0.038078860435903600 | 0.200069899 |

| | | | | | |
|---|---|---|---|---|---|
| | Mesd | 1.25 | 2.323042146 | 0.004752890998799090 | 0.131469154 |
| | Plpp1 | 1.25 | 1.425693563 | 0.037523767522188500 | 0.199541056 |
| | Rnf130 | 1.25 | 2.011347471 | 0.009742098787191260 | 0.140783376 |
| | Hadh | 1.25 | 1.611200514 | 0.024479327667331300 | 0.169826196 |
| | Derl1 | 1.25 | 2.222773272 | 0.005987240832246600 | 0.131469154 |
| | Seh1l | 1.25 | 3.108059905 | 0.000779722550192238 | 0.123542711 |
| | Dars | 1.25 | 2.040385507 | 0.009112016398311570 | 0.140488765 |
| | Prpsap2 | 1.25 | 1.736110045 | 0.018360730465597100 | 0.159370958 |
| | Rhoa | 1.25 | 2.022824057 | 0.009488027683429810 | 0.140783376 |
| | Fundc1 | 1.25 | 2.337065269 | 0.004601874077605120 | 0.131469154 |
| | Sp4 | 1.25 | 2.784417926 | 0.001642790087700600 | 0.127020757 |
| | Mtmr14 | 1.25 | 1.465816791 | 0.034212373812496600 | 0.192402580 |
| | Jrk | 1.25 | 1.51477783 | 0.030564843017635500 | 0.184774091 |
| | Smyd3 | 1.25 | 1.822236716 | 0.015057860994903600 | 0.151522906 |
| | Lrch2 | 1.25 | 1.381871078 | 0.041507724153812000 | 0.205392575 |
| | Nudcd2 | 1.25 | 1.901383244 | 0.012549220667016000 | 0.145640913 |
| | Rplp1 | 1.25 | 1.306001005 | 0.049430954313544900 | 0.221308908 |
| | Gmps | 1.24 | 1.520520486 | 0.030163345879775100 | 0.183414716 |
| | Stag1 | 1.24 | 1.423369327 | 0.037725123755487300 | 0.199593499 |
| | Eif3e | 1.24 | 2.161817732 | 0.006889413764304150 | 0.134513024 |
| | Atl3 | 1.24 | 1.664790486 | 0.021637621239456800 | 0.163994790 |
| | Ddx18 | 1.24 | 1.747715044 | 0.017876601389692400 | 0.158172814 |
| | Lman1 | 1.24 | 1.569385144 | 0.026953480629486200 | 0.176702184 |

| | | | | | |
|---|---|---|---|---|---|
| | Dcps | 1.24 | 1.406133298 | 0.039252443929330400 | 0.201979414 |
| | Adk | 1.24 | 1.708741094 | 0.019555048881841700 | 0.160482390 |
| | Mzt2 | 1.24 | 2.033351135 | 0.009260807682168720 | 0.140488765 |
| | Topbp1 | 1.24 | 1.845821106 | 0.014261949488029700 | 0.149786882 |
| | Smim15 | 1.24 | 2.050751045 | 0.008897109890973180 | 0.140488765 |
| | Zkscan16 | 1.24 | 1.37137132 | 0.042523468352437400 | 0.207645692 |
| | Rfc5 | 1.24 | 1.400873822 | 0.039730696493583900 | 0.202412377 |
| | Ddx52 | 1.24 | 1.902552228 | 0.012515487534000900 | 0.145640913 |
| | Pcdhgb6 | 1.24 | 1.319288191 | 0.047941521076580800 | 0.218206175 |
| | Stk3 | 1.24 | 1.421582933 | 0.037880618969915700 | 0.199763957 |
| | Rpl10 | 1.24 | 1.315546924 | 0.048356301446240200 | 0.218869849 |
| | Dolk | 1.24 | 1.752414499 | 0.017684203401296900 | 0.157464225 |
| | Rap1a | 1.24 | 1.316740784 | 0.048223554266352400 | 0.218577458 |
| | Tcta | 1.24 | 2.147244142 | 0.007124524073801620 | 0.134660365 |
| | Hdac9 | 1.24 | 1.546053954 | 0.028441077531999100 | 0.179474072 |
| | Snx5 | 1.24 | 2.418745707 | 0.003812890148647200 | 0.130257845 |
| | Galnt1 | 1.24 | 1.668305515 | 0.021463200639401500 | 0.163484390 |
| | Asnsd1 | 1.24 | 1.75604593 | 0.017536950254180600 | 0.157085313 |
| | Mettl2 | 1.24 | 1.547824966 | 0.028325333660335200 | 0.179474072 |
| | Osgepl1 | 1.24 | 1.641345288 | 0.022837823463929300 | 0.166831290 |
| | Fbxo6 | 1.23 | 1.64073021 | 0.022870190902531500 | 0.166831290 |
| | Pdia3 | 1.23 | 1.435983854 | 0.036645119828552000 | 0.197777137 |
| | Nsa2 | 1.23 | 1.444375662 | 0.035943828885872100 | 0.196696198 |

| | | | | | |
|---|---|---|---|---|---|
| | Snrpb2 | 1.23 | 2.05472923 | 0.008815983534876540 | 0.140488765 |
| | Mgat1 | 1.23 | 2.089672333 | 0.008134440130477590 | 0.138507239 |
| | Snrpg | 1.23 | 1.302459048 | 0.049835744679301300 | 0.221544150 |
| | Adh5 | 1.23 | 2.198428102 | 0.006332451872128930 | 0.132111948 |
| | Taf10 | 1.23 | 1.416392386 | 0.038336072153840200 | 0.200478413 |
| | Vhl | 1.23 | 2.234307411 | 0.005830322650364760 | 0.131469154 |
| | Parp8 | 1.23 | 1.424023642 | 0.037668329264087100 | 0.199593499 |
| | Txndc9 | 1.23 | 2.479265553 | 0.003316915800756050 | 0.128703904 |
| | Atg5 | 1.23 | 1.986154565 | 0.010323939122952500 | 0.141095284 |
| | Snapin | 1.23 | 2.341304916 | 0.004557168470776380 | 0.131469154 |
| | Rps24 | 1.23 | 1.798480687 | 0.015904474095210100 | 0.152122332 |
| | Bmper | 1.23 | 1.372653733 | 0.042398087507807100 | 0.207401965 |
| | Rpl35 | 1.23 | 2.134822867 | 0.007331234872225210 | 0.136112614 |
| | Cryl1 | 1.23 | 1.470506902 | 0.033844889308622500 | 0.191807783 |
| | Bet1 | 1.23 | 1.489429499 | 0.032401901784626600 | 0.188385619 |
| | Etv1 | 1.23 | 1.560008667 | 0.027541737394805600 | 0.178595240 |
| | Atr | 1.23 | 1.759834298 | 0.017384640015628500 | 0.156718571 |
| | Mmd2 | 1.23 | 1.396265671 | 0.040154509845963400 | 0.202779537 |
| | Spg21 | 1.23 | 1.47026266 | 0.033863928613865800 | 0.191807783 |
| | Klf6 | 1.23 | 1.777399887 | 0.016695526258944000 | 0.154175555 |
| | Pdgfrb | 1.23 | 1.69411459 | 0.020224854711607400 | 0.160920515 |
| | Gsta4 | 1.23 | 2.244129415 | 0.005699943954794420 | 0.131469154 |
| | Zfp869 | 1.23 | 1.637575723 | 0.023036912702071800 | 0.166831290 |

| | | | | | |
|---|---|---|---|---|---|
| | Med30 | 1.23 | 1.529007974 | 0.029579581550524500 | 0.182155577 |
| | Arsb | 1.23 | 1.945094781 | 0.011347631374846300 | 0.141961618 |
| | Rpl7a | 1.23 | 1.456558478 | 0.034949544683098800 | 0.194290171 |
| | Maged2 | 1.23 | 1.83944999 | 0.014472714984684200 | 0.150441568 |
| | Kpna4 | 1.23 | 1.723202938 | 0.018914595656770700 | 0.160468942 |
| | Ift80 | 1.23 | 1.357636466 | 0.043889793154733600 | 0.210649545 |
| | Zfp120 | 1.23 | 1.510092322 | 0.030896385693476100 | 0.185396043 |
| | Dpy30 | 1.22 | 1.617518348 | 0.024125796018251100 | 0.169658425 |
| | Rpl5 | 1.22 | 1.609566254 | 0.024571617504692400 | 0.169977260 |
| | Clcc1 | 1.22 | 1.922810659 | 0.011945087657149600 | 0.143647440 |
| | Prpf40a | 1.22 | 1.558394897 | 0.027644268521712600 | 0.178595240 |
| | Cep63 | 1.22 | 1.313858907 | 0.048544618615362700 | 0.219352714 |
| | Nkap | 1.22 | 1.341029763 | 0.045600566390007300 | 0.213460217 |
| | Bckdha | 1.22 | 1.755294448 | 0.017567321616927300 | 0.157085313 |
| | Osgep | 1.22 | 1.85438458 | 0.013983484965506900 | 0.148646776 |
| | Eif3l | 1.22 | 2.382499286 | 0.004144772659224810 | 0.131469154 |
| | Rock1 | 1.22 | 1.34714413 | 0.044963061079399000 | 0.211951088 |
| | Stam2 | 1.22 | 1.504924458 | 0.031266231729133800 | 0.185821360 |
| | Sephs1 | 1.22 | 2.067181999 | 0.008566787630867170 | 0.140472827 |
| | Hif1a | 1.22 | 1.779665012 | 0.016608675057430200 | 0.153711652 |
| | Timp3 | 1.22 | 1.413578595 | 0.038585257671378000 | 0.200781458 |
| | Me1 | 1.22 | 1.560154131 | 0.027532514047098500 | 0.178595240 |
| | Ctnnb1 | 1.22 | 1.466016338 | 0.034196657768728100 | 0.192402580 |

| | | | | | |
|---|---|---|---|---|---|
| | Magoh | 1.22 | 1.884997413 | 0.013031745411766800 | 0.146312614 |
| | Nudt10 | 1.22 | 1.442413467 | 0.036106594839175600 | 0.196819835 |
| | Tor1aip2 | 1.22 | 1.47222649 | 0.033711145501326400 | 0.191535824 |
| | Tmem43 | 1.22 | 1.378702673 | 0.041811651993456600 | 0.205721435 |
| | Pigt | 1.22 | 1.712900887 | 0.019368639357499000 | 0.160482390 |
| | Rps14 | 1.22 | 1.561908823 | 0.027421498042819400 | 0.178595240 |
| | C1d | 1.22 | 1.922008234 | 0.011967178410324700 | 0.143647440 |
| | Rnf139 | 1.22 | 2.183101396 | 0.006559920919835590 | 0.133060702 |
| | Etfb | 1.22 | 1.549514205 | 0.028215372928738300 | 0.179474072 |
| | Cntln | 1.22 | 1.313885962 | 0.048541594475048800 | 0.219352714 |
| | Tsen34 | 1.22 | 2.018131768 | 0.009591095866223330 | 0.140783376 |
| | Slc6a9 | 1.22 | 1.41574018 | 0.038393686982639600 | 0.200478413 |
| | Exoc6 | 1.21 | 1.635617934 | 0.023140996974622600 | 0.166831290 |
| | Acp2 | 1.21 | 2.758242952 | 0.001744845781231670 | 0.127020757 |
| | Pxmp4 | 1.21 | 1.61642177 | 0.024186789800637100 | 0.169664195 |
| | Mipep | 1.21 | 2.207946951 | 0.006195167443915790 | 0.131469154 |
| | Rpl19 | 1.21 | 1.438052656 | 0.036470972495157200 | 0.197424556 |
| | Pdcl | 1.21 | 1.799732259 | 0.015858705729945500 | 0.152122332 |
| | Copb2 | 1.21 | 2.096768728 | 0.008002602993635670 | 0.137560781 |
| | 2010315B03Rik | 1.21 | 1.440287026 | 0.036283817513878600 | 0.197299540 |
| | Serpinb6a | 1.21 | 2.194907186 | 0.006383999052480650 | 0.132511376 |
| | Golga7 | 1.21 | 1.859056624 | 0.013833859977646100 | 0.148058275 |
| | Arf4 | 1.21 | 1.546842789 | 0.028389465155540600 | 0.179474072 |

| | | | | | |
|---|---|---|---|---|---|
| | Dazap2 | 1.21 | 1.748978757 | 0.017824659505395300 | 0.157916600 |
| | Btf3 | 1.21 | 1.986281734 | 0.010320916524912700 | 0.141095284 |
| | Slc30a1 | 1.21 | 1.465706247 | 0.034221083255078900 | 0.192402580 |
| | Adam10 | 1.21 | 1.841549868 | 0.014402906200064700 | 0.150441568 |
| | Tmed2 | 1.21 | 2.630402244 | 0.002342058593623310 | 0.127020757 |
| | Degs1 | 1.21 | 1.573112722 | 0.026723127123741100 | 0.176398764 |
| | Qars | 1.21 | 2.922089022 | 0.001196495247616410 | 0.127020757 |
| | Asah1 | 1.21 | 1.35418501 | 0.044239986918664000 | 0.211076965 |
| | Rbm43 | 1.21 | 1.474863909 | 0.033507042092017600 | 0.191129135 |
| | Sf3b6 | 1.21 | 2.152541681 | 0.007038146768937770 | 0.134660365 |
| | 1600012H06Rik | 1.21 | 2.02634915 | 0.009411326711475610 | 0.140783376 |
| | Dcaf13 | 1.21 | 1.954660273 | 0.011100428064870000 | 0.141933868 |
| | Usp1 | 1.21 | 1.561144512 | 0.027469799385033000 | 0.178595240 |
| | Trnau1ap | 1.21 | 1.486828079 | 0.032596571337951700 | 0.188981203 |
| | Mtx1 | 1.21 | 1.863182599 | 0.013703054989918900 | 0.147964655 |
| | Samd8 | 1.21 | 1.55948183 | 0.027575168187060500 | 0.178595240 |
| | Neo1 | 1.21 | 1.703679783 | 0.019784278516586700 | 0.160671218 |
| | Rpl11 | 1.21 | 1.381972474 | 0.041498034333924400 | 0.205392575 |
| | Ankrd28 | 1.21 | 1.641122143 | 0.022849560816612700 | 0.166831290 |
| | Syf2 | 1.20 | 1.957764847 | 0.011021359100077700 | 0.141933868 |
| | Rnf2 | 1.20 | 1.811880466 | 0.015421248450855900 | 0.151749058 |
| | Fmnl2 | 1.20 | 1.350696757 | 0.044596753364814800 | 0.211647507 |
| | Mapk1ip1l | 1.20 | 1.316788522 | 0.048218253741508400 | 0.218577458 |

| | | | | | |
|---|---|---|---|---|---|
| | Tmbim1 | 1.20 | 1.537987707 | 0.028974255990058400 | 0.180151698 |
| | Aco1 | 1.20 | 1.614746967 | 0.024280243236183600 | 0.169664195 |
| | Eif3k | 1.20 | 1.355921442 | 0.044063456088647000 | 0.210746937 |
| | Arl6 | 1.20 | 1.521469098 | 0.030097533191315200 | 0.183250722 |
| | Ppil1 | 1.20 | 1.502690815 | 0.031427452973947100 | 0.186151134 |
| | Actr3 | 1.20 | 2.150174751 | 0.007076609789501350 | 0.134660365 |
| | Alg5 | 1.20 | 1.530327346 | 0.029489856166133400 | 0.181876173 |
| | Ano10 | 1.20 | 2.236607359 | 0.005799527886008470 | 0.131469154 |
| | Ufl1 | 1.20 | 1.561402361 | 0.027453494916105900 | 0.178595240 |
| | Erlin1 | 1.20 | 1.510184461 | 0.030889831509696900 | 0.185396043 |
| | Oscp1 | 1.20 | 1.916637013 | 0.012116103835770000 | 0.143647440 |
| | Entpd5 | 1.20 | 1.327386659 | 0.047055819461626700 | 0.216606590 |
| | Mettl14 | 1.20 | 1.316799504 | 0.048217034441439500 | 0.218577458 |
| | Tmem68 | 1.20 | 1.379120793 | 0.041771416886409600 | 0.205721435 |
| | Dscr3 | 1.20 | 1.926801068 | 0.011835835802641500 | 0.143647440 |
| | Mapre1 | 1.20 | 1.850254738 | 0.014117092558630500 | 0.149025511 |
| | Fdps | 1.20 | 1.366354654 | 0.043017517683528500 | 0.208649386 |
| | Cyfip1 | 1.20 | 1.440351348 | 0.036278444022239100 | 0.197299540 |
| | Hmgcl | 1.20 | 1.354657715 | 0.044191860356494200 | 0.210920581 |
| | Snrpd1 | 1.20 | 2.007751363 | 0.009823101615772120 | 0.140783376 |
| | Orai3 | 1.20 | 1.557972799 | 0.027671149536016000 | 0.178595240 |
| | Zfp943 | 1.20 | 1.620359962 | 0.023968454847959700 | 0.168887506 |
| | Ssbp1 | 1.20 | 1.336273 | 0.046102767865708000 | 0.214668636 |

| | | | | | |
|---|---|---|---|---|---|
| | Suco | 1.20 | 1.407567442 | 0.039123036746417900 | 0.201809641 |
| | Wtap | 1.20 | 1.957731865 | 0.011022196129412300 | 0.141933868 |
| | Tmed5 | 1.20 | 1.51293151 | 0.030695060247092000 | 0.184991589 |
| | Chmp5 | 1.20 | 1.8980458 | 0.012646029766589600 | 0.145739007 |
| | Hsd17b11 | 1.20 | 1.426780965 | 0.037429931756165300 | 0.199280377 |
| | Dcaf17 | 1.20 | 1.482514333 | 0.032921958797739000 | 0.189849178 |
| | Ptgr2 | 1.20 | 1.590314623 | 0.025685343421657400 | 0.173610331 |
| | Rps23 | 1.20 | 1.307988533 | 0.049205252711134900 | 0.221101824 |
| | Zfp26 | 1.20 | 1.703749264 | 0.019781113570448600 | 0.160671218 |
| | 9530068E07 Rik | 1.20 | 1.417431234 | 0.038244480515414300 | 0.200458830 |
| | Rhob | 1.19 | 1.628369177 | 0.023530481962271000 | 0.167968850 |
| | Cd200 | 1.19 | 1.670164885 | 0.021371505427202700 | 0.163057111 |
| | Brat1 | 1.19 | 1.398329688 | 0.039964125276280300 | 0.202412377 |
| | Fbxo8 | 1.19 | 1.519677937 | 0.030221920756819700 | 0.183617574 |
| | Rps17 | 1.19 | 1.816419608 | 0.015260908626332200 | 0.151522906 |
| | Rpl17 | 1.19 | 1.389505938 | 0.040784398561691000 | 0.203788517 |
| | Haus2 | 1.19 | 1.608088186 | 0.024655386434641300 | 0.170091091 |
| | Stard3nl | 1.19 | 1.829354361 | 0.014813089218021100 | 0.151437765 |
| | Atp1b3 | 1.19 | 2.11984467 | 0.007588489367274580 | 0.136287102 |
| | Txndc12 | 1.19 | 1.484675829 | 0.032758512374353100 | 0.189525776 |
| | Rbm45 | 1.19 | 1.947959159 | 0.011273034627865500 | 0.141933868 |
| | Rpl3 | 1.19 | 1.697575824 | 0.020064307565449900 | 0.160920515 |
| | Ube2e3 | 1.19 | 1.446871639 | 0.035737845027291100 | 0.196291813 |

| | | | | | |
|---|---|---|---|---|---|
| | Rdx | 1.19 | 1.560799991 | 0.027491599523638900 | 0.178595240 |
| | Klhl7 | 1.19 | 1.405712272 | 0.039290515594984600 | 0.201979414 |
| | Commd3 | 1.19 | 2.082842613 | 0.008263373574138730 | 0.139726969 |
| | Rnf128 | 1.19 | 1.639707938 | 0.022924087723046600 | 0.166831290 |
| | Rpl12 | 1.19 | 1.988636071 | 0.010265117598770500 | 0.141095284 |
| | Mcfd2 | 1.19 | 1.511435449 | 0.030800981140492800 | 0.185304852 |
| | Cc2d2a | 1.19 | 1.626548748 | 0.023629321510954700 | 0.168059269 |
| | Mydgf | 1.19 | 1.426689505 | 0.037437815059851800 | 0.199280377 |
| | Fam149a | 1.19 | 1.352867316 | 0.044374419378754700 | 0.211311132 |
| | Arxes2 | 1.19 | 1.663946377 | 0.021679717726085100 | 0.164132783 |
| | Atraid | 1.19 | 1.736588703 | 0.018340505329556800 | 0.159370958 |
| | Rpl21 | 1.19 | 1.392204465 | 0.040531766731611900 | 0.203309852 |
| | Sdcbp | 1.19 | 1.889958691 | 0.012883720930230000 | 0.146280491 |
| | Igbp1 | 1.19 | 1.688808024 | 0.020473494489962400 | 0.161588907 |
| | Spg11 | 1.19 | 1.403984101 | 0.039447174254887700 | 0.202168336 |
| | Fkbp15 | 1.19 | 1.438232552 | 0.036455868404179200 | 0.197424556 |
| | Rps29 | 1.19 | 1.398253358 | 0.039971149844679100 | 0.202412377 |
| | Tmed7 | 1.19 | 1.594494861 | 0.025439298947895500 | 0.172537713 |
| | Mrpl39 | 1.19 | 2.311862211 | 0.004876831931466530 | 0.131469154 |
| | Emc2 | 1.19 | 1.95618934 | 0.011061414325648000 | 0.141933868 |
| | Abhd17c | 1.19 | 1.574784526 | 0.026620454985454300 | 0.176115393 |
| | Ptp4a2 | 1.19 | 1.693089478 | 0.020272649979130600 | 0.161021089 |
| | Sugt1 | 1.19 | 2.041174968 | 0.009095467612568800 | 0.140488765 |

| | | | | | |
|---|---|---|---|---|---|
| | Psmf1 | 1.19 | 2.163828952 | 0.006857582601545330 | 0.134513024 |
| | Birc2 | 1.19 | 1.503647026 | 0.031358333463593600 | 0.186051321 |
| | Ppt1 | 1.19 | 1.872012048 | 0.013427277118264000 | 0.147515895 |
| | Wdr36 | 1.18 | 1.418683517 | 0.038134361845481900 | 0.200173662 |
| | Slc33a1 | 1.18 | 1.333317806 | 0.046417547871888100 | 0.215117546 |
| | Fuca2 | 1.18 | 1.595650318 | 0.025371706704654800 | 0.172334465 |
| | Hmgb1 | 1.18 | 1.430475152 | 0.037112896307863700 | 0.198740572 |
| | Acot9 | 1.18 | 1.66227264 | 0.021763430865299100 | 0.164404235 |
| | Yipf1 | 1.18 | 1.709069128 | 0.019540284018734700 | 0.160482390 |
| | Phf14 | 1.18 | 1.432712995 | 0.036922151891048800 | 0.198531875 |
| | Adnp2 | 1.18 | 1.660330019 | 0.021860997818428800 | 0.164554124 |
| | Yipf5 | 1.18 | 2.034171929 | 0.009243321760651820 | 0.140488765 |
| | Usp25 | 1.18 | 2.248735258 | 0.005639813491301470 | 0.131469154 |
| | Zfp617 | 1.18 | 1.630848583 | 0.023396528203028200 | 0.167793160 |
| | Smpd2 | 1.18 | 1.334805327 | 0.046258833016900700 | 0.214941372 |
| | Stxbp4 | 1.18 | 1.31173001 | 0.048783166867673500 | 0.219852245 |
| | Gemin8 | 1.18 | 1.319753591 | 0.047890173393266400 | 0.218121338 |
| | Tusc3 | 1.18 | 1.682087036 | 0.020792799406624600 | 0.162153154 |
| | Sec23b | 1.18 | 1.700009048 | 0.019952207480719600 | 0.160872834 |
| | Nasp | 1.18 | 1.338944274 | 0.045820067694420200 | 0.213863298 |
| | Chid1 | 1.18 | 1.708169508 | 0.019580802720094600 | 0.160482390 |
| | Timm22 | 1.18 | 1.305666425 | 0.049469050586014300 | 0.221308908 |
| | Tceal8 | 1.18 | 1.478287671 | 0.033243927710268100 | 0.190971911 |

| | | | | | |
|---|---|---|---|---|---|
| | Slc35a2 | 1.18 | 1.824916168 | 0.014965245043491100 | 0.151437765 |
| | Rnf181 | 1.18 | 1.980166348 | 0.010467275434961300 | 0.141095284 |
| | Slc20a2 | 1.18 | 1.692216073 | 0.020313461102313600 | 0.161158931 |
| | Bhlhb9 | 1.18 | 1.384806914 | 0.041228077701736500 | 0.204948695 |
| | Dhx36 | 1.18 | 1.974092775 | 0.010614687795115100 | 0.141480889 |
| | Spcs1 | 1.18 | 1.806523988 | 0.015612627985810300 | 0.151883712 |
| | Cspg5 | 1.18 | 1.351180437 | 0.044547112906402300 | 0.211585183 |
| | Pdcd10 | 1.18 | 1.403037537 | 0.039533244946565200 | 0.202168336 |
| | Snrpe | 1.18 | 1.34671675 | 0.045007330069822500 | 0.212087010 |
| | Efcab14 | 1.18 | 1.476402368 | 0.033388555615237500 | 0.191129135 |
| | C330007P06Rik | 1.18 | 1.601194493 | 0.025049871783272800 | 0.171163743 |
| | Nek9 | 1.18 | 1.376276407 | 0.042045894226596300 | 0.206192945 |
| | Mrpl3 | 1.17 | 2.037910463 | 0.009164094028462660 | 0.140488765 |
| | Nlgn1 | 1.17 | 1.521800998 | 0.030074540574743500 | 0.183250722 |
| | Sqle | 1.17 | 1.311470598 | 0.048812314592214600 | 0.219911480 |
| | Ankra2 | 1.17 | 1.857197039 | 0.013893221533105000 | 0.148565570 |
| | Cndp2 | 1.17 | 1.405161886 | 0.039340340455366000 | 0.202034437 |
| | Txndc17 | 1.17 | 1.623943372 | 0.023771502241605000 | 0.168547065 |
| | Rpl7l1 | 1.17 | 1.60987066 | 0.024554400743913500 | 0.169975829 |
| | Wdr44 | 1.17 | 1.561072872 | 0.027474331099505300 | 0.178595240 |
| | Gpkow | 1.17 | 1.630071748 | 0.023438415679963600 | 0.167830782 |
| | Use1 | 1.17 | 1.49766759 | 0.031793065918225200 | 0.187015633 |
| | Rpl37a | 1.17 | 1.402812323 | 0.039553751119558600 | 0.202197952 |

| | | | | | |
|---|---|---|---|---|---|
| | Fam213a | 1.17 | 1.956141307 | 0.011062637794281900 | 0.141933868 |
| | Commd7 | 1.17 | 1.361092155 | 0.043541946989400800 | 0.209862494 |
| | Kdelr1 | 1.17 | 1.55018496 | 0.028171828757727100 | 0.179474072 |
| | Nme3 | 1.17 | 1.321580684 | 0.047689120854600100 | 0.217851134 |
| | Capns1 | 1.17 | 1.636484224 | 0.023094883508547800 | 0.166831290 |
| | BC004004 | 1.17 | 2.101494809 | 0.007915989167151340 | 0.137039524 |
| | Nras | 1.17 | 1.443318753 | 0.036031409145722200 | 0.196819835 |
| | Bloc1s2 | 1.17 | 1.614773407 | 0.024278765077261700 | 0.169664195 |
| | Prkaa1 | 1.17 | 1.368515016 | 0.042804061938535800 | 0.207981123 |
| | Rae1 | 1.17 | 1.31976731 | 0.047888660513927400 | 0.218121338 |
| | Id2 | 1.17 | 1.84037437 | 0.014441943107636900 | 0.150441568 |
| | Glod4 | 1.17 | 1.707281673 | 0.019620872999466200 | 0.160482390 |
| | Taf9 | 1.17 | 1.993870746 | 0.010142131888749100 | 0.141095284 |
| | Ndufc2 | 1.17 | 1.37117621 | 0.042542576696440900 | 0.207659443 |
| | Sfr1 | 1.17 | 1.772571344 | 0.016882185023473200 | 0.154858548 |
| | Tsn | 1.16 | 1.531638523 | 0.029400957692795900 | 0.181444365 |
| | Cetn3 | 1.16 | 1.991489541 | 0.010197893186800200 | 0.141095284 |
| | Tmem50a | 1.16 | 1.572611364 | 0.026753994631071800 | 0.176398764 |
| | Gabra2 | 1.16 | 1.684940409 | 0.020656635752000500 | 0.161859432 |
| | Impad1 | 1.16 | 1.707503565 | 0.019610850760578800 | 0.160482390 |
| | Lin7c | 1.16 | 1.421824413 | 0.037859562189037500 | 0.199763957 |
| | Tril | 1.16 | 1.333820124 | 0.046363891011530000 | 0.214941372 |
| | Pcm1 | 1.16 | 1.902890238 | 0.012505750568373200 | 0.145640913 |

| | | | | | |
|---|---|---|---|---|---|
| | Dnttip2 | 1.16 | 1.633156147 | 0.023272543622721000 | 0.167340671 |
| | Psmd13 | 1.16 | 2.30134558 | 0.004996368023568030 | 0.131469154 |
| | Eif3h | 1.16 | 1.401699953 | 0.039655191066835200 | 0.202412377 |
| | Scfd2 | 1.16 | 1.321026044 | 0.047750063756135800 | 0.217912197 |
| | Fam45a | 1.16 | 2.25640351 | 0.005541106410596000 | 0.131469154 |
| | Sar1b | 1.16 | 1.414137187 | 0.038535660996442500 | 0.200781458 |
| | Psma4 | 1.16 | 1.689802191 | 0.020426681123782600 | 0.161497713 |
| | Tmem237 | 1.16 | 1.382922459 | 0.041407359918475100 | 0.205392575 |
| | Smarcb1 | 1.16 | 1.428864989 | 0.037250749166353000 | 0.198960061 |
| | Tgoln1 | 1.16 | 1.512290612 | 0.030740391079643400 | 0.185102416 |
| | Acbd3 | 1.16 | 1.369324353 | 0.042724367986678300 | 0.207844103 |
| | Zdhhc20 | 1.16 | 1.387264238 | 0.040995459811618500 | 0.204067695 |
| | 43166 | 1.16 | 1.399748801 | 0.039833750555725600 | 0.202412377 |
| | Pbrm1 | 1.16 | 1.353894834 | 0.044269555924437100 | 0.211144730 |
| | Stag2 | 1.16 | 1.35353342 | 0.044306411858682600 | 0.211173918 |
| | Eif2s1 | 1.16 | 1.998621898 | 0.010031782364196900 | 0.141095284 |
| | Gosr2 | 1.15 | 1.840355501 | 0.014442570585702000 | 0.150441568 |
| | Dnajc10 | 1.15 | 1.34463067 | 0.045224037183802100 | 0.212626274 |
| | Pcnx4 | 1.15 | 1.522590078 | 0.030019947033815200 | 0.183250722 |
| | Nxt2 | 1.15 | 1.304327746 | 0.049621770328923000 | 0.221308908 |
| | Gmfb | 1.15 | 1.774072249 | 0.016823941542847800 | 0.154749105 |
| | Nosip | 1.15 | 2.123548259 | 0.007524051174037540 | 0.136287102 |
| | Rap2b | 1.15 | 1.542732679 | 0.028659415001431700 | 0.179884640 |

| | | | | | |
|---|---|---|---|---|---|
| | Mettl9 | 1.15 | 1.305612883 | 0.049475149717507000 | 0.221308908 |
| | AI597479 | 1.15 | 1.611276108 | 0.024475067123643200 | 0.169826196 |
| | Gtf2b | 1.15 | 1.500676198 | 0.031573578173935800 | 0.186633806 |
| | Hibadh | 1.15 | 1.314690238 | 0.048451782832886000 | 0.219149423 |
| | Lsm14a | 1.15 | 1.486668176 | 0.032608575245127700 | 0.188981203 |
| | Pccb | 1.15 | 1.834608493 | 0.014634958899744900 | 0.151070715 |
| | Snx2 | 1.15 | 1.875236672 | 0.013327949179867500 | 0.146863953 |
| | Pex3 | 1.15 | 1.330662841 | 0.046702180629445300 | 0.215957866 |
| | Tubb4b | 1.15 | 1.404322967 | 0.039416406896208800 | 0.202168336 |
| | Saraf | 1.15 | 2.188379011 | 0.006480686141907270 | 0.133060702 |
| | Crls1 | 1.15 | 1.539914186 | 0.028846014237661600 | 0.180005941 |
| | Snx1 | 1.15 | 1.304075121 | 0.049650643159236300 | 0.221317530 |
| | Tcea1 | 1.15 | 2.117322246 | 0.007632692294018410 | 0.136287102 |
| | Cln8 | 1.15 | 1.496457716 | 0.031881759697221700 | 0.187100141 |
| | Ntan1 | 1.15 | 2.182098756 | 0.006575083071302350 | 0.133060702 |
| | Rtcb | 1.15 | 1.777532397 | 0.016690432987203500 | 0.154175555 |
| | Fech | 1.15 | 1.546823141 | 0.028390749596033700 | 0.179474072 |
| | Cept1 | 1.15 | 1.794157952 | 0.016063569187931900 | 0.152122332 |
| | Cog2 | 1.15 | 1.303328857 | 0.049736033045790900 | 0.221444337 |
| | Trim23 | 1.15 | 1.539187619 | 0.028894313513695600 | 0.180105205 |
| | Stx7 | 1.15 | 1.926456689 | 0.011845224910228400 | 0.143647440 |
| | Atp5g2 | 1.15 | 1.486382612 | 0.032630023584010500 | 0.189025782 |
| | Hnrnpk | 1.15 | 1.715076557 | 0.019271851616276500 | 0.160482390 |

| | | | | | |
|---|---|---|---|---|---|
| | Ift122 | 1.15 | 1.412763943 | 0.038657704058048300 | 0.200960331 |
| | Cct4 | 1.15 | 1.537086012 | 0.029034475721049000 | 0.180362898 |
| | Ubxn1 | 1.15 | 1.548911382 | 0.028254564513673600 | 0.179474072 |
| | Ncoa4 | 1.15 | 1.372314895 | 0.042431179602263000 | 0.207489978 |
| | Rlim | 1.15 | 1.424285317 | 0.037645639878262700 | 0.199593499 |
| | Gdi2 | 1.14 | 1.988829871 | 0.010260537891966100 | 0.141095284 |
| | Oaz1 | 1.14 | 1.637709081 | 0.023029839893345200 | 0.166831290 |
| | Eif2s2 | 1.14 | 2.203070222 | 0.006265125543586840 | 0.132069644 |
| | Bub3 | 1.14 | 1.510082397 | 0.030897091794664800 | 0.185396043 |
| | Tm9sf2 | 1.14 | 1.636655921 | 0.023085754819587400 | 0.166831290 |
| | Bzw1 | 1.14 | 2.215110533 | 0.006093817820723810 | 0.131469154 |
| | Vps28 | 1.14 | 1.442763968 | 0.036077466499444300 | 0.196819835 |
| | Cacul1 | 1.14 | 2.127789185 | 0.007450935683738940 | 0.136287102 |
| | Zzz3 | 1.14 | 1.79630256 | 0.015984440509615900 | 0.152122332 |
| | Bnip3l | 1.14 | 1.711478596 | 0.019432174614701000 | 0.160482390 |
| | Psma1 | 1.14 | 1.527663932 | 0.029671265416351000 | 0.182340276 |
| | Cnpy3 | 1.14 | 1.414760668 | 0.038480378226402200 | 0.200706096 |
| | Nsdhl | 1.14 | 1.513527093 | 0.030652994475696700 | 0.184968428 |
| | Psma6 | 1.14 | 1.624265474 | 0.023753878231221500 | 0.168509056 |
| | Rsl1d1 | 1.14 | 1.657712694 | 0.021993143397843400 | 0.164780689 |
| | Thtpa | 1.14 | 1.572058049 | 0.026788102455662400 | 0.176398764 |
| | Ndufs5 | 1.14 | 1.324783361 | 0.047338733976945700 | 0.217172036 |
| | Ttc5 | 1.13 | 1.469286008 | 0.033940168321149200 | 0.191843625 |

| | | | | | |
|---|---|---|---|---|---|
| | Cdc42se2 | 1.13 | 1.387902855 | 0.040935221553945600 | 0.204032563 |
| | Ipo11 | 1.13 | 1.678155081 | 0.020981905131964600 | 0.162610003 |
| | Tceal9 | 1.13 | 1.464141339 | 0.034344615708745500 | 0.192834605 |
| | Arl6ip1 | 1.13 | 1.817666085 | 0.015217170786449500 | 0.151522906 |
| | Anapc1 | 1.13 | 1.387177673 | 0.041003631984508000 | 0.204067695 |
| | Vps45 | 1.13 | 1.501875845 | 0.031486483146281000 | 0.186409205 |
| | Gstz1 | 1.13 | 1.45853174 | 0.034791107989715500 | 0.193941020 |
| | Cbx3 | 1.13 | 1.627490434 | 0.023578141308622400 | 0.167979916 |
| | Setd2 | 1.13 | 1.521723877 | 0.030079881636778200 | 0.183250722 |
| | Fbxo25 | 1.13 | 1.507305979 | 0.031095247741138600 | 0.185612424 |
| | Washc5 | 1.13 | 1.674899 | 0.021139806113392800 | 0.162735056 |
| | Cops5 | 1.13 | 1.809747732 | 0.015497165383878900 | 0.151749058 |
| | Exoc2 | 1.13 | 1.390193168 | 0.040719912073427500 | 0.203540310 |
| | Mcrip1 | 1.12 | 1.531554637 | 0.029406637156338500 | 0.181444365 |
| | Wwc2 | 1.12 | 1.367654164 | 0.042888991662284900 | 0.208320125 |
| | Rab18 | 1.12 | 1.319647277 | 0.047901898190053600 | 0.218121338 |
| | Matr3 | 1.12 | 1.413526336 | 0.038589900982334000 | 0.200781458 |
| | Dirc2 | 1.12 | 1.76220873 | 0.017289851790069000 | 0.156718571 |
| | Capza2 | 1.12 | 1.41623429 | 0.038350030166280300 | 0.200478413 |
| | Ccz1 | 1.12 | 1.561579017 | 0.027442330061390000 | 0.178595240 |
| | Sar1a | 1.12 | 1.614837871 | 0.024275161546639000 | 0.169664195 |
| | Vps35 | 1.12 | 1.316540559 | 0.048245792137679100 | 0.218577458 |
| | Ssb | 1.12 | 1.392640989 | 0.040491047407822800 | 0.203298902 |

| | | | | | |
|---|---|---|---|---|---|
| | Ndufb11 | 1.12 | 1.676062835 | 0.021083230906765700 | 0.162610003 |
| | Nt5c3b | 1.12 | 1.364262708 | 0.043225227917226500 | 0.209003820 |
| | Cdc42 | 1.12 | 1.625869745 | 0.023666294005236600 | 0.168197479 |
| | Bpgm | 1.12 | 1.378855334 | 0.041796957146592800 | 0.205721435 |
| | Psmd8 | 1.12 | 1.379400902 | 0.041744484086536500 | 0.205721435 |
| | Ubr7 | 1.12 | 1.597480505 | 0.025265011220688500 | 0.171864613 |
| | Vps4b | 1.11 | 1.464892973 | 0.034285226785602400 | 0.192684377 |
| | Rab5a | 1.11 | 1.357247463 | 0.043929123428975500 | 0.210667443 |
| | Rpl9 | 1.11 | 1.423261592 | 0.037734483318321700 | 0.199593499 |
| | Tmem9b | 1.11 | 1.357865912 | 0.043866611501608800 | 0.210611848 |
| | Psma3 | 1.10 | 1.345612166 | 0.045121947330988500 | 0.212408591 |
| | Rbbp7 | 1.10 | 1.546360585 | 0.028421003926886200 | 0.179474072 |
| | Ndufa3 | 1.10 | 1.378729106 | 0.041809107213129800 | 0.205721435 |
| | Arf2 | 1.10 | 1.397905689 | 0.040003161112672400 | 0.202460860 |
| | Tsg101 | 1.10 | 1.327945191 | 0.046995341379663000 | 0.216553651 |
| | Dstn | 1.10 | 1.323899994 | 0.047435120245440400 | 0.217289839 |
| | Ptdss1 | 1.10 | 1.32912776 | 0.046867548771447500 | 0.216327507 |

| DOWN p<0.05 | Gene ID | Fold change (BRAF V600E vs. BRAFwt) | log-10 P-values | P-value | FDR step up |
|---|---|---|---|---|---|
| | Ube2v1 | -1.09 | 1.405726156 | 0.039289259567660300 | 0.20197941444 |
| | Chd8 | -1.09 | 1.424512806 | 0.037625925755101700 | 0.19959349860 |
| | Pi4k2a | -1.10 | 1.357248753 | 0.043928992865401100 | 0.21066744315 |
| | Hbs1l | -1.10 | 1.372853286 | 0.042378610626430900 | 0.20740196527 |
| | Ncaph2 | -1.10 | 1.380371057 | 0.041651336667437300 | 0.20572143528 |
| | Gpat4 | -1.10 | 1.403167051 | 0.039521457173240600 | 0.20216833599 |
| | Kat2a | -1.10 | 1.356853957 | 0.043968944757966200 | 0.21066744315 |
| | Ankrd17 | -1.10 | 1.450162088 | 0.035468099036610400 | 0.19580841658 |
| | Eif1b | -1.10 | 1.320709924 | 0.047784833379982500 | 0.21799847127 |
| | Rrp7a | -1.10 | 1.436963744 | 0.036562531360266600 | 0.19756304526 |
| | Psmc5 | -1.10 | 1.339532393 | 0.045758060401404900 | 0.21379174022 |
| | Ppp2r5c | -1.10 | 1.370010588 | 0.042656911888136500 | 0.20784410277 |
| | Pdpk1 | -1.11 | 1.41258973 | 0.038673214313953900 | 0.20096033149 |
| | Cyth3 | -1.11 | 1.400519082 | 0.039763162484172300 | 0.20241237663 |
| | Ppm1f | -1.11 | 1.482485575 | 0.032924138854330500 | 0.18984917835 |
| | Pip4k2b | -1.11 | 1.603049019 | 0.024943131751707400 | 0.17077407743 |
| | Dlg3 | -1.11 | 1.365899121 | 0.043062662615891300 | 0.20870544595 |
| | Smarca4 | -1.11 | 1.636153631 | 0.023112470448338900 | 0.16683129036 |
| | Dnajc5 | -1.11 | 1.519532591 | 0.030232036873770600 | 0.18361757429 |
| | Ogt | -1.11 | 1.324132687 | 0.047409711545090300 | 0.21728983949 |
| | Lrrc59 | -1.11 | 2.013783699 | 0.009687602292284840 | 0.14078337569 |

| | | | | | |
|---|---|---|---|---|---|
| | Timm44 | -1.11 | 1.334001834 | 0.046344496247540800 | 0.21494137193 |
| | Usp7 | -1.11 | 1.723788123 | 0.018889126584700100 | 0.16046894179 |
| | Btbd2 | -1.11 | 1.721803461 | 0.018975644674095700 | 0.16048070792 |
| | Trappc10 | -1.11 | 1.574991378 | 0.026607778826232800 | 0.17611539333 |
| | Mprip | -1.11 | 1.684800944 | 0.020663270262777900 | 0.16185943212 |
| | D5Ertd579e | -1.11 | 1.364466756 | 0.043204923886896800 | 0.20900381980 |
| | Rnf187 | -1.11 | 1.467888153 | 0.034049586862387700 | 0.19202608564 |
| | Foxk2 | -1.11 | 1.303747356 | 0.049688129061513000 | 0.22131753045 |
| | Tmem57 | -1.12 | 1.353684647 | 0.044290986441094400 | 0.21117364493 |
| | Actl6b | -1.12 | 1.415916397 | 0.038378111728084900 | 0.20047841336 |
| | Tnpo2 | -1.12 | 1.333947693 | 0.046350274141272500 | 0.21494137193 |
| | Med14 | -1.12 | 1.380265505 | 0.041661460942205500 | 0.20572143528 |
| | Snx32 | -1.12 | 1.395814919 | 0.040196207676468800 | 0.20284101714 |
| | Mfsd6 | -1.12 | 1.623484536 | 0.023796630347461600 | 0.16855128742 |
| | Ncoa1 | -1.12 | 1.380446441 | 0.041644107493250500 | 0.20572143528 |
| | Wipi2 | -1.12 | 1.813406121 | 0.015367169398761300 | 0.15174905791 |
| | Slc45a4 | -1.12 | 1.606821402 | 0.024727408212638200 | 0.17028275660 |
| | Usf3 | -1.12 | 1.347654409 | 0.044910262196024500 | 0.21195108826 |
| | Tollip | -1.12 | 1.682481168 | 0.020773938063151800 | 0.16209805958 |
| | Cluh | -1.12 | 1.748334072 | 0.017851138850627600 | 0.15804929056 |
| | Iars | -1.12 | 1.553645354 | 0.027948251695490900 | 0.17888249734 |
| | Copg1 | -1.12 | 1.473907152 | 0.033580939917960000 | 0.19130833143 |
| | Arid1b | -1.12 | 1.856307959 | 0.013921692633134700 | 0.14862753048 |

| | | | | | |
|---|---|---|---|---|---|
| | Myh10 | -1.12 | 1.375135696 | 0.042156476397891800 | 0.20658778252 |
| | Creld1 | -1.12 | 1.325877434 | 0.047219628517406600 | 0.21706378366 |
| | Sgip1 | -1.13 | 1.361229299 | 0.043528199214699800 | 0.20986249400 |
| | Cdk11b | -1.13 | 1.482301473 | 0.032938098737095400 | 0.18985000619 |
| | Map2k7 | -1.13 | 1.688082837 | 0.020507709789969900 | 0.16167323019 |
| | Mcoln1 | -1.13 | 1.343345801 | 0.045358031606124900 | 0.21288007959 |
| | Pik3ca | -1.13 | 1.365418425 | 0.043110352577721600 | 0.20875838147 |
| | Zfp445 | -1.13 | 1.718016555 | 0.019141829582180100 | 0.16048070792 |
| | Zscan26 | -1.13 | 1.696540984 | 0.020112173910272200 | 0.16092051453 |
| | Emc1 | -1.13 | 2.009918739 | 0.009774200887360300 | 0.14078337569 |
| | Ndufb10 | -1.13 | 1.32725687 | 0.047069884218092200 | 0.21660659044 |
| | Vwa8 | -1.13 | 1.439064108 | 0.036386132085534200 | 0.19742455646 |
| | Dip2b | -1.13 | 1.664863499 | 0.021633983819370600 | 0.16399478955 |
| | Synj1 | -1.13 | 1.372991452 | 0.042365130467563500 | 0.20740196527 |
| | Nmnat2 | -1.13 | 2.250709897 | 0.005614228738761430 | 0.13146915442 |
| | Fam222b | -1.13 | 1.334595653 | 0.046281171864919900 | 0.21494137193 |
| | Mapre2 | -1.13 | 1.645722202 | 0.022608814882883200 | 0.16645864635 |
| | Rprd2 | -1.13 | 1.458898285 | 0.034761756631172000 | 0.19385604621 |
| | Sirt3 | -1.13 | 1.467274757 | 0.034097712366682400 | 0.19210195393 |
| | Camsap2 | -1.13 | 1.568978553 | 0.026978726616509300 | 0.17678334880 |
| | Epg5 | -1.13 | 1.625735952 | 0.023673585962435100 | 0.16819747870 |
| | Camsap1 | -1.13 | 2.026385121 | 0.009410547242779680 | 0.14078337569 |
| | Man2a2 | -1.13 | 1.806364974 | 0.015618345489080400 | 0.15188371222 |

| | | | | | |
|---|---|---|---|---|---|
| | Tef | -1.13 | 1.557714748 | 0.027687596170035600 | 0.17861749248 |
| | Slc27a4 | -1.13 | 1.351034561 | 0.044562078470757500 | 0.21158518323 |
| | Ylpm1 | -1.13 | 1.32328698 | 0.047502123066772500 | 0.21743060395 |
| | Rubcn | -1.13 | 1.867820545 | 0.013557495077323100 | 0.14796465545 |
| | Fkbp4 | -1.13 | 1.474165922 | 0.033560937000111100 | 0.19130833143 |
| | Chp1 | -1.14 | 1.737439298 | 0.018304619343396600 | 0.15929308068 |
| | Egln2 | -1.14 | 1.91690943 | 0.012108506236541700 | 0.14364743987 |
| | Rnf10 | -1.14 | 1.943674791 | 0.011384794850249300 | 0.14196161768 |
| | Zfyve27 | -1.14 | 1.322539012 | 0.047584004481083000 | 0.21755514385 |
| | Tspyl2 | -1.14 | 1.313685402 | 0.048564016539335300 | 0.21936822856 |
| | Tatdn2 | -1.14 | 1.637857884 | 0.023021950520267700 | 0.16683129036 |
| | Tmem189 | -1.14 | 1.468465048 | 0.034004387124995300 | 0.19202608564 |
| | Pde4dip | -1.14 | 1.701968143 | 0.019862406114743000 | 0.16073576114 |
| | Kansl3 | -1.14 | 1.401575263 | 0.039666578118536200 | 0.20241237663 |
| | Cdkl2 | -1.14 | 1.309624875 | 0.049020205174829700 | 0.22050961871 |
| | Mapre3 | -1.14 | 1.379512244 | 0.041733783228350200 | 0.20572143528 |
| | Rps6kc1 | -1.14 | 1.738511288 | 0.018259492898146400 | 0.15920485681 |
| | Setd3 | -1.14 | 1.510271087 | 0.030883670695882200 | 0.18539604295 |
| | Stx16 | -1.14 | 1.345239763 | 0.045160655536196700 | 0.21251800264 |
| | Nfe2l1 | -1.14 | 1.936005964 | 0.011587614440075300 | 0.14293124777 |
| | Cyth2 | -1.14 | 1.455776322 | 0.035012544844164200 | 0.19429017089 |
| | Tnfrsf21 | -1.14 | 1.675260799 | 0.021122202465399500 | 0.16273505647 |
| | Acap2 | -1.14 | 1.807387978 | 0.015581598933935400 | 0.15184875954 |

| | | | | | |
|---|---|---|---|---|---|
| | Bcas3 | -1.14 | 1.675963383 | 0.021088059418769400 | 0.16261000251 |
| | Afg3l2 | -1.14 | 2.044369422 | 0.009028811337654220 | 0.14048876521 |
| | Letm1 | -1.14 | 1.831863633 | 0.014727748772978700 | 0.15143776501 |
| | Pi4ka | -1.14 | 1.355513785 | 0.044104836327762800 | 0.21075109435 |
| | Tecpr1 | -1.14 | 1.513032717 | 0.030687907958367000 | 0.18499158897 |
| | Adcy3 | -1.14 | 1.498837683 | 0.031707523047650500 | 0.18676431809 |
| | Tcf25 | -1.14 | 1.317957722 | 0.048088616011824400 | 0.21839935441 |
| | Clip3 | -1.14 | 1.586584468 | 0.025906904942552900 | 0.17426163846 |
| | Plcg1 | -1.14 | 1.723576332 | 0.018898340413793500 | 0.16046894179 |
| | Amigo1 | -1.14 | 1.397034696 | 0.040083469320473500 | 0.20264420601 |
| | Trim8 | -1.14 | 1.579754587 | 0.026317547373463700 | 0.17487839220 |
| | Mark2 | -1.14 | 1.649038581 | 0.022436825962097900 | 0.16638123343 |
| | Arfgef2 | -1.14 | 1.635736616 | 0.023134674003866700 | 0.16683129036 |
| | Zzef1 | -1.14 | 2.162089201 | 0.006885108659860510 | 0.13451302432 |
| | Rock2 | -1.14 | 1.430673118 | 0.037095982868622800 | 0.19873305304 |
| | Vamp2 | -1.14 | 1.54164812 | 0.028731075297165200 | 0.17993235070 |
| | Myt1l | -1.14 | 1.92205108 | 0.011965997847135400 | 0.14364743987 |
| | Nfyb | -1.14 | 1.454244352 | 0.035136269412593900 | 0.19452356084 |
| | Slc25a23 | -1.14 | 1.401783359 | 0.039647576063449500 | 0.20241237663 |
| | Wasl | -1.14 | 1.859398939 | 0.013822960302833800 | 0.14805827509 |
| | Stxbp5 | -1.14 | 1.387847495 | 0.040940439959070400 | 0.20403256309 |
| | Gas6 | -1.14 | 1.926924782 | 0.011832464716998800 | 0.14364743987 |
| | Slc25a44 | -1.14 | 1.768979757 | 0.017022378504236100 | 0.15563363212 |

| | | | | | |
|---|---|---|---|---|---|
| | Dnmt3a | -1.14 | 1.539551469 | 0.028870116165828600 | 0.18007456479 |
| | Ash1l | -1.14 | 1.423232802 | 0.037736984889085100 | 0.19959349860 |
| | 3-Sep | -1.15 | 1.441349703 | 0.036195143008407300 | 0.19708779001 |
| | Trrap | -1.15 | 1.417360462 | 0.038250713280869200 | 0.20045882959 |
| | Utp4 | -1.15 | 1.306420377 | 0.049383244842682700 | 0.22130890818 |
| | Cdc37l1 | -1.15 | 1.492454771 | 0.032176976169733800 | 0.18774684907 |
| | Sfxn3 | -1.15 | 1.57054588 | 0.026881538458334000 | 0.17656750476 |
| | Ipo9 | -1.15 | 1.431477835 | 0.037027310206655400 | 0.19853187451 |
| | Dctn1 | -1.15 | 2.056716445 | 0.008775736097681120 | 0.14048876521 |
| | Srgap2 | -1.15 | 1.676122858 | 0.021080317201260200 | 0.16261000251 |
| | Plxna2 | -1.15 | 1.768418709 | 0.017044383241005400 | 0.15563363212 |
| | Peg13 | -1.15 | 1.354798783 | 0.044177508251363100 | 0.21092058116 |
| | Taok2 | -1.15 | 1.894353431 | 0.012754004587564700 | 0.14587849176 |
| | Pdxk | -1.15 | 1.653189388 | 0.022223405560852800 | 0.16526067964 |
| | Gabbr1 | -1.15 | 1.541474561 | 0.028742559478344400 | 0.17993235070 |
| | Chst10 | -1.15 | 1.683633984 | 0.020718867660916800 | 0.16194423238 |
| | Stard3 | -1.15 | 1.509464111 | 0.030941109976876000 | 0.18547374364 |
| | Spata2 | -1.15 | 1.707951856 | 0.019590618345819000 | 0.16048239041 |
| | Clcn6 | -1.15 | 1.612235999 | 0.024421031369725500 | 0.16982619623 |
| | Nedd4l | -1.15 | 1.582745395 | 0.026136931798658300 | 0.17473771883 |
| | Itsn1 | -1.15 | 1.915651263 | 0.012143635875651100 | 0.14372584028 |
| | Lrtm2 | -1.15 | 1.731199751 | 0.018569501663213900 | 0.15996749329 |
| | Pex14 | -1.15 | 1.736115452 | 0.018360501871452300 | 0.15937095786 |

| | | | | | |
|---|---|---|---|---|---|
| | Ndrg3 | -1.15 | 1.435158766 | 0.036714805687346200 | 0.19792002548 |
| | Camk2b | -1.15 | 1.756528166 | 0.017517488196743900 | 0.15705144846 |
| | Got1 | -1.15 | 1.378632826 | 0.041818377054484200 | 0.20572143528 |
| | Ppp6r2 | -1.15 | 1.626978706 | 0.023605939734469100 | 0.16797991605 |
| | Map6 | -1.15 | 1.727970723 | 0.018708082505690400 | 0.15996749329 |
| | Hsph1 | -1.15 | 1.49430461 | 0.032040212676146100 | 0.18734662229 |
| | Crebbp | -1.15 | 1.322861595 | 0.047548673382566500 | 0.21755514385 |
| | Ercc6 | -1.15 | 1.414677156 | 0.038487778470920500 | 0.20070609638 |
| | Ptms | -1.15 | 1.823288292 | 0.015021444851504400 | 0.15143776501 |
| | Zmat3 | -1.15 | 1.414056792 | 0.038542795274609100 | 0.20078145755 |
| | Pom121 | -1.15 | 1.727047959 | 0.018747874665562500 | 0.16020805086 |
| | Spock2 | -1.15 | 2.177690758 | 0.006642158610292290 | 0.13324073206 |
| | Icmt | -1.15 | 1.838106964 | 0.014517540159890100 | 0.15044156813 |
| | Smarcc2 | -1.15 | 1.855386816 | 0.013951252017098100 | 0.14862753048 |
| | Pacsin1 | -1.16 | 1.307085925 | 0.049307623920645500 | 0.22128146941 |
| | Pnpla6 | -1.16 | 1.555902765 | 0.027803356930510900 | 0.17888249734 |
| | Nt5m | -1.16 | 1.796032537 | 0.015994381945630800 | 0.15212233233 |
| | Mapt | -1.16 | 1.673949044 | 0.021186096973121000 | 0.16273689505 |
| | Ahcyl2 | -1.16 | 1.690181439 | 0.020408851298760000 | 0.16148333185 |
| | Dlg2 | -1.16 | 1.378183729 | 0.041861643141797500 | 0.20572143528 |
| | Kif3c | -1.16 | 1.718991166 | 0.019098921078809600 | 0.16048070792 |
| | Srpk2 | -1.16 | 1.443446124 | 0.036020843302710800 | 0.19681983497 |
| | Basp1 | -1.16 | 1.473271036 | 0.033630162356425500 | 0.19145320024 |

| | | | | | |
|---|---|---|---|---|---|
| | Clasp1 | -1.16 | 1.399963367 | 0.039814075228599800 | 0.20241237663 |
| | Gng3 | -1.16 | 1.480160309 | 0.033100891554178700 | 0.19038901249 |
| | Rai1 | -1.16 | 1.378046328 | 0.041874889343193100 | 0.20572143528 |
| | Prpf39 | -1.16 | 1.442626683 | 0.036088872787744800 | 0.19681983497 |
| | Snn | -1.16 | 1.40998622 | 0.038905748928019700 | 0.20150919564 |
| | Crebl2 | -1.16 | 1.419198847 | 0.038089138794579100 | 0.20006989915 |
| | Ankrd11 | -1.16 | 1.949536094 | 0.011232176169080000 | 0.14193386802 |
| | Tnik | -1.16 | 1.996698589 | 0.010076307460140100 | 0.14109528430 |
| | Hira | -1.16 | 1.882462734 | 0.013108025157517000 | 0.14665206711 |
| | Dexi | -1.16 | 1.56333835 | 0.027331385616746600 | 0.17832885554 |
| | Arhgap44 | -1.16 | 1.845604913 | 0.014269050889755400 | 0.14978688180 |
| | Atcay | -1.16 | 1.705327302 | 0.019709367995062600 | 0.16067121778 |
| | Pfdn2 | -1.16 | 1.532929829 | 0.029313668424340600 | 0.18127773079 |
| | Usp28 | -1.16 | 1.430660866 | 0.037097029404234400 | 0.19873305304 |
| | Phospho2 | -1.16 | 1.505650352 | 0.031214015926163100 | 0.18582135954 |
| | Epas1 | -1.16 | 1.731154631 | 0.018571430999149700 | 0.15996749329 |
| | Slc12a6 | -1.16 | 1.688445558 | 0.020490588994194800 | 0.16163098930 |
| | Nkrf | -1.16 | 1.707462294 | 0.019612714468082200 | 0.16048239041 |
| | Stim2 | -1.16 | 1.470377317 | 0.033854989469827900 | 0.19180778285 |
| | Fam234b | -1.16 | 2.047699811 | 0.008959838650450640 | 0.14048876521 |
| | Plekha8 | -1.16 | 1.40027268 | 0.039785728965542000 | 0.20241237663 |
| | Usp20 | -1.16 | 1.718291284 | 0.019129724535729200 | 0.16048070792 |
| | Tmed8 | -1.16 | 1.925974965 | 0.011858371049063600 | 0.14364743987 |

| | | | | | |
|---|---|---|---|---|---|
| | Ss18l1 | -1.16 | 1.548868833 | 0.028257332860486900 | 0.17947407154 |
| | Neurl4 | -1.16 | 1.695123863 | 0.020177907994670000 | 0.16092051453 |
| | Nlgn2 | -1.16 | 1.737932663 | 0.018283836827099500 | 0.15923263788 |
| | Eogt | -1.16 | 1.394608024 | 0.040308067358912300 | 0.20310713369 |
| | Nop56 | -1.16 | 1.840911905 | 0.014424079090857400 | 0.15044156813 |
| | Zrsr2 | -1.16 | 1.864140051 | 0.013672878309147800 | 0.14796465545 |
| | Lsm14b | -1.16 | 2.214815291 | 0.006097961933142910 | 0.13146915442 |
| | Rere | -1.16 | 1.407792996 | 0.039102723204418100 | 0.20180964127 |
| | Sec61a2 | -1.16 | 1.310320158 | 0.048941789175356400 | 0.22035030310 |
| | Dgkq | -1.16 | 1.810083664 | 0.015485182780943600 | 0.15174905791 |
| | Papd7 | -1.16 | 1.434139877 | 0.036801042583456400 | 0.19822937128 |
| | Arhgef9 | -1.16 | 1.55905239 | 0.027602448636143000 | 0.17859523991 |
| | Drosha | -1.16 | 1.986981076 | 0.010304310191678300 | 0.14109528430 |
| | Med12l | -1.16 | 1.579731653 | 0.026318937128266600 | 0.17487839220 |
| | Kif21a | -1.16 | 1.369262181 | 0.042730484704382000 | 0.20784410277 |
| | Golga4 | -1.16 | 1.787272838 | 0.016320263343676800 | 0.15255560449 |
| | Mapk8ip2 | -1.16 | 2.007131273 | 0.009837137168487820 | 0.14078337569 |
| | Iglon5 | -1.16 | 1.365209026 | 0.043131143657280400 | 0.20875838147 |
| | Syngap1 | -1.16 | 1.589711135 | 0.025721060152735900 | 0.17361033068 |
| | Syngr1 | -1.16 | 1.524367308 | 0.029897349780922200 | 0.18308227454 |
| | Tspan7 | -1.16 | 1.611461382 | 0.024464628074554000 | 0.16982619623 |
| | Ecsit | -1.16 | 1.449621623 | 0.035512265380608900 | 0.19597350948 |
| | Slc7a6 | -1.16 | 1.410738474 | 0.038838417465263400 | 0.20139624678 |

| | | | | | |
|---|---|---|---|---|---|
| | Smg1 | -1.16 | 1.883269504 | 0.013083697559296200 | 0.14665206711 |
| | Kbtbd2 | -1.17 | 1.985090645 | 0.010349261369084800 | 0.14109528430 |
| | Dzip1l | -1.17 | 1.343731394 | 0.045317778432561 00 | 0.21288007959 |
| | Dpcd | -1.17 | 1.793230757 | 0.016097900650842900 | 0.15221980976 |
| | Mark1 | -1.17 | 1.425445328 | 0.037545221613374700 | 0.19956638206 |
| | 1810055G02Rik | -1.17 | 1.368854117 | 0.042770653160089900 | 0.20796586874 |
| | Ube4b | -1.17 | 2.664049908 | 0.002167455011823410 | 0.12702075696 |
| | Pnkd | -1.17 | 1.636266306 | 0.023106474867728100 | 0.16683129036 |
| | Clec16a | -1.17 | 2.011076144 | 0.009748187100251860 | 0.14078337569 |
| | Impdh1 | -1.17 | 1.670962417 | 0.021332295124949900 | 0.16305711054 |
| | Numb | -1.17 | 1.978008482 | 0.010519413282366900 | 0.14109528430 |
| | Bms1 | -1.17 | 1.304571481 | 0.049593929360583700 | 0.22130890818 |
| | Map3k10 | -1.17 | 1.606377139 | 0.024752716108981800 | 0.17028275660 |
| | Snx30 | -1.17 | 1.475168134 | 0.033483578512170800 | 0.19112913454 |
| | Hdac5 | -1.17 | 1.407964468 | 0.039087287434130000 | 0.20180964127 |
| | Edrf1 | -1.17 | 1.554394633 | 0.027900074761986500 | 0.17888249734 |
| | Ptp4a3 | -1.17 | 1.427841209 | 0.037338665382489700 | 0.19917336996 |
| | Fam168b | -1.17 | 1.670644454 | 0.021347919009442400 | 0.16305711054 |
| | Clip2 | -1.17 | 2.188407788 | 0.006480256732524860 | 0.13306070174 |
| | Efr3b | -1.17 | 2.306217677 | 0.004940629908702610 | 0.13146915442 |
| | Sptan1 | -1.17 | 1.545732241 | 0.028462153694678100 | 0.17947407154 |
| | Atg2b | -1.17 | 1.49708806 | 0.031835519439963500 | 0.18710014086 |
| | Lhfpl4 | -1.17 | 2.72296271 | 0.001892506107714930 | 0.12702075696 |

| | | | | | |
|---|---|---|---|---|---|
| | Ppp2r5b | -1.17 | 2.36888561 | 0.004276755176136880 | 0.13146915442 |
| | Bcl11a | -1.17 | 1.656393257 | 0.022060062740706300 | 0.16510202730 |
| | Usp32 | -1.17 | 1.471416888 | 0.033774047665742600 | 0.19177239214 |
| | Rnf26 | -1.17 | 1.605655671 | 0.024793870524379900 | 0.17034628744 |
| | Mfhas1 | -1.17 | 1.780166442 | 0.016589509962311400 | 0.15371165215 |
| | Rapgef1 | -1.17 | 1.647768263 | 0.022502550074950100 | 0.16645864635 |
| | Abhd8 | -1.17 | 1.348084261 | 0.044865833396157000 | 0.21195108826 |
| | Fsd1 | -1.17 | 1.653493763 | 0.022207835775525200 | 0.16526067964 |
| | Gas7 | -1.17 | 1.329812605 | 0.046793700907737400 | 0.21616201339 |
| | Hmgxb3 | -1.17 | 2.223910352 | 0.005971585400847710 | 0.13146915442 |
| | Rab3a | -1.17 | 1.677319723 | 0.021022302288577900 | 0.16261000251 |
| | Ubfd1 | -1.17 | 2.550735639 | 0.002813612993987240 | 0.12870390352 |
| | Katnal1 | -1.17 | 2.160449301 | 0.006911156041595540 | 0.13451302432 |
| | Clip1 | -1.17 | 2.116938226 | 0.007639444389910700 | 0.13628710184 |
| | Scn2b | -1.17 | 1.435797057 | 0.036660884858002900 | 0.19778453822 |
| | Sh3bp5l | -1.17 | 1.986486129 | 0.010316060279495200 | 0.14109528430 |
| | Pou3f3 | -1.17 | 1.891951505 | 0.012824737817546100 | 0.14588139267 |
| | Rab11fip3 | -1.17 | 1.905530894 | 0.012429942108098800 | 0.14536156128 |
| | Pfkfb2 | -1.17 | 1.311985253 | 0.048754504533426000 | 0.21979515971 |
| | Sv2a | -1.17 | 1.791358937 | 0.016167432748198400 | 0.15237084595 |
| | Stxbp1 | -1.17 | 1.371857868 | 0.042475855283468700 | 0.20763455263 |
| | Narf | -1.17 | 1.620163548 | 0.023979297297498200 | 0.16888750598 |
| | Sugp2 | -1.17 | 1.533688789 | 0.029262485485025600 | 0.18120586438 |

| | | | | | |
|---|---|---|---|---|---|
| | Spsb3 | -1.17 | 1.538886567 | 0.028914349975892700 | 0.18010520536 |
| | Ppp1r3f | -1.17 | 1.436795521 | 0.036576696538459100 | 0.19756304526 |
| | Chrnb2 | -1.17 | 2.062823644 | 0.008653192313097260 | 0.14048876521 |
| | Dpp9 | -1.17 | 2.355202793 | 0.004413643052517180 | 0.13146915442 |
| | Fxr2 | -1.17 | 2.18509831 | 0.006529827220127890 | 0.13306070174 |
| | Klhdc3 | -1.17 | 1.540421809 | 0.028812317437215800 | 0.17995201039 |
| | Nbea | -1.18 | 1.380646628 | 0.041624916226720400 | 0.20572143528 |
| | Mib2 | -1.18 | 1.489878496 | 0.032368420248448900 | 0.18838393165 |
| | Gpc5 | -1.18 | 1.642196904 | 0.022793084286130200 | 0.16683129036 |
| | Mras | -1.18 | 1.400124355 | 0.039799319384762000 | 0.20241237663 |
| | Sec14l1 | -1.18 | 1.983096108 | 0.010396900591605300 | 0.14109528430 |
| | Nova2 | -1.18 | 1.786485487 | 0.016349877879238300 | 0.15272853293 |
| | Gtpbp2 | -1.18 | 1.752834174 | 0.017667122748002700 | 0.15743445764 |
| | Atxn2 | -1.18 | 1.752970254 | 0.017661587846321800 | 0.15743445764 |
| | Ddx19b | -1.18 | 1.569976558 | 0.026916800897784000 | 0.17663025842 |
| | Rtn2 | -1.18 | 1.911838197 | 0.012250725347649500 | 0.14449546524 |
| | Msl1 | -1.18 | 2.357122704 | 0.004394174464436850 | 0.13146915442 |
| | Ckmt1 | -1.18 | 1.552767023 | 0.028004832325983800 | 0.17898344232 |
| | Abcg4 | -1.18 | 1.96372169 | 0.010871220642293600 | 0.14193386802 |
| | Ptprs | -1.18 | 2.141665831 | 0.007216625508570930 | 0.13538768215 |
| | Slc35e2 | -1.18 | 2.057162066 | 0.008766736110963680 | 0.14048876521 |
| | Wasf3 | -1.18 | 1.68470576 | 0.020667799511950000 | 0.16185943212 |
| | Klf13 | -1.18 | 1.827518486 | 0.014875840534204000 | 0.15143776501 |

| | | | | | |
|---|---|---|---|---|---|
| | Cdk5r1 | -1.18 | 1.922542157 | 0.011952474961337700 | 0.14364743987 |
| | Stx3 | -1.18 | 1.33382956 | 0.046362883659766800 | 0.21494137193 |
| | Zrsr1 | -1.18 | 1.47674827 | 0.033361973229660300 | 0.19112913454 |
| | Ubr4 | -1.18 | 1.630390832 | 0.023421201343828700 | 0.16779495707 |
| | B4galnt1 | -1.18 | 1.626998378 | 0.023604870491972900 | 0.16797991605 |
| | Trim46 | -1.18 | 1.507538292 | 0.031078618681247000 | 0.18559378501 |
| | Ppp1r9b | -1.18 | 1.423716359 | 0.037694990748776600 | 0.19959349860 |
| | Arhgef4 | -1.18 | 1.328299082 | 0.046957062232714000 | 0.21644984641 |
| | 2810403A07Rik | -1.18 | 1.363039414 | 0.043347153774150700 | 0.20928785664 |
| | Cnnm1 | -1.18 | 1.450291054 | 0.035457568159624700 | 0.19580841658 |
| | Cdan1 | -1.18 | 1.818746942 | 0.015179345899039200 | 0.15152290628 |
| | Dennd4b | -1.18 | 1.662648301 | 0.021744613837160400 | 0.16435244155 |
| | Bicd2 | -1.18 | 2.215836994 | 0.006083632979371190 | 0.13146915442 |
| | Herc1 | -1.18 | 1.448633001 | 0.035593197046124400 | 0.19610510049 |
| | Syp | -1.18 | 1.457492775 | 0.034874438609317400 | 0.19416923052 |
| | Pik3r2 | -1.18 | 1.423518095 | 0.037712203164504100 | 0.19959349860 |
| | Srgap3 | -1.18 | 1.862350168 | 0.013729345440168500 | 0.14796465545 |
| | Sbno1 | -1.18 | 2.607815101 | 0.002467089466492850 | 0.12702075696 |
| | Rmnd5a | -1.18 | 1.660405446 | 0.021857201378794600 | 0.16455412374 |
| | Pip4k2c | -1.18 | 1.797141623 | 0.015953588169726200 | 0.15212233233 |
| | Herc2 | -1.18 | 1.73919802 | 0.018230642716834300 | 0.15915327927 |
| | Coq10b | -1.18 | 1.350482081 | 0.044618803347750100 | 0.21164750713 |
| | Atp6v1g2 | -1.18 | 1.391660438 | 0.040582571481784800 | 0.20330985190 |

| | | | | | |
|---|---|---|---|---|---|
| | Ints1 | -1.19 | 1.540259448 | 0.028823090926707000 | 0.17995201039 |
| | Cpsf7 | -1.19 | 1.679212374 | 0.020930886642621000 | 0.16246887298 |
| | Arhgap35 | -1.19 | 1.901522534 | 0.012545196445756500 | 0.14564091316 |
| | Aatk | -1.19 | 1.632391232 | 0.023313569230211500 | 0.16746040501 |
| | Mgll | -1.19 | 1.463231556 | 0.034416637983264300 | 0.19283467894 |
| | Btbd10 | -1.19 | 1.41946988 | 0.038065375648112700 | 0.20006989915 |
| | Abhd6 | -1.19 | 1.43028865 | 0.037128837342875000 | 0.19874848225 |
| | Pld3 | -1.19 | 1.370759601 | 0.042583406375162500 | 0.20766997023 |
| | Cend1 | -1.19 | 1.759358154 | 0.017403710305857100 | 0.15671857076 |
| | Ddhd2 | -1.19 | 1.852316264 | 0.014050239777265800 | 0.14864677641 |
| | Flrt1 | -1.19 | 1.426015038 | 0.037496001822566100 | 0.19947060048 |
| | Eml2 | -1.19 | 1.388453862 | 0.040883318253807200 | 0.20403256309 |
| | Atp5sl | -1.19 | 1.395909144 | 0.040187487595272000 | 0.20284101714 |
| | Dnm1 | -1.19 | 1.326483909 | 0.047153734230360000 | 0.21691980651 |
| | Cdk9 | -1.19 | 1.726602439 | 0.018767116992773400 | 0.16021558505 |
| | Zfp335 | -1.19 | 1.381922152 | 0.041502843086900300 | 0.20539257521 |
| | Atg9a | -1.19 | 2.24968739 | 0.005627462500535290 | 0.13146915442 |
| | Eef1a2 | -1.19 | 1.421634812 | 0.037876094167651500 | 0.19976395727 |
| | E2f6 | -1.19 | 1.947679258 | 0.011280302403738100 | 0.14193386802 |
| | Atg13 | -1.19 | 2.95246344 | 0.001115672068573980 | 0.12702075696 |
| | Slc6a8 | -1.19 | 2.004356931 | 0.009900179506589680 | 0.14109528430 |
| | Tmem38a | -1.19 | 2.118622315 | 0.007609877838968910 | 0.13628710184 |
| | Rab11fip4 | -1.19 | 1.838839671 | 0.014493067963579200 | 0.15044156813 |

| | | | | | |
|---|---|---|---|---|---|
| | Eno2 | -1.19 | 1.704771225 | 0.019734620312612600 | 0.16067121778 |
| | Adgra1 | -1.19 | 1.322315555 | 0.047608494109418400 | 0.21755514385 |
| | Zc3h13 | -1.19 | 2.356090074 | 0.004404635010119140 | 0.13146915442 |
| | Fry | -1.19 | 1.863507468 | 0.013692808428545600 | 0.14796465545 |
| | Sgtb | -1.19 | 1.405120871 | 0.039344055977623800 | 0.20203443738 |
| | Zbtb4 | -1.19 | 1.84750456 | 0.014206772973411900 | 0.14936133698 |
| | Jcad | -1.19 | 1.462690712 | 0.034459525129689700 | 0.19283467894 |
| | Ppip5k1 | -1.19 | 1.814271938 | 0.015336563662166800 | 0.15172046169 |
| | Ap1ar | -1.19 | 1.86487151 | 0.013649869215637200 | 0.14796465545 |
| | Habp4 | -1.19 | 1.432342203 | 0.036953688745385700 | 0.19853187451 |
| | Asphd2 | -1.19 | 1.385528293 | 0.041159653202737800 | 0.20469590831 |
| | Ypel3 | -1.19 | 1.980938238 | 0.010448688021460100 | 0.14109528430 |
| | Ing2 | -1.19 | 1.37139173 | 0.042521470047338400 | 0.20764569248 |
| | Rab6b | -1.19 | 1.370389744 | 0.042619687044115400 | 0.20774640641 |
| | Ralgapb | -1.19 | 1.768438524 | 0.017043605611315400 | 0.15563363212 |
| | Fam171a2 | -1.19 | 1.524349188 | 0.029898597195954900 | 0.18308227454 |
| | Tmem250-ps | -1.19 | 1.92399416 | 0.011912580280202600 | 0.14364743987 |
| | Map9 | -1.19 | 1.364240043 | 0.043227483871376500 | 0.20900381980 |
| | Pcnt | -1.19 | 1.436582564 | 0.036594636346315800 | 0.19758227821 |
| | Kctd17 | -1.19 | 1.334257902 | 0.046317178798149200 | 0.21494137193 |
| | 5-Sep | -1.19 | 1.404618974 | 0.039389550629188500 | 0.20216833599 |
| | Gnl1 | -1.19 | 1.740934387 | 0.018157899700209800 | 0.15903535871 |
| | Ehbp1 | -1.19 | 1.862502061 | 0.013724544462841800 | 0.14796465545 |

| | | | | | |
|---|---|---|---|---|---|
| | Pitpnm1 | -1.19 | 1.302773224 | 0.049799705638461300 | 0.22145558420 |
| | Rhot2 | -1.19 | 1.472958209 | 0.033654395241414900 | 0.19145320024 |
| | Bcl9 | -1.19 | 1.794326621 | 0.016057331711726600 | 0.15212233233 |
| | Slc7a1 | -1.19 | 1.620533151 | 0.023958898563904100 | 0.16888750598 |
| | Drp2 | -1.19 | 1.398854531 | 0.039915857998860600 | 0.20241237663 |
| | Fbxo9 | -1.20 | 1.334625066 | 0.046278037521829500 | 0.21494137193 |
| | Ppard | -1.20 | 1.342458756 | 0.045450769838875000 | 0.21322602539 |
| | Zfp523 | -1.20 | 1.978428138 | 0.010509253346435600 | 0.14109528430 |
| | Setd1b | -1.20 | 1.443378008 | 0.036026493340845100 | 0.19681983497 |
| | Tsc1 | -1.20 | 2.309599613 | 0.004902305670524800 | 0.13146915442 |
| | Plxna4 | -1.20 | 1.841110748 | 0.014417476489844300 | 0.15044156813 |
| | Ano8 | -1.20 | 2.292923809 | 0.005094202339478140 | 0.13146915442 |
| | Zfp598 | -1.20 | 1.343509913 | 0.045340894820980800 | 0.21288007959 |
| | Mapk9 | -1.20 | 1.540515576 | 0.028806097374302800 | 0.17995201039 |
| | Dock3 | -1.20 | 1.915801088 | 0.012139447245192200 | 0.14372584028 |
| | Atxn7l3 | -1.20 | 1.558117151 | 0.027661953623045300 | 0.17859523991 |
| | Agap2 | -1.20 | 1.464401011 | 0.034324086563471600 | 0.19282390575 |
| | Tubg2 | -1.20 | 1.638272086 | 0.023000004151113800 | 0.16683129036 |
| | Crebzf | -1.20 | 2.022193589 | 0.009501811506856860 | 0.14078337569 |
| | Kmt5a | -1.20 | 2.306272675 | 0.004940004274180540 | 0.13146915442 |
| | Krba1 | -1.20 | 1.505941982 | 0.031193062666266900 | 0.18579274993 |
| | Coro2b | -1.20 | 1.670923759 | 0.021334194039411500 | 0.16305711054 |
| | Zc3h3 | -1.20 | 1.508472914 | 0.031011807989965700 | 0.18558122979 |

| | | | | | |
|---|---|---|---|---|---|
| | Napa | -1.20 | 1.540685536 | 0.028794826376064100 | 0.17995201039 |
| | Spry2 | -1.20 | 1.935039239 | 0.011613436795160300 | 0.14312128700 |
| | Zfp385b | -1.20 | 1.879078536 | 0.013210567167113700 | 0.14682504272 |
| | Ice1 | -1.20 | 1.996491481 | 0.010081113830928600 | 0.14109528430 |
| | Pcdh1 | -1.20 | 1.347496914 | 0.044926551714105100 | 0.21195108826 |
| | Lin7b | -1.20 | 1.303756999 | 0.049687025796396500 | 0.22131753045 |
| | Trak2 | -1.20 | 1.943270756 | 0.011395391324211500 | 0.14196161768 |
| | Atp1a3 | -1.20 | 1.766731714 | 0.017110720041514000 | 0.15591406107 |
| | Pnpo | -1.20 | 1.571648491 | 0.026813376671790300 | 0.17639876384 |
| | Ubald1 | -1.20 | 1.470261318 | 0.033864033272371500 | 0.19180778285 |
| | Zfp251 | -1.20 | 2.550248877 | 0.002816768289403940 | 0.12870390352 |
| | Tubgcp6 | -1.20 | 1.807660212 | 0.015571834824283100 | 0.15184875954 |
| | Grik5 | -1.20 | 2.187064652 | 0.006500329153308900 | 0.13306070174 |
| | Mthfsd | -1.20 | 1.553922487 | 0.027930423010534100 | 0.17888249734 |
| | Mapk14 | -1.20 | 2.117593597 | 0.007627924808956020 | 0.13628710184 |
| | Calm3 | -1.20 | 1.792982758 | 0.016107095799988200 | 0.15221980976 |
| | Slc20a1 | -1.20 | 2.525079337 | 0.002984837299337490 | 0.12870390352 |
| | 4933427D14Rik | -1.20 | 1.887704048 | 0.012950780789708400 | 0.14628049103 |
| | Fam212b | -1.20 | 1.827283429 | 0.014883894079098400 | 0.15143776501 |
| | Zfp612 | -1.20 | 1.599700817 | 0.025136174528433400 | 0.17153549292 |
| | Smurf1 | -1.20 | 1.634002963 | 0.023227209508954400 | 0.16727729867 |
| | Arhgap23 | -1.20 | 1.682779841 | 0.020759656291979700 | 0.16207865745 |
| | Ulk1 | -1.20 | 2.430936733 | 0.003707347259204800 | 0.12985696725 |

| | | | | | |
|---|---|---|---|---|---|
| | A230050P20Rik | -1.21 | 1.589580527 | 0.025728796560439800 | 0.17361033068 |
| | Megf8 | -1.21 | 2.824487674 | 0.001498001768647780 | 0.12702075696 |
| | Lrrc4 | -1.21 | 1.34766179 | 0.044909498980762700 | 0.21195108826 |
| | Cacnb3 | -1.21 | 1.495801378 | 0.031929978191362100 | 0.18710014086 |
| | Lrrc24 | -1.21 | 1.741172617 | 0.018147942003129500 | 0.15903535871 |
| | Cmc2 | -1.21 | 1.489339784 | 0.032408595925208100 | 0.18838561902 |
| | Madd | -1.21 | 1.456457507 | 0.034957671188043900 | 0.19429017089 |
| | Clk4 | -1.21 | 1.707658622 | 0.019603850318575300 | 0.16048239041 |
| | Brwd1 | -1.21 | 3.07543533 | 0.000840552163179585 | 0.12354271087 |
| | 5330417C22Rik | -1.21 | 1.816645256 | 0.015252981515464400 | 0.15152290628 |
| | Dlg4 | -1.21 | 1.510087756 | 0.030896710540728900 | 0.18539604295 |
| | Josd1 | -1.21 | 2.223691217 | 0.005974599285021750 | 0.13146915442 |
| | Hook1 | -1.21 | 1.580973308 | 0.026243798322550800 | 0.17487839220 |
| | Zmiz2 | -1.21 | 1.547887533 | 0.028321253199812100 | 0.17947407154 |
| | Atxn1 | -1.21 | 1.827260901 | 0.014884666166708600 | 0.15143776501 |
| | Unc13b | -1.21 | 1.773590679 | 0.016842607226724100 | 0.15479465248 |
| | Ccdc9 | -1.21 | 1.880527706 | 0.013166559140056900 | 0.14682504272 |
| | Rusc2 | -1.21 | 2.507781655 | 0.003106120819310820 | 0.12870390352 |
| | Faim2 | -1.21 | 1.384521006 | 0.041255228193572500 | 0.20494869509 |
| | Elavl3 | -1.21 | 2.741669647 | 0.001812718442964970 | 0.12702075696 |
| | Dgki | -1.21 | 1.375973219 | 0.042075257399683400 | 0.20626332926 |
| | Ube2o | -1.21 | 1.760209594 | 0.017369623547678500 | 0.15671857076 |
| | Arhgef11 | -1.21 | 2.174794697 | 0.006686599374730450 | 0.13346292906 |

| | | | | | |
|---|---|---|---|---|---|
| | Adarb1 | -1.21 | 1.815089373 | 0.015307724115265800 | 0.15159717255 |
| | Schip1 | -1.21 | 1.615727742 | 0.024225472605272400 | 0.16966419499 |
| | Caskin1 | -1.21 | 1.783451594 | 0.016464494682138500 | 0.15327829247 |
| | Map3k12 | -1.21 | 2.064309515 | 0.008623637360884620 | 0.14048876521 |
| | Mturn | -1.21 | 1.398171432 | 0.039978690792956100 | 0.20241237663 |
| | Ap5z1 | -1.21 | 1.967297334 | 0.010782082888588300 | 0.14150160435 |
| | Rbfox2 | -1.21 | 2.206281113 | 0.006218976084093020 | 0.13146915442 |
| | Ankrd52 | -1.21 | 1.917245208 | 0.012099148082245600 | 0.14364743987 |
| | Prdm2 | -1.21 | 1.925426681 | 0.011873351328029800 | 0.14364743987 |
| | Fam219aos | -1.21 | 1.742153346 | 0.018107006349300300 | 0.15898298674 |
| | Cdc42bpa | -1.21 | 1.463844581 | 0.034368091756424600 | 0.19283460548 |
| | Aak1 | -1.21 | 1.752030468 | 0.017699847797995600 | 0.15746422541 |
| | Tmem151a | -1.21 | 1.366485657 | 0.043004543581551300 | 0.20864938598 |
| | Cox10 | -1.21 | 2.262465049 | 0.005464305240775960 | 0.13146915442 |
| | Trio | -1.21 | 2.690472244 | 0.002039518999887060 | 0.12702075696 |
| | Fam189b | -1.21 | 2.186092767 | 0.006514892184750220 | 0.13306070174 |
| | Iffo1 | -1.21 | 1.484058453 | 0.032805113705309200 | 0.18957195214 |
| | Prmt8 | -1.21 | 1.899709252 | 0.012597685080602900 | 0.14573900673 |
| | Ctif | -1.21 | 1.782318323 | 0.016507514102972600 | 0.15336308885 |
| | Mef2d | -1.21 | 2.295306747 | 0.005066327415284900 | 0.13146915442 |
| | Marf1 | -1.21 | 2.120976168 | 0.007568744278692790 | 0.13628710184 |
| | Chchd4 | -1.21 | 1.470563207 | 0.033840501715818100 | 0.19180778285 |
| | Prkcz | -1.21 | 2.064914883 | 0.008611625146067260 | 0.14048876521 |

| | | | | | |
|---|---|---|---|---|---|
| | St3gal3 | -1.21 | 2.046774095 | 0.008978957267468470 | 0.14048876521 |
| | Tcea2 | -1.21 | 1.350043919 | 0.044663842271721900 | 0.21164750713 |
| | B4galnt4 | -1.22 | 1.468133029 | 0.034030393501787100 | 0.19202608564 |
| | Slc6a1 | -1.22 | 1.983337373 | 0.010391126394269200 | 0.14109528430 |
| | Chst1 | -1.22 | 1.981538589 | 0.010434254162877100 | 0.14109528430 |
| | Mon1a | -1.22 | 1.838263455 | 0.014512309951698700 | 0.15044156813 |
| | Safb | -1.22 | 1.67597534 | 0.021087478829326100 | 0.16261000251 |
| | Adgrb2 | -1.22 | 1.446217527 | 0.035791712070945700 | 0.19633290043 |
| | Srpk1 | -1.22 | 2.190652621 | 0.006446847234876540 | 0.13301220399 |
| | Mllt6 | -1.22 | 2.101910226 | 0.007908420882924640 | 0.13703952374 |
| | 2410089E03Rik | -1.22 | 2.216796781 | 0.006070203053633050 | 0.13146915442 |
| | Htt | -1.22 | 2.509535905 | 0.003093599547163950 | 0.12870390352 |
| | Wbscr17 | -1.22 | 1.616614127 | 0.024176079364647400 | 0.16966419499 |
| | Nefl | -1.22 | 1.49286931 | 0.032146277542274600 | 0.18774684907 |
| | Lgi3 | -1.22 | 1.32240556 | 0.047598628623603100 | 0.21755514385 |
| | Kcnn1 | -1.22 | 1.403416858 | 0.039498730872080400 | 0.20216833599 |
| | Mrm1 | -1.22 | 1.582424569 | 0.026156247096648600 | 0.17473771883 |
| | Dynll2 | -1.22 | 1.941810499 | 0.011433771308461100 | 0.14204082704 |
| | Bcat1 | -1.22 | 1.654542391 | 0.022154278403974500 | 0.16526067964 |
| | Lyst | -1.22 | 2.141343778 | 0.007221979012308470 | 0.13538768215 |
| | Stx1b | -1.22 | 1.809199176 | 0.015516752194929800 | 0.15174905791 |
| | Snrpn | -1.22 | 1.809973017 | 0.015489128498272600 | 0.15174905791 |
| | Plbd2 | -1.22 | 1.517319222 | 0.030386506908944800 | 0.18410096624 |

| | | | | | |
|---|---|---|---|---|---|
| | Hdgfl2 | -1.22 | 1.68423737 | 0.020690101925824000 | 0.16185943212 |
| | Spire1 | -1.22 | 1.948562767 | 0.011257377584733700 | 0.14193386802 |
| | Cpeb3 | -1.22 | 1.978682232 | 0.010503106479347600 | 0.14109528430 |
| | Inafm2 | -1.22 | 1.816897241 | 0.015244134058687100 | 0.15152290628 |
| | Sult4a1 | -1.22 | 1.582209856 | 0.026169181803233800 | 0.17473771883 |
| | Irgq | -1.22 | 1.911200552 | 0.012268725449171600 | 0.14458366758 |
| | Adgrl4 | -1.22 | 2.042261138 | 0.009072748296039950 | 0.14048876521 |
| | Plk2 | -1.22 | 1.920942696 | 0.011996575841619000 | 0.14364743987 |
| | Slc7a14 | -1.22 | 2.442947666 | 0.003606220968185140 | 0.12985696725 |
| | Phf24 | -1.22 | 1.349912576 | 0.044677351933140500 | 0.21164750713 |
| | Mettl7a1 | -1.22 | 1.679990378 | 0.020893424217354700 | 0.16246887298 |
| | Mast2 | -1.22 | 1.876537166 | 0.013288098348785900 | 0.14686395323 |
| | Ttc39b | -1.22 | 1.337404945 | 0.045982762133387700 | 0.21433145674 |
| | Slx4 | -1.22 | 1.636089328 | 0.023115892798137800 | 0.16683129036 |
| | Pgbd5 | -1.22 | 2.147933283 | 0.007113227805808040 | 0.13466036492 |
| | Med1 | -1.22 | 2.048889783 | 0.008935322198869510 | 0.14048876521 |
| | Ubp1 | -1.23 | 1.839484496 | 0.014471565162113500 | 0.15044156813 |
| | Ttll5 | -1.23 | 1.637314044 | 0.023050797531448700 | 0.16683129036 |
| | Sorbs2 | -1.23 | 1.355596075 | 0.044096480157314500 | 0.21075109435 |
| | Snurf | -1.23 | 1.826634012 | 0.014906167183077200 | 0.15143776501 |
| | Ppp2r2c | -1.23 | 1.757689279 | 0.017470716697528900 | 0.15680830472 |
| | Cacna1e | -1.23 | 1.549706216 | 0.028202901101289100 | 0.17947407154 |
| | Agtpbp1 | -1.23 | 1.447326123 | 0.035700465362501400 | 0.19629181303 |

| | | | | | |
|---|---|---|---|---|---|
| | Clk3 | -1.23 | 1.942975007 | 0.011403154100975600 | 0.14196161768 |
| | Apba2 | -1.23 | 2.147440957 | 0.007121296090707600 | 0.13466036492 |
| | Braf | -1.23 | 1.395008557 | 0.040270909980230700 | 0.20310713369 |
| | Asic1 | -1.23 | 3.073073359 | 0.000845136076097478 | 0.12354271087 |
| | Slc43a2 | -1.23 | 1.823777716 | 0.015004526130863900 | 0.15143776501 |
| | Glt1d1 | -1.23 | 1.694386423 | 0.020212199535224200 | 0.16092051453 |
| | Smg7 | -1.23 | 2.512499501 | 0.003072560892399400 | 0.12870390352 |
| | Dnajb1 | -1.23 | 1.615999927 | 0.024210294551765700 | 0.16966419499 |
| | Slc25a42 | -1.23 | 1.319600952 | 0.047907007995445200 | 0.21812133760 |
| | Map3k5 | -1.23 | 1.483150575 | 0.032873763368351900 | 0.18984917835 |
| | Pde4a | -1.23 | 2.345057755 | 0.004517958581700010 | 0.13146915442 |
| | Slf2 | -1.23 | 1.639532396 | 0.022933355531171600 | 0.16683129036 |
| | Zfp106 | -1.23 | 2.992654781 | 0.001017056826882220 | 0.12702075696 |
| | Pcyox1l | -1.23 | 1.757775699 | 0.017467240558335300 | 0.15680830472 |
| | Syt7 | -1.23 | 1.874067788 | 0.013363869076123300 | 0.14714176681 |
| | Dffa | -1.23 | 2.036596785 | 0.009191856051706520 | 0.14048876521 |
| | Gga3 | -1.23 | 3.028977358 | 0.000935454443235843 | 0.12702075696 |
| | Vstm5 | -1.23 | 1.39101379 | 0.040643042337154800 | 0.20330985190 |
| | Clstn1 | -1.23 | 1.424298311 | 0.037644513514715100 | 0.19959349860 |
| | Phactr1 | -1.23 | 2.16151659 | 0.006894192572881890 | 0.13451302432 |
| | Mtmr4 | -1.23 | 3.641527779 | 0.000228282290176887 | 0.11984957964 |
| | Snapc5 | -1.23 | 2.149786478 | 0.007082939334886160 | 0.13466036492 |
| | Syt4 | -1.23 | 1.510694031 | 0.030853608805339300 | 0.18539604295 |

| | | | | | |
|---|---|---|---|---|---|
| | Ogdhl | -1.23 | 1.555920787 | 0.027802203173965000 | 0.17888249734 |
| | Sipa1l1 | -1.23 | 2.014252456 | 0.009677151583673220 | 0.14078337569 |
| | Cdr2l | -1.23 | 1.334388797 | 0.046303220988295600 | 0.21494137193 |
| | Rnf150 | -1.23 | 1.52875871 | 0.029596563656535900 | 0.18215557680 |
| | Cbx6 | -1.23 | 1.828826736 | 0.014831096618115500 | 0.15143776501 |
| | Otub2 | -1.23 | 1.591215428 | 0.025632122645771400 | 0.17350295432 |
| | Cntrob | -1.23 | 1.496012076 | 0.031914491099917600 | 0.18710014086 |
| | Mroh1 | -1.23 | 1.726218039 | 0.018783735355292000 | 0.16021558505 |
| | Gatsl2 | -1.23 | 1.744876883 | 0.017993809450337000 | 0.15859713641 |
| | Actr1b | -1.23 | 1.78542239 | 0.016389949283027300 | 0.15299884042 |
| | Ak6 | -1.23 | 1.918900571 | 0.012053118584035200 | 0.14364743987 |
| | Chn1 | -1.23 | 1.515037863 | 0.030546547913274100 | 0.18474476887 |
| | Cep170b | -1.23 | 1.49076323 | 0.032302547193328200 | 0.18831960161 |
| | Cx3cl1 | -1.23 | 2.028881196 | 0.009356615954006960 | 0.14078337569 |
| | Usp31 | -1.23 | 2.243731317 | 0.005705171219362490 | 0.13146915442 |
| | Miga2 | -1.24 | 2.737400766 | 0.001830624348599740 | 0.12702075696 |
| | Cep83 | -1.24 | 1.906333135 | 0.012407002368719900 | 0.14532375495 |
| | Arhgef17 | -1.24 | 1.725052054 | 0.018834233296704100 | 0.16044711701 |
| | Csrnp2 | -1.24 | 1.853556405 | 0.014010176117198600 | 0.14864677641 |
| | Arhgap21 | -1.24 | 2.742373389 | 0.001809783442676260 | 0.12702075696 |
| | Rnf157 | -1.24 | 2.425546915 | 0.003753644029030040 | 0.12985696725 |
| | Wnk2 | -1.24 | 1.992692774 | 0.010169678549659200 | 0.14109528430 |
| | Dusp14 | -1.24 | 1.637171391 | 0.023058370278778900 | 0.16683129036 |

| | | | | | |
|---|---|---|---|---|---|
| | Apc2 | -1.24 | 2.232246839 | 0.005858051173079600 | 0.13146915442 |
| | Adgrl1 | -1.24 | 1.947838298 | 0.011276172279409100 | 0.14193386802 |
| | Ube2ql1 | -1.24 | 1.700165361 | 0.019945027489068400 | 0.16087283358 |
| | Pfas | -1.24 | 1.422876717 | 0.037767938772075900 | 0.19960355641 |
| | Unc80 | -1.24 | 1.709284341 | 0.019530603297447700 | 0.16048239041 |
| | Zfp365 | -1.24 | 1.523119922 | 0.029983344748105300 | 0.18323315564 |
| | Pop5 | -1.24 | 2.026254559 | 0.009413376764059280 | 0.14078337569 |
| | Apba1 | -1.24 | 1.677176639 | 0.021029229496771800 | 0.16261000251 |
| | Pfkfb3 | -1.24 | 1.348508273 | 0.044822051200856500 | 0.21194074520 |
| | Zfp831 | -1.24 | 1.357893493 | 0.043863825659458800 | 0.21061184788 |
| | Zc3h11a | -1.24 | 2.134301758 | 0.007340036878597430 | 0.13611261370 |
| | Arhgef3 | -1.24 | 1.781118646 | 0.016553176803545900 | 0.15347989370 |
| | Inpp4a | -1.24 | 1.926065906 | 0.011855888176870100 | 0.14364743987 |
| | Trak1 | -1.24 | 2.545432663 | 0.002848179374221310 | 0.12870390352 |
| | Odf2 | -1.24 | 1.45497159 | 0.035077481928605200 | 0.19437919645 |
| | Dus3l | -1.24 | 1.781742434 | 0.016529418134322400 | 0.15336308885 |
| | Rapgefl1 | -1.24 | 1.649040707 | 0.022436716095482200 | 0.16638123343 |
| | Gtpbp3 | -1.24 | 1.959219349 | 0.010984509053874800 | 0.14193386802 |
| | Jade1 | -1.24 | 2.540082882 | 0.002883481157976830 | 0.12870390352 |
| | 2900092D14Rik | -1.24 | 1.712401308 | 0.019390932380090400 | 0.16048239041 |
| | Mkl2 | -1.24 | 2.345018446 | 0.004518367529875960 | 0.13146915442 |
| | Nog | -1.24 | 1.462049295 | 0.034510456547698500 | 0.19292440334 |
| | Cers6 | -1.24 | 2.359206271 | 0.004373143504097330 | 0.13146915442 |

| | | | | | |
|---|---|---|---|---|---|
| | Pde8b | -1.25 | 1.952470876 | 0.011156529639237500 | 0.14193386802 |
| | Nyap1 | -1.25 | 2.361331179 | 0.004351798927638650 | 0.13146915442 |
| | Prpf40b | -1.25 | 2.302819261 | 0.004979442698371750 | 0.13146915442 |
| | Ankrd13b | -1.25 | 1.719070359 | 0.019095438719349900 | 0.16048070792 |
| | Atxn2l | -1.25 | 1.708000806 | 0.019588410403397200 | 0.16048239041 |
| | Sez6l | -1.25 | 1.343349045 | 0.045357692778694600 | 0.21288007959 |
| | Chd5 | -1.25 | 1.514284634 | 0.030599572990892900 | 0.18490269678 |
| | Ripor1 | -1.25 | 2.428107895 | 0.003731574401778110 | 0.12985696725 |
| | Mypop | -1.25 | 1.821725609 | 0.015075592549228900 | 0.15152290628 |
| | 9330159F19Rik | -1.25 | 2.387392835 | 0.004098332261999780 | 0.13146915442 |
| | Lmtk3 | -1.25 | 1.952162562 | 0.011164452704017400 | 0.14193386802 |
| | Thy1 | -1.25 | 1.644870374 | 0.022653203454637200 | 0.16645864635 |
| | Socs7 | -1.25 | 1.939069669 | 0.011506157931250400 | 0.14243794246 |
| | Dpysl2 | -1.25 | 2.54209842 | 0.002870130077178060 | 0.12870390352 |
| | Chka | -1.25 | 2.352323861 | 0.004442998217092420 | 0.13146915442 |
| | Prickle2 | -1.25 | 2.338215496 | 0.004589702161982970 | 0.13146915442 |
| | Rufy2 | -1.25 | 2.254954201 | 0.005559628834629470 | 0.13146915442 |
| | Cplx2 | -1.25 | 2.068931388 | 0.008532349014391650 | 0.14043458802 |
| | Chgb | -1.25 | 2.424670012 | 0.003761230839411490 | 0.12985696725 |
| | Arhgef2 | -1.25 | 2.215699294 | 0.006085562210645550 | 0.13146915442 |
| | Crtc1 | -1.25 | 2.336134296 | 0.004611749441574920 | 0.13146915442 |
| | Lrrtm2 | -1.25 | 2.401821273 | 0.003964411501878480 | 0.13146915442 |
| | Prr14l | -1.25 | 1.866194405 | 0.013608353886914300 | 0.14796465545 |

| | | | | | |
|---|---|---|---|---|---|
| | Srf | -1.25 | 1.757674089 | 0.017471327744026300 | 0.15680830472 |
| | Mical3 | -1.25 | 2.576701069 | 0.002650323764890700 | 0.12734763009 |
| | Fbxl19 | -1.25 | 2.393102921 | 0.004044800248769210 | 0.13146915442 |
| | Eml5 | -1.25 | 1.755275096 | 0.017568104415862400 | 0.15708531271 |
| | Pou3f2 | -1.25 | 1.403206496 | 0.039517867786510800 | 0.20216833599 |
| | Oprd1 | -1.25 | 1.460740513 | 0.034614613553709300 | 0.19327078620 |
| | Elovl4 | -1.25 | 2.299642382 | 0.005016001041167350 | 0.13146915442 |
| | Rundc1 | -1.25 | 2.46173409 | 0.003453551290989740 | 0.12936217305 |
| | Gramd1b | -1.25 | 2.266395006 | 0.005415081462608540 | 0.13146915442 |
| | 5730409E04Rik | -1.25 | 1.801120957 | 0.015808077017102400 | 0.15212233233 |
| | Disp2 | -1.25 | 2.565816124 | 0.002717589627702300 | 0.12870390352 |
| | Esrra | -1.25 | 1.970738116 | 0.010696997236786900 | 0.14150160435 |
| | Zbtb7a | -1.26 | 2.620929234 | 0.002393705768644930 | 0.12702075696 |
| | Klhl17 | -1.26 | 2.727465617 | 0.001872985359405800 | 0.12702075696 |
| | Prrg3 | -1.26 | 1.700622754 | 0.019924032716270600 | 0.16087283358 |
| | Mlxip | -1.26 | 2.231199032 | 0.005872201751762130 | 0.13146915442 |
| | Iqsec1 | -1.26 | 1.685651784 | 0.020622827825728300 | 0.16185943212 |
| | Wipf2 | -1.26 | 1.834333831 | 0.014644217463239200 | 0.15107071484 |
| | Diras1 | -1.26 | 2.696676663 | 0.002010589159053250 | 0.12702075696 |
| | Ciz1 | -1.26 | 1.963166532 | 0.010885126182828400 | 0.14193386802 |
| | Sez6l2 | -1.26 | 1.658432178 | 0.021956738136289400 | 0.16477746517 |
| | Elfn2 | -1.26 | 1.665132639 | 0.021620581012585600 | 0.16399478955 |
| | B230217C12Rik | -1.26 | 1.546139095 | 0.028435502346843000 | 0.17947407154 |

| | | | | | |
|---|---|---|---|---|---|
| | Sncb | -1.26 | 1.906013632 | 0.012416133346557900 | 0.14532375495 |
| | Tomm34 | -1.26 | 2.494313918 | 0.003203952594778130 | 0.12870390352 |
| | Shf | -1.26 | 1.517690148 | 0.030360565185105000 | 0.18410096624 |
| | Fam53b | -1.26 | 1.71535995 | 0.019259280110813900 | 0.16048239041 |
| | Zfp382 | -1.26 | 1.398700332 | 0.039930032887646800 | 0.20241237663 |
| | Auts2 | -1.26 | 2.319456881 | 0.004792290307249950 | 0.13146915442 |
| | Pcgf6 | -1.26 | 1.68948687 | 0.020441517344988200 | 0.16152207581 |
| | Add2 | -1.26 | 3.146651436 | 0.000713425394030185 | 0.12354271087 |
| | Gpd1 | -1.26 | 1.855597934 | 0.013944471724863300 | 0.14862753048 |
| | Lrrc45 | -1.26 | 1.878091184 | 0.013240635079678100 | 0.14686395323 |
| | Ppp1r37 | -1.26 | 2.096640209 | 0.008004971512783730 | 0.13756078109 |
| | Brsk1 | -1.26 | 1.625517889 | 0.023685475677535400 | 0.16819747870 |
| | Dlgap3 | -1.26 | 1.469560701 | 0.033918707864512600 | 0.19184362522 |
| | Diexf | -1.26 | 2.155229257 | 0.006994726585843250 | 0.13466036492 |
| | Ociad2 | -1.26 | 2.33945149 | 0.004576658520880100 | 0.13146915442 |
| | Syndig1 | -1.26 | 1.536061603 | 0.029103042743977000 | 0.18061208843 |
| | Cntn2 | -1.26 | 1.499762983 | 0.031640039519652500 | 0.18675506144 |
| | Cacna1i | -1.26 | 1.715000992 | 0.019275205086687400 | 0.16048239041 |
| | Mapk8ip3 | -1.26 | 2.232855185 | 0.005849851137027160 | 0.13146915442 |
| | Tacc1 | -1.26 | 2.97066967 | 0.001069868324395140 | 0.12702075696 |
| | Gal3st3 | -1.26 | 1.480603679 | 0.033067116151555500 | 0.19035410266 |
| | Fn3krp | -1.26 | 1.813082679 | 0.015378618420219300 | 0.15174905791 |
| | Matk | -1.26 | 1.611756322 | 0.024448019205726600 | 0.16982619623 |

| | | | | | |
|---|---|---|---|---|---|
| | Foxp1 | -1.26 | 1.39296886 | 0.040460490157881100 | 0.20327882825 |
| | Vars2 | -1.26 | 2.364938925 | 0.004315797658464090 | 0.13146915442 |
| | Foxo3 | -1.26 | 1.554775757 | 0.027875601238593700 | 0.17888249734 |
| | Rab15 | -1.26 | 1.970008623 | 0.010714980295661000 | 0.14150160435 |
| | Slc45a1 | -1.27 | 2.361611018 | 0.004348995734335830 | 0.13146915442 |
| | Mafg | -1.27 | 1.761193379 | 0.017330321579132400 | 0.15671857076 |
| | Hivep2 | -1.27 | 1.482513342 | 0.032922033930697100 | 0.18984917835 |
| | Nap1l2 | -1.27 | 1.760257479 | 0.017367708487448900 | 0.15671857076 |
| | D17Wsu92e | -1.27 | 2.306147216 | 0.004941431551048000 | 0.13146915442 |
| | Epb41l1 | -1.27 | 1.977832137 | 0.010523685543564200 | 0.14109528430 |
| | Cdh7 | -1.27 | 1.301494588 | 0.049946540391892800 | 0.22163429645 |
| | Zbtb17 | -1.27 | 2.274612118 | 0.005313588053858980 | 0.13146915442 |
| | Abcc5 | -1.27 | 2.797561421 | 0.001593817457498330 | 0.12702075696 |
| | Trim66 | -1.27 | 1.340116625 | 0.045696546024998300 | 0.21359986354 |
| | Ints6l | -1.27 | 2.157214346 | 0.006962827785409210 | 0.13466036492 |
| | Dync1h1 | -1.27 | 2.62160696 | 0.002389973249211110 | 0.12702075696 |
| | Syne1 | -1.27 | 1.975043203 | 0.010591483562146300 | 0.14143593355 |
| | Kcnh3 | -1.27 | 1.609429648 | 0.024579347613758900 | 0.16997725997 |
| | Ablim1 | -1.27 | 3.12660515 | 0.000747127719762863 | 0.12354271087 |
| | R3hdm1 | -1.27 | 1.341861219 | 0.045513347741005500 | 0.21337390355 |
| | Ttc9 | -1.27 | 2.439018853 | 0.003638992382993150 | 0.12985696725 |
| | Ptprj | -1.27 | 3.645172723 | 0.000226374381531262 | 0.11984957964 |
| | Map4k2 | -1.27 | 2.219829945 | 0.006027955741504980 | 0.13146915442 |

| | | | | | |
|---|---|---|---|---|---|
| | Rbm33 | -1.27 | 2.143450668 | 0.007187027912157920 | 0.13530366435 |
| | Prkcb | -1.27 | 1.537225883 | 0.029025126214113600 | 0.18036289823 |
| | Capn15 | -1.27 | 2.077518624 | 0.008365297226676500 | 0.14038435593 |
| | Avpi1 | -1.27 | 2.183206279 | 0.006558336870797370 | 0.13306070174 |
| | Adgrf5 | -1.27 | 1.855329367 | 0.013953097614005100 | 0.14862753048 |
| | Ccdc141 | -1.27 | 1.988010218 | 0.010279921107095700 | 0.14109528430 |
| | Frmd5 | -1.27 | 1.732827087 | 0.018500050458640200 | 0.15996749329 |
| | Ddx55 | -1.27 | 2.265400353 | 0.005427497687846660 | 0.13146915442 |
| | Map3k13 | -1.27 | 1.470076578 | 0.033878441409472800 | 0.19181032691 |
| | Itpr1 | -1.27 | 1.815173345 | 0.015304764634065600 | 0.15159717255 |
| | Rusc1 | -1.27 | 2.367124915 | 0.004294128978153080 | 0.13146915442 |
| | Camta2 | -1.27 | 1.446926735 | 0.035733311464051400 | 0.19629181303 |
| | Zfp651 | -1.27 | 2.593234561 | 0.002551322969191990 | 0.12702075696 |
| | Rtn4rl1 | -1.27 | 1.520768872 | 0.030146099477434600 | 0.18341471643 |
| | Slc2a13 | -1.27 | 1.738505814 | 0.018259723042610300 | 0.15920485681 |
| | Ppm1h | -1.27 | 2.055112148 | 0.008808213891368040 | 0.14048876521 |
| | Numbl | -1.27 | 1.420130365 | 0.038007528986297800 | 0.20006989915 |
| | Fbxo41 | -1.27 | 2.214921443 | 0.006096471623758860 | 0.13146915442 |
| | Cacna1b | -1.27 | 1.572707642 | 0.026748064246271300 | 0.17639876384 |
| | Adam11 | -1.27 | 1.598092589 | 0.025229428405236900 | 0.17170756598 |
| | Lmtk2 | -1.27 | 1.606034873 | 0.024772231310292000 | 0.17028275660 |
| | Rimbp2 | -1.27 | 1.387695203 | 0.040954798811017300 | 0.20403256309 |
| | Fam160b2 | -1.27 | 2.546551514 | 0.002840851198812490 | 0.12870390352 |

| | | | | | |
|---|---|---|---|---|---|
| | Csnk1e | -1.27 | 2.615938266 | 0.002421373217768610 | 0.12702075696 |
| | Spred1 | -1.27 | 2.404187414 | 0.003942871164105900 | 0.13146915442 |
| | Luzp2 | -1.27 | 1.463961504 | 0.034358840222774600 | 0.19283460548 |
| | Rita1 | -1.27 | 2.395138005 | 0.004025890839629710 | 0.13146915442 |
| | Tmem198 | -1.28 | 1.475063232 | 0.033491667268484700 | 0.19112913454 |
| | AI854703 | -1.28 | 1.525018071 | 0.029852584013035200 | 0.18308227454 |
| | Unc79 | -1.28 | 2.919407649 | 0.001203905367625580 | 0.12702075696 |
| | Asap1 | -1.28 | 1.347289766 | 0.044947985662306800 | 0.21195108826 |
| | Mpp3 | -1.28 | 2.615272424 | 0.002425088409978710 | 0.12702075696 |
| | Ankzf1 | -1.28 | 2.166360376 | 0.006817727271992020 | 0.13451302432 |
| | Dixdc1 | -1.28 | 2.623028979 | 0.002382160509589780 | 0.12702075696 |
| | Cplx1 | -1.28 | 1.465751542 | 0.034217514311925400 | 0.19240257979 |
| | Ncor2 | -1.28 | 1.860272328 | 0.013795189558576300 | 0.14797790767 |
| | Rbfox1 | -1.28 | 1.729644112 | 0.018636136726330100 | 0.15996749329 |
| | Ank3 | -1.28 | 1.4961442 1 | 0.031904782608819900 | 0.18710014086 |
| | Rilpl1 | -1.28 | 1.356467978 | 0.044008039504728900 | 0.21069946177 |
| | Ahdc1 | -1.28 | 2.075827808 | 0.008397928867979300 | 0.14038435593 |
| | Gabbr2 | -1.28 | 1.325249433 | 0.047287958742024700 | 0.21713627016 |
| | Trpc3 | -1.28 | 1.533165176 | 0.029297787460951100 | 0.18127734780 |
| | Clvs1 | -1.28 | 2.141335212 | 0.007222121462286470 | 0.13538768215 |
| | Limk1 | -1.28 | 2.056359614 | 0.008782949502610230 | 0.14048876521 |
| | Dio2 | -1.28 | 1.614089822 | 0.024317010233993300 | 0.16966419499 |
| | Srcin1 | -1.28 | 2.178854512 | 0.006624383817494120 | 0.13324073206 |

| | | | | | |
|---|---|---|---|---|---|
| | Pde5a | -1.28 | 1.419648656 | 0.038049709363126900 | 0.20006989915 |
| | Trim37 | -1.28 | 1.545366547 | 0.028486130085969600 | 0.17947407154 |
| | Rpusd1 | -1.28 | 1.550868204 | 0.028127542884845500 | 0.17943387501 |
| | Fbxl18 | -1.28 | 1.796025098 | 0.015994655935837600 | 0.15212233233 |
| | Kcnc2 | -1.28 | 1.486842041 | 0.032595523409826200 | 0.18898120305 |
| | Spns2 | -1.28 | 2.040146594 | 0.009117030471927340 | 0.14048876521 |
| | Snph | -1.28 | 1.970216954 | 0.010709841556354500 | 0.14150160435 |
| | Sfswap | -1.28 | 1.628512663 | 0.023522709055803800 | 0.16796885012 |
| | Grip2 | -1.28 | 3.210620031 | 0.000615715333612546 | 0.12354271087 |
| | 2900026A02Rik | -1.28 | 1.708715186 | 0.019556215499916300 | 0.16048239041 |
| | Rap1gap | -1.28 | 1.89237421 | 0.012812261384724700 | 0.14588139267 |
| | Jph3 | -1.28 | 1.588075506 | 0.025818112813145400 | 0.17390523930 |
| | Phyhip | -1.28 | 2.082003605 | 0.008279352909969430 | 0.13976239353 |
| | Slc12a5 | -1.28 | 2.057803213 | 0.008753803359969580 | 0.14048876521 |
| | Mdn1 | -1.28 | 2.414688416 | 0.003848678058530500 | 0.13025784533 |
| | Pip5k1c | -1.28 | 1.613207321 | 0.024366473487966600 | 0.16978687231 |
| | Nbas | -1.28 | 1.584038105 | 0.026059248938317300 | 0.17473771883 |
| | Mapk4 | -1.28 | 2.014517379 | 0.009671250244349900 | 0.14078337569 |
| | Slc35f3 | -1.29 | 2.297959434 | 0.005035476410423780 | 0.13146915442 |
| | Bean1 | -1.29 | 1.305039276 | 0.049540538577307900 | 0.22130890818 |
| | St8sia1 | -1.29 | 1.822239207 | 0.015057774627506700 | 0.15152290628 |
| | Smpd4 | -1.29 | 1.864856804 | 0.013650331427909400 | 0.14796465545 |
| | Clstn3 | -1.29 | 2.176359431 | 0.006662551340343330 | 0.13345498246 |

| | | | | | |
|---|---|---|---|---|---|
| | Atp2b3 | -1.29 | 2.467264 6 | 0.00340985098669 4230 | 0.12936217 305 |
| | Mycn | -1.29 | 1.3727257 03 | 0.04239106201260 9600 | 0.20740196 527 |
| | Chga | -1.29 | 2.9874212 06 | 0.00102938727177 4580 | 0.12702075 696 |
| | Dopey2 | -1.29 | 3.1964786 27 | 0.00063609410760 6383 | 0.12354271 087 |
| | Jade2 | -1.29 | 2.0977621 65 | 0.00798431816822 1460 | 0.13756078 109 |
| | Pqlc1 | -1.29 | 1.3095177 17 | 0.04903230193971 0900 | 0.22050961 871 |
| | Dcbld2 | -1.29 | 1.7441383 39 | 0.01802443504637 2400 | 0.15866352 465 |
| | Zmym1 | -1.29 | 1.6842446 43 | 0.02068975542317 2500 | 0.16185943 212 |
| | Clk1 | -1.29 | 2.0767684 63 | 0.00837975915417 6810 | 0.14038435 593 |
| | Cables1 | -1.29 | 2.1373752 43 | 0.00728827509694 0460 | 0.13564617 473 |
| | D630045J12 Rik | -1.29 | 2.1685145 89 | 0.00678399331136 4630 | 0.13451302 432 |
| | Sema4f | -1.29 | 2.3272887 6 | 0.00470664280281 8760 | 0.13146915 442 |
| | Cdk5r2 | -1.29 | 2.2683158 44 | 0.00539118402200 9530 | 0.13146915 442 |
| | Clasrp | -1.29 | 1.5204757 22 | 0.03016645507061 2400 | 0.18341471 643 |
| | Map7d2 | -1.29 | 2.2628310 64 | 0.00545970196474 6120 | 0.13146915 442 |
| | Med13l | -1.29 | 2.4163251 21 | 0.00383420102572 8800 | 0.13025784 533 |
| | Ryr2 | -1.30 | 1.4381915 79 | 0.03645930798617 1100 | 0.19742455 646 |
| | Rcan2 | -1.30 | 1.3343898 54 | 0.04630310829019 8300 | 0.21494137 193 |
| | Cecr6 | -1.30 | 2.1024308 44 | 0.00789894621146 8060 | 0.13703952 374 |
| | Peg3 | -1.30 | 2.6384854 64 | 0.00229887065425 8700 | 0.12702075 696 |
| | Frmpd3 | -1.30 | 1.5540748 95 | 0.02792062300603 8100 | 0.17888249 734 |
| | Car7 | -1.30 | 1.5697464 73 | 0.02693106495920 2300 | 0.17663950 530 |
| | Acap3 | -1.30 | 2.0151237 49 | 0.00965775649495 0590 | 0.14078337 569 |

| | | | | | |
|---|---|---|---|---|---|
| | Camkk2 | -1.30 | 1.633326538 | 0.023263414652106100 | 0.16734067081 |
| | Shd | -1.30 | 1.3594572 | 0.043706174942243400 | 0.21035605950 |
| | Zgpat | -1.30 | 1.914741874 | 0.012169090642936800 | 0.14387163261 |
| | Otud3 | -1.30 | 1.742341589 | 0.018099159672797200 | 0.15898298674 |
| | Iqsec3 | -1.30 | 1.378581746 | 0.041823295828669400 | 0.20572143528 |
| | Lrp3 | -1.30 | 2.556111829 | 0.002778997592850260 | 0.12870390352 |
| | Klf9 | -1.30 | 2.876444951 | 0.001329092016718520 | 0.12702075696 |
| | Bcor | -1.31 | 1.627232936 | 0.023592125183066900 | 0.16797991605 |
| | Atmin | -1.31 | 2.376206513 | 0.004205266149145650 | 0.13146915442 |
| | Nab2 | -1.31 | 1.975698805 | 0.010575506950193200 | 0.14140280189 |
| | Lrrc3b | -1.31 | 1.740020496 | 0.018196149806571700 | 0.15903535871 |
| | Ankrd27 | -1.31 | 2.658143863 | 0.002197131937928360 | 0.12702075696 |
| | Rian | -1.31 | 2.4314013 | 0.003703383607925690 | 0.12985696725 |
| | Tnk2 | -1.31 | 2.366078454 | 0.004304488439524450 | 0.13146915442 |
| | Adrb1 | -1.31 | 1.712727993 | 0.019376351635997500 | 0.16048239041 |
| | Klc2 | -1.31 | 1.661939304 | 0.021780141461899300 | 0.16443550615 |
| | Hdac4 | -1.31 | 2.541104323 | 0.002876707309426850 | 0.12870390352 |
| | Kcnq2 | -1.31 | 2.494214429 | 0.003204686645268470 | 0.12870390352 |
| | Myo5a | -1.31 | 1.773040586 | 0.016863954207203400 | 0.15479465248 |
| | Tbr1 | -1.31 | 1.69734421 | 0.020075010902318000 | 0.16092051453 |
| | Spred2 | -1.31 | 1.399044994 | 0.039898356406826400 | 0.20241237663 |
| | Csmd2 | -1.31 | 2.597910764 | 0.002523999334966390 | 0.12702075696 |
| | Noc2l | -1.31 | 2.103956121 | 0.007871253138826570 | 0.13703952374 |

| | | | | | |
|---|---|---|---|---|---|
| | Kcnt1 | -1.31 | 2.609161743 | 0.002459451466105600 | 0.12702075696 |
| | Fbxo31 | -1.31 | 2.055782394 | 0.008794630660450160 | 0.14048876521 |
| | Grik3 | -1.31 | 3.031707326 | 0.000929592635282453 | 0.12702075696 |
| | Homer1 | -1.31 | 1.610874889 | 0.024497688673248700 | 0.16983992939 |
| | Nktr | -1.31 | 2.65275261 | 0.002224576728136180 | 0.12702075696 |
| | Brinp1 | -1.31 | 1.545360959 | 0.028486496618628000 | 0.17947407154 |
| | Kmt2a | -1.31 | 1.490331433 | 0.032334679946384400 | 0.18834711197 |
| | Fam43b | -1.31 | 1.454700938 | 0.035099349065288800 | 0.19439748307 |
| | Map1a | -1.32 | 1.991861791 | 0.010189155952498200 | 0.14109528430 |
| | Asphd1 | -1.32 | 1.722714305 | 0.018935888831907100 | 0.16046894179 |
| | Sptbn2 | -1.32 | 1.736156548 | 0.018358764566583800 | 0.15937095786 |
| | Pou6f1 | -1.32 | 1.57650725 | 0.026515068310030500 | 0.17576522503 |
| | Pcnx2 | -1.32 | 1.957873802 | 0.011018594440710900 | 0.14193386802 |
| | Acin1 | -1.32 | 2.073291851 | 0.008447109988170140 | 0.14043458802 |
| | Dmtn | -1.32 | 2.131384332 | 0.007389510452050360 | 0.13628710184 |
| | Synj2 | -1.32 | 1.379064117 | 0.041776868478124500 | 0.20572143528 |
| | Stx1a | -1.32 | 1.446457823 | 0.035771913880031600 | 0.19630266319 |
| | Cpeb1 | -1.32 | 2.331673136 | 0.004659366407018810 | 0.13146915442 |
| | Sbk1 | -1.32 | 2.349383597 | 0.004473180289716210 | 0.13146915442 |
| | Enc1 | -1.32 | 2.148039623 | 0.007111486285476440 | 0.13466036492 |
| | Adora1 | -1.32 | 2.207948375 | 0.006195147131075020 | 0.13146915442 |
| | Fgf9 | -1.32 | 1.623167279 | 0.023814020408044700 | 0.16858756024 |
| | Fzd4 | -1.32 | 1.383928315 | 0.041311568583234400 | 0.20515441413 |

| | | | | | |
|---|---|---|---|---|---|
| | Hectd4 | -1.32 | 3.421504546 | 0.000378874568250960 | 0.12354271087 |
| | Lypd6 | -1.32 | 1.615492818 | 0.024238580457203600 | 0.16966419499 |
| | Lrrc49 | -1.32 | 2.768628877 | 0.001703613696967500 | 0.12702075696 |
| | Robo4 | -1.32 | 1.431456627 | 0.037029118487562600 | 0.19853187451 |
| | Tet3 | -1.32 | 2.50579161 | 0.003120386500384650 | 0.12870390352 |
| | Pagr1a | -1.32 | 2.76897562 | 0.001702254064146700 | 0.12702075696 |
| | Rell2 | -1.32 | 1.750603253 | 0.017758110215688300 | 0.15755428328 |
| | Slc36a1 | -1.33 | 2.944329037 | 0.001136765706115840 | 0.12702075696 |
| | Pcdhb3 | -1.33 | 1.577413869 | 0.026459774027039700 | 0.17556619480 |
| | Git1 | -1.33 | 1.96195515 | 0.010915530558550400 | 0.14193386802 |
| | D130043K22Rik | -1.33 | 1.848814543 | 0.014163984975748000 | 0.14902551114 |
| | Sidt1 | -1.33 | 1.951189371 | 0.011189498658177700 | 0.14193386802 |
| | Plcb4 | -1.33 | 1.442745832 | 0.036078973105035300 | 0.19681983497 |
| | Shank2 | -1.33 | 1.818281757 | 0.015195613657568600 | 0.15152290628 |
| | Lnx1 | -1.33 | 3.359511011 | 0.000437007599096855 | 0.12354271087 |
| | Bsn | -1.33 | 1.996949914 | 0.010070478015412900 | 0.14109528430 |
| | Prrt2 | -1.33 | 1.828930258 | 0.014827561760854200 | 0.15143776501 |
| | Cry2 | -1.33 | 2.663575239 | 0.002169825259909530 | 0.12702075696 |
| | Chrm1 | -1.33 | 2.689564817 | 0.002043784883652140 | 0.12702075696 |
| | Kctd1 | -1.33 | 3.773480549 | 0.000168468787993190 | 0.11984957964 |
| | Ccdc92b | -1.33 | 3.096428775 | 0.000800886964150339 | 0.12354271087 |
| | Srrm4 | -1.33 | 1.498820276 | 0.031708793968899300 | 0.18676431809 |
| | Zfp983 | -1.33 | 1.823838511 | 0.015002425859057500 | 0.15143776501 |

| | | | | | |
|---|---|---|---|---|---|
| | E4f1 | -1.33 | 1.646437303 | 0.022571618295236400 | 0.16645864635 |
| | Grin1 | -1.33 | 2.309980155 | 0.004898012000678940 | 0.13146915442 |
| | Tsc22d3 | -1.33 | 1.996732651 | 0.010075517197881500 | 0.14109528430 |
| | Snrnp70 | -1.33 | 2.069324308 | 0.008524633018358640 | 0.14043458802 |
| | Notch4 | -1.33 | 1.34046245 | 0.045660172731767500 | 0.21355222379 |
| | Cacna1a | -1.33 | 1.703828546 | 0.019777502775324400 | 0.16067121778 |
| | Ngef | -1.33 | 2.110795656 | 0.007748262834335930 | 0.13702558508 |
| | Tmem151b | -1.33 | 2.265279478 | 0.005429008507694080 | 0.13146915442 |
| | Adgrb1 | -1.33 | 2.530321722 | 0.002949023804237080 | 0.12870390352 |
| | Dab2ip | -1.34 | 2.222541445 | 0.005990437693368110 | 0.13146915442 |
| | Ggt7 | -1.34 | 2.814045879 | 0.001534454874697380 | 0.12702075696 |
| | Kcnip2 | -1.34 | 1.981047469 | 0.010446060352525700 | 0.14109528430 |
| | Syt12 | -1.34 | 2.251741033 | 0.005600914814943200 | 0.13146915442 |
| | Znrf1 | -1.34 | 2.891272496 | 0.001284480466138260 | 0.12702075696 |
| | Entpd7 | -1.34 | 1.598586044 | 0.025200778433994500 | 0.17159757010 |
| | Mn1 | -1.34 | 2.92113652 | 0.001199122301750280 | 0.12702075696 |
| | Sptbn4 | -1.34 | 3.247949626 | 0.000565002506395465 | 0.12354271087 |
| | Camta1 | -1.34 | 1.870971633 | 0.013459482657938800 | 0.14751589520 |
| | Prrt3 | -1.34 | 1.750632068 | 0.017756932033966200 | 0.15755428328 |
| | Csmd1 | -1.34 | 2.006630456 | 0.009848487627059430 | 0.14078337569 |
| | Igf1r | -1.34 | 3.957046437 | 0.000110396057326945 | 0.11276116861 |
| | Plch2 | -1.34 | 2.811298444 | 0.001544192915803530 | 0.12702075696 |
| | Speg | -1.34 | 3.170681025 | 0.000675023627684268 | 0.12354271087 |

| | | | | | |
|---|---|---|---|---|---|
| | Brsk2 | -1.34 | 1.72870693 | 0.018676395823560300 | 0.15996749329 |
| | Cacnb1 | -1.34 | 2.699355574 | 0.001998225173039860 | 0.12702075696 |
| | Amer2 | -1.34 | 2.202967736 | 0.006266604178512740 | 0.13206964420 |
| | Shank3 | -1.34 | 2.432220676 | 0.003696403084615310 | 0.12985696725 |
| | Dennd6b | -1.34 | 1.870535795 | 0.013472996759554700 | 0.14751589520 |
| | Klhl34 | -1.34 | 2.475314585 | 0.003347228917708270 | 0.12870390352 |
| | Fam193b | -1.34 | 1.805054144 | 0.015665557531347800 | 0.15202007489 |
| | Dbp | -1.34 | 2.850860525 | 0.001409741468105530 | 0.12702075696 |
| | Vcpkmt | -1.34 | 1.990380215 | 0.010223975149272200 | 0.14109528430 |
| | Ankrd33b | -1.35 | 1.548022059 | 0.028312481853532800 | 0.17947407154 |
| | Efna3 | -1.35 | 1.724218838 | 0.018870402415933500 | 0.16046894179 |
| | Ccdc159 | -1.35 | 1.639548373 | 0.022932511880356800 | 0.16683129036 |
| | Camk2a | -1.35 | 1.415019268 | 0.038457471943544400 | 0.20070038814 |
| | Rims4 | -1.35 | 1.587443774 | 0.025855695574324900 | 0.17407305874 |
| | Adam23 | -1.35 | 1.644992502 | 0.022646834052680300 | 0.16645864635 |
| | Kcnab2 | -1.35 | 2.100243965 | 0.007938821466152140 | 0.13721678713 |
| | Tnks1bp1 | -1.35 | 1.974061895 | 0.010615442566770700 | 0.14148088876 |
| | Map3k9 | -1.35 | 1.808907015 | 0.015527194194006700 | 0.15174905791 |
| | Syt13 | -1.35 | 1.969513164 | 0.010727211312427300 | 0.14150160435 |
| | Kcnj9 | -1.35 | 1.318767071 | 0.047999081765525200 | 0.21825403979 |
| | Hpca | -1.35 | 1.325759782 | 0.047232422178322700 | 0.21706378366 |
| | Prodh | -1.35 | 1.542973093 | 0.028643554266522300 | 0.17988464033 |
| | C130074G19Rik | -1.35 | 1.882178268 | 0.013116613818508600 | 0.14665206711 |

| | | | | | |
|---|---|---|---|---|---|
| | Hs3st4 | -1.35 | 1.923787729 | 0.011918243950235100 | 0.14364743987 |
| | Pitpnm2 | -1.35 | 2.951103026 | 0.001119172354929320 | 0.12702075696 |
| | Adcy9 | -1.35 | 2.356170268 | 0.004403821748236440 | 0.13146915442 |
| | Etv5 | -1.35 | 3.258872716 | 0.000550969152312654 | 0.12354271087 |
| | Ankrd9 | -1.35 | 1.39139407 | 0.040607469762316500 | 0.20330985190 |
| | Tbc1d30 | -1.35 | 1.700750188 | 0.019918187291610200 | 0.16087283358 |
| | Scn8a | -1.35 | 1.623489964 | 0.023796332890489700 | 0.16855128742 |
| | Tle2 | -1.35 | 1.377518255 | 0.041925837362962700 | 0.20585780012 |
| | Gm996 | -1.36 | 2.460938318 | 0.003459885145095430 | 0.12936217305 |
| | Prickle1 | -1.36 | 1.803588628 | 0.015718509882269100 | 0.15212233233 |
| | Mast1 | -1.36 | 2.510019849 | 0.003090154198739700 | 0.12870390352 |
| | Mcf2l | -1.36 | 4.715717008 | 0.000019243452509147 | 0.06610607023 |
| | Nrg3 | -1.36 | 2.207140454 | 0.006206682725605750 | 0.13146915442 |
| | Gpt | -1.36 | 2.853331522 | 0.001401743263464310 | 0.12702075696 |
| | Slc29a2 | -1.36 | 2.315950626 | 0.004831137231414660 | 0.13146915442 |
| | Zfp692 | -1.36 | 1.46977945 | 0.033901627673899400 | 0.19184362522 |
| | Lhx6 | -1.36 | 1.995127661 | 0.010112821437341700 | 0.14109528430 |
| | Nfic | -1.37 | 1.783451328 | 0.016464504743396300 | 0.15327829247 |
| | Mfsd2a | -1.37 | 2.305045805 | 0.004953979379186120 | 0.13146915442 |
| | Cish | -1.37 | 1.544560059 | 0.028539078174935600 | 0.17972294831 |
| | Sertad1 | -1.37 | 1.444505086 | 0.035933118874628300 | 0.19669619782 |
| | Ccdc184 | -1.37 | 1.962840143 | 0.010893309853607100 | 0.14193386802 |
| | Hrh3 | -1.37 | 2.175639721 | 0.006673601639246190 | 0.13346292906 |

| | | | | | |
|---|---|---|---|---|---|
| | Lpcat4 | -1.37 | 2.485376217 | 0.003270572517478800 | 0.12870390352 |
| | Dpf1 | -1.37 | 1.955376046 | 0.011082148216125200 | 0.14193386802 |
| | Tmem25 | -1.37 | 1.886773655 | 0.012978555103544700 | 0.14628049103 |
| | Panx2 | -1.37 | 2.317832103 | 0.004810252753143470 | 0.13146915442 |
| | Cdc7 | -1.37 | 1.588828844 | 0.025773366876240900 | 0.17368898197 |
| | Inha | -1.37 | 2.300468003 | 0.005006474368468430 | 0.13146915442 |
| | Nphp4 | -1.37 | 2.103018924 | 0.007888257451808750 | 0.13703952374 |
| | Ccdc92 | -1.37 | 2.556241855 | 0.002778165698348220 | 0.12870390352 |
| | A930015D03Rik | -1.37 | 1.323857666 | 0.047439743721236500 | 0.21728983949 |
| | Ftx | -1.37 | 1.528687145 | 0.029601441160798600 | 0.18215557680 |
| | Kalrn | -1.37 | 1.691579214 | 0.020343271062743700 | 0.16130230102 |
| | Jag2 | -1.37 | 2.264846929 | 0.005434418385939030 | 0.13146915442 |
| | Cit | -1.37 | 3.705394829 | 0.000197063036585440 | 0.11984957964 |
| | Dmpk | -1.37 | 1.544134801 | 0.028567037083424100 | 0.17981660860 |
| | Osbpl6 | -1.37 | 2.309090544 | 0.004908055396421350 | 0.13146915442 |
| | Pdp1 | -1.37 | 2.641036437 | 0.002285407051925580 | 0.12702075696 |
| | Unc13a | -1.37 | 2.363848335 | 0.004326649008226690 | 0.13146915442 |
| | Slc25a37 | -1.37 | 2.62444066 | 0.002374429833946180 | 0.12702075696 |
| | Nat8l | -1.38 | 1.80932128 | 0.015512390187257000 | 0.15174905791 |
| | Srrm2 | -1.38 | 2.393158862 | 0.004044279274730830 | 0.13146915442 |
| | Plekha6 | -1.38 | 2.80512321 | 0.001566306643289070 | 0.12702075696 |
| | Nptx1 | -1.38 | 2.126772101 | 0.007468405646381520 | 0.13628710184 |
| | Extl1 | -1.38 | 1.955410909 | 0.011081258612952100 | 0.14193386802 |

| | | | | | |
|---|---|---|---|---|---|
| | Sh3rf3 | -1.38 | 1.90224503 | 0.012524343503895300 | 0.14564091316 |
| | Plcl2 | -1.38 | 2.270587825 | 0.005363054066608150 | 0.13146915442 |
| | Ece1 | -1.38 | 3.374188581 | 0.000422485121334359 | 0.12354271087 |
| | Kcnb1 | -1.38 | 2.847973703 | 0.001419143448911060 | 0.12702075696 |
| | Adamts17 | -1.38 | 1.39260851 | 0.040494075651790100 | 0.20329890155 |
| | Prdm8 | -1.38 | 1.325179249 | 0.047295601236839100 | 0.21713627016 |
| | Mast3 | -1.38 | 1.896368284 | 0.012694971079400400 | 0.14576593494 |
| | Nuak1 | -1.38 | 1.653868193 | 0.022188697378846100 | 0.16526067964 |
| | Ttc14 | -1.38 | 3.276350709 | 0.000529235892967766 | 0.12354271087 |
| | Them6 | -1.38 | 1.643753818 | 0.022711519011011500 | 0.16678647237 |
| | Spata2l | -1.38 | 2.24185535 | 0.005729868438391930 | 0.13146915442 |
| | Il17ra | -1.39 | 2.034793867 | 0.009230094199381860 | 0.14048876521 |
| | Cntnap1 | -1.39 | 2.726702452 | 0.001876279563069000 | 0.12702075696 |
| | Arhgef25 | -1.39 | 1.982448849 | 0.010412407380462500 | 0.14109528430 |
| | Fam57b | -1.39 | 2.331373134 | 0.004662586114665370 | 0.13146915442 |
| | Synpo | -1.39 | 2.647200953 | 0.002253196391710010 | 0.12702075696 |
| | Pakap | -1.39 | 2.018520851 | 0.009582507083867890 | 0.14078337569 |
| | Fam43a | -1.39 | 1.698868169 | 0.020004690228900600 | 0.16092051453 |
| | Fbxo27 | -1.39 | 1.463387825 | 0.034404256318260400 | 0.19283467894 |
| | Nos1ap | -1.39 | 1.645899438 | 0.022599590077451500 | 0.16645864635 |
| | Tspoap1 | -1.40 | 1.800405962 | 0.015834123877625800 | 0.15212233233 |
| | Lynx1 | -1.40 | 1.696765603 | 0.020101774529398500 | 0.16092051453 |
| | Trerf1 | -1.40 | 3.638181159 | 0.000230048201003184 | 0.11984957964 |

| | | | | | |
|---|---|---|---|---|---|
| | Epha10 | -1.40 | 2.308699636 | 0.004912475116295620 | 0.13146915442 |
| | Lingo1 | -1.40 | 1.622888535 | 0.023829309874617900 | 0.16860893254 |
| | Rbfox3 | -1.40 | 2.594970622 | 0.002541144593828600 | 0.12702075696 |
| | Ksr2 | -1.40 | 2.328692875 | 0.004691450353236130 | 0.13146915442 |
| | Hecw1 | -1.40 | 1.358875594 | 0.043764745333317800 | 0.21049050249 |
| | Mccc1os | -1.40 | 1.331685014 | 0.046592389699636800 | 0.21565324658 |
| | Adamtsl5 | -1.40 | 1.438276972 | 0.036452139857649000 | 0.19742455646 |
| | Neurod2 | -1.40 | 2.137789923 | 0.007281319318855560 | 0.13564617473 |
| | Clstn2 | -1.40 | 1.699811245 | 0.019961296940189000 | 0.16087283358 |
| | Zswim4 | -1.40 | 2.198711406 | 0.006328322360743330 | 0.13211194818 |
| | Akap8l | -1.40 | 2.753458289 | 0.001764175193651620 | 0.12702075696 |
| | Hsf4 | -1.41 | 1.952728837 | 0.011149904883389300 | 0.14193386802 |
| | Srrm3 | -1.41 | 2.747939739 | 0.001786735479304030 | 0.12702075696 |
| | Rasgrp4 | -1.41 | 1.865402624 | 0.013633186494405500 | 0.14796465545 |
| | Ttbk1 | -1.41 | 2.876632646 | 0.001328517728306050 | 0.12702075696 |
| | Luzp1 | -1.41 | 2.815413502 | 0.001529630370075900 | 0.12702075696 |
| | Arhgap39 | -1.41 | 2.421343866 | 0.003790147698935080 | 0.13025784533 |
| | Unc5a | -1.41 | 3.179759478 | 0.000661059456554119 | 0.12354271087 |
| | Mapk6 | -1.41 | 1.941457526 | 0.011443067865313800 | 0.14204082704 |
| | Wdr90 | -1.41 | 1.610619388 | 0.024512105228549200 | 0.16985417950 |
| | Zfhx2 | -1.41 | 3.312167895 | 0.000487340051612562 | 0.12354271087 |
| | Camk2n1 | -1.41 | 1.367129923 | 0.042940794632893500 | 0.20849804207 |
| | Rnf165 | -1.41 | 1.782028419 | 0.016518537021917600 | 0.15336308885 |

| | | | | | |
|---|---|---|---|---|---|
| | Shank1 | -1.42 | 1.693359319 | 0.020260057837351500 | 0.16101414387 |
| | Carmil2 | -1.42 | 2.026284845 | 0.009412720316632600 | 0.14078337569 |
| | Dusp7 | -1.42 | 3.173748978 | 0.000670271914781631 | 0.12354271087 |
| | Slc30a2 | -1.42 | 1.955752267 | 0.011072552119691200 | 0.14193386802 |
| | Nsun5 | -1.42 | 2.077506151 | 0.008365537467880660 | 0.14038435593 |
| | 11-Mar | -1.42 | 1.393310091 | 0.040428712285938400 | 0.20327882825 |
| | Tchh | -1.42 | 2.221942181 | 0.005998709337685130 | 0.13146915442 |
| | Pla2g4e | -1.42 | 1.369605292 | 0.042696739098750500 | 0.20784410277 |
| | Trmt44 | -1.42 | 1.451607159 | 0.035350278570855900 | 0.19547210376 |
| | Malat1 | -1.43 | 3.243423011 | 0.000570922276598487 | 0.12354271087 |
| | Gm9899 | -1.43 | 1.862507901 | 0.013724359918514900 | 0.14796465545 |
| | Tarbp1 | -1.43 | 1.791382303 | 0.016166562911710400 | 0.15237084595 |
| | Lrfn2 | -1.43 | 1.361086293 | 0.043542534778268000 | 0.20986249400 |
| | Kcnk3 | -1.43 | 2.72878967 | 0.001867283800691310 | 0.12702075696 |
| | Cdr1 | -1.43 | 2.062034183 | 0.008668936411065140 | 0.14048876521 |
| | Ephb3 | -1.43 | 2.014106072 | 0.009680413925277520 | 0.14078337569 |
| | Ccdc57 | -1.43 | 1.328475588 | 0.046937981795473400 | 0.21643449928 |
| | Ina | -1.43 | 4.111533627 | 0.000077351078345636 | 0.11276116861 |
| | Hlf | -1.43 | 1.934492295 | 0.011628071798938600 | 0.14317323888 |
| | Camkk1 | -1.43 | 2.040657286 | 0.009106315948307010 | 0.14048876521 |
| | 9330102E08Rik | -1.43 | 2.196876811 | 0.006355111708653110 | 0.13211194818 |
| | Bicdl1 | -1.43 | 2.60729401 | 0.002470051393238020 | 0.12702075696 |
| | Car4 | -1.44 | 1.474021674 | 0.033572085946814300 | 0.19130833143 |

| | | | | | |
|---|---|---|---|---|---|
| | Pdgfb | -1.44 | 1.640492766 | 0.022882698246824100 | 0.16683129036 |
| | Zmynd8 | -1.44 | 4.430345367 | 0.000037123988787007 | 0.08502012165 |
| | Rhobtb2 | -1.44 | 4.046781457 | 0.000089788050713414 | 0.11276116861 |
| | Dlc1 | -1.44 | 3.102398733 | 0.000789953025185523 | 0.12354271087 |
| | C030018K13Rik | -1.44 | 1.459523349 | 0.034711761320808100 | 0.19373448916 |
| | Gp1bb | -1.44 | 1.885112725 | 0.013028285733673500 | 0.14631261435 |
| | Banp | -1.44 | 1.922111304 | 0.011964338613383100 | 0.14364743987 |
| | Sik2 | -1.44 | 1.589168898 | 0.025753194169897500 | 0.17363819484 |
| | Vwa5b2 | -1.44 | 1.476895444 | 0.033350669448455100 | 0.19112913454 |
| | Atp10a | -1.44 | 2.774458125 | 0.001680899991807030 | 0.12702075696 |
| | Gse1 | -1.44 | 3.364287554 | 0.000432227550478145 | 0.12354271087 |
| | Klhl29 | -1.44 | 3.693651005 | 0.000202464551304462 | 0.11984957964 |
| | Nrsn1 | -1.45 | 1.407660552 | 0.039114649918981400 | 0.20180964127 |
| | C77080 | -1.45 | 1.701663901 | 0.019876325445462900 | 0.16075373040 |
| | Fgd5 | -1.45 | 1.996349054 | 0.010084420487216700 | 0.14109528430 |
| | Kcnc1 | -1.45 | 2.431917815 | 0.003698981719404630 | 0.12985696725 |
| | Cacng2 | -1.45 | 2.105093252 | 0.007850670464098120 | 0.13703952374 |
| | Hspa1b | -1.45 | 1.555380001 | 0.027836844206685300 | 0.17888249734 |
| | Tinagl1 | -1.45 | 1.681265998 | 0.020832145606565200 | 0.16236784616 |
| | Mrs2 | -1.46 | 1.419843769 | 0.038032618800776100 | 0.20006989915 |
| | Astn2 | -1.46 | 2.264528406 | 0.005438405592704190 | 0.13146915442 |
| | Atp2b2 | -1.46 | 1.806774236 | 0.015603634309350800 | 0.15188371222 |
| | Aifm3 | -1.46 | 1.907538326 | 0.012372619982646600 | 0.14518545788 |

| | | | | | |
|---|---|---|---|---|---|
| | Stac2 | -1.46 | 1.337610013 | 0.045961054824393800 | 0.21430297059 |
| | Kndc1 | -1.47 | 2.34017802 | 0.004569008647823840 | 0.13146915442 |
| | Zbed6 | -1.47 | 2.65875552 | 0.002194039691250770 | 0.12702075696 |
| | Gls2 | -1.47 | 1.919657056 | 0.012032141882056200 | 0.14364743987 |
| | Tub | -1.47 | 3.450403511 | 0.000354483878476640 | 0.12354271087 |
| | Cd209c | -1.47 | 1.446828848 | 0.035741366435633800 | 0.19629181303 |
| | Nefm | -1.47 | 1.830129736 | 0.014786666030245800 | 0.15143776501 |
| | Nefh | -1.47 | 1.595335508 | 0.025390104690989400 | 0.17237422360 |
| | Sptb | -1.47 | 3.267399 | 0.000540257743541001 | 0.12354271087 |
| | Snapc4 | -1.47 | 2.610255449 | 0.002453265495220920 | 0.12702075696 |
| | Snhg14 | -1.48 | 2.007933686 | 0.009818978615738550 | 0.14078337569 |
| | Nrgn | -1.48 | 1.517498626 | 0.030373957006333100 | 0.18410096624 |
| | Mapk11 | -1.48 | 1.770075083 | 0.016979500768495300 | 0.15554354671 |
| | Rnf112 | -1.48 | 2.68940285 | 0.002044547240620960 | 0.12702075696 |
| | Sowahb | -1.49 | 1.468963202 | 0.033965405082406200 | 0.19190733192 |
| | Nr1d1 | -1.49 | 2.36114251 | 0.004353689872407620 | 0.13146915442 |
| | Ciart | -1.49 | 2.532446471 | 0.002934631192846000 | 0.12870390352 |
| | Epop | -1.49 | 1.722795432 | 0.018932351911154500 | 0.16046894179 |
| | Mertk | -1.50 | 4.90011322 | 0.000012585972534158 | 0.05764794953 |
| | Kctd16 | -1.50 | 1.641145611 | 0.022848326125666600 | 0.16683129036 |
| | Rtn4r | -1.50 | 1.717145173 | 0.019180274895211800 | 0.16048239041 |
| | Hr | -1.50 | 1.453855692 | 0.035167727687720400 | 0.19461930977 |
| | Vasn | -1.51 | 1.954344161 | 0.011108510721984300 | 0.14193386802 |

| | | | | | |
|---|---|---|---|---|---|
| | Sgsm1 | -1.51 | 1.590927143 | 0.025649142874918700 | 0.17353267959 |
| | Lime1 | -1.51 | 2.296351892 | 0.005054149783931390 | 0.13146915442 |
| | Dusp1 | -1.51 | 1.437954468 | 0.036479219041568300 | 0.19742455646 |
| | Lrrc14b | -1.52 | 2.663071016 | 0.002172345926641020 | 0.12702075696 |
| | Fmn1 | -1.52 | 1.44068729 | 0.036250392195092300 | 0.19727391650 |
| | 4933439C10Rik | -1.53 | 1.756434239 | 0.017521277234776900 | 0.15705144846 |
| | Sema3f | -1.53 | 1.790138208 | 0.016212940611611500 | 0.15251082377 |
| | Pvalb | -1.53 | 1.304632817 | 0.049586925652657300 | 0.22130890818 |
| | Gfod1 | -1.53 | 2.68891469 | 0.002046846668267630 | 0.12702075696 |
| | Serpinb8 | -1.53 | 2.007661102 | 0.009825143414138870 | 0.14078337569 |
| | Scrt1 | -1.53 | 1.853094428 | 0.014025087272781800 | 0.14864677641 |
| | Cux2 | -1.53 | 2.544720509 | 0.002852853636120140 | 0.12870390352 |
| | Kcns1 | -1.53 | 1.672709875 | 0.021246633412606600 | 0.16273689505 |
| | Chrm4 | -1.53 | 3.727796544 | 0.000187155871140574 | 0.11984957964 |
| | Spag5 | -1.53 | 1.839334503 | 0.014476564061226700 | 0.15044156813 |
| | Tmem181b-ps | -1.54 | 1.302920697 | 0.049782798156383900 | 0.22145206522 |
| | Trank1 | -1.54 | 3.62801806 | 0.000235495135027173 | 0.11984957964 |
| | Kcnk12 | -1.54 | 1.581838624 | 0.026191560581704500 | 0.17473771883 |
| | Mtcl1 | -1.54 | 3.720270745 | 0.000190427319743057 | 0.11984957964 |
| | Cdc25b | -1.54 | 1.69587524 | 0.020143028173584600 | 0.16092051453 |
| | Cabp1 | -1.54 | 2.232797779 | 0.005850624434337890 | 0.13146915442 |
| | Mtus2 | -1.55 | 2.697693379 | 0.002005887726713880 | 0.12702075696 |
| | Lzts3 | -1.55 | 4.194664299 | 0.000063875704170158 | 0.10971450638 |

| | | | | | |
|---|---|---|---|---|---|
| | Slc25a34 | -1.55 | 1.454916876 | 0.035081901404838700 | 0.19437919645 |
| | Bhlhe40 | -1.55 | 1.750353511 | 0.017768324996279200 | 0.15755428328 |
| | Galnt9 | -1.55 | 1.69793345 | 0.020047792097534900 | 0.16092051453 |
| | Sstr2 | -1.56 | 3.064022782 | 0.000862933279136401 | 0.12481648620 |
| | Hipk4 | -1.56 | 2.270887335 | 0.005359356717071850 | 0.13146915442 |
| | Ankrd24 | -1.56 | 2.650346191 | 0.002236937290732670 | 0.12702075696 |
| | Ank1 | -1.56 | 2.115950862 | 0.007656832355030440 | 0.13628710184 |
| | Dnajb5 | -1.56 | 3.389158929 | 0.000408169990123288 | 0.12354271087 |
| | Cbfa2t3 | -1.57 | 2.722298832 | 0.001895401273235770 | 0.12702075696 |
| | Celsr3 | -1.57 | 3.168508604 | 0.000678408679471483 | 0.12354271087 |
| | Rgs11 | -1.57 | 3.16202017 | 0.000688620314274778 | 0.12354271087 |
| | Phospho1 | -1.57 | 1.6706209 | 0.021349076831078500 | 0.16305711054 |
| | Rims3 | -1.57 | 1.457682244 | 0.034859227317401900 | 0.19416321142 |
| | Dok3 | -1.57 | 1.72187672 | 0.018972444036371900 | 0.16048070792 |
| | Myo5c | -1.57 | 1.462294075 | 0.034491011032909300 | 0.19289417281 |
| | Pygm | -1.57 | 1.505405083 | 0.031231649154713400 | 0.18582135954 |
| | Kcnmb4os2 | -1.58 | 1.463404825 | 0.034402909683046600 | 0.19283467894 |
| | Osbpl3 | -1.58 | 2.580761124 | 0.002625662346674400 | 0.12734763009 |
| | Tmem145 | -1.58 | 1.645944726 | 0.022597233548162500 | 0.16645864635 |
| | Camk1g | -1.59 | 1.339039825 | 0.045809987664955800 | 0.21386329830 |
| | Arid3b | -1.59 | 2.76333889 | 0.001724491706597180 | 0.12702075696 |
| | Zbtb40 | -1.59 | 3.204205382 | 0.000624877112229142 | 0.12354271087 |
| | Plekhg5 | -1.59 | 2.458514414 | 0.003479249599537500 | 0.12936217305 |

| | | | | | |
|---|---|---|---|---|---|
| | Map3k6 | -1.60 | 1.427095394 | 0.037402842284820100 | 0.19928037673 |
| | Dagla | -1.60 | 3.162711166 | 0.000687525536342074 | 0.12354271087 |
| | D030047H15Rik | -1.60 | 1.467802581 | 0.034056296536446400 | 0.19202608564 |
| | Fcrls | -1.60 | 1.401279285 | 0.039693620645310100 | 0.20241237663 |
| | Slc2a4rg-ps | -1.60 | 1.799793517 | 0.015856468996605300 | 0.15212233233 |
| | Fgf5 | -1.60 | 1.567360598 | 0.027079422723390400 | 0.17718968935 |
| | Trhr2 | -1.61 | 1.408339554 | 0.039053543563818800 | 0.20180964127 |
| | Ephb2 | -1.61 | 3.443096004 | 0.000360498943081844 | 0.12354271087 |
| | Jdp2 | -1.61 | 2.692657424 | 0.002029282805764640 | 0.12702075696 |
| | Chrm2 | -1.61 | 1.983034046 | 0.010398386467878400 | 0.14109528430 |
| | Zfp57 | -1.61 | 1.471190939 | 0.033791623756087500 | 0.19179293764 |
| | Plk3 | -1.62 | 3.89124553 | 0.000128456022359367 | 0.11767428022 |
| | Gm14827 | -1.62 | 1.411737143 | 0.038749210368866500 | 0.20115334329 |
| | Per1 | -1.62 | 2.038235927 | 0.009157228956197660 | 0.14048876521 |
| | Ccl28 | -1.62 | 1.309437451 | 0.049041364877400500 | 0.22050961871 |
| | 2410018L13Rik | -1.63 | 2.016796078 | 0.009620639065962330 | 0.14078337569 |
| | Has3 | -1.63 | 2.023922646 | 0.009464057156364520 | 0.14078337569 |
| | Egr1 | -1.63 | 1.720164541 | 0.019047389326618500 | 0.16048070792 |
| | 2900055J20Rik | -1.63 | 1.455724896 | 0.035016691048053900 | 0.19429017089 |
| | Tspan11 | -1.64 | 1.418456962 | 0.038154260281044300 | 0.20017366189 |
| | Kirrel2 | -1.64 | 1.716303692 | 0.019217474241679500 | 0.16048239041 |
| | Tmem86b | -1.64 | 1.431794363 | 0.037000333371986300 | 0.19853187451 |
| | Vamp1 | -1.64 | 1.540241594 | 0.028824275880102200 | 0.17995201039 |

| | | | | | |
|---|---|---|---|---|---|
| | Satb2 | -1.65 | 1.638332245 | 0.022996818346637000 | 0.16683129036 |
| | Cyp4f15 | -1.65 | 2.120523844 | 0.007576631342461810 | 0.13628710184 |
| | Abcc8 | -1.66 | 2.5810406 | 0.002623973231878040 | 0.12734763009 |
| | Epha8 | -1.67 | 2.280186988 | 0.005245815505948510 | 0.13146915442 |
| | Myo19 | -1.67 | 1.69009082 | 0.020413110211605100 | 0.16148333185 |
| | Igsf9b | -1.68 | 3.287273167 | 0.000516091649995626 | 0.12354271087 |
| | Tcap | -1.68 | 1.406147856 | 0.039251128204669400 | 0.20197941444 |
| | Chrd | -1.68 | 2.177756384 | 0.006641154999459550 | 0.13324073206 |
| | Kcnab3 | -1.68 | 1.559671176 | 0.027563148461066500 | 0.17859523991 |
| | Kcnc3 | -1.68 | 2.231467822 | 0.005868568497150970 | 0.13146915442 |
| | Fmnl1 | -1.69 | 3.384080771 | 0.000412970689440053 | 0.12354271087 |
| | Catsperz | -1.69 | 1.583949741 | 0.026064551679056800 | 0.17473771883 |
| | Klhl33 | -1.70 | 2.288405366 | 0.005147479598867480 | 0.13146915442 |
| | Hsbp1l1 | -1.71 | 1.369577958 | 0.042699426491674000 | 0.20784410277 |
| | Dnajc21 | -1.71 | 2.437365711 | 0.003652870611836450 | 0.12985696725 |
| | Col19a1 | -1.71 | 2.002230157 | 0.009948780351951980 | 0.14109528430 |
| | Psrc1 | -1.71 | 2.498242755 | 0.003175098811438480 | 0.12870390352 |
| | Kcnh4 | -1.71 | 2.246747474 | 0.005665686317687880 | 0.13146915442 |
| | Vipr1 | -1.71 | 3.364132281 | 0.000432382112444280 | 0.12354271087 |
| | Tbx3 | -1.72 | 1.472150458 | 0.033717047816078100 | 0.19153582404 |
| | Snhg11 | -1.72 | 2.859843511 | 0.001380881748102440 | 0.12702075696 |
| | Plec | -1.72 | 1.631791206 | 0.023345801795708900 | 0.16751816558 |
| | 2010111I01Rik | -1.72 | 4.031194866 | 0.000093069018520189 | 0.11276116861 |

| | | | | | |
|---|---|---|---|---|---|
| | Proser2 | -1.73 | 1.336727549 | 0.046054540220400400 | 0.21452048718 |
| | Cpne9 | -1.75 | 2.012861244 | 0.009708200914549660 | 0.14078337569 |
| | 9430037G07Rik | -1.75 | 1.553432037 | 0.027961982738112300 | 0.17888249734 |
| | Gadd45a | -1.76 | 2.797759939 | 0.001593089079941100 | 0.12702075696 |
| | Trim17 | -1.77 | 2.669004678 | 0.002142867518937170 | 0.12702075696 |
| | Adam33 | -1.77 | 1.571376516 | 0.026830173671759400 | 0.17639876384 |
| | Ier5 | -1.79 | 4.470807704 | 0.000033821455715184 | 0.08502012165 |
| | Rnf223 | -1.80 | 1.354798152 | 0.044177572489819900 | 0.21092058116 |
| | 4933428G20Rik | -1.80 | 1.488288485 | 0.032487142656339900 | 0.18859561776 |
| | Ccer2 | -1.80 | 1.400726732 | 0.039744154978700400 | 0.20241237663 |
| | D330050G23Rik | -1.80 | 1.360565968 | 0.043594733966341500 | 0.20996678564 |
| | Meg3 | -1.80 | 2.467066208 | 0.003411409006371380 | 0.12936217305 |
| | Aoc2 | -1.80 | 2.451581759 | 0.003535234613003970 | 0.12954042351 |
| | Syt2 | -1.80 | 1.942924879 | 0.011404470360820400 | 0.14196161768 |
| | Tbc1d2 | -1.81 | 2.773847674 | 0.001683264351507550 | 0.12702075696 |
| | Zfp85os | -1.82 | 2.587045785 | 0.002587940067622080 | 0.12734763009 |
| | A330023F24Rik | -1.82 | 3.512586408 | 0.000307194610734970 | 0.12354271087 |
| | Il12a | -1.86 | 1.693575776 | 0.020249962547378500 | 0.16101414387 |
| | Kcnj14 | -1.87 | 2.674654996 | 0.002115168663101620 | 0.12702075696 |
| | Mirg | -1.87 | 3.305381178 | 0.000495015527399041 | 0.12354271087 |
| | Robo3 | -1.88 | 1.679029611 | 0.020939696823263000 | 0.16246887298 |
| | BC030499 | -1.88 | 1.895049038 | 0.012733592928625200 | 0.14587849176 |
| | Phf21b | -1.89 | 2.23744824 | 0.005788309701849640 | 0.13146915442 |

| | | | | | |
|---|---|---|---|---|---|
| | Rnf39 | -1.90 | 1.407303356 | 0.039146833989983300 | 0.20184489526 |
| | Serinc2 | -1.92 | 3.018273672 | 0.000958796253701395 | 0.12702075696 |
| | Tmem200b | -1.92 | 2.968155551 | 0.001076079726877270 | 0.12702075696 |
| | Nphs1 | -1.93 | 1.490044572 | 0.032356044777707300 | 0.18838393165 |
| | Col24a1 | -1.97 | 1.950302158 | 0.011212380867579300 | 0.14193386802 |
| | Miat | -2.00 | 2.635430033 | 0.002315101132048240 | 0.12702075696 |
| | Gpr3 | -2.01 | 3.380528451 | 0.000416362443403200 | 0.12354271087 |
| | Coro6 | -2.01 | 3.025635229 | 0.000942681036330567 | 0.12702075696 |
| | Per2 | -2.04 | 3.182896953 | 0.000656300971888273 | 0.12354271087 |
| | Rtl1 | -2.05 | 1.564782874 | 0.027240628657908700 | 0.17799024174 |
| | Mir124a-1hg | -2.15 | 3.783138731 | 0.000164763598678915 | 0.11984957964 |
| | BC002163 | -2.17 | 2.165678458 | 0.006828440688619860 | 0.13451302432 |
| | Plekhg4 | -2.19 | 2.118368849 | 0.007614320456245580 | 0.13628710184 |
| | Grin1os | -2.19 | 3.086253081 | 0.000819873630906080 | 0.12354271087 |
| | Dnase1l2 | -2.27 | 2.591309978 | 0.002562654288754300 | 0.12712430535 |
| | Alox12b | -2.29 | 1.799039734 | 0.015884014191478000 | 0.15212233233 |
| | Zbtb16 | -2.35 | 2.876392401 | 0.001329252847513210 | 0.12702075696 |
| | Arc | -2.37 | 3.015092401 | 0.000965845361942554 | 0.12702075696 |
| | Gm13830 | -2.46 | 2.678739796 | 0.002095367502594090 | 0.12702075696 |
| | Chn1os3 | -2.57 | 2.360363103 | 0.004361510243533220 | 0.13146915442 |
| | Six4 | -2.94 | 2.910913738 | 0.001227683055021350 | 0.12702075696 |
| | Tnfrsf25 | -3.01 | 2.574208438 | 0.002665579022182740 | 0.12762272245 |
| | Fam227b | -3.23 | 1.39949042 | 0.039857456432702500 | 0.20241237663 |

| | | | | | |
|---|---|---|---|---|---|
| | Gh | -3.25 | 1.340852918 | 0.045619138742993400 | 0.21346021711 |
| | Pomc | -3.42 | 1.713856551 | 0.019326065583706800 | 0.16048239041 |
| | Prl | -6.64 | 2.095093625 | 0.008033529178611810 | 0.13764180105 |
