## Supplementary material for "BRAFV600E Expression in Mouse Neuroglial Progenitors Increase Neuronal Excitability, Cause Appearance of Balloon-like cells, Neuronal Mislocalization, and Inflammatory Immune response": Sup. Table 3. DE genes in BRAFwt to control-FP

| UP p<0.05 | Gene ID | Fold change (BRAFwt vs. control-FP) | log-10 P-values | P-value | FDR step up |
|---|---|---|---|---|---|
| | C3 | 8.49E+01 | 1.561382357645 | 0.0274547594 | 0.35570414585 |
| | C4b | 1.21E+01 | 1.444399121843 | 0.0359418873 | 0.37716131557 |
| | A2m | 8.88E+00 | 2.432021004077 | 0.0036981029 | 0.22143440510 |
| | Gfap | 7.99E+00 | 1.779052342932 | 0.0166321218 | 0.32205632977 |
| | Plin4 | 6.01E+00 | 2.783803853598 | 0.0016451146 | 0.19673992847 |
| | Serpina3n | 5.27E+00 | 1.407810369416 | 0.0391011590 | 0.38989997216 |
| | Serpina3h | 4.72E+00 | 1.450565757269 | 0.0354351473 | 0.37517732932 |
| | Prl | 4.35E+00 | 1.512668571167 | 0.0307136499 | 0.36169282125 |
| | Ggta1 | 4.26E+00 | 2.175041706300 | 0.0066827974 | 0.25823683896 |
| | Cybrd1 | 4.06E+00 | 1.697028469017 | 0.0200896112 | 0.34145893141 |
| | S100a4 | 3.99E+00 | 2.203238129451 | 0.0062627038 | 0.25527338269 |
| | Gadl1 | 3.54E+00 | 1.540272785960 | 0.0288222057 | 0.35982823218 |
| | BC002163 | 3.50E+00 | 2.812585441475 | 0.0015396236 | 0.19673992847 |
| | Plscr4 | 3.32E+00 | 1.593235343740 | 0.0255131837 | 0.35302467272 |
| | Ccr1 | 3.31E+00 | 1.864130818740 | 0.0136731690 | 0.30416998758 |
| | Osmr | 3.29E+00 | 1.613158833506 | 0.0243691941 | 0.34940735647 |
| | Angpt1 | 3.25E+00 | 1.620998632574 | 0.0239332329 | 0.34940735647 |
| | Vim | 3.13E+00 | 1.595515302574 | 0.0253795956 | 0.35302467272 |
| | Aspg | 3.11E+00 | 1.816065403724 | 0.0152733603 | 0.31840303817 |
| | Slc14a1 | 3.03E+00 | 1.976366603831 | 0.0105592579 | 0.28670727701 |
| | Gm28042 | 2.91E+00 | 2.047206702615 | 0.0089700176 | 0.27519811275 |
| | Mgst1 | 2.84E+00 | 1.598009135753 | 0.0252342769 | 0.35302467272 |

| | | | | | |
|---|---|---|---|---|---|
| | Txlnb | 2.79E+00 | 1.508659230312 | 0.0309985065 | 0.36235522207 |
| | Fxyd1 | 2.78E+00 | 1.762040357455 | 0.0172965562 | 0.32736724913 |
| | Tnnt2 | 2.78E+00 | 1.940340169862 | 0.0114725466 | 0.28966274854 |
| | Meox1 | 2.75E+00 | 2.093906560183 | 0.0080555174 | 0.26995021897 |
| | Plce1 | 2.74E+00 | 1.614257924051 | 0.0243075997 | 0.34940735647 |
| | Tmem252 | 2.71E+00 | 1.557185129986 | 0.0277213815 | 0.35681741541 |
| | Paqr6 | 2.63E+00 | 1.582063299802 | 0.0261780143 | 0.35302467272 |
| | Gm1653 | 2.58E+00 | 1.856775232588 | 0.0139067218 | 0.30438581455 |
| | Stac3 | 2.47E+00 | 1.910116424955 | 0.0122993901 | 0.29368070731 |
| | Fas | 2.45E+00 | 2.163290053016 | 0.0068660972 | 0.25895522490 |
| | Pipox | 2.41E+00 | 1.700502604598 | 0.0199295455 | 0.34145893141 |
| | Trim34a | 2.35E+00 | 1.984771887518 | 0.0103568602 | 0.28214902540 |
| | Mt2 | 2.33E+00 | 3.327358709589 | 0.0004705885 | 0.15651963832 |
| | Gm5643 | 2.31E+00 | 1.878607602084 | 0.0132249000 | 0.29799400500 |
| | Dnase1l1 | 2.30E+00 | 2.004288791047 | 0.0099017329 | 0.27933254996 |
| | Cort | 2.27E+00 | 1.459958970286 | 0.0346769610 | 0.37517732932 |
| | Slc37a2 | 2.25E+00 | 1.635427384680 | 0.0231511525 | 0.34928420415 |
| | Cthrc1 | 2.22E+00 | 1.775287170378 | 0.0167769430 | 0.32205632977 |
| | 2700038G22Rik | 2.22E+00 | 2.732591230673 | 0.0018510100 | 0.19673992847 |
| | Col16a1 | 2.21E+00 | 1.669070062860 | 0.0214254493 | 0.34484103836 |
| | Fgf1 | 2.19E+00 | 1.965082230623 | 0.0108372170 | 0.28743859126 |
| | Prrx2 | 2.15E+00 | 1.357790325056 | 0.0438742469 | 0.40581230583 |
| | Mroh5 | 2.14E+00 | 1.893965632327 | 0.0127653982 | 0.29799400500 |

| | | | | | |
|---|---|---|---|---|---|
| | Ifi208 | 2.10E+00 | 2.403398457504 | 0.0039500404 | 0.22301019851 |
| | Cntd1 | 2.07E+00 | 1.550471900427 | 0.0281532217 | 0.35778052517 |
| | Ly6a | 2.04E+00 | 1.490899937000 | 0.0322923806 | 0.36612491461 |
| | Slc22a4 | 2.04E+00 | 1.613199498867 | 0.0243669124 | 0.34940735647 |
| | D030028A08Rik | 2.04E+00 | 1.859213194983 | 0.0138288735 | 0.30421038291 |
| | Cd38 | 2.03E+00 | 1.612012693504 | 0.0244335914 | 0.34940735647 |
| | Gm17750 | 2.01E+00 | 2.598474242445 | 0.0025207267 | 0.20681751624 |
| | Nostrin | 1.99E+00 | 2.722022362828 | 0.0018966083 | 0.19737886765 |
| | Trim16 | 1.99E+00 | 2.753194342318 | 0.0017652477 | 0.19673992847 |
| | Gm973 | 1.97E+00 | 1.477639496144 | 0.0332935806 | 0.37160615371 |
| | 2810459M11Rik | 1.97E+00 | 1.535652273984 | 0.0291304857 | 0.35982823218 |
| | Itih3 | 1.96E+00 | 2.021773297917 | 0.0095110114 | 0.27933254996 |
| | Slc2a4 | 1.96E+00 | 1.388011349603 | 0.0409249964 | 0.39782718298 |
| | Naprt | 1.96E+00 | 1.347041202287 | 0.0449737186 | 0.40639873371 |
| | Pm20d1 | 1.96E+00 | 1.552653988677 | 0.0280121221 | 0.35760698238 |
| | Mterf1b | 1.95E+00 | 1.612294758928 | 0.0244177274 | 0.34940735647 |
| | Tmie | 1.95E+00 | 1.589265585745 | 0.0257474613 | 0.35302467272 |
| | Zfp185 | 1.95E+00 | 2.025901024449 | 0.0094210428 | 0.27883484032 |
| | Igfbp3 | 1.94E+00 | 1.515452566119 | 0.0305173932 | 0.36169282125 |
| | Pla1a | 1.93E+00 | 1.487168889852 | 0.0325710013 | 0.36793168158 |
| | Acer2 | 1.92E+00 | 1.701118941526 | 0.0199012822 | 0.34145893141 |
| | Xlr | 1.92E+00 | 1.415248859542 | 0.0384371466 | 0.38774483584 |
| | Synpo2 | 1.92E+00 | 1.333965842059 | 0.0463483372 | 0.41182214750 |

| | | | | | |
|---|---|---|---|---|---|
| | Ifit3 | 1.92E+00 | 1.515744426690 | 0.0304968914 | 0.36169282125 |
| | Gsap | 1.91E+00 | 2.402378571428 | 0.0039593275 | 0.22301019851 |
| | Esyt3 | 1.91E+00 | 1.776065720000 | 0.0167468943 | 0.32205632977 |
| | Tmem176a | 1.90E+00 | 1.800528453578 | 0.0158296585 | 0.32047634791 |
| | Ttc41 | 1.90E+00 | 1.329060246723 | 0.0468748351 | 0.41286871138 |
| | Klf4 | 1.90E+00 | 2.406592918169 | 0.0039210924 | 0.22301019851 |
| | Mt1 | 1.89E+00 | 3.326932109087 | 0.0004710510 | 0.15651963832 |
| | Gch1 | 1.89E+00 | 2.218314186174 | 0.0060490311 | 0.25265694675 |
| | F2r | 1.89E+00 | 1.489813715687 | 0.0323732488 | 0.36628764513 |
| | Dera | 1.88E+00 | 1.905995607840 | 0.0124166486 | 0.29549462766 |
| | Aldh1l2 | 1.86E+00 | 1.621401844018 | 0.0239110229 | 0.34940735647 |
| | Pros1 | 1.86E+00 | 1.944808471739 | 0.0113551148 | 0.28910669889 |
| | BC026585 | 1.85E+00 | 1.596717749741 | 0.0253094233 | 0.35302467272 |
| | Dapp1 | 1.83E+00 | 1.327819678194 | 0.0470089252 | 0.41298385768 |
| | C4a | 1.83E+00 | 1.344943884515 | 0.0451914333 | 0.40639873371 |
| | Cyp4f14 | 1.82E+00 | 3.857326176801 | 0.0001388909 | 0.12414735192 |
| | Abcc4 | 1.82E+00 | 2.713853689762 | 0.0019326193 | 0.19737886765 |
| | Kcnj16 | 1.82E+00 | 1.430623827164 | 0.0371001933 | 0.38314395894 |
| | Apobec3 | 1.82E+00 | 1.353458615193 | 0.0443140440 | 0.40590171168 |
| | Chn1os3 | 1.81E+00 | 1.588728603820 | 0.0257793164 | 0.35302467272 |
| | AI506816 | 1.81E+00 | 1.550881892096 | 0.0281266564 | 0.35776698947 |
| | Spin4 | 1.81E+00 | 1.895332131951 | 0.0127252953 | 0.29799400500 |
| | Neat1 | 1.81E+00 | 2.170562211454 | 0.0067520833 | 0.25823683896 |

| | | | | | |
|---|---|---|---|---|---|
| | Maff | 1.80E+00 | 1.443793884183 | 0.0359920112 | 0.37740502073 |
| | Il13ra1 | 1.80E+00 | 1.740002457972 | 0.0181969056 | 0.33250133049 |
| | Cd14 | 1.80E+00 | 1.386680766185 | 0.0410505740 | 0.39791751326 |
| | Lgals1 | 1.80E+00 | 2.072464384899 | 0.0084632197 | 0.27170265038 |
| | Park2 | 1.78E+00 | 2.630985357383 | 0.0023389161 | 0.20580985051 |
| | Sntb1 | 1.78E+00 | 1.321292171036 | 0.0477208125 | 0.41341683535 |
| | Sdc4 | 1.78E+00 | 2.663489127892 | 0.0021702555 | 0.20573435897 |
| | Tmem86b | 1.78E+00 | 2.049700271534 | 0.0089186625 | 0.27519811275 |
| | Rida | 1.77E+00 | 1.687904288199 | 0.0205161427 | 0.34343029754 |
| | Etohd2 | 1.77E+00 | 2.158328263299 | 0.0069449918 | 0.25914424154 |
| | Pxmp2 | 1.77E+00 | 1.490707742945 | 0.0323066746 | 0.36612491461 |
| | Mr1 | 1.77E+00 | 2.350301128164 | 0.0044637398 | 0.23131440987 |
| | Bmpr1b | 1.76E+00 | 1.634294598131 | 0.0232116173 | 0.34928420415 |
| | Tspan12 | 1.76E+00 | 3.656188534244 | 0.0002207046 | 0.12414735192 |
| | Sparc | 1.75E+00 | 1.428568139553 | 0.0372762195 | 0.38370165784 |
| | Ngfr | 1.75E+00 | 1.301827244816 | 0.0499082975 | 0.41690373122 |
| | S100a13 | 1.75E+00 | 2.599363357520 | 0.0025155714 | 0.20681751624 |
| | Aqp4 | 1.75E+00 | 1.373887200712 | 0.0422778408 | 0.39997175100 |
| | Skap1 | 1.75E+00 | 1.304610177833 | 0.0495895106 | 0.41641904240 |
| | Tfcp2l1 | 1.75E+00 | 1.378237305410 | 0.0418564792 | 0.39867372940 |
| | Gm14403 | 1.74E+00 | 1.454124977864 | 0.0351459286 | 0.37517732932 |
| | Tnfrsf1a | 1.74E+00 | 1.530855793605 | 0.0294539948 | 0.36081204708 |
| | Plekhd1 | 1.73E+00 | 1.945850737698 | 0.0113278962 | 0.28910669889 |

| | | | | | |
|---|---|---|---|---|---|
| | Adgrv1 | 1.73E+00 | 1.881419336888 | 0.0131395552 | 0.29799400500 |
| | Ormdl2 | 1.73E+00 | 3.240685529958 | 0.0005745323 | 0.15892434038 |
| | Glp2r | 1.72E+00 | 1.349319154238 | 0.0447384410 | 0.40598617309 |
| | Dbi | 1.72E+00 | 1.493260375672 | 0.0321173441 | 0.36603760992 |
| | Nek3 | 1.72E+00 | 1.755628687912 | 0.0175538067 | 0.32896966427 |
| | Tlr1 | 1.72E+00 | 1.395396524076 | 0.0402349509 | 0.39455570492 |
| | Plcd4 | 1.71E+00 | 1.969715284793 | 0.0107222200 | 0.28670727701 |
| | Aox1 | 1.70E+00 | 1.709201464550 | 0.0195343307 | 0.34145893141 |
| | Ahnak | 1.70E+00 | 1.987702946974 | 0.0102871969 | 0.28134380644 |
| | Prex2 | 1.70E+00 | 2.423865046327 | 0.0037682087 | 0.22143440510 |
| | Havcr2 | 1.69E+00 | 1.969403012476 | 0.0107299324 | 0.28670727701 |
| | Arhgef26 | 1.69E+00 | 2.241922302829 | 0.0057289852 | 0.24753450361 |
| | Lgi4 | 1.68E+00 | 1.568656327757 | 0.0269987509 | 0.35335118973 |
| | Cnn3 | 1.68E+00 | 1.459018771326 | 0.0347521140 | 0.37517732932 |
| | Smoc1 | 1.67E+00 | 1.350041932634 | 0.0446640465 | 0.40590171168 |
| | Rcn3 | 1.67E+00 | 1.344672892844 | 0.0452196407 | 0.40639873371 |
| | Ccdc102a | 1.66E+00 | 1.507895989756 | 0.0310530320 | 0.36235522207 |
| | Mlph | 1.66E+00 | 1.316676622350 | 0.0482306792 | 0.41349382326 |
| | Sema3d | 1.66E+00 | 2.000566466680 | 0.0099869651 | 0.28044040061 |
| | Phtf1os | 1.65E+00 | 2.612671326756 | 0.0024396564 | 0.20619505997 |
| | Slc1a5 | 1.65E+00 | 1.417787771810 | 0.0382130962 | 0.38774483584 |
| | P2ry14 | 1.65E+00 | 2.006443776729 | 0.0098527219 | 0.27933254996 |
| | Xdh | 1.65E+00 | 2.451364580563 | 0.0035370029 | 0.21798760175 |

| | | | | | |
|---|---|---|---|---|---|
| | Fam46a | 1.64E+00 | 2.005068797435 | 0.0098839651 | 0.27933254996 |
| | Krt12 | 1.64E+00 | 1.313696718742 | 0.0485627510 | 0.41349382326 |
| | Ccdc80 | 1.64E+00 | 1.465391472065 | 0.0342458956 | 0.37448815689 |
| | Arhgap18 | 1.63E+00 | 2.618053686869 | 0.0024096075 | 0.20580985051 |
| | Gm11266 | 1.63E+00 | 1.706945765397 | 0.0196360548 | 0.34145893141 |
| | Ttc32 | 1.63E+00 | 1.779185897033 | 0.0166270079 | 0.32205632977 |
| | Prdx6 | 1.63E+00 | 2.192812507279 | 0.0064148646 | 0.25527338269 |
| | Ptges | 1.63E+00 | 2.076407294475 | 0.0083867308 | 0.27149590175 |
| | H2-L | 1.63E+00 | 1.556319470232 | 0.0277766924 | 0.35687453723 |
| | Serhl | 1.62E+00 | 1.915259913460 | 0.0121545836 | 0.29368070731 |
| | Mlc1 | 1.62E+00 | 1.893835045796 | 0.0127692372 | 0.29799400500 |
| | Gdpd2 | 1.61E+00 | 1.625531258804 | 0.0236847465 | 0.34940735647 |
| | Tmem176b | 1.61E+00 | 1.728223859713 | 0.0186971813 | 0.33416745766 |
| | Fam107a | 1.60E+00 | 2.767015003261 | 0.0017099562 | 0.19673992847 |
| | Ccdc141 | 1.60E+00 | 4.254439897614 | 0.0000556622 | 0.12414735192 |
| | Gatsl3 | 1.60E+00 | 1.555740800843 | 0.0278137277 | 0.35696699505 |
| | Cyp2j9 | 1.59E+00 | 3.319020379778 | 0.0004797109 | 0.15651963832 |
| | Dnase1l2 | 1.59E+00 | 1.475078580620 | 0.0334904836 | 0.37233576573 |
| | Ppp1r3c | 1.59E+00 | 2.547727031641 | 0.0028331722 | 0.21635817222 |
| | Dbx2 | 1.59E+00 | 2.605439053638 | 0.0024806240 | 0.20681751624 |
| | Hhatl | 1.59E+00 | 2.595652606997 | 0.0025371573 | 0.20695533061 |
| | Map3k6 | 1.59E+00 | 1.712917547176 | 0.0193678964 | 0.34145893141 |
| | Marveld2 | 1.58E+00 | 1.607382289120 | 0.0246954936 | 0.35033142072 |

| | | | | | |
|---|---|---|---|---|---|
| | Hsd11b1 | 1.58E+00 | 1.415184518637 | 0.0384428415 | 0.38774483584 |
| | Itga6 | 1.57E+00 | 2.296000138747 | 0.0050582450 | 0.23794313665 |
| | Myh15 | 1.57E+00 | 1.499387558208 | 0.0316674025 | 0.36455475965 |
| | Gpt2 | 1.57E+00 | 3.353272154367 | 0.0004433307 | 0.15651963832 |
| | Oaf | 1.56E+00 | 2.485316043169 | 0.0032710257 | 0.21635817222 |
| | Scrg1 | 1.56E+00 | 2.090877493691 | 0.0081118985 | 0.27041155915 |
| | Zfp36l1 | 1.56E+00 | 1.401482554269 | 0.0396750466 | 0.39200063666 |
| | Naaa | 1.56E+00 | 1.580866167734 | 0.0262502735 | 0.35302467272 |
| | Clu | 1.56E+00 | 1.520395789407 | 0.0301720077 | 0.36156289629 |
| | Nxn | 1.56E+00 | 1.689107952512 | 0.0204593602 | 0.34310300466 |
| | Fkbp5 | 1.56E+00 | 2.655374355362 | 0.0022111879 | 0.20580985051 |
| | Mtm1 | 1.56E+00 | 2.264565102335 | 0.0054379461 | 0.24609625966 |
| | Cldn10 | 1.56E+00 | 1.780753219820 | 0.0165671109 | 0.32205632977 |
| | Paqr5 | 1.56E+00 | 1.989668224093 | 0.0102407503 | 0.28116971896 |
| | Npy | 1.55E+00 | 2.039809292454 | 0.0091241141 | 0.27519811275 |
| | Hmgn5 | 1.55E+00 | 3.769546059756 | 0.0001700020 | 0.12414735192 |
| | Phkg1 | 1.55E+00 | 2.046625347910 | 0.0089820331 | 0.27519811275 |
| | Als2cl | 1.55E+00 | 1.753637832334 | 0.0176344601 | 0.32988196686 |
| | Rgcc | 1.55E+00 | 2.086776438705 | 0.0081888622 | 0.27126220899 |
| | Gjb6 | 1.55E+00 | 2.347153792710 | 0.0044962061 | 0.23131440987 |
| | Fam114a1 | 1.54E+00 | 2.135437918309 | 0.0073208597 | 0.26540480894 |
| | Id4 | 1.54E+00 | 1.457547821112 | 0.0348700186 | 0.37517732932 |
| | Lrig1 | 1.54E+00 | 1.321300571804 | 0.0477198894 | 0.41341683535 |

| | | | | | |
|---|---|---|---|---|---|
| | Afap1l2 | 1.54E+00 | 1.533186979319 | 0.0292963166 | 0.35982823218 |
| | Arhgef37 | 1.54E+00 | 1.510719509729 | 0.0308517988 | 0.36169282125 |
| | Mterf1a | 1.54E+00 | 1.658872011413 | 0.0219345126 | 0.34768003377 |
| | Car5b | 1.54E+00 | 1.513331078528 | 0.0306668325 | 0.36169282125 |
| | Fabp7 | 1.54E+00 | 1.666798208669 | 0.0215378224 | 0.34494937007 |
| | Bvht | 1.53E+00 | 1.387796023253 | 0.0409452924 | 0.39782718298 |
| | Tmem47 | 1.53E+00 | 2.011922898460 | 0.0097291993 | 0.27933254996 |
| | F3 | 1.53E+00 | 2.430338336282 | 0.0037124590 | 0.22143440510 |
| | Smim1 | 1.53E+00 | 2.302548856071 | 0.0049825440 | 0.23696641539 |
| | Gna15 | 1.53E+00 | 1.588674234818 | 0.0257825439 | 0.35302467272 |
| | Adora2b | 1.52E+00 | 1.932425460657 | 0.0116835424 | 0.29197224661 |
| | Crybg3 | 1.52E+00 | 1.894478673169 | 0.0127503271 | 0.29799400500 |
| | Emx2 | 1.52E+00 | 2.637476159566 | 0.0023042195 | 0.20580985051 |
| | Per2 | 1.52E+00 | 1.946758077479 | 0.0113042544 | 0.28910669889 |
| | Nfe2l2 | 1.51E+00 | 1.450398960549 | 0.0354487593 | 0.37517732932 |
| | Clk1 | 1.51E+00 | 3.819229648428 | 0.0001516248 | 0.12414735192 |
| | Gja1 | 1.51E+00 | 1.844797092163 | 0.0142956171 | 0.30761887753 |
| | Ggt1 | 1.51E+00 | 1.302833824147 | 0.0497927573 | 0.41670901250 |
| | Vwa1 | 1.51E+00 | 2.261511773471 | 0.0054763125 | 0.24609625966 |
| | Atp10a | 1.50E+00 | 3.637938530085 | 0.0002301768 | 0.12414735192 |
| | Dhrs4 | 1.50E+00 | 2.043932880636 | 0.0090378914 | 0.27519811275 |
| | Rreb1 | 1.50E+00 | 2.314477451638 | 0.0048475528 | 0.23452126144 |
| | Cyp4v3 | 1.50E+00 | 1.654028850635 | 0.0221804907 | 0.34768003377 |

| | | | | | |
|---|---|---|---|---|---|
| | Gpld1 | 1.50E+00 | 1.705989589879 | 0.0196793346 | 0.34145893141 |
| | Mrvi1 | 1.50E+00 | 1.643522172186 | 0.0227236362 | 0.34911739477 |
| | Slfn5 | 1.49E+00 | 2.356831227758 | 0.0043971246 | 0.23131440987 |
| | Mfsd2a | 1.49E+00 | 3.208863208989 | 0.0006182111 | 0.16062039967 |
| | Slc18a2 | 1.49E+00 | 2.414551833362 | 0.0038498886 | 0.22301019851 |
| | S100a1 | 1.49E+00 | 2.099201179117 | 0.0079579063 | 0.26838804157 |
| | Sox17 | 1.49E+00 | 1.462247426732 | 0.0344947160 | 0.37517732932 |
| | Tiparp | 1.49E+00 | 2.623930348147 | 0.0023772215 | 0.20580985051 |
| | Spag1 | 1.49E+00 | 2.267420829679 | 0.0054023059 | 0.24609625966 |
| | Pop4 | 1.48E+00 | 1.637271202491 | 0.0230530715 | 0.34928420415 |
| | Renbp | 1.48E+00 | 1.585427438240 | 0.0259760171 | 0.35302467272 |
| | Zfp760 | 1.48E+00 | 3.026342909960 | 0.0009411462 | 0.18658819479 |
| | Prdm5 | 1.48E+00 | 1.324591530298 | 0.0473596484 | 0.41341683535 |
| | Flt4 | 1.48E+00 | 2.072386424079 | 0.0084647391 | 0.27170265038 |
| | Sugct | 1.48E+00 | 1.389927967664 | 0.0407447852 | 0.39642811095 |
| | Tbc1d2 | 1.47E+00 | 1.846771877934 | 0.0142307609 | 0.30716550080 |
| | Mro | 1.47E+00 | 1.862114343522 | 0.0137368026 | 0.30416998758 |
| | Rapgef3 | 1.47E+00 | 1.953593732616 | 0.0111277220 | 0.28910669889 |
| | Selenop | 1.47E+00 | 2.803275595940 | 0.0015729844 | 0.19673992847 |
| | Fgfrl1 | 1.47E+00 | 2.459659893700 | 0.0034700849 | 0.21798760175 |
| | Slc15a2 | 1.47E+00 | 3.004364227092 | 0.0009900013 | 0.18863065563 |
| | Maob | 1.47E+00 | 1.736399912748 | 0.0183484798 | 0.33259582828 |
| | Adgrl4 | 1.47E+00 | 4.277577606825 | 0.0000527743 | 0.12414735192 |

| | | | | | |
|---|---|---|---|---|---|
| | Zfp85os | 1.47E+00 | 1.576851717228 | 0.0264940458 | 0.35302467272 |
| | Cd63 | 1.47E+00 | 1.317826746892 | 0.0481031208 | 0.41349382326 |
| | Gm14327 | 1.47E+00 | 2.206884939195 | 0.0062103355 | 0.25407779976 |
| | Rasl11a | 1.47E+00 | 1.525416966388 | 0.0298251773 | 0.36081204708 |
| | Lonrf3 | 1.47E+00 | 2.201970366061 | 0.0062810122 | 0.25527338269 |
| | Pnpla7 | 1.47E+00 | 2.864363327262 | 0.0013665851 | 0.19673992847 |
| | Ednrb | 1.47E+00 | 1.360622463792 | 0.0435890633 | 0.40500301841 |
| | Lsm5 | 1.47E+00 | 2.241243135405 | 0.0057379514 | 0.24753450361 |
| | Mertk | 1.46E+00 | 4.822018123654 | 0.0000150654 | 0.10568407527 |
| | Bhlhe41 | 1.46E+00 | 3.077996627962 | 0.0008356095 | 0.18036309803 |
| | Gm7827 | 1.45E+00 | 1.386134770643 | 0.0411022153 | 0.39793007993 |
| | Pbxip1 | 1.45E+00 | 1.780535052209 | 0.0165754355 | 0.32205632977 |
| | Mfap3l | 1.45E+00 | 2.508462748196 | 0.0031012534 | 0.21635817222 |
| | C030023E24Rik | 1.45E+00 | 2.019577584356 | 0.0095592191 | 0.27933254996 |
| | Ccr5 | 1.45E+00 | 1.558362014268 | 0.0276463617 | 0.35669740320 |
| | Layn | 1.45E+00 | 1.729265901658 | 0.0186523733 | 0.33416745766 |
| | Gpr3 | 1.45E+00 | 1.644303784141 | 0.0226827766 | 0.34911739477 |
| | Fli1 | 1.45E+00 | 1.381587643331 | 0.0415348223 | 0.39793007993 |
| | E130114P18Rik | 1.45E+00 | 1.366001009104 | 0.0430525610 | 0.40376165176 |
| | Cyp2d22 | 1.44E+00 | 2.735141321557 | 0.0018401731 | 0.19673992847 |
| | Klf15 | 1.44E+00 | 3.264107160006 | 0.0005443683 | 0.15892434038 |
| | Gm16973 | 1.44E+00 | 1.460472364740 | 0.0346359924 | 0.37517732932 |
| | Zwilch | 1.44E+00 | 1.629692246240 | 0.0234589059 | 0.34940735647 |

| | | | | | |
|---|---|---|---|---|---|
| | Hsd3b7 | 1.44E+00 | 1.402691158601 | 0.0395647879 | 0.39158035901 |
| | Dio2 | 1.44E+00 | 2.900352847575 | 0.0012579030 | 0.19673992847 |
| | Prodh | 1.43E+00 | 2.256907330788 | 0.0055346819 | 0.24609625966 |
| | Idh1 | 1.43E+00 | 2.861050376343 | 0.0013770497 | 0.19673992847 |
| | Gpam | 1.43E+00 | 1.946399190438 | 0.0113135997 | 0.28910669889 |
| | Slc39a12 | 1.43E+00 | 2.955908333351 | 0.0011068574 | 0.19673992847 |
| | Dok3 | 1.43E+00 | 1.307145004454 | 0.0493009168 | 0.41624308297 |
| | Myo6 | 1.43E+00 | 2.797872650144 | 0.0015926757 | 0.19673992847 |
| | Grap | 1.43E+00 | 1.651095596600 | 0.0223308062 | 0.34772609504 |
| | Amy1 | 1.43E+00 | 2.316037651652 | 0.0048301692 | 0.23452126144 |
| | Ssfa2 | 1.43E+00 | 2.506775181158 | 0.0031133276 | 0.21635817222 |
| | Pgm5 | 1.43E+00 | 1.307256186934 | 0.0492882970 | 0.41624308297 |
| | Il18 | 1.42E+00 | 3.572837913327 | 0.0002674004 | 0.12414735192 |
| | Plpp3 | 1.42E+00 | 2.196428556436 | 0.0063616745 | 0.25527338269 |
| | Heph | 1.42E+00 | 1.609609335744 | 0.0245691801 | 0.34959999709 |
| | Adam1a | 1.42E+00 | 1.904061567960 | 0.0124720669 | 0.29557955866 |
| | Sox21 | 1.42E+00 | 1.479480018761 | 0.0331527823 | 0.37151240819 |
| | Gm12709 | 1.42E+00 | 1.335165257113 | 0.0462205110 | 0.41120720974 |
| | Paqr8 | 1.41E+00 | 1.974743235905 | 0.0105988016 | 0.28670727701 |
| | Stat3 | 1.41E+00 | 1.612132370382 | 0.0244268592 | 0.34940735647 |
| | Abca1 | 1.41E+00 | 2.314836486715 | 0.0048435469 | 0.23452126144 |
| | Bbs2 | 1.41E+00 | 1.924047678025 | 0.0119111124 | 0.29197224661 |
| | Slc1a3 | 1.41E+00 | 1.841457477485 | 0.0144059705 | 0.30857368972 |

| | | | | | |
|---|---|---|---|---|---|
| | Hopx | 1.41E+00 | 1.396321179844 | 0.0401493779 | 0.39446482604 |
| | Wwtr1 | 1.41E+00 | 1.807891574928 | 0.0155635414 | 0.32047634791 |
| | Fzd1 | 1.41E+00 | 1.506475291749 | 0.0311547814 | 0.36273990287 |
| | Lef1 | 1.41E+00 | 1.492247501627 | 0.0321923365 | 0.36603760992 |
| | Tsen54 | 1.41E+00 | 2.025017023642 | 0.0094402387 | 0.27883484032 |
| | Pmaip1 | 1.41E+00 | 1.319604709339 | 0.0479065935 | 0.41349382326 |
| | Ehmt1 | 1.41E+00 | 1.643228529663 | 0.0227390057 | 0.34911739477 |
| | Prkd1 | 1.41E+00 | 1.665465331806 | 0.0216040249 | 0.34522149051 |
| | Cpe | 1.41E+00 | 1.988261753965 | 0.0102739689 | 0.28134380644 |
| | Rhoj | 1.40E+00 | 1.636161835321 | 0.0231120338 | 0.34928420415 |
| | Ctla2a | 1.40E+00 | 2.449686553621 | 0.0035506956 | 0.21798760175 |
| | Phka1 | 1.40E+00 | 2.831658630850 | 0.0014734702 | 0.19673992847 |
| | Ctso | 1.40E+00 | 1.654350064114 | 0.0221640916 | 0.34768003377 |
| | Fam198a | 1.40E+00 | 2.240373805864 | 0.0057494486 | 0.24753450361 |
| | Tex30 | 1.40E+00 | 1.655363291256 | 0.0221124421 | 0.34768003377 |
| | Ghr | 1.40E+00 | 1.684308491723 | 0.0206867139 | 0.34416989597 |
| | Scd1 | 1.40E+00 | 2.831803969242 | 0.0014729772 | 0.19673992847 |
| | Zfp972 | 1.40E+00 | 1.744187039923 | 0.0180224139 | 0.33215831652 |
| | Rom1 | 1.40E+00 | 1.538483859881 | 0.0289411737 | 0.35982823218 |
| | Atp13a4 | 1.40E+00 | 1.671240261589 | 0.0213186519 | 0.34442385739 |
| | Lamb2 | 1.40E+00 | 1.611172108762 | 0.0244809288 | 0.34940735647 |
| | Gm20594 | 1.40E+00 | 2.106586935003 | 0.0078237158 | 0.26784898316 |
| | Tlr3 | 1.40E+00 | 1.453890154574 | 0.0351649371 | 0.37517732932 |

| | | | | | |
|---|---|---|---|---|---|
| | Axl | 1.40E+00 | 2.291902385191 | 0.0051061976 | 0.23805503318 |
| | Antxr1 | 1.40E+00 | 1.325631804602 | 0.0472463427 | 0.41316085601 |
| | Nod1 | 1.40E+00 | 1.746998117661 | 0.0179061361 | 0.33215831652 |
| | Kctd12b | 1.40E+00 | 1.649148316218 | 0.0224311574 | 0.34890148449 |
| | Atp6v0e | 1.40E+00 | 1.515700463716 | 0.0304999787 | 0.36169282125 |
| | Nde1 | 1.39E+00 | 1.562790859844 | 0.0273658625 | 0.35550282459 |
| | Sox18 | 1.39E+00 | 1.633890160717 | 0.0232332432 | 0.34928420415 |
| | Mfsd7b | 1.39E+00 | 1.386557777450 | 0.0410622008 | 0.39791751326 |
| | Phactr4 | 1.39E+00 | 2.432966472838 | 0.0036900608 | 0.22143440510 |
| | Per1 | 1.39E+00 | 1.475663819934 | 0.0334453835 | 0.37233576573 |
| | Itpr2 | 1.39E+00 | 1.604150436929 | 0.0248799534 | 0.35188079293 |
| | Mirg | 1.39E+00 | 1.899104114229 | 0.0126152507 | 0.29788534939 |
| | Sft2d2 | 1.39E+00 | 1.786319308134 | 0.0163561352 | 0.32205632977 |
| | Suclg2 | 1.39E+00 | 3.590502011187 | 0.0002567426 | 0.12414735192 |
| | Cpne3 | 1.39E+00 | 1.764739940017 | 0.0171893740 | 0.32671938921 |
| | Pdlim5 | 1.39E+00 | 1.879291868293 | 0.0132040795 | 0.29799400500 |
| | Pdk4 | 1.39E+00 | 2.399059225267 | 0.0039897049 | 0.22301019851 |
| | Zfp36l2 | 1.39E+00 | 1.674163494446 | 0.0211756381 | 0.34442385739 |
| | Ephb4 | 1.38E+00 | 1.508255325030 | 0.0310273493 | 0.36235522207 |
| | A930015D03Rik | 1.38E+00 | 1.330901844180 | 0.0466764863 | 0.41258358190 |
| | Aldoc | 1.38E+00 | 3.095518342151 | 0.0008025677 | 0.18036309803 |
| | Pdia4 | 1.38E+00 | 2.077241174987 | 0.0083706431 | 0.27149590175 |
| | Sfxn5 | 1.38E+00 | 2.452196030552 | 0.0035302379 | 0.21798760175 |

| | | | | | |
|---|---|---|---|---|---|
| | Elovl5 | 1.38E+00 | 2.074970463076 | 0.0084145237 | 0.27149590175 |
| | Tirap | 1.38E+00 | 1.702541666942 | 0.0198361934 | 0.34145893141 |
| | Psat1 | 1.38E+00 | 2.781578042628 | 0.0016535676 | 0.19673992847 |
| | Acadl | 1.38E+00 | 2.875240074731 | 0.0013327845 | 0.19673992847 |
| | Cachd1 | 1.38E+00 | 1.729492601993 | 0.0186426393 | 0.33416745766 |
| | Sdf2l1 | 1.38E+00 | 1.600712090614 | 0.0250777119 | 0.35302467272 |
| | Gpr37l1 | 1.37E+00 | 2.215275333113 | 0.0060915059 | 0.25265694675 |
| | Gdf11 | 1.37E+00 | 2.890365447924 | 0.0012871660 | 0.19673992847 |
| | 1190005I06Rik | 1.37E+00 | 1.355462614930 | 0.0441100332 | 0.40581230583 |
| | Wdr92 | 1.37E+00 | 2.252161974923 | 0.0055954887 | 0.24609625966 |
| | Spry4 | 1.37E+00 | 1.883517091928 | 0.0130762408 | 0.29799400500 |
| | Smtn | 1.37E+00 | 1.672966835977 | 0.0212340661 | 0.34442385739 |
| | Nadk2 | 1.37E+00 | 2.849105613049 | 0.0014154495 | 0.19673992847 |
| | Nr2e1 | 1.37E+00 | 1.669158810291 | 0.0214210714 | 0.34484103836 |
| | Cdk5rap3 | 1.37E+00 | 2.585657312956 | 0.0025962271 | 0.20934435885 |
| | Dbhos | 1.37E+00 | 1.706712702627 | 0.0196465952 | 0.34145893141 |
| | Cbs | 1.37E+00 | 2.617718737997 | 0.0024114667 | 0.20580985051 |
| | Trim56 | 1.37E+00 | 1.533085040217 | 0.0293031940 | 0.35982823218 |
| | Arsk | 1.37E+00 | 1.822158697666 | 0.0150605663 | 0.31606491893 |
| | Cpt2 | 1.37E+00 | 1.917274598354 | 0.0120983293 | 0.29366705924 |
| | Slc7a10 | 1.37E+00 | 3.221591383879 | 0.0006003557 | 0.15892434038 |
| | Pgghg | 1.36E+00 | 1.558128828038 | 0.0276612099 | 0.35669740320 |
| | Cyp4f15 | 1.36E+00 | 1.511163379234 | 0.0308202829 | 0.36169282125 |

| | | | | | |
|---|---|---|---|---|---|
| | C030018K13 Rik | 1.36E+00 | 1.460582855 849 | 0.0346271816 | 0.37517732 932 |
| | Rhobtb3 | 1.36E+00 | 1.891016378 157 | 0.0128523819 | 0.29799400 500 |
| | Tns3 | 1.36E+00 | 2.423314565 951 | 0.0037729881 | 0.22143440 510 |
| | Acss1 | 1.36E+00 | 2.222499975 370 | 0.0059910097 | 0.25165828 280 |
| | Dock11 | 1.36E+00 | 1.332445991 934 | 0.0465108213 | 0.41222161 860 |
| | Stk17b | 1.36E+00 | 1.469809325 453 | 0.0338992956 | 0.37426250 252 |
| | Tlcd1 | 1.36E+00 | 2.963674446 778 | 0.0010872403 | 0.19673992 847 |
| | Gimap8 | 1.36E+00 | 1.322152895 593 | 0.0476263286 | 0.41341683 535 |
| | Pcolce2 | 1.36E+00 | 1.401166221 916 | 0.0397039557 | 0.39201020 342 |
| | Clk4 | 1.36E+00 | 3.344448331 480 | 0.0004524303 | 0.15651963 832 |
| | H2-T10 | 1.36E+00 | 1.362295411 693 | 0.0434214766 | 0.40451747 478 |
| | 1810026B05 Rik | 1.36E+00 | 2.681374470 873 | 0.0020826943 | 0.20013836 353 |
| | Mettl7a1 | 1.36E+00 | 2.919630589 110 | 0.0012032875 | 0.19673992 847 |
| | P4ha1 | 1.36E+00 | 3.690062821 222 | 0.0002041443 | 0.12414735 192 |
| | Gstm1 | 1.36E+00 | 2.174342388 488 | 0.0066935669 | 0.25823683 896 |
| | Snap23 | 1.36E+00 | 1.757140859 672 | 0.0174927923 | 0.32896966 427 |
| | Prkd3 | 1.35E+00 | 2.200238177 502 | 0.0063061141 | 0.25527338 269 |
| | Arid3b | 1.35E+00 | 1.671625420 631 | 0.0212997536 | 0.34442385 739 |
| | Ezr | 1.35E+00 | 2.501232276 165 | 0.0031533177 | 0.21635817 222 |
| | Qtrt2 | 1.35E+00 | 2.068267581 347 | 0.0085454004 | 0.27248174 582 |
| | Ptprz1 | 1.35E+00 | 2.206796831 083 | 0.0062115955 | 0.25407779 976 |
| | Prom1 | 1.35E+00 | 1.633531849 025 | 0.0232524196 | 0.34928420 415 |
| | Scg3 | 1.35E+00 | 2.403356724 673 | 0.0039504200 | 0.22301019 851 |

| | | | | | |
|---|---|---|---|---|---|
| | Car8 | 1.35E+00 | 3.080734062068 | 0.0008303591 | 0.18036309803 |
| | Acsf2 | 1.35E+00 | 2.238959765617 | 0.0057681990 | 0.24753450361 |
| | Rin2 | 1.35E+00 | 1.561340287345 | 0.0274574191 | 0.35570414585 |
| | Kcnj10 | 1.35E+00 | 2.023842989582 | 0.0094657932 | 0.27900226483 |
| | Dbp | 1.35E+00 | 2.883185449381 | 0.0013086230 | 0.19673992847 |
| | Adal | 1.35E+00 | 2.151749841193 | 0.0070509910 | 0.26033000890 |
| | Pcdhgc3 | 1.34E+00 | 1.586748868466 | 0.0258970999 | 0.35302467272 |
| | Szrd1 | 1.34E+00 | 1.775478744443 | 0.0167695441 | 0.32205632977 |
| | Slco2b1 | 1.34E+00 | 1.392095685000 | 0.0405419202 | 0.39610246594 |
| | Galnt10 | 1.34E+00 | 1.318554754577 | 0.0480225531 | 0.41349382326 |
| | Ifih1 | 1.34E+00 | 1.847997322243 | 0.0141906627 | 0.30683961242 |
| | Ugdh | 1.34E+00 | 1.524109640081 | 0.0299150932 | 0.36088457206 |
| | Chpt1 | 1.34E+00 | 2.144300726677 | 0.0071729743 | 0.26207507574 |
| | Ccdc190 | 1.34E+00 | 1.328011903976 | 0.0469881229 | 0.41298385768 |
| | Esam | 1.34E+00 | 2.043352026962 | 0.0090499874 | 0.27519811275 |
| | Rnf182 | 1.34E+00 | 1.636183890020 | 0.0231108602 | 0.34928420415 |
| | Gli3 | 1.34E+00 | 1.693074192746 | 0.0202733635 | 0.34145893141 |
| | Serpine2 | 1.34E+00 | 2.479077767370 | 0.0033183503 | 0.21635817222 |
| | Manba | 1.34E+00 | 2.288614967124 | 0.0051449959 | 0.23823198830 |
| | Sardh | 1.34E+00 | 2.353127867884 | 0.0044347805 | 0.23131440987 |
| | Gadd45a | 1.34E+00 | 1.402560539963 | 0.0395766892 | 0.39158035901 |
| | Ccdc28b | 1.34E+00 | 2.501428102712 | 0.0031518961 | 0.21635817222 |
| | Sparcl1 | 1.34E+00 | 2.532142432549 | 0.0029366864 | 0.21635817222 |

| | | | | | |
|---|---|---|---|---|---|
| | Atp11c | 1.34E+00 | 2.638808682541 | 0.0022971604 | 0.20580985051 |
| | Ikbip | 1.34E+00 | 1.513965170974 | 0.0306220900 | 0.36169282125 |
| | Abhd4 | 1.34E+00 | 1.513043035079 | 0.0306871789 | 0.36169282125 |
| | Cd209c | 1.34E+00 | 1.345980234450 | 0.0450837223 | 0.40639873371 |
| | Gna13 | 1.34E+00 | 1.676394231320 | 0.0210671491 | 0.34442385739 |
| | Notch1 | 1.34E+00 | 1.691748876076 | 0.0203353253 | 0.34168217226 |
| | Mmd2 | 1.33E+00 | 2.086302520380 | 0.0081978030 | 0.27126220899 |
| | Gpr34 | 1.33E+00 | 2.528467808576 | 0.0029616395 | 0.21635817222 |
| | Fbxo2 | 1.33E+00 | 1.821750680115 | 0.0150747223 | 0.31606491893 |
| | Glud1 | 1.33E+00 | 2.966433233075 | 0.0010803557 | 0.19673992847 |
| | Ston2 | 1.33E+00 | 1.481834432450 | 0.0329735394 | 0.37094768274 |
| | Hmgn1 | 1.33E+00 | 2.339292820442 | 0.0045783309 | 0.23336997860 |
| | Ociad2 | 1.33E+00 | 3.281399904151 | 0.0005231185 | 0.15892434038 |
| | Fads1 | 1.33E+00 | 2.812497313203 | 0.0015399361 | 0.19673992847 |
| | Gng5 | 1.33E+00 | 1.405545551610 | 0.0393056016 | 0.39082749195 |
| | Abhd3 | 1.33E+00 | 2.699468673510 | 0.0019977049 | 0.19737886765 |
| | Selenbp1 | 1.33E+00 | 1.619703717182 | 0.0240047000 | 0.34940735647 |
| | Tsc22d4 | 1.33E+00 | 1.913531456265 | 0.0122030543 | 0.29368070731 |
| | Slc41a1 | 1.33E+00 | 2.102308778108 | 0.0079011666 | 0.26784898316 |
| | Abca9 | 1.33E+00 | 1.305979066000 | 0.0494334514 | 0.41624308297 |
| | Apoe | 1.33E+00 | 1.838704066804 | 0.0144975940 | 0.31006287159 |
| | Timp3 | 1.33E+00 | 2.853379510402 | 0.0014015884 | 0.19673992847 |
| | Pxdn | 1.33E+00 | 1.360332615492 | 0.0436181643 | 0.40500519244 |

| | | | | | |
|---|---|---|---|---|---|
| | Med7 | 1.33E+00 | 2.750702848749 | 0.0017754038 | 0.19673992847 |
| | Igsf11 | 1.33E+00 | 1.311853368026 | 0.0487693123 | 0.41456148115 |
| | Thoc1 | 1.33E+00 | 2.767645437056 | 0.0017074758 | 0.19673992847 |
| | Ndrg2 | 1.33E+00 | 2.616107569611 | 0.0024204295 | 0.20580985051 |
| | Gng12 | 1.32E+00 | 1.434223539501 | 0.0367939540 | 0.38097356011 |
| | Zbtb41 | 1.32E+00 | 2.684172972734 | 0.0020693170 | 0.20013836353 |
| | Nat8f1 | 1.32E+00 | 2.159101423147 | 0.0069326389 | 0.25914424154 |
| | Krcc1 | 1.32E+00 | 1.417382696445 | 0.0382487550 | 0.38774483584 |
| | Rhbdd1 | 1.32E+00 | 1.735554363721 | 0.0183842381 | 0.33277754144 |
| | Trp53inp1 | 1.32E+00 | 1.638889636541 | 0.0229673222 | 0.34928420415 |
| | Zcchc24 | 1.32E+00 | 1.943580908586 | 0.0113872562 | 0.28910669889 |
| | Zfp983 | 1.32E+00 | 2.107861585062 | 0.0078007869 | 0.26784898316 |
| | Timp4 | 1.32E+00 | 2.550625015609 | 0.0028143298 | 0.21635817222 |
| | Trib1 | 1.32E+00 | 1.569878230921 | 0.0269228957 | 0.35302467272 |
| | Daam2 | 1.32E+00 | 1.928544679146 | 0.0117884124 | 0.29197224661 |
| | Dnajc3 | 1.32E+00 | 2.239025367975 | 0.0057673277 | 0.24753450361 |
| | Adipor2 | 1.32E+00 | 3.940844520265 | 0.0001145923 | 0.12414735192 |
| | Nwd1 | 1.32E+00 | 2.103852466789 | 0.0078731320 | 0.26784898316 |
| | Ppib | 1.32E+00 | 2.644275089298 | 0.0022684275 | 0.20580985051 |
| | Dnajc1 | 1.32E+00 | 2.166902719624 | 0.0068092187 | 0.25868674163 |
| | Rmdn1 | 1.32E+00 | 1.480641294171 | 0.0330642523 | 0.37122592199 |
| | Cmtm6 | 1.32E+00 | 1.574957251227 | 0.0266098698 | 0.35302467272 |
| | Taf13 | 1.32E+00 | 1.751021695061 | 0.0177410085 | 0.33101568689 |

| | | | | | |
|---|---|---|---|---|---|
| | Appl2 | 1.31E+00 | 2.311928580570 | 0.0048760867 | 0.23462092332 |
| | Myo10 | 1.31E+00 | 1.501082649918 | 0.0315440426 | 0.36394976741 |
| | S100b | 1.31E+00 | 2.070178355345 | 0.0085078857 | 0.27190349843 |
| | Kdr | 1.31E+00 | 1.744725566233 | 0.0180000799 | 0.33215831652 |
| | Nkain4 | 1.31E+00 | 1.705337073913 | 0.0197089245 | 0.34145893141 |
| | A330023F24Rik | 1.31E+00 | 1.374196258006 | 0.0422477653 | 0.39995691455 |
| | Tsc22d3 | 1.31E+00 | 3.614670824643 | 0.0002428450 | 0.12414735192 |
| | Gm5607 | 1.31E+00 | 1.666079353159 | 0.0215735019 | 0.34512683163 |
| | Fam181b | 1.31E+00 | 1.821206573559 | 0.0150936205 | 0.31606491893 |
| | Banp | 1.31E+00 | 1.564989937268 | 0.0272276439 | 0.35469252049 |
| | Plekhf2 | 1.31E+00 | 1.501746968569 | 0.0314958281 | 0.36369256642 |
| | Tspan4 | 1.31E+00 | 1.522354670876 | 0.0300362236 | 0.36156289629 |
| | Dock1 | 1.31E+00 | 1.858382780127 | 0.0138553410 | 0.30421038291 |
| | Cflar | 1.31E+00 | 1.367459187210 | 0.0429082510 | 0.40348710590 |
| | Igdcc4 | 1.31E+00 | 1.667873074900 | 0.0214845828 | 0.34488409262 |
| | Idh2 | 1.31E+00 | 1.491513489353 | 0.0322467916 | 0.36603760992 |
| | Gt(ROSA)26Sor | 1.31E+00 | 2.432439356545 | 0.0036945423 | 0.22143440510 |
| | Rnd3 | 1.31E+00 | 1.878362097273 | 0.0132323781 | 0.29799400500 |
| | Glrx3 | 1.31E+00 | 3.308182428056 | 0.0004918329 | 0.15682762534 |
| | Cyp7b1 | 1.31E+00 | 1.748757671871 | 0.0178337358 | 0.33140041469 |
| | Slc4a4 | 1.31E+00 | 1.744621969548 | 0.0180043742 | 0.33215831652 |
| | Tmod3 | 1.31E+00 | 1.775208454771 | 0.0167799841 | 0.32205632977 |
| | Gm14322 | 1.31E+00 | 1.660863382165 | 0.0218341665 | 0.34692339303 |

| | | | | | |
|---|---|---|---|---|---|
| | Erbin | 1.31E+00 | 1.315538055969 | 0.0483572888 | 0.41349382326 |
| | Pfkfb3 | 1.31E+00 | 1.786035727807 | 0.0163668187 | 0.32205632977 |
| | Plgrkt | 1.30E+00 | 1.884792552765 | 0.0130378940 | 0.29799400500 |
| | Magt1 | 1.30E+00 | 2.311127131669 | 0.0048850934 | 0.23462092332 |
| | Mt3 | 1.30E+00 | 2.192277432287 | 0.0064227729 | 0.25527338269 |
| | Arl5a | 1.30E+00 | 1.931098872840 | 0.0117192853 | 0.29197224661 |
| | Smc4 | 1.30E+00 | 1.436921186106 | 0.0365661144 | 0.37997200643 |
| | Sgk3 | 1.30E+00 | 1.542588975059 | 0.0286688997 | 0.35982823218 |
| | Sox2 | 1.30E+00 | 1.364200572368 | 0.0432314127 | 0.40394442986 |
| | Eif2ak2 | 1.30E+00 | 1.366093684229 | 0.0430433749 | 0.40376165176 |
| | Ddah1 | 1.30E+00 | 1.303593936868 | 0.0497056850 | 0.41670901250 |
| | Arhgap31 | 1.30E+00 | 2.061672766821 | 0.0086761536 | 0.27415863813 |
| | Tulp3 | 1.30E+00 | 1.306640918736 | 0.0493581736 | 0.41624308297 |
| | Vcpkmt | 1.30E+00 | 2.006430933108 | 0.0098530133 | 0.27933254996 |
| | Zbtb40 | 1.30E+00 | 1.945931203167 | 0.0113257976 | 0.28910669889 |
| | Tcn2 | 1.30E+00 | 1.341687632597 | 0.0455315429 | 0.40649422025 |
| | Rgs5 | 1.30E+00 | 2.050021495950 | 0.0089120683 | 0.27519811275 |
| | Kdelc1 | 1.30E+00 | 2.099849875259 | 0.0079460286 | 0.26838804157 |
| | Apcdd1 | 1.30E+00 | 1.620737704358 | 0.0239476166 | 0.34940735647 |
| | Kdm6b | 1.30E+00 | 1.568039223263 | 0.0270371417 | 0.35347594190 |
| | Ccdc39 | 1.30E+00 | 1.551084832342 | 0.0281135163 | 0.35776698947 |
| | Lrp4 | 1.30E+00 | 1.554530497081 | 0.0278913479 | 0.35708187720 |
| | Atp1b2 | 1.29E+00 | 1.676197067540 | 0.0210767155 | 0.34442385739 |

| | | | | | |
|---|---|---|---|---|---|
| | Atp7a | 1.29E+00 | 1.613661946499 | 0.0243409797 | 0.34940735647 |
| | Idi1 | 1.29E+00 | 2.760352302023 | 0.0017363917 | 0.19673992847 |
| | Sox1 | 1.29E+00 | 1.768426112949 | 0.0170440927 | 0.32534506161 |
| | Cyp2j6 | 1.29E+00 | 1.341882600936 | 0.0455111070 | 0.40649422025 |
| | Zhx2 | 1.29E+00 | 1.615143220416 | 0.0242580999 | 0.34940735647 |
| | Itga7 | 1.29E+00 | 1.344424295422 | 0.0452455326 | 0.40639873371 |
| | Anks1 | 1.29E+00 | 1.543625571036 | 0.0286005529 | 0.35982823218 |
| | Smox | 1.29E+00 | 1.447111808536 | 0.0357180871 | 0.37678064837 |
| | Zfp959 | 1.29E+00 | 1.793127628155 | 0.0161017238 | 0.32134734631 |
| | Arap1 | 1.29E+00 | 1.367118750508 | 0.0429418993 | 0.40353305267 |
| | Cnbd2 | 1.29E+00 | 1.533485884042 | 0.0292761603 | 0.35982823218 |
| | Rrp8 | 1.29E+00 | 1.868203567885 | 0.0135455434 | 0.30307269491 |
| | Galnt7 | 1.29E+00 | 1.537464707469 | 0.0290091693 | 0.35982823218 |
| | Htra1 | 1.29E+00 | 4.108836087050 | 0.0000778330 | 0.12414735192 |
| | Oat | 1.28E+00 | 1.631408431072 | 0.0233663872 | 0.34940735647 |
| | Decr1 | 1.28E+00 | 2.093559963978 | 0.0080619488 | 0.26995021897 |
| | Hepacam | 1.28E+00 | 1.898491561052 | 0.0126330565 | 0.29788534939 |
| | Nfia | 1.28E+00 | 1.317132572936 | 0.0481800700 | 0.41349382326 |
| | Nefh | 1.28E+00 | 1.507391245184 | 0.0310891433 | 0.36235522207 |
| | Srsf10 | 1.28E+00 | 3.893971162425 | 0.0001276524 | 0.12414735192 |
| | Aldh6a1 | 1.28E+00 | 2.555610434404 | 0.0027822078 | 0.21635817222 |
| | Ptch1 | 1.28E+00 | 2.894098466223 | 0.0012761494 | 0.19673992847 |
| | Rcn2 | 1.28E+00 | 2.129333096599 | 0.0074244947 | 0.26606167588 |

| | | | | | |
|---|---|---|---|---|---|
| | Hspa5 | 1.28E+00 | 2.599474817447 | 0.0025149258 | 0.20681751624 |
| | Qk | 1.28E+00 | 2.064133700985 | 0.0086271291 | 0.27380488757 |
| | Mpp6 | 1.27E+00 | 2.674142186607 | 0.0021176677 | 0.20211481558 |
| | Ccdc191 | 1.27E+00 | 1.578783078057 | 0.0263764851 | 0.35302467272 |
| | Golim4 | 1.27E+00 | 1.889207432048 | 0.0129060270 | 0.29799400500 |
| | Cd2ap | 1.27E+00 | 1.674013321055 | 0.0211829616 | 0.34442385739 |
| | Nmrk1 | 1.27E+00 | 1.596297807045 | 0.0253339082 | 0.35302467272 |
| | Cdc7 | 1.27E+00 | 1.347116803866 | 0.0449658902 | 0.40639873371 |
| | Rab31 | 1.27E+00 | 3.002213677057 | 0.0009949158 | 0.18863065563 |
| | Slc5a3 | 1.27E+00 | 1.341005282998 | 0.0456031368 | 0.40649422025 |
| | Hsd17b11 | 1.27E+00 | 2.774872179205 | 0.0016792982 | 0.19673992847 |
| | Pantr1 | 1.27E+00 | 2.039925986026 | 0.0091216628 | 0.27519811275 |
| | Amot | 1.27E+00 | 2.139174937729 | 0.0072581353 | 0.26449776328 |
| | S1pr1 | 1.27E+00 | 2.386109590183 | 0.0041104598 | 0.22794368224 |
| | Fgfr3 | 1.27E+00 | 2.580223246814 | 0.0026289163 | 0.21017196812 |
| | Dclre1c | 1.27E+00 | 1.919278168189 | 0.0120426435 | 0.29333036282 |
| | Soat1 | 1.27E+00 | 1.372477250837 | 0.0424153201 | 0.40073194706 |
| | Tgfbr1 | 1.27E+00 | 1.540986502410 | 0.0287748784 | 0.35982823218 |
| | 6430573F11Rik | 1.27E+00 | 1.351546220872 | 0.0445096090 | 0.40590171168 |
| | Ryk | 1.26E+00 | 2.259661720721 | 0.0054996909 | 0.24609625966 |
| | Pnrc2 | 1.26E+00 | 2.180306076316 | 0.0066022798 | 0.25823683896 |
| | Ptpn12 | 1.26E+00 | 3.123626160137 | 0.0007522702 | 0.18036309803 |
| | Rock1 | 1.26E+00 | 1.928871671578 | 0.0117795399 | 0.29197224661 |

| | | | | | |
|---|---|---|---|---|---|
| | Atp1a2 | 1.26E+00 | 1.827185995415 | 0.0148872337 | 0.31577974654 |
| | Kif1c | 1.26E+00 | 2.446898041604 | 0.0035735672 | 0.21798760175 |
| | P2ry12 | 1.26E+00 | 1.503058989274 | 0.0314008215 | 0.36353444340 |
| | Hdac8 | 1.26E+00 | 1.868460524062 | 0.0135375314 | 0.30307269491 |
| | Ugp2 | 1.26E+00 | 2.328555853744 | 0.0046929308 | 0.23349226603 |
| | Smim11 | 1.26E+00 | 1.601456297382 | 0.0250347756 | 0.35302467272 |
| | Wsb1 | 1.26E+00 | 1.781536626173 | 0.0165372531 | 0.32205632977 |
| | Pygb | 1.26E+00 | 1.710536659538 | 0.0194743666 | 0.34145893141 |
| | Uimc1 | 1.26E+00 | 1.683942343839 | 0.0207041620 | 0.34416989597 |
| | Snapc5 | 1.26E+00 | 2.467200601653 | 0.0034103535 | 0.21721559168 |
| | Ttc14 | 1.26E+00 | 2.626539362126 | 0.0023629832 | 0.20580985051 |
| | Echdc1 | 1.26E+00 | 1.450603007581 | 0.0354321081 | 0.37517732932 |
| | Ints13 | 1.26E+00 | 2.054276821534 | 0.0088251720 | 0.27519811275 |
| | Cox6c | 1.26E+00 | 3.037039672789 | 0.0009182487 | 0.18658819479 |
| | Ankdd1b | 1.26E+00 | 1.857654854410 | 0.0138785836 | 0.30424457471 |
| | Slc12a4 | 1.26E+00 | 1.590991947757 | 0.0256453158 | 0.35302467272 |
| | Cep95 | 1.25E+00 | 1.498935067568 | 0.0317004139 | 0.36455475965 |
| | Zfp948 | 1.25E+00 | 1.834543193257 | 0.0146371595 | 0.31209627424 |
| | Cep76 | 1.25E+00 | 1.849564289260 | 0.0141395540 | 0.30683961242 |
| | Asrgl1 | 1.25E+00 | 1.794106291115 | 0.0160654801 | 0.32114897231 |
| | Pld1 | 1.25E+00 | 1.329608039503 | 0.0468157473 | 0.41286871138 |
| | Slc2a1 | 1.25E+00 | 2.018662170167 | 0.0095793894 | 0.27933254996 |
| | Chka | 1.25E+00 | 2.368320061415 | 0.0042823281 | 0.23131440987 |

| | | | | | |
|---|---|---|---|---|---|
| | Etfrf1 | 1.25E+00 | 1.700927370677 | 0.0199100628 | 0.34145893141 |
| | Lamp2 | 1.25E+00 | 1.419370375522 | 0.0380740981 | 0.38748059271 |
| | Serp1 | 1.25E+00 | 2.349492640266 | 0.0044720573 | 0.23131440987 |
| | AI987944 | 1.25E+00 | 1.514300261982 | 0.0305984719 | 0.36169282125 |
| | Hadhb | 1.25E+00 | 1.954297442603 | 0.0111097058 | 0.28910669889 |
| | Ints6l | 1.25E+00 | 2.174601290985 | 0.0066895778 | 0.25823683896 |
| | Mettl23 | 1.25E+00 | 1.474144664952 | 0.0335625798 | 0.37233576573 |
| | Taf1d | 1.25E+00 | 1.535660834792 | 0.0291299115 | 0.35982823218 |
| | Ostc | 1.25E+00 | 2.727261152958 | 0.0018738674 | 0.19737886765 |
| | Chic2 | 1.25E+00 | 1.726249152656 | 0.0187823897 | 0.33526326670 |
| | Tmem38b | 1.25E+00 | 1.965056923271 | 0.0108378485 | 0.28743859126 |
| | Purg | 1.25E+00 | 2.111601265026 | 0.0077339033 | 0.26784898316 |
| | Agpat5 | 1.25E+00 | 2.490116638274 | 0.0032350676 | 0.21635817222 |
| | Ptgs1 | 1.25E+00 | 1.469394242528 | 0.0339317108 | 0.37426250252 |
| | Zswim4 | 1.25E+00 | 1.360855146052 | 0.0435657158 | 0.40500301841 |
| | 2-Sep | 1.25E+00 | 1.448887297043 | 0.0355723620 | 0.37609663793 |
| | Sowahc | 1.25E+00 | 1.527093212468 | 0.0297102829 | 0.36081204708 |
| | Prpf39 | 1.25E+00 | 2.653640998077 | 0.0022200308 | 0.20580985051 |
| | Coq10b | 1.25E+00 | 2.353396485094 | 0.0044320384 | 0.23131440987 |
| | Lrrc58 | 1.25E+00 | 2.262752597453 | 0.0054606885 | 0.24609625966 |
| | 2010111I01Rik | 1.24E+00 | 1.952103668073 | 0.0111659668 | 0.28910669889 |
| | Acsl3 | 1.24E+00 | 2.040732421507 | 0.0091047406 | 0.27519811275 |
| | Tvp23b | 1.24E+00 | 1.848464454394 | 0.0141754073 | 0.30683961242 |

| | | | | | |
|---|---|---|---|---|---|
| | Ehd2 | 1.24E+00 | 1.418598951703 | 0.0381417880 | 0.38774483584 |
| | Srp54a | 1.24E+00 | 1.643139987221 | 0.0227436421 | 0.34911739477 |
| | Ephx2 | 1.24E+00 | 2.078520124502 | 0.0083460287 | 0.27149590175 |
| | Ktn1 | 1.24E+00 | 2.769095562838 | 0.0017017840 | 0.19673992847 |
| | Rbm25 | 1.24E+00 | 3.035239656824 | 0.0009220625 | 0.18658819479 |
| | Pon2 | 1.24E+00 | 1.315807780437 | 0.0483272652 | 0.41349382326 |
| | Sil1 | 1.24E+00 | 1.415623256670 | 0.0384040250 | 0.38774483584 |
| | Flt1 | 1.24E+00 | 1.655952889837 | 0.0220824426 | 0.34768003377 |
| | Tmem229a | 1.24E+00 | 1.581642905067 | 0.0262033667 | 0.35302467272 |
| | Caprin2 | 1.24E+00 | 1.332929424897 | 0.0464590767 | 0.41202329127 |
| | Fam114a2 | 1.24E+00 | 2.820323415415 | 0.0015124345 | 0.19673992847 |
| | Igip | 1.24E+00 | 2.167109212340 | 0.0068059819 | 0.25868674163 |
| | Zfp51 | 1.24E+00 | 1.536623748347 | 0.0290653965 | 0.35982823218 |
| | Hapln1 | 1.24E+00 | 2.019898327887 | 0.0095521618 | 0.27933254996 |
| | Smad5 | 1.24E+00 | 1.486612631060 | 0.0326127461 | 0.36810686029 |
| | Stk40 | 1.24E+00 | 1.352509501409 | 0.0444109945 | 0.40590171168 |
| | Kat2b | 1.24E+00 | 1.629564187777 | 0.0234658242 | 0.34940735647 |
| | Acad11 | 1.24E+00 | 2.912428376097 | 0.0012234089 | 0.19673992847 |
| | Gnai3 | 1.24E+00 | 1.808682634131 | 0.0155352185 | 0.32047634791 |
| | Lix1l | 1.24E+00 | 1.510844333325 | 0.0308429327 | 0.36169282125 |
| | Tmed5 | 1.24E+00 | 2.110039767427 | 0.0077617604 | 0.26784898316 |
| | Gcsh | 1.24E+00 | 1.861852053110 | 0.0137451014 | 0.30416998758 |
| | Mettl16 | 1.24E+00 | 1.922876829218 | 0.0119432678 | 0.29197224661 |

| | | | | | |
|---|---|---|---|---|---|
| | Stk38 | 1.24E+00 | 1.506066152124 | 0.0311841455 | 0.36276953226 |
| | Phkb | 1.24E+00 | 1.744065676163 | 0.0180274510 | 0.33215831652 |
| | Npc2 | 1.24E+00 | 1.326597486627 | 0.0471414041 | 0.41316085601 |
| | Luc7l3 | 1.24E+00 | 1.733924065188 | 0.0184533804 | 0.33277754144 |
| | Slc1a4 | 1.23E+00 | 1.468931485140 | 0.0339678857 | 0.37428753475 |
| | Pts | 1.23E+00 | 1.331882564831 | 0.0465712007 | 0.41249617801 |
| | Slco1a4 | 1.23E+00 | 1.347118065642 | 0.0449657596 | 0.40639873371 |
| | Hmgn3 | 1.23E+00 | 2.538516761785 | 0.0028938981 | 0.21635817222 |
| | Ak3 | 1.23E+00 | 2.701087724854 | 0.0019902713 | 0.19737886765 |
| | Gpm6b | 1.23E+00 | 1.880683152118 | 0.0131618473 | 0.29799400500 |
| | Eci1 | 1.23E+00 | 1.856036118603 | 0.0139304094 | 0.30442935277 |
| | Pik3c2a | 1.23E+00 | 1.606708494483 | 0.0247338376 | 0.35050364559 |
| | Ccdc117 | 1.23E+00 | 1.617010948334 | 0.0241539994 | 0.34940735647 |
| | Ap1s2 | 1.23E+00 | 1.663638639912 | 0.0216950852 | 0.34628219045 |
| | Garem2 | 1.23E+00 | 1.356135295339 | 0.0440417639 | 0.40581230583 |
| | Lrrc27 | 1.23E+00 | 1.582543072550 | 0.0261491109 | 0.35302467272 |
| | Pou3f3 | 1.23E+00 | 2.824370010537 | 0.0014984077 | 0.19673992847 |
| | Ccnl2 | 1.23E+00 | 1.574361223056 | 0.0266464144 | 0.35302467272 |
| | Sh3bgrl | 1.23E+00 | 2.404645189974 | 0.0039387173 | 0.22301019851 |
| | Tubb2b | 1.23E+00 | 2.913014288175 | 0.0012217595 | 0.19673992847 |
| | Nr1d2 | 1.23E+00 | 1.749580350599 | 0.0177999855 | 0.33121193245 |
| | Fam189a2 | 1.23E+00 | 1.329353582379 | 0.0468431851 | 0.41286871138 |
| | D830031N03Rik | 1.23E+00 | 1.321209818902 | 0.0477298623 | 0.41341683535 |

| | | | | | |
|---|---|---|---|---|---|
| | Acaa2 | 1.23E+00 | 1.904920600694 | 0.0124474216 | 0.29549462766 |
| | 2900092D14Rik | 1.23E+00 | 2.105981462460 | 0.0078346308 | 0.26784898316 |
| | Glul | 1.23E+00 | 1.478529182997 | 0.0332254458 | 0.37160615371 |
| | Acadvl | 1.22E+00 | 2.339224074062 | 0.0045790557 | 0.23336997860 |
| | Zc3h11a | 1.22E+00 | 2.691048358859 | 0.0020368153 | 0.19885714633 |
| | Styx | 1.22E+00 | 1.331234251012 | 0.0466407740 | 0.41258358190 |
| | Abcd4 | 1.22E+00 | 1.830129304282 | 0.0147866807 | 0.31433999327 |
| | Taf4 | 1.22E+00 | 1.359962984578 | 0.0436553039 | 0.40508195312 |
| | Lrrc57 | 1.22E+00 | 1.619348225889 | 0.0240243571 | 0.34940735647 |
| | Snrnp48 | 1.22E+00 | 1.333367330761 | 0.0464122550 | 0.41195512346 |
| | Acsbg1 | 1.22E+00 | 1.825199366987 | 0.0149554895 | 0.31606491893 |
| | Tpp1 | 1.22E+00 | 1.995953995914 | 0.0100935980 | 0.28044040061 |
| | Zfp383 | 1.22E+00 | 1.446903036192 | 0.0357352614 | 0.37678064837 |
| | Ppp2r1b | 1.22E+00 | 1.380382714369 | 0.0416502186 | 0.39793007993 |
| | St8sia1 | 1.22E+00 | 1.375308118144 | 0.0421397429 | 0.39995691455 |
| | Ptpn1 | 1.22E+00 | 2.014871941130 | 0.0096633578 | 0.27933254996 |
| | Chuk | 1.22E+00 | 2.161074414670 | 0.0069012154 | 0.25914424154 |
| | Anapc10 | 1.22E+00 | 1.862402258896 | 0.0137276988 | 0.30416998758 |
| | Slc7a2 | 1.22E+00 | 1.569627991140 | 0.0269384131 | 0.35302467272 |
| | Nab1 | 1.22E+00 | 2.772848468932 | 0.0016871416 | 0.19673992847 |
| | Gm2a | 1.22E+00 | 1.644798382667 | 0.0226569589 | 0.34911739477 |
| | Cry1 | 1.22E+00 | 1.542296764461 | 0.0286881957 | 0.35982823218 |
| | Aldh1l1 | 1.22E+00 | 2.104237005687 | 0.0078661640 | 0.26784898316 |

| | | | | | |
|---|---|---|---|---|---|
| | Cyp51 | 1.22E+00 | 1.674232971191 | 0.0211722507 | 0.34442385739 |
| | Adgrf5 | 1.22E+00 | 1.764156979316 | 0.0172124630 | 0.32671938921 |
| | Asah1 | 1.22E+00 | 2.004838098106 | 0.0098892169 | 0.27933254996 |
| | Msmo1 | 1.22E+00 | 1.784543051416 | 0.0164231685 | 0.32205632977 |
| | Zfp420 | 1.22E+00 | 1.789823167596 | 0.0162247059 | 0.32196976428 |
| | Rbm41 | 1.22E+00 | 1.571347255476 | 0.0268319814 | 0.35302467272 |
| | St3gal4 | 1.22E+00 | 1.991266021840 | 0.0102031431 | 0.28112592435 |
| | Rb1 | 1.22E+00 | 2.836299157788 | 0.0014578097 | 0.19673992847 |
| | Slc30a2 | 1.22E+00 | 1.439909422999 | 0.0363153787 | 0.37922603407 |
| | Per3 | 1.21E+00 | 1.322758013230 | 0.0475600154 | 0.41341683535 |
| | Orc4 | 1.21E+00 | 3.019274949097 | 0.0009565883 | 0.18658819479 |
| | Zfp970 | 1.21E+00 | 1.533055413415 | 0.0293051930 | 0.35982823218 |
| | Katnbl1 | 1.21E+00 | 1.393157537335 | 0.0404429161 | 0.39541053181 |
| | Stxbp2 | 1.21E+00 | 1.585455682991 | 0.0259743278 | 0.35302467272 |
| | Ndufaf4 | 1.21E+00 | 1.745785443882 | 0.0179562050 | 0.33215831652 |
| | Desi2 | 1.21E+00 | 1.734872327140 | 0.0184131323 | 0.33277754144 |
| | Ybx3 | 1.21E+00 | 1.868126455951 | 0.0135479487 | 0.30307269491 |
| | B4galt4 | 1.21E+00 | 1.761464271806 | 0.0173195151 | 0.32736724913 |
| | Pnpt1 | 1.21E+00 | 1.526455012725 | 0.0297539746 | 0.36081204708 |
| | Crebzf | 1.21E+00 | 2.618726413177 | 0.0024058779 | 0.20580985051 |
| | Gprc5b | 1.21E+00 | 1.316549339294 | 0.0482448167 | 0.41349382326 |
| | Malat1 | 1.21E+00 | 1.575919661908 | 0.0265509667 | 0.35302467272 |
| | Slc29a2 | 1.21E+00 | 1.381969390435 | 0.0414983290 | 0.39793007993 |

| | | | | | |
|---|---|---|---|---|---|
| | Arhgap12 | 1.21E+00 | 1.412272082816 | 0.0387015106 | 0.38885839955 |
| | Rab33b | 1.21E+00 | 1.478284004766 | 0.0332442083 | 0.37160615371 |
| | Scd2 | 1.21E+00 | 1.383340874361 | 0.0413674857 | 0.39793007993 |
| | Dtna | 1.21E+00 | 1.755882916867 | 0.0175435340 | 0.32896966427 |
| | Pin4 | 1.21E+00 | 2.113858531943 | 0.0076938102 | 0.26784898316 |
| | Sbk1 | 1.21E+00 | 1.410184171899 | 0.0388880197 | 0.38971351173 |
| | Fam118b | 1.21E+00 | 1.675407236294 | 0.0211150816 | 0.34442385739 |
| | Ppara | 1.21E+00 | 1.403342598954 | 0.0395054853 | 0.39142793684 |
| | Mageh1 | 1.21E+00 | 1.608192409725 | 0.0246494703 | 0.35003245720 |
| | Ccnd2 | 1.21E+00 | 1.572168190112 | 0.0267813096 | 0.35302467272 |
| | Eci2 | 1.21E+00 | 1.716965525942 | 0.0191882105 | 0.34119749140 |
| | Ralb | 1.20E+00 | 1.570599889724 | 0.0268781956 | 0.35302467272 |
| | Dynlt1a | 1.20E+00 | 1.822573259224 | 0.0150461969 | 0.31606491893 |
| | Plxnb1 | 1.20E+00 | 1.736576355047 | 0.0183410268 | 0.33259582828 |
| | Camk2n1 | 1.20E+00 | 1.682733246277 | 0.0207618837 | 0.34442385739 |
| | Sppl2a | 1.20E+00 | 1.601207748077 | 0.0250491072 | 0.35302467272 |
| | Itgb1 | 1.20E+00 | 1.496136973053 | 0.0319053143 | 0.36534960967 |
| | Pla2g16 | 1.20E+00 | 1.415745845428 | 0.0383931861 | 0.38774483584 |
| | Hsd17b10 | 1.20E+00 | 1.541049510010 | 0.0287707041 | 0.35982823218 |
| | Hbp1 | 1.20E+00 | 1.883580218594 | 0.0130743402 | 0.29799400500 |
| | Lpin1 | 1.20E+00 | 1.671767518667 | 0.0212927856 | 0.34442385739 |
| | Gpt | 1.20E+00 | 1.798090111007 | 0.0159187840 | 0.32096290667 |
| | Ankzf1 | 1.20E+00 | 1.655970585127 | 0.0220815429 | 0.34768003377 |

| | | | | | |
|---|---|---|---|---|---|
| | Zfp932 | 1.20E+00 | 1.855056206398 | 0.0139618765 | 0.30464250071 |
| | Memo1 | 1.20E+00 | 2.215025301420 | 0.0060950139 | 0.25265694675 |
| | Znrf1 | 1.20E+00 | 1.912955460004 | 0.0122192497 | 0.29368070731 |
| | Eng | 1.20E+00 | 1.525896762095 | 0.0297922455 | 0.36081204708 |
| | Fam213a | 1.20E+00 | 2.237215328398 | 0.0057914148 | 0.24772423670 |
| | Ocln | 1.20E+00 | 1.353529860933 | 0.0443067749 | 0.40590171168 |
| | Suox | 1.20E+00 | 1.810427259502 | 0.0154729364 | 0.32047634791 |
| | Ppig | 1.20E+00 | 2.261728721828 | 0.0054735776 | 0.24609625966 |
| | Atad2b | 1.20E+00 | 1.615160704724 | 0.0242571233 | 0.34940735647 |
| | Traf3 | 1.20E+00 | 1.615198469959 | 0.0242550140 | 0.34940735647 |
| | Rpl10a | 1.20E+00 | 1.371493419757 | 0.0425115148 | 0.40137049295 |
| | Eps8 | 1.20E+00 | 1.778138645108 | 0.0166671504 | 0.32205632977 |
| | Taf12 | 1.20E+00 | 1.525149301539 | 0.0298435648 | 0.36081204708 |
| | Hadha | 1.20E+00 | 2.477334843694 | 0.0033316944 | 0.21635817222 |
| | Syap1 | 1.20E+00 | 1.847901548731 | 0.0141937925 | 0.30683961242 |
| | Zmpste24 | 1.20E+00 | 1.739903063540 | 0.0182010707 | 0.33250133049 |
| | Anp32b | 1.20E+00 | 1.862042652637 | 0.0137390704 | 0.30416998758 |
| | B3glct | 1.20E+00 | 1.611527488957 | 0.0244609044 | 0.34940735647 |
| | Clic4 | 1.19E+00 | 1.781911090362 | 0.0165230003 | 0.32205632977 |
| | Rassf2 | 1.19E+00 | 1.452758364983 | 0.0352566980 | 0.37517732932 |
| | Tmco1 | 1.19E+00 | 1.409383187525 | 0.0389598084 | 0.38971786946 |
| | Myh14 | 1.19E+00 | 1.860749947789 | 0.0137800265 | 0.30421038291 |
| | Chst2 | 1.19E+00 | 1.693937902718 | 0.0202330846 | 0.34145893141 |

| | | | | | |
|---|---|---|---|---|---|
| | Ptges3 | 1.19E+00 | 2.752484992930 | 0.0017681333 | 0.19673992847 |
| | St6gal1 | 1.19E+00 | 1.442409361926 | 0.0361069362 | 0.37804501067 |
| | Ier5 | 1.19E+00 | 1.311721650473 | 0.0487841059 | 0.41456148115 |
| | Phip | 1.19E+00 | 1.585957522290 | 0.0259443311 | 0.35302467272 |
| | Afg1l | 1.19E+00 | 1.609900962122 | 0.0245526876 | 0.34959999709 |
| | Avpi1 | 1.19E+00 | 1.801084709599 | 0.0158093965 | 0.32047634791 |
| | Ints8 | 1.19E+00 | 2.414160803686 | 0.0038533566 | 0.22301019851 |
| | Upf3b | 1.19E+00 | 1.820290297995 | 0.0151254987 | 0.31626042693 |
| | Ccnl1 | 1.19E+00 | 1.502311339482 | 0.0314549255 | 0.36369256642 |
| | Slc38a3 | 1.19E+00 | 1.575286706592 | 0.0265896912 | 0.35302467272 |
| | Phf20l1 | 1.19E+00 | 2.129145343189 | 0.0074277052 | 0.26606167588 |
| | Polr2h | 1.19E+00 | 1.373433292603 | 0.0423220511 | 0.40012020025 |
| | Cd164 | 1.19E+00 | 1.598655416767 | 0.0251967533 | 0.35302467272 |
| | N4bp2l2 | 1.19E+00 | 1.518403557909 | 0.0303107332 | 0.36156289629 |
| | Etfa | 1.19E+00 | 1.537737885291 | 0.0289909278 | 0.35982823218 |
| | Mid1ip1 | 1.19E+00 | 1.614613950420 | 0.0242876810 | 0.34940735647 |
| | Srsf11 | 1.19E+00 | 2.212197818735 | 0.0061348250 | 0.25296335739 |
| | Mrpl24 | 1.19E+00 | 1.990586598507 | 0.0102191177 | 0.28112592435 |
| | Csrp1 | 1.19E+00 | 1.851945260814 | 0.0140622476 | 0.30588113681 |
| | Nnt | 1.19E+00 | 1.640665600241 | 0.0228735936 | 0.34928420415 |
| | Ctsd | 1.19E+00 | 1.696109564695 | 0.0201321629 | 0.34145893141 |
| | Tmem181a | 1.19E+00 | 1.485688322189 | 0.0326822297 | 0.36859460006 |
| | Vamp3 | 1.19E+00 | 1.706753844857 | 0.0196447341 | 0.34145893141 |

| | | | | | |
|---|---|---|---|---|---|
| | Tnfrsf19 | 1.19E+00 | 1.655548037479 | 0.0221030376 | 0.34768003377 |
| | Inppl1 | 1.18E+00 | 1.402206842081 | 0.0396089343 | 0.39162321959 |
| | Epas1 | 1.18E+00 | 2.254240889672 | 0.0055687678 | 0.24609625966 |
| | Dpysl2 | 1.18E+00 | 2.690153976090 | 0.0020410142 | 0.19885714633 |
| | Ptprb | 1.18E+00 | 1.966402396924 | 0.0108043241 | 0.28743859126 |
| | Slc25a33 | 1.18E+00 | 1.421726633815 | 0.0378680870 | 0.38695503319 |
| | Metrn | 1.18E+00 | 1.444665908049 | 0.0359198151 | 0.37716131557 |
| | Dusp11 | 1.18E+00 | 1.670944668245 | 0.0213331669 | 0.34442385739 |
| | Pcyt1a | 1.18E+00 | 1.909804927492 | 0.0123082150 | 0.29368070731 |
| | Slc31a1 | 1.18E+00 | 1.750905261851 | 0.0177457655 | 0.33101568689 |
| | Sarnp | 1.18E+00 | 2.020404038935 | 0.0095410454 | 0.27933254996 |
| | Cxcl14 | 1.18E+00 | 1.673597291719 | 0.0212032634 | 0.34442385739 |
| | Rrbp1 | 1.18E+00 | 1.704639219626 | 0.0197406196 | 0.34145893141 |
| | Mrrf | 1.18E+00 | 1.343282745400 | 0.0453646176 | 0.40649422025 |
| | Kank3 | 1.18E+00 | 1.412093367215 | 0.0387174399 | 0.38885839955 |
| | Map3k2 | 1.18E+00 | 1.549339258976 | 0.0282267412 | 0.35817906857 |
| | Nktr | 1.18E+00 | 1.621881877417 | 0.0238846083 | 0.34940735647 |
| | Elf2 | 1.18E+00 | 1.790879190003 | 0.0161853021 | 0.32196976428 |
| | Crkl | 1.18E+00 | 2.078192407038 | 0.0083523290 | 0.27149590175 |
| | Acadm | 1.18E+00 | 1.554311747982 | 0.0279054000 | 0.35708187720 |
| | Pomt1 | 1.18E+00 | 1.619577393944 | 0.0240116833 | 0.34940735647 |
| | Commd1 | 1.18E+00 | 1.473346300585 | 0.0336243347 | 0.37233576573 |
| | Fth1 | 1.18E+00 | 1.518755655334 | 0.0302861692 | 0.36156289629 |

| | | | | | |
|---|---|---|---|---|---|
| | Mcl1 | 1.18E+00 | 1.929542954437 | 0.0117613465 | 0.29197224661 |
| | E130307A14Rik | 1.18E+00 | 1.499785125939 | 0.0316384263 | 0.36455475965 |
| | Fam162a | 1.18E+00 | 1.455372056231 | 0.0350451517 | 0.37517732932 |
| | Srek1 | 1.17E+00 | 1.770123928194 | 0.0169775912 | 0.32495989684 |
| | Plod3 | 1.17E+00 | 1.305648448836 | 0.0494710982 | 0.41624308297 |
| | Prpf3 | 1.17E+00 | 1.970250117227 | 0.0107090238 | 0.28670727701 |
| | Vegfa | 1.17E+00 | 1.455447074448 | 0.0350390986 | 0.37517732932 |
| | Xndc1 | 1.17E+00 | 1.356580255120 | 0.0439966637 | 0.40581230583 |
| | Tra2a | 1.17E+00 | 1.676063391772 | 0.0210832039 | 0.34442385739 |
| | Pdcd11 | 1.17E+00 | 1.437925088732 | 0.0364816869 | 0.37970182986 |
| | Cpsf6 | 1.17E+00 | 1.866305706081 | 0.0136048668 | 0.30345990638 |
| | Rpl39 | 1.17E+00 | 1.363629255799 | 0.0432883213 | 0.40394442986 |
| | Chrm4 | 1.17E+00 | 1.422398183800 | 0.0378095768 | 0.38663874804 |
| | Itm2a | 1.17E+00 | 1.351893051570 | 0.0444740775 | 0.40590171168 |
| | Rian | 1.17E+00 | 2.211245675349 | 0.0061482897 | 0.25296335739 |
| | Git2 | 1.17E+00 | 1.581367340884 | 0.0262199983 | 0.35302467272 |
| | Xbp1 | 1.17E+00 | 1.813564678555 | 0.0153615600 | 0.31960333310 |
| | Adh5 | 1.17E+00 | 2.356415502793 | 0.0044013357 | 0.23131440987 |
| | Vcl | 1.17E+00 | 1.693226027895 | 0.0202662769 | 0.34145893141 |
| | Fnta | 1.17E+00 | 2.359409104031 | 0.0043711015 | 0.23131440987 |
| | Med21 | 1.17E+00 | 1.301090157155 | 0.0499930741 | 0.41700524975 |
| | Naa50 | 1.17E+00 | 1.879525313438 | 0.0131969839 | 0.29799400500 |
| | Mrps35 | 1.17E+00 | 1.559738630628 | 0.0275588677 | 0.35668903445 |

| | | | | | |
|---|---|---|---|---|---|
| | Ptgr2 | 1.17E+00 | 1.307455566889 | 0.0492656745 | 0.41624308297 |
| | Scp2 | 1.17E+00 | 1.571452848226 | 0.0268254584 | 0.35302467272 |
| | Rbl2 | 1.17E+00 | 1.305659050617 | 0.0494698905 | 0.41624308297 |
| | Rfk | 1.17E+00 | 1.420207765740 | 0.0380007558 | 0.38734157611 |
| | Smc5 | 1.17E+00 | 1.409268323104 | 0.0389701141 | 0.38971786946 |
| | Arglu1 | 1.17E+00 | 1.612496361200 | 0.0244063952 | 0.34940735647 |
| | Cabp1 | 1.17E+00 | 1.351279392084 | 0.0445369639 | 0.40590171168 |
| | Lrrc49 | 1.17E+00 | 1.384437593178 | 0.0412631527 | 0.39793007993 |
| | Gstm5 | 1.17E+00 | 2.421601329550 | 0.0037879014 | 0.22143440510 |
| | Yars2 | 1.17E+00 | 1.311267775734 | 0.0488351161 | 0.41474375209 |
| | Pitrm1 | 1.17E+00 | 1.313631437192 | 0.0485700514 | 0.41349382326 |
| | Abat | 1.17E+00 | 2.010792579700 | 0.0097545541 | 0.27933254996 |
| | Alkbh5 | 1.16E+00 | 1.737749679299 | 0.0182915421 | 0.33256492555 |
| | Enah | 1.16E+00 | 2.331180358060 | 0.0046646562 | 0.23349226603 |
| | Pop5 | 1.16E+00 | 1.453604618886 | 0.0351880646 | 0.37517732932 |
| | Nemf | 1.16E+00 | 1.835890995014 | 0.0145918046 | 0.31160276768 |
| | Bnip2 | 1.16E+00 | 1.879219512128 | 0.0132062796 | 0.29799400500 |
| | Ppp4r3b | 1.16E+00 | 1.636978061192 | 0.0230686372 | 0.34928420415 |
| | Maoa | 1.16E+00 | 1.451822716776 | 0.0353327372 | 0.37517732932 |
| | Rps24 | 1.16E+00 | 2.003706782306 | 0.0099150114 | 0.27933254996 |
| | Carnmt1 | 1.16E+00 | 1.363488210913 | 0.0433023822 | 0.40394442986 |
| | Spry2 | 1.16E+00 | 1.728736378198 | 0.0186751295 | 0.33416745766 |
| | Smurf2 | 1.16E+00 | 1.368171669777 | 0.0428379155 | 0.40309587861 |

| | | | | | |
|---|---|---|---|---|---|
| | Plekha8 | 1.16E+00 | 1.399046567492 | 0.0398982119 | 0.39337449959 |
| | Bfar | 1.16E+00 | 2.051362921192 | 0.0088845836 | 0.27519811275 |
| | Sec24a | 1.16E+00 | 1.344034572706 | 0.0452861528 | 0.40649422025 |
| | Acin1 | 1.16E+00 | 1.498972482520 | 0.0316976830 | 0.36455475965 |
| | Srsf7 | 1.16E+00 | 2.005125748494 | 0.0098826690 | 0.27933254996 |
| | Mbd2 | 1.16E+00 | 1.614238464016 | 0.0243086889 | 0.34940735647 |
| | Rbm39 | 1.15E+00 | 2.122867067243 | 0.0075358619 | 0.26699025975 |
| | Cdk8 | 1.15E+00 | 1.397895239654 | 0.0400041236 | 0.39386516069 |
| | Uqcrh | 1.15E+00 | 2.189265831754 | 0.0064674662 | 0.25632359011 |
| | Nsrp1 | 1.15E+00 | 1.441449700638 | 0.0361868099 | 0.37831664932 |
| | Ccar1 | 1.15E+00 | 1.547940798257 | 0.0283177799 | 0.35889652382 |
| | Ccni | 1.15E+00 | 2.081990870073 | 0.0082795957 | 0.27149590175 |
| | Map2 | 1.15E+00 | 1.404824525327 | 0.0393709120 | 0.39119964228 |
| | Ptp4a1 | 1.15E+00 | 1.571460481932 | 0.0268249869 | 0.35302467272 |
| | Mfhas1 | 1.15E+00 | 1.492030934728 | 0.0322083936 | 0.36603760992 |
| | Epb41 | 1.15E+00 | 1.420029615382 | 0.0380163471 | 0.38734157611 |
| | Ptma | 1.15E+00 | 1.533274778337 | 0.0292903945 | 0.35982823218 |
| | Arxes2 | 1.15E+00 | 1.327070099867 | 0.0470901312 | 0.41316085601 |
| | Acap2 | 1.15E+00 | 2.074805135807 | 0.0084177275 | 0.27149590175 |
| | Atl2 | 1.15E+00 | 1.404379860410 | 0.0394112437 | 0.39121861966 |
| | Ndufa5 | 1.15E+00 | 1.376510282507 | 0.0420232578 | 0.39972796706 |
| | Atp5j | 1.15E+00 | 2.154704048803 | 0.0070031907 | 0.26033000890 |
| | Srsf1 | 1.15E+00 | 1.887917988633 | 0.0129444026 | 0.29799400500 |

| | | | | | |
|---|---|---|---|---|---|
| | Bag1 | 1.15E+00 | 2.434995330105 | 0.0036728625 | 0.22143440510 |
| | Prkcsh | 1.15E+00 | 1.404188745147 | 0.0394285907 | 0.39121861966 |
| | Rpl5 | 1.15E+00 | 1.635791052014 | 0.0231317744 | 0.34928420415 |
| | Aldh7a1 | 1.15E+00 | 2.081559656359 | 0.0082878206 | 0.27149590175 |
| | Snx5 | 1.15E+00 | 1.949747490441 | 0.0112267101 | 0.28910669889 |
| | Epb41l2 | 1.15E+00 | 1.466619026240 | 0.0341492346 | 0.37434910630 |
| | Hypk | 1.15E+00 | 1.589874926343 | 0.0257113614 | 0.35302467272 |
| | Pfkfb2 | 1.15E+00 | 1.570171566234 | 0.0269047173 | 0.35302467272 |
| | Cript | 1.15E+00 | 1.409249680355 | 0.0389717869 | 0.38971786946 |
| | 9330159F19Rik | 1.15E+00 | 1.564080433254 | 0.0272847241 | 0.35477727456 |
| | Gm12191 | 1.15E+00 | 1.379638229161 | 0.0417216784 | 0.39793007993 |
| | Srsf3 | 1.15E+00 | 2.028234634329 | 0.0093705561 | 0.27853580989 |
| | Cetn3 | 1.15E+00 | 1.738490772907 | 0.0182603554 | 0.33256492555 |
| | Slc38a2 | 1.15E+00 | 1.565408310335 | 0.0272014271 | 0.35468031815 |
| | Rapgef4 | 1.15E+00 | 1.687457104069 | 0.0205372787 | 0.34343029754 |
| | E2f6 | 1.15E+00 | 1.785667506166 | 0.0163807014 | 0.32205632977 |
| | Sgms1 | 1.15E+00 | 1.635518152627 | 0.0231463144 | 0.34928420415 |
| | Bmt2 | 1.15E+00 | 1.391105373443 | 0.0406344725 | 0.39642811095 |
| | D430019H16Rik | 1.15E+00 | 1.600539366119 | 0.0250876876 | 0.35302467272 |
| | Bckdha | 1.14E+00 | 1.408581795303 | 0.0390317663 | 0.38976205057 |
| | Ddrgk1 | 1.14E+00 | 1.888728092398 | 0.0129202795 | 0.29799400500 |
| | Prpf38b | 1.14E+00 | 1.620112265475 | 0.0239821290 | 0.34940735647 |
| | Mfn1 | 1.14E+00 | 1.501768959452 | 0.0314942333 | 0.36369256642 |

| | | | | | |
|---|---|---|---|---|---|
| | Npm1 | 1.14E+00 | 1.973349042563 | 0.0106328811 | 0.28670727701 |
| | Ss18 | 1.14E+00 | 1.316232236994 | 0.0482800558 | 0.41349382326 |
| | Srsf6 | 1.14E+00 | 1.766537517656 | 0.0171183729 | 0.32631898318 |
| | Rps18 | 1.14E+00 | 1.378549666427 | 0.0418263853 | 0.39865773467 |
| | Spred1 | 1.14E+00 | 1.321925402353 | 0.0476512829 | 0.41341683535 |
| | Kmt5a | 1.14E+00 | 1.621037563054 | 0.0239310876 | 0.34940735647 |
| | Cyb5a | 1.14E+00 | 1.636630500304 | 0.0230871061 | 0.34928420415 |
| | Stx8 | 1.14E+00 | 1.460231785128 | 0.0346551845 | 0.37517732932 |
| | Eif1b | 1.14E+00 | 1.969574853317 | 0.0107256877 | 0.28670727701 |
| | Sp3 | 1.14E+00 | 1.661699776468 | 0.0217921572 | 0.34681617379 |
| | 0610037L13Rik | 1.14E+00 | 1.473730272548 | 0.0335946196 | 0.37233576573 |
| | Tle1 | 1.14E+00 | 1.452549134069 | 0.0352736877 | 0.37517732932 |
| | Fuca2 | 1.14E+00 | 1.973566017950 | 0.0106275702 | 0.28670727701 |
| | Hnrnpf | 1.14E+00 | 1.518097351542 | 0.0303321118 | 0.36156289629 |
| | Hspe1 | 1.14E+00 | 1.324463381389 | 0.0473736251 | 0.41341683535 |
| | Zfand6 | 1.14E+00 | 1.644939409656 | 0.0226496028 | 0.34911739477 |
| | Rps6 | 1.14E+00 | 1.778089525394 | 0.0166690356 | 0.32205632977 |
| | Ndufs4 | 1.14E+00 | 1.797993587051 | 0.0159223224 | 0.32096290667 |
| | 1110059E24Rik | 1.14E+00 | 1.510720408167 | 0.0308517350 | 0.36169282125 |
| | Rap1b | 1.14E+00 | 1.321084802482 | 0.0477436038 | 0.41341683535 |
| | Svip | 1.13E+00 | 1.761033628234 | 0.0173366975 | 0.32736724913 |
| | Ppp4r3a | 1.13E+00 | 1.386496055312 | 0.0410680370 | 0.39791751326 |
| | Tmem129 | 1.13E+00 | 1.416636940471 | 0.0383144909 | 0.38774483584 |

| | | | | | |
|---|---|---|---|---|---|
| | Hsd17b4 | 1.13E+00 | 1.507296659588 | 0.0310959150 | 0.36235522207 |
| | Luc7l2 | 1.13E+00 | 2.041459034028 | 0.0090895203 | 0.27519811275 |
| | Rps4x | 1.13E+00 | 1.576971049504 | 0.0264867670 | 0.35302467272 |
| | Rmdn3 | 1.13E+00 | 1.892550879656 | 0.0128070505 | 0.29799400500 |
| | Paip2 | 1.13E+00 | 1.382995735596 | 0.0414003740 | 0.39793007993 |
| | Zfp651 | 1.13E+00 | 1.408161144282 | 0.0390695902 | 0.38986226900 |
| | Ldhb | 1.13E+00 | 1.519187648217 | 0.0302560585 | 0.36156289629 |
| | Tma7 | 1.13E+00 | 1.618911135013 | 0.0240485483 | 0.34940735647 |
| | Cnot6 | 1.13E+00 | 1.453658505102 | 0.0351836989 | 0.37517732932 |
| | Serpinb6a | 1.13E+00 | 1.341502813019 | 0.0455509236 | 0.40649422025 |
| | Cdc42se1 | 1.13E+00 | 1.453864577960 | 0.0351670081 | 0.37517732932 |
| | Mcf2l | 1.13E+00 | 1.498151048557 | 0.0317576934 | 0.36461574327 |
| | Mfsd11 | 1.13E+00 | 1.454620242692 | 0.0351058714 | 0.37517732932 |
| | Picalm | 1.12E+00 | 1.675359173898 | 0.0211174185 | 0.34442385739 |
| | Pcca | 1.12E+00 | 1.330695231009 | 0.0466986977 | 0.41258358190 |
| | Mapkapk5 | 1.12E+00 | 1.318497750897 | 0.0480288568 | 0.41349382326 |
| | Spire1 | 1.12E+00 | 1.361218996361 | 0.0435292319 | 0.40498350350 |
| | Rpl23a | 1.12E+00 | 1.513554604747 | 0.0306510527 | 0.36169282125 |
| | Txlna | 1.12E+00 | 1.652437469005 | 0.0222619156 | 0.34768003377 |
| | Igf1r | 1.12E+00 | 1.357622216549 | 0.0438912332 | 0.40581230583 |
| | Atg3 | 1.12E+00 | 1.585420003804 | 0.0259764618 | 0.35302467272 |
| | Eloc | 1.12E+00 | 1.344445185997 | 0.0452433562 | 0.40639873371 |
| | Phospho2 | 1.12E+00 | 1.436313356244 | 0.0366173275 | 0.37997200643 |

| | | | | | |
|---|---|---|---|---|---|
| | Mat2a | 1.12E+00 | 1.495858553150 | 0.0319257749 | 0.36534960967 |
| | Metap2 | 1.12E+00 | 1.345186070577 | 0.0451662391 | 0.40639873371 |
| | Ide | 1.12E+00 | 1.322817871930 | 0.0475534607 | 0.41341683535 |
| | Cops9 | 1.12E+00 | 1.543466026693 | 0.0286110617 | 0.35982823218 |
| | Pdia3 | 1.12E+00 | 1.341680571170 | 0.0455322832 | 0.40649422025 |
| | Smarca5 | 1.12E+00 | 1.643393634259 | 0.0227303627 | 0.34911739477 |
| | Wnk1 | 1.11E+00 | 1.522278349138 | 0.0300415026 | 0.36156289629 |
| | Eif2s2 | 1.11E+00 | 1.730482578954 | 0.0186001917 | 0.33416745766 |
| | Zfp365 | 1.11E+00 | 1.323478744591 | 0.0474811529 | 0.41341683535 |
| | Eif1 | 1.11E+00 | 1.432403514796 | 0.0369484722 | 0.38229134580 |
| | Sf3b6 | 1.11E+00 | 1.314049769662 | 0.0485232890 | 0.41349382326 |
| | Trak2 | 1.11E+00 | 1.391327507268 | 0.0406136940 | 0.39642811095 |
| | Sec62 | 1.10E+00 | 1.596289938187 | 0.0253343672 | 0.35302467272 |
| | Egln2 | 1.10E+00 | 1.405974776699 | 0.0392667740 | 0.39082749195 |
| | Eif5b | 1.10E+00 | 1.545419645926 | 0.0284826474 | 0.35982823218 |
| | Rpl21 | 1.10E+00 | 1.630421005561 | 0.0234195742 | 0.34940735647 |
| | Hdac2 | 1.10E+00 | 1.333275892581 | 0.0464220278 | 0.41195512346 |
| | Myl6 | 1.10E+00 | 1.351399318624 | 0.0445246671 | 0.40590171168 |

| DOWN p<0.05 | Gene ID | Fold change (BRAFwt vs. control-FP) | log-10 P-values | P-value | FDR step up |
|---|---|---|---|---|---|
| | Ipo5 | -1.09E+00 | 1.3535328258635 | 0.0443064724 | 0.4059017117 |
| | Arhgap44 | -1.09E+00 | 1.3706907277327 | 0.0425901601 | 0.4015725442 |
| | Trappc11 | -1.09E+00 | 1.3341877025689 | 0.0463246661 | 0.4118221475 |
| | Efr3b | -1.09E+00 | 1.3195996605927 | 0.0479071504 | 0.4134938233 |
| | Fbxw11 | -1.09E+00 | 1.3414179215064 | 0.0455598283 | 0.4064942202 |
| | Gbf1 | -1.09E+00 | 1.3902470800200 | 0.0407148576 | 0.3964281109 |
| | Gnb1 | -1.09E+00 | 1.3635440223869 | 0.0432968178 | 0.4039444299 |
| | Wdr82 | -1.09E+00 | 1.5053780187932 | 0.0312335955 | 0.3627695323 |
| | Clptm1l | -1.09E+00 | 1.3838202692456 | 0.0413218475 | 0.3979300799 |
| | Rad23b | -1.09E+00 | 1.3636943110675 | 0.0432818374 | 0.4039444299 |
| | Rab5b | -1.09E+00 | 1.6269358210845 | 0.0236082708 | 0.3494073565 |
| | Ncoa1 | -1.09E+00 | 1.3086093431111 | 0.0491349657 | 0.4162175084 |
| | Brap | -1.09E+00 | 1.4294161130203 | 0.0372035075 | 0.3837016578 |
| | Uba1 | -1.09E+00 | 1.5036155055776 | 0.0313606095 | 0.3635344434 |
| | Pcnx | -1.10E+00 | 1.3434068080794 | 0.0453516604 | 0.4064942202 |
| | Tex264 | -1.10E+00 | 1.3028879995046 | 0.0497865463 | 0.4167090125 |
| | Mapk8ip2 | -1.10E+00 | 1.4537551319055 | 0.0351758717 | 0.3751773293 |
| | Zzef1 | -1.10E+00 | 1.3942682880715 | 0.0403396116 | 0.3949509771 |
| | Ehd3 | -1.10E+00 | 1.3092819156363 | 0.0490589314 | 0.4161407544 |
| | Gria3 | -1.10E+00 | 1.3290299234497 | 0.0468781081 | 0.4128687114 |
| | Ciapin1 | -1.10E+00 | 1.4122359483866 | 0.0387047308 | 0.3888583996 |

| | | | | | |
|---|---|---|---|---|---|
| | Dip2b | -1.10E+00 | 1.3817651766678 | 0.0415178469 | 0.3979300799 |
| | Stx6 | -1.10E+00 | 1.3814355576928 | 0.0415493699 | 0.3979300799 |
| | Kif3c | -1.10E+00 | 1.3256456390906 | 0.0472448376 | 0.4131608560 |
| | Copa | -1.10E+00 | 1.4165745444854 | 0.0383199960 | 0.3877448358 |
| | Atg9a | -1.10E+00 | 1.3514517398401 | 0.0445192931 | 0.4059017117 |
| | Psmd8 | -1.10E+00 | 1.3798196944345 | 0.0417042491 | 0.3979300799 |
| | Dstyk | -1.10E+00 | 1.3755014598556 | 0.0421209871 | 0.3999569146 |
| | Ap3b2 | -1.10E+00 | 1.4117590152272 | 0.0387472589 | 0.3888583996 |
| | Atp6v0e2 | -1.10E+00 | 1.3842617218651 | 0.0412798659 | 0.3979300799 |
| | Rab3gap1 | -1.10E+00 | 1.4166822444708 | 0.0383104943 | 0.3877448358 |
| | Slc35e1 | -1.10E+00 | 1.5694631411217 | 0.0269486404 | 0.3530246727 |
| | Pcdh10 | -1.10E+00 | 1.3258102452096 | 0.0472269343 | 0.4131608560 |
| | Atp6v0d1 | -1.11E+00 | 1.7375639269049 | 0.0182993673 | 0.3325649255 |
| | Jak1 | -1.11E+00 | 1.5459188953285 | 0.0284499236 | 0.3598282322 |
| | Maged1 | -1.11E+00 | 1.3762067787013 | 0.0420526357 | 0.3997279671 |
| | Rnft2 | -1.11E+00 | 1.5329078267121 | 0.0293151535 | 0.3598282322 |
| | Ttc3 | -1.11E+00 | 1.3456709801126 | 0.0451158371 | 0.4063987337 |
| | Dpp9 | -1.11E+00 | 1.3830169848922 | 0.0413983484 | 0.3979300799 |
| | Sel1l | -1.11E+00 | 1.5182608676500 | 0.0303206936 | 0.3615628963 |
| | Tnpo2 | -1.11E+00 | 1.6397117812811 | 0.0229238849 | 0.3492842042 |
| | Dusp8 | -1.11E+00 | 1.5707015669525 | 0.0268719036 | 0.3530246727 |
| | Ikbkap | -1.11E+00 | 1.4311887882029 | 0.0370519622 | 0.3830796087 |
| | Rusc2 | -1.11E+00 | 1.3708914095919 | 0.0425704842 | 0.4015725442 |

| | | | | | |
|---|---|---|---|---|---|
| | Rab3a | -1.11E+00 | 1.3218938678629 | 0.0476547430 | 0.4134168354 |
| | Rap1gds1 | -1.11E+00 | 1.3097269695868 | 0.0490086828 | 0.4159660133 |
| | Cdc42se2 | -1.11E+00 | 1.4683421226191 | 0.0340140133 | 0.3742875348 |
| | Cdipt | -1.11E+00 | 1.6126622297286 | 0.0243970755 | 0.3494073565 |
| | Prrc2b | -1.11E+00 | 1.3838525390707 | 0.0413187773 | 0.3979300799 |
| | Prpf8 | -1.11E+00 | 1.9400902321636 | 0.0114791510 | 0.2896627485 |
| | Dctn2 | -1.11E+00 | 1.4356874045342 | 0.0366701423 | 0.3799720064 |
| | Dcaf5 | -1.11E+00 | 1.6204936623843 | 0.0239610772 | 0.3494073565 |
| | Eef1a2 | -1.11E+00 | 1.3947024395767 | 0.0402993054 | 0.3948318816 |
| | Magi2 | -1.11E+00 | 1.7092286457285 | 0.0195331081 | 0.3414589314 |
| | Snrnp200 | -1.12E+00 | 1.9125247586211 | 0.0122313739 | 0.2936807073 |
| | Dctn1 | -1.12E+00 | 1.6748541530977 | 0.0211419892 | 0.3444238574 |
| | Vps16 | -1.12E+00 | 1.3500184795014 | 0.0446664586 | 0.4059017117 |
| | Ncstn | -1.12E+00 | 1.7147456609608 | 0.0192865407 | 0.3412234131 |
| | Camk1d | -1.12E+00 | 1.5827879732869 | 0.0261343695 | 0.3530246727 |
| | Ppm1e | -1.12E+00 | 1.5438799413212 | 0.0285838062 | 0.3598282322 |
| | Kat2a | -1.12E+00 | 1.6689222419570 | 0.0214327431 | 0.3448410384 |
| | Mgrn1 | -1.12E+00 | 1.4805093634641 | 0.0330742981 | 0.3712259220 |
| | Trim32 | -1.12E+00 | 1.5885642498952 | 0.0257890741 | 0.3530246727 |
| | Myl12b | -1.12E+00 | 1.7314673463180 | 0.0185580634 | 0.3341674577 |
| | Srr | -1.12E+00 | 1.8023571423839 | 0.0157631445 | 0.3204763479 |
| | Tmem8b | -1.12E+00 | 1.5030094864261 | 0.0314044010 | 0.3635344434 |
| | Syn1 | -1.12E+00 | 1.6769322956491 | 0.0210410643 | 0.3444238574 |

| | | | | | |
|---|---|---|---|---|---|
| | Tnfrsf21 | -1.12E+00 | 1.3625672530284 | 0.0433943060 | 0.4045174748 |
| | Ap2m1 | -1.12E+00 | 1.6204440309965 | 0.0239638156 | 0.3494073565 |
| | Mapre3 | -1.12E+00 | 1.3499894922940 | 0.0446694400 | 0.4059017117 |
| | Atp6v1c1 | -1.12E+00 | 1.5199339792964 | 0.0302041084 | 0.3615628963 |
| | Mgat3 | -1.12E+00 | 1.8131425983299 | 0.0153764968 | 0.3196033331 |
| | Magee1 | -1.12E+00 | 1.6459763236053 | 0.0225955895 | 0.3491173948 |
| | Rcc2 | -1.12E+00 | 1.4534831101202 | 0.0351979111 | 0.3751773293 |
| | Spata2 | -1.12E+00 | 1.4726202135022 | 0.0336805974 | 0.3726646546 |
| | Apba2 | -1.12E+00 | 1.3291448403459 | 0.0468657056 | 0.4128687114 |
| | Atcay | -1.12E+00 | 1.5282360783058 | 0.0296322017 | 0.3608120471 |
| | Rock2 | -1.12E+00 | 1.5348009629323 | 0.0291876438 | 0.3598282322 |
| | Rusc1 | -1.12E+00 | 1.3518961019894 | 0.0444737651 | 0.4059017117 |
| | Map1lc3a | -1.12E+00 | 1.6183672310151 | 0.0240786852 | 0.3494073565 |
| | Wdr7 | -1.12E+00 | 1.6521566146683 | 0.0222763168 | 0.3476800338 |
| | Pex5 | -1.12E+00 | 1.6889076917136 | 0.0204687965 | 0.3431030047 |
| | Nmnat2 | -1.12E+00 | 2.1021673279186 | 0.0079037405 | 0.2678489832 |
| | Smcr8 | -1.12E+00 | 1.6325124019640 | 0.0233070656 | 0.3494073565 |
| | Slc12a5 | -1.12E+00 | 1.4801003231528 | 0.0331054638 | 0.3712787029 |
| | Dcaf7 | -1.13E+00 | 1.5807404124883 | 0.0262578757 | 0.3530246727 |
| | Zfp318 | -1.13E+00 | 1.4736447648210 | 0.0336012346 | 0.3723357657 |
| | Sv2a | -1.13E+00 | 1.3155316396136 | 0.0483580033 | 0.4134938233 |
| | Tsen34 | -1.13E+00 | 1.5360928154176 | 0.0291009512 | 0.3598282322 |
| | Slc9a8 | -1.13E+00 | 1.3580623353070 | 0.0438467759 | 0.4058123058 |

| | | | | | |
|---|---|---|---|---|---|
| | Celsr2 | -1.13E+00 | 1.3641702850295 | 0.0432344277 | 0.4039444299 |
| | Pef1 | -1.13E+00 | 1.4127311481582 | 0.0386606233 | 0.3888583996 |
| | Necap1 | -1.13E+00 | 1.9857095159913 | 0.0103345241 | 0.2820882756 |
| | Usp12 | -1.13E+00 | 1.5746627522986 | 0.0266279203 | 0.3530246727 |
| | Bag4 | -1.13E+00 | 1.6426437905221 | 0.0227696424 | 0.3491345164 |
| | Syngr1 | -1.13E+00 | 1.5513594905069 | 0.0280957422 | 0.3577669895 |
| | Nisch | -1.13E+00 | 1.5112734677562 | 0.0308124713 | 0.3616928212 |
| | Tmem184b | -1.13E+00 | 1.9918436036264 | 0.0101895827 | 0.2811259244 |
| | Thrb | -1.13E+00 | 1.5730953118287 | 0.0267241985 | 0.3530246727 |
| | Fbxw7 | -1.13E+00 | 1.3579110728192 | 0.0438620502 | 0.4058123058 |
| | Reep1 | -1.13E+00 | 1.4839842458994 | 0.0328107195 | 0.3697465017 |
| | Klhdc3 | -1.13E+00 | 1.3138116508846 | 0.0485499011 | 0.4134938233 |
| | Rangap1 | -1.13E+00 | 1.9250910445903 | 0.0118825310 | 0.2919722466 |
| | Prkaca | -1.13E+00 | 2.3540501007448 | 0.0044253732 | 0.2313144099 |
| | Katnb1 | -1.13E+00 | 1.8227636957345 | 0.0150396006 | 0.3160649189 |
| | Cog7 | -1.13E+00 | 1.4033844403039 | 0.0395016794 | 0.3914279368 |
| | Cds2 | -1.13E+00 | 1.7437575689941 | 0.0180402450 | 0.3321583165 |
| | Hmgxb3 | -1.13E+00 | 1.5807610502377 | 0.0262566279 | 0.3530246727 |
| | Wbp2 | -1.13E+00 | 1.6938378875187 | 0.0202377447 | 0.3414589314 |
| | Atp6v1b2 | -1.13E+00 | 1.7070122996348 | 0.0196330467 | 0.3414589314 |
| | Dock3 | -1.13E+00 | 1.3409985680610 | 0.0456038420 | 0.4064942202 |
| | Nol6 | -1.13E+00 | 1.6283570342312 | 0.0235311399 | 0.3494073565 |
| | Wdfy3 | -1.13E+00 | 1.3045506400216 | 0.0495963093 | 0.4164190424 |

| | | | | | |
|---|---|---|---|---|---|
| | Ccdc97 | -1.13E+00 | 1.6572650845153 | 0.0220158225 | 0.3476800338 |
| | Myh10 | -1.13E+00 | 1.5765527346765 | 0.0265122915 | 0.3530246727 |
| | Acsl1 | -1.13E+00 | 1.3851686556372 | 0.0411937515 | 0.3979300799 |
| | Rnf130 | -1.13E+00 | 1.5930420794308 | 0.0255245398 | 0.3530246727 |
| | Chmp6 | -1.13E+00 | 1.3939335141608 | 0.0403707191 | 0.3949799091 |
| | Phf24 | -1.13E+00 | 2.1189107388985 | 0.0076048256 | 0.2678489832 |
| | Ube3c | -1.13E+00 | 1.9318813506911 | 0.0116981894 | 0.2919722466 |
| | Fam131b | -1.13E+00 | 1.5535960241411 | 0.0279514264 | 0.3571571155 |
| | Trub2 | -1.13E+00 | 1.4528445917152 | 0.0352496986 | 0.3751773293 |
| | Spock2 | -1.14E+00 | 2.0038280651797 | 0.0099122429 | 0.2793325500 |
| | Tecpr1 | -1.14E+00 | 1.4971035596430 | 0.0318343832 | 0.3651973810 |
| | 6030458C11Rik | -1.14E+00 | 1.7727305327346 | 0.0168759981 | 0.3234566298 |
| | Slc24a3 | -1.14E+00 | 1.5106236315005 | 0.0308586106 | 0.3616928212 |
| | AI837181 | -1.14E+00 | 1.7858881029295 | 0.0163723831 | 0.3220563298 |
| | Klhl22 | -1.14E+00 | 1.8534739331224 | 0.0140128369 | 0.3052796609 |
| | Clstn3 | -1.14E+00 | 1.6943232829073 | 0.0202151383 | 0.3414589314 |
| | Gprasp1 | -1.14E+00 | 1.5338290820852 | 0.0292530341 | 0.3598282322 |
| | L3mbtl2 | -1.14E+00 | 1.6184346334570 | 0.0240749485 | 0.3494073565 |
| | Myt1l | -1.14E+00 | 1.8602005935835 | 0.0137974683 | 0.3042103829 |
| | Usp5 | -1.14E+00 | 1.5343771475914 | 0.0292161410 | 0.3598282322 |
| | Smap1 | -1.14E+00 | 1.8039369208061 | 0.0157059091 | 0.3204763479 |
| | Arhgef11 | -1.14E+00 | 1.3802700499579 | 0.0416610249 | 0.3979300799 |
| | Ldoc1l | -1.14E+00 | 1.3255048534444 | 0.0472601555 | 0.4131608560 |

| | | | | | |
|---|---|---|---|---|---|
| | Agk | -1.14E+00 | 1.3964528077178 | 0.0401372111 | 0.3944648260 |
| | Ube2o | -1.14E+00 | 1.3184376563750 | 0.0480355031 | 0.4134938233 |
| | Tbc1d24 | -1.14E+00 | 1.3574155199600 | 0.0439121276 | 0.4058123058 |
| | 3110035E14Rik | -1.14E+00 | 1.3491134956269 | 0.0447596317 | 0.4059861731 |
| | Slc22a17 | -1.14E+00 | 1.3014541755783 | 0.0499511883 | 0.4169037312 |
| | Prkar1b | -1.14E+00 | 1.4927607463350 | 0.0321543144 | 0.3660376099 |
| | Mical3 | -1.14E+00 | 1.4125849622887 | 0.0386736389 | 0.3888583996 |
| | Ndufa11 | -1.14E+00 | 1.7067534378796 | 0.0196447525 | 0.3414589314 |
| | Pld3 | -1.14E+00 | 1.4595979476359 | 0.0347057994 | 0.3751773293 |
| | Spock1 | -1.14E+00 | 1.4276058341311 | 0.0373589074 | 0.3837082505 |
| | Camsap1 | -1.14E+00 | 2.1723946113993 | 0.0067236545 | 0.2582368390 |
| | Gstz1 | -1.14E+00 | 1.5790483137121 | 0.0263603812 | 0.3530246727 |
| | Sh3bp5 | -1.14E+00 | 1.5573091732141 | 0.0277134649 | 0.3568174154 |
| | Abcg4 | -1.14E+00 | 1.5325364114162 | 0.0293402350 | 0.3598282322 |
| | Gpr85 | -1.14E+00 | 1.3810667548818 | 0.0415846686 | 0.3979300799 |
| | Rad54l2 | -1.14E+00 | 1.5617271212713 | 0.0274329732 | 0.3557041458 |
| | Ap2a2 | -1.14E+00 | 1.3478270592082 | 0.0448924121 | 0.4063987337 |
| | Kcnb1 | -1.14E+00 | 1.5806398545693 | 0.0262639562 | 0.3530246727 |
| | Ipo4 | -1.14E+00 | 1.4966212213075 | 0.0318697590 | 0.3653045083 |
| | Snx12 | -1.14E+00 | 1.4524080000594 | 0.0352851526 | 0.3751773293 |
| | Mul1 | -1.14E+00 | 1.3612445473700 | 0.0435266710 | 0.4049835035 |
| | Umad1 | -1.14E+00 | 1.6709517814249 | 0.0213328175 | 0.3444238574 |
| | Nsf | -1.14E+00 | 1.6417968190554 | 0.0228140916 | 0.3492842042 |

| | | | | | |
|---|---|---|---|---|---|
| | Ubap2l | -1.14E+00 | 2.4472976302807 | 0.0035702808 | 0.2179876018 |
| | Ralgapa1 | -1.14E+00 | 1.6062914771199 | 0.0247575989 | 0.3505036456 |
| | Cbfa2t2 | -1.14E+00 | 1.9440250087012 | 0.0113756178 | 0.2891066989 |
| | Dscam | -1.14E+00 | 1.3083446452987 | 0.0491649220 | 0.4162175084 |
| | Map6 | -1.14E+00 | 2.1487402701579 | 0.0071000226 | 0.2607678450 |
| | Igsf8 | -1.14E+00 | 1.3685316562291 | 0.0428024219 | 0.4030322011 |
| | Erc1 | -1.14E+00 | 1.5109658623584 | 0.0308343031 | 0.3616928212 |
| | Vps33a | -1.14E+00 | 1.5787663055815 | 0.0263775038 | 0.3530246727 |
| | Tubg2 | -1.14E+00 | 1.3965211521176 | 0.0401308952 | 0.3944648260 |
| | Trp53bp1 | -1.14E+00 | 1.5417682663058 | 0.0287231280 | 0.3598282322 |
| | Vti1a | -1.14E+00 | 1.9686332108255 | 0.0107489685 | 0.2867072770 |
| | Kpna1 | -1.14E+00 | 1.5932868169497 | 0.0255101600 | 0.3530246727 |
| | Snap47 | -1.15E+00 | 1.8590368804233 | 0.0138344889 | 0.3042103829 |
| | Rundc3b | -1.15E+00 | 1.5192871434436 | 0.0302491278 | 0.3615628963 |
| | Ptpn3 | -1.15E+00 | 1.4360428120139 | 0.0366401454 | 0.3799720064 |
| | Bloc1s6 | -1.15E+00 | 1.4278525790120 | 0.0373376879 | 0.3837082505 |
| | Slc23a2 | -1.15E+00 | 1.4574192130167 | 0.0348803462 | 0.3751773293 |
| | Ttl | -1.15E+00 | 1.5295223343502 | 0.0295445695 | 0.3608120471 |
| | Ipo11 | -1.15E+00 | 1.9959257140834 | 0.0100942553 | 0.2804404006 |
| | Scarb1 | -1.15E+00 | 1.3422651917636 | 0.0454710317 | 0.4064942202 |
| | Atp2b3 | -1.15E+00 | 1.4445249689565 | 0.0359314738 | 0.3771613156 |
| | Fem1a | -1.15E+00 | 1.4891149318358 | 0.0324253795 | 0.3665818492 |
| | Pds5b | -1.15E+00 | 1.3058323609141 | 0.0494501529 | 0.4162430830 |

| | | | | | |
|---|---|---|---|---|---|
| | Atp6v1a | -1.15E+00 | 1.7636510427877 | 0.0172325266 | 0.3267193892 |
| | Srcap | -1.15E+00 | 1.9993324453115 | 0.0100153828 | 0.2804404006 |
| | Basp1 | -1.15E+00 | 1.5259152053107 | 0.0297909803 | 0.3608120471 |
| | Ndfip1 | -1.15E+00 | 1.6521182281332 | 0.0222782858 | 0.3476800338 |
| | Med14 | -1.15E+00 | 2.1090227357627 | 0.0077799582 | 0.2678489832 |
| | Ica1l | -1.15E+00 | 1.3953097099972 | 0.0402429946 | 0.3945557049 |
| | Syngr3 | -1.15E+00 | 1.4737041928597 | 0.0335966370 | 0.3723357657 |
| | Mfn2 | -1.15E+00 | 1.6025984481669 | 0.0249690231 | 0.3527848889 |
| | Faim2 | -1.15E+00 | 1.3564300004816 | 0.0440118880 | 0.4058123058 |
| | Elac2 | -1.15E+00 | 1.3809196018145 | 0.0415987613 | 0.3979300799 |
| | Gls | -1.15E+00 | 2.3613456223108 | 0.0043516542 | 0.2313144099 |
| | Nol11 | -1.15E+00 | 1.6993056793930 | 0.0199845476 | 0.3414589314 |
| | Pim2 | -1.15E+00 | 1.3666246208545 | 0.0429907854 | 0.4037220341 |
| | Anks1b | -1.15E+00 | 1.4390176806694 | 0.0363900221 | 0.3793105572 |
| | Faah | -1.15E+00 | 1.6988395689763 | 0.0200060077 | 0.3414589314 |
| | Slc25a12 | -1.15E+00 | 1.7060527086999 | 0.0196764747 | 0.3414589314 |
| | Exoc2 | -1.15E+00 | 1.8044680628536 | 0.0156867125 | 0.3204763479 |
| | Dmwd | -1.15E+00 | 1.7803904549884 | 0.0165809552 | 0.3220563298 |
| | Btbd11 | -1.15E+00 | 1.3256133213183 | 0.0472483535 | 0.4131608560 |
| | Zmynd19 | -1.15E+00 | 1.3455984696764 | 0.0451233703 | 0.4063987337 |
| | Zfp532 | -1.15E+00 | 1.3055145951582 | 0.0494863480 | 0.4162430830 |
| | Sec22b | -1.15E+00 | 1.5701337446594 | 0.0269070605 | 0.3530246727 |
| | Mkrn2 | -1.15E+00 | 1.5905387368436 | 0.0256720922 | 0.3530246727 |

| | | | | | |
|---|---|---|---|---|---|
| | Ambra1 | -1.15E+00 | 1.8058278541436 | 0.0156376737 | 0.3204763479 |
| | Champ1 | -1.15E+00 | 1.9571793871520 | 0.0110362267 | 0.2891066989 |
| | Mgat1 | -1.15E+00 | 1.3170907522689 | 0.0481847098 | 0.4134938233 |
| | Dnajc6 | -1.15E+00 | 1.8245003109978 | 0.0149795818 | 0.3160649189 |
| | Cops7a | -1.15E+00 | 2.1780373869449 | 0.0066368593 | 0.2582368390 |
| | Rai1 | -1.15E+00 | 1.3902525067385 | 0.0407143488 | 0.3964281109 |
| | Pcgf3 | -1.15E+00 | 1.5837195468604 | 0.0260783706 | 0.3530246727 |
| | Usp22 | -1.15E+00 | 2.0714896856670 | 0.0084822353 | 0.2717026504 |
| | Med24 | -1.15E+00 | 1.3352969576383 | 0.0462064967 | 0.4112072097 |
| | Syt16 | -1.15E+00 | 1.4236425328978 | 0.0377013991 | 0.3858137340 |
| | Gas7 | -1.15E+00 | 1.3030060773738 | 0.0497730120 | 0.4167090125 |
| | Abtb1 | -1.15E+00 | 1.6785855660756 | 0.0209611176 | 0.3444238574 |
| | Prmt8 | -1.15E+00 | 1.8010754979974 | 0.0158097318 | 0.3204763479 |
| | Tmem151a | -1.15E+00 | 1.6942551523014 | 0.0202183098 | 0.3414589314 |
| | Atp6v0a1 | -1.15E+00 | 2.0173494109080 | 0.0096083893 | 0.2793325500 |
| | Trmt112 | -1.15E+00 | 1.3057096769103 | 0.0494641241 | 0.4162430830 |
| | Scamp5 | -1.15E+00 | 1.7152247186547 | 0.0192652780 | 0.3412234131 |
| | Clmp | -1.16E+00 | 1.3184070702260 | 0.0480388862 | 0.4134938233 |
| | Trmt6 | -1.16E+00 | 1.3181858235537 | 0.0480633654 | 0.4134938233 |
| | Lonrf2 | -1.16E+00 | 1.8813078782657 | 0.0131429278 | 0.2979940050 |
| | Oaz1-ps | -1.16E+00 | 1.5181735697760 | 0.0303267890 | 0.3615628963 |
| | Xxylt1 | -1.16E+00 | 1.3874543568715 | 0.0409775174 | 0.3978647535 |
| | Prepl | -1.16E+00 | 2.1271528793102 | 0.0074618604 | 0.2660616759 |

| | | | | | |
|---|---|---|---|---|---|
| | Heatr3 | -1.16E+00 | 1.7206340167138 | 0.0190268101 | 0.3391945940 |
| | Tssc1 | -1.16E+00 | 1.6336440332581 | 0.0232464139 | 0.3492842042 |
| | Lrrc28 | -1.16E+00 | 1.4632205204181 | 0.0344175126 | 0.3751773293 |
| | Fam169a | -1.16E+00 | 1.4601105566258 | 0.0346648594 | 0.3751773293 |
| | Naa25 | -1.16E+00 | 1.7422033641693 | 0.0181049211 | 0.3325013305 |
| | Kif3b | -1.16E+00 | 2.1738356606707 | 0.0067013815 | 0.2582368390 |
| | Il34 | -1.16E+00 | 1.3145686436252 | 0.0484653503 | 0.4134938233 |
| | Lrfn4 | -1.16E+00 | 1.6127446696878 | 0.0243924448 | 0.3494073565 |
| | Cyb561 | -1.16E+00 | 1.3086285123395 | 0.0491327970 | 0.4162175084 |
| | Sgtb | -1.16E+00 | 1.4662317406307 | 0.0341797010 | 0.3743491063 |
| | Ywhag | -1.16E+00 | 2.1524839876651 | 0.0070390818 | 0.2603300089 |
| | Mettl2 | -1.16E+00 | 1.3991219340597 | 0.0398912886 | 0.3933744996 |
| | Pdxp | -1.16E+00 | 1.9909230389689 | 0.0102112042 | 0.2811259244 |
| | Fkrp | -1.16E+00 | 1.8675512165695 | 0.0135659054 | 0.3030726949 |
| | Stxbp1 | -1.16E+00 | 1.6526299122895 | 0.0222520531 | 0.3476800338 |
| | Fam78b | -1.16E+00 | 1.5588586008812 | 0.0276147680 | 0.3566974032 |
| | Arpc3 | -1.16E+00 | 1.5878252201441 | 0.0258329962 | 0.3530246727 |
| | Adar | -1.16E+00 | 2.4963962300467 | 0.0031886274 | 0.2163581722 |
| | Isy1 | -1.16E+00 | 1.4452796812696 | 0.0358690867 | 0.3771613156 |
| | Pi4ka | -1.16E+00 | 1.9160841218397 | 0.0121315384 | 0.2936807073 |
| | Dlgap1 | -1.16E+00 | 1.8947973902940 | 0.0127409734 | 0.2979940050 |
| | Wasf1 | -1.16E+00 | 2.0435265433579 | 0.0090463515 | 0.2751981128 |
| | Cnnm3 | -1.16E+00 | 1.3592567831748 | 0.0437263490 | 0.4054730182 |

| | | | | | |
|---|---|---|---|---|---|
| | Cnnm1 | -1.16E+00 | 1.4518548646586 | 0.0353301219 | 0.3751773293 |
| | Sh3gl2 | -1.16E+00 | 1.3150075501464 | 0.0484163950 | 0.4134938233 |
| | Ppp1r3f | -1.16E+00 | 1.3208864979874 | 0.0477654091 | 0.4134168354 |
| | Gnaq | -1.16E+00 | 2.0423400966412 | 0.0090710989 | 0.2751981128 |
| | Cyfip2 | -1.16E+00 | 1.8052197045150 | 0.0156595867 | 0.3204763479 |
| | Stx1b | -1.16E+00 | 1.5116994452028 | 0.0307822637 | 0.3616928212 |
| | Tbr1 | -1.16E+00 | 1.3490561145226 | 0.0447655460 | 0.4059861731 |
| | Rexo1 | -1.16E+00 | 1.4288357547603 | 0.0372532567 | 0.3837016578 |
| | Olfm2 | -1.16E+00 | 1.3187265402023 | 0.0480035615 | 0.4134938233 |
| | Phlpp2 | -1.16E+00 | 1.4087250041834 | 0.0390188977 | 0.3897620506 |
| | Bace1 | -1.16E+00 | 2.1265791269033 | 0.0074717249 | 0.2660616759 |
| | Pam | -1.16E+00 | 1.9623036958423 | 0.0109067737 | 0.2878886486 |
| | Sema6b | -1.16E+00 | 1.7960382022508 | 0.0159941733 | 0.3209658675 |
| | Arfgap1 | -1.16E+00 | 1.4192434494664 | 0.0380852272 | 0.3874805927 |
| | Hgs | -1.17E+00 | 1.7916792037747 | 0.0161555146 | 0.3219628834 |
| | Slc35b4 | -1.17E+00 | 1.8089723176977 | 0.0155248596 | 0.3204763479 |
| | Trim23 | -1.17E+00 | 1.5554122856593 | 0.0278347750 | 0.3569669951 |
| | Pom121 | -1.17E+00 | 2.0907503229536 | 0.0081142742 | 0.2704115592 |
| | Hid1 | -1.17E+00 | 1.5769731007125 | 0.0264866419 | 0.3530246727 |
| | Slc41a2 | -1.17E+00 | 1.5118047466178 | 0.0307748010 | 0.3616928212 |
| | Zfp574 | -1.17E+00 | 1.8836321391323 | 0.0130727772 | 0.2979940050 |
| | Unc13a | -1.17E+00 | 1.5881414757143 | 0.0258141913 | 0.3530246727 |
| | Lrrc24 | -1.17E+00 | 1.4686181063939 | 0.0339924051 | 0.3742875348 |

| | | | | | |
|---|---|---|---|---|---|
| | Ppp1r10 | -1.17E+00 | 1.6435587299513 | 0.0227217235 | 0.3491173948 |
| | Ptprn2 | -1.17E+00 | 1.6090688988172 | 0.0245997731 | 0.3496806651 |
| | Gnl3l | -1.17E+00 | 1.6983209882036 | 0.0200299106 | 0.3414589314 |
| | Map3k5 | -1.17E+00 | 1.4924710820168 | 0.0321757677 | 0.3660376099 |
| | Rasgrp1 | -1.17E+00 | 1.7804001053595 | 0.0165805867 | 0.3220563298 |
| | Mre11a | -1.17E+00 | 1.3236150636197 | 0.0474662515 | 0.4134168354 |
| | Zfp9 | -1.17E+00 | 1.4265707154117 | 0.0374480566 | 0.3843425268 |
| | Ergic1 | -1.17E+00 | 2.6361973997636 | 0.0023110141 | 0.2058098505 |
| | Zfyve28 | -1.17E+00 | 2.2262456955642 | 0.0059395604 | 0.2506111110 |
| | Oscp1 | -1.17E+00 | 1.3822134621929 | 0.0414750137 | 0.3979300799 |
| | Ddx56 | -1.17E+00 | 1.6211552067648 | 0.0239246059 | 0.3494073565 |
| | Mpped1 | -1.17E+00 | 2.2143267292533 | 0.0061048257 | 0.2526569467 |
| | Dnm1 | -1.17E+00 | 1.7902347192623 | 0.0162093381 | 0.3219697643 |
| | Lmo4 | -1.17E+00 | 1.4509365710615 | 0.0354049046 | 0.3751773293 |
| | Grid1 | -1.17E+00 | 1.4304762319008 | 0.0371128040 | 0.3831439589 |
| | Coa7 | -1.17E+00 | 1.4358887227559 | 0.0366531477 | 0.3799720064 |
| | Mlycd | -1.17E+00 | 1.3763362385014 | 0.0420401020 | 0.3997279671 |
| | Plekhm3 | -1.17E+00 | 1.3744324662601 | 0.0422247934 | 0.3999569146 |
| | Setd2 | -1.17E+00 | 2.3302580759663 | 0.0046745728 | 0.2334922660 |
| | Epha4 | -1.17E+00 | 1.9244783872211 | 0.0118993054 | 0.2919722466 |
| | Zfp277 | -1.17E+00 | 2.0370450988112 | 0.0091823724 | 0.2758644208 |
| | Prkacb | -1.17E+00 | 1.9973907920603 | 0.0100602601 | 0.2804404006 |
| | Fam219a | -1.17E+00 | 2.3009299068361 | 0.0050011524 | 0.2370478677 |

| | | | | | |
|---|---|---|---|---|---|
| | Psmf1 | -1.17E+00 | 2.0413918127298 | 0.0090909274 | 0.2751981128 |
| | Cap2 | -1.17E+00 | 2.5315084526001 | 0.0029409765 | 0.2163581722 |
| | Pcbp3 | -1.17E+00 | 1.5517237806296 | 0.0280721851 | 0.3577669895 |
| | Cnksr2 | -1.17E+00 | 1.6449305340401 | 0.0226500657 | 0.3491173948 |
| | Med15 | -1.17E+00 | 1.5727850361812 | 0.0267432980 | 0.3530246727 |
| | St3gal5 | -1.17E+00 | 2.2851048269028 | 0.0051867483 | 0.2393752586 |
| | Fam217b | -1.17E+00 | 1.5259620601668 | 0.0297877664 | 0.3608120471 |
| | Wdr11 | -1.17E+00 | 2.3720368493110 | 0.0042458354 | 0.2313144099 |
| | Exosc4 | -1.17E+00 | 1.3824260405352 | 0.0414547175 | 0.3979300799 |
| | Erf | -1.18E+00 | 1.5878091611540 | 0.0258339514 | 0.3530246727 |
| | Wdr91 | -1.18E+00 | 2.0954617613266 | 0.0080267223 | 0.2699502190 |
| | Cep83os | -1.18E+00 | 1.4898594487654 | 0.0323698399 | 0.3662876451 |
| | Rps6kc1 | -1.18E+00 | 2.4021822211260 | 0.0039611180 | 0.2230101985 |
| | Pcnx4 | -1.18E+00 | 1.6840368222114 | 0.0206996584 | 0.3441698960 |
| | Elmo2 | -1.18E+00 | 1.8953050134665 | 0.0127260899 | 0.2979940050 |
| | Mta3 | -1.18E+00 | 2.4526378901414 | 0.0035266480 | 0.2179876018 |
| | Trim3 | -1.18E+00 | 2.0049065060843 | 0.0098876593 | 0.2793325500 |
| | Dazap1 | -1.18E+00 | 1.4695040065836 | 0.0339231360 | 0.3742625025 |
| | Synj1 | -1.18E+00 | 2.3083436010568 | 0.0049165040 | 0.2346209233 |
| | Irf2bp1 | -1.18E+00 | 1.3451534943943 | 0.0451696271 | 0.4063987337 |
| | Rpap1 | -1.18E+00 | 1.5589841149407 | 0.0276067883 | 0.3566974032 |
| | Ap2a1 | -1.18E+00 | 2.3153306620514 | 0.0048380387 | 0.2345212614 |
| | Pacs1 | -1.18E+00 | 1.4782338840256 | 0.0332480452 | 0.3716061537 |

| | | | | | |
|---|---|---|---|---|---|
| | Sez6l | -1.18E+00 | 1.3983578189374 | 0.0399615368 | 0.3937221634 |
| | Dock4 | -1.18E+00 | 2.3930114509260 | 0.0040456522 | 0.2252400833 |
| | Vps51 | -1.18E+00 | 1.5980495723095 | 0.0252319275 | 0.3530246727 |
| | Cbwd1 | -1.18E+00 | 1.3173593041341 | 0.0481549233 | 0.4134938233 |
| | Faxc | -1.18E+00 | 1.4815304232383 | 0.0329966292 | 0.3709476827 |
| | Sphk2 | -1.18E+00 | 2.0894207535840 | 0.0081391536 | 0.2705979278 |
| | Zfp507 | -1.18E+00 | 1.5394400226268 | 0.0288775256 | 0.3598282322 |
| | Hace1 | -1.18E+00 | 1.6259053986449 | 0.0236643512 | 0.3494073565 |
| | Diras2 | -1.18E+00 | 1.3954969351297 | 0.0402256495 | 0.3945557049 |
| | Brpf3 | -1.18E+00 | 1.6614897432770 | 0.0218026989 | 0.3468161738 |
| | Egln3 | -1.18E+00 | 1.3224992256338 | 0.0475883640 | 0.4134168354 |
| | Cbx4 | -1.18E+00 | 1.7175788465140 | 0.0191611316 | 0.3411556805 |
| | Ephb6 | -1.18E+00 | 1.4120600252356 | 0.0387204124 | 0.3888583996 |
| | Tmem115 | -1.18E+00 | 1.3503456513806 | 0.0446328222 | 0.4059017117 |
| | Trpc1 | -1.18E+00 | 1.9555849765623 | 0.0110768181 | 0.2891066989 |
| | Oprl1 | -1.18E+00 | 2.1926486563087 | 0.0064172852 | 0.2552733827 |
| | Fam136a | -1.18E+00 | 1.4743860234887 | 0.0335439326 | 0.3723357657 |
| | Brinp2 | -1.18E+00 | 1.5566628817810 | 0.0277547371 | 0.3568745372 |
| | Tnks | -1.19E+00 | 1.8448902300567 | 0.0142925516 | 0.3076188775 |
| | Celf5 | -1.19E+00 | 1.3636754914552 | 0.0432837130 | 0.4039444299 |
| | Fam8a1 | -1.19E+00 | 1.8166389792690 | 0.0152532020 | 0.3184030382 |
| | Sigmar1 | -1.19E+00 | 1.4650545595200 | 0.0342724728 | 0.3744881569 |
| | Nup210 | -1.19E+00 | 1.5053599316888 | 0.0312348963 | 0.3627695323 |

| | | | | | |
|---|---|---|---|---|---|
| | Cd200 | -1.19E+00 | 1.9104688102402 | 0.0122894144 | 0.2936807073 |
| | Soga3 | -1.19E+00 | 1.9908882091672 | 0.0102120232 | 0.2811259244 |
| | Ldlrad4 | -1.19E+00 | 1.6955762907307 | 0.0201568985 | 0.3414589314 |
| | Ptcd1 | -1.19E+00 | 1.4716388865861 | 0.0337567878 | 0.3732133432 |
| | Rfng | -1.19E+00 | 1.4536926756956 | 0.0351809307 | 0.3751773293 |
| | Sarm1 | -1.19E+00 | 1.8842704600811 | 0.0130535771 | 0.2979940050 |
| | Armcx5 | -1.19E+00 | 1.8094101844184 | 0.0155092150 | 0.3204763479 |
| | Syngap1 | -1.19E+00 | 2.3365329155734 | 0.0046075185 | 0.2333699786 |
| | Lmtk3 | -1.19E+00 | 1.3026405500922 | 0.0498149215 | 0.4167090125 |
| | Bin3 | -1.19E+00 | 1.3035334510974 | 0.0497126082 | 0.4167090125 |
| | Nosip | -1.19E+00 | 2.7328776652218 | 0.0018497896 | 0.1967399285 |
| | Herc3 | -1.19E+00 | 1.9960275668873 | 0.0100918883 | 0.2804404006 |
| | Snca | -1.19E+00 | 1.3254634657348 | 0.0472646596 | 0.4131608560 |
| | Elk1 | -1.19E+00 | 2.4888946556466 | 0.0032441830 | 0.2163581722 |
| | Hdac9 | -1.19E+00 | 1.4173521566111 | 0.0382514448 | 0.3877448358 |
| | Peg13 | -1.19E+00 | 2.2256131157217 | 0.0059482181 | 0.2506111110 |
| | Wbscr17 | -1.20E+00 | 2.1964298507729 | 0.0063616555 | 0.2552733827 |
| | Svop | -1.20E+00 | 1.5245705132927 | 0.0298833641 | 0.3608120471 |
| | Tmem178 | -1.20E+00 | 1.6201233705477 | 0.0239815158 | 0.3494073565 |
| | Ino80e | -1.20E+00 | 1.5282780379229 | 0.0296293389 | 0.3608120471 |
| | Fam171a2 | -1.20E+00 | 1.6569160337000 | 0.0220335242 | 0.3476800338 |
| | Bcl7a | -1.20E+00 | 2.5702297975655 | 0.0026901110 | 0.2132330933 |
| | Grk5 | -1.20E+00 | 1.3555310032200 | 0.0441030878 | 0.4058123058 |

| | | | | | |
|---|---|---|---|---|---|
| | Armcx4 | -1.20E+00 | 1.6537635922225 | 0.0221940422 | 0.3476800338 |
| | Chpf2 | -1.20E+00 | 1.8894098436725 | 0.0129000132 | 0.2979940050 |
| | Dnajc16 | -1.20E+00 | 2.0066416386641 | 0.0098482340 | 0.2793325500 |
| | Ncoa6 | -1.20E+00 | 2.4946825891856 | 0.0032012339 | 0.2163581722 |
| | Rgmb | -1.20E+00 | 2.1525469447910 | 0.0070380615 | 0.2603300089 |
| | Ptprk | -1.20E+00 | 1.7685196553645 | 0.0170404220 | 0.3253450616 |
| | Endov | -1.20E+00 | 1.3646829772498 | 0.0431834189 | 0.4039444299 |
| | Smyd3 | -1.20E+00 | 1.5492023413027 | 0.0282356415 | 0.3581790686 |
| | Tmem18 | -1.20E+00 | 1.7651212373016 | 0.0171742888 | 0.3267193892 |
| | Mboat7 | -1.20E+00 | 1.7006078221728 | 0.0199247177 | 0.3414589314 |
| | Vps18 | -1.20E+00 | 2.4990434113507 | 0.0031692507 | 0.2163581722 |
| | Zfp827 | -1.20E+00 | 2.2410682085847 | 0.0057402630 | 0.2475345036 |
| | Cstf1 | -1.20E+00 | 1.7409481833118 | 0.0181573229 | 0.3325013305 |
| | Napb | -1.20E+00 | 2.2388741749606 | 0.0057693359 | 0.2475345036 |
| | Jph1 | -1.20E+00 | 1.3560097214180 | 0.0440545002 | 0.4058123058 |
| | Ptprn | -1.20E+00 | 2.1599884794422 | 0.0069184932 | 0.2591442415 |
| | Lrfn5 | -1.20E+00 | 2.3659680658481 | 0.0043055827 | 0.2313144099 |
| | Trmt5 | -1.20E+00 | 1.5742103853521 | 0.0266556707 | 0.3530246727 |
| | Atp6v0c | -1.20E+00 | 2.5129441097659 | 0.0030694170 | 0.2163581722 |
| | Gm9855 | -1.20E+00 | 1.7959253301307 | 0.0159983307 | 0.3209658675 |
| | Ext1 | -1.21E+00 | 2.5152736944325 | 0.0030529965 | 0.2163581722 |
| | Scn2a | -1.21E+00 | 1.5421079335933 | 0.0287006721 | 0.3598282322 |
| | Morn4 | -1.21E+00 | 1.9175478620051 | 0.0120907193 | 0.2936670592 |

| | | | | | |
|---|---|---|---|---|---|
| | Large1 | -1.21E+00 | 2.1703734098336 | 0.0067550192 | 0.2582368390 |
| | Rab11fip2 | -1.21E+00 | 1.8676659673524 | 0.0135623214 | 0.3030726949 |
| | Cmip | -1.21E+00 | 2.3307328317451 | 0.0046694655 | 0.2334922660 |
| | Sprn | -1.21E+00 | 2.0518783838280 | 0.0088740448 | 0.2751981128 |
| | Slc25a51 | -1.21E+00 | 2.4278029140839 | 0.0037341958 | 0.2214344051 |
| | Arl4a | -1.21E+00 | 2.4771313086796 | 0.0033332562 | 0.2163581722 |
| | Nell2 | -1.21E+00 | 1.9483582215060 | 0.0112626809 | 0.2891066989 |
| | Uhmk1 | -1.21E+00 | 1.6341964373827 | 0.0232168643 | 0.3492842042 |
| | Gng2 | -1.21E+00 | 1.6544037999776 | 0.0221613494 | 0.3476800338 |
| | Kcnc4 | -1.21E+00 | 1.6851090239454 | 0.0206486173 | 0.3441698960 |
| | Cadm3 | -1.21E+00 | 2.2894437130681 | 0.0051351873 | 0.2382319883 |
| | Klhdc1 | -1.21E+00 | 1.4668381350542 | 0.0341320101 | 0.3743491063 |
| | Nlgn1 | -1.21E+00 | 2.5176761726560 | 0.0030361542 | 0.2163581722 |
| | Otud7a | -1.21E+00 | 1.6672767383318 | 0.0215141039 | 0.3449493701 |
| | Rgs8 | -1.21E+00 | 1.6282489457887 | 0.0235369971 | 0.3494073565 |
| | Hint3 | -1.21E+00 | 1.5874684895990 | 0.0258542242 | 0.3530246727 |
| | Rcl1 | -1.21E+00 | 1.3081527997344 | 0.0491866450 | 0.4162175084 |
| | Slc39a10 | -1.21E+00 | 2.8088162892241 | 0.0015530438 | 0.1967399285 |
| | Rimbp2 | -1.21E+00 | 1.3461388149570 | 0.0450672632 | 0.4063987337 |
| | Tmem132a | -1.21E+00 | 2.1946944381182 | 0.0063871272 | 0.2552733827 |
| | Fbxl18 | -1.21E+00 | 1.3140967091519 | 0.0485180448 | 0.4134938233 |
| | Omg | -1.21E+00 | 1.9202464171440 | 0.0120158247 | 0.2931861221 |
| | Naxe | -1.21E+00 | 2.0348016530812 | 0.0092299287 | 0.2760516419 |

| | | | | | |
|---|---|---|---|---|---|
| | Epg5 | -1.21E+00 | 2.8642784358721 | 0.0013668522 | 0.1967399285 |
| | Sac3d1 | -1.21E+00 | 1.5881278918972 | 0.0258149987 | 0.3530246727 |
| | Slc17a7 | -1.22E+00 | 1.8829015664462 | 0.0130947868 | 0.2979940050 |
| | Mex3b | -1.22E+00 | 1.8037334008960 | 0.0157132709 | 0.3204763479 |
| | Ak5 | -1.22E+00 | 1.6736339854199 | 0.0212014720 | 0.3444238574 |
| | Hs3st2 | -1.22E+00 | 1.4432675554780 | 0.0360356570 | 0.3775804837 |
| | Pacsin1 | -1.22E+00 | 2.3276990890567 | 0.0047021980 | 0.2334922660 |
| | Caln1 | -1.22E+00 | 1.4068888040254 | 0.0391842191 | 0.3904507058 |
| | Slc6a17 | -1.22E+00 | 2.8460608416362 | 0.0014254079 | 0.1967399285 |
| | Lrrc4c | -1.22E+00 | 2.1155085688959 | 0.0076646342 | 0.2678489832 |
| | Flrt2 | -1.22E+00 | 2.0169286929974 | 0.0096177018 | 0.2793325500 |
| | Sema4f | -1.22E+00 | 2.2550685326140 | 0.0055581654 | 0.2460962597 |
| | Ankrd34a | -1.22E+00 | 1.5894950936366 | 0.0257338584 | 0.3530246727 |
| | Appbp2 | -1.22E+00 | 2.0489582135747 | 0.0089339144 | 0.2751981128 |
| | Rnf144b | -1.22E+00 | 1.6278605565387 | 0.0235580557 | 0.3494073565 |
| | Lrfn3 | -1.22E+00 | 1.6924211691852 | 0.0203038703 | 0.3415627104 |
| | D130043K22Rik | -1.22E+00 | 1.4584763622598 | 0.0347955446 | 0.3751773293 |
| | Fhod3 | -1.22E+00 | 2.1038353158922 | 0.0078734429 | 0.2678489832 |
| | Flrt1 | -1.22E+00 | 2.0663279858286 | 0.0085836503 | 0.2730807555 |
| | Arhgap26 | -1.22E+00 | 2.7407592548007 | 0.0018165223 | 0.1967399285 |
| | Atrn | -1.22E+00 | 2.5670316119991 | 0.0027099944 | 0.2136023649 |
| | Pcdhgc5 | -1.22E+00 | 2.0338255086341 | 0.0092506977 | 0.2760516419 |
| | Pygo1 | -1.22E+00 | 1.3546582216814 | 0.0441918088 | 0.4059017117 |

| | | | | | |
|---|---|---|---|---|---|
| | Rasgrf1 | -1.22E+00 | 2.1230118502500 | 0.0075333501 | 0.2669902597 |
| | Etl4 | -1.22E+00 | 2.2165755872825 | 0.0060732955 | 0.2526569467 |
| | Samd8 | -1.23E+00 | 1.6369600084679 | 0.0230695961 | 0.3492842042 |
| | Sstr3 | -1.23E+00 | 2.6307740539915 | 0.0023400544 | 0.2058098505 |
| | Cacng8 | -1.23E+00 | 1.3530987077201 | 0.0443507831 | 0.4059017117 |
| | Lrrc7 | -1.23E+00 | 1.9546854533665 | 0.0110997845 | 0.2891066989 |
| | Fhl2 | -1.23E+00 | 1.8038813414318 | 0.0157079192 | 0.3204763479 |
| | Arhgap1 | -1.23E+00 | 2.5140638347378 | 0.0030615134 | 0.2163581722 |
| | Slc25a22 | -1.23E+00 | 2.0330464293912 | 0.0092673074 | 0.2760516419 |
| | A730017C20Rik | -1.23E+00 | 1.4677444124370 | 0.0340608583 | 0.3743491063 |
| | Tiam1 | -1.23E+00 | 1.5079421783503 | 0.0310497295 | 0.3623552221 |
| | Ypel1 | -1.23E+00 | 1.3283183151607 | 0.0469549827 | 0.4129838577 |
| | Ntng1 | -1.23E+00 | 1.4634270589552 | 0.0344011484 | 0.3751773293 |
| | Kcnv1 | -1.23E+00 | 2.0535712019158 | 0.0088395223 | 0.2751981128 |
| | Paqr9 | -1.23E+00 | 1.7086791652476 | 0.0195578376 | 0.3414589314 |
| | Zfp810 | -1.23E+00 | 1.3282187935649 | 0.0469657440 | 0.4129838577 |
| | Nol4 | -1.23E+00 | 1.9480914140204 | 0.0112696022 | 0.2891066989 |
| | Slc38a7 | -1.23E+00 | 2.1096960365793 | 0.0077679060 | 0.2678489832 |
| | Jrk | -1.23E+00 | 1.7403711313154 | 0.0181814648 | 0.3325013305 |
| | Arntl | -1.24E+00 | 1.7849172983259 | 0.0164090222 | 0.3220563298 |
| | Glg1 | -1.24E+00 | 2.3593917955563 | 0.0043712758 | 0.2313144099 |
| | 2310057M21Rik | -1.24E+00 | 1.9345314147493 | 0.0116270244 | 0.2918195937 |
| | Chrna4 | -1.24E+00 | 1.9630813130613 | 0.0108872623 | 0.2878886486 |

| | | | | | |
|---|---|---|---|---|---|
| | Fen1 | -1.24E+00 | 1.4651092915427 | 0.0342681539 | 0.3744881569 |
| | Ccsap | -1.24E+00 | 1.7570754425032 | 0.0174954274 | 0.3289696643 |
| | Adgrl3 | -1.24E+00 | 3.5196301750404 | 0.0003022524 | 0.1285030856 |
| | 1700030J22Rik | -1.24E+00 | 1.5455132520850 | 0.0284765091 | 0.3598282322 |
| | Smyd4 | -1.24E+00 | 1.4662813442493 | 0.0341757973 | 0.3743491063 |
| | 1700020I14Rik | -1.24E+00 | 1.7851042438567 | 0.0164019603 | 0.3220563298 |
| | Mib1 | -1.24E+00 | 2.4245077055738 | 0.0037626368 | 0.2214344051 |
| | Arhgap32 | -1.24E+00 | 2.2949133003912 | 0.0050709193 | 0.2379431367 |
| | Mtmr9 | -1.24E+00 | 3.0943680468251 | 0.0008046962 | 0.1803630980 |
| | St3gal1 | -1.24E+00 | 1.9228042070364 | 0.0119452651 | 0.2919722466 |
| | 2610524H06Rik | -1.25E+00 | 1.4930486058234 | 0.0321330089 | 0.3660376099 |
| | Gda | -1.25E+00 | 2.3254274511842 | 0.0047268579 | 0.2334922660 |
| | Tma16 | -1.25E+00 | 1.4106269075596 | 0.0388483960 | 0.3895947079 |
| | Tmeff2 | -1.25E+00 | 1.7127818827659 | 0.0193739474 | 0.3414589314 |
| | A930017M01Rik | -1.25E+00 | 1.9719065891259 | 0.0106682556 | 0.2867072770 |
| | Gfra2 | -1.25E+00 | 1.5328939229450 | 0.0293160921 | 0.3598282322 |
| | Far2 | -1.25E+00 | 1.5077949317126 | 0.0310602587 | 0.3623552221 |
| | Nos1ap | -1.25E+00 | 2.3266962340511 | 0.0047130687 | 0.2334922660 |
| | Ears2 | -1.25E+00 | 1.5261194083797 | 0.0297769760 | 0.3608120471 |
| | Tcaf1 | -1.25E+00 | 3.8268582769279 | 0.0001489847 | 0.1241473519 |
| | Lrrc8d | -1.25E+00 | 2.7048789766926 | 0.0019729725 | 0.1973788677 |
| | 2900079G21Rik | -1.25E+00 | 1.3024328211400 | 0.0498387543 | 0.4167090125 |
| | Alkbh8 | -1.25E+00 | 1.4250168846304 | 0.0375822793 | 0.3848754585 |

| | | | | | |
|---|---|---|---|---|---|
| | Smg9 | -1.25E+00 | 2.3474422927157 | 0.0044932202 | 0.2313144099 |
| | Taf4b | -1.25E+00 | 1.5034831261166 | 0.0313701701 | 0.3635344434 |
| | Dkkl1 | -1.25E+00 | 1.3168676884606 | 0.0482094650 | 0.4134938233 |
| | Pex5l | -1.25E+00 | 2.0125949566606 | 0.0097141553 | 0.2793325500 |
| | Fam19a1 | -1.26E+00 | 1.9703276155751 | 0.0107071130 | 0.2867072770 |
| | Ncald | -1.26E+00 | 1.7405966651249 | 0.0181720254 | 0.3325013305 |
| | Sema3a | -1.26E+00 | 1.6855050844381 | 0.0206297952 | 0.3441698960 |
| | Cntnap4 | -1.26E+00 | 1.4445420573948 | 0.0359300600 | 0.3771613156 |
| | Slc7a4 | -1.26E+00 | 1.6131566321024 | 0.0243693176 | 0.3494073565 |
| | Tnip1 | -1.26E+00 | 1.8418538378018 | 0.0143928289 | 0.3085736897 |
| | Pcdhga12 | -1.26E+00 | 1.7384925685242 | 0.0182602800 | 0.3325649255 |
| | 6430584L05Rik | -1.26E+00 | 1.3358875137559 | 0.0461437076 | 0.4110452171 |
| | Ppm1l | -1.26E+00 | 1.4983309195326 | 0.0317445431 | 0.3646157433 |
| | Slc9a7 | -1.26E+00 | 1.5255602460201 | 0.0298153392 | 0.3608120471 |
| | Usp46 | -1.26E+00 | 2.6488157745529 | 0.0022448340 | 0.2058098505 |
| | Tnr | -1.26E+00 | 1.6963375260639 | 0.0201215983 | 0.3414589314 |
| | Dscaml1 | -1.26E+00 | 1.5339082354787 | 0.0292477030 | 0.3598282322 |
| | Ubl7 | -1.26E+00 | 2.2595203282672 | 0.0055014817 | 0.2460962597 |
| | Zfp536 | -1.27E+00 | 1.5280311381637 | 0.0296461882 | 0.3608120471 |
| | Creg2 | -1.27E+00 | 2.5631734844470 | 0.0027341763 | 0.2143044334 |
| | Slc16a7 | -1.27E+00 | 1.4513660785614 | 0.0353699073 | 0.3751773293 |
| | Tmem120b | -1.27E+00 | 1.3125878854166 | 0.0486868991 | 0.4142372309 |
| | Asic2 | -1.27E+00 | 1.7040375044420 | 0.0197679892 | 0.3414589314 |

| | | | | | |
|---|---|---|---|---|---|
| | D430041D05Rik | -1.27E+00 | 3.5724570746202 | 0.0002676350 | 0.1241473519 |
| | Gnaz | -1.27E+00 | 2.0269901188774 | 0.0093974469 | 0.2787445671 |
| | Frrs1l | -1.27E+00 | 1.7816366491733 | 0.0165334448 | 0.3220563298 |
| | Erc2 | -1.27E+00 | 3.1582500927716 | 0.0006946242 | 0.1740281692 |
| | Dgkg | -1.27E+00 | 1.3537295511269 | 0.0442864072 | 0.4059017117 |
| | Necab1 | -1.27E+00 | 2.2669228738811 | 0.0054085036 | 0.2460962597 |
| | Cdon | -1.27E+00 | 1.8034358808380 | 0.0157240392 | 0.3204763479 |
| | Sox12 | -1.28E+00 | 1.5962515474512 | 0.0253366069 | 0.3530246727 |
| | Osbpl10 | -1.28E+00 | 1.9432743848986 | 0.0113952961 | 0.2891066989 |
| | Lrfn1 | -1.28E+00 | 2.0595715329912 | 0.0087182329 | 0.2748692308 |
| | Tmem159 | -1.28E+00 | 1.5267135990240 | 0.0297362638 | 0.3608120471 |
| | Zbtb34 | -1.28E+00 | 2.1736353869451 | 0.0067044725 | 0.2582368390 |
| | Ccdc177 | -1.28E+00 | 1.6468098750338 | 0.0225522629 | 0.3491173948 |
| | Gatad2b | -1.28E+00 | 1.7940140223009 | 0.0160688937 | 0.3211489723 |
| | Plekha7 | -1.28E+00 | 1.5695197450890 | 0.0269451282 | 0.3530246727 |
| | Nckap1 | -1.28E+00 | 2.5229253523224 | 0.0029996781 | 0.2163581722 |
| | Glce | -1.28E+00 | 2.8996942388173 | 0.0012598121 | 0.1967399285 |
| | Kcnf1 | -1.28E+00 | 1.9999163263916 | 0.0100019268 | 0.2804404006 |
| | Xpo4 | -1.28E+00 | 1.5963903378221 | 0.0253285112 | 0.3530246727 |
| | Adnp2 | -1.29E+00 | 3.0817607835712 | 0.0008283983 | 0.1803630980 |
| | Ksr2 | -1.29E+00 | 1.9357103514687 | 0.0115955045 | 0.2915500503 |
| | Adra1d | -1.29E+00 | 2.1167309733756 | 0.0076430909 | 0.2678489832 |
| | Megf9 | -1.29E+00 | 2.5385704727295 | 0.0028935402 | 0.2163581722 |

| | | | | | |
|---|---|---|---|---|---|
| | Exoc6 | -1.29E+00 | 2.3469357318014 | 0.0044984642 | 0.2313144099 |
| | Gabbr2 | -1.29E+00 | 2.8259848159730 | 0.0014928466 | 0.1967399285 |
| | Dusp16 | -1.29E+00 | 1.4252187524574 | 0.0375648144 | 0.3848754585 |
| | Slc36a4 | -1.29E+00 | 1.7988149366878 | 0.0158922381 | 0.3209629067 |
| | Nupl2 | -1.29E+00 | 1.4467910264303 | 0.0357444792 | 0.3767806484 |
| | Chl1 | -1.29E+00 | 2.3492176988184 | 0.0044748894 | 0.2313144099 |
| | Fbxl4 | -1.29E+00 | 1.7815141012515 | 0.0165381109 | 0.3220563298 |
| | Rnf126 | -1.29E+00 | 1.9501829321983 | 0.0112154594 | 0.2891066989 |
| | Nrip1 | -1.29E+00 | 1.5661768259808 | 0.0271533348 | 0.3543825924 |
| | Pxylp1 | -1.30E+00 | 1.8268154097767 | 0.0148999424 | 0.3157797465 |
| | Ncam2 | -1.30E+00 | 2.1107014566137 | 0.0077499436 | 0.2678489832 |
| | Amer3 | -1.30E+00 | 2.7365498260207 | 0.0018342147 | 0.1967399285 |
| | Fv1 | -1.30E+00 | 1.4567463194686 | 0.0349344315 | 0.3751773293 |
| | Alg10b | -1.30E+00 | 2.7826146619695 | 0.0016496254 | 0.1967399285 |
| | Arhgap20 | -1.30E+00 | 2.3745664505887 | 0.0042211769 | 0.2313144099 |
| | Nectin4 | -1.30E+00 | 1.5357514322158 | 0.0291238354 | 0.3598282322 |
| | Tmem132d | -1.30E+00 | 1.3152480296527 | 0.0483895931 | 0.4134938233 |
| | Sorl1 | -1.30E+00 | 2.7102782945581 | 0.0019485955 | 0.1973788677 |
| | Nt5dc3 | -1.30E+00 | 1.5168962283113 | 0.0304161171 | 0.3616928212 |
| | Greb1l | -1.30E+00 | 1.3746466033845 | 0.0422039788 | 0.3999569146 |
| | Tmem245 | -1.30E+00 | 1.4292021043855 | 0.0372218449 | 0.3837016578 |
| | Cdh4 | -1.30E+00 | 1.6347754018685 | 0.0231859342 | 0.3492842042 |
| | Zdhhc8 | -1.30E+00 | 1.9570670751805 | 0.0110390811 | 0.2891066989 |

| | | | | | |
|---|---|---|---|---|---|
| | Tspyl3 | -1.30E+00 | 3.2347061883915 | 0.0005824972 | 0.1589243404 |
| | Epha6 | -1.30E+00 | 1.4942761801995 | 0.0320423101 | 0.3660376099 |
| | Zbed5 | -1.31E+00 | 1.4627743261507 | 0.0344528913 | 0.3751773293 |
| | Slitrk3 | -1.31E+00 | 3.6215188874366 | 0.0002390458 | 0.1241473519 |
| | Parm1 | -1.31E+00 | 3.1042344950974 | 0.0007866209 | 0.1803630980 |
| | Rnf113a1 | -1.31E+00 | 1.4477416966583 | 0.0356663201 | 0.3767806484 |
| | Dyrk2 | -1.31E+00 | 1.5786123721187 | 0.0263868548 | 0.3530246727 |
| | Dact2 | -1.31E+00 | 1.7752416527640 | 0.0167787015 | 0.3220563298 |
| | Rai2 | -1.31E+00 | 1.4593705809451 | 0.0347239737 | 0.3751773293 |
| | Acvr1b | -1.31E+00 | 3.7896450566431 | 0.0001623136 | 0.1241473519 |
| | Prrg2 | -1.31E+00 | 1.3548128740199 | 0.0441760749 | 0.4059017117 |
| | Trim66 | -1.31E+00 | 2.2324012676496 | 0.0058559685 | 0.2482346736 |
| | Zc3h6 | -1.32E+00 | 1.5540832342359 | 0.0279200869 | 0.3570818772 |
| | Xylt1 | -1.32E+00 | 1.7960266750298 | 0.0159945978 | 0.3209658675 |
| | Pcsk1 | -1.32E+00 | 1.4822686873926 | 0.0329405854 | 0.3709120488 |
| | Slc4a8 | -1.32E+00 | 3.6310032717683 | 0.0002338820 | 0.1241473519 |
| | Foxp4 | -1.32E+00 | 1.9111341436541 | 0.0122706016 | 0.2936807073 |
| | Ramp3 | -1.32E+00 | 1.9243784089558 | 0.0119020451 | 0.2919722466 |
| | Sstr1 | -1.32E+00 | 1.9535904669237 | 0.0111278057 | 0.2891066989 |
| | Zfp697 | -1.33E+00 | 2.0764067356879 | 0.0083867416 | 0.2714959017 |
| | Jun | -1.33E+00 | 2.4497632164409 | 0.0035500689 | 0.2179876018 |
| | Wnt9a | -1.33E+00 | 2.2528706052116 | 0.0055863661 | 0.2460962597 |
| | Gnb1l | -1.33E+00 | 1.3413648714573 | 0.0455653939 | 0.4064942202 |

| | | | | | |
|---|---|---|---|---|---|
| | Slitrk4 | -1.33E+00 | 2.7363841127290 | 0.0018349147 | 0.1967399285 |
| | Adam19 | -1.33E+00 | 2.0426101588610 | 0.0090654599 | 0.2751981128 |
| | Pde1a | -1.33E+00 | 2.5438661509549 | 0.0028584714 | 0.2163581722 |
| | Riox2 | -1.33E+00 | 1.9498998686493 | 0.0112227718 | 0.2891066989 |
| | Sertm1 | -1.33E+00 | 2.4788483045128 | 0.0033201041 | 0.2163581722 |
| | Vps37d | -1.34E+00 | 1.4160453270056 | 0.0383667200 | 0.3877448358 |
| | 1700001L19Rik | -1.34E+00 | 1.4502775593876 | 0.0354586699 | 0.3751773293 |
| | Cdh6 | -1.34E+00 | 1.4369058646255 | 0.0365674045 | 0.3799720064 |
| | Hs6st3 | -1.34E+00 | 1.6732244514116 | 0.0212214741 | 0.3444238574 |
| | Nxph3 | -1.34E+00 | 1.4254115112999 | 0.0375481452 | 0.3848754585 |
| | Fbxo34 | -1.34E+00 | 3.7043517363822 | 0.0001975369 | 0.1241473519 |
| | Pnmal1 | -1.34E+00 | 2.1646192559904 | 0.0068451149 | 0.2588597369 |
| | Bid | -1.34E+00 | 1.4395599682780 | 0.0363446116 | 0.3792260341 |
| | Rps2 | -1.34E+00 | 1.8301140959477 | 0.0147871985 | 0.3143399933 |
| | Grin2d | -1.34E+00 | 1.4385781153381 | 0.0364268724 | 0.3794127840 |
| | Mrnip | -1.34E+00 | 1.4636284378741 | 0.0343852006 | 0.3751773293 |
| | Grm2 | -1.34E+00 | 2.3092832674811 | 0.0049058779 | 0.2346209233 |
| | Vwc2l | -1.34E+00 | 1.7303361904542 | 0.0186064624 | 0.3341674577 |
| | Dcx | -1.35E+00 | 1.3798257167617 | 0.0417036708 | 0.3979300799 |
| | 9430083A17Rik | -1.35E+00 | 1.8437032416215 | 0.0143316686 | 0.3079223752 |
| | Cntnap5a | -1.35E+00 | 1.9049807736469 | 0.0124456971 | 0.2954946277 |
| | Bmp3 | -1.35E+00 | 1.9386577052801 | 0.0115170776 | 0.2900980240 |
| | Dlgap2 | -1.35E+00 | 2.7909015165524 | 0.0016184470 | 0.1967399285 |

| | | | | | |
|---|---|---|---|---|---|
| | Pcdhac2 | -1.35E+00 | 2.7187536109010 | 0.0019109371 | 0.1973788677 |
| | Hdhd3 | -1.35E+00 | 1.5301596447252 | 0.0295012457 | 0.3608120471 |
| | Pcdha12 | -1.35E+00 | 1.6681198946330 | 0.0214723761 | 0.3448840926 |
| | Rasl11b | -1.36E+00 | 2.9023953357458 | 0.0012520010 | 0.1967399285 |
| | Ubb | -1.36E+00 | 1.8906824649512 | 0.0128622674 | 0.2979940050 |
| | Kirrel3 | -1.36E+00 | 1.5245862954801 | 0.0298822782 | 0.3608120471 |
| | Zc3h12b | -1.36E+00 | 1.7003371893188 | 0.0199371378 | 0.3414589314 |
| | B230209E15Rik | -1.36E+00 | 3.2325219803793 | 0.0005854341 | 0.1589243404 |
| | Alg6 | -1.36E+00 | 2.1362213854458 | 0.0073076647 | 0.2654048089 |
| | Chrm3 | -1.36E+00 | 2.5146631543109 | 0.0030572915 | 0.2163581722 |
| | Zfp174 | -1.36E+00 | 1.9247542485169 | 0.0118917495 | 0.2919722466 |
| | Aldh1b1 | -1.36E+00 | 2.1506027287565 | 0.0070696395 | 0.2603334457 |
| | Fem1c | -1.37E+00 | 2.1277054681726 | 0.0074523721 | 0.2660616759 |
| | Ccdc87 | -1.37E+00 | 1.4614119793652 | 0.0345611369 | 0.3751773293 |
| | Gabra5 | -1.37E+00 | 2.1978087401066 | 0.0063414892 | 0.2552733827 |
| | Sorcs3 | -1.37E+00 | 2.4860144845124 | 0.0032657694 | 0.2163581722 |
| | Camk2d | -1.37E+00 | 1.4777230081273 | 0.0332871790 | 0.3716061537 |
| | Klf11 | -1.37E+00 | 1.6554444734751 | 0.0221083090 | 0.3476800338 |
| | Ikzf4 | -1.37E+00 | 1.3686935266355 | 0.0427864716 | 0.4030322011 |
| | Dnajb14 | -1.37E+00 | 2.5789702120883 | 0.0026365122 | 0.2101719681 |
| | Slitrk1 | -1.38E+00 | 3.9055509694782 | 0.0001242937 | 0.1241473519 |
| | Alpl | -1.38E+00 | 1.5713836296544 | 0.0268297342 | 0.3530246727 |
| | Slc7a8 | -1.38E+00 | 3.2573546367742 | 0.0005528984 | 0.1589243404 |

| | | | | | |
|---|---|---|---|---|---|
| | Rnf141 | -1.38E+00 | 3.2224677931298 | 0.0005991454 | 0.1589243404 |
| | Rab40b | -1.38E+00 | 1.8988407882175 | 0.0126229020 | 0.2978853494 |
| | Zfp458 | -1.38E+00 | 1.5217474885874 | 0.0300782463 | 0.3615628963 |
| | Nectin3 | -1.38E+00 | 2.0849288975083 | 0.0082237728 | 0.2714812521 |
| | D130017N08Rik | -1.38E+00 | 1.7075824206101 | 0.0196072903 | 0.3414589314 |
| | H6pd | -1.38E+00 | 1.3899973610500 | 0.0407382753 | 0.3964281109 |
| | Atp8a2 | -1.39E+00 | 3.0255146827261 | 0.0009429427 | 0.1865881948 |
| | Atp2b4 | -1.39E+00 | 1.5181824227650 | 0.0303261708 | 0.3615628963 |
| | D11Wsu47e | -1.39E+00 | 1.6625800126745 | 0.0217480332 | 0.3467328475 |
| | Pstpip2 | -1.39E+00 | 1.7013183698658 | 0.0198921456 | 0.3414589314 |
| | Tmem163 | -1.39E+00 | 2.3466923357347 | 0.0045009860 | 0.2313144099 |
| | Pdzrn3 | -1.39E+00 | 2.2327246587318 | 0.0058516096 | 0.2482346736 |
| | Gnpda1 | -1.40E+00 | 2.4657744017637 | 0.0034215713 | 0.2172155917 |
| | Rad18 | -1.40E+00 | 1.5119150682037 | 0.0307669844 | 0.3616928212 |
| | Lrrc8b | -1.40E+00 | 3.3884095705639 | 0.0004088749 | 0.1565196383 |
| | Zfp551 | -1.40E+00 | 1.3750468613018 | 0.0421651004 | 0.3999569146 |
| | Fstl4 | -1.40E+00 | 2.1446896574865 | 0.0071665534 | 0.2620750757 |
| | 1190002N15Rik | -1.40E+00 | 1.9407496760259 | 0.0114617340 | 0.2896627485 |
| | Klhl11 | -1.40E+00 | 2.0632127485457 | 0.0086454430 | 0.2738048876 |
| | Rprm | -1.40E+00 | 1.6175066186186 | 0.0241264476 | 0.3494073565 |
| | Igsf21 | -1.40E+00 | 1.7164247503162 | 0.0192121182 | 0.3411974914 |
| | Igsf3 | -1.41E+00 | 1.5939828584685 | 0.0254693078 | 0.3530246727 |
| | Ccdc142 | -1.41E+00 | 1.5804162668891 | 0.0262774811 | 0.3530246727 |

| | | | | | |
|---|---|---|---|---|---|
| | Mas1 | -1.41E+00 | 2.2570016808206 | 0.0055334797 | 0.2460962597 |
| | Emx1 | -1.41E+00 | 2.5856471601954 | 0.0025962878 | 0.2093443588 |
| | Fxyd6 | -1.41E+00 | 1.4418505542518 | 0.0361534249 | 0.3782494794 |
| | Gabrg3 | -1.41E+00 | 1.5800799286532 | 0.0262978396 | 0.3530246727 |
| | Rapgef5 | -1.41E+00 | 4.0135077489761 | 0.0000969376 | 0.1241473519 |
| | Atp23 | -1.42E+00 | 1.7601773089436 | 0.0173709148 | 0.3275724935 |
| | Snhg7 | -1.42E+00 | 1.3454914578536 | 0.0451344903 | 0.4063987337 |
| | Nwd2 | -1.42E+00 | 3.5387917779720 | 0.0002892066 | 0.1267990252 |
| | Aff2 | -1.43E+00 | 2.3366148913466 | 0.0046066489 | 0.2333699786 |
| | Ndufs5 | -1.43E+00 | 2.4921962248213 | 0.0032196138 | 0.2163581722 |
| | 1700019D03Rik | -1.43E+00 | 1.3049828396202 | 0.0495469768 | 0.4164190424 |
| | Npr3 | -1.43E+00 | 1.6101742103384 | 0.0245372445 | 0.3495999971 |
| | 4930539E08Rik | -1.43E+00 | 2.1826704778189 | 0.0065664331 | 0.2582368390 |
| | Lmbrd2 | -1.43E+00 | 3.0188410115992 | 0.0009575445 | 0.1865881948 |
| | Ccnf | -1.43E+00 | 1.4757557555643 | 0.0334383042 | 0.3723357657 |
| | 6530402F18Rik | -1.43E+00 | 2.1830353739026 | 0.0065609182 | 0.2582368390 |
| | Vstm2l | -1.44E+00 | 2.2918138072677 | 0.0051072391 | 0.2380550332 |
| | Myo5b | -1.44E+00 | 1.4464056113075 | 0.0357762147 | 0.3768320517 |
| | Kcnj6 | -1.45E+00 | 3.0362380384999 | 0.0009199452 | 0.1865881948 |
| | Zbtb46 | -1.45E+00 | 1.5835614141669 | 0.0260878678 | 0.3530246727 |
| | Rxfp1 | -1.45E+00 | 1.5402283356834 | 0.0288251559 | 0.3598282322 |
| | Bbs10 | -1.45E+00 | 1.5676935292619 | 0.0270586715 | 0.3534759419 |
| | Ankrd34c | -1.45E+00 | 1.7046053333135 | 0.0197421600 | 0.3414589314 |

| | | | | | |
|---|---|---|---|---|---|
| | Nav3 | -1.46E+00 | 2.2493422780715 | 0.0056319361 | 0.2469252002 |
| | Gpr137c | -1.46E+00 | 2.4891826308002 | 0.0032420325 | 0.2163581722 |
| | Prox1 | -1.46E+00 | 1.4202918162122 | 0.0379934021 | 0.3873415761 |
| | Tmem203 | -1.46E+00 | 1.5641735215956 | 0.0272788764 | 0.3547772746 |
| | Zfp941 | -1.46E+00 | 3.1137637465024 | 0.0007695490 | 0.1803630980 |
| | Ddi2 | -1.46E+00 | 2.7446631500609 | 0.0018002667 | 0.1967399285 |
| | AU023762 | -1.47E+00 | 1.3224897534697 | 0.0475894019 | 0.4134168354 |
| | Rasgef1b | -1.47E+00 | 3.0962671044901 | 0.0008011852 | 0.1803630980 |
| | Samd12 | -1.47E+00 | 1.4394372424531 | 0.0363548835 | 0.3792260341 |
| | Arhgef28 | -1.48E+00 | 3.2500193498446 | 0.0005623163 | 0.1589243404 |
| | Col26a1 | -1.48E+00 | 1.3509037113834 | 0.0445755067 | 0.4059017117 |
| | Arfgef3 | -1.48E+00 | 2.1784990833301 | 0.0066298075 | 0.2582368390 |
| | Scube1 | -1.49E+00 | 4.2059318587109 | 0.0000622398 | 0.1241473519 |
| | Grin2b | -1.49E+00 | 2.3631651544425 | 0.0043334605 | 0.2313144099 |
| | LOC432842 | -1.49E+00 | 2.5511961604051 | 0.0028106310 | 0.2163581722 |
| | Rgs12 | -1.49E+00 | 1.8088461620624 | 0.0155293700 | 0.3204763479 |
| | Zfp599 | -1.50E+00 | 1.6710362914297 | 0.0213286667 | 0.3444238574 |
| | Kcnq3 | -1.50E+00 | 3.4353251053157 | 0.0003670075 | 0.1471944524 |
| | Prdm9 | -1.51E+00 | 1.8896742908217 | 0.0128921607 | 0.2979940050 |
| | Kcnh7 | -1.51E+00 | 1.6815248858730 | 0.0208197310 | 0.3444238574 |
| | Pcdha3 | -1.51E+00 | 1.9112183976444 | 0.0122682213 | 0.2936807073 |
| | Mgat5 | -1.51E+00 | 3.1591438612440 | 0.0006931961 | 0.1740281692 |
| | Tmem178b | -1.52E+00 | 2.5444562227922 | 0.0028545902 | 0.2163581722 |

| | | | | | |
|---|---|---|---|---|---|
| | Mob3b | -1.52E+00 | 1.7340933840997 | 0.0184461874 | 0.3327775414 |
| | Nudt17 | -1.52E+00 | 1.5411333245363 | 0.0287651521 | 0.3598282322 |
| | Dgkh | -1.52E+00 | 2.0818115506324 | 0.0082830150 | 0.2714959017 |
| | Grem2 | -1.52E+00 | 3.3444165872748 | 0.0004524634 | 0.1565196383 |
| | Enpp1 | -1.53E+00 | 2.2738935824042 | 0.0053223866 | 0.2448297839 |
| | Fam135b | -1.53E+00 | 2.7913192581394 | 0.0016168910 | 0.1967399285 |
| | Zfp369 | -1.53E+00 | 2.8987091563072 | 0.0012626729 | 0.1967399285 |
| | Gm10052 | -1.53E+00 | 1.6804357158962 | 0.0208720105 | 0.3444238574 |
| | Adamtsl2 | -1.53E+00 | 2.4663625903896 | 0.0034169404 | 0.2172155917 |
| | Efcc1 | -1.54E+00 | 1.7029762665642 | 0.0198163532 | 0.3414589314 |
| | Sh2d3c | -1.54E+00 | 3.4350981843441 | 0.0003671993 | 0.1471944524 |
| | Gpr26 | -1.56E+00 | 3.7822764946936 | 0.0001650910 | 0.1241473519 |
| | Rasgef1c | -1.57E+00 | 3.9256670263963 | 0.0001186678 | 0.1241473519 |
| | Lrch4 | -1.57E+00 | 2.0336943761518 | 0.0092534914 | 0.2760516419 |
| | Mef2c | -1.58E+00 | 2.5034212265213 | 0.0031374642 | 0.2163581722 |
| | Kcnk9 | -1.58E+00 | 2.1295650004456 | 0.0074205313 | 0.2660616759 |
| | 1700016K19Rik | -1.59E+00 | 1.7155797795501 | 0.0192495340 | 0.3412234131 |
| | Barx2 | -1.59E+00 | 1.9253583162492 | 0.0118752205 | 0.2919722466 |
| | Slco4c1 | -1.59E+00 | 1.4527495664876 | 0.0352574122 | 0.3751773293 |
| | Sacs | -1.60E+00 | 1.4282228567778 | 0.0373058675 | 0.3837082505 |
| | Doc2a | -1.60E+00 | 2.6386583470453 | 0.0022979557 | 0.2058098505 |
| | Cyp4x1 | -1.61E+00 | 2.7643914612274 | 0.0017203172 | 0.1967399285 |
| | Epb41l4b | -1.61E+00 | 2.7149398695909 | 0.0019277918 | 0.1973788677 |

| | | | | | |
|---|---|---|---|---|---|
| | Xkr4 | -1.61E+00 | 2.9534495556988 | 0.0011131417 | 0.1967399285 |
| | Zfp738 | -1.61E+00 | 1.5975403583679 | 0.0252615295 | 0.3530246727 |
| | Pcdh20 | -1.61E+00 | 1.4765703102907 | 0.0333756467 | 0.3722260122 |
| | Htr2a | -1.63E+00 | 2.8387094547195 | 0.0014497414 | 0.1967399285 |
| | Fancd2 | -1.63E+00 | 1.6212384064681 | 0.0239200230 | 0.3494073565 |
| | Kcns2 | -1.64E+00 | 2.4754246332956 | 0.0033463809 | 0.2163581722 |
| | Adgrd1 | -1.65E+00 | 1.5216858517294 | 0.0300825154 | 0.3615628963 |
| | Zkscan16 | -1.65E+00 | 3.5617585167783 | 0.0002743099 | 0.1241473519 |
| | Klhdc8a | -1.65E+00 | 2.4696737542812 | 0.0033909879 | 0.2172155917 |
| | Grin2a | -1.66E+00 | 3.3195773810838 | 0.0004790961 | 0.1565196383 |
| | Tnfaip8l3 | -1.66E+00 | 1.8230781540220 | 0.0150287149 | 0.3160649189 |
| | Kcna3 | -1.67E+00 | 1.7955007803527 | 0.0160139777 | 0.3209658675 |
| | Xlr3b | -1.68E+00 | 1.3565024393939 | 0.0440045476 | 0.4058123058 |
| | Htr1a | -1.68E+00 | 3.6225964176370 | 0.0002384534 | 0.1241473519 |
| | Plcxd3 | -1.69E+00 | 1.9619214976766 | 0.0109163764 | 0.2878886486 |
| | Map3k21 | -1.69E+00 | 2.2323672332685 | 0.0058564274 | 0.2482346736 |
| | Gpr27 | -1.70E+00 | 1.5427277242860 | 0.0286597419 | 0.3598282322 |
| | Nyap2 | -1.70E+00 | 2.7083396244521 | 0.0019573134 | 0.1973788677 |
| | Cdkl5 | -1.71E+00 | 5.0958684711750 | 0.0000080192 | 0.1056840753 |
| | C2cd4c | -1.72E+00 | 1.8018400911083 | 0.0157819226 | 0.3204763479 |
| | Usp43 | -1.73E+00 | 1.7554200017274 | 0.0175622437 | 0.3289696643 |
| | Kcnh5 | -1.73E+00 | 2.0579755040092 | 0.0087503313 | 0.2751981128 |
| | Capn6 | -1.73E+00 | 1.4674742199377 | 0.0340820556 | 0.3743491063 |

| | | | | | |
|---|---|---|---|---|---|
| | Gm5468 | -1.73E+00 | 2.3239403110348 | 0.0047430717 | 0.2334922660 |
| | Zfp652os | -1.74E+00 | 1.3855469252932 | 0.0411578874 | 0.3979300799 |
| | Myh1 | -1.74E+00 | 1.6749594092882 | 0.0211368658 | 0.3444238574 |
| | Zfp580 | -1.75E+00 | 2.7618984647157 | 0.0017302208 | 0.1967399285 |
| | Dsg2 | -1.76E+00 | 1.3315270103913 | 0.0466093439 | 0.4125735615 |
| | Zdhhc22 | -1.76E+00 | 2.3186785079499 | 0.0048008871 | 0.2345212614 |
| | Tmem232 | -1.77E+00 | 1.6481834238043 | 0.0224810492 | 0.3491173948 |
| | Glra3 | -1.78E+00 | 1.4918088360154 | 0.0322248693 | 0.3660376099 |
| | Ninj2 | -1.79E+00 | 1.4055960224679 | 0.0393010341 | 0.3908274919 |
| | Sstr4 | -1.85E+00 | 3.5765881110616 | 0.0002651013 | 0.1241473519 |
| | Rimbp3 | -1.86E+00 | 2.4581085514644 | 0.0034825026 | 0.2179876018 |
| | Dynlt1f | -1.87E+00 | 1.4210834499800 | 0.0379242106 | 0.3872464885 |
| | Xkr7 | -1.92E+00 | 1.9275064904771 | 0.0118166265 | 0.2919722466 |
| | Syt10 | -1.94E+00 | 1.6218898670668 | 0.0238841689 | 0.3494073565 |
| | Tenm1 | -1.96E+00 | 2.6068436520893 | 0.0024726141 | 0.2068175162 |
| | Adnp | -1.97E+00 | 2.9051556482700 | 0.0012440687 | 0.1967399285 |
| | Oprk1 | -1.98E+00 | 1.6961069615494 | 0.0201322835 | 0.3414589314 |
| | Hunk | -1.98E+00 | 2.0390263030573 | 0.0091405788 | 0.2751981128 |
| | LOC100861615 | -1.98E+00 | 1.6515829882636 | 0.0223057593 | 0.3477220035 |
| | Gm21949 | -1.99E+00 | 2.1190394045825 | 0.0076025729 | 0.2678489832 |
| | Pbld1 | -2.01E+00 | 1.7504141192700 | 0.0177658455 | 0.3310156869 |
| | 4930413G21Rik | -2.04E+00 | 2.0062823470191 | 0.0098563849 | 0.2793325500 |
| | A230001M10Rik | -2.04E+00 | 2.4895162623172 | 0.0032395429 | 0.2163581722 |

| | | | | | |
|---|---|---|---|---|---|
| | Lyrm7 | -2.10E+00 | 2.6332001750350 | 0.0023270184 | 0.2058098505 |
| | Hcn4 | -2.10E+00 | 1.4929463277339 | 0.0321405772 | 0.3660376099 |
| | Pla2g4b | -2.12E+00 | 3.9895026866874 | 0.0001024465 | 0.1241473519 |
| | Cfap206 | -2.13E+00 | 2.1660817764065 | 0.0068221022 | 0.2586867416 |
| | Zcchc12 | -2.14E+00 | 2.5232411170373 | 0.0029974979 | 0.2163581722 |
| | Nptxr | -2.16E+00 | 2.8305502687379 | 0.0014772355 | 0.1967399285 |
| | Npbwr1 | -2.18E+00 | 2.1338855907898 | 0.0073470739 | 0.2656686780 |
| | Melk | -2.37E+00 | 1.3015339982022 | 0.0499420082 | 0.4169037312 |
| | Gpr39 | -2.62E+00 | 2.2964626514417 | 0.0050528610 | 0.2379431367 |
| | Sp6 | -4.50E+00 | 2.4108302932525 | 0.0038830207 | 0.2230101985 |
| | Tgoln2 | -4.64E+00 | 2.4004441729062 | 0.0039770022 | 0.2230101985 |
