## Supplementary material for "BRAFV600E Expression in Mouse Neuroglial Progenitors Increase Neuronal Excitability, Cause Appearance of Balloon-like cells, Neuronal Mislocalization, and Inflammatory Immune response": Sup. Table 4 and 5

Table S4. Calcium currents average peak and current density

| Condition | $I_{Ca}$ peak (pA) | $I_{Ca}$ peak (pA/pF) | $C_m$ (pF) | n |
|---|---|---|---|---|
| **GLAST+ untransfected neighbor** | **-837.27 ± 390.4** | **-9.39 ± 2.99** | **84.61 ± 14.61** | **2** |
| **GLAST+ BRAFV600E** | **-1036.67 ± 216.72** | **-6.75 ± 1.33** | **168.44 ± 41.64** | **4** |
| **NESTIN+ control-FP** | **-1263.42 ± 512.77** | **-6.45 ± 3.22** | **220.93 ± 32.93** | **3** |
| **NESTIN+ untransfected neighbor** | **-1760.51** | **-6.16** | **285.8** | **1** |
| **NESTIN+ BRAFV600E** | **-367.65** | **-2.26** | **162.4** | **1** |

Table S5. Potassium peak currents insensitive to 100 µM 4AP and sensitive to 100 µM 4AP with current densities

| Condition | $I_K$ peak (pA) with pre-pulse inactivation subtraction after 100 µM 4AP at +20 mV | $I_K$ peak (pA/pF) density with pre-pulse inactivation subtraction after 100 µM 4AP | $C_m$ (pF) | n | $I_K$ peak sensitive to 100 µM 4AP (pA) at +20 mV | $I_K$ peak density sensitive to 100 µM 4AP (pA/pF) | n |
|---|---|---|---|---|---|---|---|
| **GLAST+ untransfected neighbor** | **854.37 ± 172.75** | **6.18 ± 2.58** | **169.08 ± 31.36** | **4** | **553.55 ± 9.59** | **3.05 ± 0.68** | **2** |
| **GLAST+ BRAFV600E** | **1752.05 ± 213.19*** | **15.50 ± 4.84** | **137.27 ± 21.29** | **5** | **405.11 ± 53.14** | **3.58 ± 0.64** | **3** |

***  - t(7)=3.15, p=0.016**
